# Comparative evaluation of computational methods for reconstruction of human viral genomes

**DOI:** 10.1101/2025.01.17.633368

**Authors:** Maria J. P. Sousa, Mari Toppinen, Lari Pyöriä, Klaus Hedman, Antti Sajantila, Maria F. Perdomo, Diogo Pratas

**Affiliations:** Institute of Electronics and Informatics Engineering of Aveiro and Intelligent Systems Associate Laboratory, University of Aveiro, Portugal; Department of Electronics, Telecommunications and Informatics, University of Aveiro, Portugal; Department of Forensic Medicine, University of Helsinki, Finland; Department of Virology and Helsinki University Hospital, University of Helsinki, Finland; Forensic Medicine Unit, Finnish Institute for Health and Welfare, Finland

**Keywords:** Human viral genomes, viral reconstruction, sequence assembly, survey, reproducibility

## Abstract

The increasing availability of viral sequences has led to the emergence of many optimized viral genome reconstruction tools. Given that the number of new tools is steadily increasing, it is complex to identify functional and optimized tools that offer an equilibrium between accuracy and computational resources as well as the features that each tool provides. In this paper, we surveyed open-source computational tools (including pipelines) used for human viral genome reconstruction, identifying specific characteristics, features, similarities, and dissimilarities between these tools. For quantitative comparison, we create an open-source reconstruction benchmark based on viral data. The benchmark was executed using both synthetic and real datasets. With the former, we evaluated the effects to the reconstruction process of using different human viruses with simulated mutation rates, contamination and mitochondrial DNA inclusion, and various coverage depths. Each reconstruction program was also evaluated using real datasets, demonstrating their performance in real-life scenarios. The evaluation measures include the identity, a Normalized Compression Semi-Distance, and the Normalized Relative Compression between the genomes before and after reconstruction, as well as metrics regarding the length of the genomes reconstructed, computational time and resources spent by each tool. The benchmark is fully reproducible and freely available at https://github.com/viromelab/HVRS.

## Introduction

With the advancement of high throughput sequencing technologies, it has become easier to study the genetic makeup of viral genomes (1). However, the reconstruction of human viral genomes from short sequence reads remains a challenging and complex process both at the laboratory and computational levels.

Particularly, human samples can be characterized by complex mixtures of viral and host cells, as well as bacteria, fungi, protozoa, and other organisms. This composition can make it difficult to accurately identify and reconstruct viral genomes from metagenomic data.

At a laboratory level, there are additional challenges related to tissues and samples, which need special protocols, such as bone, teeth and bone marrow (2, 3), or which are in sub optimally preserved (4).

Additionally, viruses can exhibit a high degree of genetic diversity, with different strains and variants that may be present within a single infection. This diversity can complicate the assembly and annotation of viral genomes, particularly when reference genomes are not available (5).

The reconstruction of viral sequences is crucial because they can be responsible for diseases such as cancer (6) or pandemics such as Covid-19 (7). Additionally, their lifelong persistence may be associated with human evolution (8, 9).

Human viral genomes can range from a few thousand to over a hundred thousand base pairs in length and contain regions with average high-complexity interspaced with alternated low-complexity (10). These characteristics can make it challenging to generate long and contiguous sequences from short-read sequencing data. Moreover, viral sequences are mostly present in low abundance within metagenomic data, hindering the recovery of viral sequences from noise and other non-viral sequences (11).

At a technical level, the accuracy and completeness of viral genome reconstruction can be affected by sequencing errors, PCR bias, cross similarity, the quality and quantity of the input data, the existence of local low complexity regions, and ambiguous read mapping.

Despite these challenges, many computer programs have been created using one of three main types of methodologies, namely reference-free (RF), reference-based (RB), and hybrid (HB).

The RF (or *de novo*) reconstruction has no prior knowledge of the DNA sequence to be reconstructed (12). This methodology is ideal for reconstructing novel viral genomes, especially with high coverage but computationally intensive for reconstructing enriched samples. A well-known tool is SPAdes (13) that has been extended to metaSPAdes (14), which is used in metaviralSPAdes (15) and, for the reconstruction of coronaviruses, in coronaSPAdes (16).

The RB (or targeted) reconstruction uses a reference or a database of reference sequences upon which the DNA sequence is aligned while spending on average lower computational resources than reference-free approaches, especially when the reconstruction aims to assemble already known targets. However, the database containing the references is critical for accurate reconstruction because, without the proper reference genome, the reconstructed sequence will be inaccurately predicted or unreconstructed. Therefore, choosing a diverse and high-quality database is a critical part of the methodology to accomplish accurate results.

Finally, the HB approaches combine both previous methods. For example, TRACESPipe (17) is a hybrid pipeline that provides the reconstruction using metaSPAdes (14), Bowtie (18) and BWA (19) combined with iterative refinement. These methodologies allow better adaptation to different sequences but require much more computational resources.

In this paper, we provide a survey on computational tools for human viral genome reconstruction, comparing the features and characteristics of these tools. For quantitative comparison purposes, we created a fully reproducible benchmark for human viral reconstruction assessment. This benchmark provides the installation of all programs necessary and the reconstruction of human viral genomes using the selected opensource tools or pipelines.

The benchmark was tested using both generated and real datasets. The generated datasets contain several human DNA viral genomes mutated at different rates, with contamination and human DNA included, followed by the sequencing simulation using different coverage depths. The contamination sequences inserted in the datasets were generated using a pseudo-random algorithm and the human DNA sequences were retrieved from a database. Although the human viral genomes are not synthetic, the mutation rates are generated in silico. This provides a controlled environment to benchmark-test following recommendation practices (20). The mutated viral genomes are compared with the genomes reconstructed by each program using the identity, the Normalized Compression Semi-Distance (NCSD) and the Normalized Relative Compression (NRC). Additionally, the length of the genome reconstructed and of the scaffolds generated, as well as the computational time and the resources spent by each tool are evaluated.

Using real datasets, the genomes are classified so that suitable references, necessary for the execution of some RB and HB programs, can be retrieved from a database. For these datasets, the evaluation process is done using metrics that assess the length of the genome reconstructed and of the scaffolds generated, the computational time and the resources spent by each tool.

## Methodology

This paper contains two main frameworks: a systematic review with feature and qualitative comparison and the benchmark of the tools with quantitative measures.

### Systematic review methodology

The search strategy targeted studies that focused on viral genome reconstruction tools. The databases of articles searched were: Pubmed (https://pubmed.ncbi.nlm.nih.gov), IEEE Xplore Digital Library (https://ieeexplore.ieee.org) and Google Scholar (https://scholar.google.com/).

The search strategies used Boolean logic with MeSH terminology, including terms of reconstruction and general computational terms. The term “assembly or reconstruction” was used to search IEEE Xplore Digital Library and the top results from Google Scholar, including related references and studies/reviews, were screened.

More specifically, when searching for articles in the Pubmed database, two main MeSH searches were made. A more general one ((“Genome, Viral”[MAJR]) AND (“Software”[MAJR])) yielded 98 results, and a more specific search ((“Genome, Viral”[MAJR]) AND (“Software”[MAJR]) AND assembly) yielded 16 results. Moreover, IEEE Xplore Digital Library was searched, using the term “viral reconstruction”, generating 33 results. Lastly, additional articles were considered when mentioned in one or more of the articles selected or if they were found through Google Scholar. All results obtained were filtered using the criteria previously stipulated. This review exclusively included studies that provided an open-source computational tool capable of reconstructing human viral genomes. This review excluded any tools that were not able to be installed locally, tools that were unable to be installed or executed without a registration, tools only accessible through a graphical user interface (GUI) and tools that required aligned reads, contigs or the result of other tools as input. Only articles with full texts (abstracts were excluded) and in the English language were included. The publication dates were specified as 01-2000 to 03-2023.

To compare the reconstruction programs found in the systematic review, several aspects were analysed, namely their programming language, licence, operating system and reconstruction methodology. Additionally, the connections that the reconstruction tools and pipelines shared with each other, as well as the alignment tools used by the programs were taken into consideration.

### Reconstruction benchmark methodology

The benchmark is open-source and fully reproducible, including the installation of each selected tool from the survey. The benchmark contains a total of sixty-five different synthetic datasets, as well as six *real* datasets.

Out of the sixty-five generated datasets, sixty-two consist of four *real* viral sequences (B19V - human parvovirus B19, HPV - human papillomavirus, VZV - Varicella-Zoster virus and MCPyV - Merkel cell polyomavirus) followed by different mutations rates. Additionally, there are three more generated datasets composed of *real* viral sequences and explore different viral compositions, containing the viruses B19V, HPV, VZV, MCPyV, HPyV7 - human polyomavirus 7, HHV6B - human herpesvirus 6B, EBV - Epstein-Barr virus and HCMV - Cytomegalovirus.

Each of the viral genomes was mutated with GTO (21). In many of the datasets generated, viral contamination generated using AlcoR (22) was added, along with human mitochondrial DNA sequences retrieved from the NCBI database (reference sequence NC_012920.1). Each dataset then followed a read simulation process using ART (23).

The characteristics of each generated dataset are available in the Supplementary Table 1 and the viral sequences in each generated dataset can be seen in the Supplementary Table 2. To test the performance of the reconstruction programs under real conditions, six real datasets were considered. The datasets are available in SRA (24) under the codes PRJNA644600 and PRJNA924035 and contain the human viral metagenome, both from single and multiple tissues.

**Table 1.**
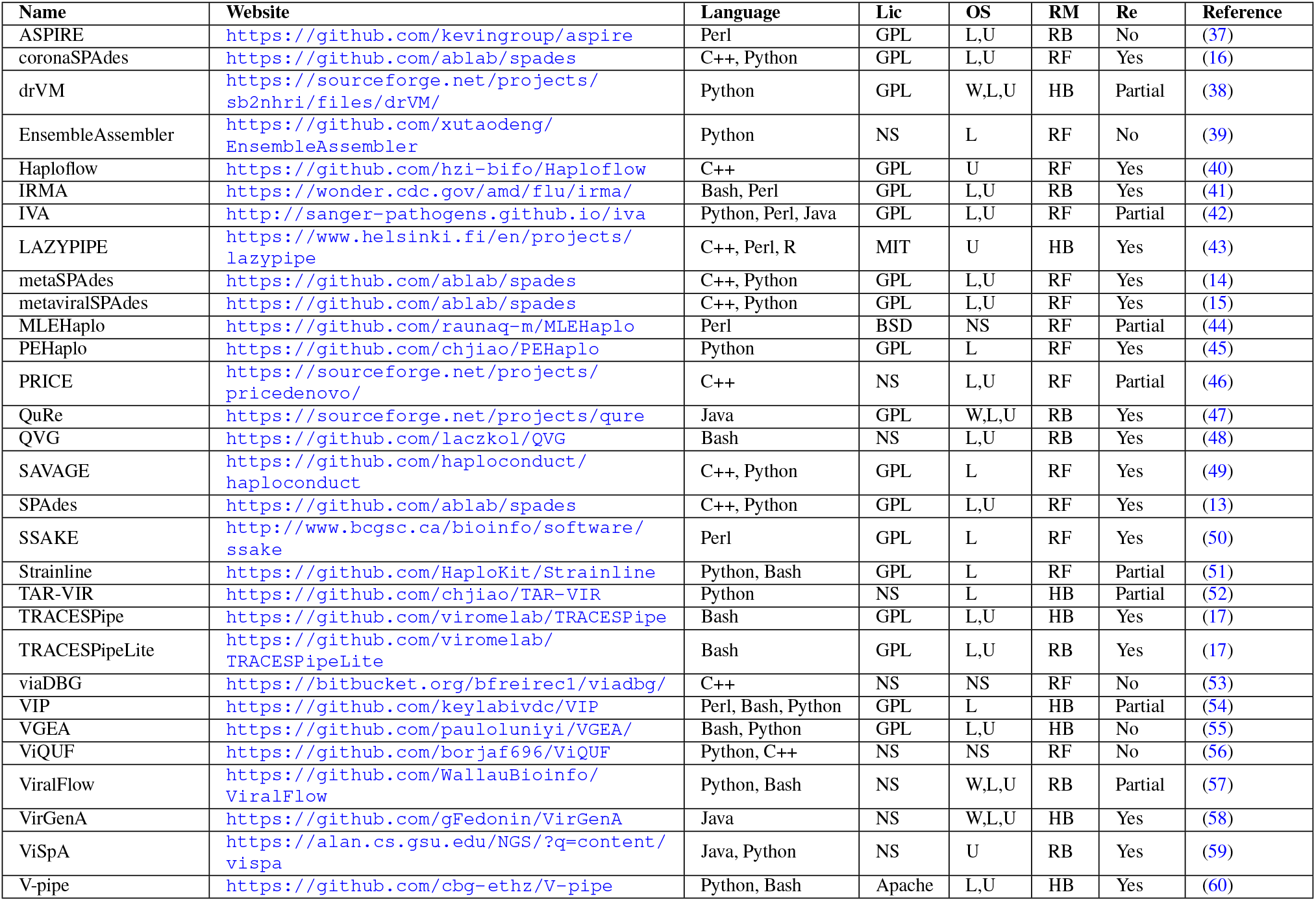
Computational tools used for viral genome reconstruction and their characteristics. Fields OS, RM and Re stand for Operating System, Reconstruction Methodology and Reproducible, respectively. The W, L, and U (OS) stand for Windows, Linux, and Unix, respectively. The RF, RB, and HB (RM) stand for reference-free, reference-based, and hybrid. NS and Lic stand for not specified and License, respectively.

The complete benchmark and scripts to replicate the results are publicly available at the repository https://github.com/viromelab/HVRS. The benchmark is flexible to add or remove more viral sequences, datasets, and tools.

As depicted in Figure 1, after the reconstruction process for each of the benchmarked tools, the reconstructed sequences were compared with the mutated sequences using different approaches: identity, Normalized Compression Semi-Distance (NCSD), Normalized Relative Compression (NRC), length of the genome reconstructed and metrics regarding the length of the scaffolds.

**Fig. 1.**
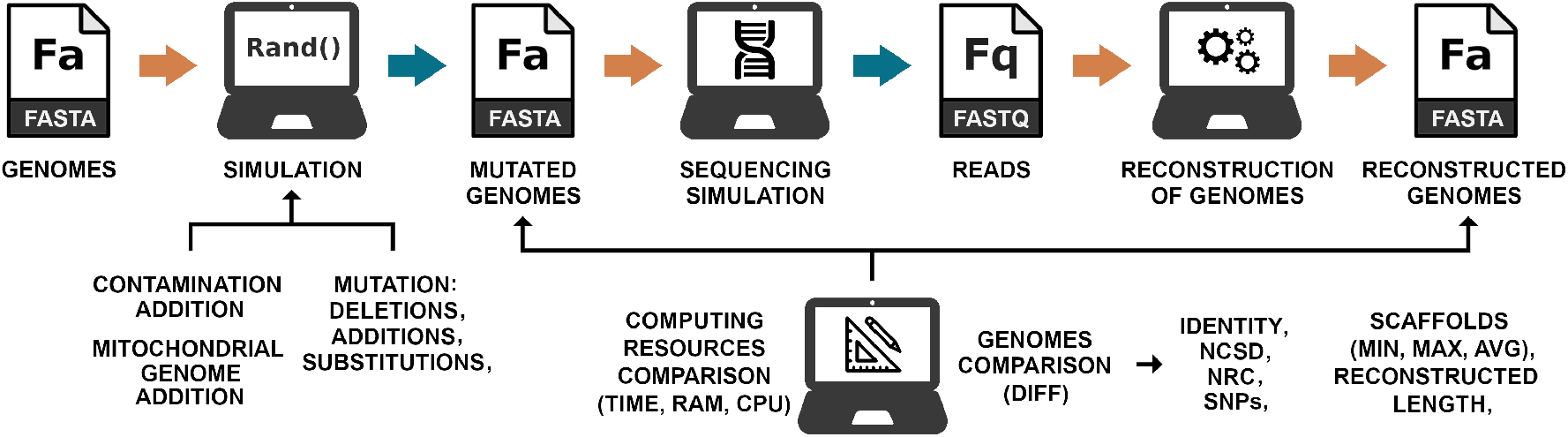
Benchmark methodology depicting the different phases. In the reconstruction phase the genomes are used as references specifically for reference-based and hybrid reconstruction approaches that require them, while the mutated genomes are used only for evaluation in the genomes comparison phase. The blue arrow indicates that the step is optional in the execution of HVRS.

The identity, also referred to as the average identity, was computed with dnadiff, from MUMmer 4 (25). This method has been widely adopted to compare genomes, especially to highlight and provide statistics on the differences between two sequences. Although very practical and fast, because it is not based on lossless data compression approximations and it is based on alignments, it may underestimate or perform ambiguous analyses in the presence of regions of low complexity.

The NCSD is a particular case of the Normalized Compression Distance (NCD) (26). The NCD is an approximation of the Normalized Information Distance (NID) (26–28), a normalized distance derived from the information distance (27) that contains the other known distances, and is defined as

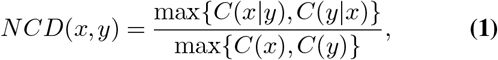

where *x* and *y* are two strings and *C*(*x*) and *C*(*y*) represent the number of bits from the lossless compression on the input *x* and *y*, respectively. *C*(*x* | *y*) and *C*(*y* | *x*) represent the number of bits from the lossless compression on the input *x* given *y* and *y* given *x*, respectively.

However, in our case, we are required to have a relative comparison assuming that if some reconstruction tools provide the reconstructed sequence of the contamination or mitochondrial DNA in *y*, then *y* contains more information than *x* in a perfect assembly. To avoid this constrain, we use the minimum through a semi-distance (29) defined as

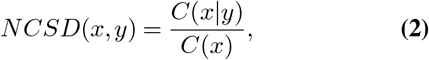

where *x* is the sequence with the conjoint sequences of the mutated viruses before the NGS sequencing simulation process and *y* the sequence with the conjoint reconstructed sequences from a given reconstruction tool. Using the NCSD, the theoretical range of values is between slightly above 0 and close to 1, with 0 indicating that the reference and the file being evaluated are similar and therefore, the genome is correctly reconstructed, and with values close to 1 indicating that the reference and the file being evaluated are completely different, according to the compressor used.

The Normalized Relative Compression (NRC) (30–32) is also a semi-distance and is described as

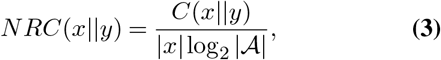

where *x* is the sequence with the conjoint sequences of the mutated viruses before the NGS sequencing simulation process and *y* the sequence with the conjoint reconstructed sequences from a given reconstruction tool.

In this study, we considered the alphabet (𝒜) to have 4 symbols, 𝒜 = {*A, C, G, T*}, corresponding to the bases that compose the genomes in this study. We make sure that symbols outside this alphabet (very rarely appear in full assembled genomes considered references) are substituted by symbols from the 𝒜 using a pseudo-random generation provided by GTO (21). Using the NRC, the theoretical range of values is between slightly above 0 and close to 1, with 0 indicating that the reference and the file being evaluated are similar and therefore, the genome is correctly reconstructed, and with values close to 1 indicating that the reference and the file being evaluated are completely different, according to the compressor used.

The NCSD and NRC contain conditional and relative compression modes that are not supported by many data compression tools, especially for genomic sequences (33, 34). Fortunately, the GeCo3 (35) compression tool has been shown to provide state-of-the-art compression results, at the expense of more computational resources, and it can support, besides the usual compression as *C*(*x*) and *C*(*y*), both relative *C*(*x*||*y*) and conditional *C*(*x*|*y*) compression modes. Therefore, we use GeCo3 to compute these measures.

Additionally, metrics regarding the length of the scaffolds and the number of bases reconstructed were retrieved using SeqKit (36). These metrics can provide us with an upper bound of the amount of information reconstructed by the programs and, when combined with the number of scaffolds generated, can indicate the degree of fragmentation of the reconstruction process. These metrics are especially important when evaluating real datasets, as they can be used even when the true composition of a dataset is unknown. In the context of this review, it was considered that each scaffold is a sequence of nucleotide bases present in the FASTA file output by a reconstruction program that comes after a header.

Each one of the tools was also evaluated taking into consideration the number of SNPs and ratio of SNPs in relation to the number of bases reconstructed. The number of SNPs is a metric retrieved using dnadiff (25) and when combined with the number of bases reconstructed, retrieved by SeqKit (36), it can indicate if a RB or HB program is relying heavily on the reference genome used in the reconstruction process.

Lastly, the computational resources used were quantified through the computational time needed to reconstruct the genomes, the maximum amount of RAM used and the CPU usage. The CPU usage is a percentage calculated as the user time plus system time divided by the total running time.

In order to obtain the statistics, each of the tools was executed three times for each synthetic dataset and two times for each real dataset. Specifically, for obtaining the time of execution of each reconstruction tool for each of the datasets, if three results were obtained, the two closest measures were averaged. If only two results were obtained, the two values were averaged and if only one result was retrieved, it was considered as such. This process is aimed at eliminating possible outliers that may have occurred in cases where the tools successfully reconstructed the datasets in every execution cycle. For the remaining measures, the results obtained were averaged.

In the context of this review, it was considered that a tool performed a metagenomic analysis, as the identification of the sequence organisms contained in a FASTQ sample, if the genomes were reconstructed without the contents of the sample being explicitly given. This can be achieved by reference-free tools that reconstruct sequences of more than one species and by reference-based/hybrid tools that use a database of references from which they measure and select the most suitable ones automatically. It was also considered that metagenomic classification indicates that the scaffolds generated by the tools are classified by their provenance, without relying on previous annotations.

### Reconstruction tools review

In this Section, we describe the human viral genome reconstruction tools found using the search methodology previously defined. Table 1 includes a summary of the characteristics of each tool, including the website, programming language, License, Operating System (OS), reconstruction methodology, and whether or not the tool was reproducible. In the context of this review, it was considered that a tool is reproducible if it was successfully installed and executed, partially reproducible if it was only able to be partially installed or if it was only able to perform some of the tasks desired but did not output results, and not reproducible if was not able to be installed or executed. It should be noted that the inability to install or execute some of the tools considered may be due to version conflicts between dependencies required in the execution process, conflicts with other programs installed, conflicts with the version of the operating system used, or the dependencies being no longer available. According to the type of assembly strategy, the computational tools listed in Table 1 are divided into three categories: reference-based (RB), reference-free (RF), and hybrid (HB).

#### Reference-Based (RB) reconstruction tools

The RB category includes several computational tools such as QuRe (47), QVG (48), TRACESPipeLite (17), ViSpA (59), ViralFlow (57) and IRMA (41).

QVG (48) is a pipeline prepared to deal with single-end and pair-end reads. QVG checks the quality and adapter content of the input reads, and filters them using fastp (61). Afterwards, the reads are aligned to the reference with BWA (62) and duplications are marked with sambamba (63). The variant call is performed by freebayes (64) through parallel (65) for calling variant positions of multiple samples simultaneously. QVG includes multiple statistic outputs, for example, breadth and depth coverage values are provided along with R plots.

TRACESPipeLite (17) is a variation of TRACESPipe for single-end and pair-end sequencing. TRACESPipeLite includes a high quality curated human viral database. TRACESPipeLite uses AdapterRemoval (66) for trimming followed by FALCON-meta (67) for classification, specifically to identify the genomes with the highest similarity (best reference) in the database according to the reads. Then, the reads are aligned to the best references using BWA (62) while the consensus sequences are generated using SAMtools (68) and bcftools (69).

QuRe (47) reconstructs viral quasispecies and corrects errors, using the Poisson distribution, while providing support for reads longer than 100 bp. QuRe is platform-independent as it has been implemented in Java. QuRe can align sequence fragments with a reference genome and partition the genome into sliding windows based on coverage and diversity. Using a heuristic algorithm, QuRe reconstructs the individual sequences of the viral quasispecies while including their prevalence. This feature is achieved by matching multinomial distributions of distinct viral variants that overlap across the genome partition. Additionally, QuRe has a built-in Poisson error correction method and a post-reconstruction probabilistic clustering, both parameterized on given error rates in homopolymeric and non-homopolymeric regions.

ViSpA (59) focuses on reconstructing quasispecies from 454 pyrosequencing reads. ViSpA uses MOSAIK (70), in alternative to SEGEMEHL (71), to align the reads to the reference and extend the reference. Afterwards, it creates a consensus sequence, constructs the read graph, assembles the contigs and estimates the candidate quasispecies sequence frequencies. ViSpA also uses an error correction algorithm, assembles viral variants based on maximum-bandwidth paths in weighted read graphs and does frequency estimation via Expectation Maximization.

ViralFlow (57) is a pipeline created to analyse and assemble the SARS-CoV-2 virus from Illumina paired-end amplicon sequencing data. This pipeline trims the reads using fastp (61) and aligns them against a reference genome with BWA (19). The aligned reads are sorted and indexed with SAM-tools (68) and the minor variants are analysed using both SAMtools and iVar (72). ViralFlow is also capable of identifying intrahost variants, evaluating the quality of the consensus and of the set of mutations retrieved using nextclade (73) and retrieving the assembly metrics with bamdst (74).

IRMA (41) is a pipeline designed to assemble highly variable viral RNA genomes, detect indels and perform variant calling and phasing. The pipeline begins with the filtering of the input reads based on their length or quality. The filtered reads are then aligned to a reference genome, using SAM (75) or BLAT (76), creating a new consensus sequence which allows more reads to be aligned. After this process, the pipeline enhances the consensus sequence generated using the implementation of the Striped Smith-Waterman (SSW) provided by (77).

#### Reference-Free (RF) reconstruction tools

SPAdes (13) is a tool for *de novo* assembly that is capable of single-cell and multicell assembly and can be applied to single, pairedend, and mate-pairs reads. SPAdes uses k-mers for building the initial de Bruijn graph (78, 79) and in the following stages it performs graph-theoretical operations which are based on graph structure, coverage and sequence lengths. The errors are minimized iteratively. Four main phases are used in SPAdes. The first phase is the assembly graph construction, where SPAdes employs different k-mer de Bruijn graphs, which detect and remove bubble and chimeric reads. In the second phase, the pairs of k-mers are adjusted and exact distances between k-mers in the genome are estimated. The third phase is the paired assembly graph construction. The last phase is the contig construction. Here, SPAdes constructs the contigs and maps the reads back to their positions in the assembly graph after graph simplification. SPAdes serves as a base for a myriad of other assembling pipelines, including the following pipelines that come with the SPAdes package, namely metaSPAdes (14), metaviralSPAdes (15) and coronaSPAdes (16).

The metaSPAdes (14) is a pipeline developed to assemble genomes from metagenomic datasets. To reconstruct the genomes, metaSPAdes uses a de Bruijn graph generated by SPAdes (13) and, from it, creates the assembly graph. Afterwards, it uses a modified version of exSPAnder (80) to resolve repeats and scaffolding in the graph.

The metaviralSPAdes (15) is a pipeline made to identify and reconstruct viral genomes in metagenomic samples. It starts by using metaSPAdes (14) to construct the assembly graph and modifies the graph with viralAssembly. The provenance of the contigs assembled is then assessed by viralVerify (81). Finally, viralComplete (82) determines whether the viral contigs correspond to the entirety of the viral genome by comparing them to a database, using a Naive Bayesian Classifier. The coronaSPAdes (16) is another pipeline variation of SPAdes that focuses on the recovery of coronaviruses, but is also capable of making RNA species recovery. It is a pipeline that uses rnaviralSPAdes (83) to assemble the input data and afterwards uses HMMPathExtension to align HMMs to the assembly graph, which is then used to create the assembly graph paths. Although HMMs based on Pfam SARS-CoV-2 (84) are included in coronaSPAdes, there is a possibility to create a custom set, providing additional flexibility.

Another RF reconstruction program is SAVAGE (49), specifically for quasispecies - the ensemble of viral strains populating an infected person. SAVAGE is based on overlap graphs and relies on deep coverage datasets (≥ 20, 000x). It has two main modes of operation: RF reconstruction, which uses FM-index-based techniques, and RB, which aligns the reads to the reference provided. In detail, SAVAGE performs overlap graph construction using pairwise overlaps with FM-index or read-to-reference alignment followed by BLAST (85), then checks for the quality of the overlap using an overlap score and a mismatch rate. This process provides an undirected overlap graph that through read orientations enables a directed overlap graph. Finally, the processes of transitive edge removal and read clustering are made recursively until convergence is achieved, and the final contigs are output.

As SAVAGE, viaDBG (53) is a tool that focuses on reconstructing viral quasispecies, however, it uses a de Bruijn graph-based approach. The viaDBG has two main phases, error correction and haplotype inference. The error correction phase identifies solid k-mers via the LoRDEC algorithm (86). The haplotype inference phase builds a de Bruijn graph and obtains the unary paths of the graph. The paired-end information is then added to the graph and some heuristics are used to polish the paired-end information. Finally, the haplotypes are obtained by splitting the graph nodes based on the paired-end information and obtaining the unary paths from the modified graph.

ViQUF (56) is an assembler designed specifically for reconstructing quasispecies, and it is able to provide frequency estimations for the contigs. The ViQUF methodology involves several steps. Firstly, it selects k-mers above a predefined frequency threshold and utilizes them to construct a de Bruijn graph. Subsequently, it solves a min-cost flow problem on a flow network created for each pair of adjacent vertices, using paired-end information. This process generates an approximate paired assembly graph, where the suggested frequency values serve as edge labels. Finally, the original haplotypes are obtained through a greedy path reconstruction, guided by a min-cost flow solution within the approximate paired assembly graph.

SSAKE (50) is an assembler that uses an overlap-based strategy specialized for short reads. The SSAKE methodology involves loading the sequence reads in a hash table keyed by uniqueness along with values representing the number of occurrences of each sequence in the set. Afterwards, the sequences are organized using a prefix tree, including their reverse complement. After, the sequences are sorted by decreasing occurrence. Then, the most frequent sequences are progressively extended by the longest sequences that can be aligned to them. When this process is no longer possible, the extended sequence is complemented and the extension process is repeated.

PRICE (46) is a specialized *de novo* tool designed for paired-reads, which employs a combination of overlap and de Bruijn graph-based strategies to extend contigs. Initially, it identifies and merges identical or closely similar reads to form contigs through overlap graphs. Subsequently, contigs that fall below a user-defined threshold are further extended using a de Bruijn graph approach. Finally, the sequences generated from both assemblies are combined, and redundant information is removed.

Haploflow (40) is a tool for strain-resolved assembly of viral genomes that uses information on differential coverage between strains to deconvolute the assembly graph into strain resolved genome assemblies. The Haploflow methodology involves creating a de Bruijn graph, finding the connected components and turning them into unitig graphs. From the unitig graphs, a set of contigs is generated, based on the flows of the graphs.

Another RF pipeline is Strainline (51) which assembles viral haplotypes from noisy long reads data. To mitigate the drawbacks of noisy long reads, the errors are corrected using a local de Bruijn graph strategy, utilizing both Daccord (87) and Daligner (88). Then, the reads are organized in clusters, determined by Minimap2 (89), where the reads of each cluster are ordered by Spoa (90) and are iteratively extended using an overlap based strategy. Lastly, the resulting contigs are filtered to remove low divergence and low abundance haplotypes.

MLEHaplo (44) is a pipeline designed to reconstruct viral haplotypes, from paired ended data. It corrects errors with BLESS (91) and represents the reads in a de Bruijn graph. The de Bruijn graph is then used by ViPRA (44) to compute a path cover of the graph, retaining the paths that are candidates to being haplotype, from which the haplotypes are chosen.

EnsembleAssembler (39) is a pipeline designed to analyse metagenomic reads and assemble small viral, bacterial, and eukaryotic mitochondrial genomes. The tool assembles the reads using de Bruijn graph based assemblers, in particular, SOAPdenovo2 (92), ABySS (93) and MetaVelvet (94). This assembly is performed by splitting the input data into chunks with 100K reads followed by the assembly of each chunk. All contigs generated are combined and short contigs are filtered. Finally, an overlap graph based assembler, either CAP3 (95) or Minimo from the AMOS package (96), is used to generate the final contigs.

PEHaplo (45) is a pipeline that assembles viral haplotypes from deep sequencing data. It starts by trimming and correcting the input reads using Karect (97), and, afterwards, it constructs an overlap graph. Then, using a path finding algorithm, the contigs are retrieved.

IVA (42) is an assembler developed to reconstruct RNA viruses from short read pairs with highly variable depth. The input reads can be trimmed using Trimmomatic (98). Then, the most abundant kmer is found using kmc (99) and it is extended with reads that have a perfect match to it, which generates the contigs. The contigs are then extended using SMALT (100) and SAMtools (68). Lastly, the contig ends are trimmed and overlapping contigs are merged.

#### Hybrid (HB) reconstruction tools

The hybrid approaches are rare (not to be confused with short and long reads hybrid assembly) and usually provide a unique architecture that goes beyond the diversity of internal tools chosen.

V-pipe (60) is a pipeline that supports quality control, read mapping and alignment, low-frequency mutation calling, and inference of viral haplotypes. Reads are aligned employing a reference-guided approach using ngshmmalign (60). This approach is based on profile hidden Markov models that are tailored to small and highly diverse viral genomes. For the read alignment, the reference sequence can be provided or it can be built *de novo* from the read data using VICUNA (101). Alternatively, reads can be aligned using BWA (62) or Bowtie2 (18). Intermediate results are provided in the form of a consensus sequence per sample, a multiple sequence alignment of all consensus sequences. Finally, variants are called by different tools, namely LoFreq (102) and ShoRAH (103).

TRACESPipe (17) is a pipeline for the reconstruction of viral genomes from single and multiple organ samples. It includes modules for quality control, filtering, assembly, and annotation of viral genomes. TRACESPipe uses a hybrid approach that combines both *de novo* and reference-based assembly methods, which can increase the accuracy and completeness of the reconstructed viral genomes. This pipeline can handle sensitive data and the genome assembly uses the reference with the highest similarity to provide reference-based (RB) assembly with Bowtie2 (18), in addition to reference-free (RF) assembly with metaSPAdes (14), followed by iterative refinement using BWA alignment (104).

ASPIRE (37) is a pipeline that uses a RF assembler, either SPAdes (13) or SGA (105) for generating scaffolds and, after, aligns the scaffolds obtained to a reference using MUMmer 3 (25), while filtering the viral unlikely contigs. Then, the gaps present in the genome are filled using GapFiller (106). ASPIRE uses an iterative refinement procedure involving repeated alignments of scaffolds to the latest version of the reconstructed viral genome followed by gap filling and a correction step (using Bowtie2 (18), SAMtools (68), and bcftools (69)) based on allele frequencies derived from read alignments.

TAR-VIR (52) is a pipeline that was designed to reconstruct and classify RNA viral reads. TAR-VIR starts by eliminating reads that do not have viral origin, using BMTagger (107), Bowtie2 (18) and Karrect (97). Then, it maps the reads to a reference using BWA (19) or Bowtie2 and obtains a seed set. Reads that have a significant overlap with the seed set are iteratively added to the seed set, which is used by PEHaplo (45) to do the strain-level assembly.

The drVM (38) is a pipeline that identifies, reconstructs, and annotates known viral genomes from NGS reads. The drVM aligns the reads to a viral database using SNAP (108), partitions the aligned reads into genus groups, and reconstructs the reads of each genus group using SPAdes (13). Lastly, the annotation of the reconstructed genomes is made using BLAST (109).

VIP (54) is a pipeline that can identify and discover viruses from metagenomic NGS data. VIP includes quality controls and is able to filter reads with similarity to the host using Bowtie2 (18). The input reads are aligned against the Virus Pathogen Resource (110) and Influenza Research Database (111) or against the NCBI Refseq (112) and neighbour genomes, which allows the taxonomic classification of each read. Afterwards, a RF reconstruction is done using Velvet-Oases (113, 114) and the resulting contig is added to the phylogenetic tree.

LAZYPIPE (version 2) (43) is a pipeline designed to discover and reconstruct viruses from metagenomic NGS data obtained from clinical, animal and environmental samples. LAZYPIPE includes the filtering of the input reads based on their quality as well as filtering the reads that contain the genome of the host, using Trimmomatic (98), fastp (61), BWA-MEM (62), SAMtools (68), and SeqKit (36). Afterwards, it assembles the filtered reads using MEGAHIT (115) or SPAdes (13), which are scanned to check for genelike regions with MetaGeneAnnotator (116), and subsequently translated to amino acid sequences with SeqKit (36) and used to assign NCBI taxonomy ids to the contigs. Later, the reads remaining after the filtering process are aligned to the contigs using BWA-MEM. In addition to assembling the genomes, LAZYPIPE also annotates, estimates the abundance and generates statistics.

VirGenA (58) is a software capable of separating strains into genetic groups, creating a consensus sequence for each group, and detecting evidence of cross contamination. VirGenA removes the adapter and polymerase chain reaction primer sequences, using Trimmomatic (98), then aligns the reads to one or more reference sequences and clusters adjacent reads using Usearch (117), which are then joined generating contigs. The contigs are then used to generate an overlap graph, which along with the reference is used to merge the contigs.

VGEA (55) is a pipeline that focuses on the reconstruction of RNA viruses. It starts by trimming and filtering the reads using fastp (61) and, using BWA (19), it aligns the reads to a reference human genome. Afterwards, using SAMtools (68), the unmapped reads are extracted and split into FASTQ files, which are reconstructed using IVA (42). The reads are then processed in terms of quality and contamination and mapped to a reference, using shiver (118). Lastly, VGEA cleans up the reconstruction made using SeqKit (36). In addition to reconstructing the genomes, VGEA is also capable of evaluating its performance with QUAST (119).

#### Uncategorized reconstruction tools

In this subsection, we include several tools that were not in accordance with the criteria defined in the methodology or that could not be categorized entirely into one of the reconstruction methodologies.

HAPHPIPE (120) is a modular pipeline that assembles genomes using either a RF or RB strategy. It is designed to assemble viral consensus sequences and haplotypes. It includes quality controls, such as trimming the reads with Trimmomatic (98) and correcting them using SPAdes (13). Then, if the RF strategy is used, the contigs are generated with SPAdes (13) and then joined into scaffolds using MUMmer 3 (or after) (25). Otherwise, if the RB strategy is chosen, the reads are aligned to the reference sequence using Bowtie2 (18) and then are realigned with Picard (121).

There are many tools to reconstruct viral genomes that require the output of other tools or pre-aligned reads in order to perform the reconstruction process. Examples of these tools are Virus-VG (122), VG-Flow (123), QSdpR (124), PredictHaplo (125), aBayesQR (126), HaploClique (127), QuasiRecomb (128), ViQuaS (129), RegressHaplo (130), CliqueSNV (131), TenSQR (132), ShoRaH (103), viralFlye (133), Arapan-S (134) and ContigExtender (135).

Moreover, there are tools that can reconstruct viral genomes, but do not comply with other criteria stipulated, such as VirAmp (136), Vipie (137) and EDGE COVID-19 (138). VirAmp can only be installed via Amazon Web Services, Vipie requires a registration to be used and EDGE COVID-19 is only accessible through a GUI. We acknowledge viral-ngs (139) which, unfortunately, does not have a complete article describing the methodology. We also acknowledge viralrecon (140), which is a pipeline capable of performing variant calling for viral samples for both Illumina and Nanopore sequencing data, however, it only supports the assembly of Illumina sequencing data. Additionally, both Genome Detective Virus tool (141) and EzCOVID19 (142) do not meet the criteria stipulated as they are only available online and require a registration.

Furthermore, there are other tools that while not being specifically designed to reconstruct viral genomes, can be used for that purpose. Examples include Falcon (143), HiCanu (144), hifiasm (145) and IPA (146), which are tools that reconstruct HiFi reads. Currently, long reads are starting to be included also in viral reconstruction pipelines and the integration of these assemblers is of significant importance. Additionally, there are assembly tools that are included directly or by option in some of the described pipelines, such as MEGAHIT (115), Velvet (113), MetaVelvet (94), SOAPdenovo2 (92), ABySS (93), CAP3 (95), Minimo (96) and SGA (105).

#### Considerations between tools and pipelines

In the realm of human viral genome reconstruction, there are two primary categories of programs: individual tools and pipelines. Tools are standalone programs that are meticulously optimized using programming languages known for their efficiency, such as C, C++, Rust, and others. However, their flexibility is constrained to specific functions, such as trimming, assembly, or classification. On the other hand, pipelines are comprehensive programs that facilitate the integration of multiple instructions and tools in a unique and customized manner. Typically, pipelines employ programming languages like Bash, Python, or Perl. The flexibility of pipelines surpasses that of individual tools due to the wider array of available sub-programs and parameters, thereby offering a greater number of sequential options. Additionally, pipelines often prioritize experimentation, allowing for optional usage and combination of specific tools. However, it is important to note that this heightened flexibility often comes at a higher computational cost.

Figure 2 provides an illustration of the connections between the reconstruction tools and pipelines found in the present systematic review, as well as the alignment tools used by these programs.

**Fig. 2.**
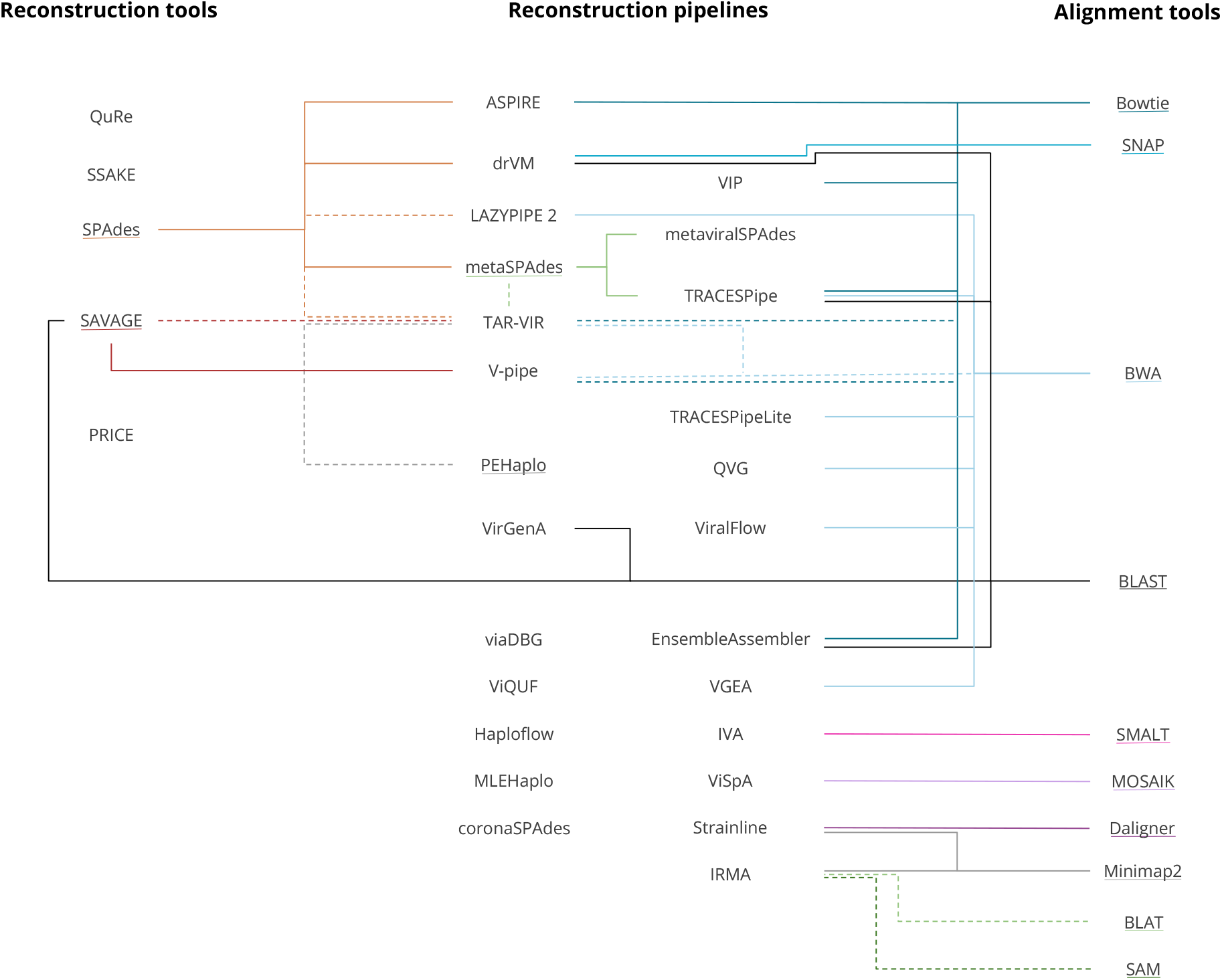
Classification of programs that perform genome reconstruction in three groups: reconstruction tools, reconstruction pipelines and alignment tools. The reconstruction pipelines contain the two columns in the middle. Programs that have their names underlined are the basis tool. Connections made with a dashed line indicate that the tool is optional. The tools or pipelines include all versions in the name as proxies. The order in which the tools appear follows no particular criteria.

Accordingly, there are two types of tools, reconstruction (*de novo*) tools, which may contain small alignment options, and exclusively alignment tools. From the included pipelines, the two most used reconstruction tools are SPAdes and SAVAGE, while the most commonly used alignment tools are BWA and Bowtie.

The alignment tools can be used for several purposes depending on the program that includes them. The following paragraphs outline the purposes each alignment tool has in each reconstruction pipeline considered.

Bowtie is used by ASPIRE to correct the reconstruction made, but can also be used to filter out the host genome, as seen in the reconstruction pipelines VIP and Ensemble-Assembler. Additionally, it can be used to align reads to a reference, as is the case in TRACESPipe and V-pipe. Furthermore, this tool can be used by TAR-VIR to obtain seed reads.

SNAP is only included in one of the pipelines considered in this review, drVM, and it is used to generate a viral database and to align the input reads to it.

BWA is used by LAZYPIPE and VGEA to filter out the host genome, and by TRACESPipe to combine scaffolds. BWA is also used to obtain seed reads, as is the case of the TAR-VIR pipeline, and to align the reads to a reference, as shown in TRACESPipeLite, V-pipe, QVG and ViralFlow.

BLAST is used for several different purposes, namely to identify to what species a scaffold belongs to, as observed in TRACESPipe, to construct overlap graphs, as seen in SAVAGE, and to perform contig annotation, as shown in drVM. BLAST is also used by VirGenA, which uses this alignment tool to identify chimeras and by EnsembleAssembler to improve the quality of the reads and remove the adaptors present.

SMALT is included in only one of the pipelines considered, IVA, to extend the contigs previously obtained, with reads that do not have a perfect match to them.

MOSAIK is used by the pipeline ViSpA to align the input reads to a reference sequence provided by the user.

Minimap2 is utilised by Strainline to determine pairwise alignments between seed reads and the corrected reads and by IRMA in the final assembly step. Strainline also uses another alignment tool, Daligner, in the error correction process to compute read overlaps.

BLAT and SAM are included in the IRMA pipeline to align the input reads to a reference genome.

### Benchmark results

The results reported in this section follow the methodology described in the Section “Methodology”, specifically in the subsection “Reconstruction benchmark methodology”. The results obtained regarding the synthetic datasets generated and accompanying figures are freely available either in the present section or in the Supplementary Material and are fully reproducible using the tool available at https://github.com/viromelab/HVRS. The benchmark was executed on a computer running Linux Ubuntu 22.04.4 LTS with an Intel® Core™ i7-6700 CPU @ 3.40GHz × 8 processor and with 64 GB of RAM and the computational resources were limited, when possible, to 48 GB of RAM and 6 CPU threads. For each program, the execution time was capped at six hours, either overall or per reference genome provided. The analysis focused on six characteristics of the datasets (existence of mitochondrial DNA and contamination, variation in the percentage of SNP and depth coverage, read length and viral composition) and takes into consideration several metrics, focusing especially on the identity, NCSD, NRC, number of nucleotide bases reconstructed, the number of scaffolds generated, the average length of the scaffolds and the ratio between the number of SNPs and number of bases reconstructed. When comparing length-based metrics, the gaps (“N” bases) present in the reconstructed genome were removed for the comparison of the results to be fairer. When comparing sets of datasets, the average performance in a given metric was the average between the values obtained in the datasets considered.

Additionally, a comparison of the overall performance obtained by the reconstruction programs across all synthetic datasets considered will be provided.

#### Mitochondrial DNA and contamination

In order to assess the impact of the inclusion of mitochondrial DNA and contamination on the performance of the reconstruction programs considered, datasets with a low percentage of SNPs, specifically 1%, and a depth coverage varying between 2x and 40x were generated and evaluated. A low percentage of SNPs was considered as the data obtained from real-life scenarios often contains diversity and as the viral genomes contained in the samples may have mutations, making them different from reference genomes.

This analysis focused on datasets 1 to 16 and the performance obtained in terms of the identity, NCSD and NRC in these datasets is represented in Figure 3. The top three plots of Figure 3 represent the results obtained in datasets 1 through 8, which contain no contamination or mitochondrial DNA, whereas the bottom plots represent the results obtained in datasets 9 to 16, which contain both contamination and mito-chondrial DNA.

**Fig. 3.**
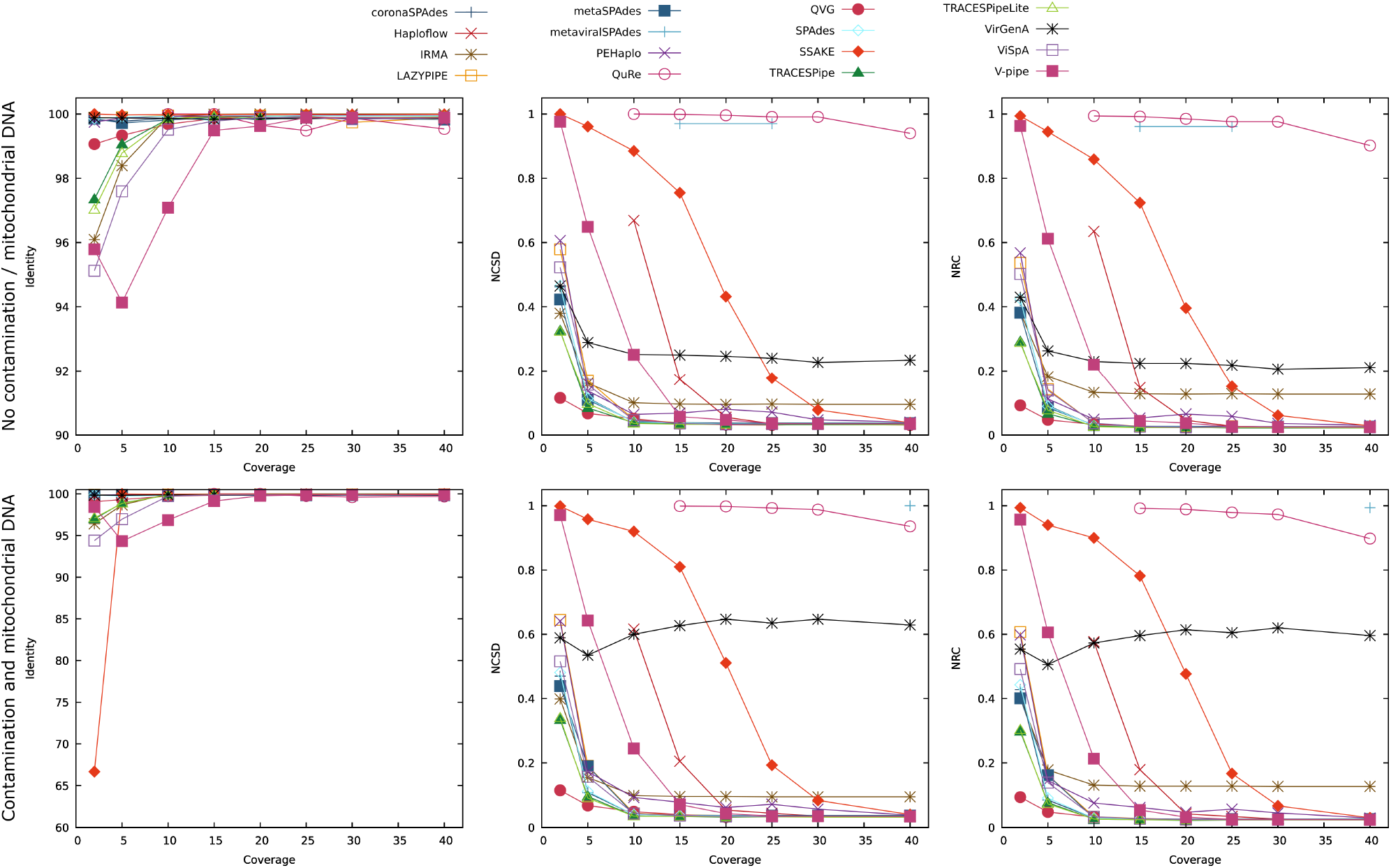
Comparison of the performance of the reconstruction programs according to the identity, NCSD and NRC for datasets without contamination and mitochondrial DNA (DS1 to DS8) and datasets with contamination and mitochondrial DNA (DS9 to DS16), with depth coverage ranging between 2x and 40x. Optimal identity values are close to 100, while lower NCSD and NRC values indicate better results.

Not all reconstruction programs were able to reconstruct datasets 1 through 8, with Haploflow and QuRe unable to reconstruct datasets with a depth coverage equal to or lower than 5x and metaviralSPAdes only reconstructing two of the datasets considered (DS4 and DS6).

The identity tended to improve at higher depth coverage, often stabilizing at values close to 100. It should be noted that the identity is an alignment-based metric, and, as such, it may provide ambiguous analyses of the data.

The results obtained regarding the NCSD and NCD are similar to each other, as they are both compression-based evaluation metrics that assess the reconstructed genomes as a whole. Similar to the identity, the tools tend to improve their performance as the depth coverage increases. Exceptions to this trend are QuRe and metaviralSPAdes, whose performance remained close to 1, and VirGenA, the performance of which stabilized at around 0.25 for depth coverages of 5x and over. QuRe and metaviralSPAdes are incapable of reconstructing many bases and therefore the results obtained using them are more limited than the remaining tools. VirGenA does not significantly alter its performance, however, according to the data obtained, it is only capable of reconstructing scaffolds corresponding to VZV in these datasets. The limitations in the results obtained using QuRe, metaviralSPAdes and Vir-Gena may be due to these programs being designed to work in specific datasets or the capping of the computational resources used to 48 GB of the RAM and 6 threads of CPU, when possible.

In terms of the number of reconstructed bases (excluding gaps), the majority of the programs reconstructed on average between 75,000 bp and 210,000 bp and the performance of each tool has maintained itself stable throughout the datasets considered. However, both QuRe and metaviralSPAdes reconstructed considerably fewer bases per dataset than the remaining programs, reconstructing less than 6,000 bp on average. On the other hand, ViSpA outputs significantly more bases on average than the remaining programs (over 880,000 bp per dataset) as it reconstructed the quasispecies, generating several different scaffolds for a single viral genome. This phenomenon is particularly noticeable in datasets with depth coverage ranging between 2x and 10x. Regarding the average length of the scaffolds and fragmentation of the genomes, IRMA, QVG, TRACESPipe, TRACESPipeLite and V-pipe were able to reconstruct scaffolds with an average length of over 20,000 bp (excluding gaps), while producing the expected number of scaffolds (between 4 and 5) considering there are four viral genomes contained in the datasets, indicating that these tools were able to identify the individual genomes and reconstruct them whole.

The ratio between the number of SNPs and the number of bases reconstructed was stable throughout the datasets. However, there are some outliers, namely PEHaplo, QVG and QuRe. PEHaplo and QVG improved their performances as the depth coverage of the datasets increased, both due to the reduction in the number of SNPs contained in the reconstructed genome and the increase in the number of bases reconstructed. Conversely, the performance of QuRe decreased at depth coverages of 20x and over due to a significant increase in the number of SNPs in relation to the number of nucleotide bases reconstructed.

The bottom three plots of Figure 3 represent the results obtained in terms of the identity, NCSD and NRC for datasets 9 through 16, which contain both contamination and mitochondrial DNA. Again, not all datasets were able to be reconstructed by all of the reconstruction tools, with Haploflow not reconstructing datasets with a depth coverage below 10x, QuRe requiring a depth coverage of at least 15x and metaviralSPAdes only reconstructing the dataset with 40x depth coverage. This means that the addition of contamination and mitochondrial DNA to the datasets can affect the reconstruction ability of the programs.

With the inclusion of contamination and mitochondrial DNA, the results obtained followed the same trend as the ones obtained in datasets 1 to 8. However, the average performance of the tools was inferior with a decrease in performance of 1.1% in terms of the identity, 6.7% in terms of NCSD and 6.9% in terms of the NRC. It should be noted that the results obtained using VirGenA were not as good in terms of NCSD and NRC in relation to the ones obtained in datasets without the addition of contamination and mitochondrial DNA, indicating that this tool may be sensitive to these additions.

The number of reconstructed bases followed the same trend observed in datasets 1 to 8 and most programs reconstructed between 50,000 bp and 250,000 bp per dataset, on average. However, the number of reconstructed bases increased by 20.7% in relation to the previous group of datasets, which may indicate that some tools reconstructed part of the mitochondrial DNA and contamination present in the samples.

In terms of the average scaffold length, PEHaplo, QuRe, SSAKE, and VirGenA produced scaffolds with an average length of less than 1,000 bp. In contrast, IRMA, QVG, TRACESPipe and V-pipe were able to reconstruct the genomes using the expected number of scaffolds and output scaffolds with over 25,000 bp. Overall, the average length of the scaffolds increased 2.0% in relation to the previous set of datasets.

The ratio of SNPs in relation to the number of bases reconstructed decreased by 11.7%, but followed the same trend observed in the previous group of datasets.

To assess the effect of the addition of contamination and mitochondrial DNA on the results obtained, datasets 5, 13, 57 and 58 were considered. The datasets have a read length of 150 bp, a depth coverage of 20x and the same viral composition (B19V, HPV, MCPyV and VZV), differing only on whether or not contamination and/or mitochondrial DNA were added. Dataset 5 contained no contamination or mitochondrial DNA, dataset 57 contained contamination, dataset 58 contained mitochondrial DNA, and dataset 13 contained both contamination and mitochondrial DNA. Figure 4 shows the effects that contamination and mitochondrial DNA have on the performance of the reconstruction programs.

**Fig. 4.**
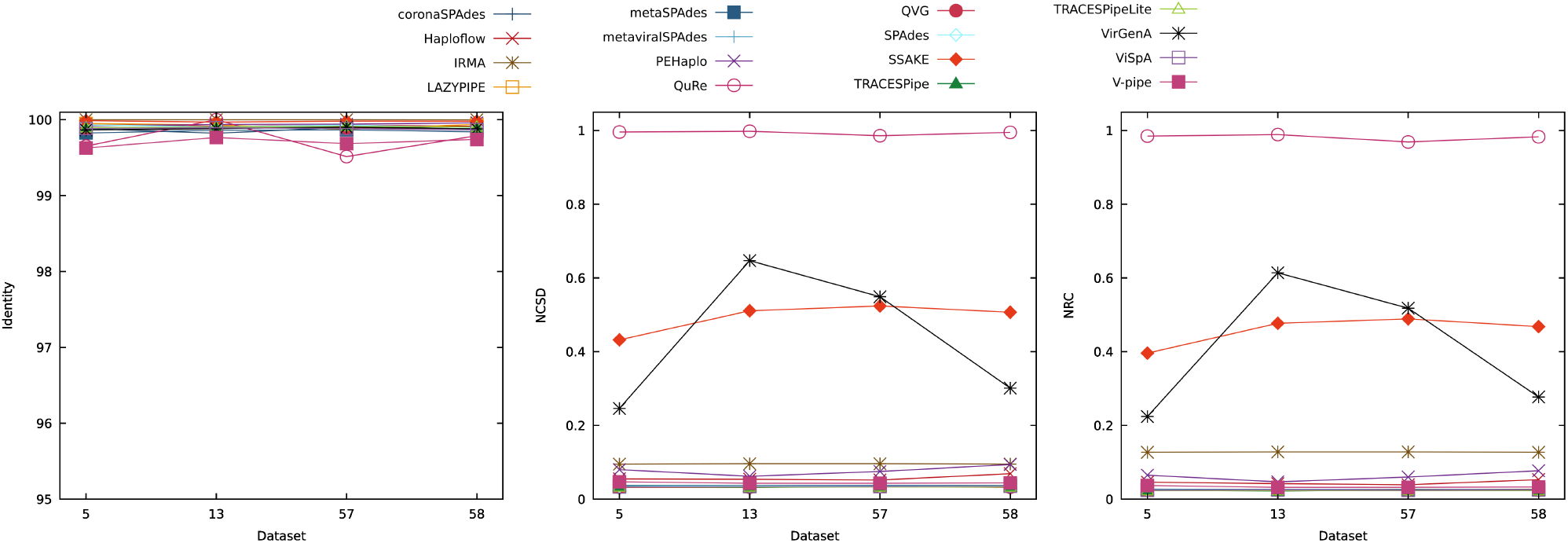
Comparison of the performance of the reconstruction programs according to the identity, NCSD and NRC for datasets DS5, DS13, DS57 and DS58, which show the effects that contamination and mitochondrial DNA have on the performance of the reconstruction programs. Optimal identity values are close to 100, while lower NCSD and NRC values indicate better results.

All reconstruction programs were able to reconstruct the four datasets considered, except for metaviralSPAdes which output no results.

Regarding the identity, the tools displayed a stable performance throughout all datasets considered. The values of the identity ranged between 99.5 and 100 and the most variance in performance was observed in the tool QuRe. In this metric, the best results overall were obtained for dataset 13, followed by datasets 58, 5 and 57.

In terms of the NCSD and NRC, the performance was best overall for dataset 5, which did not contain contamination or mitochondrial DNA. The next best results were obtained for dataset 58, which contained just mitochondrial DNA, followed by dataset 57, with just contamination and the worst performance, on average, was obtained in dataset 13, containing both contamination and mitochondrial DNA. These plots also show that VirGenA has the most variation in its performance, indicating susceptibility to these additions to the datasets. Additionally, QuRe obtained the least favourable performances across all datasets considered.

In contrast to the results obtained using the NCSD and NRC, the tools overall reconstructed the most bases from dataset 13, indicating that some tools may have reconstructed the contamination and mitochondrial DNA added. This can also be observed in datasets 57 and 58, in which most tools also increased the number of reconstructed bases in relation to dataset 5, which has no contamination or mitochondrial DNA added. Exceptions to this behaviour can be observed in the performances of QVG, QuRe and VirGenA. QVG had a stable performance in datasets 5, 13 and 58, decreasing its performance in dataset 57. QuRe maintained a stable performance throughout all datasets considered, however, it did not reconstruct many bases in any of the datasets considered. VirGenA is susceptible to the additions present, as previously discussed, and its performance declines especially when contamination or contamination and mitochondrial DNA are added to the datasets.

In terms of the average length of the scaffolds, it was observed that the scaffolds reconstructed in dataset 13 had the greatest length on average, followed by dataset 58, dataset 5 and lastly, dataset 57. The best results were obtained in dataset 13 which may be explained by some tools reconstructing contamination and mitochondrial DNA, as previously indicated.

The ratio of SNPs in relation to the number of bases reconstructed is generally low and the programs had little variance in their performance throughout the datasets considered.

It is important to note that when human samples are sequenced, it is unrealistic for only viral genomes to be present in a sample. Therefore, analysing datasets that contain only viral genomes does not accurately represent real-life scenarios. Taking this aspect into consideration, the following analyses were done using datasets containing both mitochondrial DNA and contamination.

#### Percentage of SNPs and depth coverage

To analyse the effects that the percentage of SNPs and depth coverage can have on the performance of the reconstruction programs considered, several datasets containing varying percentages of SNPs, ranging between 0% and 15%, were considered. The depth coverage of the datasets varied between 2x and 40x and all datasets contained both mitochondrial DNA and contamination.

SNPs are single nucleotide bases that are not equal to the correspondent base in another genome. In the reconstruction process, SNPs correspond to ambiguities in the genome, which make it more difficult for a dataset to be reconstructed accurately.

The datasets considered for this analysis were datasets 17 through 56, plus datasets 9, 10, 11, 13 and 16. Figure 5 represents the results obtained by the reconstruction programs when analysing datasets with depth coverage of 2x and 40x. The top three plots of Figure 5 represent the results obtained in terms of identity, NCSD and NRC with a depth coverage of 2x. At this depth coverage, each part of the genome is present on a dataset, on average, only two times, which makes the reconstruction process difficult.

**Fig. 5.**
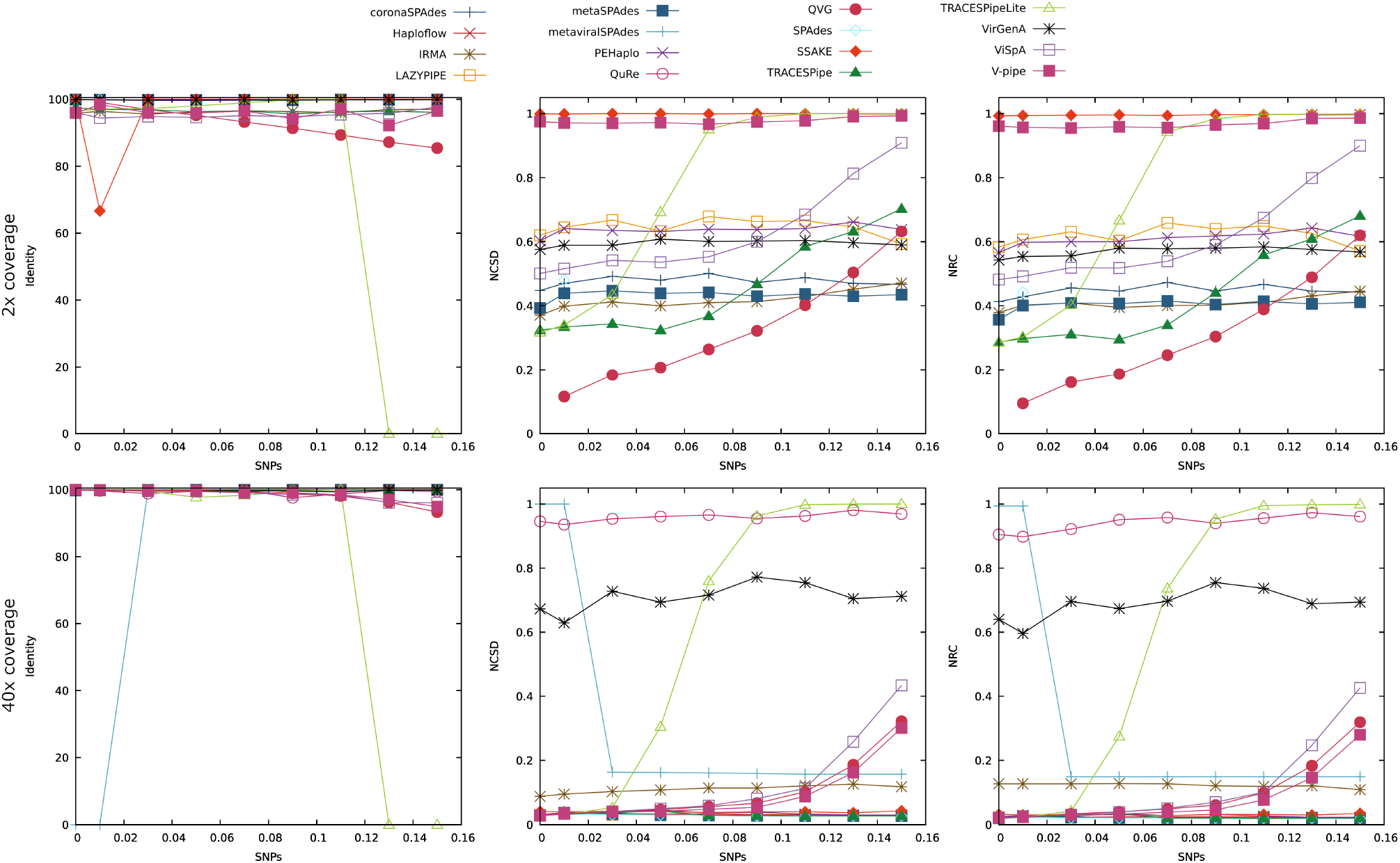
Comparison of the performance of the reconstruction programs according to the identity, NCSD and NRC for datasets DS17 to DS24 plus DS9 (2x coverage) and datasets DS49 to DS56 plus DS16 (40x coverage). The x-axis represents the ratio of SNPs added to the viruses in the datasets. Optimal identity values are close to 100, while lower NCSD and NRC values indicate better results.

It should be noted that metaviralSPAdes and QuRe were not able to reconstruct data from any of the datasets considered with depth coverage 2x. Haploflow reconstructed only the datasets with 13% or 15% of SNPs (datasets 23 and 24), QVG reconstructed all datasets except dataset 17, and SPAdes was only able to recover dataset 9.

The identity correlated inversely with the percentage of SNPs present in the datasets. This can be explained as the identity is calculated based on the genome fragments that can be aligned, which declines as the quality of the reconstruction made decreases. The drop in performance suffered by SSAKE to 66.7% in dataset 9 can be explained as in one of the reconstructed files dnadiff was not able to align any fragment of the reconstructed genomes to the viral genomes. This shows that SSAKE does not have a deterministic behaviour. The drop observed in TRACESPipeLite’s performance when the percentage of SNPs contained in the datasets was 13% or over was due to the pipeline only reconstructing mitochondrial DNA in those datasets.

The NCSD and NRC showed two trends in the performance of the reconstruction tools: either it was relatively stable or it decreased as the percentage of SNPs increased. The NCSD and NRC confirmed the drop in performance of TRACESPipeLite, as in the datasets where the identity drops to 0, the NCSD and NRC have values close to 1. Haploflow, SSAKE and V-pipe had a lower performance across all datasets reconstructed based on the NCSD and NRC as they did not reconstruct many bases from the datasets (on average less than 10,000 bp per dataset). QVG obtained better results in relation to the remaining programs until the percentage of SNPs reached 11%, from which point forward the best performance was obtained by metaSPAdes. It should be noted that the ratio of SNPs in relation to the number of bases reconstructed obtained by QVG followed an almost linear increase as the percentage of SNPs increased, indicating that this tool may be over-reliant on the reference genome when the depth coverage is low, making TRACESPipe, IRMA or metaSPAdes more reliable in this scenario. For the remaining tools, the ratio of SNPs in relation to the number of bases reconstructed was low.

Regarding the number of bases reconstructed by each of the programs, most programs had a stable performance. However, ViSpA reconstructed a significantly greater amount of bases than the other tools throughout most datasets, providing multiple genome sequences for a single virus. The drop in the number of bases reconstructed by ViSpA coincides with the decrease in its performance in terms of the NCSD and NRC. As previously mentioned, the tools Haploflow, SSAKE and V-pipe reconstructed fewer bases than the remaining tools, outputting, on average, less than 10,000 bp per dataset. When taking into consideration the number of bases reconstructed and the number of scaffolds produced, IRMA, TRACESPipe and QVG performed the best as they output the most bases using the expected number of scaffolds, considering there were four viral genomes in the datasets considered.

The average number of bases per scaffold reconstructed was either low or tended to lower as the percentage of SNPS contained in the datasets increased, with all programs except for QVG, TRACESPipe, ViSpA and IRMA outputting less than 10,000 bp per scaffold, on average.

The bottom three plots of Figure 5 show the results obtained in terms of the identity, NCSD and NRC with depth coverage of 40x, which is much higher than the one shown in the plots with 2x coverage. Thus the tools have to handle a greater amount of data, however, they have less difficulty in discovering a consensus sequence as there are, on average, forty copies of every part of the genome. It is worth noting that with a depth coverage of 40x, all programs were able to reconstruct all datasets considered, except for QVG that could not reconstruct dataset 49. This shows a significant improvement in the tools’ capacity to reconstruct datasets in relation to when the depth coverage considered was 2x.

The identity decreased slightly with the increase in the percentage of SNPs present in the sample, confirming the previously obtained results. Significant changes to the values of the identity are observed in both metaviralSPAdes and TRACESPipeLite, caused by the reconstruction of exclusively contamination and/or mitochondrial DNA, which are not considered in this metric. Overall, the identity increased by 1.0% in relation to the average performance obtained using datasets with depth coverage of 2x.

Regarding the NCSD and NRC, most tools maintained their performance throughout the datasets considered. However, some tools decreased their performance as the percentage of SNPs increased, namely, TRACESPipeLite, ViSpA, QVG and V-pipe. With a depth coverage of 40x, coronaSPAdes, Haploflow, LAZYPIPE, metaSPAdes, PEHaplo, SPAdes and TRACESPipe obtained a consistently good performance, while QuRe and VirGenA had the least favourable performances. The performance of the reconstruction programs in these datasets has improved by 66.9% in terms of the NCSD and by 67.3% in terms of the NRC, in comparison to the average performance obtained with a depth coverage of 2x.

In relation to the number of bases reconstructed, most programs had a stable performance throughout the datasets considered. ViSpA, however, reconstructed a considerably greater amount of bases than the other tools, especially in the datasets with a percentage of SNPs of 7% or greater. This may be attributed to higher ambiguity in the genome, interpreted by ViSpA as different variants. The increase in the depth coverage has led to a rise in the number of bases reconstructed by 38.5% on average, in relation to the previous group of datasets.

The average length of the scaffolds was overall stable. However, it was lower for VirGenA, SSAKE, PEHaplo and QuRe, which have output scaffolds with an average length under 2,500 bp. With the increase in depth coverage from 2x to 40x, the average length of the scaffolds increased by 178.8%. This improvement is especially significant, as it was much greater than the increase in the number of reconstructed bases, indicating that the reconstructed genomes were less fragmented. The ratio of SNPs in relation to the number of bases reconstructed was low and stable for most programs, with the exception of QVG and V-pipe, the performance of which declined in datasets with over 9% of SNPs added. The ratio of SNPs in relation to the number of bases reconstructed decreased by about 81.6% in relation to the datasets with 2x depth coverage.

Overall, as the percentage of SNPs increased, the performance of the reconstruction tools tended to decline, suggesting that higher levels of ambiguity make the reconstruction more difficult. Additionally, the performance of the programs considered improved at higher depth coverages. These findings can be explained as lower depth coverages mean that every part of the genome is present on the sequenced files, on average, fewer times, which makes the reconstruction process harder due to the reduced amount of data available. In addition to lower performance, low-depth coverages can affect the reconstruction programs’ ability to reconstruct genomic sequences.

SPAdes, coronaSPAdes, LAZYPIPE and TRACESPipe maintained constant performances at increased SNPs and depths higher than 5x. In scenarios of low-depth coverage and low percentage of SNPs, good performances were obtained with TRACESPipe and QVG, although QVG may be over-reliant on the reference genome provided. In scenarios with low-depth coverage and a high percentage of SNPs, better performance was obtained with metaSPAdes.

#### Read length

To assess how different read lengths affect the reconstruction process, datasets 13, 61 and 62 were considered. These datasets have a depth coverage of 20x, 1% of SNPs and contain both mitochondrial DNA and contamination. The reads included in dataset 61 have a length of 75 bp, in dataset 13 the reads have a length of 150 bp and in dataset 62 the reads have a length of 250 bp, all of which are considered to be short reads. Figure 6 represents the results obtained in terms of the identity, NCSD and NRC in these datasets.

**Fig. 6.**
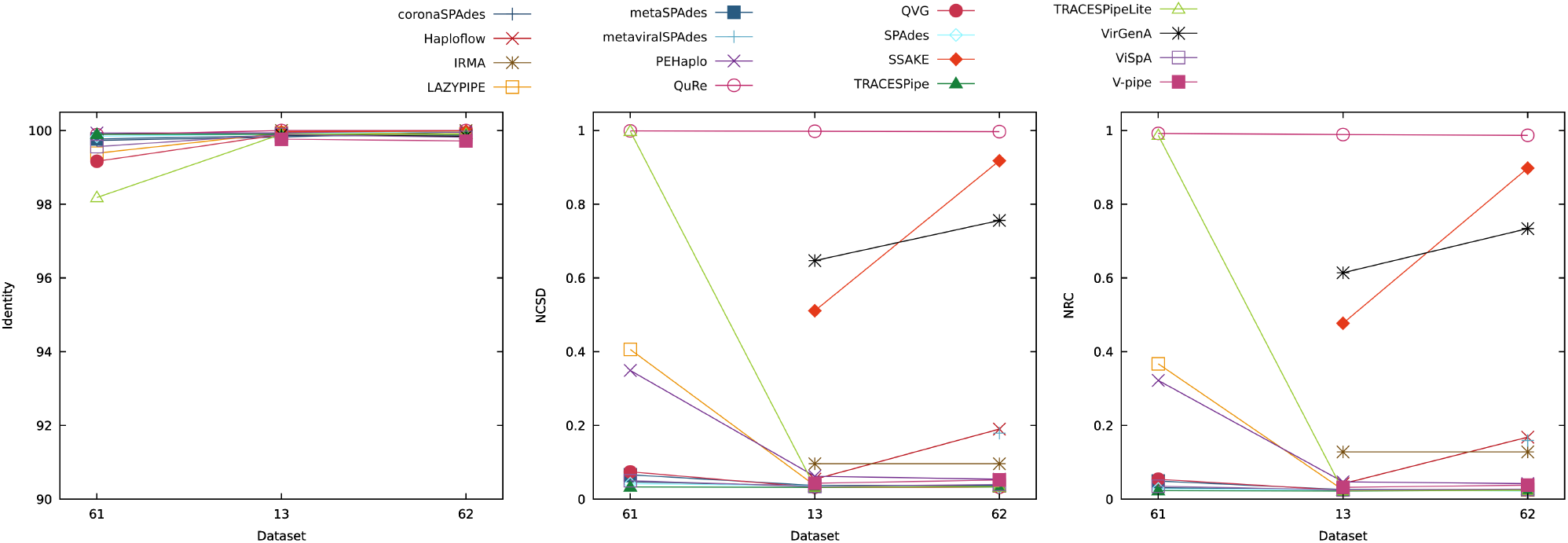
Comparison of the performance of the reconstruction programs according to the identity, NCSD and NRC for datasets 13, 61 and 62, which show the effects that different read lengths have on the performance of the reconstruction programs. Optimal identity values are close to 100, while lower NCSD and NRC values indicate better results.

The effect of the read length on the performance of the programs was evidenced by the fact that at 75 bp (dataset 61) only 10 out of the sixteen tools were able to reconstruct the genomes. In contrast, at 150 bp, all reconstruction programs except metaviralSPAdes were able to reconstruct the dataset, and at 250 bp the dataset was reconstructed by all tools.

In terms of the identity, the performance was generally constant throughout the datasets considered, varying between 98% and 100%. The best overall performance was obtained in dataset 62, with a read length of 250 bp, the second-best average performance was obtained in dataset 13, with reads of 150 bp, and the least favourable overall performance was obtained in dataset 61 with reads with 75 bp of length.

Regarding the NCSD and NRC, the reconstruction programs either improved or maintained their performance from dataset 61 to dataset 13, and maintained or decreased their performance between datasets 13 and 62. Regarding these metrics, the best overall performance was obtained for dataset 13, followed by dataset 62 and lastly, dataset 61.

The number of reconstructed bases increased or leveled from dataset 61 to dataset 13, with the exception of ViSpA, which reconstructed more bases in dataset 61, perhaps because it found the dataset to be more ambiguous. From datasets 13 to 62, most tools maintained their performance or decreased the number of bases reconstructed. Regarding this metric, the best performance was obtained when using the dataset with 150 bp, followed by the dataset with reads of 75 bp and lastly, the dataset with reads of 250 bp.

The average scaffold length also increased or stabilized between datasets 61 and 13, with the exception of ViSpA and SPAdes, whose performance decreased. Between datasets 13 and 62, most tools maintained their performance, with Haploflow and LAZYPIPE having the most significant decreases and coronaSPAdes and SPAdes increasing the average scaffold length the most. Overall, the best results were obtained with dataset 62, followed by dataset 13 and then dataset 61.

The ratio between the number of SNPs and the number of bases reconstructed decreased between datasets 61 and 13, and leveled or increased between datasets 13 and 62. The best overall performances for this metric were observed for dataset 13, followed by datasets 62 and 61.

Although the performance was best overall for the dataset with reads of length 150 bp based on the NCSD, NRC, number of reconstructed bases, and ratio of SNPs in relation to the number of bases reconstructed, the best results regarding the identity and average length of the scaffolds were obtained in the dataset with read length 250 bp. This indicates that while the performance of the programs in the dataset containing reads with a length of 75 bp is surpassed by the performance in the remaining two datasets, there may not be a read length that is best for every metric considered. It should be noted that these tests were made on simulated datasets with a constant percentage of SNPs and that an increase in length read in real-life scenarios often leads to an increase in the percentage of sequencing errors (147). It should also be noted that the reads considered in these tests were all considered to be short reads and contained the same genomes and characteristics, except for the read length. This comparison may have different results if different read lengths, viral genomes or read characteristics are tested.

#### Viral composition

To study the effect that different viral compositions can have on the performance of the reconstruction programs, four different combinations of viruses were considered. The datasets 13, 63, 64 and 65 were considered for this analysis and the viruses contained in each dataset can be found in Supplementary Table 2. All of these datasets have a depth coverage of 20x, 1% of SNPs and a read length of 150 bp. Figure 7 illustrates the performances obtained based on the identity, NCSD and NRC for each dataset considered. All reconstruction programs reconstructed these datasets except for metaviralSPAdes which only output results for dataset 64.

**Fig. 7.**
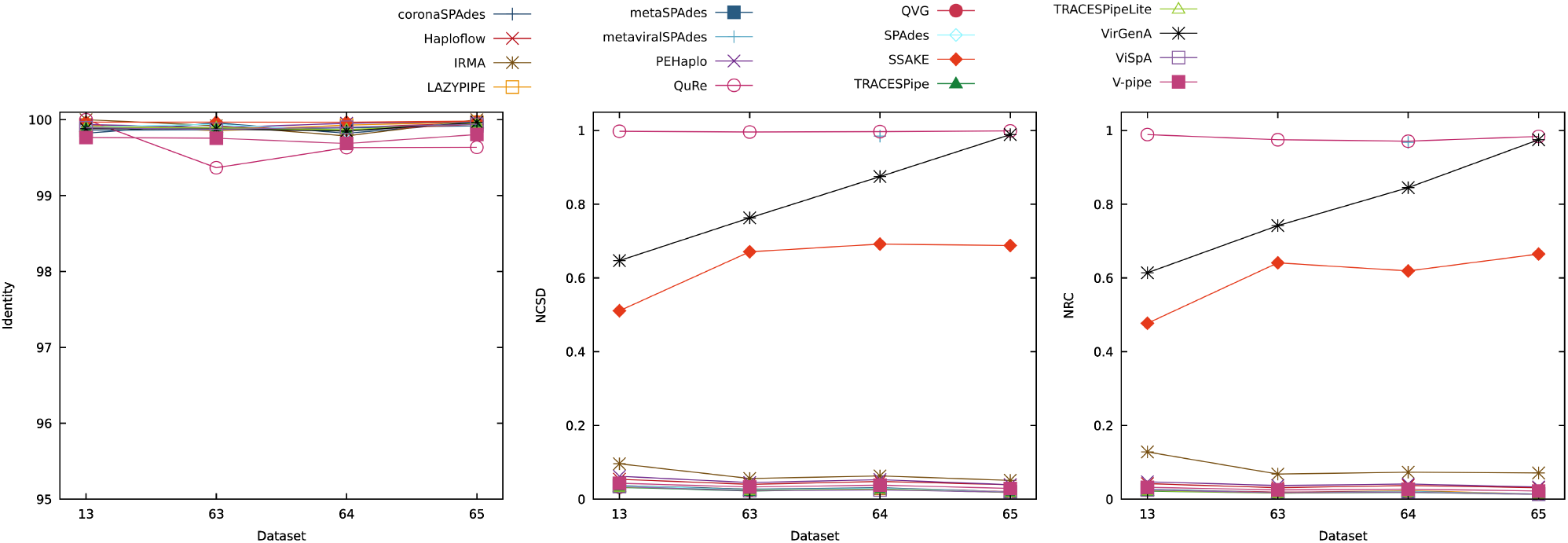
Comparison of the performance of the reconstruction programs according to the identity, NCSD and NRC for datasets 13, 63, 64 and 65, which show the effects that different viral compositions have on the performance of the reconstruction programs. Optimal identity values are close to 100, while lower NCSD and NRC values indicate better results.

Overall the identity had little variation, with values ranging between 99% and 100%.

Regarding the NCSD and NRC, the performance was overall stable, with the most variation belonging to SSAKE and VirGenA. The best overall results were observed for TRACESPipe, TRACESPipeLite, ViSpA, coronaSPAdes, QVG and SPAdes. Although QuRe had a stable performance throughout all datasets that performance was low as it reconstructed fewer bases than the remaining programs considered.

The number of reconstructed bases was greater for ViSpA, QVG and PEHaplo, reconstructing over 400,000 bp per dataset, on average. On the other hand, metaviralSPAdes and QuRe had a lower performance as they reconstructed, on average, less than 6,000 bp per dataset.

The average length of the scaffolds was greatest for QVG, TRACESPipe, ViSpA and V-pipe, all outputting scaffolds with, on average, over 50,000 bp. The reconstruction programs VirGenA, PEHaplo, SSAKE and QuRe obtained the least favourable performances in this metric averaging under 1,000 bp per scaffold.

Most reconstruction programs had little variance on the ratio of SNPs in relation to the number of bases reconstructed, which may indicate that under these scenarios there was generally little reliance on the reference genomes provided.

It should be taken into consideration that the datasets considered still have in common at least three viruses (B19V, HPV and VZV) and that different viral compositions and characteristics of the datasets may affect the results.

#### General comparisons

To evaluate the performance of each of the programs the number of datasets reconstructed by each program was calculated. In the context of this review, the number of datasets reconstructed is the count of datasets reconstructed by a tool, regardless of whether or not the results were obtained in a single execution cycle. This aspect is important as some tools, namely metaSPAdes, QuRe, SSAKE and VirGenA sometimes did not output results for a dataset in at least one of the execution cycles. Based on the number of datasets reconstructed by each program, we averaged the performance obtained in terms of the identity, NCSD, NRC, number of bases reconstructed, average scaffold length, ratio of SNPs in relation to the number of bases reconstructed, number of scaffolds generated, execution time and computational resources. Some of the results obtained for each reconstruction program are illustrated in Figure 8, whereas the remaining are available in the Supplementary Material.

**Fig. 8.**
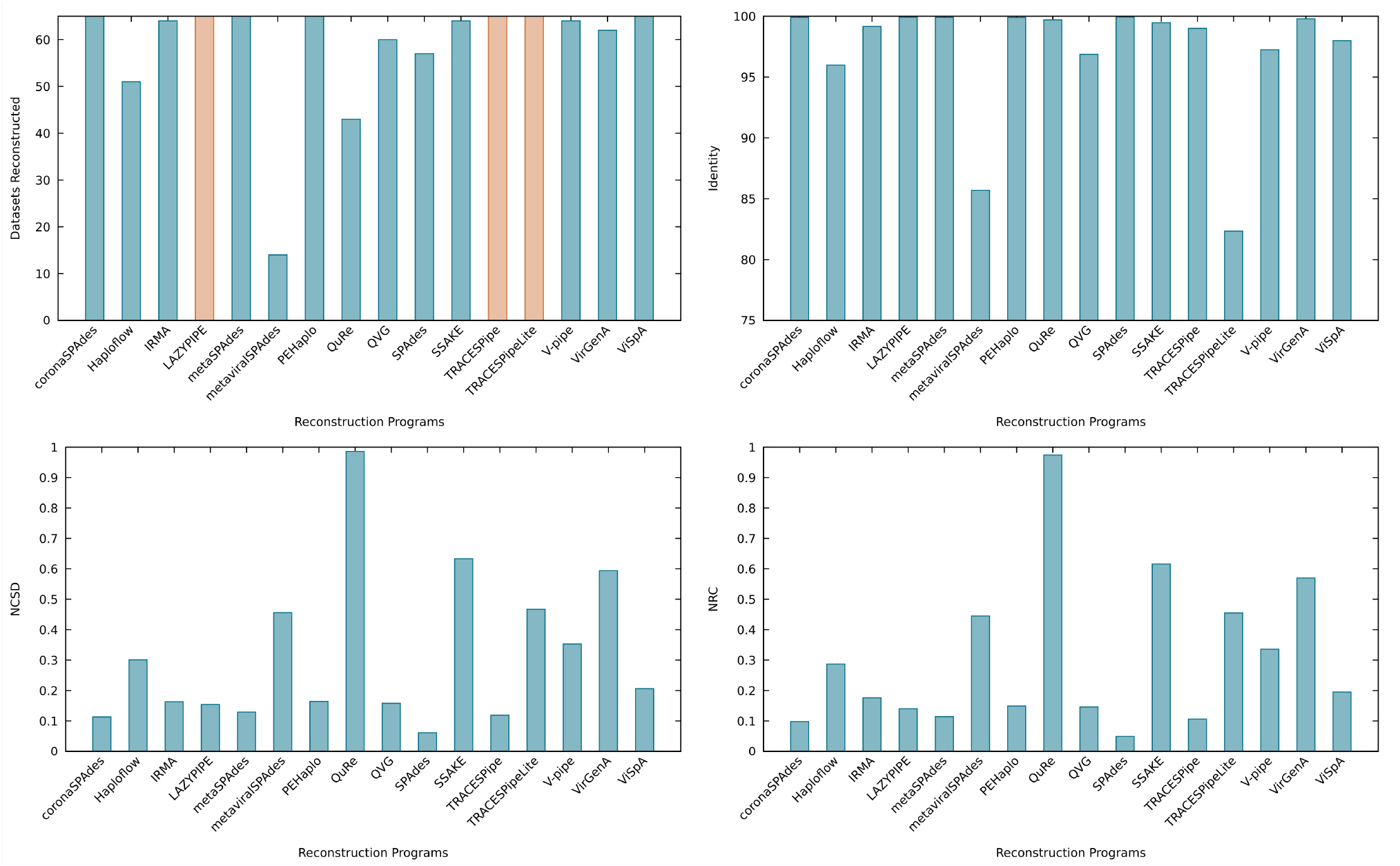
Comparison of the average performance of the reconstruction programs according to the number of datasets reconstructed, identity, NCSD and NRC. In the “Datasets Reconstructed” plot, the bars that appear in orange indicate that the corresponding pipeline does metagenomic classification. For the number of datasets reconstructed the results are best the closest they are to 65 (number of synthetic datasets considered). Optimal identity values are close to 100, while lower NCSD and NRC values indicate better results.

The coronaSPAdes, LAZYPIPE, metaSPAdes, PEHaplo, TRACESPipe, TRACESPipeLite and ViSpA were able to reconstruct all datasets. In contrast, Haploflow, metavi-ralSPAdes, QuRe and SPAdes often failed are reconstructing genomes in low-depth coverage scenarios and QVG did not output a result if no SNPs were added to the dataset.

The identity measures the correctness of the parts of the genome that could be aligned to the viral genomes contained in the datasets. Low identity values indicate that the parts of the genomes that could be aligned did not correspond to the reference and a value of zero means that no part of the genome could be aligned. In this metric, all reconstruction tools except for metaviralSPAdes and TRACESPipelite obtained an average performance above 90%. TRACESPipeLite is negatively impacted in this metric because although it is able to reconstruct parts of every dataset, for some it is only able to reconstruct mitochondrial DNA, which is not considered in this study, obtaining an identity of zero. The same phenomenon happened to metaviralSPAdes although the number of datasets reconstructed by this pipeline is lower.

The NCSD evaluates the dissimilarity of the reconstructed genomes against a reference. The programs that obtained the best results overall were SPAdes, coronaSPAdes, TRACESPipe and metaSPAdes, all of which obtained an average value of NCSD below 0.15. Conversely, QuRe obtained the least favourable performance, with an NCSD value nearing 1, which indicates that it either did not reconstruct much data from the datasets or that the data was reconstructed incorrectly. It should be noted that the average was made by dividing the sum of all NCSD values obtained by the number of datasets reconstructed by each tool, which penalizes tools that can reconstruct more datasets, albeit incorrectly or that only reconstruct parts of the genome that are not being taken into consideration, namely mitochondrial DNA or contamination.

The NRC corroborates the findings from the NCSD, as the best performances were obtained by SPAdes, coronaSPAdes, TRACESPipe and metaSPAdes. Similar to the NCSD, QuRe obtained the lowest performance in this metric with an average performance close to 1.

The overall number of bases reconstructed (excluding gaps) is a metric that should not be considered by itself as some programs may reconstruct bases that do not belong to viral genomes or may reconstruct the quasispecies spectrum, outputting several different genomic sequences for a single virus. Hence, the number of scaffolds generated should also be taken into consideration, as tools that reconstruct many bases and output an appropriate number of scaffolds (taking into consideration the number of viruses in the sample) correctly identified the genomes and were able to reconstruct them in one piece. Although ViSpA, PEHaplo, QVG, SPAdes, coronaSPAdes, metaSPAdes, LAZYPIPE, Haploflow, TRACESPipe and IRMA reconstructed on average over 140,000 bp per dataset, out of those only IRMA, QVG and TRACESPipe produced a suitable number of scaffolds (about 4 to 6) considering the number of viral genomes contained in the datasets. We observed that QVG, TRACESPipe, TRACESPipeLite and V-pipe had the most discrepancy between the number of bases considering and disregarding gaps, which indicates that these tools output scaffolds with the most gaps.

In terms of the average length of each scaffold (excluding gaps), VirGenA, PEHaplo, QuRe and SSAKE obtained a lower performance, each reconstructing scaffolds with, on average, less than 2,000 bp. On the other hand, the programs metaviralSPAdes, QVG, TRACESPipe and ViSpA were capable of outputting scaffolds with over 35,000 bp, on average. Regarding the ratio of SNPs in relation to the number of bases reconstructed, QVG obtained the least favourable performances out of the programs considered obtaining results over 0.02. Although this metric may perform ambiguous analyses, as it is based on alignments, it can be used as an indicator of the degree of reliance on the reference genomes for reconstruction programs that follow RB or HB methodologies.

The time needed for a program to reconstruct a dataset was, on average, less than 180 seconds. On average, metaviralSPAdes, SPAdes, SSAKE, LAZYPIPE, Haploflow and coronaSPAdes required under 10 seconds to reconstruct a dataset, whereas QuRe and VirGenA required over 700 seconds.

We analyzed the CPU usage and found that VirGenA and QuRe were the most resource-intensive tools, each requiring over 5 execution cores on average to reconstruct a dataset. On the other hand, SSAKE, Haploflow and ViSpA required the least amount of resources, requiring, on average, less than 1 execution core.

Regarding the maximum RAM used by the tools, QuRe, TRACESPipe and ViSpA utilised the most resources, each requiring, on average, over 6 GB to reconstruct a dataset, while coronaSPAdes, V-pipe, Haploflow and SSAKE required the least resources, each using under 0.15 GB of RAM.

To analyse the computational resources utilized by each reconstruction program taking into consideration the reconstruction performance, the metric *P* was introduced. The metric *P* is defined as

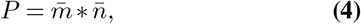

where 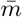 represents the average value obtained by the reconstruction program in the metric chosen (either the execution time, RAM usage or CPU usage) and the 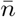 represents the average value of the NCSD for the reconstruction program considered. In this case, the NCSD acts as an attenuating factor to the metric chosen. The values of the *P* range between slightly above 0 to infinity, with lower values indicating a better performance.

Haploflow, LAZYPIPE, SPAdes and coronaSPAdes performed the best in terms of the weighted performance of the time, achieving a value under 2. On the other hand, the tools QuRe and VirGenA had a lower performance, both obtaining values over 400.

Regarding the weighted performance of the CPU, Haploflow, QVG, SPAdes, ViSpA and PEHaplo achieved the best performances, obtaining values under 30. Conversely, QuRe and VirGenA had less-than-ideal performances, obtaining values over 400.

Lastly, in terms of the weighted performance of the RAM, SSAKE, V-pipe, Haploflow, metaSPAdes, coronaSPAdes and SPAdes demonstrated the best performance with values under 0.1, while QuRe had the least satisfactory performance in this metric obtaining a value over 20.

Comparing the amount of computational resources utilized by reconstruction programs belonging to a reconstruction methodology (RF, RB, or HB), we observed that the programs that followed the RF methodology were overall the most efficient. The programs following the HB methodology were the second most efficient in terms of the execution time and RAM usage, while programs using the RB methodology were the second most efficient in regards to the CPU usage. These findings seem to contradict the description of the methodologies provided in the Section “Introduction”, however, it must be taken into consideration that some programs, namely the QuRe and VirGenA tend to consume considerably more resources than the remaining and that some tools provide additional outputs, which may have affected the results obtained.

### Performance in real datasets

In order to measure the performance of each reconstruction program in real-life scenarios, six real datasets were considered. In these scenarios, there is no detailed information on the contents of the sample, and therefore, in addition to the reconstruction process, it was necessary to classify the datasets using FALCON-meta (67). FALCON-meta provides information on the viruses present in the sample and predicts which are the most suitable references to be used in the reconstruction process. The references were extracted from a viral database, included in the benchmark, and are used by programs that follow the RB and HB methodologies and that require references to be provided by the user. When analysing these datasets with a value of top of similarity set to 8,000, FALCON-meta found a minimum of 26 and a maximum of 36 suitable references per dataset, averaging 30.8 references per dataset. The viruses found in each dataset can be found in Table 133 of the Supplementary Material.

To assess the performance of the reconstruction programs in these datasets, the metrics identity, NCSD, NRC and the ratio of SNPs in relation to the number of bases reconstructed could not be used, as these metrics require the true composition of the datasets to be known (requires a gold standard). Hence, the evaluation was based on other metrics available, namely the number of datasets reconstructed, the number of bases reconstructed, the number of scaffolds generated, the minimum, maximum and average length of the scaffolds and the computational resources used by each of the programs.

Some of the results obtained using these datasets are available in Figure 9 while the remaining are in the Supplementary Material.

**Fig. 9.**
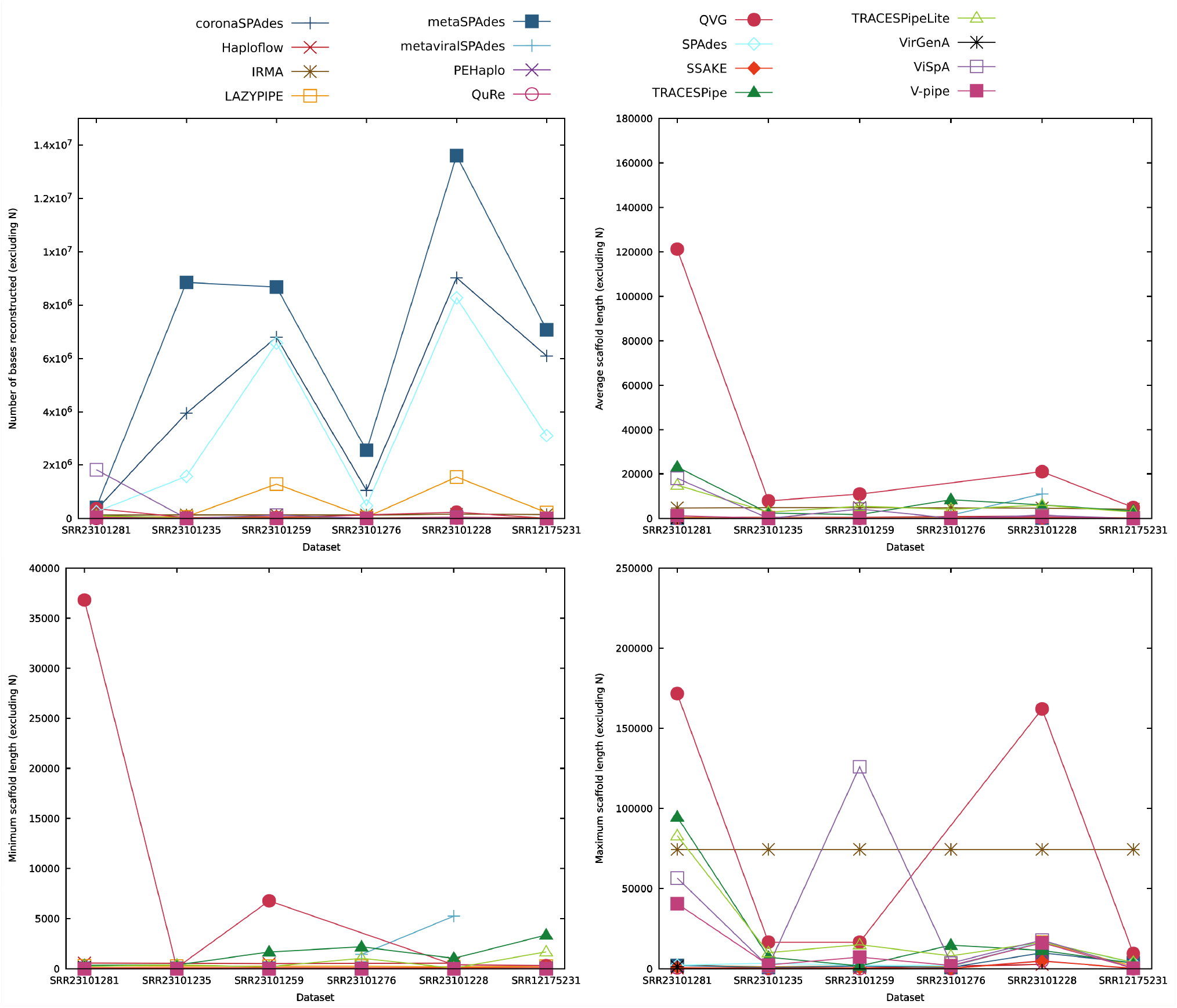
Comparison of the performance of the reconstruction programs according to the number of reconstructed bases and the average, minimum and maximum number of bases per scaffold generated, excluding non-reconstructed bases (N) in real datasets. For these metrics, higher values indicate a better performance.

All of the reconstruction programs except for PEHaplo were able to reconstruct at least one of the real datasets provided. Additionally, coronaSPAdes, IRMA, LAZYPIPE, metaSPAdes, QuRe, SPAdes, SSAKE, TRACESPipe, TRACESPipeLite, V-pipe and ViSpA were able to output genome sequences for all datasets provided.

Regarding the number of bases reconstructed (excluding gaps), metaSPAdes has output the most bases in every dataset considered except SRR23101281, where the best performance was obtained by ViSpA. In these datasets, metaSPAdes, coronaSPAdes and SPAdes reconstructed the most bases, outputting, on average, over 3,000,000 bp per dataset. The average length of the scaffolds was especially great for the tool QVG, which has obtained a better performance in datasets SRR23101281, SRR23101235, SRR23101259, SRR23101228 and SRR12175231. The dataset SRR23101276 was not reconstructed by QVG and the best performance was obtained by TRACESPipe. TRACESPipe, metaviralSPAdes, TRACESPipeLite and QVG had the best performances in this metric and output on average over 5,000 bp per scaffold. Conversely, V-pipe, LAZYPIPE, VirGenA, SPAdes, coronaSPAdes, SSAKE, metaSPAdes and QuRe obtained a lower performance in this metric, outputting scaffolds with an average length under 500 bp.

In terms of the minimum length of the scaffolds, QVG performed better than the remaining programs in datasets SRR23101281 and SRR23101259. In dataset SRR23101235 the best performance was obtained by both TRACESPipe and TRACESPipeLite. TRACESPipe also obtained the best performances in datasets SRR23101276 and SRR12175231. Additionally, metaviralSPAdes obtained the best results in this metric in dataset SRR23101228. Overall, the best reconstruction programs regarding this metric were QVG, TRACESPipe and metaviralSPAdes with a minimum scaffold length above 1,000 bp, on average.

Regarding the maximum length of the scaffolds, QVG obtained the best results in datasets SRR23101281 and SRR23101228, IRMA had the best performance in datasets SRR23101235, SRR23101276 and SRR12175231, and ViSpA in dataset SRR23101259. Overall, the programs that obtained the best performances in this metric were QVG, IRMA and ViSpA, whose maximum length scaffolds had, on average, over 30,000 bp.

Using real datasets without a gold standard, it is complex to accurately determine how many different viral genomes are contained in the dataset, and as such, the exact number of scaffolds that should have been generated by each of the programs is unknown. On average the reconstruction programs output about 4,870 scaffolds per dataset, with coronaSPAdes, LAZYPIPE, metaSPAdes and SPAdes outputting an average of over 10,000 scaffolds per dataset, while the remaining programs generated, on average, less than 1,500 scaffolds per dataset.

Regarding the reconstruction time, the most efficient programs were coronaSPAdes, Haploflow, TRACESPipeLite and LAZYPIPE, which reconstructed the datasets in under 2 minutes, on average, and the least efficient programs were QuRe and VirGenA, which required, on average, over 4.5 hours to reconstruct a dataset.

In terms of the CPU usage, Haploflow and SSAKE required, on average, less than a core to execute, while VirGenA, QuRe and IRMA spent the most resources, using on average over 6 execution cores.

Regarding the maximum RAM used, SSAKE, IRMA and V-pipe were the most efficient tools, using less than 1 GB of RAM, whereas the most resource intensive tool in this metric was QuRe, requiring on average, over 34 GB of maximum RAM.

## Discussion

Every day the number of genomic samples sequenced is rising, which makes the reconstruction of the sequenced genomes and the evaluation of the programs that reconstruct them increasingly important.

Throughout the synthetic datasets analysed, we observed that the addition of contamination and mitochondrial DNA can have a negative effect on the performance of the reconstruction programs. We also observed that the performance of the reconstruction programs generally deteriorated at low coverage and at higher percentages of SNPs in the sample. Furthermore, we showed that different values of read length and different viral compositions can affect the performance of the reconstruction programs.

Regarding the metrics considered, the identity was a good indicator of whether or not a dataset was reconstructed. However, the NCSD and NRC were better at describing the performance of the reconstruction programs. We found that the NCSD and NRC were inherently linked, as their performance is highly consistent with each other and both rely on compression. Additionally, we found that the number of reconstructed bases is more informative when combined with the number of scaffolds reconstructed, as that provides information, not only about the quantity of genome that was reconstructed, but also the degree of fragmentation of the reconstructed genome.

In terms of the computational resources, it was estimated that RB methods were the most efficient, followed by RF methods, and lastly, HB methods. In reality, on average, tools that followed the RF methodology were found to be the most efficient in terms of execution time, maximum RAM usage and CPU usage. Pipelines based on the HB methodology achieved the second-best results in terms of execution time and maximum RAM usage, while the programs using the RB methodology achieved the second-best results for the CPU usage and the least favourable results in the remaining two metrics. This may be due to other tasks that the programs that follow the RB and HB methodologies perform, namely analysing and plotting the results obtained.

Employing authentic datasets facilitated the comparison of the performance of the reconstruction programs within reallife scenarios. However, it is crucial to note the absence of a definitive gold standard in this context, rendering the findings merely indicative. A potential method to benchmark such datasets involves synthesizing a virus, integrating it into a sample, and subsequently subjecting it to sequencing and computational analysis. This approach would yield the complete viral sequence, yet it is essential to acknowledge that sequencing processes may introduce errors. Analysis pipelines might interpret these errors as variants unless guided by explicit instructions to handle low-quality sequencing data.

While it can be argued that high coverage depth sequencing minimizes uncertainties, challenges persist at smaller depth coverages. Even with the proposed methodology, complete certainty in the benchmarks for real datasets remains elusive.

Existing pipelines and tools rely on rigid or adaptable parameters, which can significantly impact their accuracy. For instance, employing an aligner with high sensitivity differs markedly from one with lower sensitivity, thereby influencing the quality of the reconstructed sequence. Throughout this review, default parameters or settings conducive to reasonable processing times were predominantly employed. Regrettably, benchmarking these pipelines under various parameter sets becomes impractical without a substantial increase in computational resources. Such an expansion could easily reach an order of magnitude when exploring diverse parameter combinations. It is imperative to comprehend that altering the parameters of these tools and pipelines can markedly influence the accuracy and, consequently, the outcomes presented in this article.

Considering the reconstruction programs and the parameters used, there is no reconstruction program that is better than the remaining for all different scenarios and across all metrics.

When the datasets had low coverage and a low percentage of SNPs, the programs TRACESPipe and QVG obtained better performances in terms of the correctness of the reconstruction (NCSD and NRC) and the length of the scaffolds produced, although QVG may be over-relying on the reference genomes provided. With a low-depth coverage and a high percentage of SNPs, metaSPAdes had better performance according to the NCSD and the NRC, however, it output significantly shorter scaffolds.

Using datasets with a higher depth coverage (at least 5x), the reconstruction programs SPAdes, coronaSPAdes, LAZYP-IPE and TRACESPipe should be considered as had better performances considering the correctness of the genome. In terms of the fragmentation of the genome output, TRACESPipe reconstructs the expected number of scaffolds (4), whereas the remaining tools output considerably more scaffolds, especially with depth coverages of 5x and 10x.

If there is a preference for tools that output long scaffolds, metaviralSPAdes, QVG and TRACESPipe should be considered as they reconstructed few scaffolds with significant average length. It should be noted that although metaviralSPAdes outputs long scaffolds, the tool reconstructed only some of the datasets considered.

If the execution time is a priority, the tools coronaSPAdes and LAZYPIPE should be considered as they have reconstructed at least part of each dataset, had a low average reconstruction time in both real and synthetic datasets, and obtained some of the best values in terms of the the weighted time performance.

In cases where computational resources are limited, Haploflow, SSAKE, and V-pipe should be considered as they had the lowest CPU and RAM requirements.

We have successfully surveyed reconstruction programs that focused on the reconstruction of viral genomes, summarized their methodology and highlighted their characteristics. Additionally, we created a publicly available benchmark capable of automatically installing all programs necessary for its execution, as well as reconstructing viral genomes and evaluating the performance of each tool for datasets with and without a gold standard. Furthermore, we provide scripts capable of generating synthetic datasets and retrieving real datasets from the NCBI. Using the benchmark, the tools were executed using both real and synthetic datasets and the results were provided and described according to each of the characteristics tested. Lastly, we compared the performances of the reconstruction programs as a whole and provided recommendations about what tools should be used in given scenarios.

## Conclusions

Although the fast and accurate reconstruction of human viral genomes is fundamental for biological, medical and forensic applications, identifying the best assembly tool is challenging.

In this paper, we surveyed some of the existent viral genome reconstruction methods and identified some features, similarities, and dissimilarities between these tools. Moreover, we provided a reconstruction benchmark and evaluated the reconstruction process in sixty-five synthetic datasets and six real datasets.

The reconstructing programs were evaluated based on the correctness of the reconstruction process, using the metrics identity, NCSD, NRC, number of SNPs and the ratio of SNPs in relation to the number of bases reconstructed. Also, some metrics were used to evaluate the genomes and scaffolds reconstructed, namely the number of bases reconstructed, the number of scaffolds output, and the minimum, maximum, and average number of bases per scaffold. Lastly, the execution time and computational resources (RAM and CPU usage) needed to execute the reconstruction programs were evaluated using both weighted and unweighted methods. This methodology is publicly available and is flexible to the augmentation of search engines to increase the number of programs being considered. Moreover, it is possible to include other datasets so that the performance of each program can be tested under different conditions.

## Declarations

### List of abbreviations

B19V: human parvovirus B19
DNA: deoxyribonucleic acid
EBV: Epstein-Barr virus
GUI: graphical user interface
HB: hybrid
HCMV: cytomegalovirus
HiFi reads: highly accurate long reads
HHV6B: human herpesvirus 6B
HMMs: Hidden Markov models
HPV: human papillomavirus
HPyV7: human polyomavirus 7
MCPyV: Merkel cell polyomavirus
NCSD: Normalized Compression Semi-Distance
NRC: Normalized Relative Compression
PCR: polymerase chain reaction
RB: reference-based
RF: reference-free
SNPs: single-nucleotide polymorphisms
VZV: Varicella-Zoster virus

### Author’s Contributions

Conceptualization: KH, AS, MFP, DP; Methodology: MS, KH, AS, MFP, DP; Investigation: MS, MT, LP, DP; Visualization: MS, DP; Software: MS; Supervision: DP; Writing—original draft: MS, DP; Writing—review and editing: all authors.

## Acknowledgements

The authors wish to thank the Finnish Computing Competence Infrastructure (FCCI) for supporting this project with computational and data storage resources.

