## Supplementary Material for "Comparative evaluation of computational methods for reconstruction of human viral genomes"

#### 1 Characteristics of the datasets generated

Table S1: Main characteristics of the synthetic datasets used in the benchmark. The fold coverage and read length of the datasets were defined when simulating the sequencing process with ART [1]. The number of SNPs refers to the ratio of mutation added to each one of the viral sequences, which were added using GTO [2] and the substitution mutation refers to the ratio of mutations in the contamination genome, defined when using AlcoR [3].

| Identifier | Read length | Fold coverage | SNPs (GTO) | Substitution mutation (AlcoR) | Includes contamination genome | Includes mitochondrial genome |
|---|---|---|---|---|---|---|
| DS1 | 150 | 2 | 0.01 | – | No | No |
| DS2 | 150 | 5 | 0.01 | – | No | No |
| DS3 | 150 | 10 | 0.01 | – | No | No |
| DS4 | 150 | 15 | 0.01 | – | No | No |
| DS5 | 150 | 20 | 0.01 | – | No | No |
| DS6 | 150 | 25 | 0.01 | – | No | No |
| DS7 | 150 | 30 | 0.01 | – | No | No |
| DS8 | 150 | 40 | 0.01 | – | No | No |
| DS9 | 150 | 2 | 0.01 | 0.3 | Yes | Yes |
| DS10 | 150 | 5 | 0.01 | 0.3 | Yes | Yes |
| DS11 | 150 | 10 | 0.01 | 0.3 | Yes | Yes |
| DS12 | 150 | 15 | 0.01 | 0.3 | Yes | Yes |
| DS13 | 150 | 20 | 0.01 | 0.3 | Yes | Yes |
| DS14 | 150 | 25 | 0.01 | 0.3 | Yes | Yes |
| DS15 | 150 | 30 | 0.01 | 0.3 | Yes | Yes |
| DS16 | 150 | 40 | 0.01 | 0.3 | Yes | Yes |
| DS17 | 150 | 2 | – | 0.3 | Yes | Yes |
| DS18 | 150 | 2 | 0.03 | 0.3 | Yes | Yes |
| DS19 | 150 | 2 | 0.05 | 0.3 | Yes | Yes |
| DS20 | 150 | 2 | 0.07 | 0.3 | Yes | Yes |
| DS21 | 150 | 2 | 0.09 | 0.3 | Yes | Yes |
| DS22 | 150 | 2 | 0.11 | 0.3 | Yes | Yes |
| DS23 | 150 | 2 | 0.13 | 0.3 | Yes | Yes |
| DS24 | 150 | 2 | 0.15 | 0.3 | Yes | Yes |
| DS25 | 150 | 5 | – | 0.3 | Yes | Yes |
| DS26 | 150 | 5 | 0.03 | 0.3 | Yes | Yes |
| DS27 | 150 | 5 | 0.05 | 0.3 | Yes | Yes |
| DS28 | 150 | 5 | 0.07 | 0.3 | Yes | Yes |
| DS29 | 150 | 5 | 0.09 | 0.3 | Yes | Yes |
| DS30 | 150 | 5 | 0.11 | 0.3 | Yes | Yes |
| DS31 | 150 | 5 | 0.13 | 0.3 | Yes | Yes |
| DS32 | 150 | 5 | 0.15 | 0.3 | Yes | Yes |
| DS33 | 150 | 10 | – | 0.3 | Yes | Yes |
| DS34 | 150 | 10 | 0.03 | 0.3 | Yes | Yes |
| DS35 | 150 | 10 | 0.05 | 0.3 | Yes | Yes |
| DS36 | 150 | 10 | 0.07 | 0.3 | Yes | Yes |
| DS37 | 150 | 10 | 0.09 | 0.3 | Yes | Yes |
| DS38 | 150 | 10 | 0.11 | 0.3 | Yes | Yes |
| DS39 | 150 | 10 | 0.13 | 0.3 | Yes | Yes |
| DS40 | 150 | 10 | 0.15 | 0.3 | Yes | Yes |
| DS41 | 150 | 20 | – | 0.3 | Yes | Yes |
| DS42 | 150 | 20 | 0.03 | 0.3 | Yes | Yes |
| DS43 | 150 | 20 | 0.05 | 0.3 | Yes | Yes |
| DS44 | 150 | 20 | 0.07 | 0.3 | Yes | Yes |
| DS45 | 150 | 20 | 0.09 | 0.3 | Yes | Yes |

| | | | | | | |
|---|---|---|---|---|---|---|
| DS46 | 150 | 20 | 0.11 | 0.3 | Yes | Yes |
| DS47 | 150 | 20 | 0.13 | 0.3 | Yes | Yes |
| DS48 | 150 | 20 | 0.15 | 0.3 | Yes | Yes |
| DS49 | 150 | 40 | – | 0.3 | Yes | Yes |
| DS50 | 150 | 40 | 0.03 | 0.3 | Yes | Yes |
| DS51 | 150 | 40 | 0.05 | 0.3 | Yes | Yes |
| DS52 | 150 | 40 | 0.07 | 0.3 | Yes | Yes |
| DS53 | 150 | 40 | 0.09 | 0.3 | Yes | Yes |
| DS54 | 150 | 40 | 0.11 | 0.3 | Yes | Yes |
| DS55 | 150 | 40 | 0.13 | 0.3 | Yes | Yes |
| DS56 | 150 | 40 | 0.15 | 0.3 | Yes | Yes |
| DS57 | 150 | 20 | 0.01 | 0.3 | Yes | No |
| DS58 | 150 | 20 | 0.01 | – | No | Yes |
| DS59 | 150 | 30 | 0.03 | 0.3 | Yes | Yes |
| DS60 | 150 | 40 | 0.05 | 0.6 | Yes | Yes |
| DS61 | 75 | 20 | 0.01 | 0.3 | Yes | Yes |
| DS62 | 250 | 20 | 0.01 | 0.3 | Yes | Yes |
| DS63 | 150 | 20 | 0.01 | 0.3 | Yes | Yes |
| DS64 | 150 | 20 | 0.01 | 0.3 | Yes | Yes |
| DS65 | 150 | 20 | 0.01 | 0.3 | Yes | Yes |

Table S2: Viral sequences present in each of the generated datasets used in the benchmark.

| **Dataset** | **B19V** | **HPV** | **VZV** | **MCPyV** | **HPyV7** | **HHV6B** | **EBV** | **HCMV** |
|---|---|---|---|---|---|---|---|---|
| DS1 - DS62 | x | x | x | x | | | | |
| DS63 | x | x | x | x | | x | | |
| DS64 | x | x | x | | x | | x | |
| DS65 | x | x | x | x | | | | x |

### 2 Supplementary Figures and Tables

#### 2.1 Generated datasets

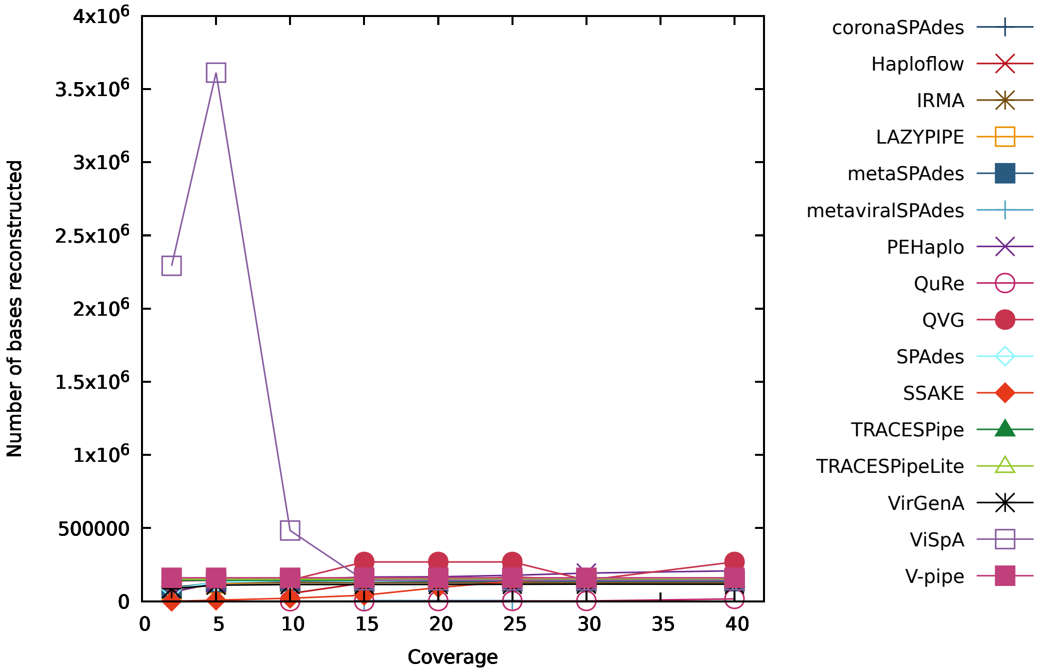


Figure S1: Figure comparing the performance of the reconstruction programs in terms of the number of bases reconstructed for datasets without contamination or mitochondrial DNA (DS1 to DS8).

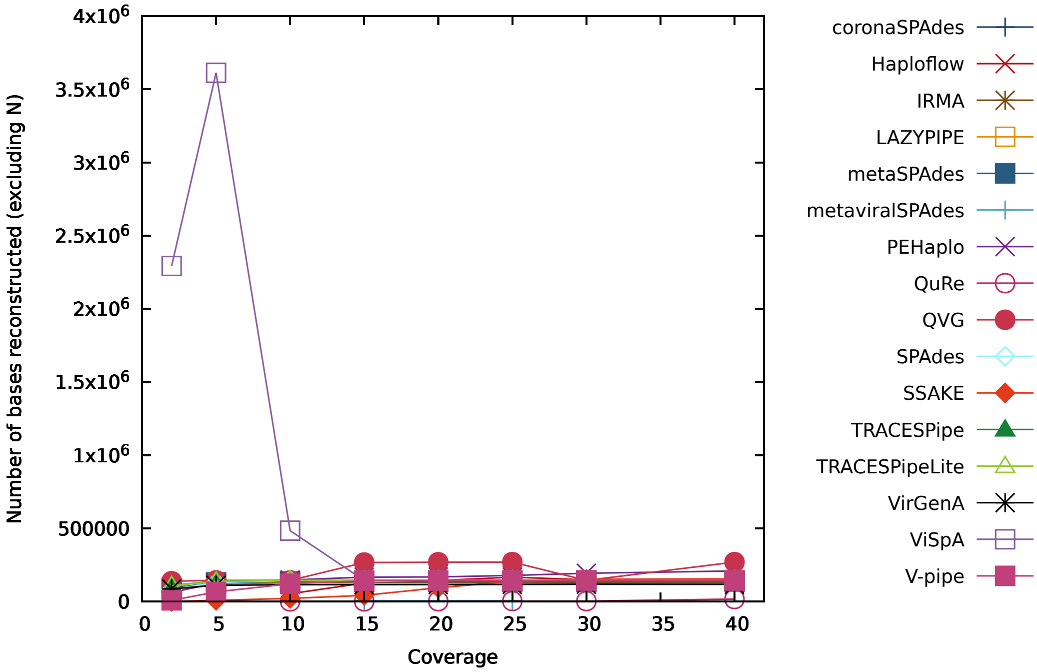


Figure S2: Figure comparing the performance of the reconstruction programs in terms of the number of bases reconstructed (excluding "N") for datasets without contamination or mitochondrial DNA (DS1 to DS8).

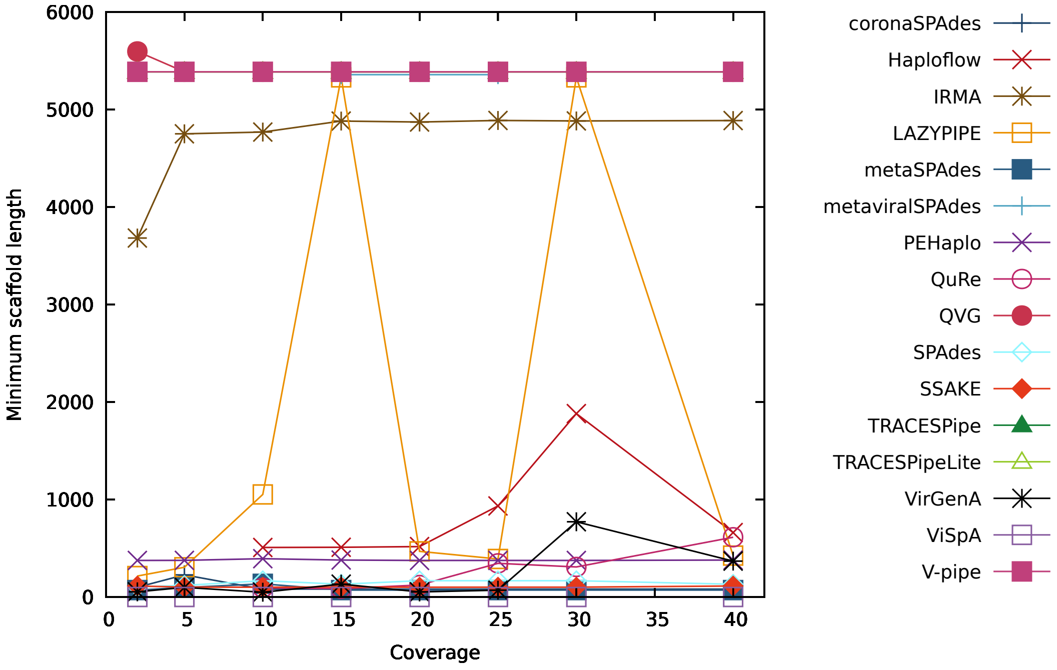


Figure S3: Figure comparing the performance of the reconstruction programs in terms of the minimum number of reconstructed bases per scaffold for datasets without contamination or mitochondrial DNA (DS1 to DS8).

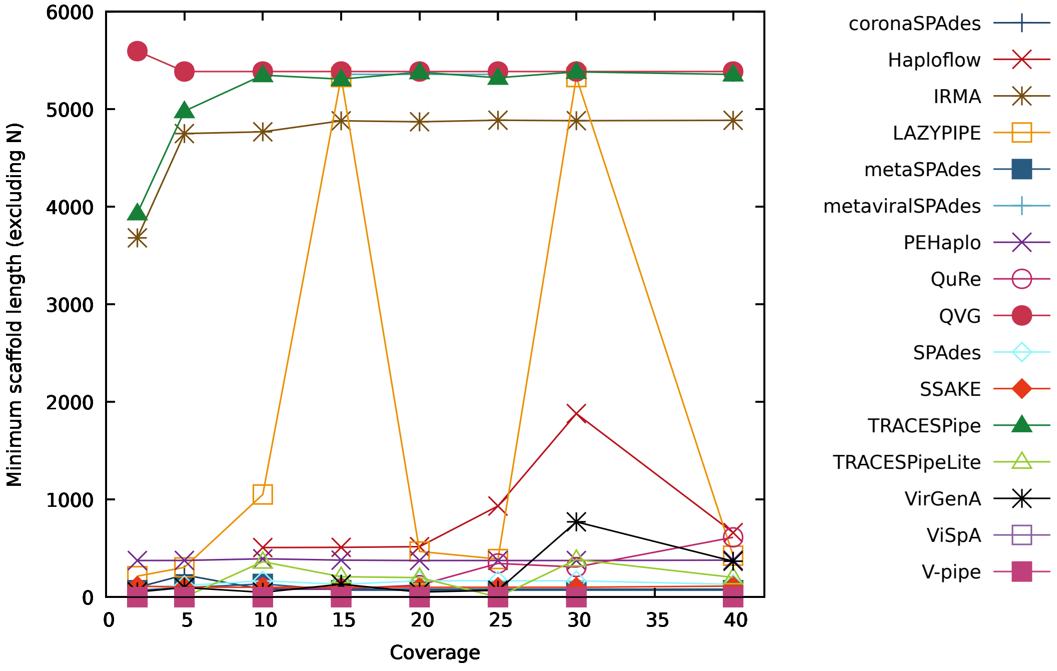


Figure S4: Figure comparing the performance of the reconstruction programs in terms of the minimum number of reconstructed bases per scaffold (excluding "N") for datasets without contamination or mitochondrial DNA (DS1 to DS8).

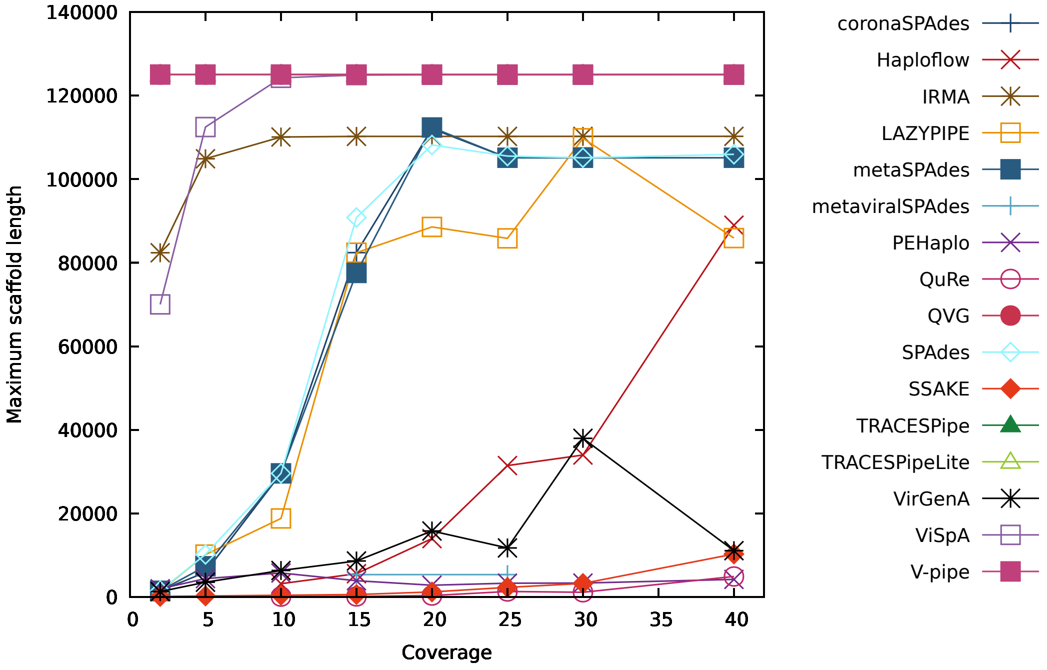


Figure S5: Figure comparing the performance of the reconstruction programs in terms of the maximum number of reconstructed bases per scaffold for datasets without contamination or mitochondrial DNA (DS1 to DS8).

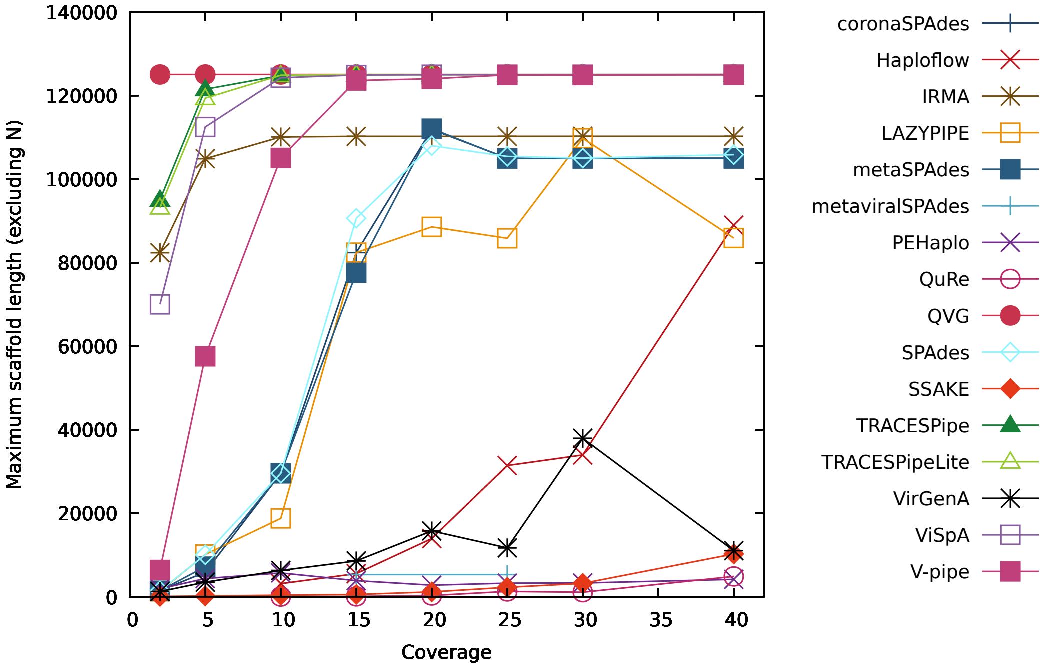


Figure S6: Figure comparing the performance of the reconstruction programs in terms of the maximum number of reconstructed bases per scaffold (excluding "N") for datasets without contamination or mitochondrial DNA (DS1 to DS8).

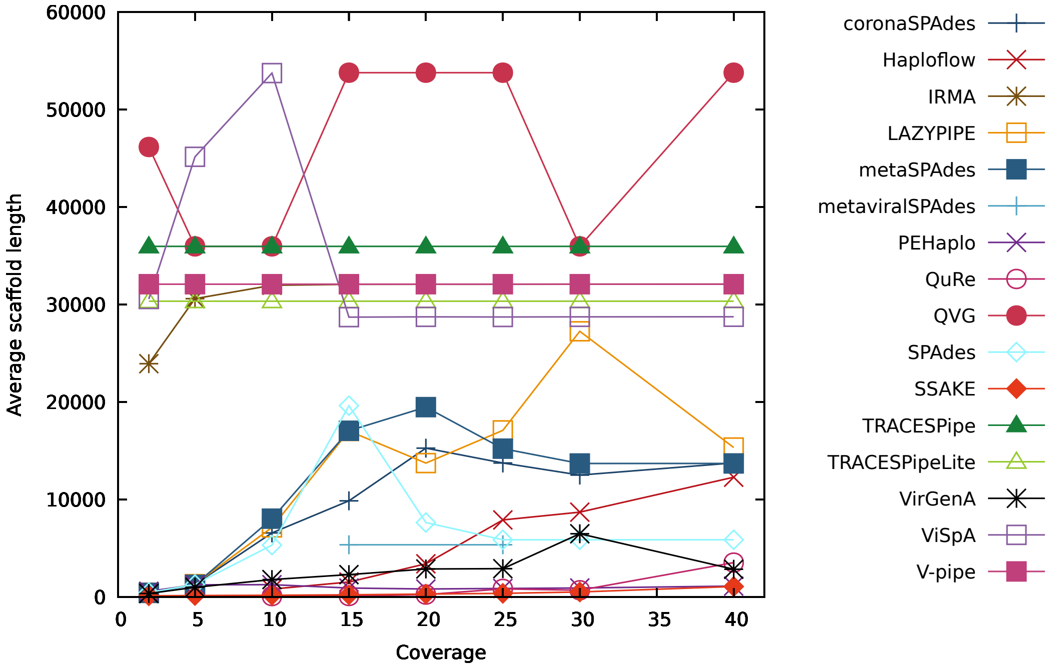


Figure S7: Figure comparing the performance of the reconstruction programs in terms of the average number of reconstructed bases per scaffold for datasets without contamination or mitochondrial DNA (DS1 to DS8).

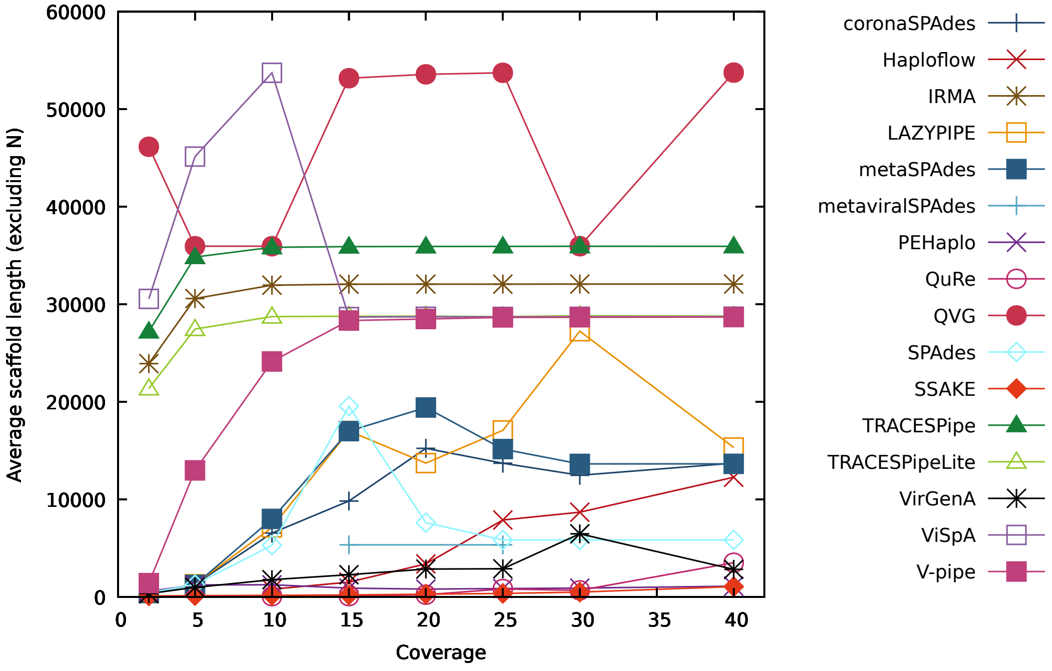


Figure S8: Figure comparing the performance of the reconstruction programs in terms of the average number of reconstructed bases per scaffold (excluding "N") for datasets without contamination or mitochondrial DNA (DS1 to DS8).

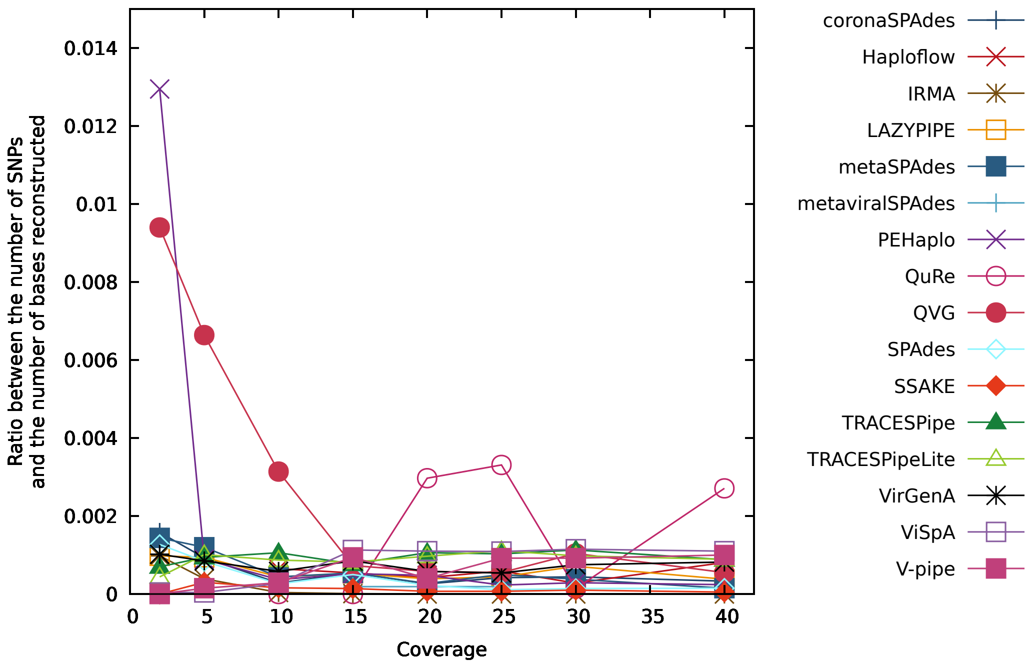


Figure S9: Figure comparing the performance of the reconstruction programs in terms of the ratio between the number of SNPs and the number of bases reconstructed for datasets without contamination or mitochondrial DNA (DS1 to DS8).

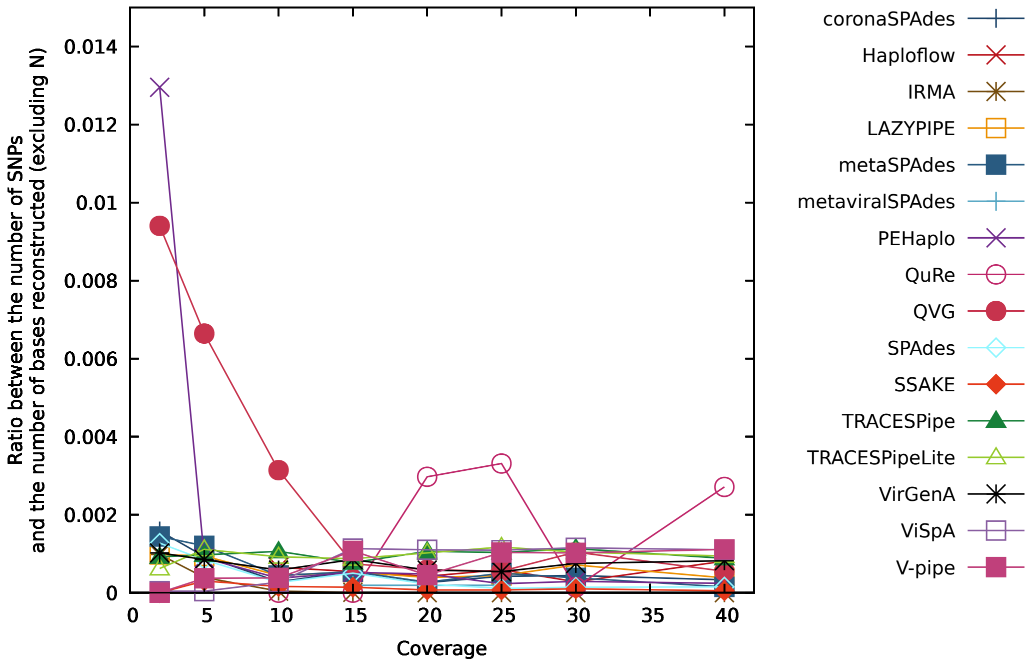


Figure S10: Figure comparing the performance of the reconstruction programs in terms of the ratio between the number of SNPs and the number of bases reconstructed (excluding "N") for datasets without contamination or mitochondrial DNA (DS1 to DS8).

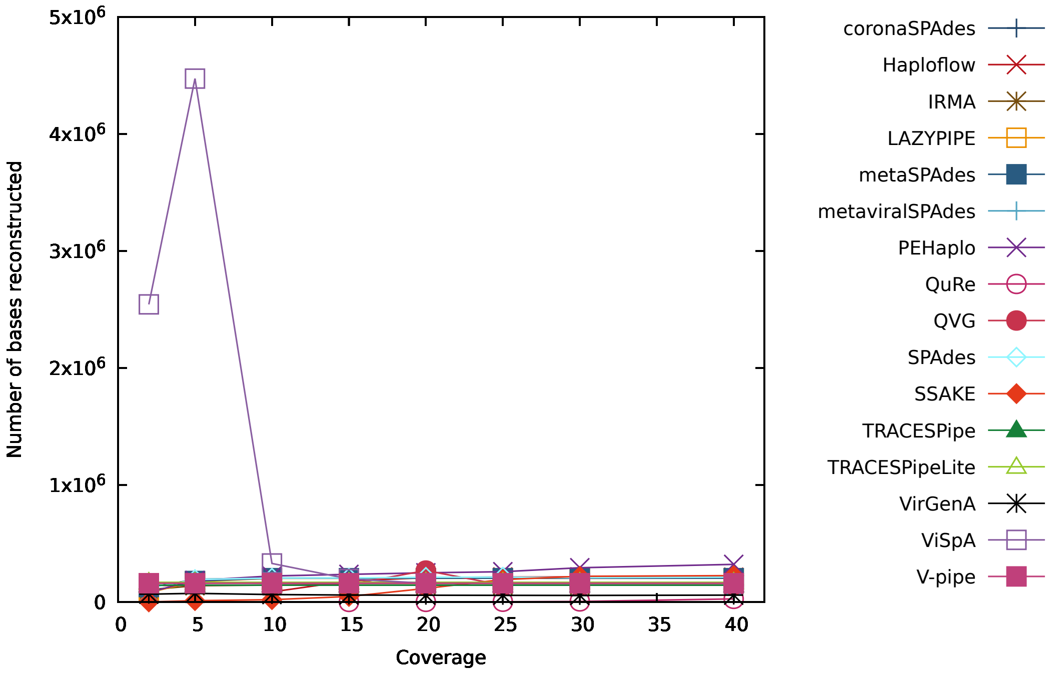


Figure S11: Figure comparing the performance of the reconstruction programs in terms of the number of bases reconstructed for datasets with contamination and mitochondrial DNA (DS9 to DS16).

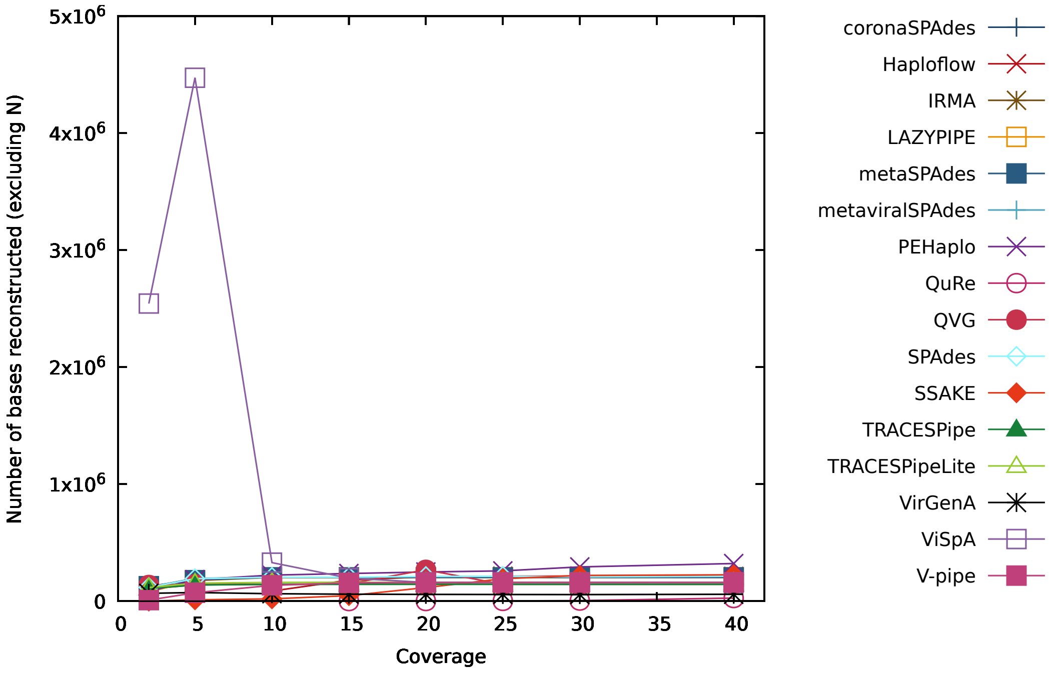


Figure S12: Figure comparing the performance of the reconstruction programs in terms of the number of bases reconstructed (excluding "N") for datasets with contamination and mitochondrial DNA (DS9 to DS16).

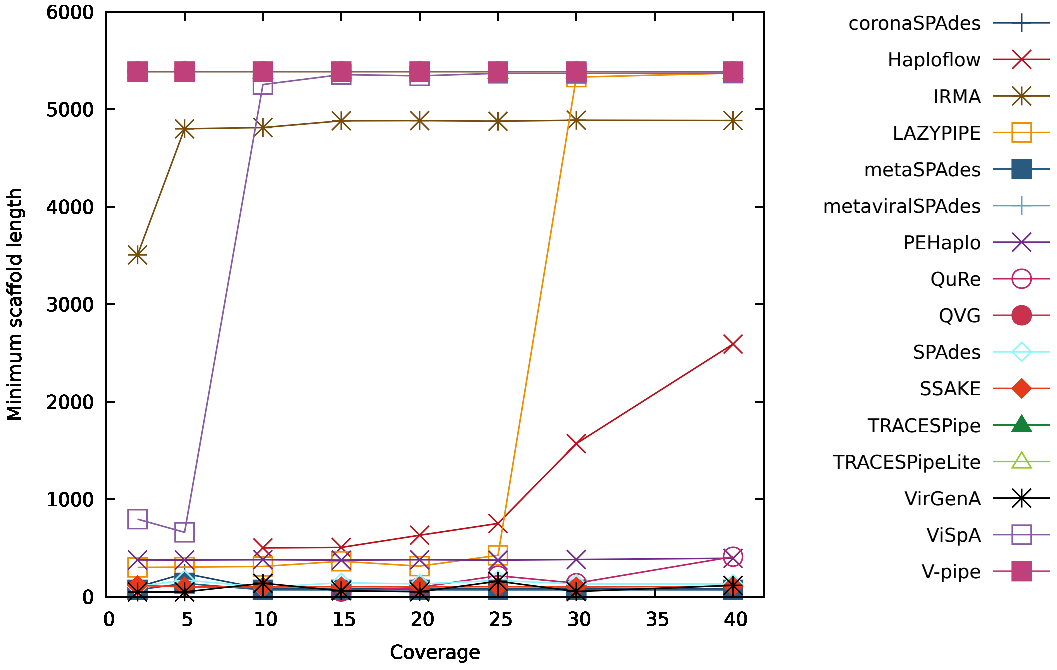


Figure S13: Figure comparing the performance of the reconstruction programs in terms of the minimum number of reconstructed bases per scaffold for datasets with contamination and mitochondrial DNA (DS9 to DS16).

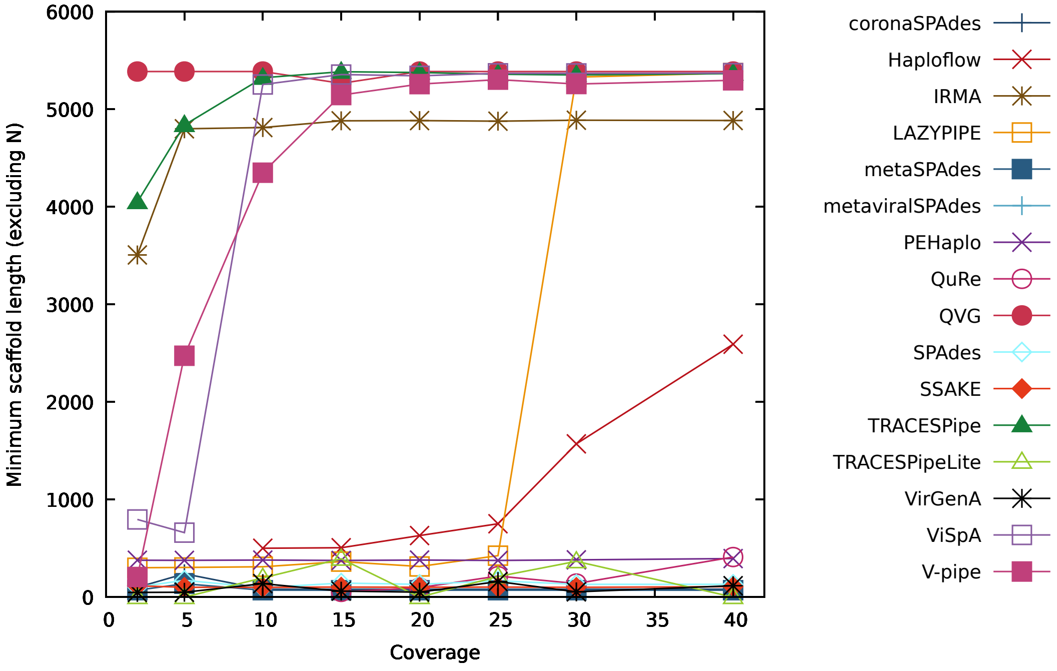


Figure S14: Figure comparing the performance of the reconstruction programs in terms of the minimum number of reconstructed bases per scaffold (excluding "N") for datasets with contamination and mitochondrial DNA (DS9 to DS16).

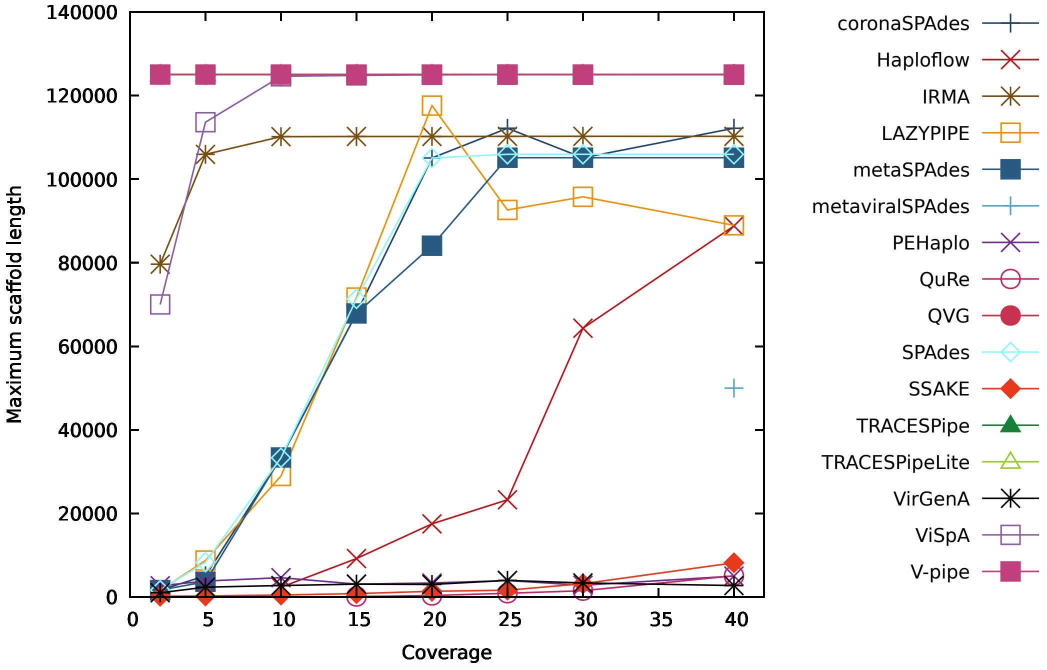


Figure S15: Figure comparing the performance of the reconstruction programs in terms of the maximum number of reconstructed bases per scaffold for datasets with contamination and mitochondrial DNA (DS9 to DS16).

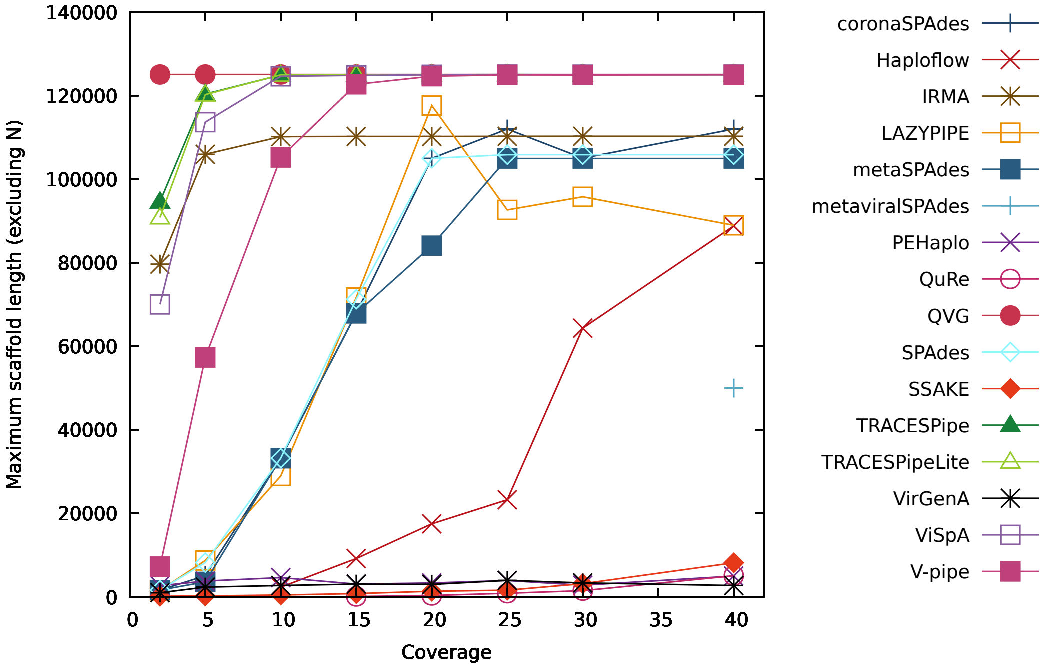


Figure S16: Figure comparing the performance of the reconstruction programs in terms of the maximum number of reconstructed bases per scaffold (excluding "N") for datasets with contamination and mitochondrial DNA (DS9 to DS16).

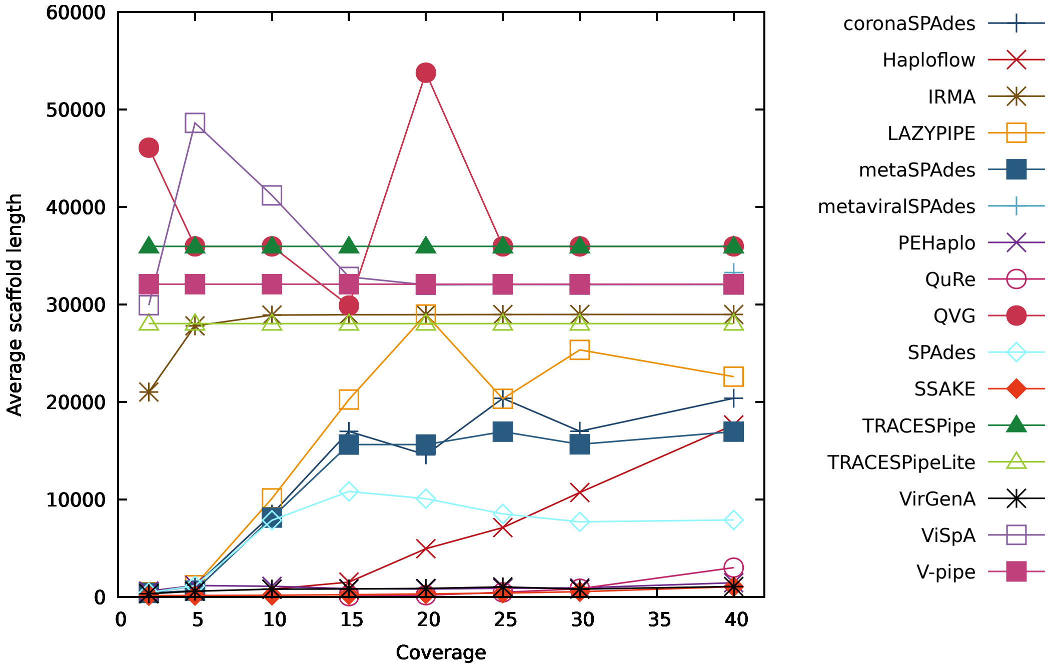


Figure S17: Figure comparing the performance of the reconstruction programs in terms of the average number of reconstructed bases per scaffold for datasets with contamination and mitochondrial DNA (DS9 to DS16).

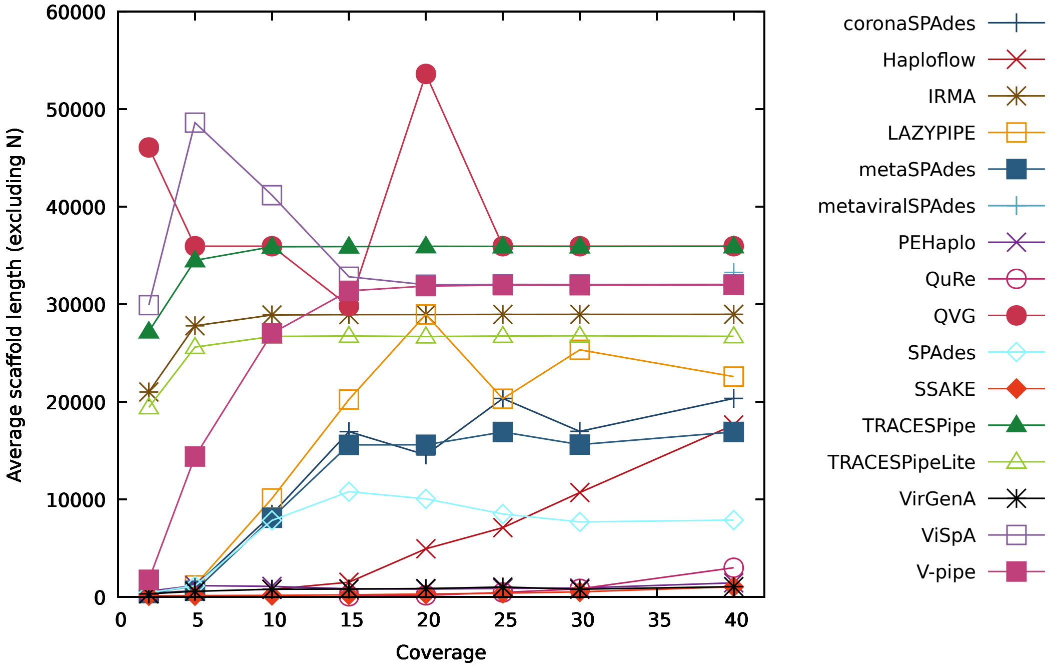


Figure S18: Figure comparing the performance of the reconstruction programs in terms of the average number of reconstructed bases per scaffold (excluding "N") for datasets with contamination and mitochondrial DNA (DS9 to DS16).

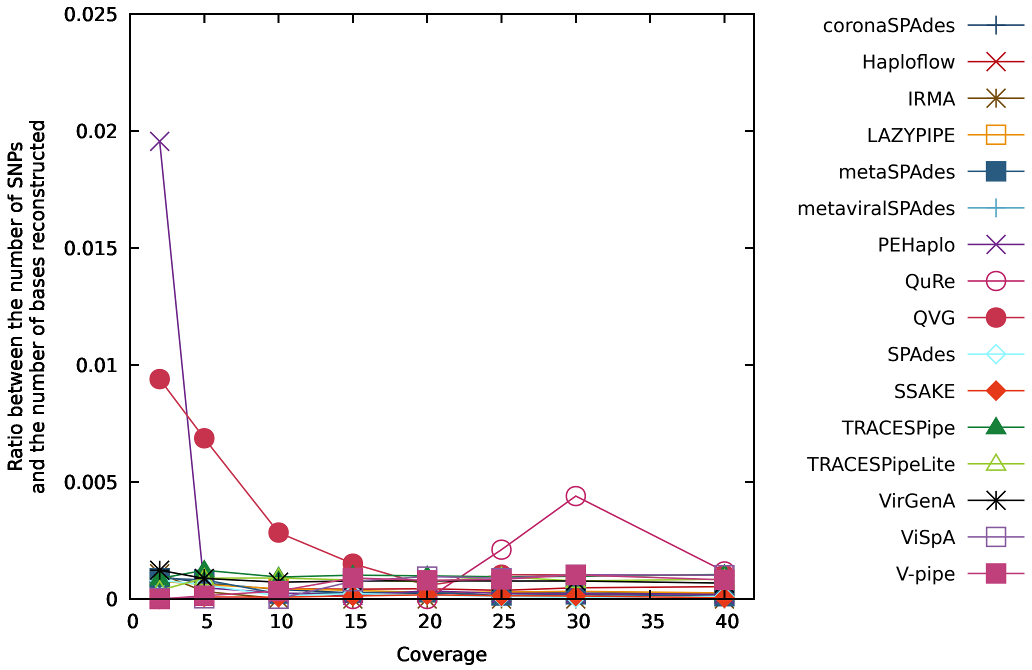


Figure S19: Figure comparing the performance of the reconstruction programs in terms of the ratio between the number of SNPs and the number of bases reconstructed for datasets with contamination and mitochondrial DNA (DS9 to DS16).

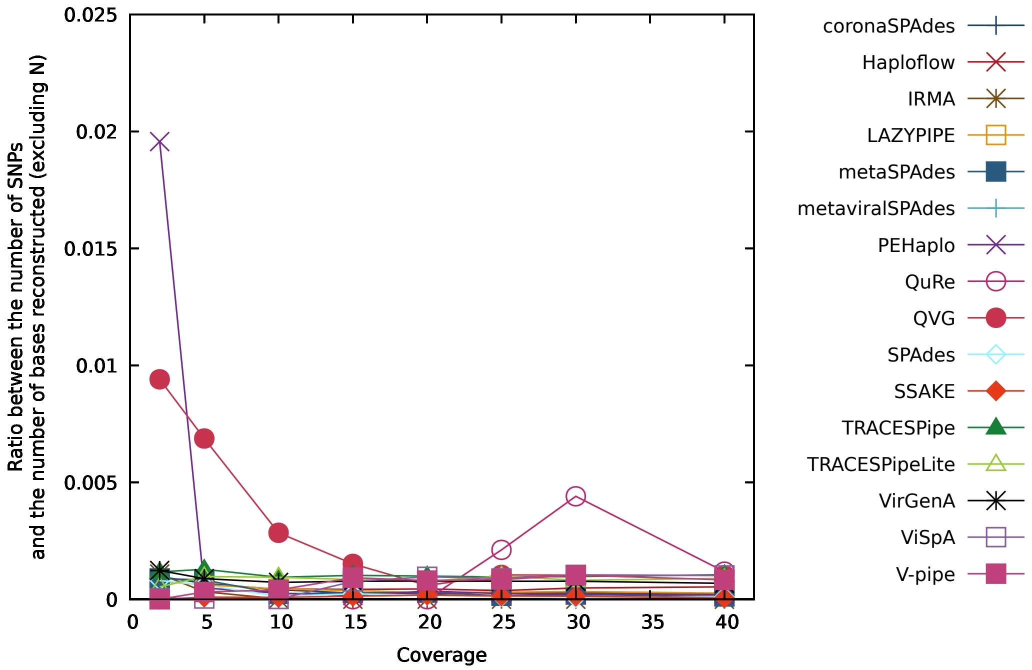


Figure S20: Figure comparing the performance of the reconstruction programs in terms of the ratio between the number of SNPs and the number of bases reconstructed (excluding "N") for datasets with contamination and mitochondrial DNA (DS9 to DS16).

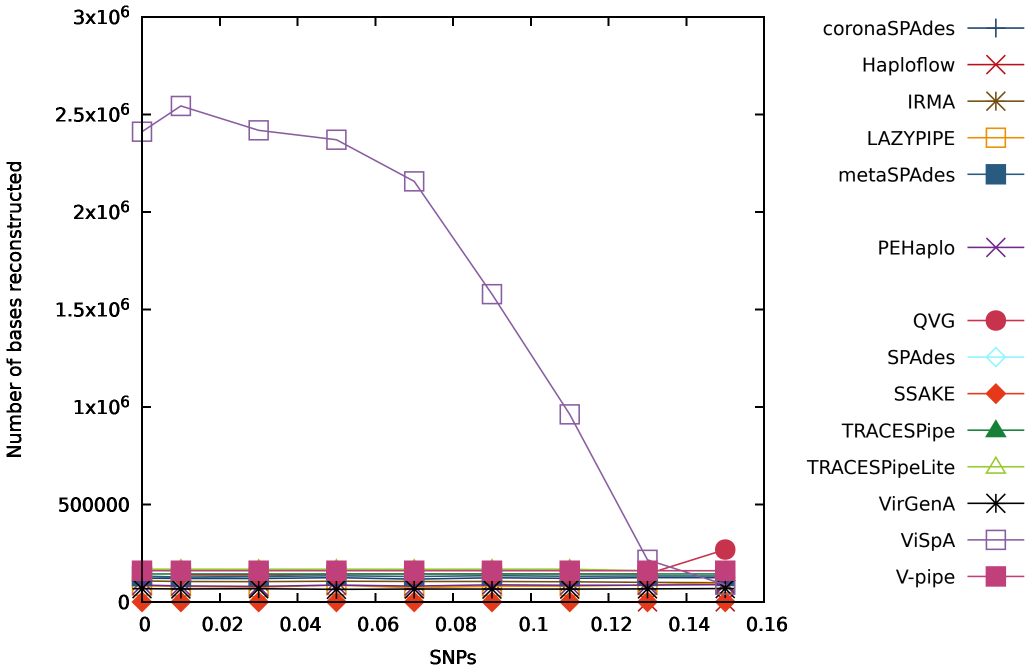


Figure S21: Figure comparing the performance of the reconstruction programs in terms of the number of bases reconstructed for datasets with depth coverage equal to 2x (DS17 to DS24 plus DS9).

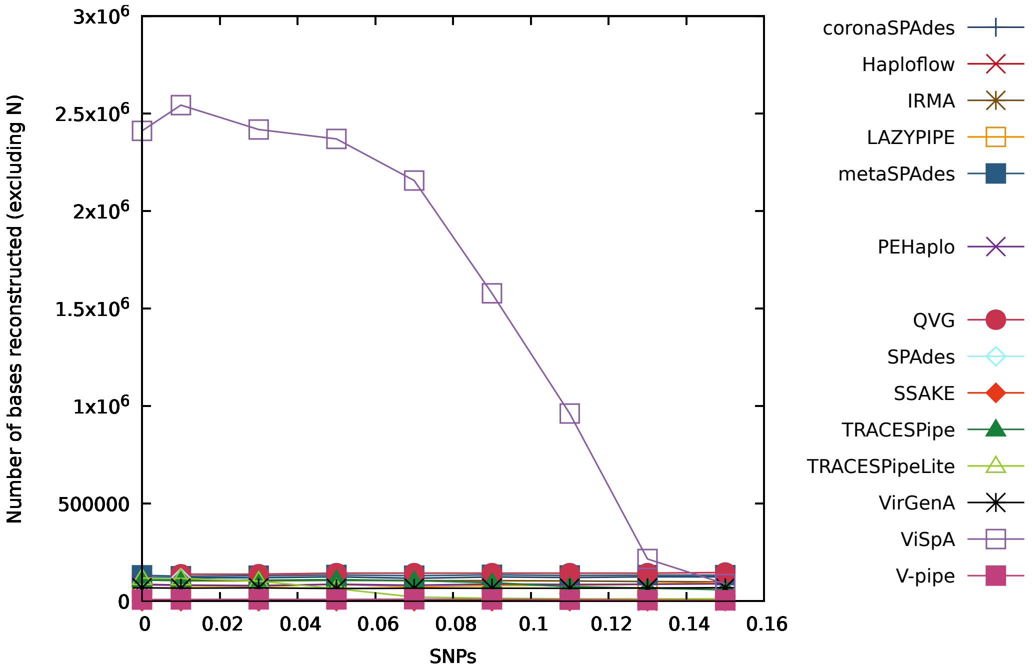


Figure S22: Figure comparing the performance of the reconstruction programs in terms of the number of bases reconstructed (excluding "N") for datasets with depth coverage equal to 2x (DS17 to DS24 plus DS9).

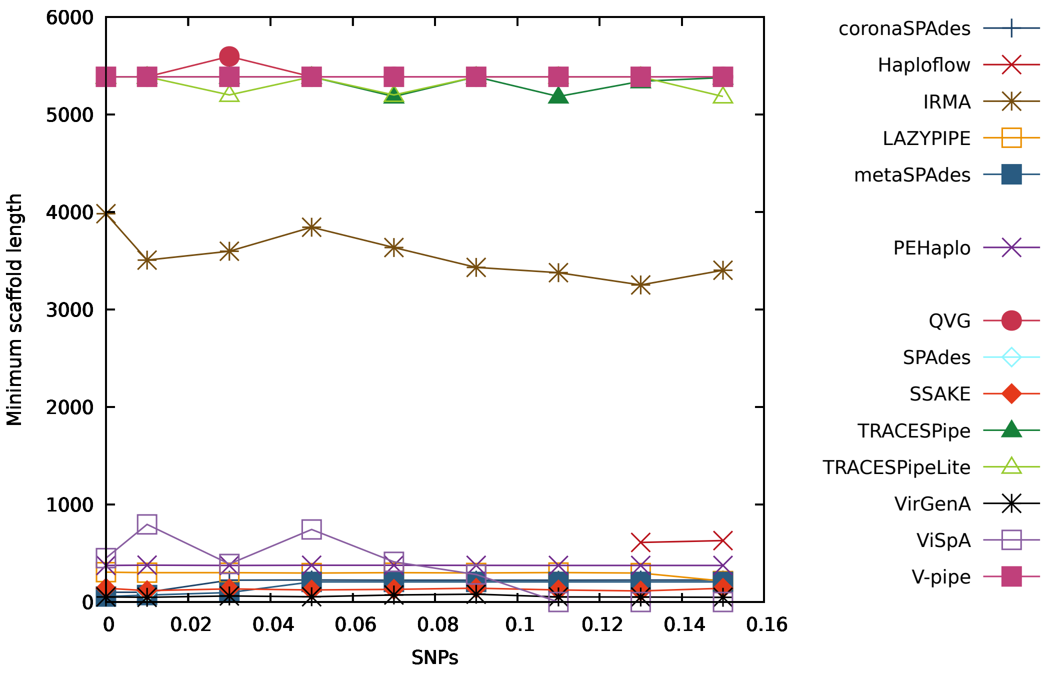


Figure S23: Figure comparing the performance of the reconstruction programs in terms of the minimum number of reconstructed bases per scaffold for datasets with depth coverage equal to 2x (DS17 to DS24 plus DS9).

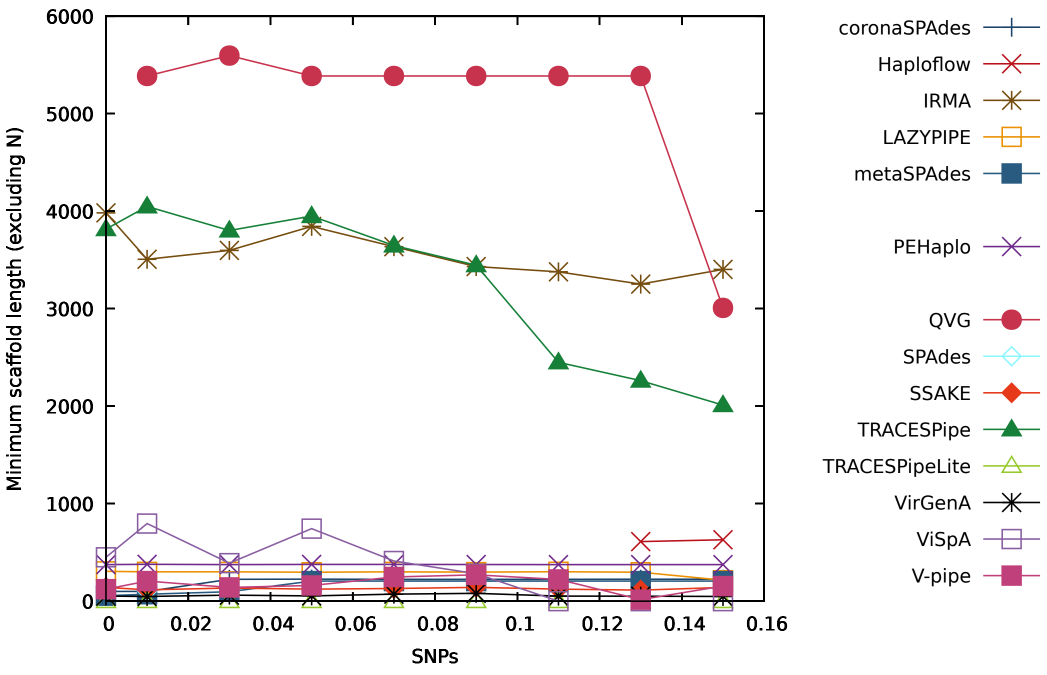


Figure S24: Figure comparing the performance of the reconstruction programs in terms of the minimum number of reconstructed bases per scaffold (excluding "N") for datasets with depth coverage equal to 2x (DS17 to DS24 plus DS9).

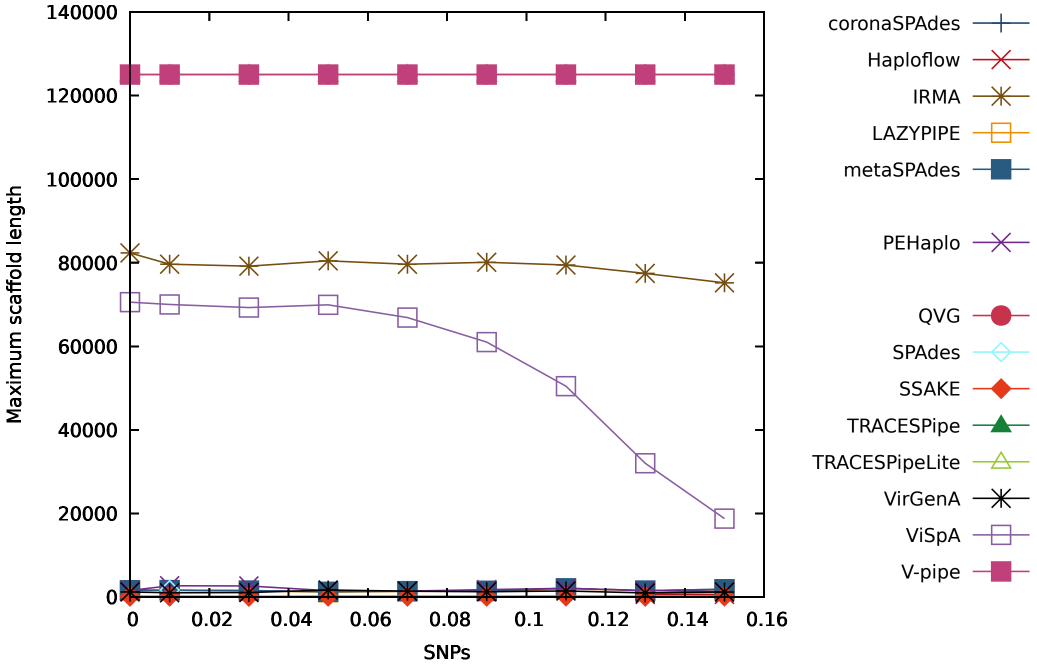


Figure S25: Figure comparing the performance of the reconstruction programs in terms of the maximum number of reconstructed bases per scaffold for datasets with depth coverage equal to 2x (DS17 to DS24 plus DS9).

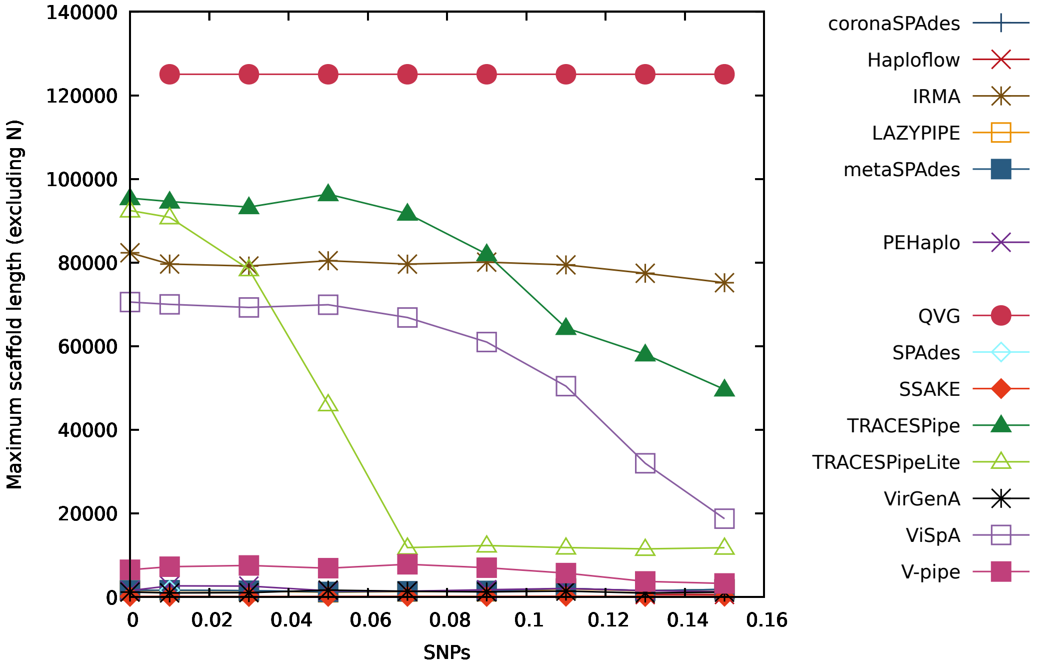


Figure S26: Figure comparing the performance of the reconstruction programs in terms of the maximum number of reconstructed bases per scaffold (excluding "N") for datasets with depth coverage equal to 2x (DS17 to DS24 plus DS9).

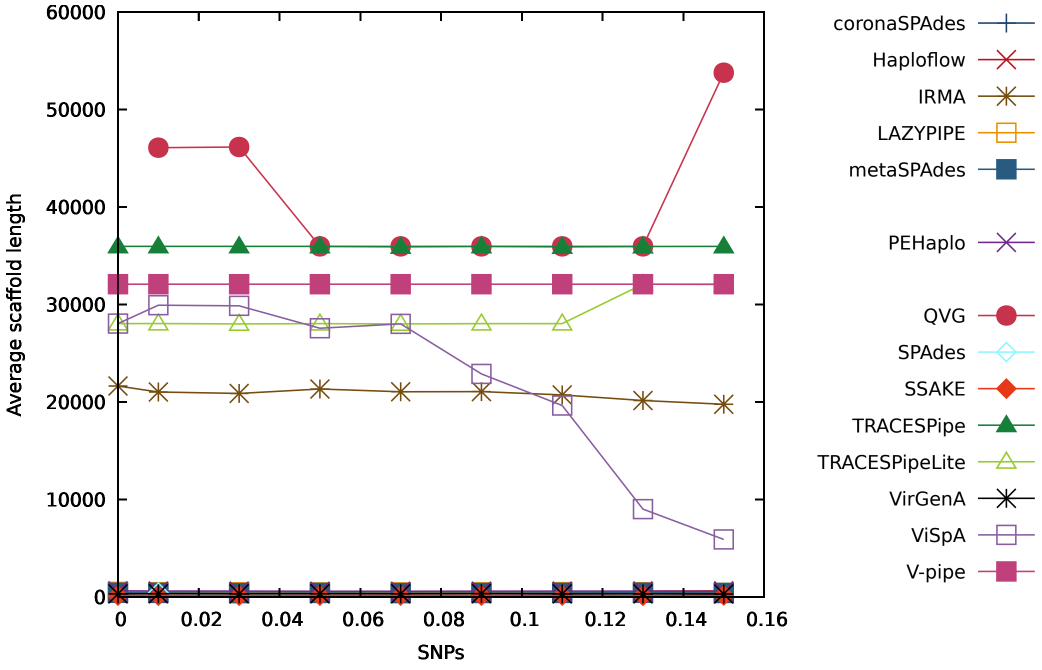


Figure S27: Figure comparing the performance of the reconstruction programs in terms of the average number of reconstructed bases per scaffold for datasets with depth coverage equal to 2x (DS17 to DS24 plus DS9).

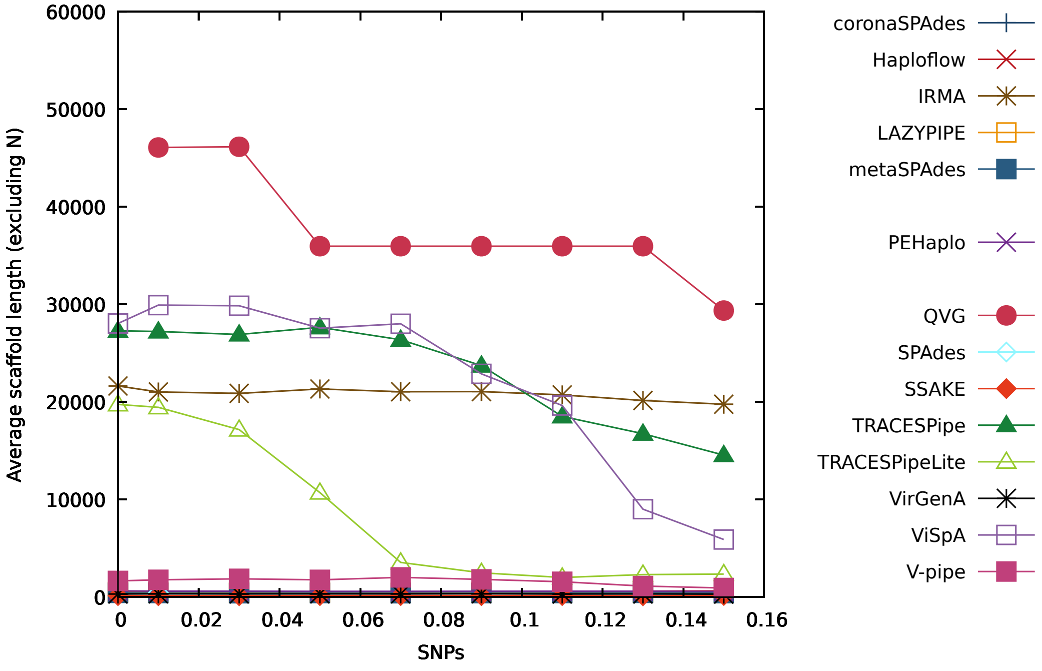


Figure S28: Figure comparing the performance of the reconstruction programs in terms of the average number of reconstructed bases per scaffold (excluding "N") for datasets with depth coverage equal to 2x (DS17 to DS24 plus DS9).

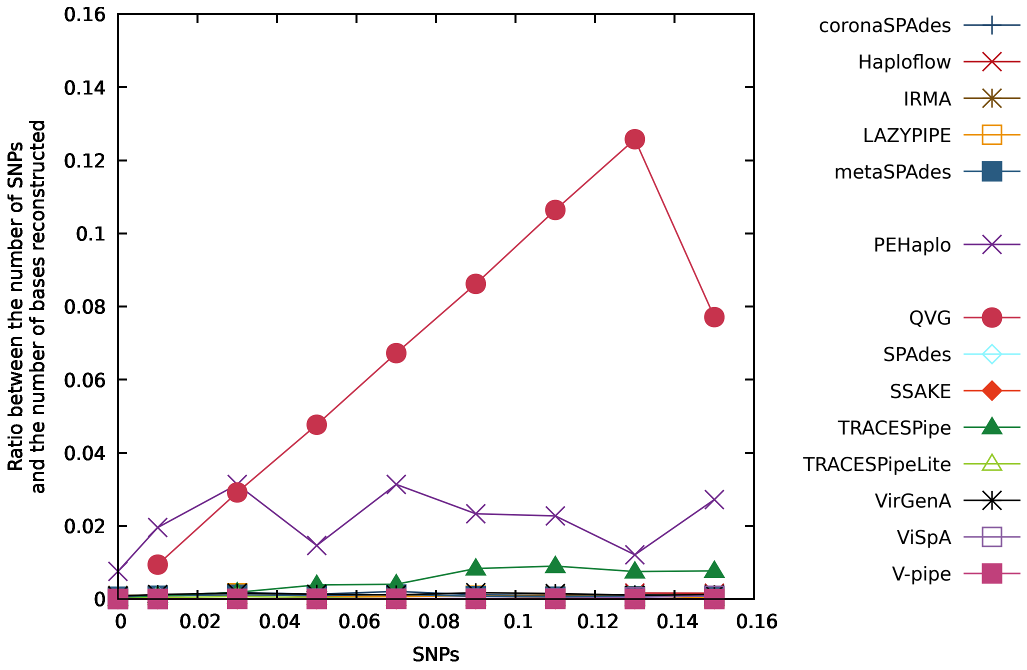


Figure S29: Figure comparing the performance of the reconstruction programs in terms of the ratio between the number of SNPs and the number of bases reconstructed for datasets with depth coverage equal to 2x (DS17 to DS24 plus DS9).

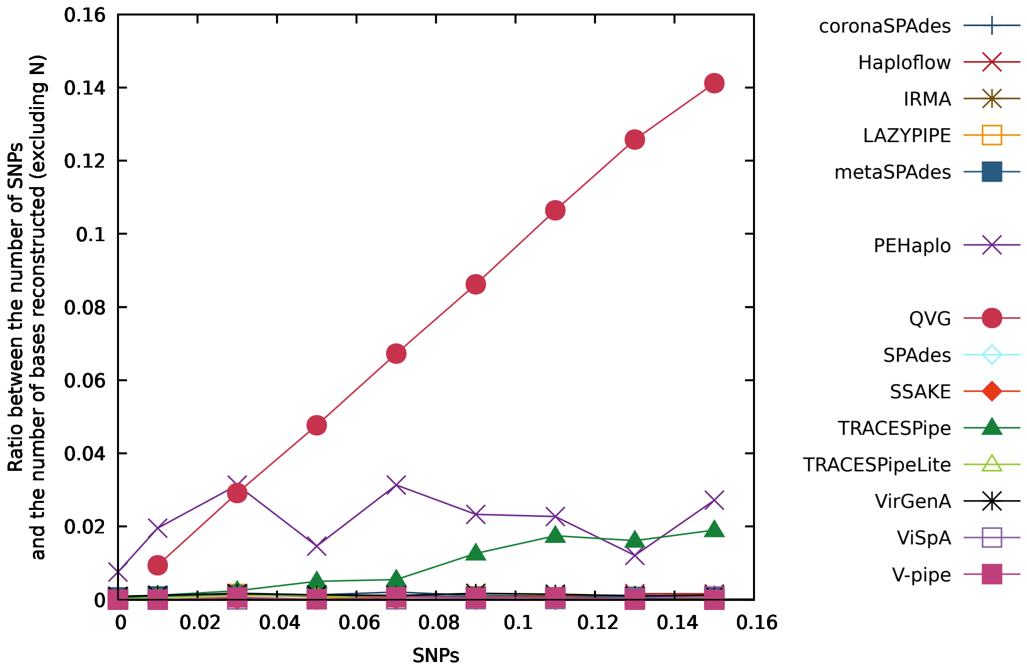


Figure S30: Figure comparing the performance of the reconstruction programs in terms of the ratio between the number of SNPs and the number of bases reconstructed (excluding "N") for datasets with depth coverage equal to 2x (DS17 to DS24 plus DS9).

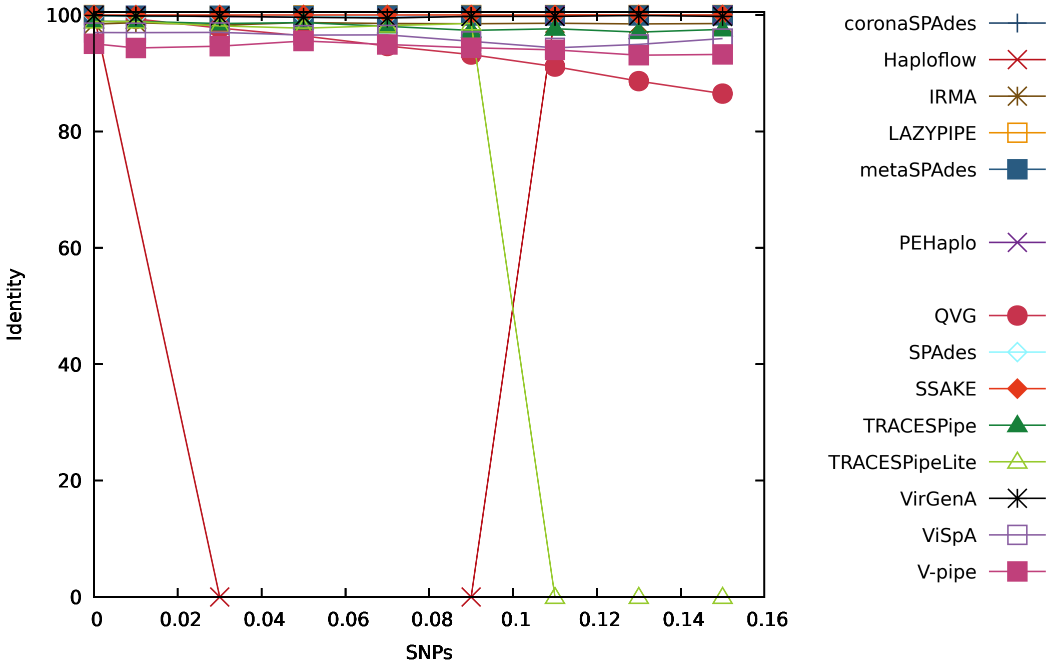


Figure S31: Figure comparing the performance of the reconstruction programs in terms of the identity for datasets with depth coverage equal to 5x (DS25 to DS32 plus DS10).

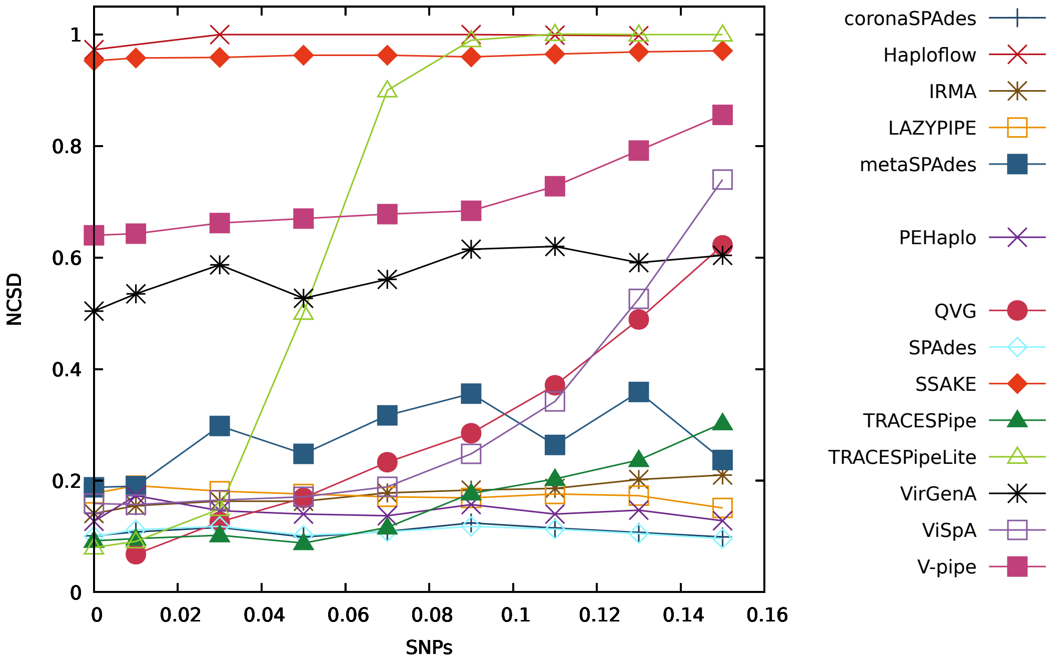


Figure S32: Figure comparing the performance of the reconstruction programs in terms of the NCSD for datasets with depth coverage equal to 5x (DS25 to DS32 plus DS10).

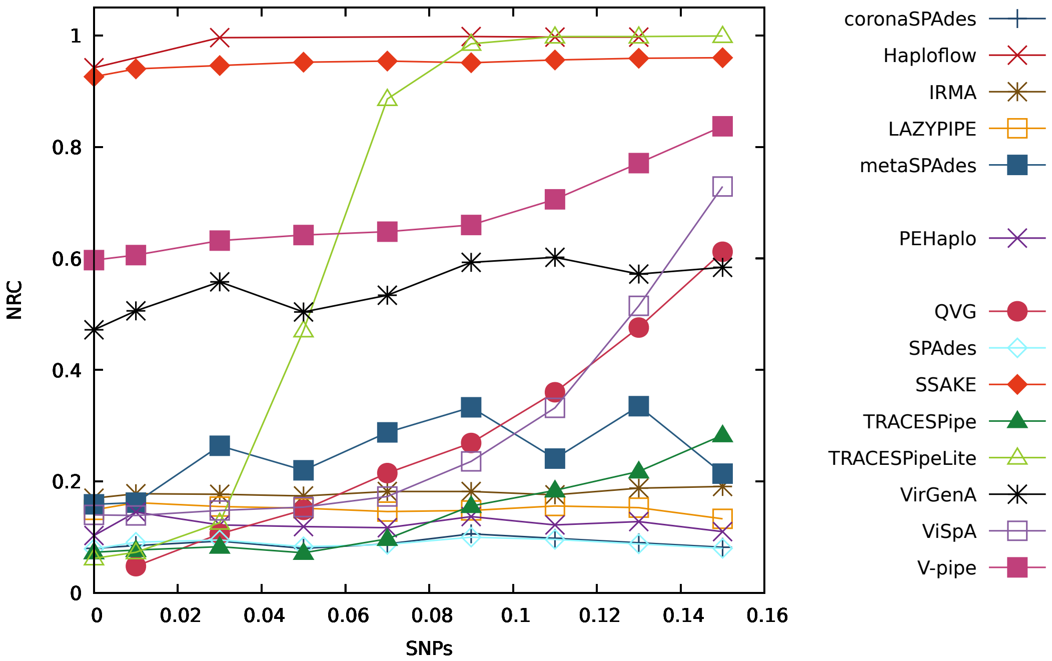


Figure S33: Figure comparing the performance of the reconstruction programs in terms of the NRC for datasets with depth coverage equal to 5x (DS25 to DS32 plus DS10).

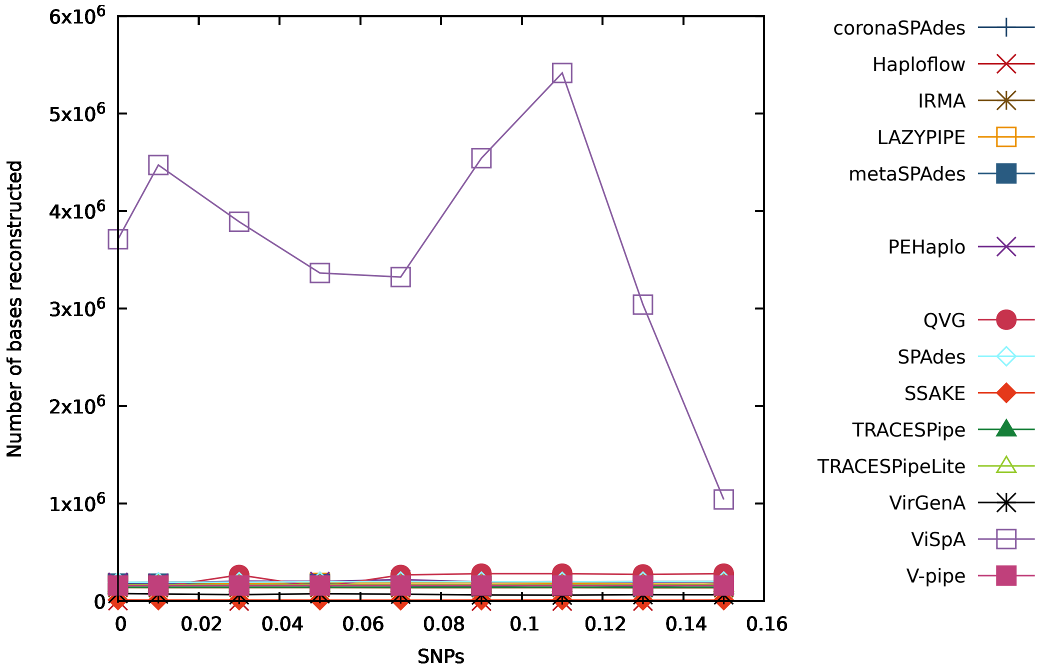


Figure S34: Figure comparing the performance of the reconstruction programs in terms of the number of bases reconstructed for datasets with depth coverage equal to 5x (DS25 to DS32 plus DS10).

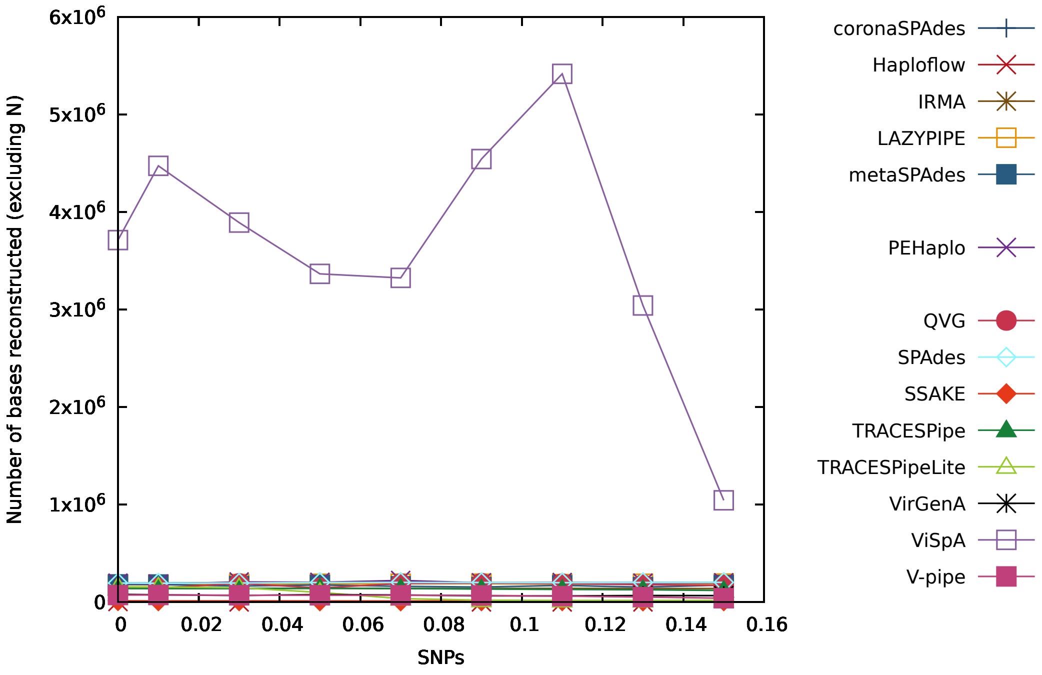


Figure S35: Figure comparing the performance of the reconstruction programs in terms of the number of bases reconstructed (excluding "N") for datasets with depth coverage equal to 5x (DS25 to DS32 plus DS10).

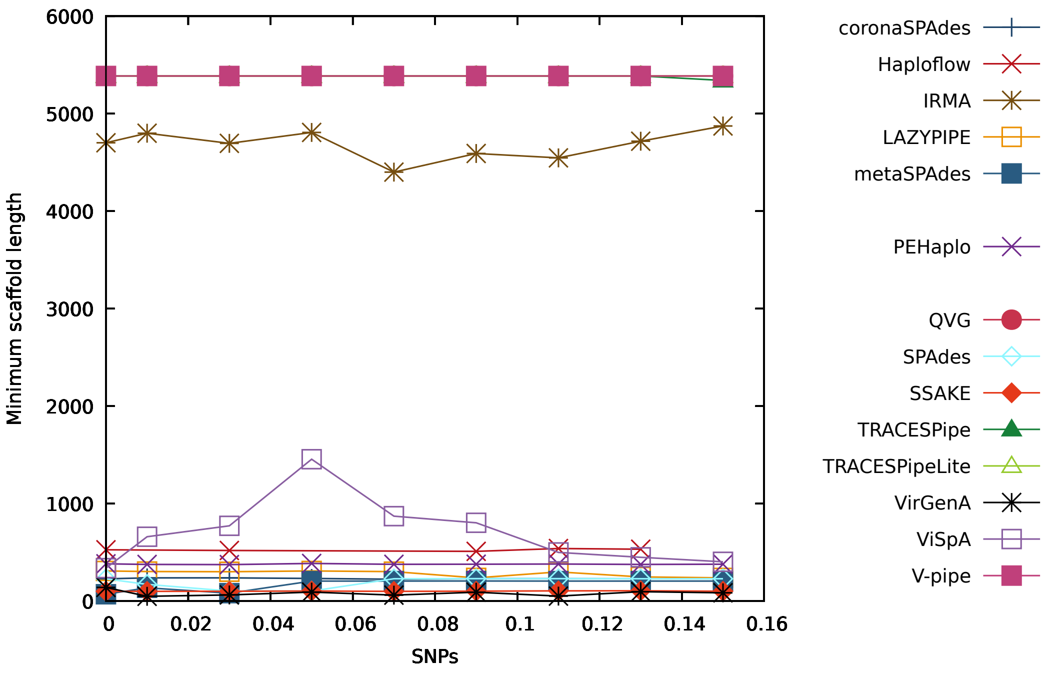


Figure S36: Figure comparing the performance of the reconstruction programs in terms of the minimum number of reconstructed bases per scaffold for datasets with depth coverage equal to 5x (DS25 to DS32 plus DS10).

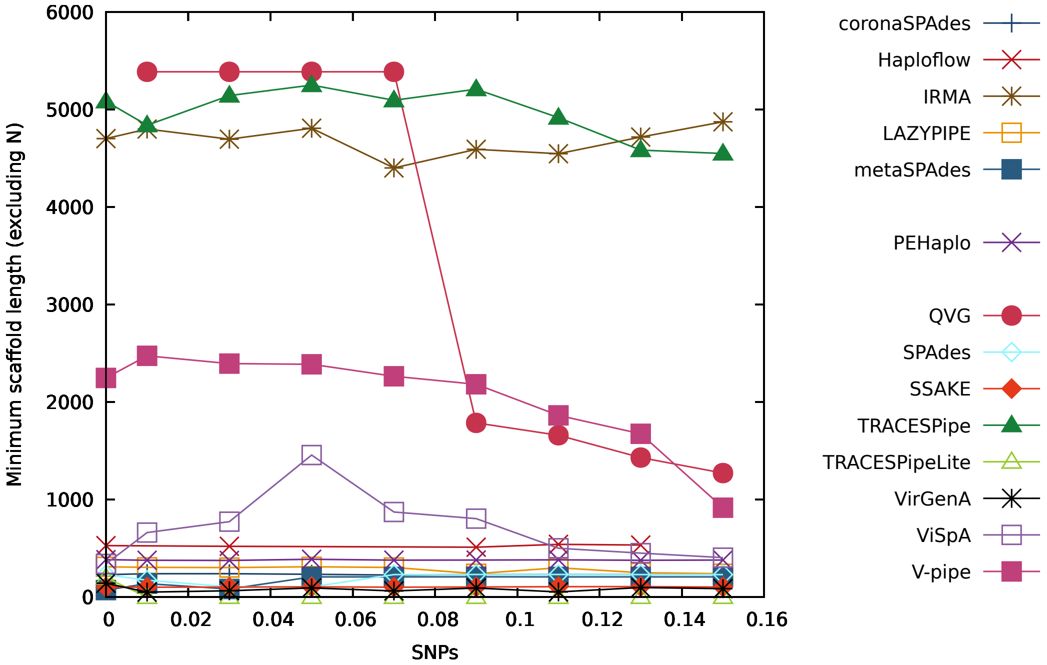


Figure S37: Figure comparing the performance of the reconstruction programs in terms of the minimum number of reconstructed bases per scaffold (excluding "N") for datasets with depth coverage equal to 5x (DS25 to DS32 plus DS10).

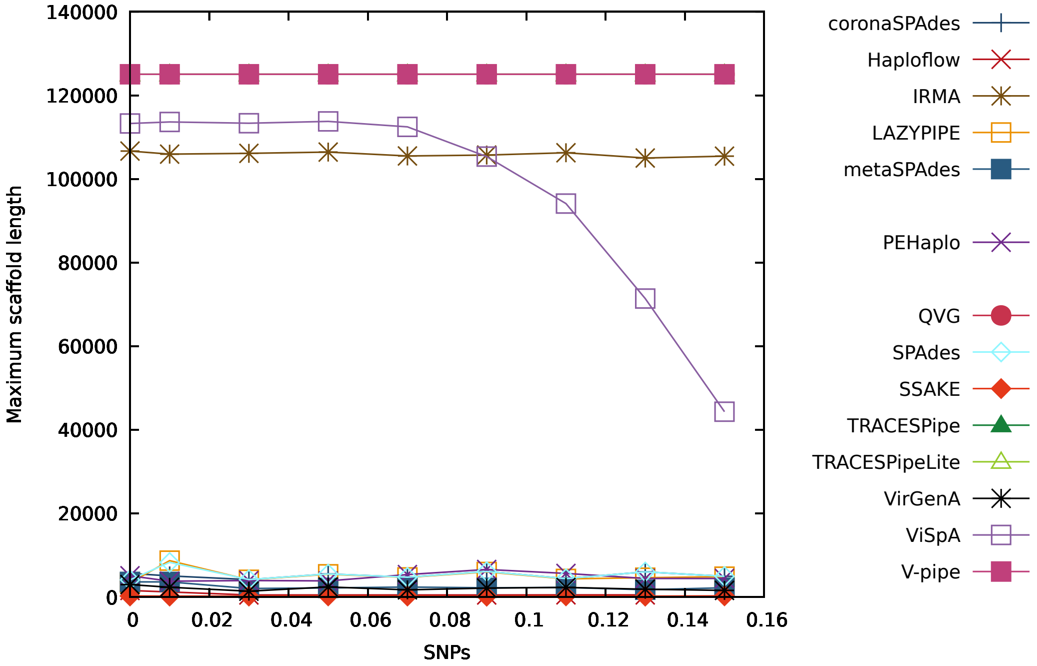


Figure S38: Figure comparing the performance of the reconstruction programs in terms of the maximum number of reconstructed bases per scaffold for datasets with depth coverage equal to 5x (DS25 to DS32 plus DS10).

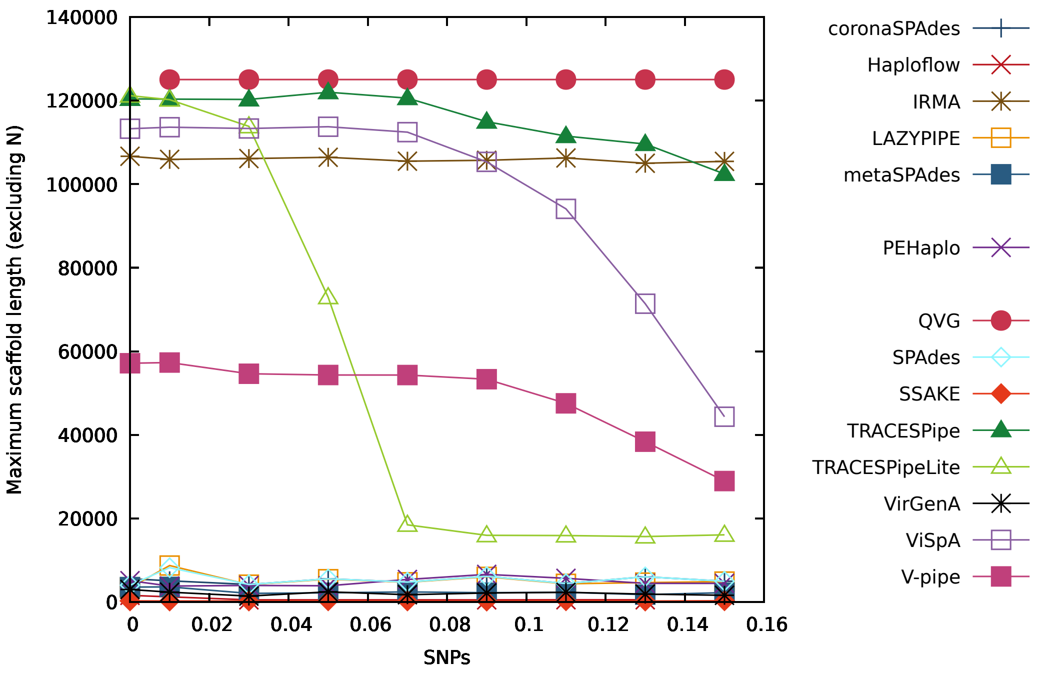


Figure S39: Figure comparing the performance of the reconstruction programs in terms of the maximum number of reconstructed bases per scaffold (excluding "N") for datasets with depth coverage equal to 5x (DS25 to DS32 plus DS10).

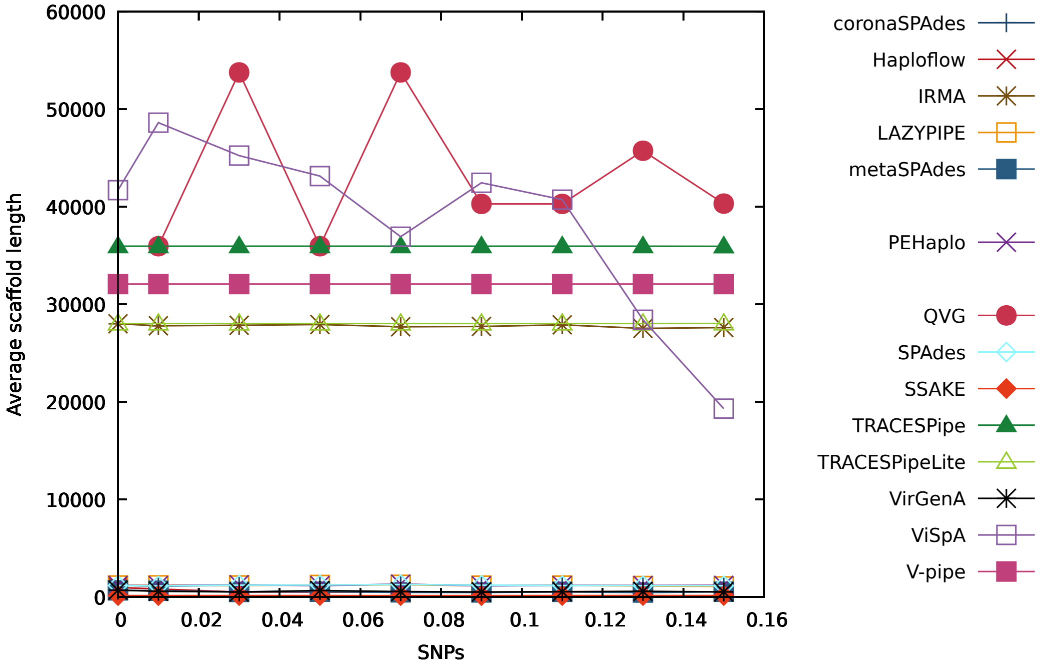


Figure S40: Figure comparing the performance of the reconstruction programs in terms of the average number of reconstructed bases per scaffold for datasets with depth coverage equal to 5x (DS25 to DS32 plus DS10).

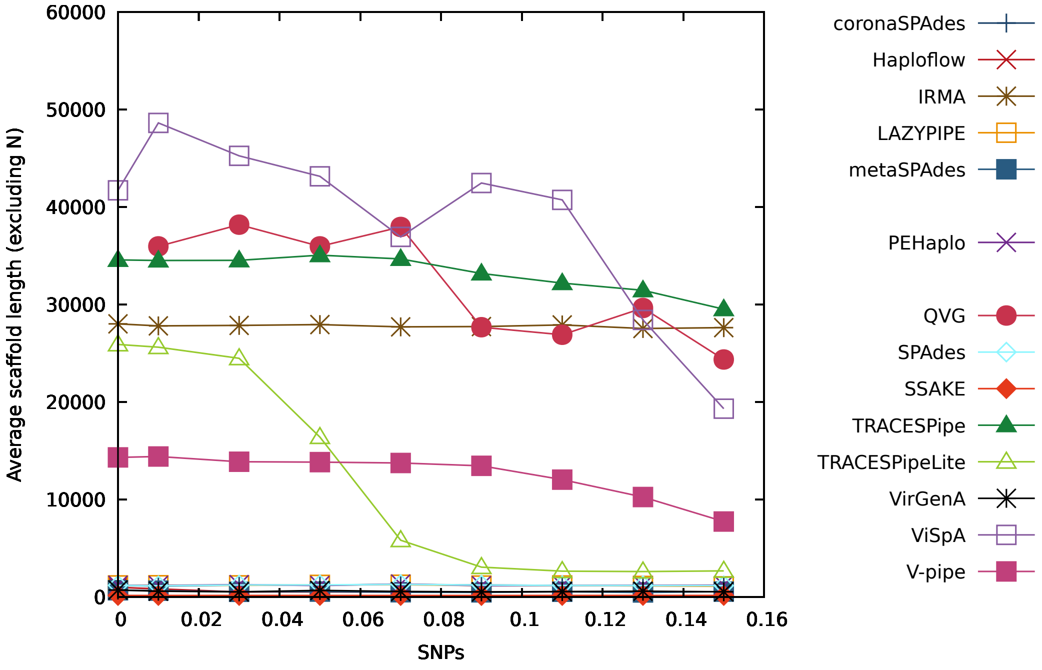


Figure S41: Figure comparing the performance of the reconstruction programs in terms of the average number of reconstructed bases per scaffold (excluding "N") for datasets with depth coverage equal to 5x (DS25 to DS32 plus DS10).

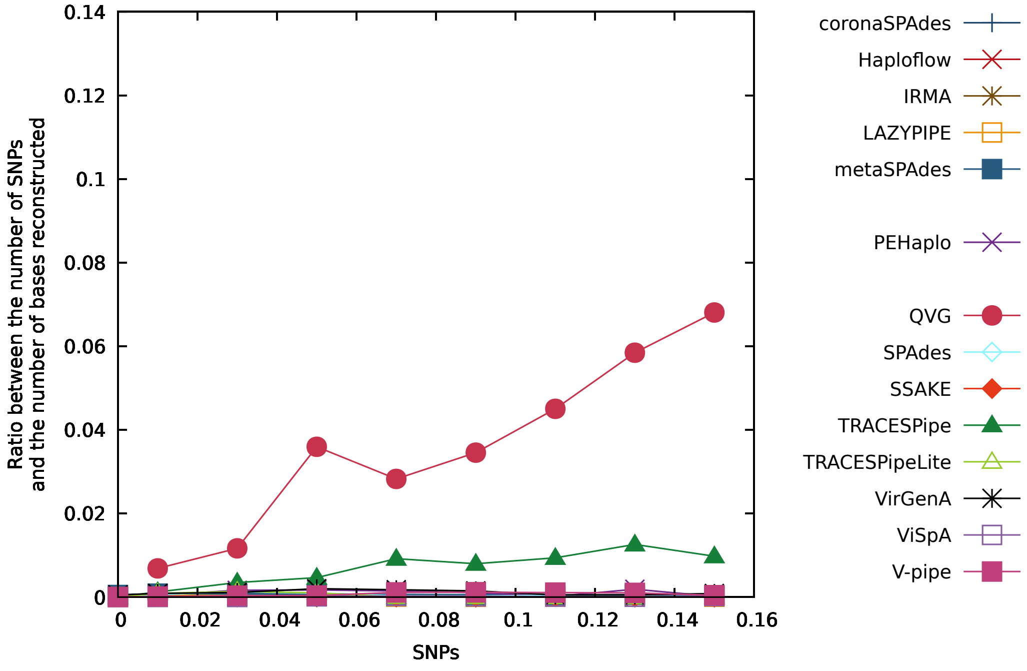


Figure S42: Figure comparing the performance of the reconstruction programs in terms of the ratio between the number of SNPs and the number of bases reconstructed for datasets with depth coverage equal to 5x (DS25 to DS32 plus DS10).

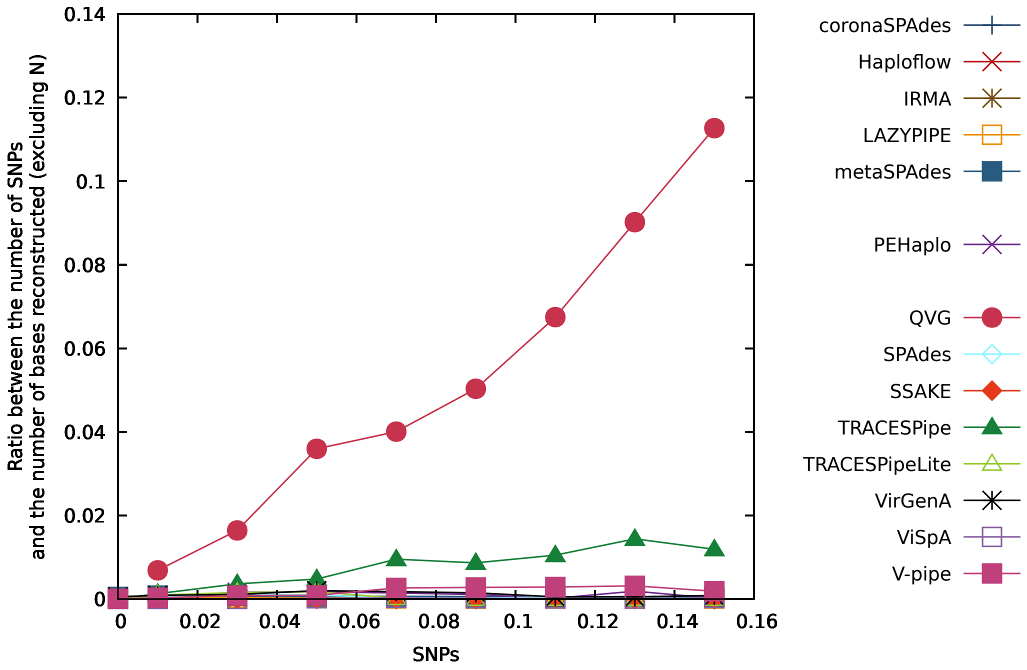


Figure S43: Figure comparing the performance of the reconstruction programs in terms of the ratio between the number of SNPs and the number of bases reconstructed (excluding "N") for datasets with depth coverage equal to 5x (DS25 to DS32 plus DS10).

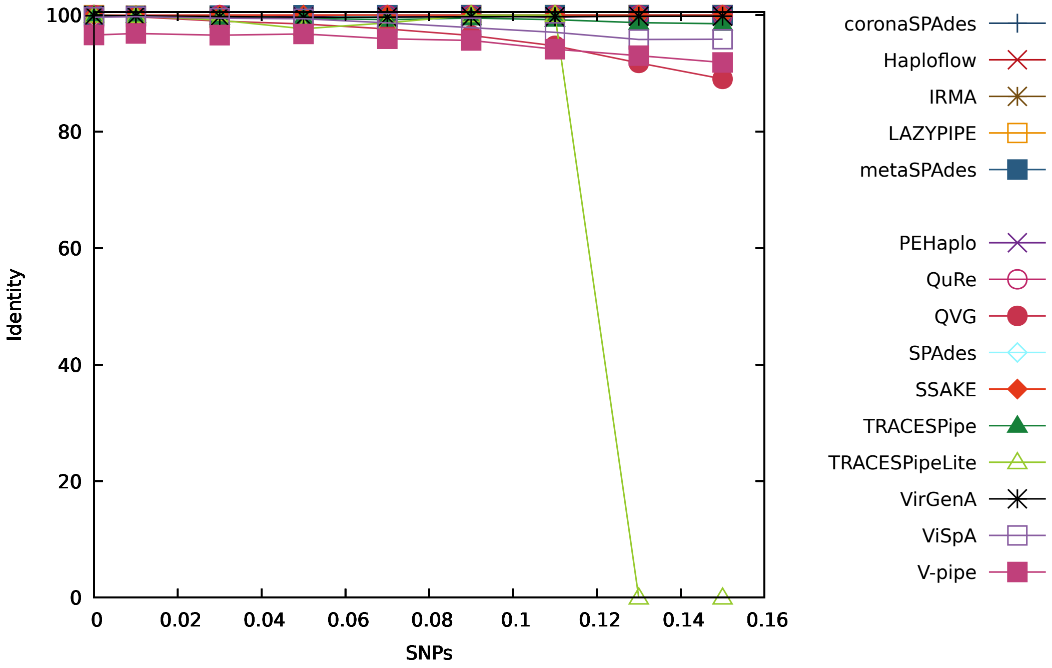


Figure S44: Figure comparing the performance of the reconstruction programs in terms of the identity for datasets with depth coverage equal to 10x (DS33 to DS40 plus DS11).

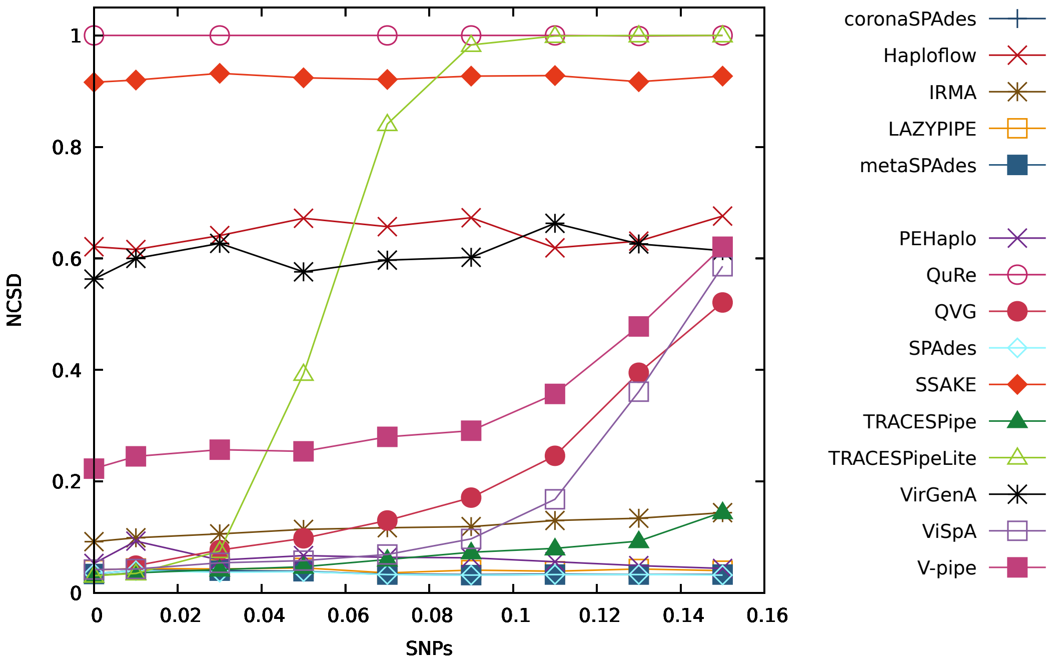


Figure S45: Figure comparing the performance of the reconstruction programs in terms of the NCSD for datasets with depth coverage equal to 10x (DS33 to DS40 plus DS11).

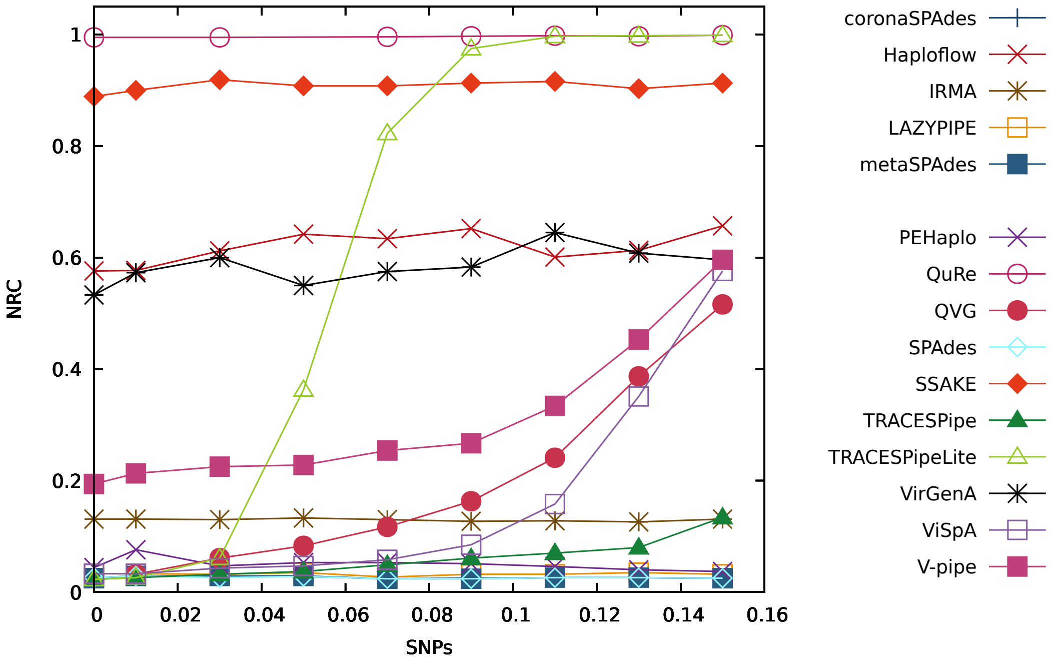


Figure S46: Figure comparing the performance of the reconstruction programs in terms of the NRC for datasets with depth coverage equal to 10x (DS33 to DS40 plus DS11).

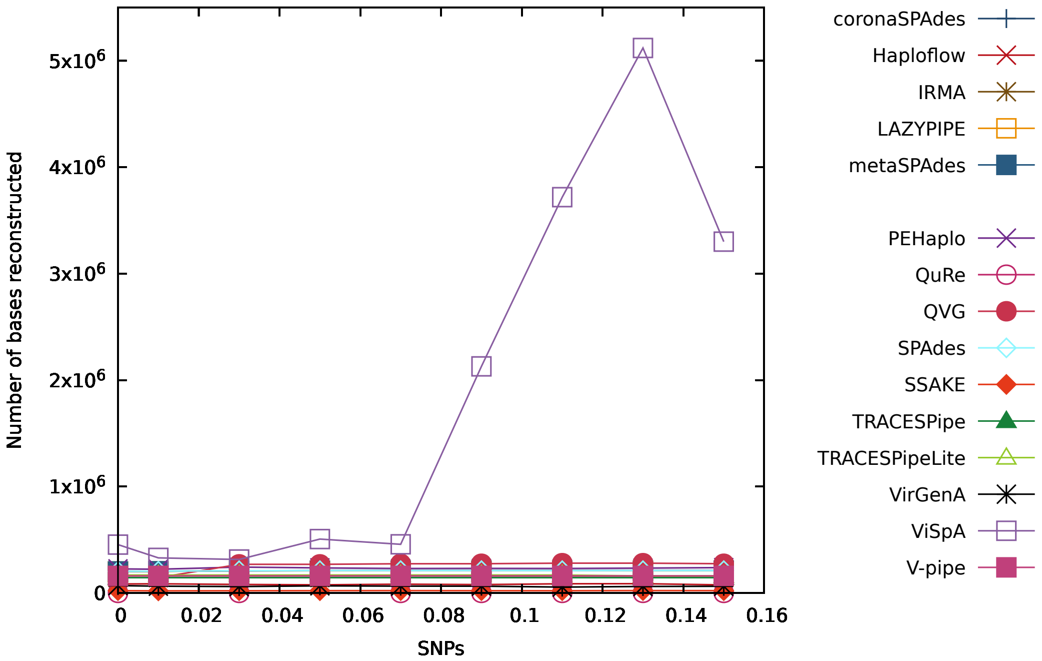


Figure S47: Figure comparing the performance of the reconstruction programs in terms of the number of bases reconstructed for datasets with depth coverage equal to 10x (DS33 to DS40 plus DS11).

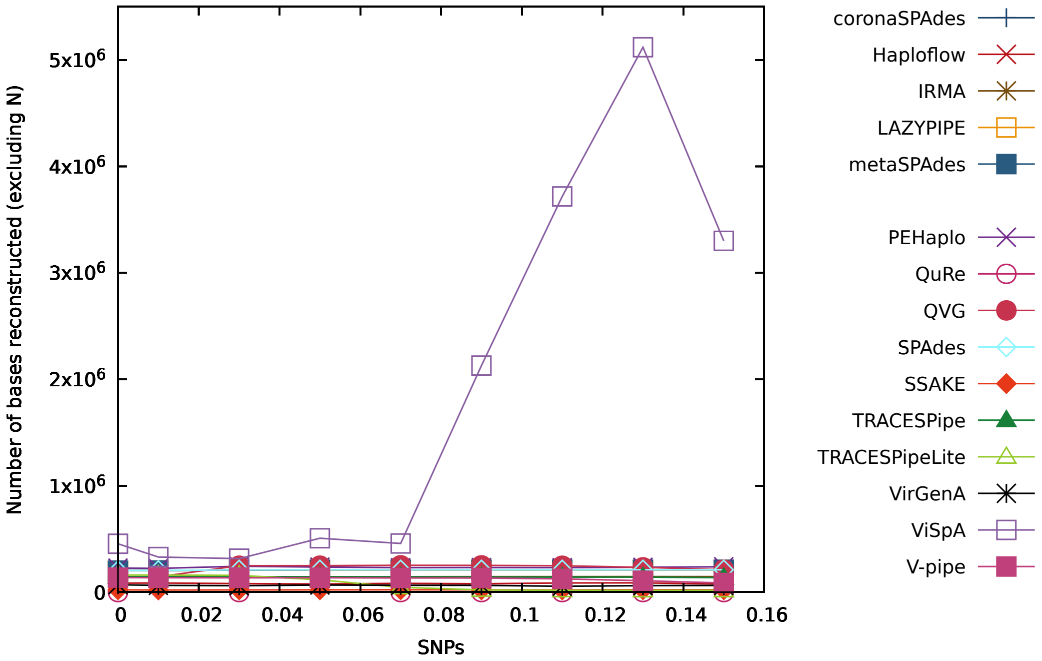


Figure S48: Figure comparing the performance of the reconstruction programs in terms of the number of bases reconstructed (excluding "N") for datasets with depth coverage equal to 10x (DS33 to DS40 plus DS11).

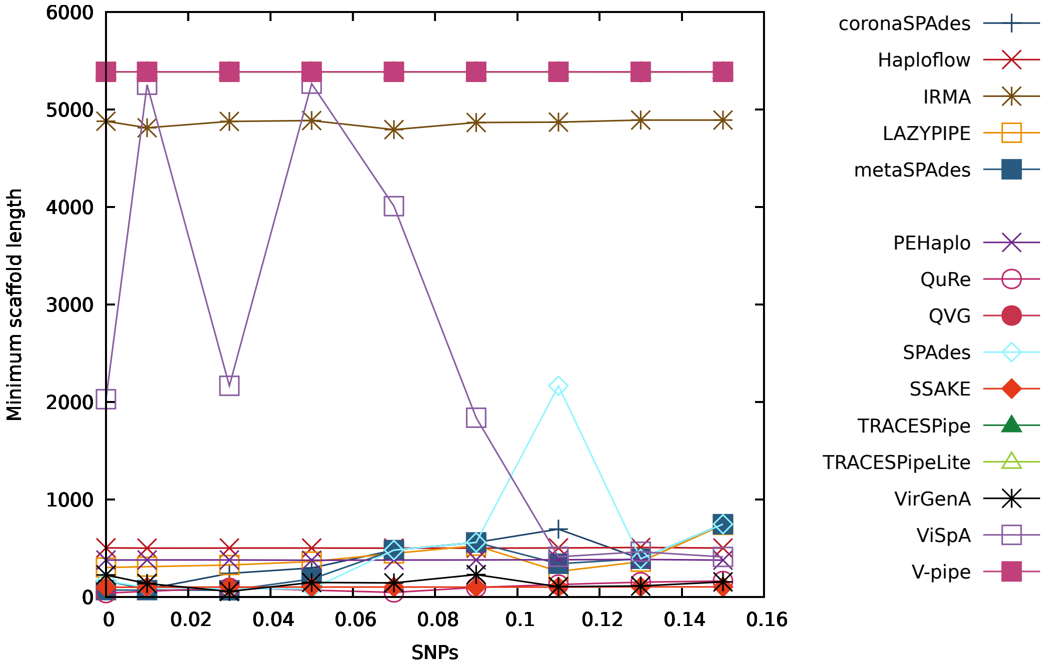


Figure S49: Figure comparing the performance of the reconstruction programs in terms of the minimum number of reconstructed bases per scaffold for datasets with depth coverage equal to 10x (DS33 to DS40 plus DS11).

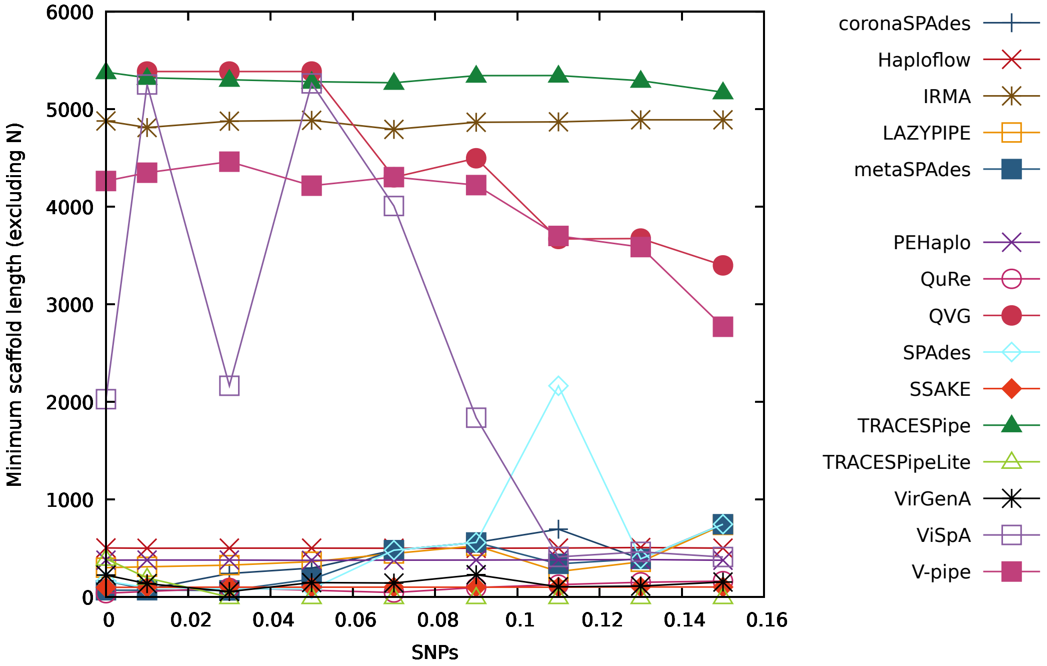


Figure S50: Figure comparing the performance of the reconstruction programs in terms of the minimum number of reconstructed bases per scaffold (excluding "N") for datasets with depth coverage equal to 10x (DS33 to DS40 plus DS11).

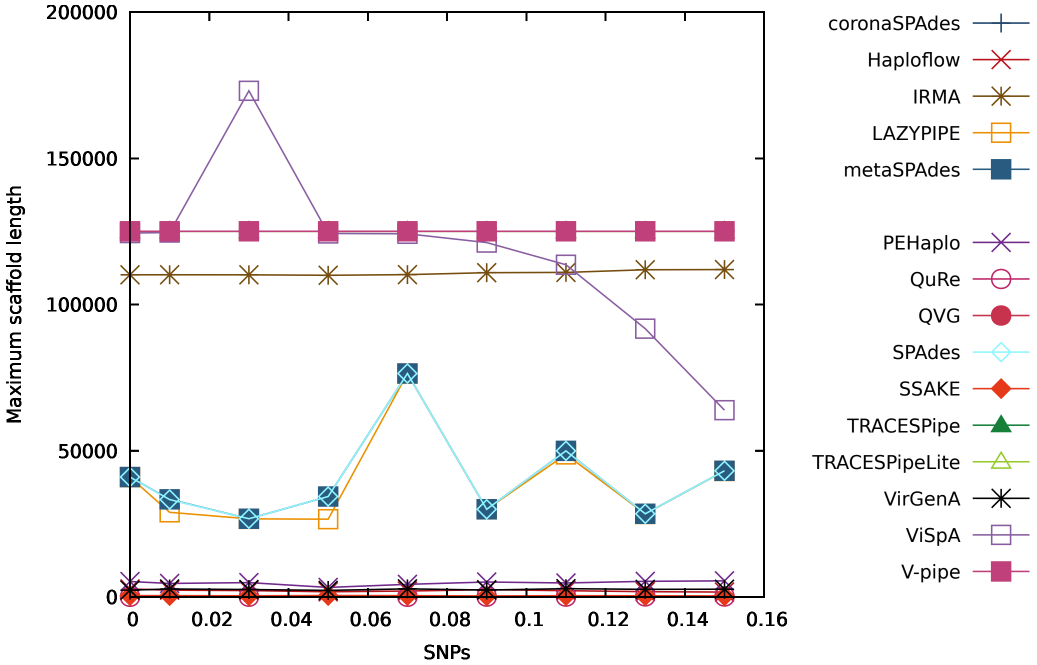


Figure S51: Figure comparing the performance of the reconstruction programs in terms of the maximum number of reconstructed bases per scaffold for datasets with depth coverage equal to 10x (DS33 to DS40 plus DS11).

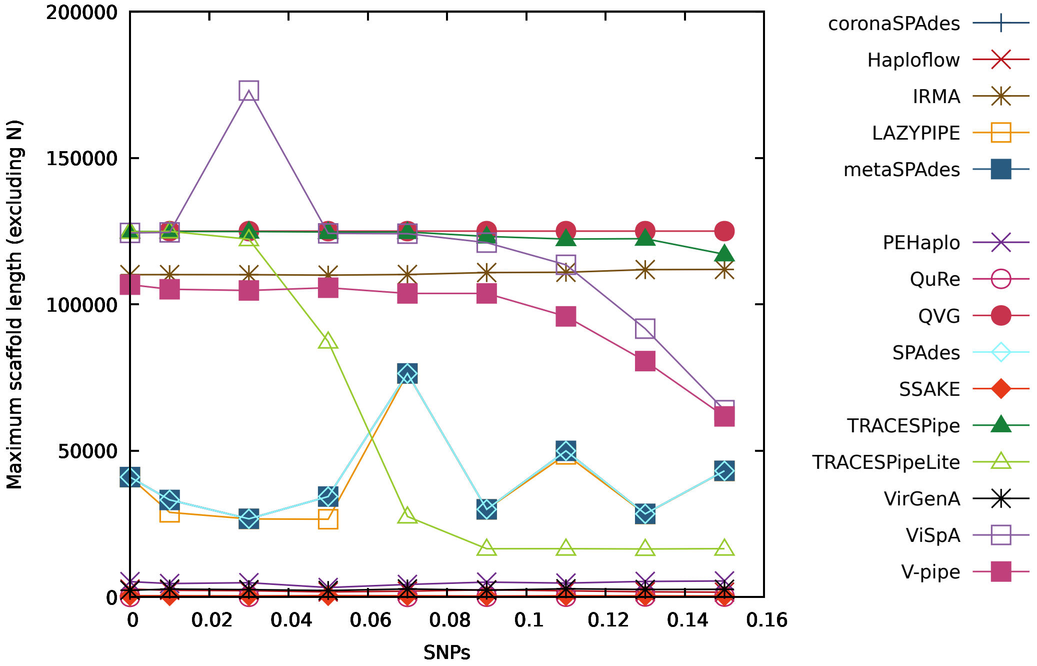


Figure S52: Figure comparing the performance of the reconstruction programs in terms of the maximum number of reconstructed bases per scaffold (excluding "N") for datasets with depth coverage equal to 10x (DS33 to DS40 plus DS11).

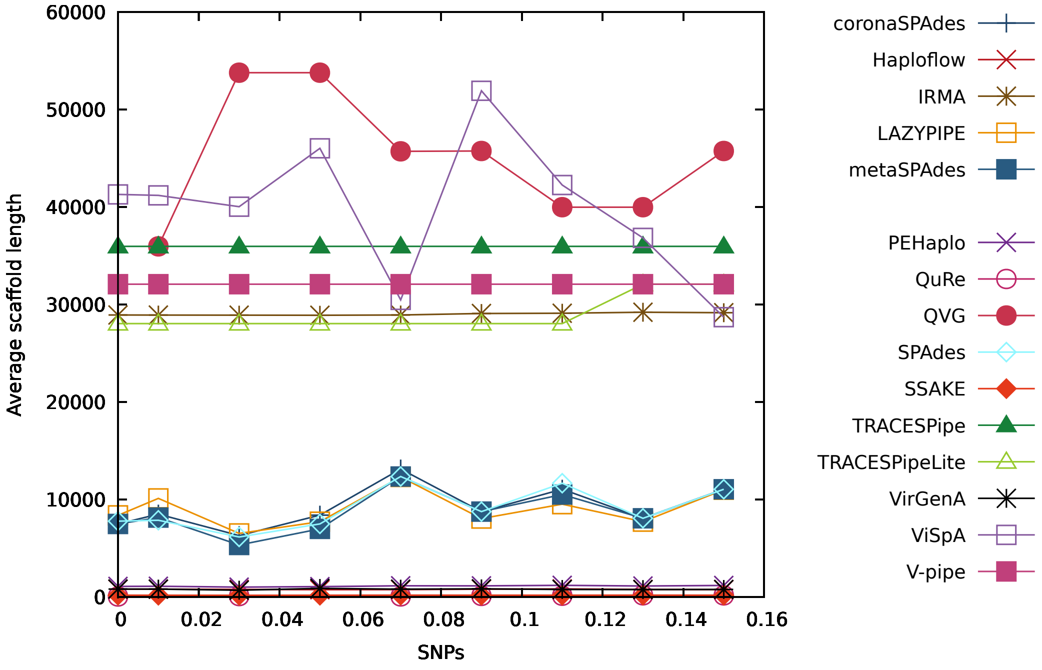


Figure S53: Figure comparing the performance of the reconstruction programs in terms of the average number of reconstructed bases per scaffold for datasets with depth coverage equal to 10x (DS33 to DS40 plus DS11).

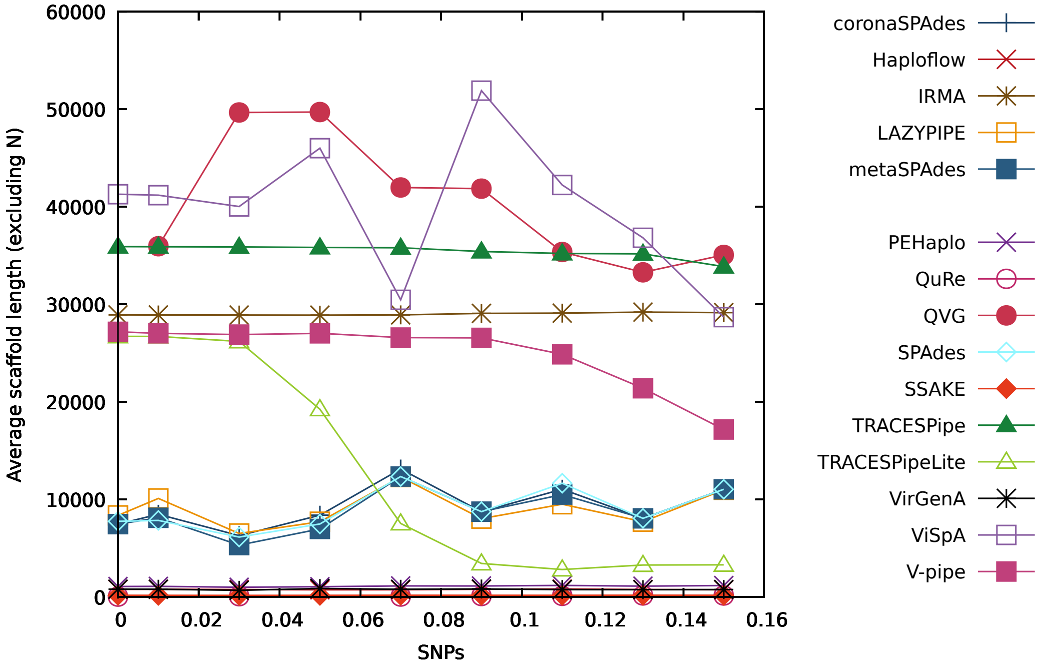


Figure S54: Figure comparing the performance of the reconstruction programs in terms of the average number of reconstructed bases per scaffold (excluding "N") for datasets with depth coverage equal to 10x (DS33 to DS40 plus DS11).

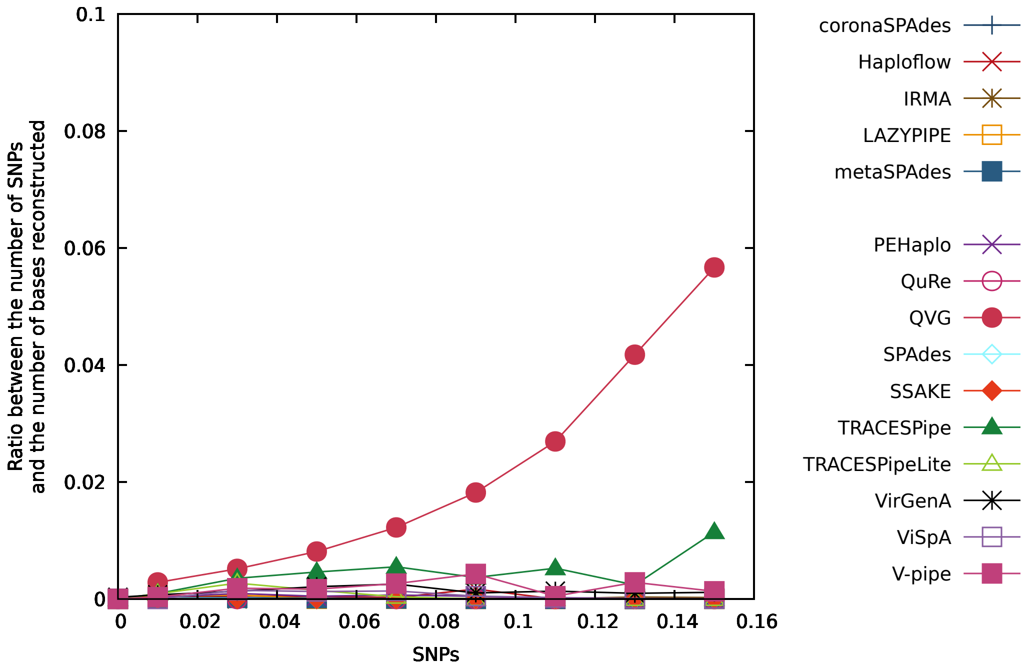


Figure S55: Figure comparing the performance of the reconstruction programs in terms of the ratio between the number of SNPs and the number of bases reconstructed for datasets with depth coverage equal to 10x (DS33 to DS40 plus DS11).

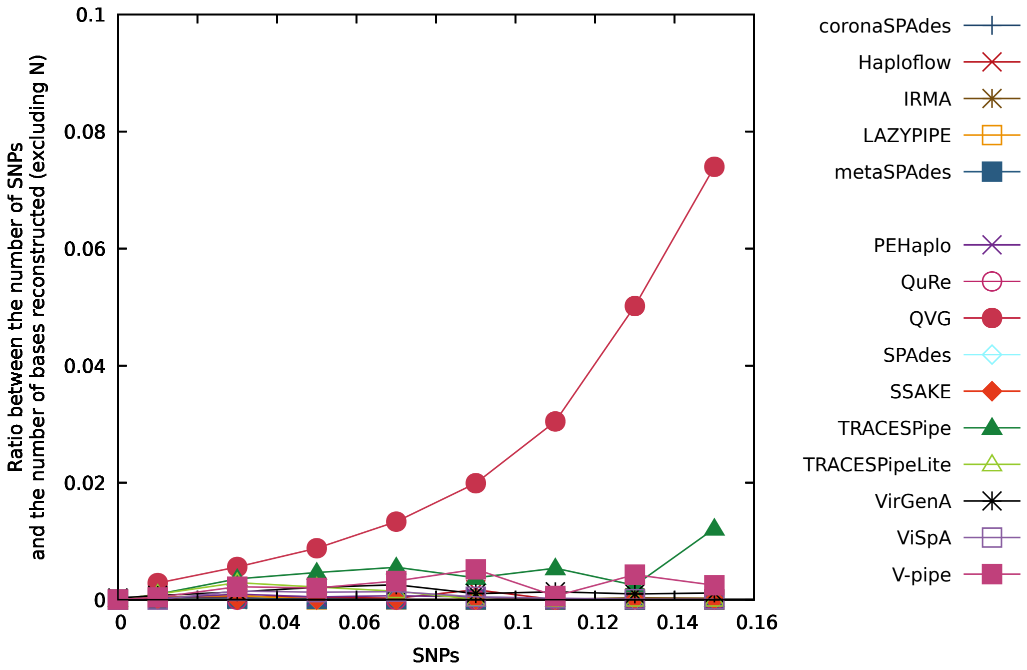


Figure S56: Figure comparing the performance of the reconstruction programs in terms of the ratio between the number of SNPs and the number of bases reconstructed (excluding "N") for datasets with depth coverage equal to 10x (DS33 to DS40 plus DS11).

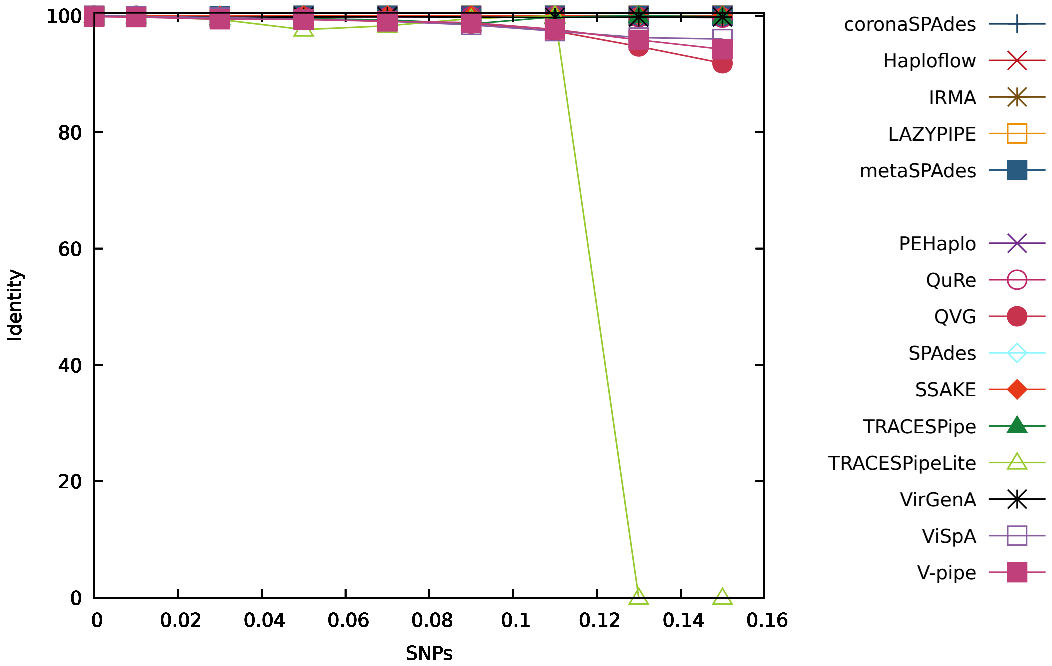


Figure S57: Figure comparing the performance of the reconstruction programs in terms of the identity for datasets with depth coverage equal to 20x (DS41 to DS48 plus DS13).

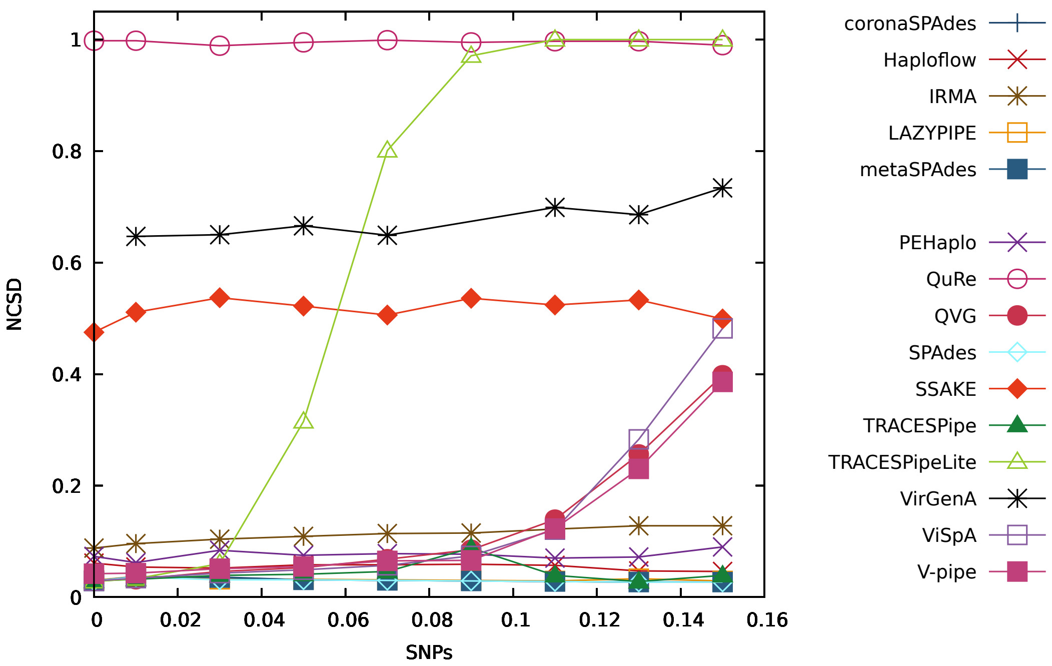


Figure S58: Figure comparing the performance of the reconstruction programs in terms of the NCSD for datasets with depth coverage equal to 20x (DS41 to DS48 plus DS13).

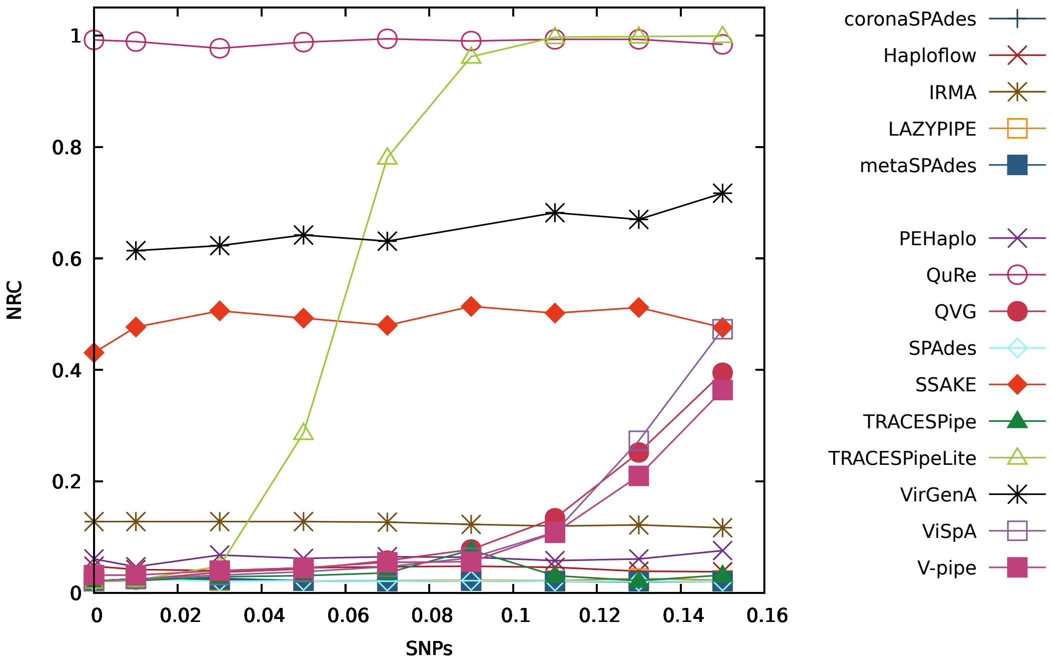


Figure S59: Figure comparing the performance of the reconstruction programs in terms of the NRC for datasets with depth coverage equal to 20x (DS41 to DS48 plus DS13).

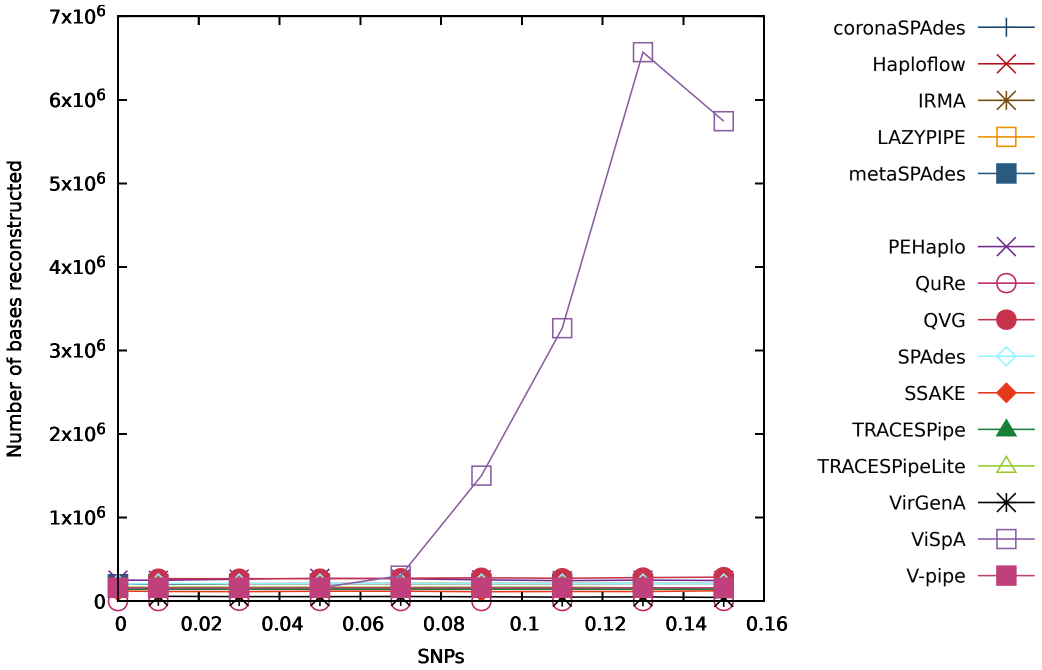


Figure S60: Figure comparing the performance of the reconstruction programs in terms of the number of bases reconstructed for datasets with depth coverage equal to 20x (DS41 to DS48 plus DS13).

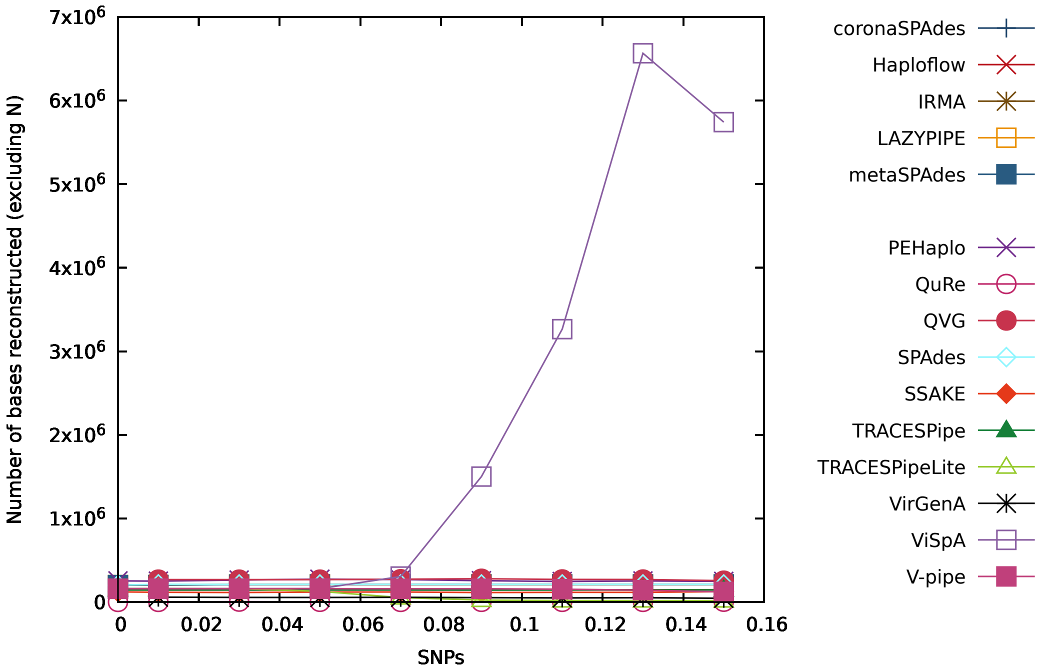


Figure S61: Figure comparing the performance of the reconstruction programs in terms of the number of bases reconstructed (excluding "N") for datasets with depth coverage equal to 20x (DS41 to DS48 plus DS13).

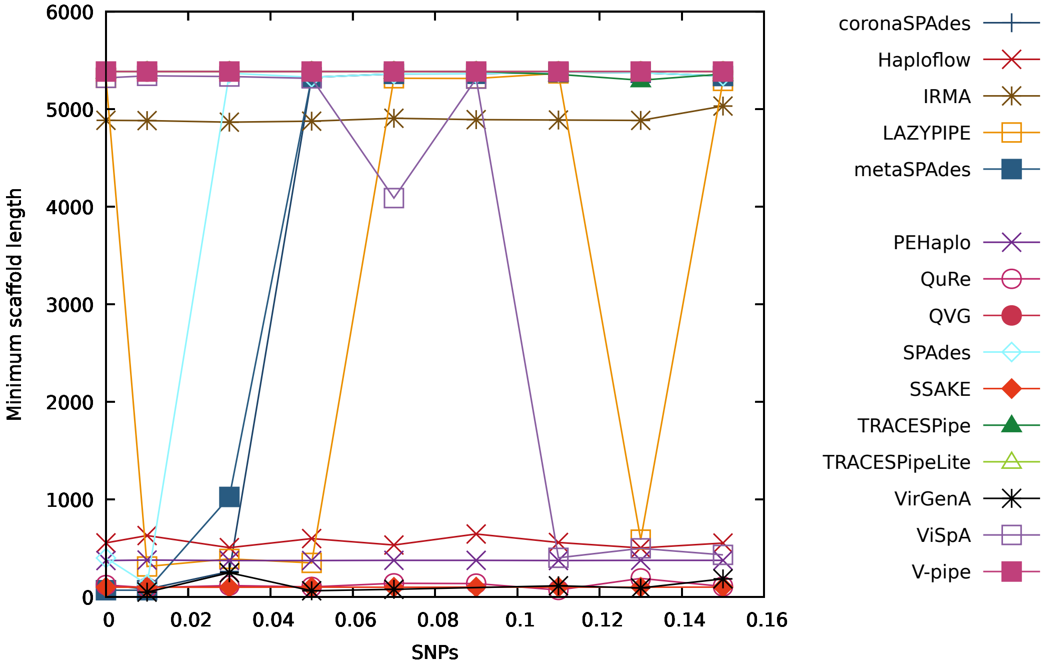


Figure S62: Figure comparing the performance of the reconstruction programs in terms of the minimum number of reconstructed bases per scaffold for datasets with depth coverage equal to 20x (DS41 to DS48 plus DS13).

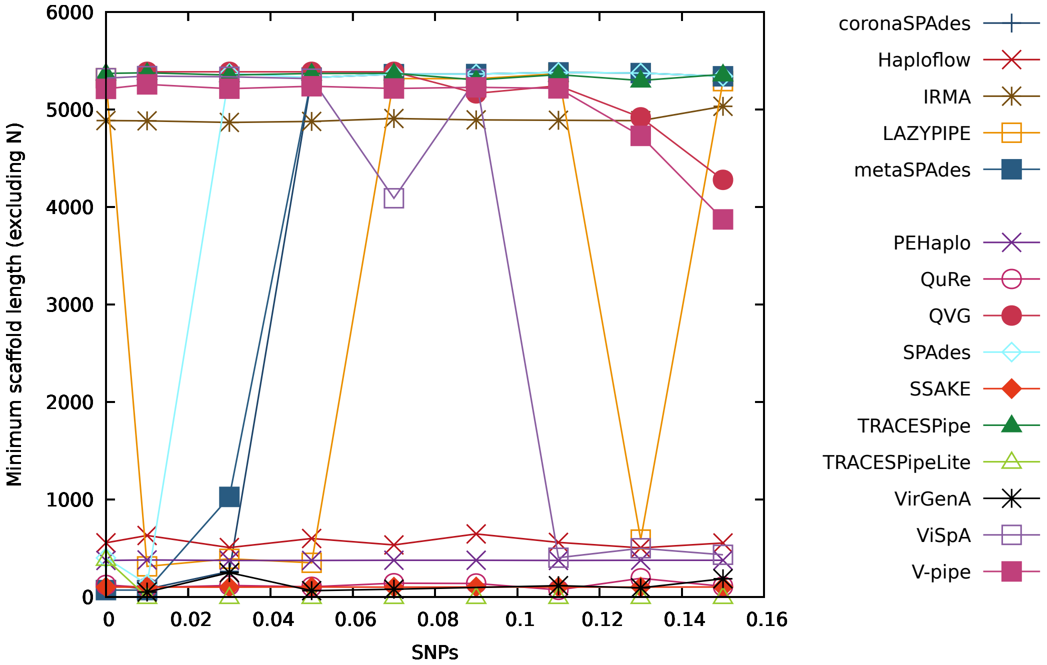


Figure S63: Figure comparing the performance of the reconstruction programs in terms of the minimum number of reconstructed bases per scaffold (excluding "N") for datasets with depth coverage equal to 20x (DS41 to DS48 plus DS13).

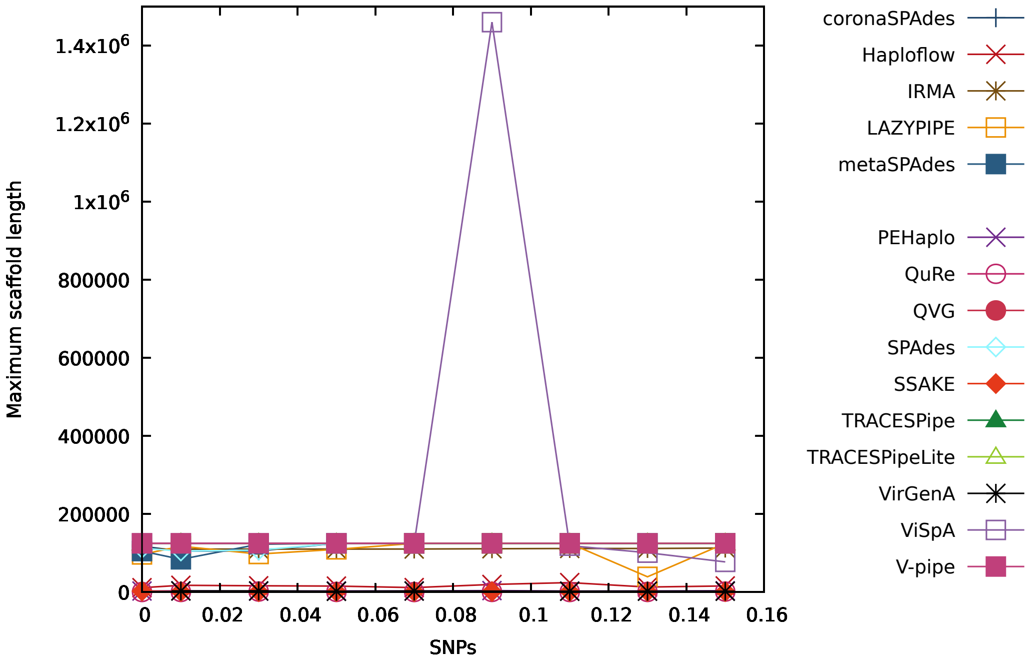


Figure S64: Figure comparing the performance of the reconstruction programs in terms of the maximum number of reconstructed bases per scaffold for datasets with depth coverage equal to 20x (DS41 to DS48 plus DS13).

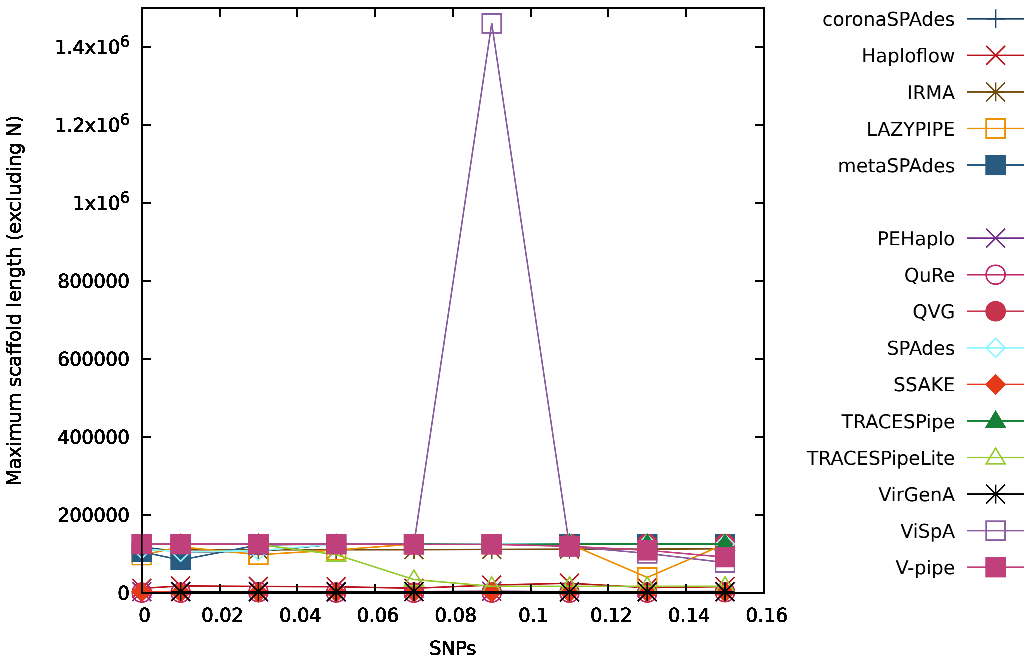


Figure S65: Figure comparing the performance of the reconstruction programs in terms of the maximum number of reconstructed bases per scaffold (excluding "N") for datasets with depth coverage equal to 20x (DS41 to DS48 plus DS13).

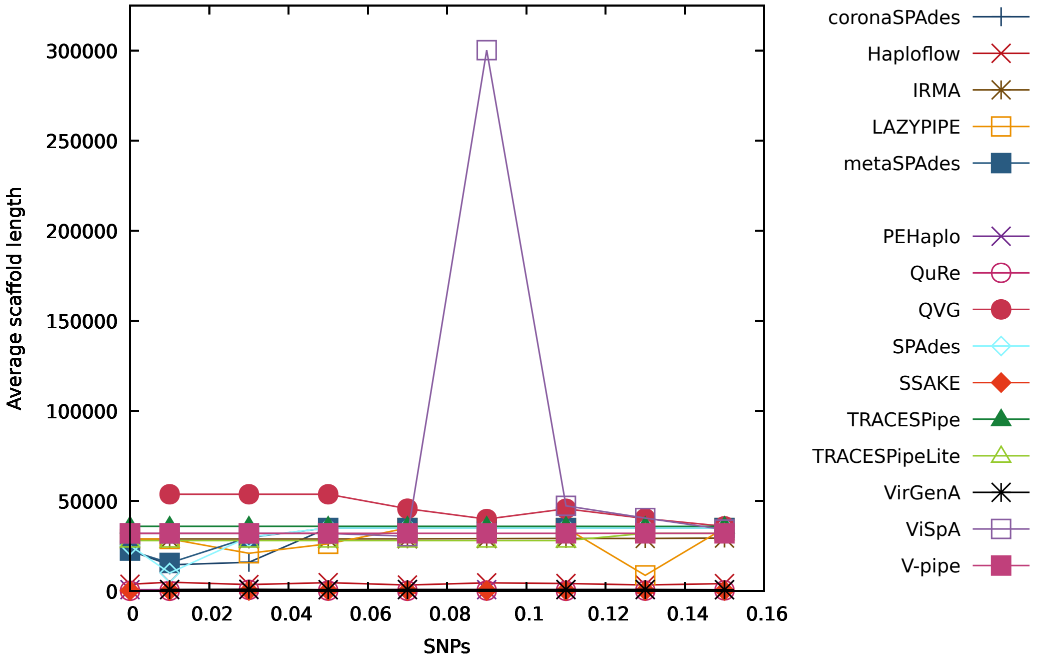


Figure S66: Figure comparing the performance of the reconstruction programs in terms of the average number of reconstructed bases per scaffold for datasets with depth coverage equal to 20x (DS41 to DS48 plus DS13).

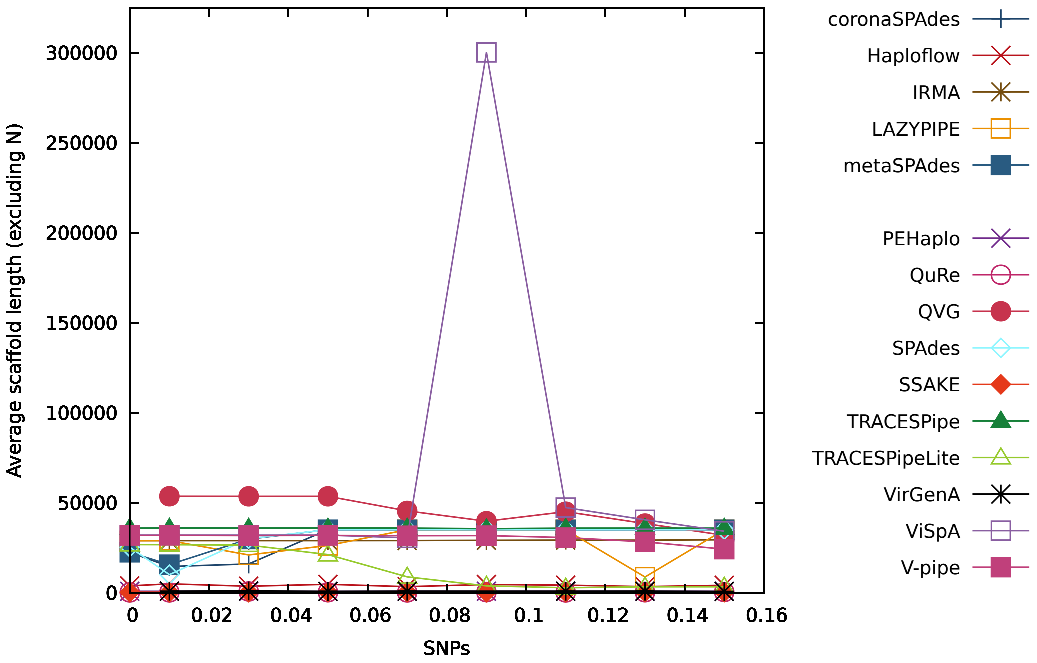


Figure S67: Figure comparing the performance of the reconstruction programs in terms of the average number of reconstructed bases per scaffold (excluding "N") for datasets with depth coverage equal to 20x (DS41 to DS48 plus DS13).

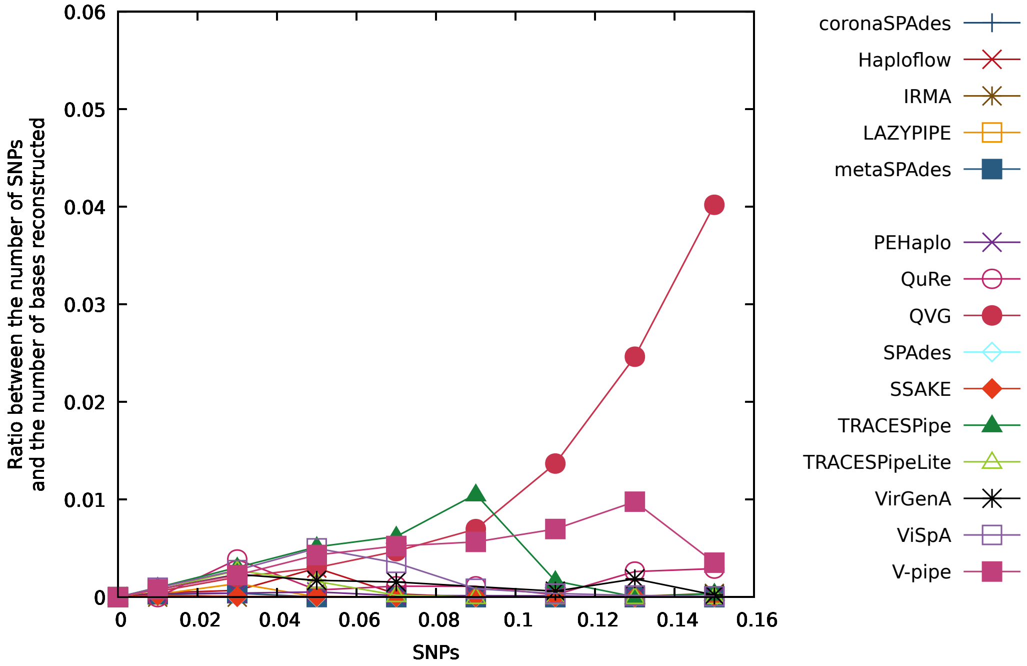


Figure S68: Figure comparing the performance of the reconstruction programs in terms of the ratio between the number of SNPs and the number of bases reconstructed for datasets with depth coverage equal to 20x (DS41 to DS48 plus DS13).

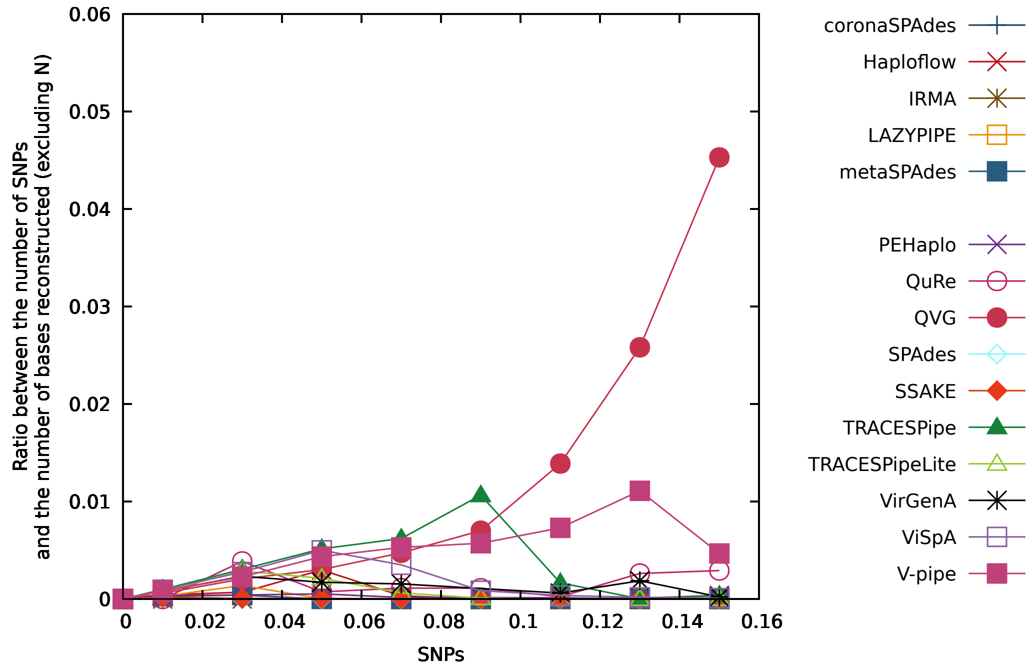


Figure S69: Figure comparing the performance of the reconstruction programs in terms of the ratio between the number of SNPs and the number of bases reconstructed (excluding "N") for datasets with depth coverage equal to 20x (DS41 to DS48 plus DS13).

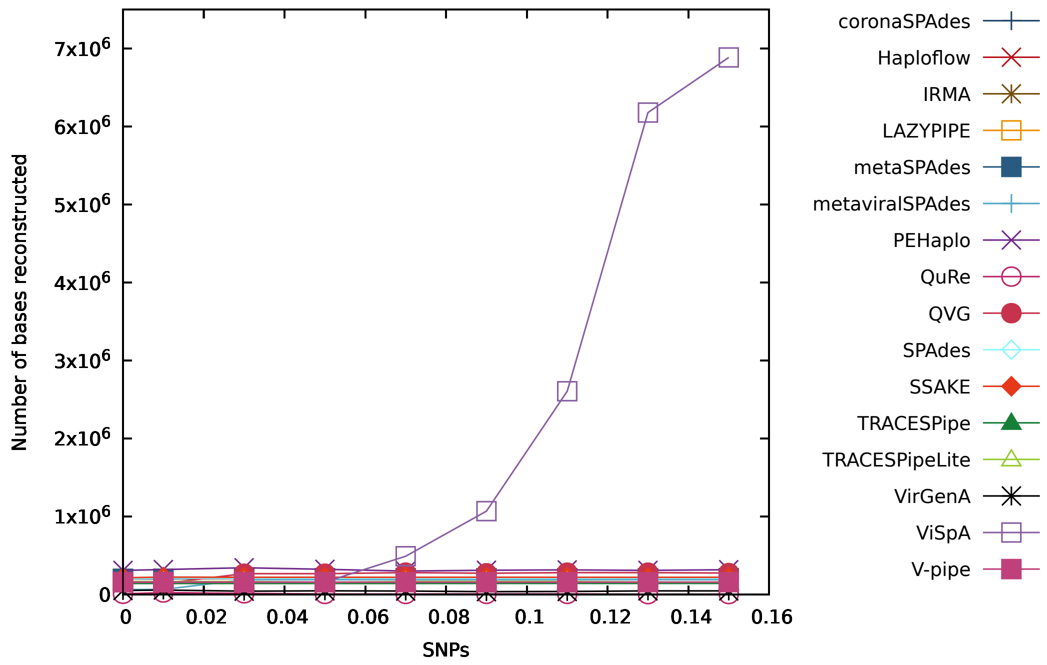


Figure S70: Figure comparing the performance of the reconstruction programs in terms of the number of bases reconstructed for datasets with depth coverage equal to 40x (DS49 to DS56 plus DS16).

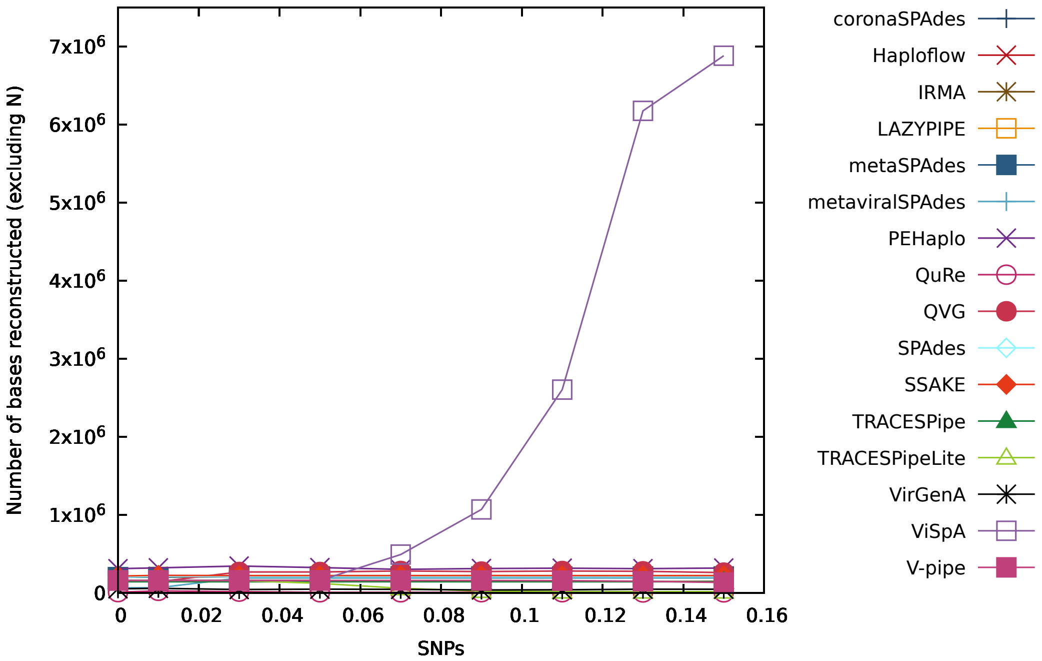


Figure S71: Figure comparing the performance of the reconstruction programs in terms of the number of bases reconstructed (excluding "N") for datasets with depth coverage equal to 40x (DS49 to DS56 plus DS16).

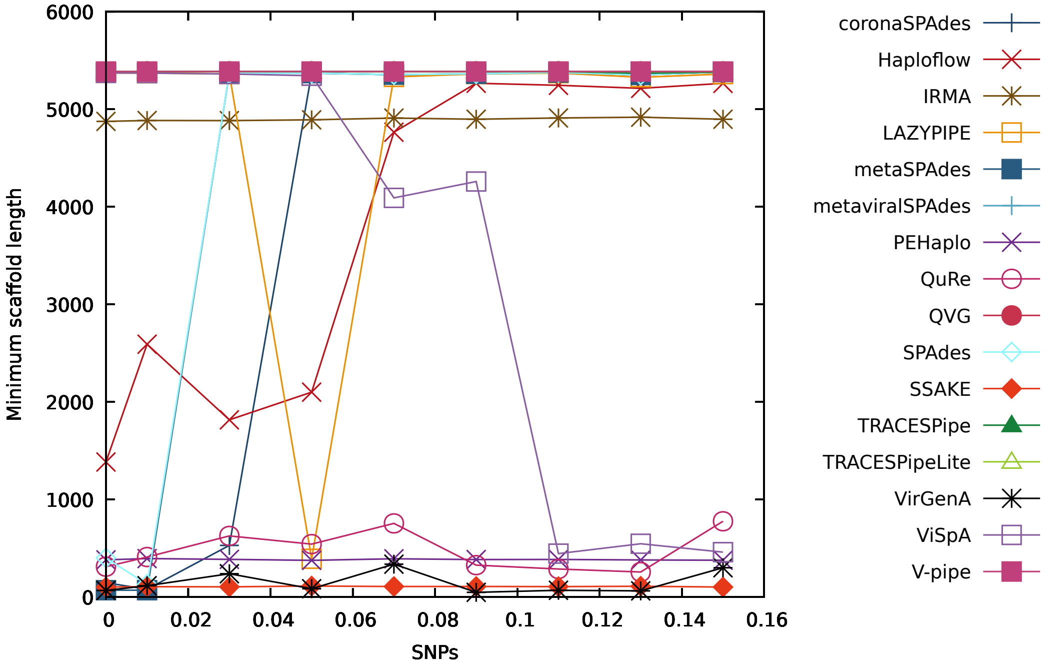


Figure S72: Figure comparing the performance of the reconstruction programs in terms of the minimum number of reconstructed bases per scaffold for datasets with depth coverage equal to 40x (DS49 to DS56 plus DS16).

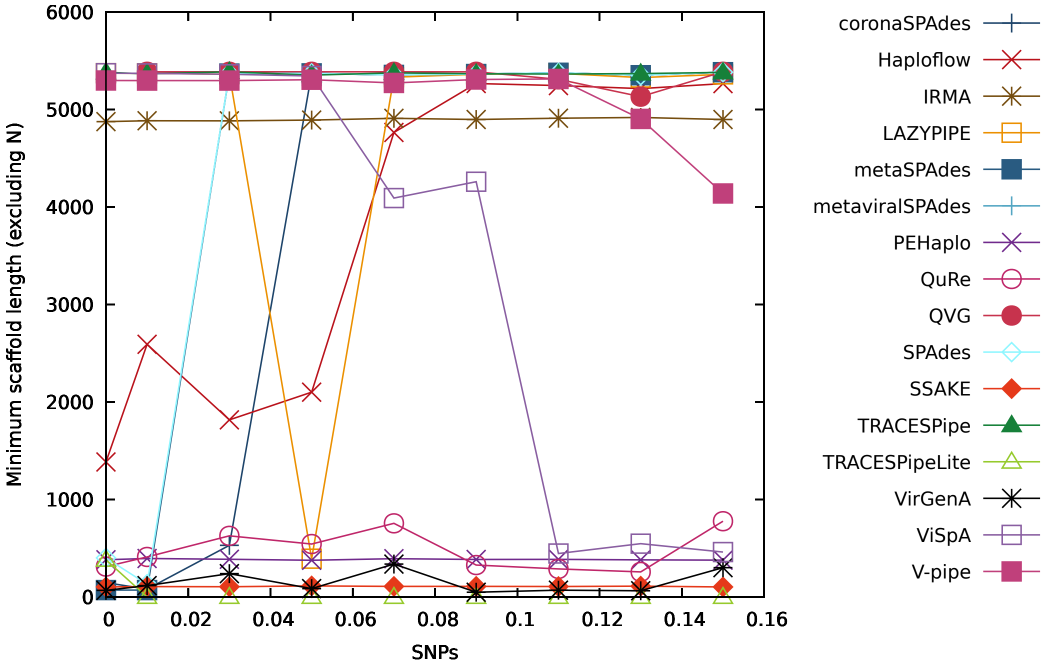


Figure S73: Figure comparing the performance of the reconstruction programs in terms of the minimum number of reconstructed bases per scaffold (excluding "N") for datasets with depth coverage equal to 40x (DS49 to DS56 plus DS16).

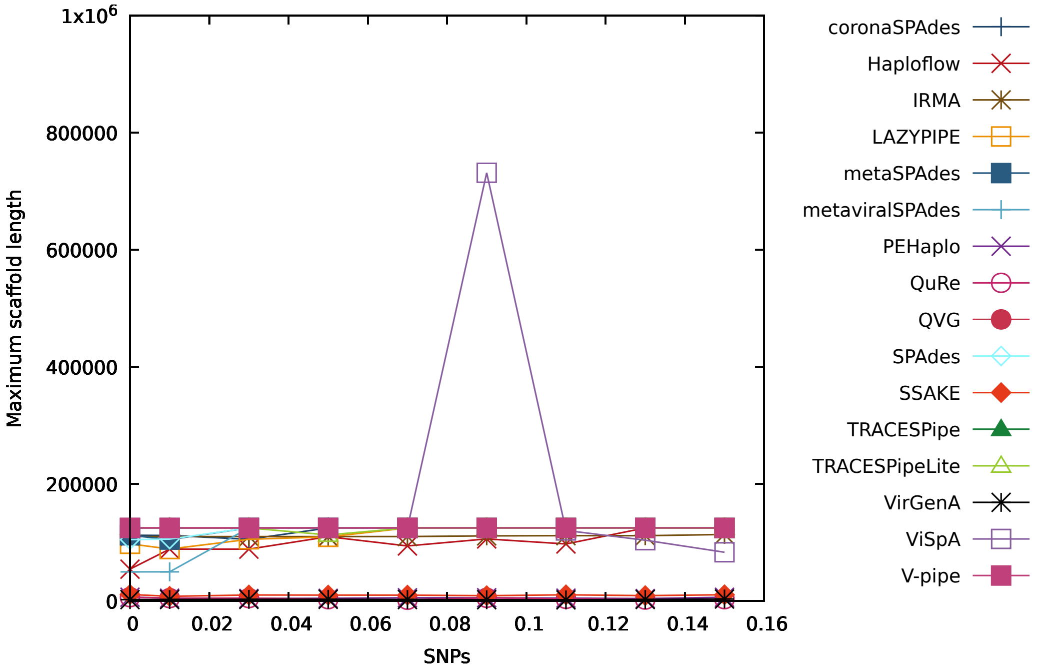


Figure S74: Figure comparing the performance of the reconstruction programs in terms of the maximum number of reconstructed bases per scaffold for datasets with depth coverage equal to 40x (DS49 to DS56 plus DS16).

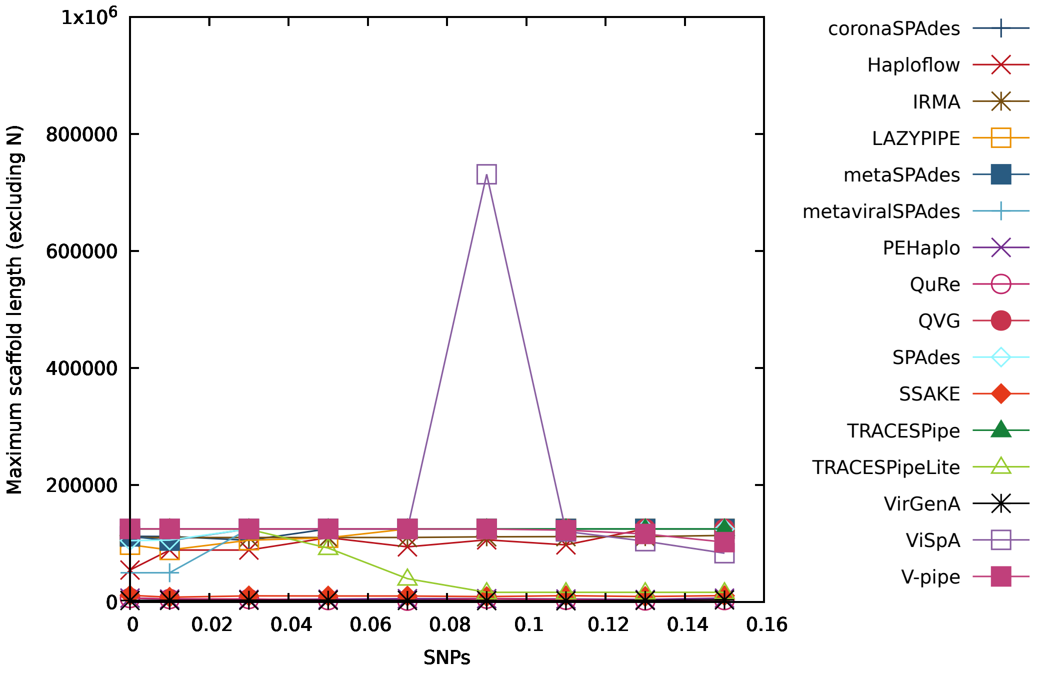


Figure S75: Figure comparing the performance of the reconstruction programs in terms of the maximum number of reconstructed bases per scaffold (excluding "N") for datasets with depth coverage equal to 40x (DS49 to DS56 plus DS16).

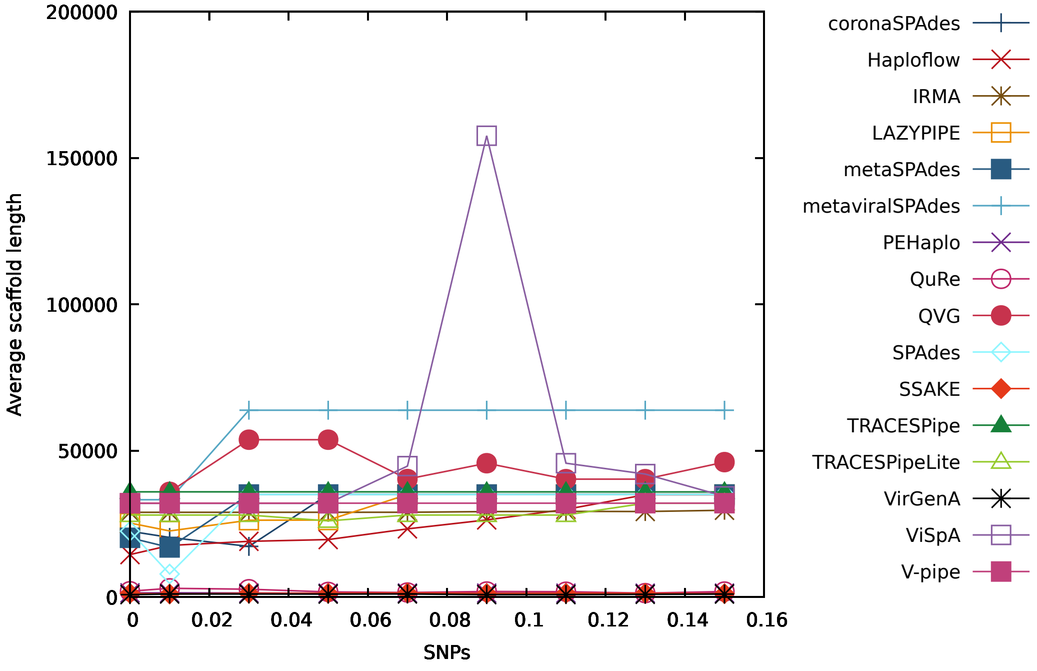


Figure S76: Figure comparing the performance of the reconstruction programs in terms of the average number of reconstructed bases per scaffold for datasets with depth coverage equal to 40x (DS49 to DS56 plus DS16).

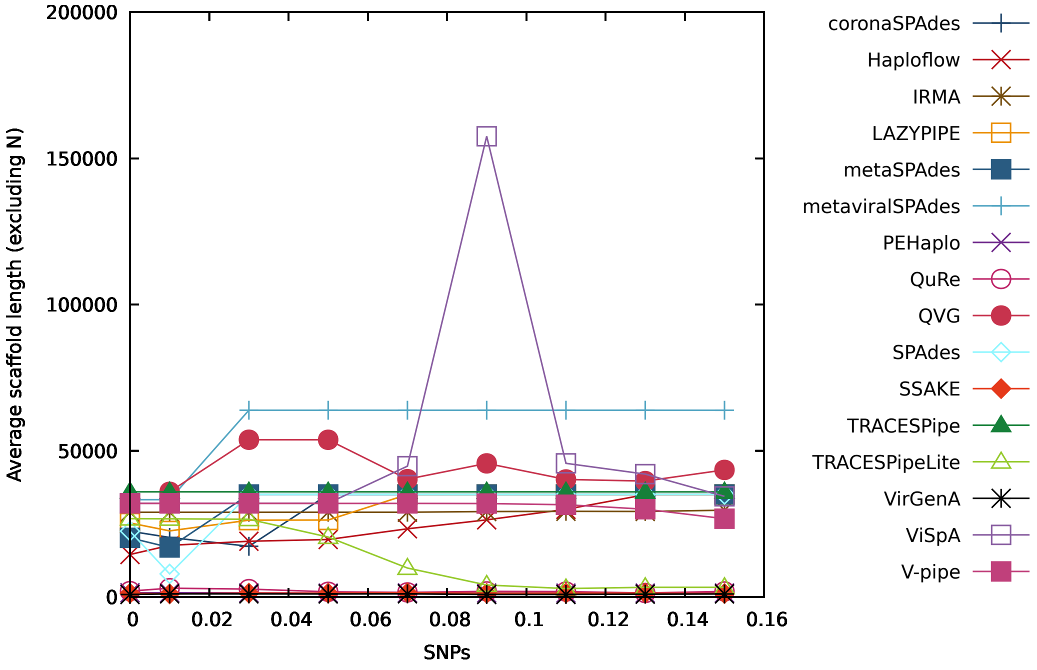


Figure S77: Figure comparing the performance of the reconstruction programs in terms of the average number of reconstructed bases per scaffold (excluding "N") for datasets with depth coverage equal to 40x (DS49 to DS56 plus DS16).

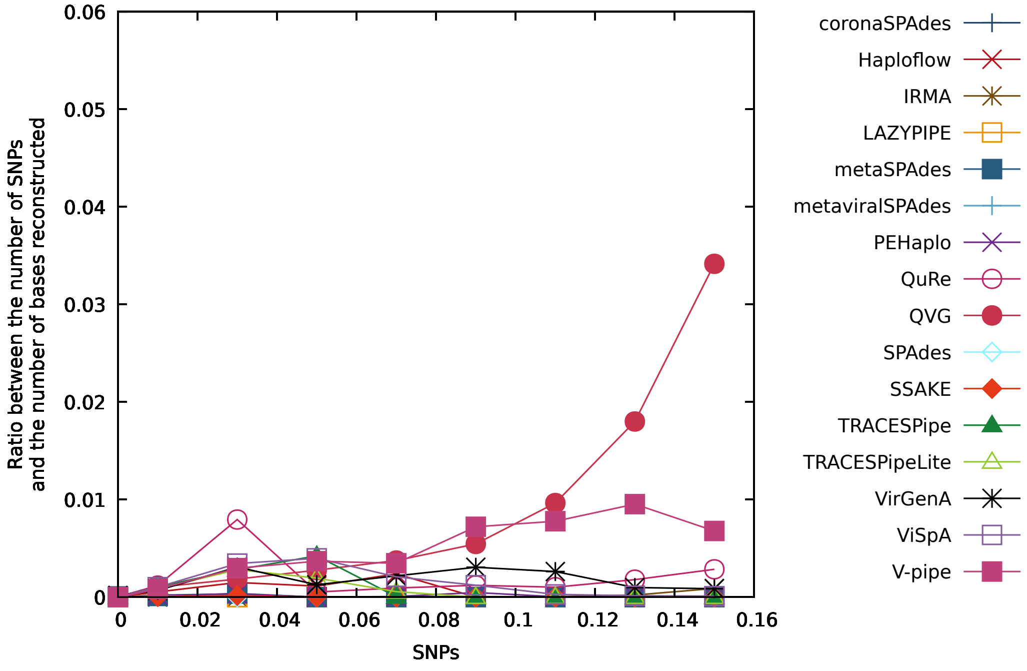


Figure S78: Figure comparing the performance of the reconstruction programs in terms of the ratio between the number of SNPs and the number of bases reconstructed for datasets with depth coverage equal to 40x (DS49 to DS56 plus DS16).

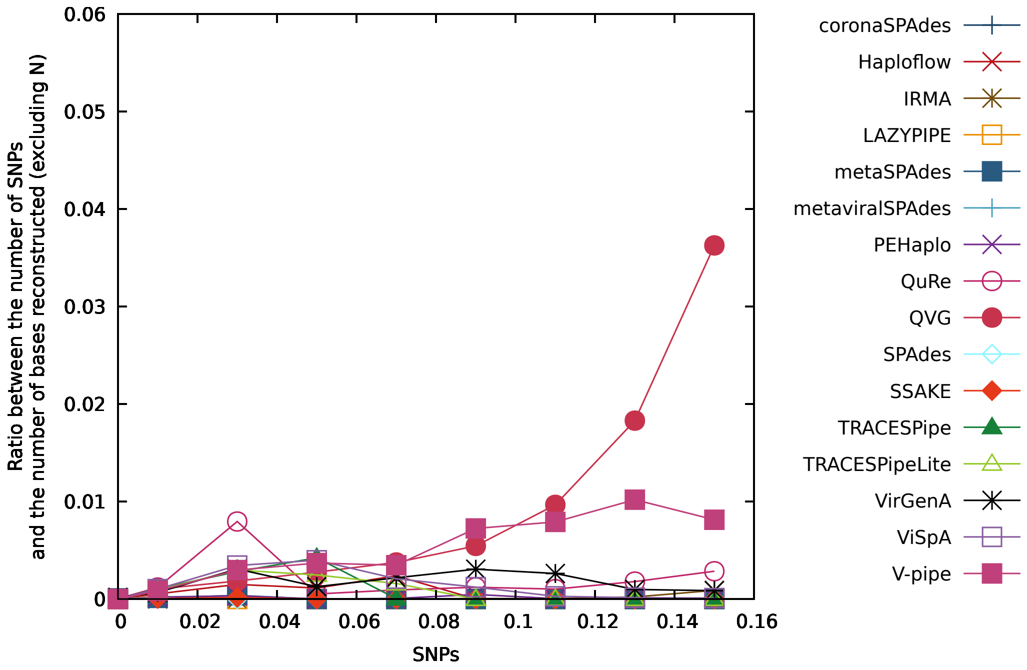


Figure S79: Figure comparing the performance of the reconstruction programs in terms of the ratio between the number of SNPs and the number of bases reconstructed (excluding "N") for datasets with depth coverage equal to 40x (DS49 to DS56 plus DS16).

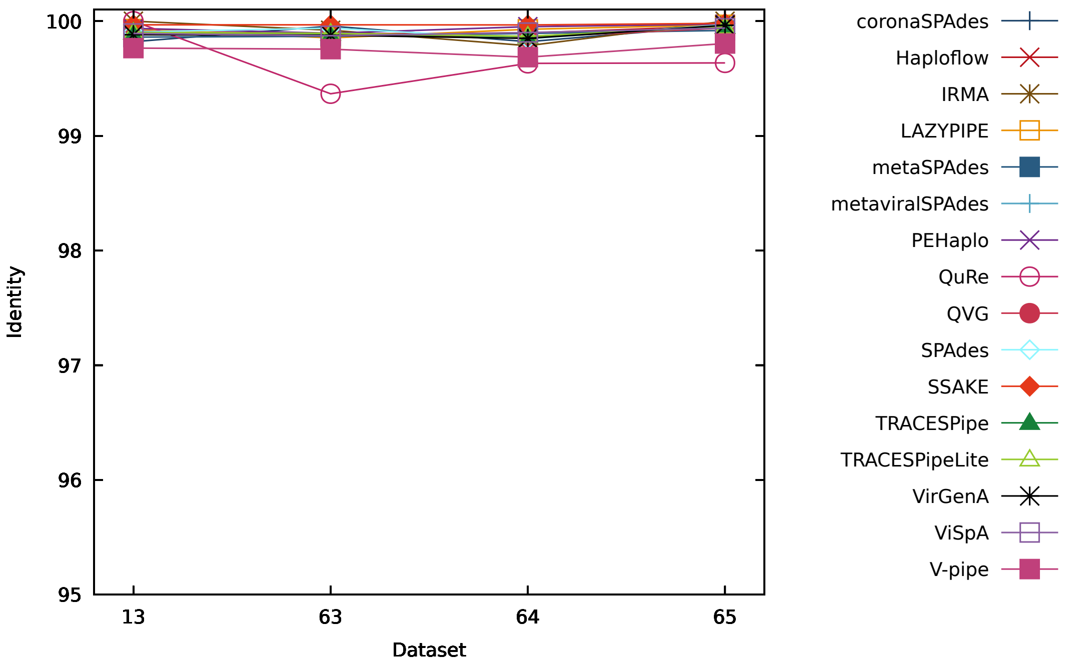


Figure S80: Figure comparing the performance of the reconstruction programs in terms of the identity for datasets 13, 63, 64 and 65.

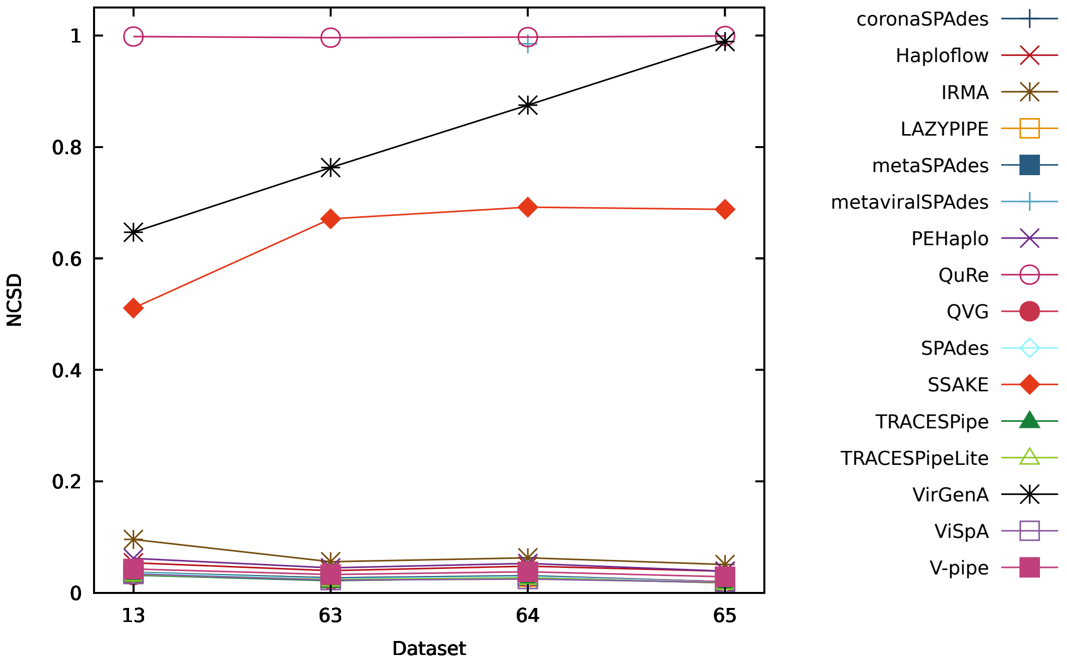


Figure S81: Figure comparing the performance of the reconstruction programs in terms of the NCSD for datasets 13, 63, 64 and 65.

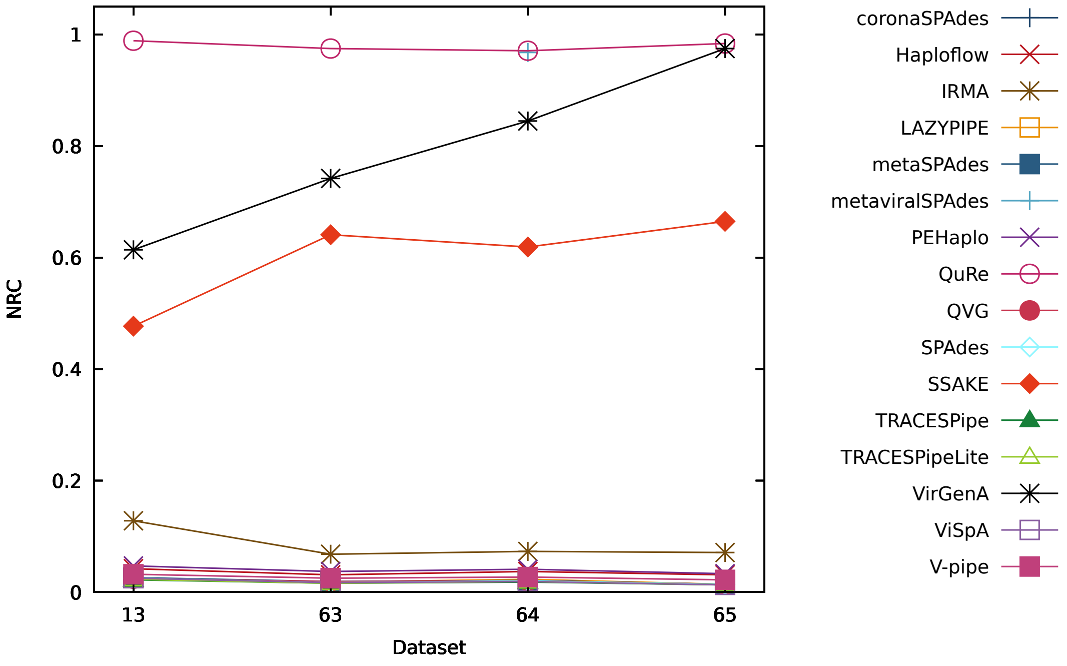


Figure S82: Figure comparing the performance of the reconstruction programs in terms of the NRC for datasets 13, 63, 64 and 65.

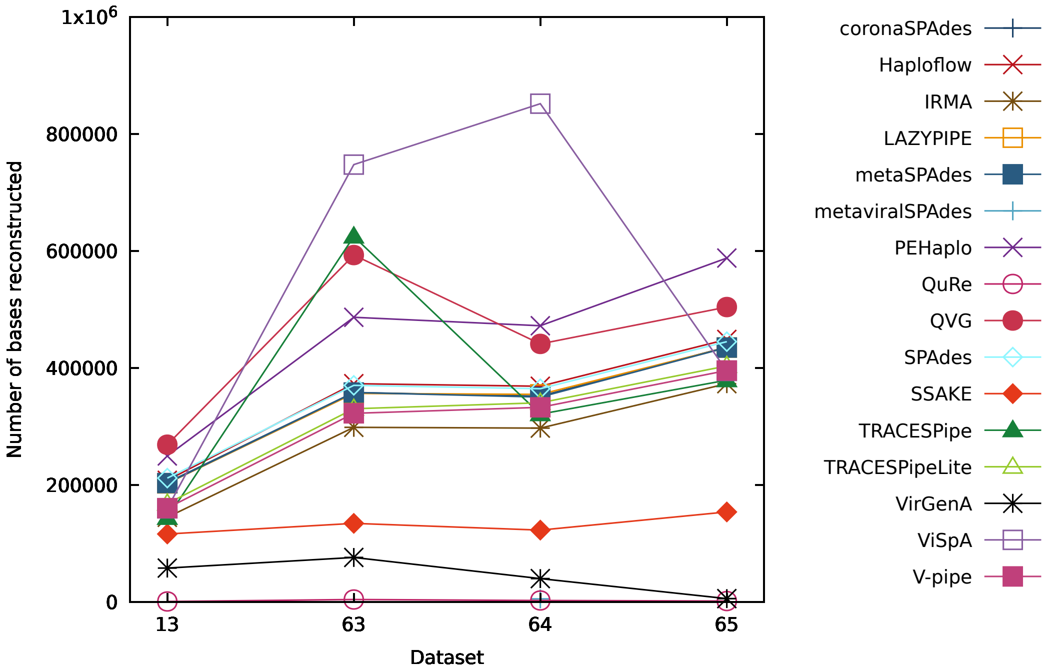


Figure S83: Figure comparing the performance of the reconstruction programs in terms of the number of bases reconstructed for datasets 13, 63, 64 and 65.

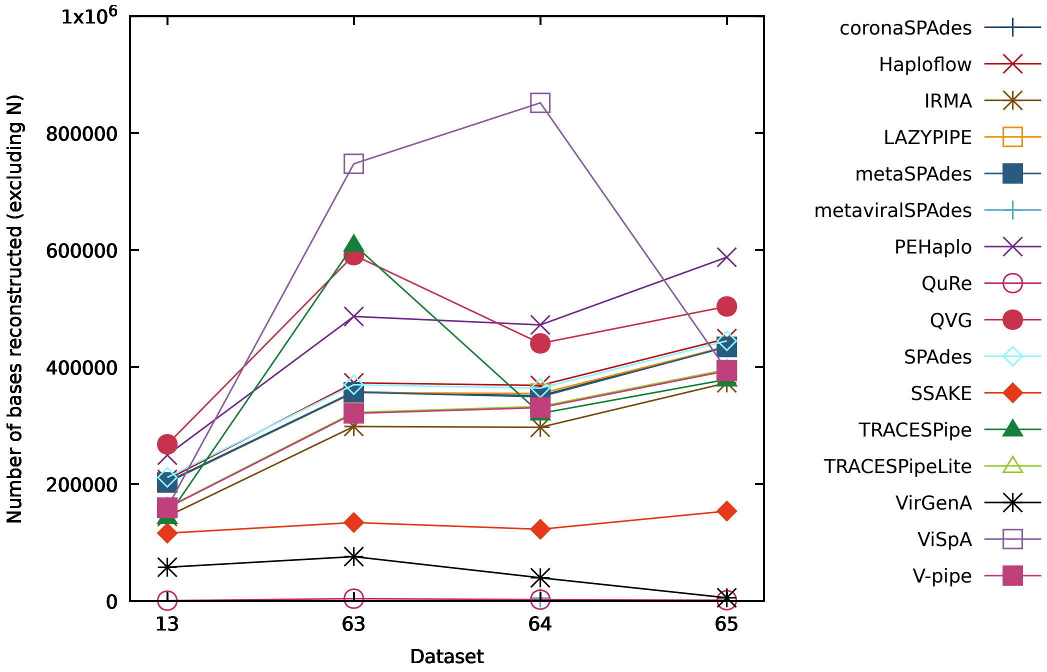


Figure S84: Figure comparing the performance of the reconstruction programs in terms of the number of bases reconstructed (excluding "N") for datasets 13, 63, 64 and 65.

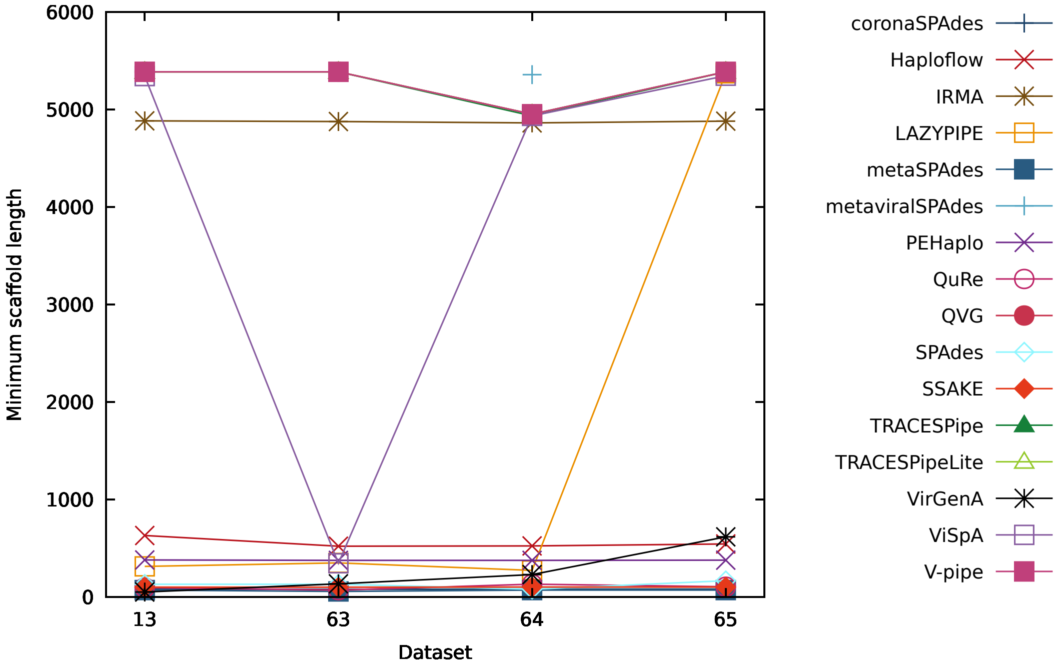


Figure S85: Figure comparing the performance of the reconstruction programs in terms of the minimum number of reconstructed bases per scaffold for datasets 13, 63, 64 and 65.

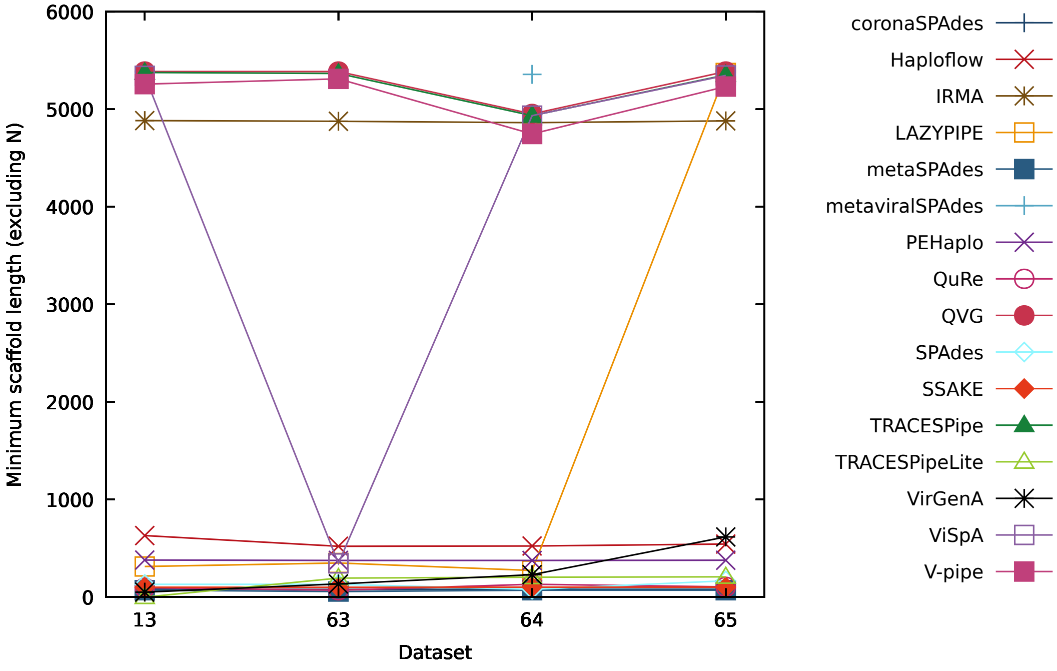


Figure S86: Figure comparing the performance of the reconstruction programs in terms of the minimum number of reconstructed bases per scaffold (excluding "N") for datasets 13, 63, 64 and 65.

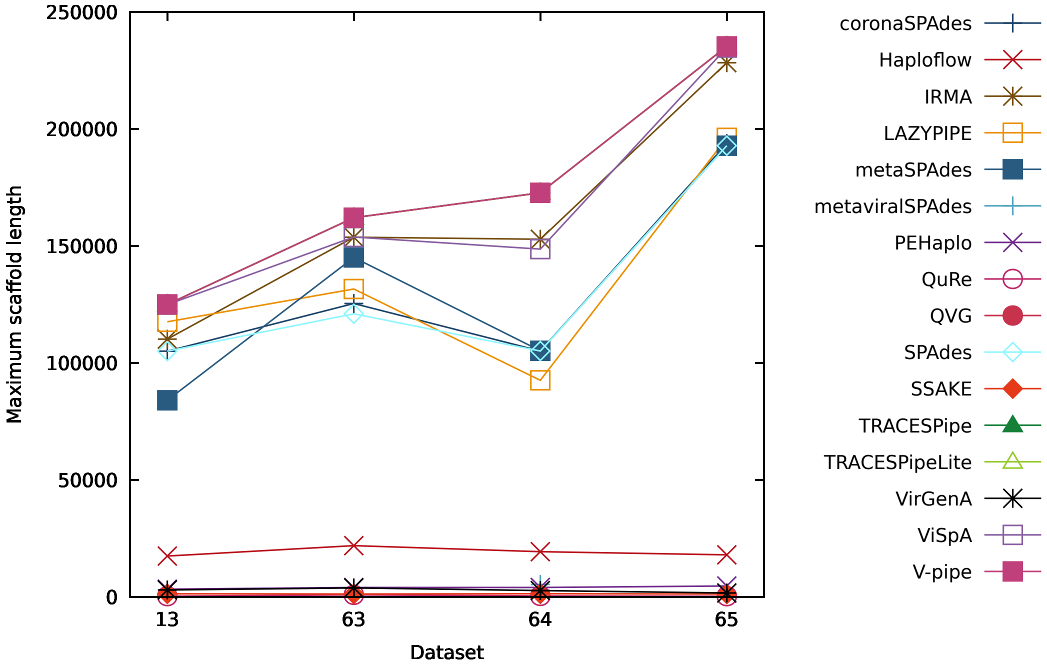


Figure S87: Figure comparing the performance of the reconstruction programs in terms of the maximum number of reconstructed bases per scaffold for datasets 13, 63, 64 and 65.

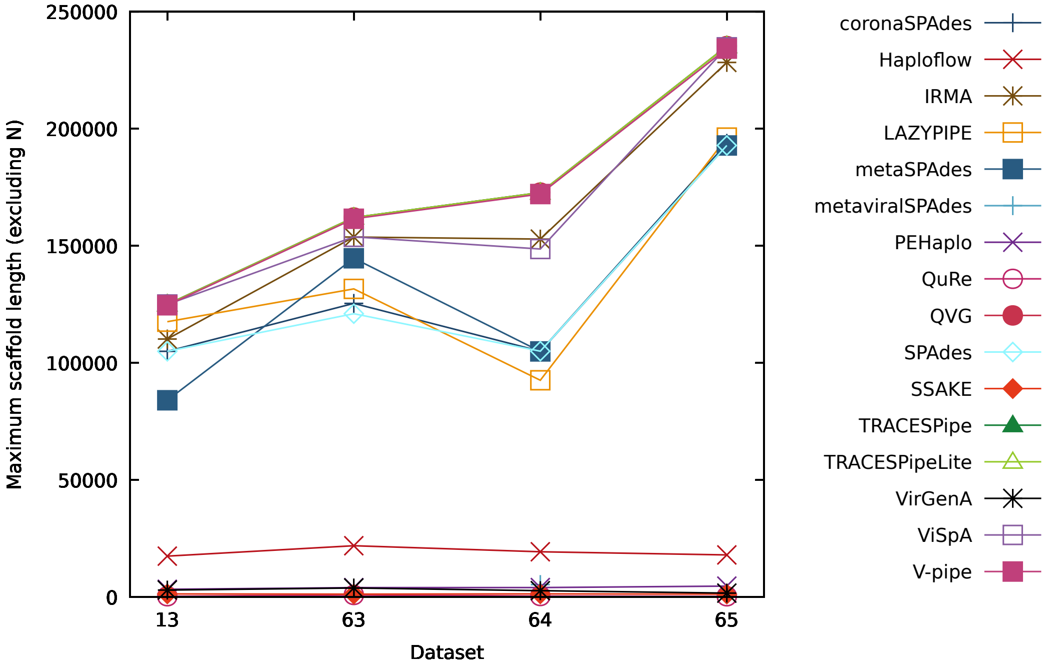


Figure S88: Figure comparing the performance of the reconstruction programs in terms of the maximum number of reconstructed bases per scaffold (excluding "N") for datasets 13, 63, 64 and 65.

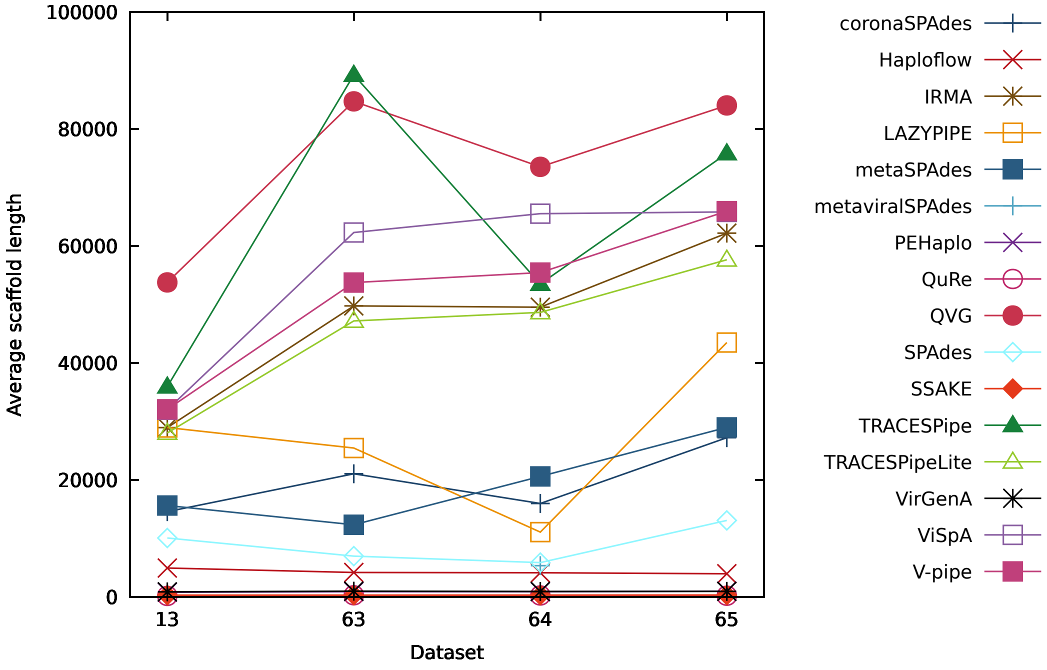


Figure S89: Figure comparing the performance of the reconstruction programs in terms of the average number of reconstructed bases per scaffold for datasets 13, 63, 64 and 65.

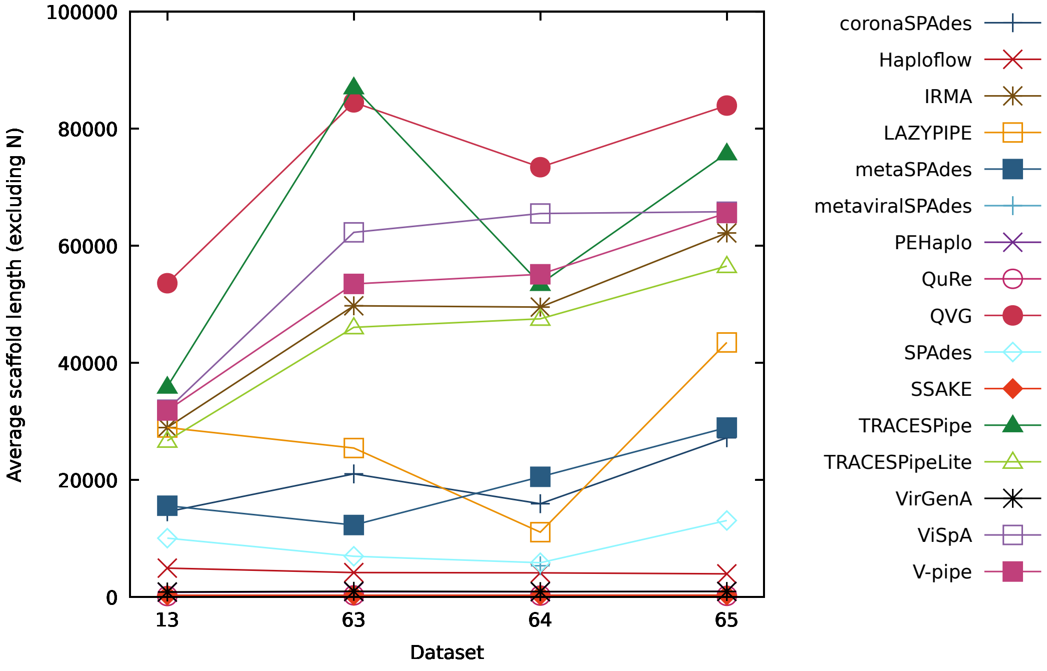


Figure S90: Figure comparing the performance of the reconstruction programs in terms of the average number of reconstructed bases per scaffold (excluding "N") for datasets 13, 63, 64 and 65.

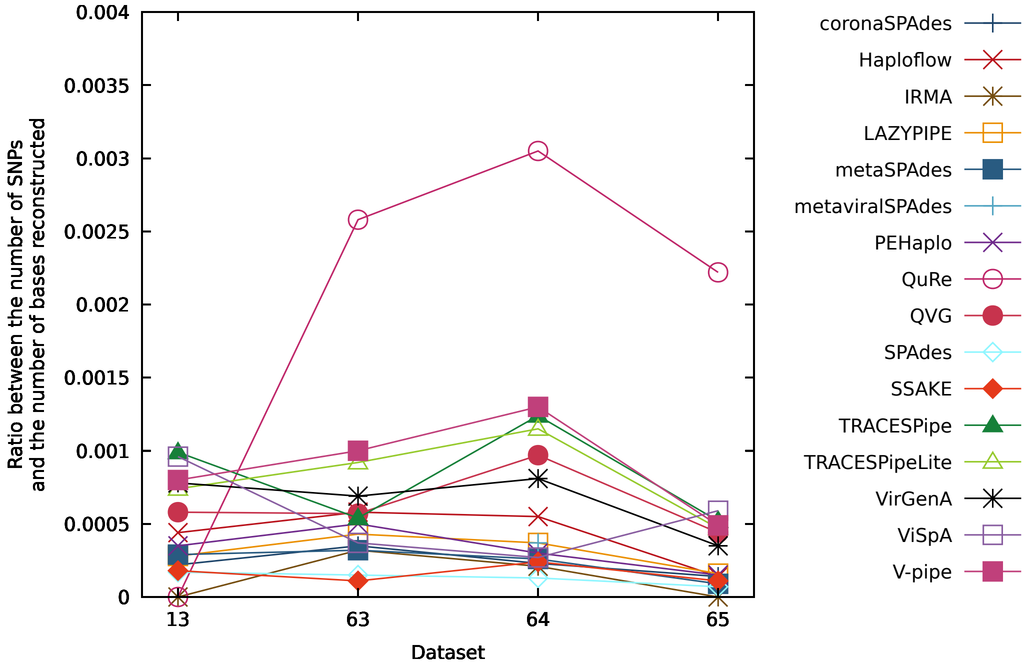


Figure S91: Figure comparing the performance of the reconstruction programs in terms of the ratio between the number of SNPs and the number of bases reconstructed for datasets 13, 63, 64 and 65.

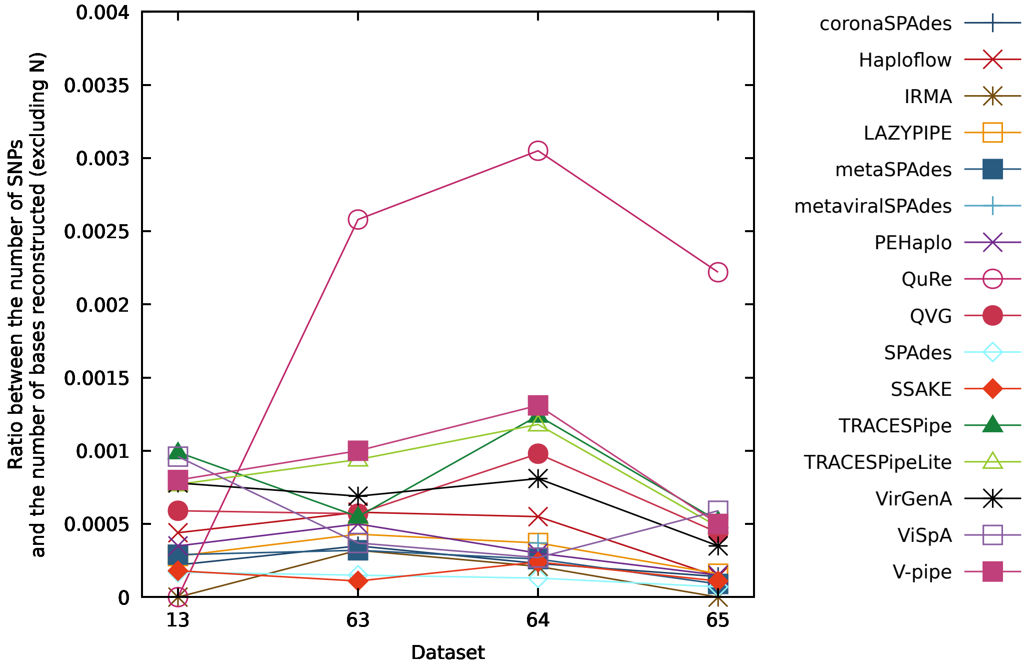


Figure S92: Figure comparing the performance of the reconstruction programs in terms of the ratio between the number of SNPs and the number of bases reconstructed (excluding "N") for datasets 13, 63, 64 and 65.

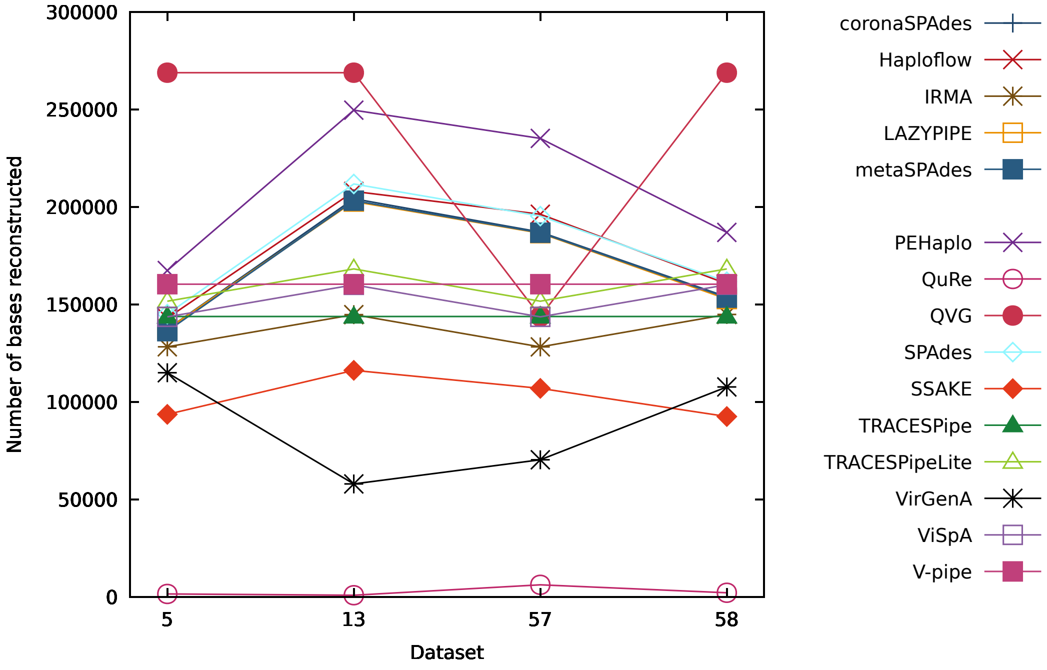


Figure S93: Figure comparing the performance of the reconstruction programs in terms of the number of bases reconstructed for datasets 5, 13, 57 and 58.

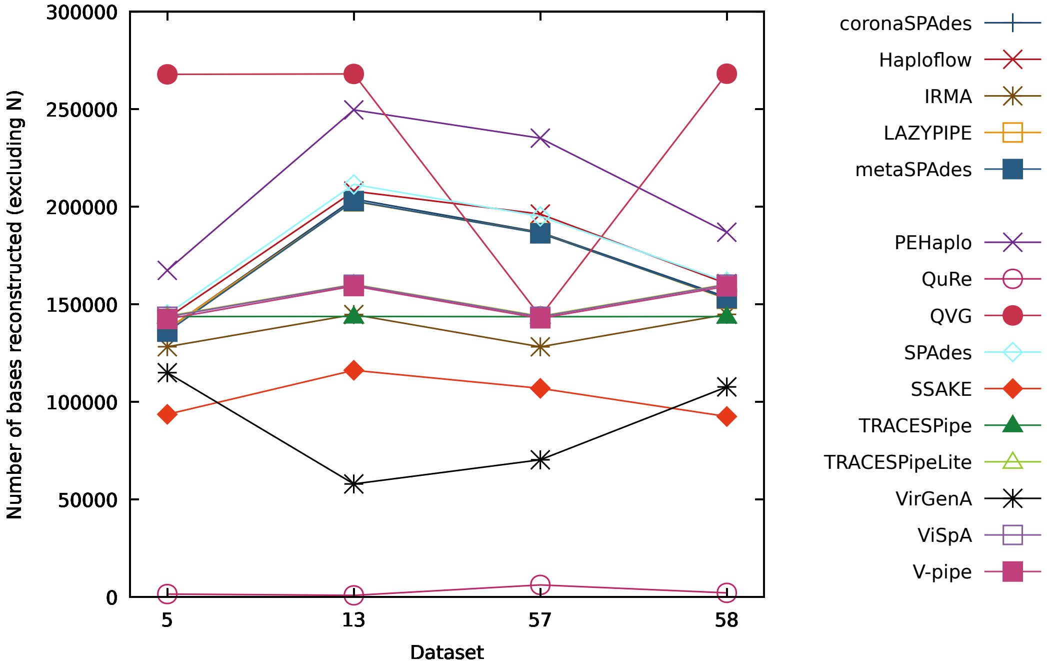


Figure S94: Figure comparing the performance of the reconstruction programs in terms of the number of bases reconstructed (excluding "N") for datasets 5, 13, 57 and 58.

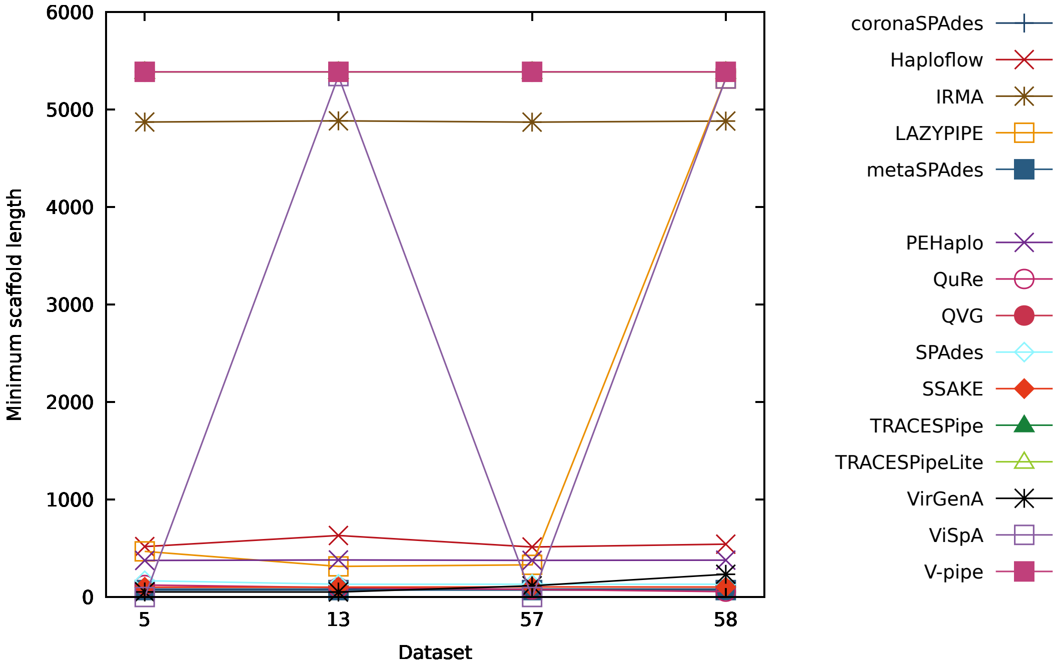


Figure S95: Figure comparing the performance of the reconstruction programs in terms of the minimum number of reconstructed bases per scaffold for datasets 5, 13, 57 and 58.

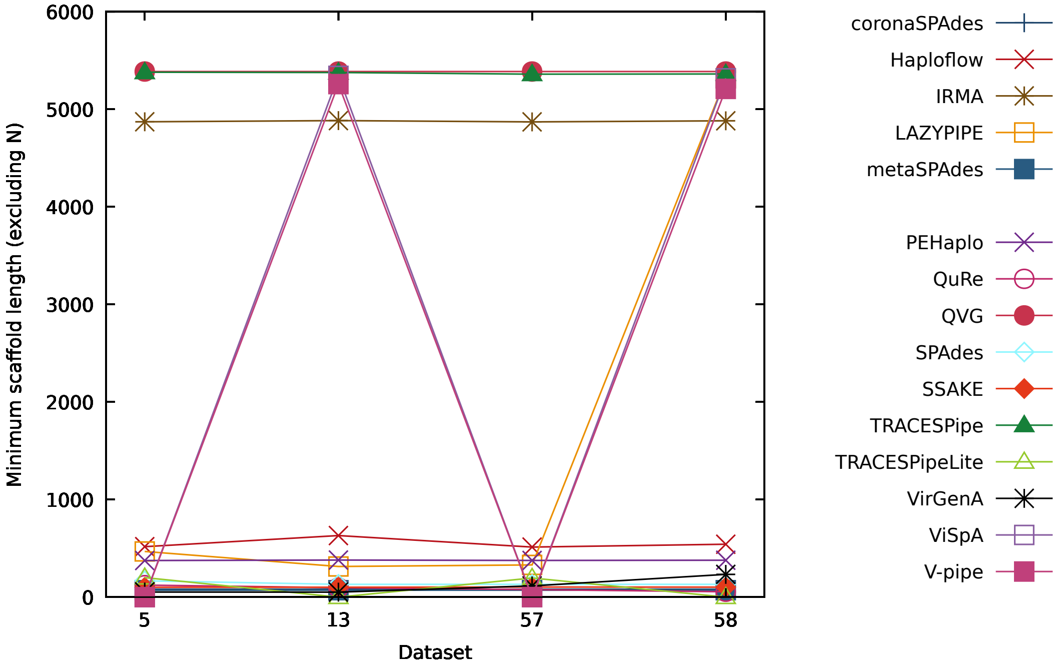


Figure S96: Figure comparing the performance of the reconstruction programs in terms of the minimum number of reconstructed bases per scaffold (excluding "N") for datasets 5, 13, 57 and 58.

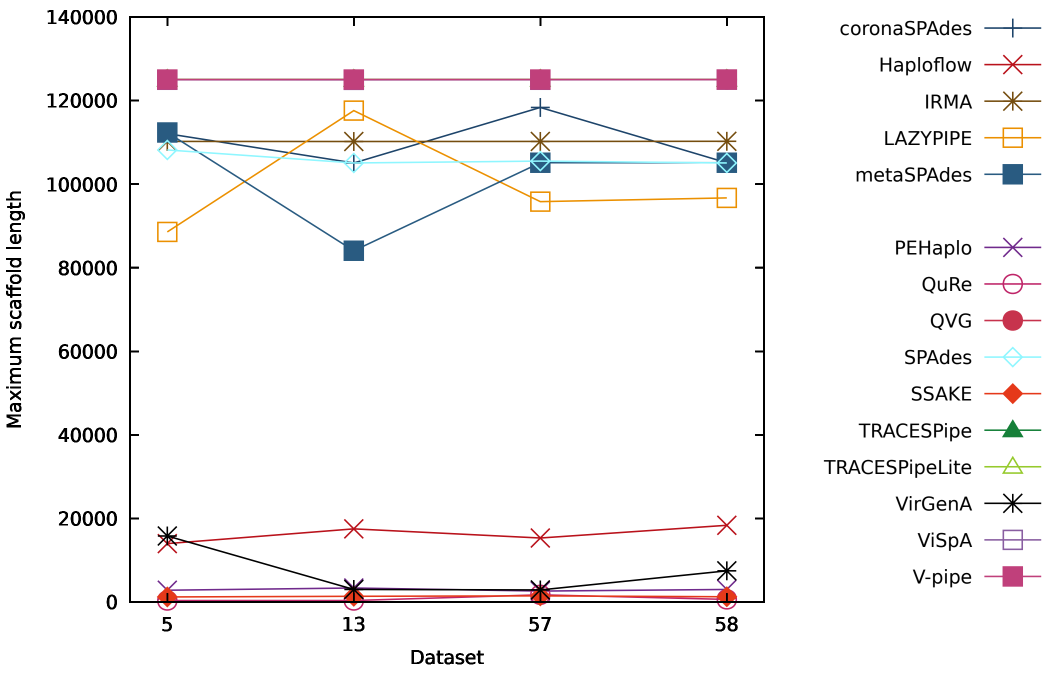


Figure S97: Figure comparing the performance of the reconstruction programs in terms of the maximum number of reconstructed bases per scaffold for datasets 5, 13, 57 and 58.

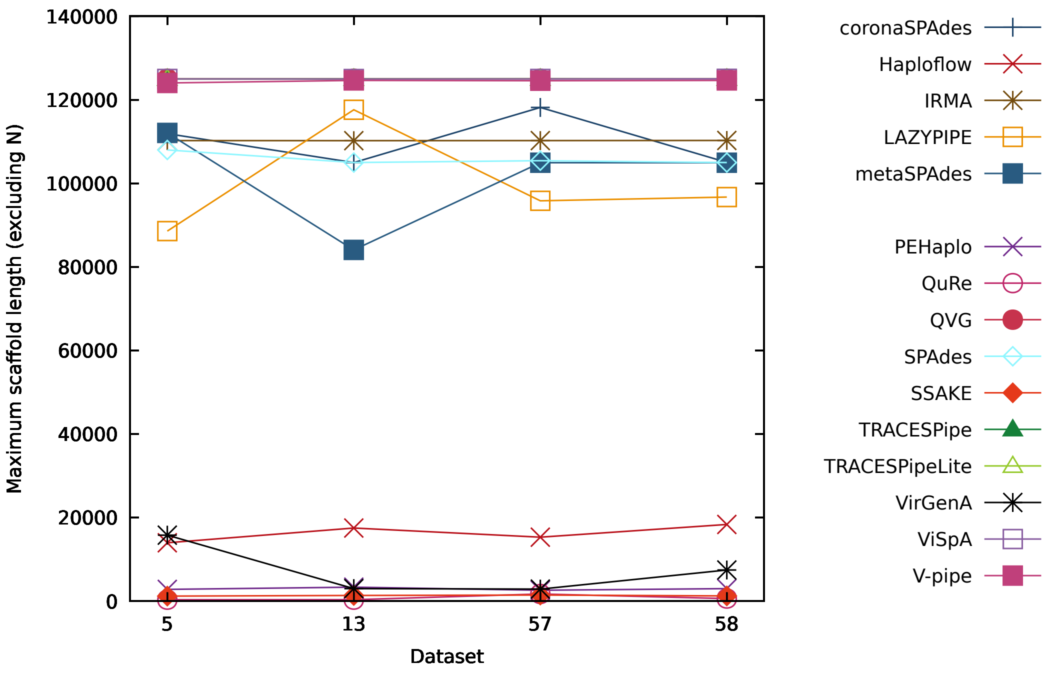


Figure S98: Figure comparing the performance of the reconstruction programs in terms of the maximum number of reconstructed bases per scaffold (excluding "N") for datasets 5, 13, 57 and 58.

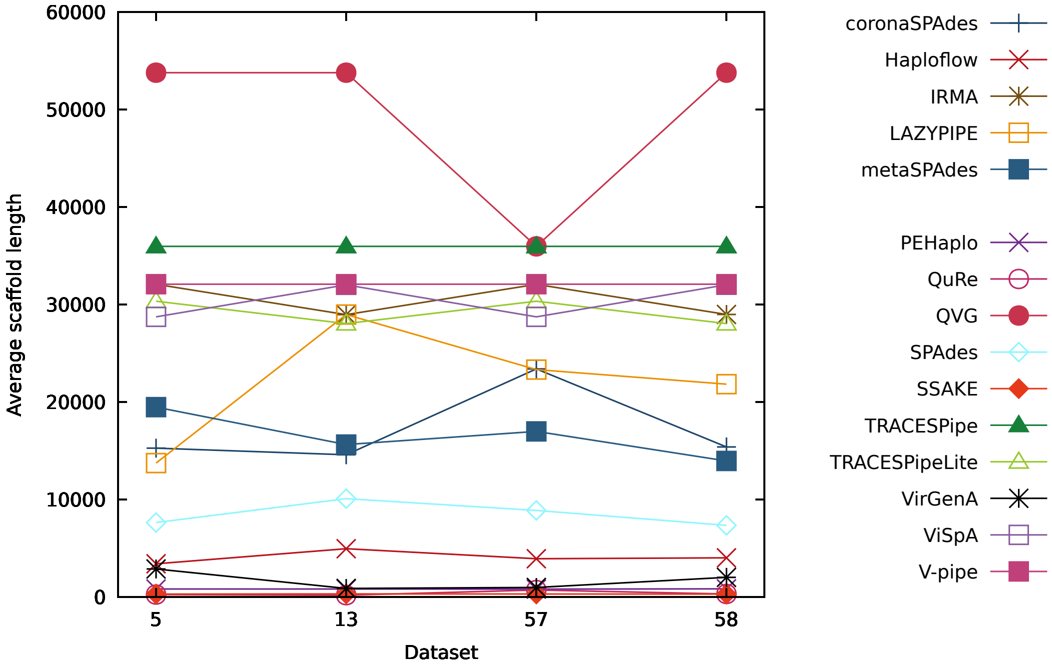


Figure S99: Figure comparing the performance of the reconstruction programs in terms of the average number of reconstructed bases per scaffold for datasets 5, 13, 57 and 58.

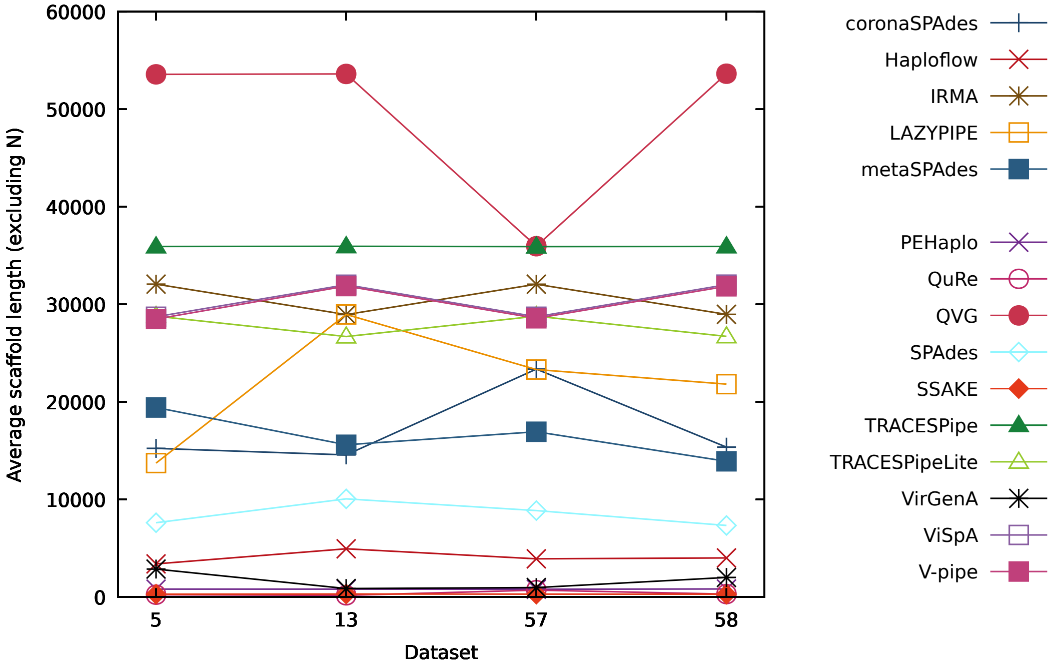


Figure S100: Figure comparing the performance of the reconstruction programs in terms of the average number of reconstructed bases per scaffold (excluding "N") for datasets 5, 13, 57 and 58.

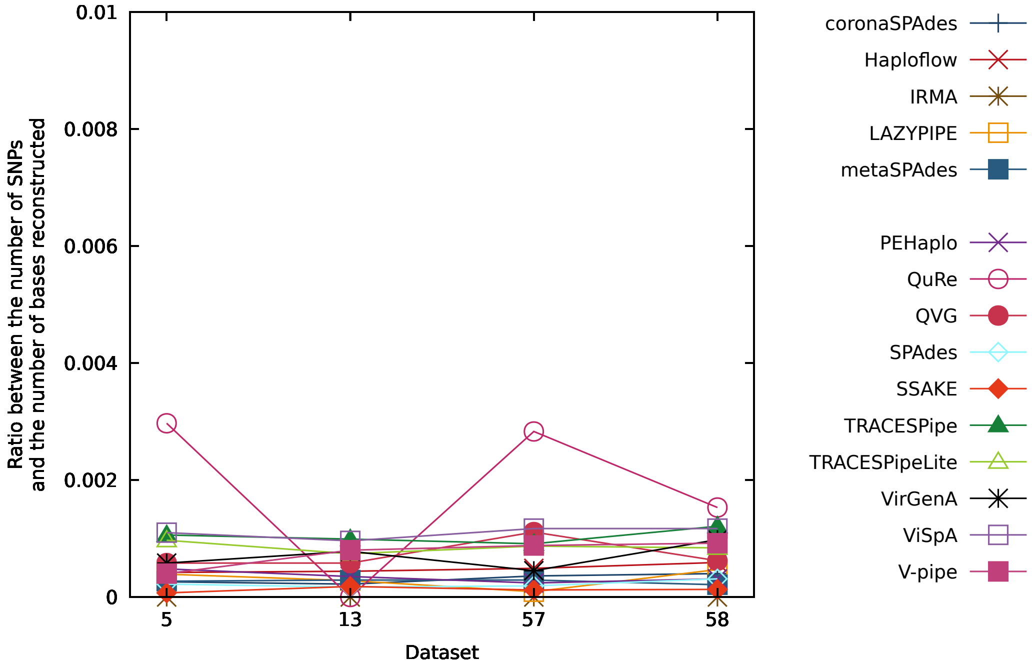


Figure S101: Figure comparing the performance of the reconstruction programs in terms of the ratio between the number of SNPs and the number of bases reconstructed for datasets 5, 13, 57 and 58.

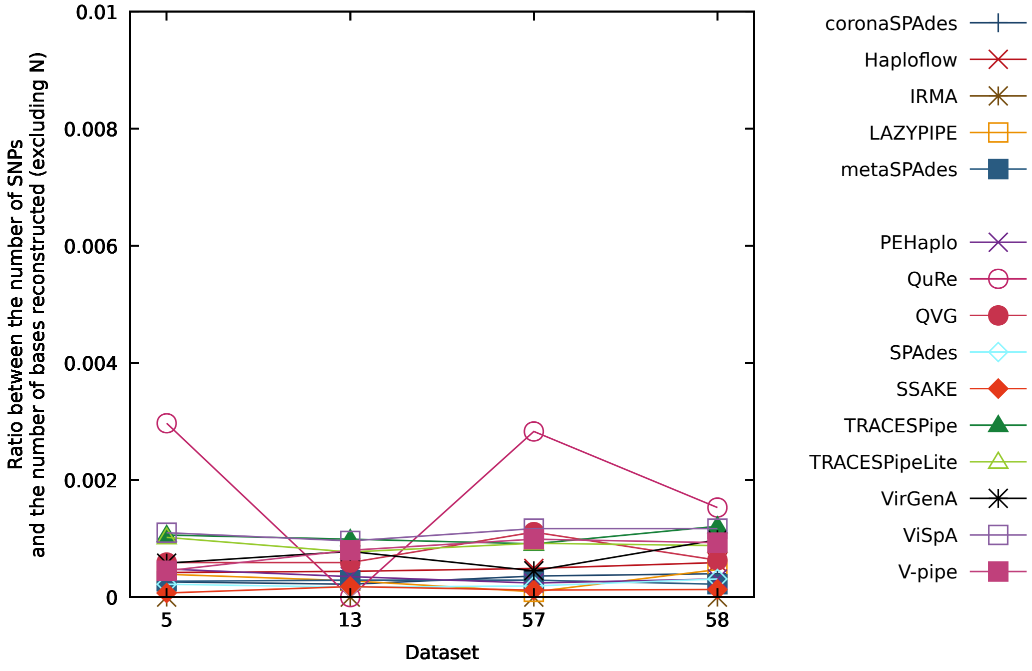


Figure S102: Figure comparing the performance of the reconstruction programs in terms of the ratio between the number of SNPs and the number of bases reconstructed (excluding "N") for datasets 5, 13, 57 and 58.

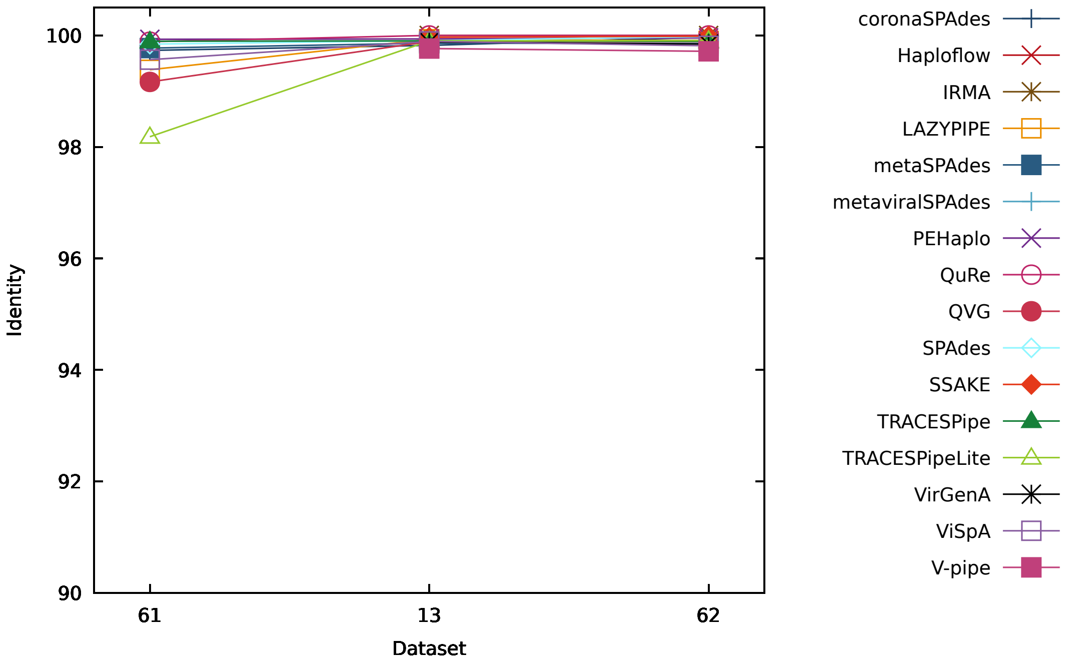


Figure S103: Figure comparing the performance of the reconstruction programs in terms of the identity for datasets 61, 13 and 62.

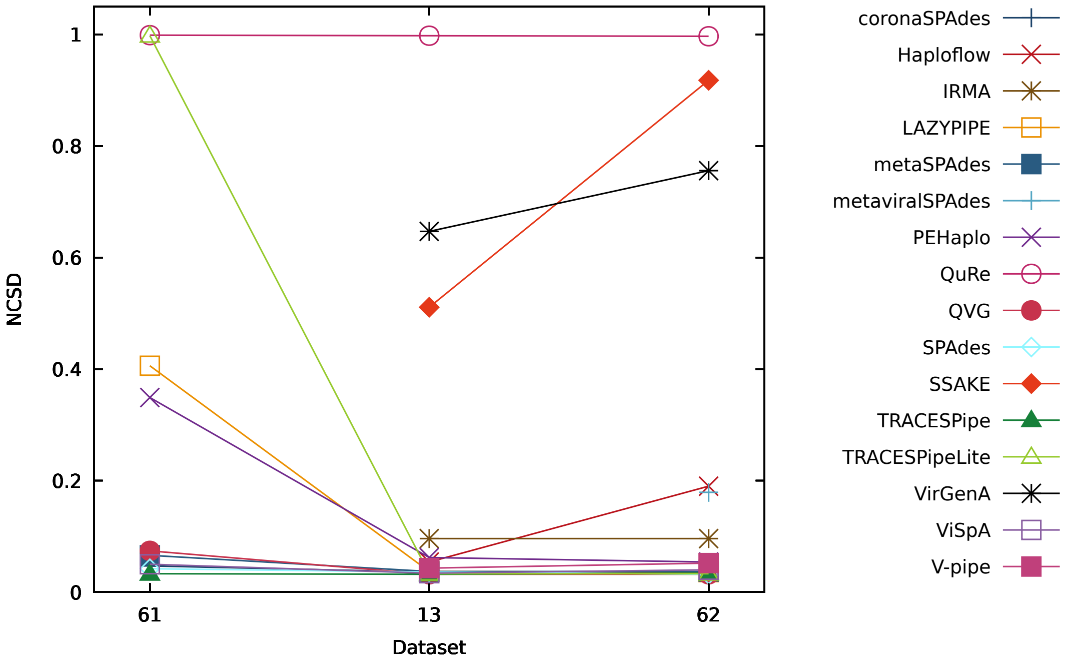


Figure S104: Figure comparing the performance of the reconstruction programs in terms of the NCSD for datasets 61, 13 and 62.

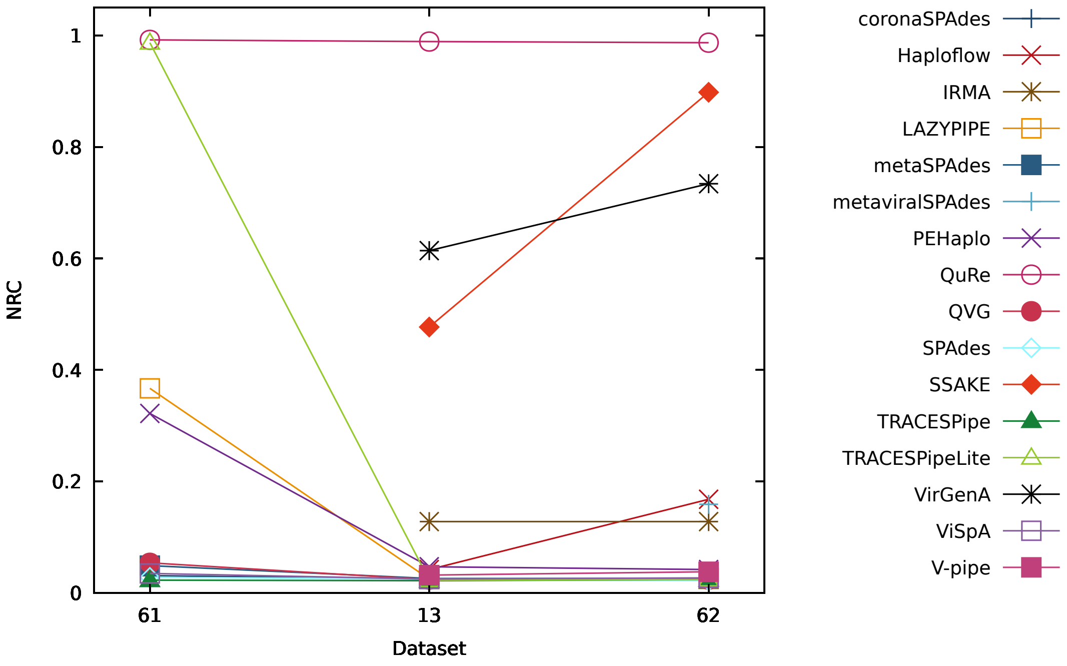


Figure S105: Figure comparing the performance of the reconstruction programs in terms of the NRC for datasets 61, 13 and 62.

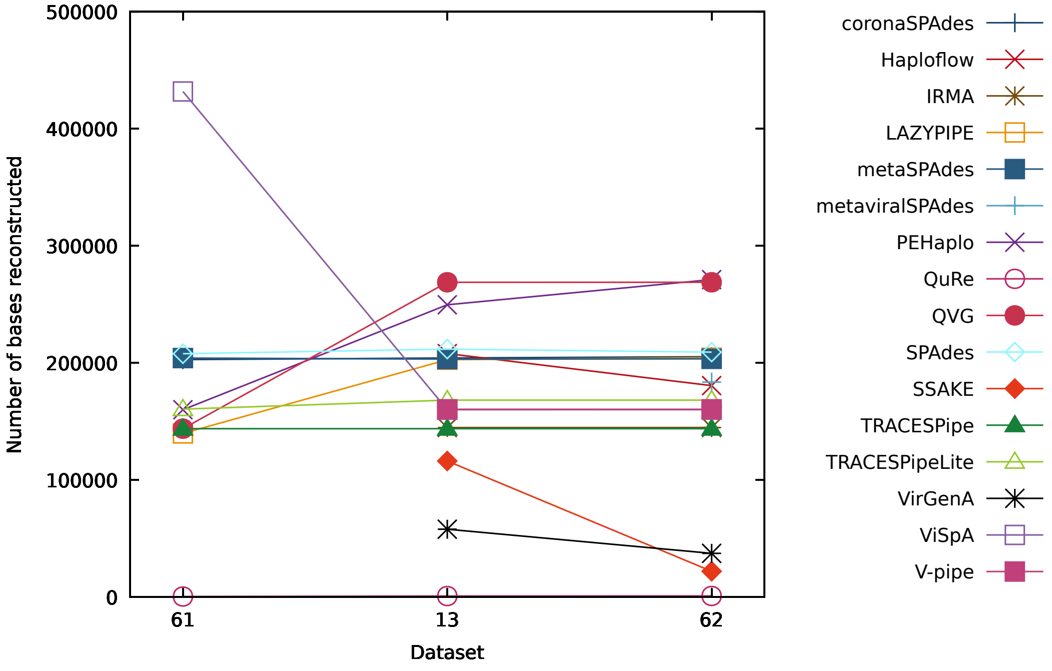


Figure S106: Figure comparing the performance of the reconstruction programs in terms of the number of bases reconstructed for datasets 61, 13 and 62.

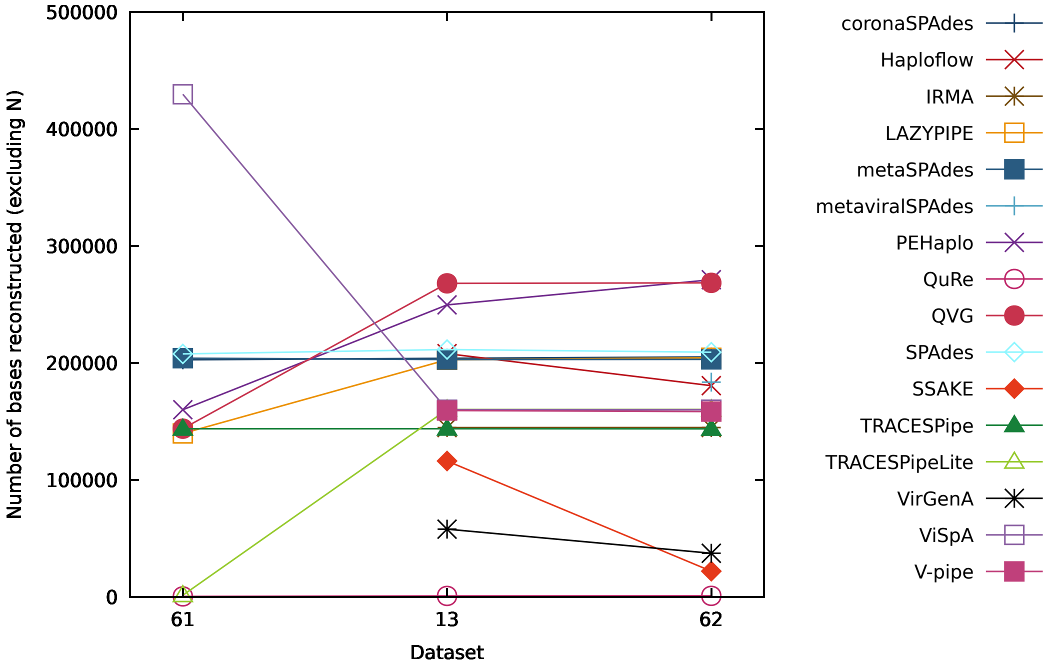


Figure S107: Figure comparing the performance of the reconstruction programs in terms of the number of bases reconstructed (excluding "N") for datasets 61, 13 and 62.

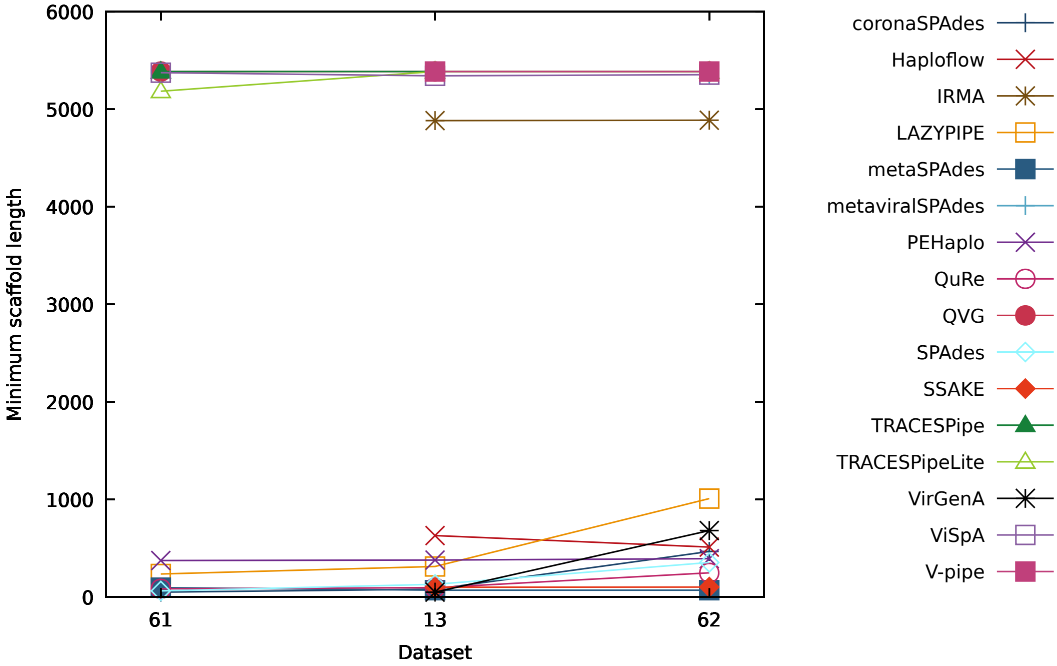


Figure S108: Figure comparing the performance of the reconstruction programs in terms of the minimum number of reconstructed bases per scaffold for datasets 61, 13 and 62.

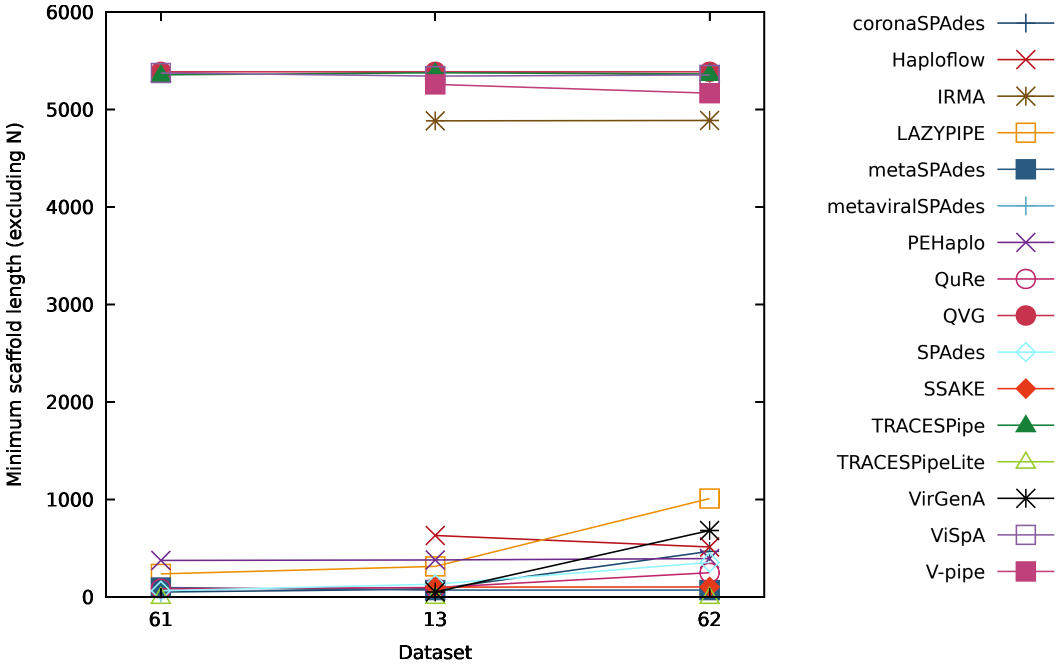


Figure S109: Figure comparing the performance of the reconstruction programs in terms of the minimum number of reconstructed bases per scaffold (excluding "N") for datasets 61, 13 and 62.

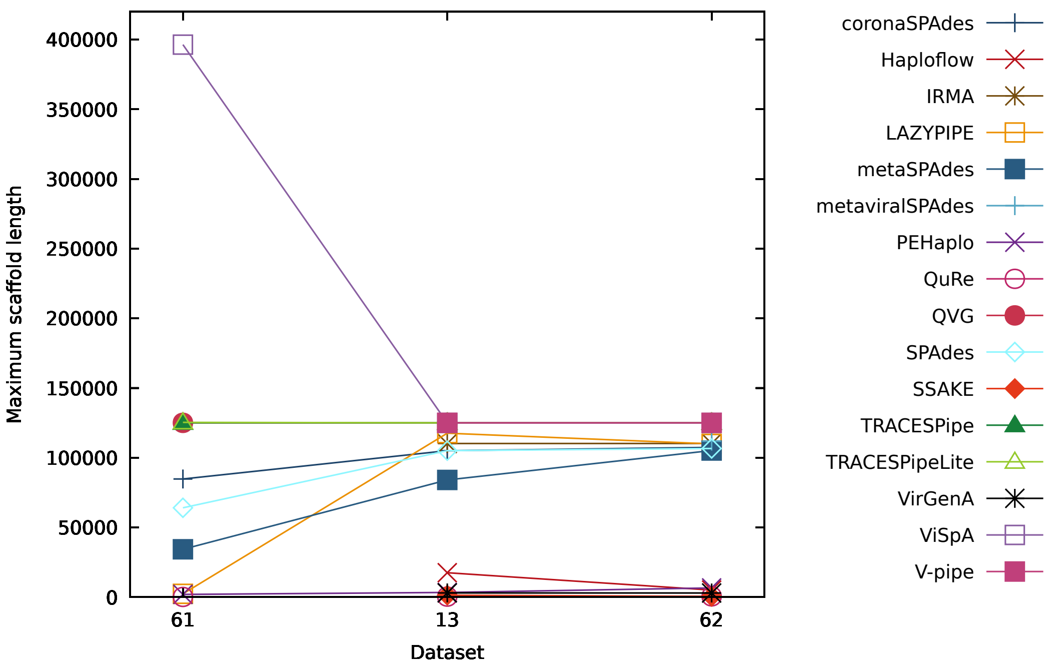


Figure S110: Figure comparing the performance of the reconstruction programs in terms of the maximum number of reconstructed bases per scaffold for datasets 61, 13 and 62.

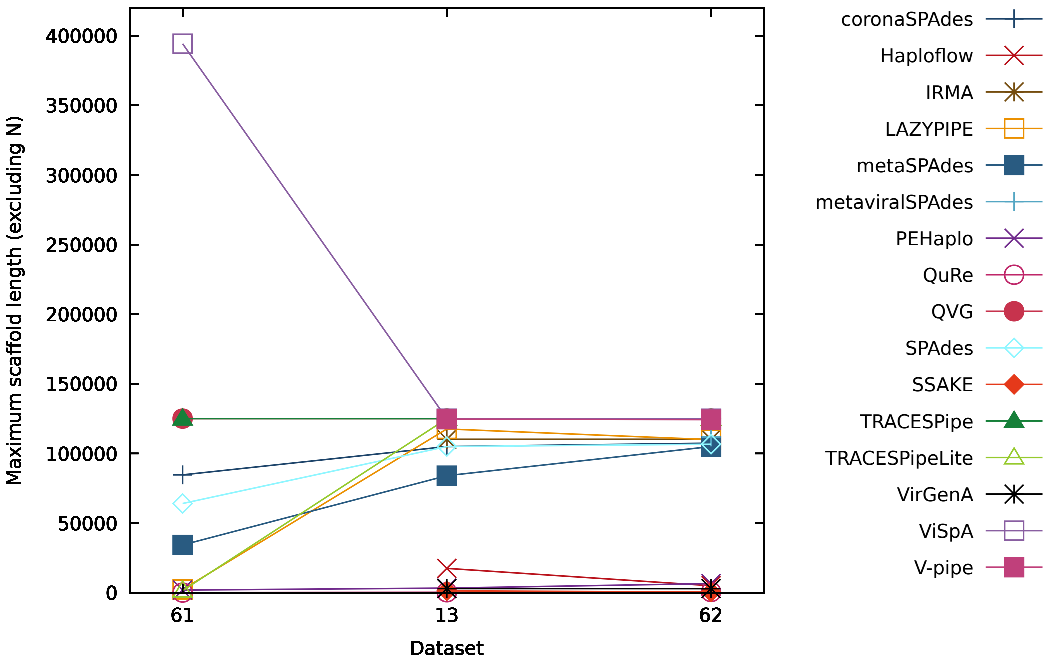


Figure S111: Figure comparing the performance of the reconstruction programs in terms of the maximum number of reconstructed bases per scaffold (excluding "N") for datasets 61, 13 and 62.

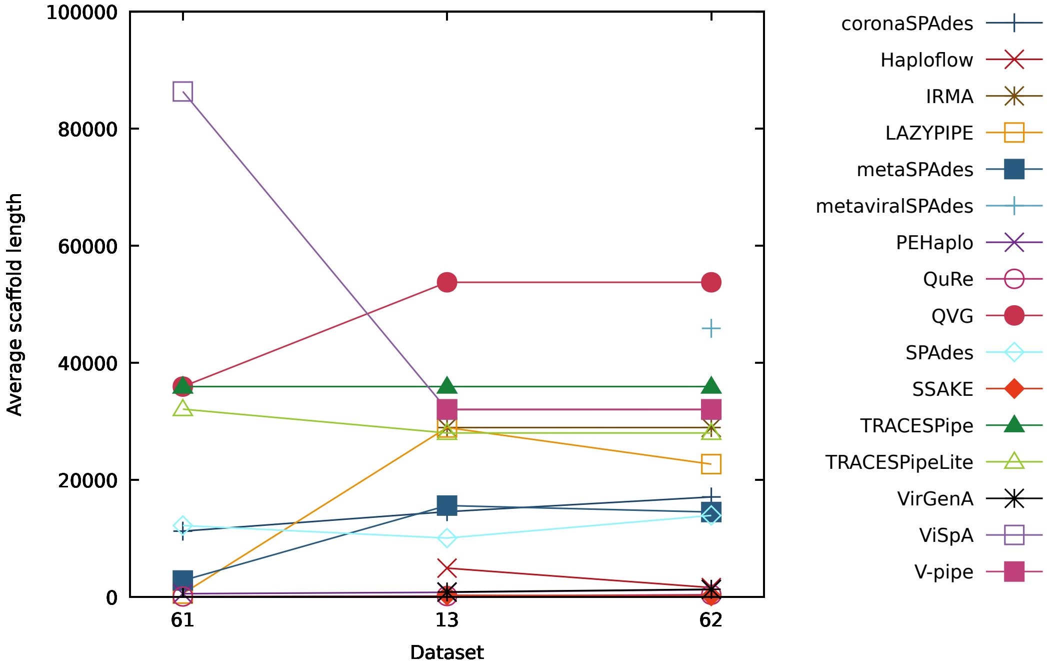


Figure S112: Figure comparing the performance of the reconstruction programs in terms of the average number of reconstructed bases per scaffold for datasets 61, 13 and 62.

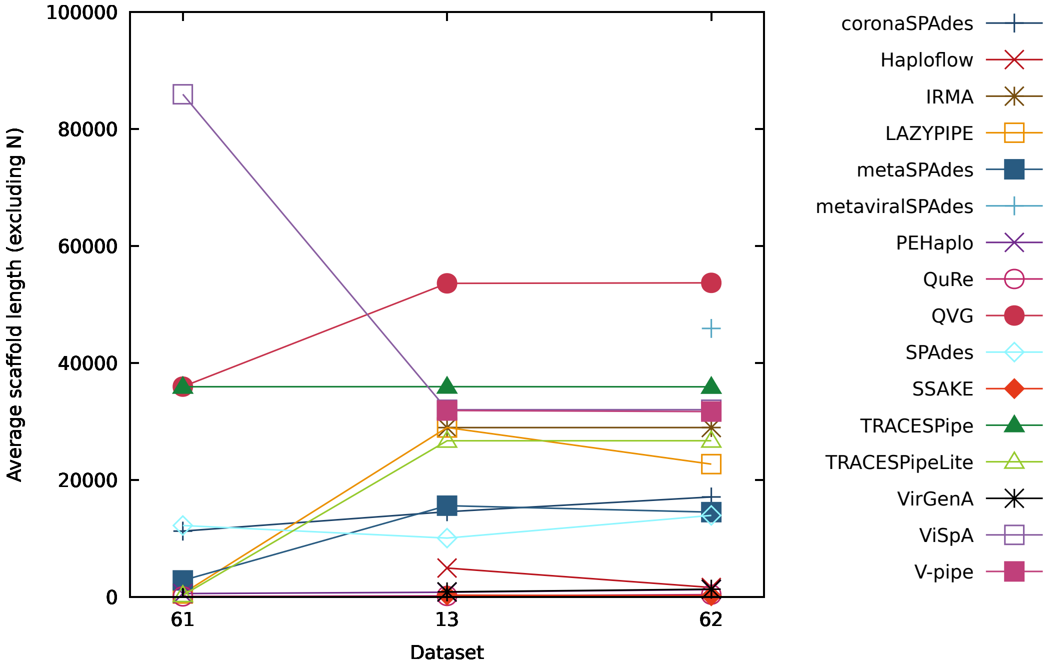


Figure S113: Figure comparing the performance of the reconstruction programs in terms of the average number of reconstructed bases per scaffold (excluding "N") for datasets 61, 13 and 62.

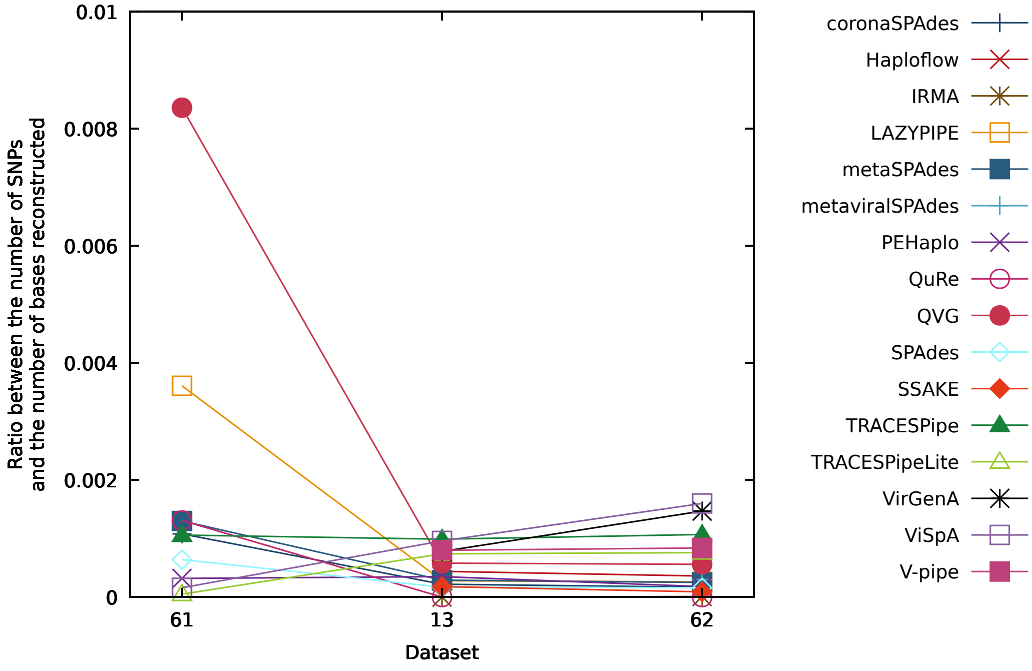


Figure S114: Figure comparing the performance of the reconstruction programs in terms of the ratio between the number of SNPs and the number of bases reconstructed for datasets 61, 13 and 62.

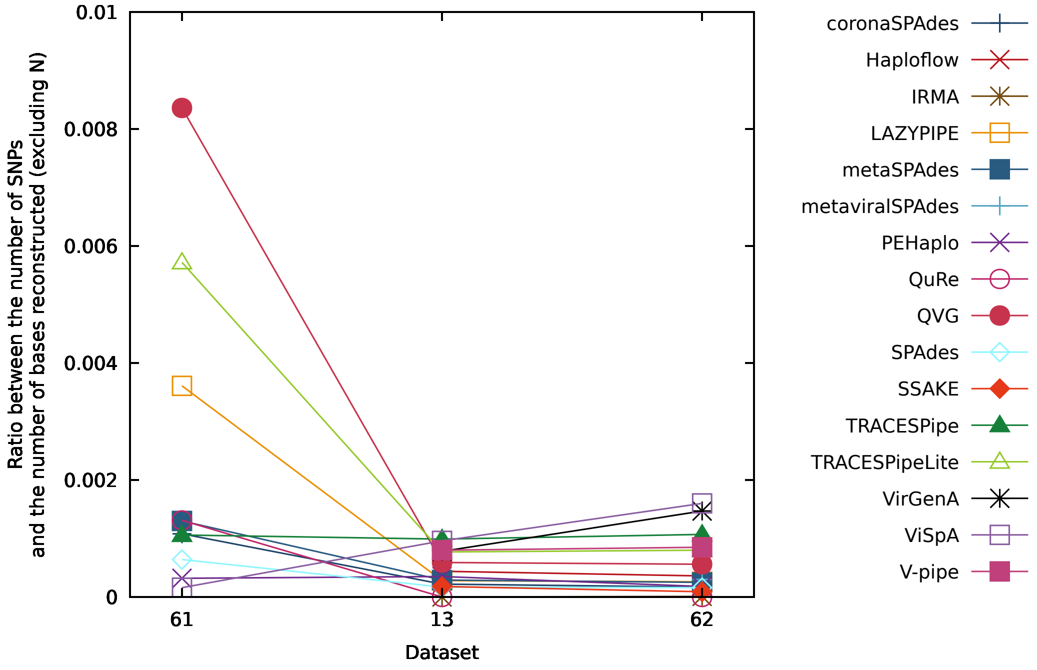


Figure S115: Figure comparing the performance of the reconstruction programs in terms of the ratio between the number of SNPs and the number of bases reconstructed (excluding "N") for datasets 61, 13 and 62.

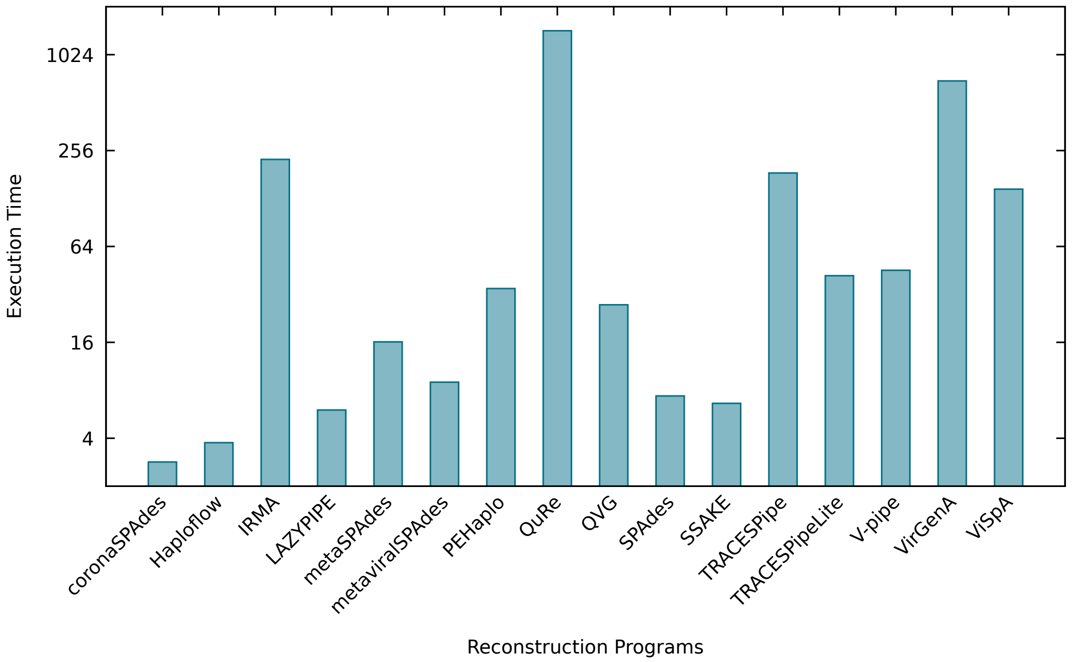


Figure S116: Figure comparing the average performance of the reconstruction programs in terms of the execution time across all datasets reconstructed. The y-axis is presented in a logarithmic scale of base 2.

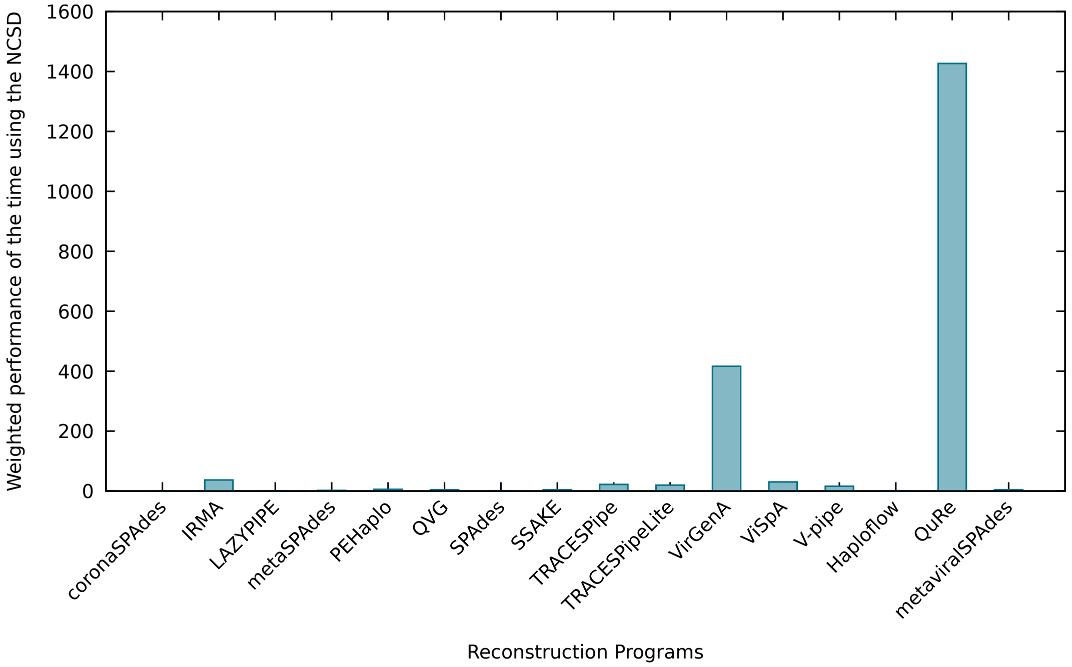


Figure S117: Figure comparing the average performance of the reconstruction programs in terms of the weighted execution time across all datasets reconstructed.

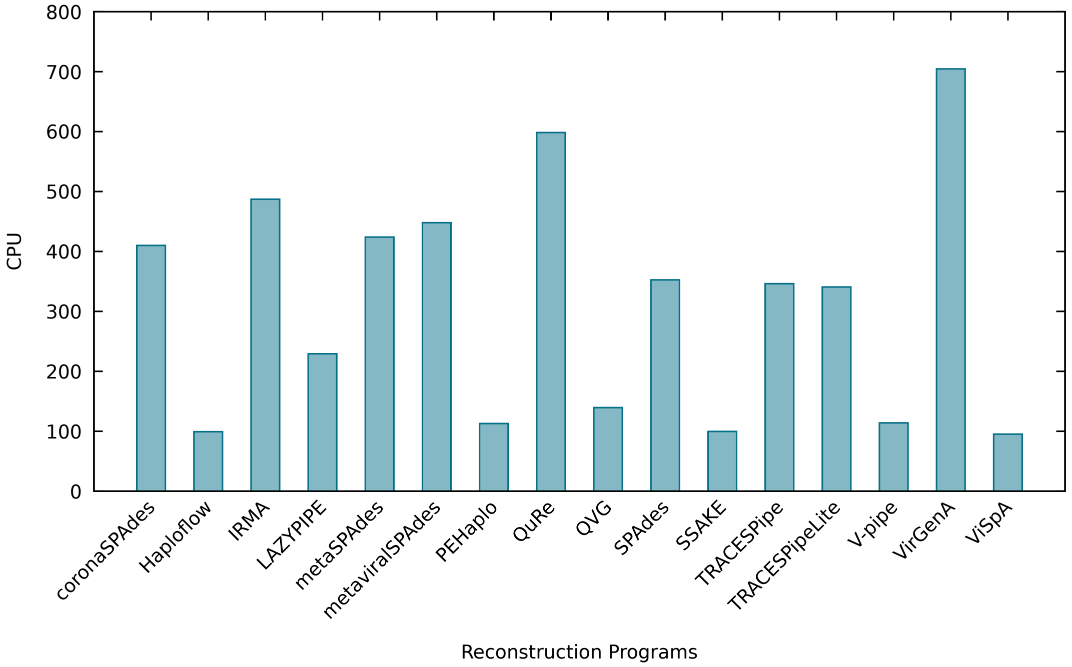


Figure S118: Figure comparing the average performance of the reconstruction programs in terms of the percentage of CPU used across all datasets reconstructed.

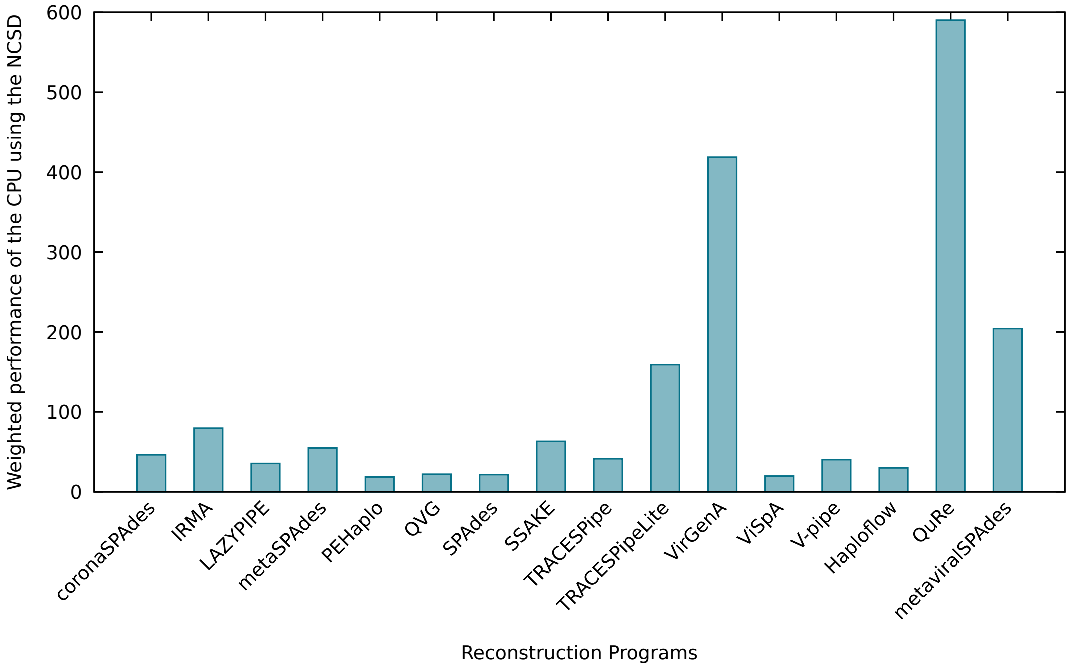


Figure S119: Figure comparing the average performance of the reconstruction programs in terms of the weighted percentage of CPU used across all datasets reconstructed.

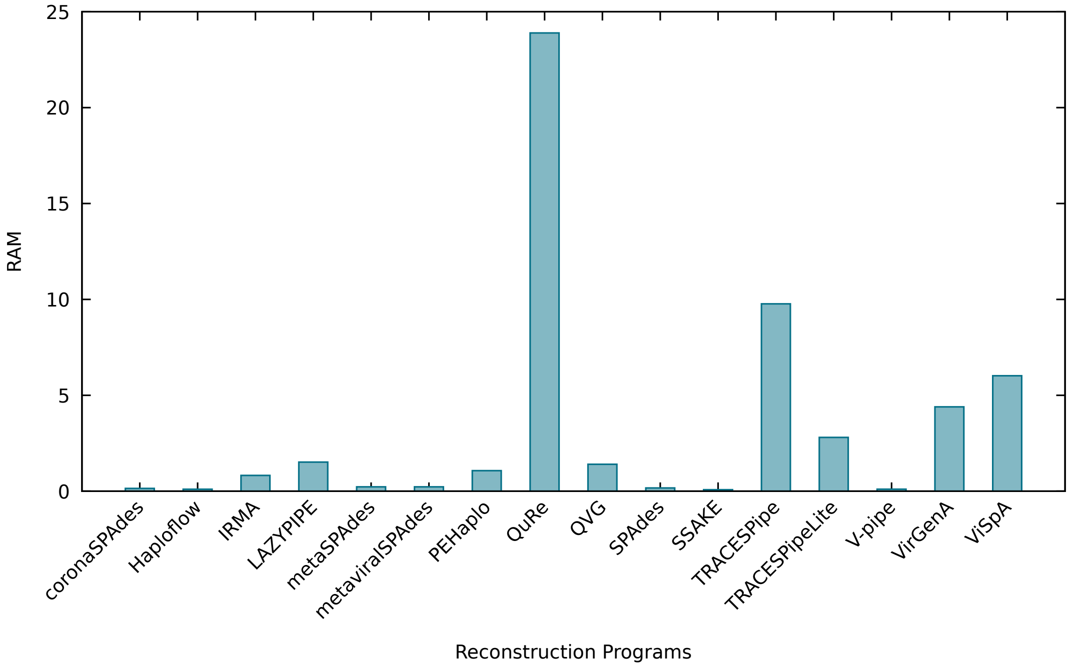


Figure S120: Figure comparing the average performance of the reconstruction programs in terms of the RAM used across all datasets reconstructed.

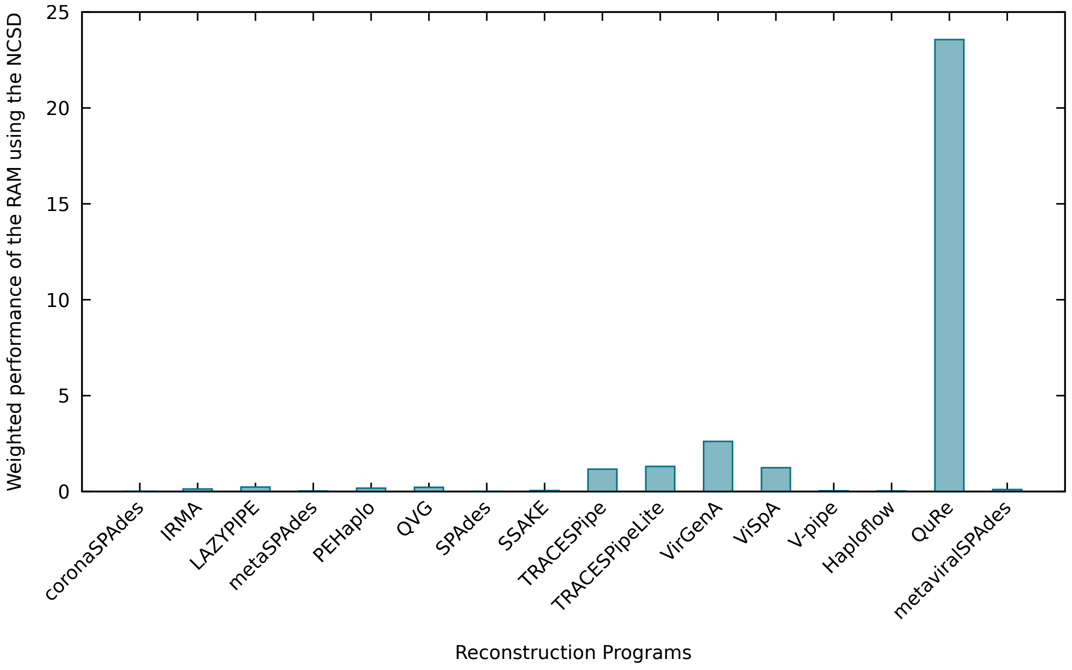


Figure S121: Figure comparing the average performance of the reconstruction programs in terms of the weighted RAM used across all datasets reconstructed.

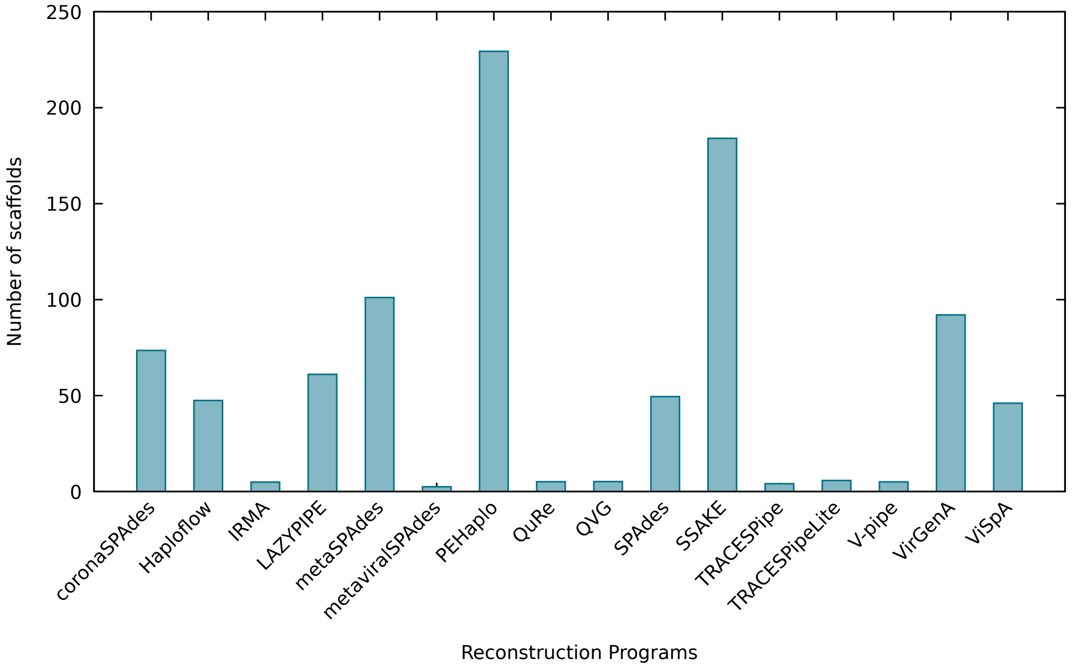


Figure S122: Figure comparing the average performance of the reconstruction programs in terms of the number of scaffolds reconstructed across all datasets reconstructed.

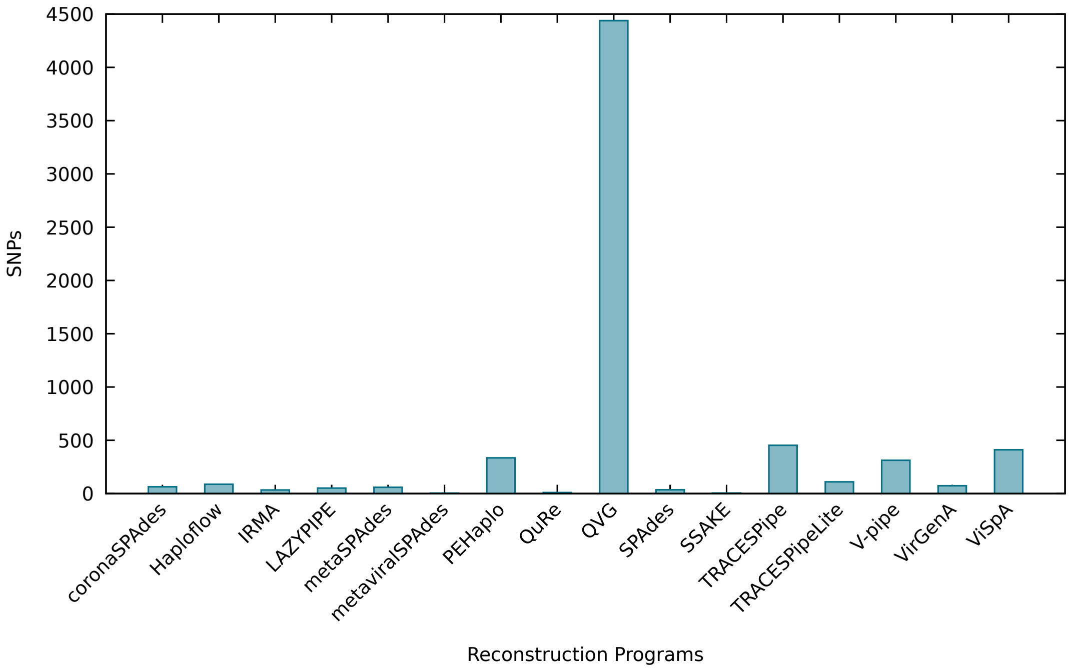


Figure S123: Figure comparing the average performance of the reconstruction programs in terms of the number of SNPs across all datasets reconstructed.

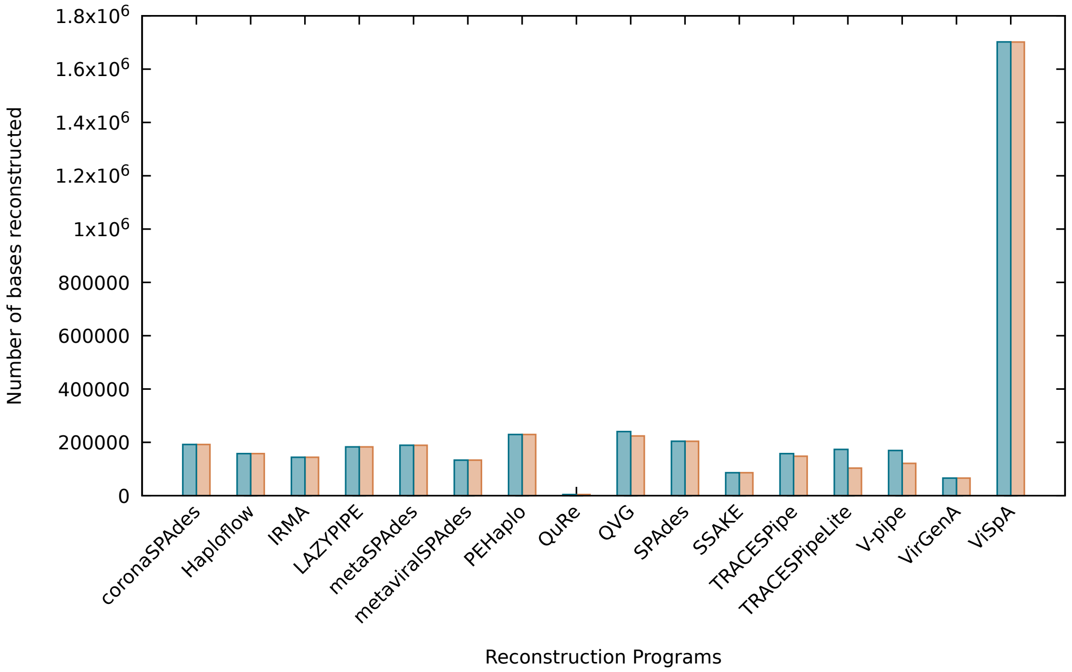


Figure S124: Figure comparing the average performance of the reconstruction programs in terms of the overall number of bases reconstructed across all datasets reconstructed. The blue bars show the performance using all bases present in the reconstructed file, whereas the orange bars show the results excluding non-reconstructed bases (N).

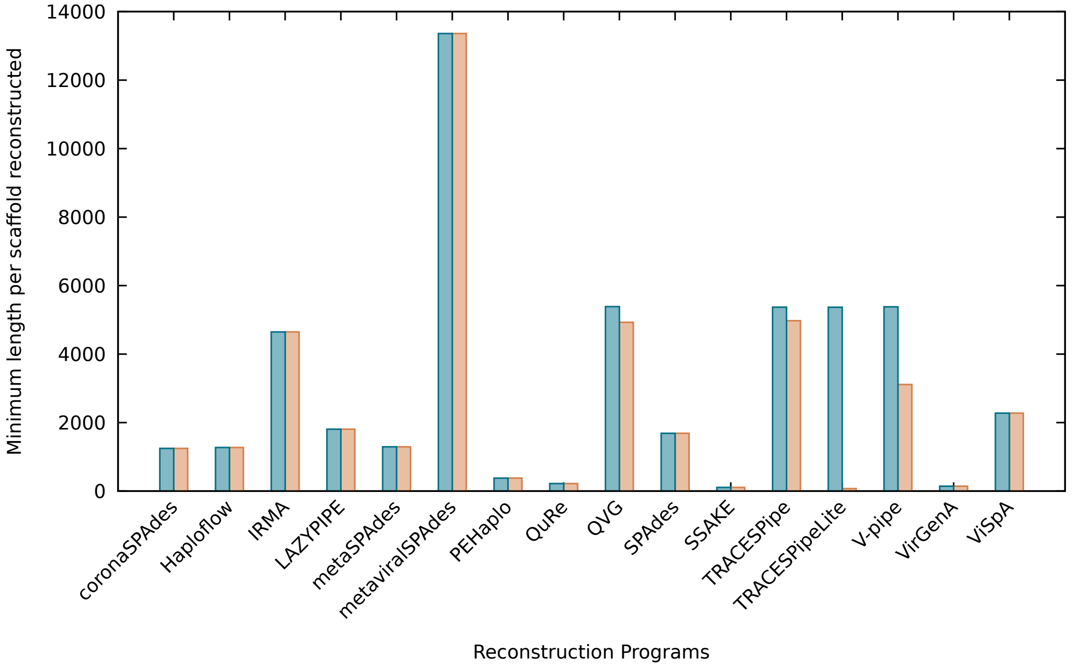


Figure S125: Figure comparing the average performance of the reconstruction programs in terms of the minimum number of bases reconstructed per scaffold across all datasets reconstructed. The blue bars show the performance using all bases present in the reconstructed file, whereas the orange bars show the results excluding non-reconstructed bases (N).

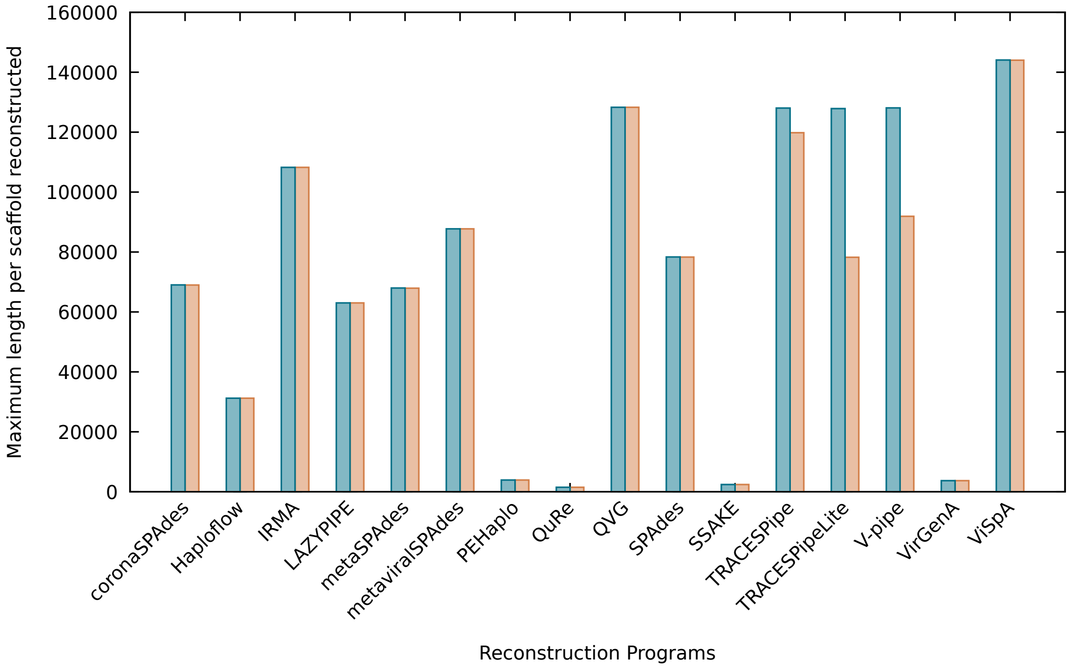


Figure S126: Figure comparing the average performance of the reconstruction programs in terms of the maximum number of bases reconstructed by scaffold across all datasets reconstructed. The blue bars show the performance using all bases present in the reconstructed file, whereas the orange bars show the results excluding non-reconstructed bases (N).

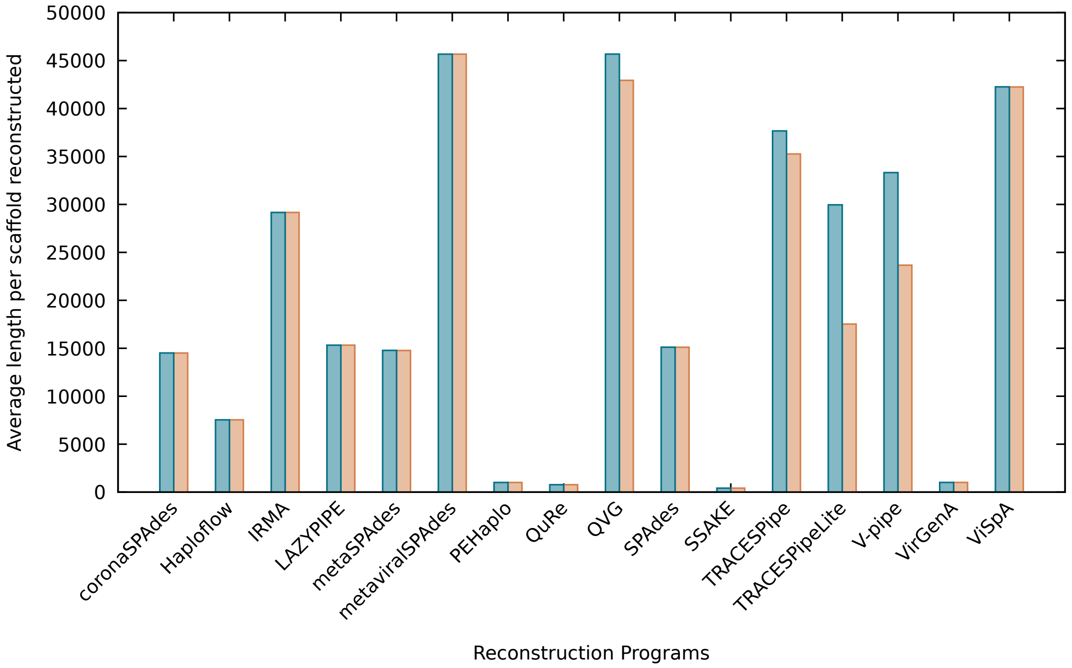


Figure S127: Figure comparing the average performance of the reconstruction programs in terms of the average number of bases reconstructed by scaffold across all datasets reconstructed. The blue bars show the performance using all bases present in the reconstructed file, whereas the orange bars show the results excluding non-reconstructed bases (N).

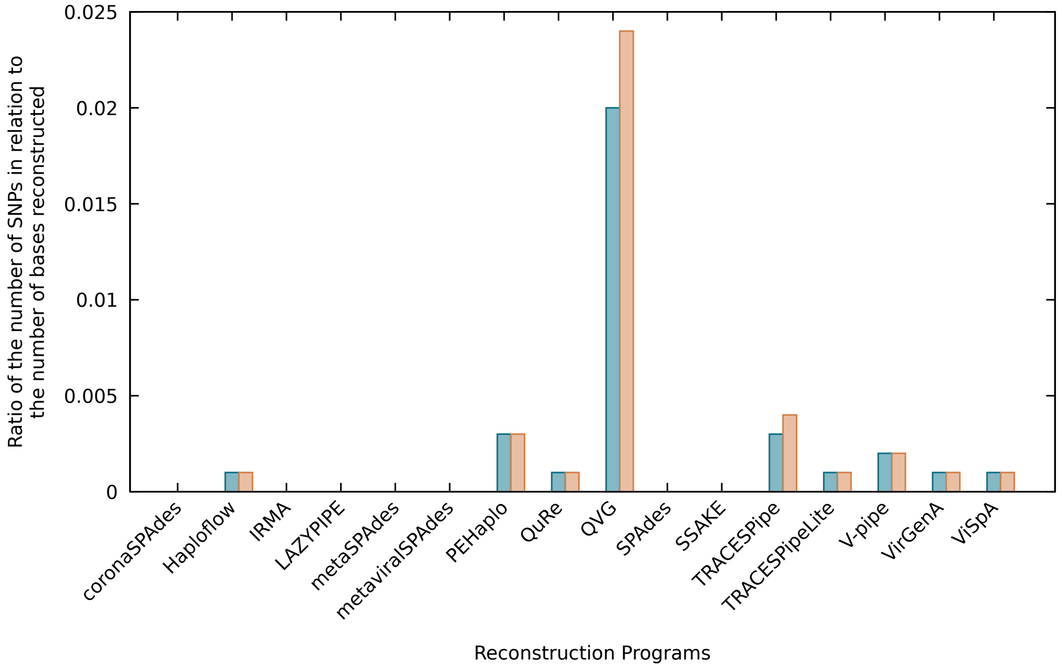


Figure S128: Figure comparing the average performance of the reconstruction programs in terms of the number of SNPs in relation to the number of bases reconstructed across all datasets reconstructed. The blue bars show the performance using all bases present in the reconstructed file, whereas the orange bars show the results excluding non-reconstructed bases (N).

Table S3: Results obtained for DS1 using the benchmark proposed. The execution time was measured in seconds, the RAM usage was measured in GB and the CPU usage is presented as a percentage. The executions were, when possible, capped at 6 threads and 48 GB of RAM.

| Reconstruction tool | Execution time | SNPs | Identity | NCSD | NRC | RAM usage | CPU usage | Number of scaffolds | Reconstructed bases | Minimum scaffold length | Maximum scaffold length | Average scaffold length |
|---|---|---|---|---|---|---|---|---|---|---|---|---|
| coronaSPAdes | 1.12 | 131.0 | 99.824 | 0.462 | 0.42 | 0.133 | 313.667 | 182.0 | 83087.0 | 100.0 | 1489.0 | 456.5 |
| Haploflow | – | – | – | – | – | – | – | – | – | – | – | – |
| IRMA | 35.28 | 93.667 | 96.09 | 0.38 | 0.383 | 0.128 | 317.0 | 4.0 | 95730.0 | 3682.0 | 82389.0 | 23932.5 |
| LAZYPIPE | 5.455 | 64.0 | 99.881 | 0.579 | 0.537 | 1.494 | 108.667 | 152.0 | 65307.0 | 215.0 | 1352.0 | 429.7 |
| metaSPAdes | 1.865 | 128.0 | 99.854 | 0.423 | 0.382 | 0.225 | 295.333 | 210.0 | 88863.0 | 71.0 | 1489.0 | 423.2 |
| metaviralSPAdes | – | – | – | – | – | – | – | – | – | – | – | – |
| PEHaplo | 11.15 | 793.0 | 99.749 | 0.606 | 0.568 | 0.091 | 46.0 | 104.0 | 61220.0 | 375.0 | 1961.0 | 588.7 |
| QuRe | – | – | – | – | – | – | – | – | – | – | – | – |
| QVG | 18.815 | 1301.0 | 99.064 | 0.115 | 0.092 | 1.387 | 82.333 | 3.0 | 138473.0 | 5596.0 | 125041.0 | 46157.7 |
| SPAdes | 2.315 | 104.0 | 99.858 | 0.466 | 0.425 | 0.141 | 240.667 | 179.0 | 82068.0 | 104.0 | 1489.0 | 458.5 |
| SSAKE | 2.445 | 0.0 | 100.0 | 1.0 | 0.994 | 0.041 | 84.0 | 2.0 | 241.0 | 113.0 | 128.0 | 120.5 |
| TRACESPipe | 180.42 | 100.0 | 97.349 | 0.326 | 0.292 | 9.768 | 319.333 | 4.0 | 143860.0 | 5387.0 | 125041.0 | 35965.0 |
| TRACESPipeLite | 35.58 | 66.0 | 97.028 | 0.323 | 0.288 | 2.707 | 364.0 | 5.0 | 151693.0 | 5387.0 | 125041.0 | 30338.6 |
| VirGenA | 200.22 | 88.0 | 99.884 | 0.465 | 0.43 | 3.857 | 567.667 | 238.0 | 86031.0 | 53.0 | 1293.0 | 361.5 |
| ViSpA | 43.835 | 94.0 | 95.123 | 0.523 | 0.502 | 4.913 | 51.0 | 75.0 | 2292352.0 | 0.0 | 70025.0 | 30564.7 |
| V-pipe | 32.89 | 0.0 | 95.791 | 0.976 | 0.963 | 0.104 | 106.667 | 5.0 | 160429.0 | 5387.0 | 125041.0 | 32085.8 |

Table S4: Results obtained for DS2 using the benchmark proposed. The execution time was measured in seconds, the RAM usage was measured in GB and the CPU usage is presented as a percentage. The executions were, when possible, capped at 6 threads and 48 GB of RAM.

| Reconstruction tool | Execution time | SNPs | Identity | NCSD | NRC | RAM usage | CPU usage | Number of scaffolds | Reconstructed bases | Minimum scaffold length | Maximum scaffold length | Average scaffold length |
|---|---|---|---|---|---|---|---|---|---|---|---|---|
| coronaSPAdes | 1.37 | 120.0 | 99.775 | 0.115 | 0.091 | 0.132 | 370.667 | 100.0 | 129540.0 | 227.0 | 5926.0 | 1295.4 |
| Haploflow | – | – | – | – | – | – | – | – | – | – | – | – |
| IRMA | 76.93 | 48.333 | 98.395 | 0.161 | 0.182 | 0.273 | 371.667 | 4.0 | 122405.0 | 4751.0 | 104886.0 | 30601.3 |
| LAZYPIPE | 4.255 | 114.0 | 99.896 | 0.17 | 0.142 | 1.499 | 184.667 | 92.0 | 121351.0 | 305.0 | 10147.0 | 1319.0 |
| metaSPAdes | 2.335 | 156.0 | 99.721 | 0.109 | 0.085 | 0.225 | 341.0 | 104.0 | 130243.0 | 97.0 | 7357.0 | 1252.3 |
| metaviralSPAdes | – | – | – | – | – | – | – | – | – | – | – | – |
| PEHaplo | 10.64 | 114.0 | 99.877 | 0.137 | 0.112 | 0.114 | 68.667 | 102.0 | 127512.0 | 376.0 | 4405.0 | 1250.1 |
| QuRe | – | – | – | – | – | – | – | – | – | – | – | – |
| QVG | 15.445 | 955.0 | 99.333 | 0.067 | 0.047 | 1.393 | 100.0 | 4.0 | 143860.0 | 5387.0 | 125041.0 | 35965.0 |
| SPAdes | 3.345 | 109.0 | 99.909 | 0.113 | 0.089 | 0.148 | 317.667 | 101.0 | 129841.0 | 119.0 | 10193.0 | 1285.6 |
| SSAKE | 4.09 | 2.333 | 99.972 | 0.961 | 0.945 | 0.054 | 100.0 | 51.0 | 7772.333 | 101.333 | 250.667 | 152.2 |
| TRACESPipe | 177.915 | 135.0 | 99.06 | 0.084 | 0.066 | 9.768 | 341.0 | 4.0 | 143860.0 | 5387.0 | 125041.0 | 35965.0 |
| TRACESPipeLite | 34.735 | 153.0 | 98.787 | 0.097 | 0.077 | 2.749 | 370.0 | 5.0 | 151693.0 | 5387.0 | 125041.0 | 30338.6 |
| VirGenA | 309.26 | 94.0 | 99.878 | 0.289 | 0.263 | 3.6 | 643.667 | 111.333 | 110418.667 | 98.333 | 3544.667 | 993.667 |
| ViSpA | 69.81 | 137.0 | 97.596 | 0.156 | 0.138 | 5.02 | 63.333 | 80.0 | 3612120.0 | 0.0 | 112477.0 | 45151.5 |
| V-pipe | 32.22 | 24.0 | 94.134 | 0.649 | 0.612 | 0.104 | 115.333 | 5.0 | 160429.0 | 5387.0 | 125041.0 | 32085.8 |

Table S5: Results obtained for DS3 using the benchmark proposed. The execution time was measured in seconds, the RAM usage was measured in GB and the CPU usage is presented as a percentage. The executions were, when possible, capped at 6 threads and 48 GB of RAM.

| Reconstruction tool | Execution time | SNPs | Identity | NCSD | NRC | RAM usage | CPU usage | Number of scaffolds | Reconstructed bases | Minimum scaffold length | Maximum scaffold length | Average scaffold length |
|---|---|---|---|---|---|---|---|---|---|---|---|---|
| coronaSPAdes | 1.74 | 39.0 | 99.868 | 0.04 | 0.029 | 0.138 | 388.667 | 21.0 | 138037.0 | 81.0 | 29589.0 | 6573.2 |
| Haploflow | 1.41 | 34.0 | 99.882 | 0.669 | 0.635 | 0.05 | 99.667 | 65.0 | 52460.0 | 508.0 | 3184.0 | 807.1 |
| IRMA | 87.815 | 5.0 | 99.921 | 0.1 | 0.132 | 0.483 | 426.667 | 4.0 | 127894.0 | 4769.0 | 110094.0 | 31973.5 |
| LAZYPIPE | 4.905 | 60.0 | 99.95 | 0.048 | 0.036 | 1.505 | 220.0 | 19.0 | 135584.0 | 1052.0 | 18805.0 | 7136.0 |
| metaSPAdes | 3.02 | 60.0 | 99.813 | 0.042 | 0.031 | 0.226 | 367.333 | 17.0 | 136629.0 | 135.0 | 29589.0 | 8037.0 |
| metaviralSPAdes | – | – | – | – | – | – | – | – | – | – | – | – |
| PEHaplo | 14.385 | 55.0 | 99.931 | 0.064 | 0.049 | 0.322 | 96.0 | 116.0 | 146720.0 | 393.0 | 5751.0 | 1264.8 |
| QuRe | 129.93 | 0.0 | 100.0 | 1.0 | 0.994 | 12.4 | 622.333 | 3.0 | 287.0 | 95.0 | 96.0 | 95.7 |
| QVG | 17.295 | 452.0 | 99.685 | 0.051 | 0.033 | 1.397 | 114.667 | 4.0 | 143860.0 | 5387.0 | 125041.0 | 35965.0 |
| SPAdes | 11.17 | 37.0 | 99.926 | 0.04 | 0.029 | 0.151 | 194.333 | 27.0 | 143682.0 | 167.0 | 29589.0 | 5321.6 |
| SSAKE | 1.46 | 3.333 | 99.984 | 0.885 | 0.859 | 0.043 | 100.0 | 116.0 | 20861.333 | 100.333 | 418.0 | 179.767 |
| TRACESPipe | 179.45 | 152.0 | 99.85 | 0.038 | 0.028 | 9.768 | 341.333 | 4.0 | 143860.0 | 5387.0 | 125041.0 | 35965.0 |
| TRACESPipeLite | 36.65 | 132.333 | 99.857 | 0.036 | 0.026 | 2.759 | 357.333 | 5.0 | 151693.0 | 5387.0 | 125041.0 | 30338.6 |
| VirGenA | 448.105 | 66.0 | 99.864 | 0.252 | 0.23 | 3.879 | 696.0 | 64.0 | 114483.0 | 49.0 | 6344.0 | 1788.8 |
| ViSpA | 44.995 | 132.0 | 99.514 | 0.046 | 0.037 | 5.215 | 73.333 | 9.0 | 483660.0 | 0.0 | 124259.0 | 53740.0 |
| V-pipe | 34.765 | 46.0 | 97.085 | 0.251 | 0.22 | 0.104 | 116.667 | 5.0 | 160429.0 | 5387.0 | 125041.0 | 32085.8 |

Table S6: Results obtained for DS4 using the benchmark proposed. The execution time was measured in seconds, the RAM usage was measured in GB and the CPU usage is presented as a percentage. The executions were, when possible, capped at 6 threads and 48 GB of RAM.

| Reconstruction tool | Execution time | SNPs | Identity | NCSD | NRC | RAM usage | CPU usage | Number of scaffolds | Reconstructed bases | Minimum scaffold length | Maximum scaffold length | Average scaffold length |
|---|---|---|---|---|---|---|---|---|---|---|---|---|
| coronaSPAdes | 2.23 | 74.0 | 99.837 | 0.036 | 0.026 | 0.139 | 390.0 | 14.0 | 138101.0 | 81.0 | 82423.0 | 9864.4 |
| Haploflow | 1.9 | 68.0 | 99.908 | 0.173 | 0.147 | 0.059 | 100.0 | 82.0 | 126008.0 | 510.0 | 5576.0 | 1536.7 |
| IRMA | 121.98 | 1.0 | 99.992 | 0.096 | 0.128 | 0.683 | 470.333 | 4.0 | 128279.0 | 4882.0 | 110245.0 | 32069.8 |
| LAZYPIPE | 4.965 | 72.0 | 99.889 | 0.038 | 0.027 | 1.508 | 225.0 | 8.0 | 136217.0 | 5330.0 | 82405.0 | 17027.1 |
| metaSPAdes | 3.745 | 73.0 | 99.899 | 0.038 | 0.027 | 0.226 | 388.0 | 8.0 | 136413.0 | 71.0 | 77567.0 | 17051.6 |
| metaviralSPAdes | 4.145 | 1.0 | 99.94 | 0.97 | 0.961 | 0.226 | 422.333 | 1.0 | 5358.0 | 5358.0 | 5358.0 | 5358.0 |
| PEHaplo | 19.74 | 86.0 | 99.935 | 0.068 | 0.053 | 0.558 | 123.0 | 180.0 | 165861.0 | 379.0 | 3898.0 | 921.5 |
| QuRe | 225.47 | 0.0 | 100.0 | 0.999 | 0.992 | 14.053 | 561.0 | 6.333 | 658.333 | 86.0 | 136.0 | 104.067 |
| QVG | 20.035 | 197.0 | 99.859 | 0.035 | 0.026 | 1.399 | 127.667 | 5.0 | 268901.0 | 5387.0 | 125041.0 | 53780.2 |
| SPAdes | 7.145 | 68.0 | 99.791 | 0.037 | 0.026 | 0.154 | 269.667 | 7.0 | 137370.0 | 131.0 | 90831.0 | 19624.3 |
| SSAKE | 2.375 | 5.667 | 99.986 | 0.755 | 0.724 | 0.051 | 100.0 | 183.333 | 41342.0 | 102.667 | 587.0 | 225.3 |
| TRACESPipe | 185.445 | 111.0 | 99.92 | 0.034 | 0.024 | 9.768 | 336.667 | 4.0 | 143860.0 | 5387.0 | 125041.0 | 35965.0 |
| TRACESPipeLite | 38.555 | 124.667 | 99.913 | 0.033 | 0.023 | 2.763 | 350.333 | 5.0 | 151693.0 | 5387.0 | 125041.0 | 30338.6 |
| VirGenA | 506.295 | 98.0 | 99.831 | 0.25 | 0.224 | 3.962 | 734.333 | 50.0 | 114803.0 | 132.0 | 8622.0 | 2296.1 |
| ViSpA | 86.255 | 162.0 | 99.782 | 0.036 | 0.026 | 4.907 | 85.667 | 5.0 | 143528.0 | 0.0 | 124918.0 | 28705.6 |
| V-pipe | 38.185 | 151.0 | 99.49 | 0.057 | 0.044 | 0.104 | 115.667 | 5.0 | 160429.0 | 5387.0 | 125041.0 | 32085.8 |

Table S7: Results obtained for DS5 using the benchmark proposed. The execution time was measured in seconds, the RAM usage was measured in GB and the CPU usage is presented as a percentage. The executions were, when possible, capped at 6 threads and 48 GB of RAM.

| Reconstruction tool | Execution time | SNPs | Identity | NCSD | NRC | RAM usage | CPU usage | Number of scaffolds | Reconstructed bases | Minimum scaffold length | Maximum scaffold length | Average scaffold length |
|---|---|---|---|---|---|---|---|---|---|---|---|---|
| coronaSPAdes | 2.445 | 36.0 | 99.884 | 0.036 | 0.025 | 0.147 | 402.667 | 9.0 | 137420.0 | 81.0 | 112095.0 | 15268.9 |
| Haploflow | 2.39 | 60.0 | 99.949 | 0.055 | 0.046 | 0.07 | 99.333 | 42.0 | 143037.0 | 517.0 | 13976.0 | 3405.6 |
| IRMA | 154.425 | 0.0 | 100.0 | 0.095 | 0.127 | 0.876 | 503.0 | 4.0 | 128314.0 | 4872.0 | 110237.0 | 32078.5 |
| LAZYPIPE | 5.73 | 53.0 | 99.947 | 0.036 | 0.026 | 1.512 | 245.333 | 10.0 | 137370.0 | 468.0 | 88573.0 | 13737.0 |
| metaSPAdes | 4.265 | 37.0 | 99.824 | 0.037 | 0.027 | 0.226 | 395.667 | 7.0 | 136333.0 | 71.0 | 112370.0 | 19476.1 |
| metaviralSPAdes | – | – | – | – | – | – | – | – | – | – | – | – |
| PEHaplo | 25.725 | 80.0 | 99.919 | 0.08 | 0.065 | 0.828 | 129.0 | 204.0 | 167438.0 | 375.0 | 2820.0 | 820.8 |
| QuRe | 284.305 | 4.667 | 99.652 | 0.996 | 0.985 | 13.86 | 554.667 | 6.333 | 1569.333 | 123.0 | 347.0 | 247.4 |
| QVG | 21.065 | 157.0 | 99.891 | 0.032 | 0.024 | 1.401 | 138.333 | 5.0 | 268901.0 | 5387.0 | 125041.0 | 53780.2 |
| SPAdes | 6.355 | 32.0 | 99.931 | 0.035 | 0.025 | 0.153 | 320.0 | 19.0 | 145107.0 | 167.0 | 108192.0 | 7637.2 |
| SSAKE | 4.295 | 6.667 | 99.987 | 0.432 | 0.396 | 0.065 | 100.0 | 313.333 | 93668.0 | 100.667 | 1211.0 | 299.1 |
| TRACESPipe | 188.335 | 153.0 | 99.893 | 0.034 | 0.024 | 9.768 | 338.667 | 4.0 | 143860.0 | 5387.0 | 125041.0 | 35965.0 |
| TRACESPipeLite | 40.195 | 147.0 | 99.894 | 0.033 | 0.023 | 2.766 | 342.0 | 5.0 | 151693.0 | 5387.0 | 125041.0 | 30338.6 |
| VirGenA | 611.85 | 67.0 | 99.866 | 0.246 | 0.224 | 3.914 | 709.0 | 40.0 | 115017.0 | 52.0 | 15798.0 | 2875.4 |
| ViSpA | 149.83 | 158.0 | 99.884 | 0.033 | 0.024 | 5.002 | 90.667 | 5.0 | 143678.0 | 0.0 | 124984.0 | 28735.6 |
| V-pipe | 40.725 | 64.0 | 99.627 | 0.047 | 0.037 | 0.104 | 115.667 | 5.0 | 160429.0 | 5387.0 | 125041.0 | 32085.8 |

Table S8: Results obtained for DS6 using the benchmark proposed. The execution time was measured in seconds, the RAM usage was measured in GB and the CPU usage is presented as a percentage. The executions were, when possible, capped at 6 threads and 48 GB of RAM.

| Reconstruction tool | Execution time | SNPs | Identity | NCSD | NRC | RAM usage | CPU usage | Number of scaffolds | Reconstructed bases | Minimum scaffold length | Maximum scaffold length | Average scaffold length |
|---|---|---|---|---|---|---|---|---|---|---|---|---|
| coronaSPAdes | 2.735 | 58.0 | 99.912 | 0.036 | 0.026 | 0.148 | 410.333 | 10.0 | 137441.0 | 81.0 | 105120.0 | 13744.1 |
| Haploflow | 2.985 | 98.0 | 99.923 | 0.035 | 0.027 | 0.08 | 100.0 | 21.0 | 165970.0 | 934.0 | 31458.0 | 7903.3 |
| IRMA | 188.2 | 0.0 | 100.0 | 0.096 | 0.128 | 1.071 | 533.667 | 4.0 | 128304.0 | 4888.0 | 110240.0 | 32076.0 |
| LAZYPIPE | 6.575 | 60.667 | 99.946 | 0.037 | 0.027 | 1.521 | 238.333 | 8.0 | 136723.0 | 390.0 | 85843.0 | 17090.4 |
| metaSPAdes | 4.91 | 68.0 | 99.904 | 0.037 | 0.026 | 0.227 | 403.333 | 9.0 | 136939.0 | 71.0 | 105136.0 | 15215.4 |
| metaviralSPAdes | 5.37 | 1.0 | 99.94 | 0.97 | 0.961 | 0.227 | 433.333 | 1.0 | 5358.0 | 5358.0 | 5358.0 | 5358.0 |
| PEHaplo | 31.53 | 41.0 | 99.962 | 0.071 | 0.058 | 1.159 | 136.333 | 203.0 | 177772.0 | 375.0 | 3301.0 | 875.7 |
| QuRe | 453.73 | 8.333 | 99.483 | 0.991 | 0.976 | 14.141 | 594.333 | 2.667 | 2514.333 | 345.333 | 1321.0 | 833.167 |
| QVG | 25.5 | 149.0 | 99.893 | 0.031 | 0.024 | 1.407 | 141.667 | 5.0 | 268901.0 | 5387.0 | 125041.0 | 53780.2 |
| SPAdes | 5.505 | 16.0 | 99.944 | 0.034 | 0.023 | 0.162 | 384.667 | 25.0 | 146735.0 | 167.0 | 105533.0 | 5869.4 |
| SSAKE | 6.125 | 9.667 | 99.988 | 0.177 | 0.151 | 0.075 | 100.0 | 363.667 | 136568.0 | 101.0 | 2305.0 | 375.533 |
| TRACESPipe | 199.76 | 148.0 | 99.894 | 0.032 | 0.022 | 9.768 | 333.667 | 4.0 | 143860.0 | 5387.0 | 125041.0 | 35965.0 |
| TRACESPipeLite | 42.275 | 167.667 | 99.882 | 0.033 | 0.023 | 2.768 | 335.0 | 5.0 | 151693.0 | 5387.0 | 125041.0 | 30338.6 |
| VirGenA | 685.01 | 63.0 | 99.851 | 0.24 | 0.218 | 4.125 | 730.667 | 40.0 | 116189.0 | 70.0 | 11738.0 | 2904.7 |
| ViSpA | 219.8 | 156.0 | 99.891 | 0.034 | 0.025 | 5.223 | 97.333 | 5.0 | 143613.0 | 0.0 | 124985.0 | 28722.6 |
| V-pipe | 44.63 | 148.0 | 99.886 | 0.036 | 0.026 | 0.104 | 114.333 | 5.0 | 160429.0 | 5387.0 | 125041.0 | 32085.8 |

Table S9: Results obtained for DS7 using the benchmark proposed. The execution time was measured in seconds, the RAM usage was measured in GB and the CPU usage is presented as a percentage. The executions were, when possible, capped at 6 threads and 48 GB of RAM.

| Reconstruction tool | Execution time | SNPs | Identity | NCSD | NRC | RAM usage | CPU usage | Number of scaffolds | Reconstructed bases | Minimum scaffold length | Maximum scaffold length | Average scaffold length |
|---|---|---|---|---|---|---|---|---|---|---|---|---|
| coronaSPAdes | 2.96 | 60.0 | 99.864 | 0.036 | 0.025 | 0.149 | 417.333 | 11.0 | 137558.0 | 81.0 | 105120.0 | 12505.3 |
| Haploflow | 3.445 | 42.0 | 99.862 | 0.035 | 0.025 | 0.089 | 99.333 | 17.0 | 147954.0 | 1881.0 | 33983.0 | 8703.2 |
| IRMA | 220.62 | 0.0 | 100.0 | 0.095 | 0.127 | 1.268 | 553.0 | 4.0 | 128337.0 | 4883.0 | 110250.0 | 32084.3 |
| LAZYPIPE | 6.5 | 96.333 | 99.74 | 0.036 | 0.025 | 1.525 | 235.667 | 5.0 | 136271.0 | 5330.0 | 109786.0 | 27254.2 |
| metaSPAdes | 5.45 | 52.0 | 99.869 | 0.037 | 0.026 | 0.228 | 411.333 | 10.0 | 136939.0 | 71.0 | 105136.0 | 13693.9 |
| metaviralSPAdes | – | – | – | – | – | – | – | – | – | – | – | – |
| PEHaplo | 38.6 | 55.0 | 99.957 | 0.047 | 0.036 | 1.461 | 146.0 | 208.0 | 192617.0 | 375.0 | 3338.0 | 926.0 |
| QuRe | 6581.505 | 0.333 | 99.869 | 0.991 | 0.976 | 14.994 | 619.333 | 3.667 | 2340.333 | 308.333 | 1131.333 | 693.767 |
| QVG | 25.245 | 149.0 | 99.893 | 0.034 | 0.024 | 1.409 | 145.667 | 4.0 | 143860.0 | 5387.0 | 125041.0 | 35965.0 |
| SPAdes | 6.555 | 20.0 | 99.941 | 0.034 | 0.024 | 0.162 | 374.0 | 25.0 | 146717.0 | 167.0 | 105128.0 | 5868.7 |
| SSAKE | 10.24 | 15.667 | 99.983 | 0.078 | 0.061 | 0.087 | 100.0 | 294.333 | 151735.667 | 101.0 | 3245.0 | 515.867 |
| TRACESPipe | 198.46 | 162.0 | 99.885 | 0.032 | 0.022 | 9.768 | 330.0 | 4.0 | 143860.0 | 5387.0 | 125041.0 | 35965.0 |
| TRACESPipeLite | 43.595 | 149.333 | 99.893 | 0.033 | 0.023 | 2.771 | 328.667 | 5.0 | 151693.0 | 5387.0 | 125041.0 | 30338.6 |
| VirGenA | 735.6 | 88.0 | 99.89 | 0.227 | 0.206 | 4.215 | 742.0 | 20.333 | 117296.0 | 771.667 | 37979.667 | 6486.867 |
| ViSpA | 282.775 | 165.0 | 99.84 | 0.034 | 0.025 | 5.546 | 106.0 | 5.0 | 143678.0 | 0.0 | 124993.0 | 28735.6 |
| V-pipe | 46.825 | 147.0 | 99.886 | 0.035 | 0.025 | 0.104 | 114.667 | 5.0 | 160429.0 | 5387.0 | 125041.0 | 32085.8 |

Table S10: Results obtained for DS8 using the benchmark proposed. The execution time was measured in seconds, the RAM usage was measured in GB and the CPU usage is presented as a percentage. The executions were, when possible, capped at 6 threads and 48 GB of RAM.

| Reconstruction tool | Execution time | SNPs | Identity | NCSD | NRC | RAM usage | CPU usage | Number of scaffolds | Reconstructed bases | Minimum scaffold length | Maximum scaffold length | Average scaffold length |
|---|---|---|---|---|---|---|---|---|---|---|---|---|
| coronaSPAdes | 3.465 | 46.0 | 99.876 | 0.036 | 0.026 | 0.15 | 430.667 | 10.0 | 137518.0 | 81.0 | 105120.0 | 13751.8 |
| Haploflow | 4.445 | 119.0 | 99.87 | 0.034 | 0.024 | 0.106 | 99.667 | 12.0 | 147444.0 | 661.0 | 88978.0 | 12287.0 |
| IRMA | 285.545 | 0.0 | 100.0 | 0.095 | 0.127 | 1.65 | 582.667 | 4.0 | 128349.0 | 4887.0 | 110254.0 | 32087.3 |
| LAZYPIPE | 7.815 | 52.0 | 99.879 | 0.035 | 0.025 | 1.531 | 257.0 | 9.0 | 137996.0 | 425.0 | 85904.0 | 15332.9 |
| metaSPAdes | 6.63 | 21.0 | 99.829 | 0.037 | 0.026 | 0.229 | 421.0 | 10.0 | 136997.0 | 71.0 | 105136.0 | 13699.7 |
| metaviralSPAdes | – | – | – | – | – | – | – | – | – | – | – | – |
| PEHaplo | 61.135 | 51.0 | 99.951 | 0.04 | 0.03 | 2.199 | 143.333 | 186.0 | 208292.0 | 379.0 | 4211.0 | 1119.8 |
| QuRe | 1778.55 | 44.0 | 99.536 | 0.94 | 0.902 | 15.574 | 615.333 | 4.333 | 16232.0 | 613.0 | 4914.0 | 3528.0 |
| QVG | 29.54 | 147.0 | 99.894 | 0.031 | 0.024 | 1.412 | 157.667 | 5.0 | 268901.0 | 5387.0 | 125041.0 | 53780.2 |
| SPAdes | 7.41 | 22.0 | 99.94 | 0.034 | 0.024 | 0.165 | 394.0 | 25.0 | 146751.0 | 131.0 | 105954.0 | 5870.0 |
| SSAKE | 11.49 | 7.333 | 99.99 | 0.038 | 0.028 | 0.098 | 100.0 | 144.333 | 154382.0 | 113.0 | 10289.333 | 1074.2 |
| TRACESPipe | 203.685 | 126.0 | 99.912 | 0.032 | 0.022 | 9.768 | 335.333 | 4.0 | 143860.0 | 5387.0 | 125041.0 | 35965.0 |
| TRACESPipeLite | 46.81 | 135.667 | 99.906 | 0.032 | 0.022 | 2.778 | 316.333 | 5.0 | 151693.0 | 5387.0 | 125041.0 | 30338.6 |
| VirGenA | 834.305 | 96.0 | 99.869 | 0.234 | 0.211 | 4.144 | 733.0 | 41.0 | 116748.0 | 368.0 | 11086.0 | 2847.5 |
| ViSpA | 472.815 | 158.0 | 99.884 | 0.034 | 0.024 | 5.217 | 113.333 | 5.0 | 143720.0 | 0.0 | 124998.0 | 28744.0 |
| V-pipe | 52.95 | 160.0 | 99.887 | 0.035 | 0.025 | 0.104 | 114.333 | 5.0 | 160429.0 | 5387.0 | 125041.0 | 32085.8 |

Table S11: Results obtained for DS9 using the benchmark proposed. The execution time was measured in seconds, the RAM usage was measured in GB and the CPU usage is presented as a percentage. The executions were, when possible, capped at 6 threads and 48 GB of RAM.

| Reconstruction tool | Execution time | SNPs | Identity | NCSD | NRC | RAM usage | CPU usage | Number of scaffolds | Reconstructed bases | Minimum scaffold length | Maximum scaffold length | Average scaffold length |
|---|---|---|---|---|---|---|---|---|---|---|---|---|
| coronaSPAdes | 1.26 | 117.0 | 99.856 | 0.47 | 0.428 | 0.134 | 357.333 | 267.0 | 120540.0 | 100.0 | 1650.0 | 451.5 |
| Haploflow | – | – | – | – | – | – | – | – | – | – | – | – |
| IRMA | 37.405 | 113.667 | 96.415 | 0.399 | 0.402 | 0.124 | 332.0 | 5.0 | 105134.0 | 3507.0 | 79651.0 | 21026.8 |
| LAZYPIPE | 4.345 | 61.0 | 99.89 | 0.645 | 0.607 | 1.496 | 184.333 | 162.0 | 79968.0 | 301.0 | 1571.0 | 493.6 |
| metaSPAdes | 2.075 | 114.0 | 99.866 | 0.439 | 0.401 | 0.225 | 326.667 | 305.0 | 127880.0 | 71.0 | 1650.0 | 419.3 |
| metaviralSPAdes | – | – | – | – | – | – | – | – | – | – | – | – |
| PEHaplo | 9.67 | 1616.0 | 99.819 | 0.641 | 0.598 | 0.091 | 62.667 | 135.0 | 82627.0 | 378.0 | 2690.0 | 612.1 |
| QuRe | – | – | – | – | – | – | – | – | – | – | – | – |
| QVG | 15.65 | 1300.0 | 99.059 | 0.115 | 0.094 | 1.389 | 100.667 | 3.0 | 138264.0 | 5387.0 | 125041.0 | 46088.0 |
| SPAdes | 2.15 | 94.0 | 99.88 | 0.481 | 0.443 | 0.148 | 368.0 | 258.0 | 118081.0 | 104.0 | 1650.0 | 457.7 |
| SSAKE | 2.74 | 0.0 | 66.667 | 0.999 | 0.994 | 0.049 | 100.0 | 3.667 | 538.333 | 116.333 | 177.0 | 145.767 |
| TRACESPipe | 159.42 | 127.0 | 96.979 | 0.334 | 0.298 | 9.768 | 354.333 | 4.0 | 143860.0 | 5387.0 | 125041.0 | 35965.0 |
| TRACESPipeLite | 35.185 | 62.0 | 97.105 | 0.34 | 0.303 | 2.777 | 367.0 | 6.0 | 168289.0 | 5387.0 | 125041.0 | 28048.2 |
| VirGenA | 252.57 | 82.0 | 99.846 | 0.589 | 0.554 | 3.654 | 599.667 | 199.333 | 66905.0 | 48.333 | 1010.0 | 335.633 |
| ViSpA | 43.605 | 91.667 | 94.393 | 0.516 | 0.492 | 5.573 | 58.333 | 85.0 | 2544032.0 | 796.0 | 70014.0 | 29929.8 |
| V-pipe | 30.595 | 0.0 | 98.428 | 0.971 | 0.957 | 0.104 | 115.0 | 5.0 | 160429.0 | 5387.0 | 125041.0 | 32085.8 |

Table S12: Results obtained for DS10 using the benchmark proposed. The execution time was measured in seconds, the RAM usage was measured in GB and the CPU usage is presented as a percentage. The executions were, when possible, capped at 6 threads and 48 GB of RAM.

| **Reconstruction tool** | **Execution time** | **SNPs** | **Identity** | **NCSD** | **NRC** | **RAM usage** | **CPU usage** | **Number of scaffolds** | **Reconstructed bases** | **Minimum scaffold length** | **Maximum scaffold length** | **Average scaffold length** |
|---|---|---|---|---|---|---|---|---|---|---|---|---|
| coronaSPAdes | 1.65 | 132.0 | 99.856 | 0.108 | 0.085 | 0.139 | 397.333 | 174.0 | 194352.0 | 239.0 | 5075.0 | 1117.0 |
| Haploflow | – | – | – | – | – | – | – | – | – | – | – | – |
| IRMA | 65.315 | 45.0 | 98.672 | 0.155 | 0.178 | 0.275 | 396.0 | 5.0 | 139050.333 | 4800.0 | 105942.0 | 27810.067 |
| LAZYPIPE | 4.82 | 109.0 | 99.867 | 0.191 | 0.162 | 1.502 | 211.667 | 145.0 | 175180.0 | 304.0 | 8726.0 | 1208.1 |
| metaSPAdes | 68.465 | 147.333 | 99.871 | 0.19 | 0.162 | 0.227 | 589.0 | 275.0 | 179576.0 | 137.0 | 3643.333 | 653.6 |
| metaviralSPAdes | – | – | – | – | – | – | – | – | – | – | – | – |
| PEHaplo | 11.785 | 88.0 | 99.917 | 0.173 | 0.145 | 0.171 | 82.0 | 156.0 | 184050.0 | 377.0 | 3812.0 | 1179.8 |
| QuRe | – | – | – | – | – | – | – | – | – | – | – | – |
| QVG | 17.165 | 989.0 | 99.317 | 0.068 | 0.048 | 1.394 | 112.0 | 4.0 | 143860.0 | 5387.0 | 125041.0 | 35965.0 |
| SPAdes | 4.205 | 118.0 | 99.716 | 0.112 | 0.091 | 0.151 | 320.0 | 167.0 | 192461.0 | 172.0 | 8313.0 | 1152.5 |
| SSAKE | 6.03 | 1.0 | 99.988 | 0.958 | 0.94 | 0.068 | 100.0 | 74.667 | 11568.667 | 100.667 | 249.333 | 154.867 |
| TRACESPipe | 174.135 | 177.0 | 98.896 | 0.096 | 0.077 | 9.768 | 350.333 | 4.0 | 143860.0 | 5387.0 | 125041.0 | 35965.0 |
| TRACESPipeLite | 37.15 | 149.667 | 98.858 | 0.091 | 0.073 | 2.81 | 356.333 | 6.0 | 168262.0 | 5387.0 | 125041.0 | 28043.7 |
| VirGenA | 378.31 | 65.0 | 99.83 | 0.535 | 0.506 | 4.446 | 681.0 | 124.0 | 73908.0 | 50.0 | 2356.0 | 596.0 |
| ViSpA | 82.4 | 125.0 | 96.964 | 0.157 | 0.139 | 6.633 | 75.667 | 92.0 | 4473795.0 | 661.0 | 113638.0 | 48628.2 |
| V-pipe | 34.08 | 22.0 | 94.328 | 0.643 | 0.606 | 0.104 | 115.0 | 5.0 | 160429.0 | 5387.0 | 125041.0 | 32085.8 |

Table S13: Results obtained for DS11 using the benchmark proposed. The execution time was measured in seconds, the RAM usage was measured in GB and the CPU usage is presented as a percentage. The executions were, when possible, capped at 6 threads and 48 GB of RAM.

| **Reconstruction tool** | **Execution time** | **SNPs** | **Identity** | **NCSD** | **NRC** | **RAM usage** | **CPU usage** | **Number of scaffolds** | **Reconstructed bases** | **Minimum scaffold length** | **Maximum scaffold length** | **Average scaffold length** |
|---|---|---|---|---|---|---|---|---|---|---|---|---|
| coronaSPAdes | 2.205 | 83.0 | 99.825 | 0.04 | 0.029 | 0.141 | 411.667 | 24.0 | 203993.0 | 81.0 | 33349.0 | 8499.7 |
| Haploflow | 2.06 | 38.0 | 99.933 | 0.616 | 0.577 | 0.07 | 99.333 | 107.0 | 86028.0 | 501.0 | 2339.0 | 804.0 |
| IRMA | 94.69 | 3.333 | 99.975 | 0.099 | 0.131 | 0.482 | 460.0 | 5.0 | 144593.0 | 4813.0 | 110179.0 | 28918.6 |
| LAZYPIPE | 5.515 | 89.0 | 99.911 | 0.044 | 0.033 | 1.509 | 248.0 | 20.0 | 202280.0 | 311.0 | 28959.0 | 10114.0 |
| metaSPAdes | 3.82 | 47.0 | 99.853 | 0.041 | 0.029 | 0.227 | 398.667 | 25.0 | 203651.0 | 71.0 | 33395.0 | 8146.0 |
| metaviralSPAdes | – | – | – | – | – | – | – | – | – | – | – | – |
| PEHaplo | 17.03 | 58.0 | 99.952 | 0.093 | 0.076 | 0.427 | 103.667 | 202.0 | 222692.0 | 380.0 | 4629.0 | 1102.4 |
| QuRe | – | – | – | – | – | – | – | – | – | – | – | – |
| QVG | 19.385 | 408.0 | 99.715 | 0.049 | 0.032 | 1.399 | 126.667 | 4.0 | 143860.0 | 5387.0 | 125041.0 | 35965.0 |
| SPAdes | 12.11 | 27.0 | 99.862 | 0.04 | 0.029 | 0.158 | 210.0 | 26.0 | 204678.0 | 97.0 | 33317.0 | 7872.2 |
| SSAKE | 1.67 | 1.667 | 99.956 | 0.92 | 0.9 | 0.048 | 100.0 | 109.333 | 19881.333 | 100.667 | 472.0 | 181.767 |
| TRACESPipe | 171.255 | 135.0 | 99.861 | 0.036 | 0.026 | 9.768 | 355.0 | 4.0 | 143860.0 | 5387.0 | 125041.0 | 35965.0 |
| TRACESPipeLite | 39.35 | 151.0 | 99.861 | 0.035 | 0.026 | 2.816 | 344.667 | 6.0 | 168262.0 | 5387.0 | 125041.0 | 28043.7 |
| VirGenA | 576.06 | 46.0 | 99.872 | 0.6 | 0.573 | 4.565 | 730.0 | 82.667 | 63628.0 | 138.333 | 2743.0 | 808.1 |
| ViSpA | 41.54 | 3.0 | 99.732 | 0.043 | 0.033 | 5.923 | 91.0 | 8.0 | 329578.0 | 5254.0 | 124613.0 | 41197.3 |
| V-pipe | 37.86 | 54.0 | 96.844 | 0.245 | 0.213 | 0.104 | 114.667 | 5.0 | 160429.0 | 5387.0 | 125041.0 | 32085.8 |

Table S14: Results obtained for DS12 using the benchmark proposed. The execution time was measured in seconds, the RAM usage was measured in GB and the CPU usage is presented as a percentage. The executions were, when possible, capped at 6 threads and 48 GB of RAM.

| **Reconstruction tool** | **Execution time** | **SNPs** | **Identity** | **NCSD** | **NRC** | **RAM usage** | **CPU usage** | **Number of scaffolds** | **Reconstructed bases** | **Minimum scaffold length** | **Maximum scaffold length** | **Average scaffold length** |
|---|---|---|---|---|---|---|---|---|---|---|---|---|
| coronaSPAdes | 2.67 | 53.0 | 99.867 | 0.037 | 0.026 | 0.148 | 415.333 | 12.0 | 203872.0 | 81.0 | 67883.0 | 16989.3 |
| Haploflow | 2.89 | 75.0 | 99.932 | 0.205 | 0.179 | 0.086 | 99.0 | 118.0 | 180100.0 | 506.0 | 9199.0 | 1526.3 |
| IRMA | 130.94 | 0.0 | 99.984 | 0.096 | 0.128 | 0.678 | 505.333 | 5.0 | 144771.0 | 4882.0 | 110219.0 | 28954.2 |
| LAZYPIPE | 6.225 | 74.0 | 99.938 | 0.039 | 0.028 | 1.518 | 261.0 | 10.0 | 202548.0 | 364.0 | 71673.0 | 20254.8 |
| metaSPAdes | 4.78 | 55.0 | 99.823 | 0.038 | 0.027 | 0.229 | 408.667 | 13.0 | 203274.0 | 71.0 | 67865.0 | 15636.5 |
| metaviralSPAdes | – | – | – | – | – | – | – | – | – | – | – | – |
| PEHaplo | 23.755 | 38.0 | 99.969 | 0.078 | 0.062 | 0.762 | 119.667 | 276.0 | 236245.0 | 376.0 | 3084.0 | 856.0 |
| QuRe | 867.745 | 0.0 | 100.0 | 0.999 | 0.992 | 43.794 | 557.333 | 6.333 | 551.667 | 53.667 | 117.333 | 88.933 |
| QVG | 22.37 | 225.0 | 99.843 | 0.04 | 0.027 | 1.405 | 143.667 | 5.0 | 149456.0 | 5387.0 | 125041.0 | 29891.2 |
| SPAdes | 8.97 | 47.0 | 99.918 | 0.037 | 0.027 | 0.162 | 280.0 | 19.0 | 205739.0 | 144.0 | 71345.0 | 10828.4 |
| SSAKE | 2.64 | 6.0 | 99.979 | 0.81 | 0.782 | 0.061 | 100.0 | 202.0 | 47187.0 | 103.0 | 809.667 | 233.567 |
| TRACESPipe | 180.16 | 147.0 | 99.896 | 0.034 | 0.024 | 9.768 | 358.333 | 4.0 | 143860.0 | 5387.0 | 125041.0 | 35965.0 |
| TRACESPipeLite | 41.75 | 135.667 | 99.897 | 0.034 | 0.024 | 2.818 | 332.667 | 6.0 | 168262.0 | 5387.0 | 125041.0 | 28043.7 |
| VirGenA | 692.265 | 46.0 | 99.864 | 0.627 | 0.596 | 4.444 | 718.333 | 72.0 | 59531.0 | 61.0 | 3075.0 | 826.8 |
| ViSpA | 68.98 | 150.0 | 99.889 | 0.037 | 0.027 | 5.775 | 103.667 | 6.0 | 197005.0 | 5356.0 | 124834.0 | 32834.2 |
| V-pipe | 42.695 | 145.0 | 99.134 | 0.071 | 0.054 | 0.104 | 114.667 | 5.0 | 160429.0 | 5387.0 | 125041.0 | 32085.8 |

Table S15: Results obtained for DS13 using the benchmark proposed. The execution time was measured in seconds, the RAM usage was measured in GB and the CPU usage is presented as a percentage. The executions were, when possible, capped at 6 threads and 48 GB of RAM.

| Reconstruction tool | Execution time | SNPs | Identity | NCSD | NRC | RAM usage | CPU usage | Number of scaffolds | Reconstructed bases | Minimum scaffold length | Maximum scaffold length | Average scaffold length |
|---|---|---|---|---|---|---|---|---|---|---|---|---|
| coronaSPAdes | 3.07 | 44.0 | 99.821 | 0.036 | 0.025 | 0.149 | 422.333 | 14.0 | 204248.0 | 81.0 | 105072.0 | 14589.1 |
| Haploflow | 3.55 | 92.0 | 99.913 | 0.054 | 0.042 | 0.099 | 99.333 | 42.0 | 208025.0 | 631.0 | 17503.0 | 4953.0 |
| IRMA | 166.675 | 0.0 | 100.0 | 0.096 | 0.128 | 0.877 | 540.667 | 5.0 | 144809.0 | 4884.0 | 110215.0 | 28961.8 |
| LAZYPIPE | 6.745 | 56.0 | 99.932 | 0.037 | 0.025 | 1.525 | 265.0 | 7.0 | 202811.0 | 314.0 | 117612.0 | 28973.0 |
| metaSPAdes | 5.575 | 59.0 | 99.861 | 0.037 | 0.026 | 0.229 | 416.667 | 13.0 | 203277.0 | 71.0 | 84069.0 | 15636.7 |
| metaviralSPAdes | – | – | – | – | – | – | – | – | – | – | – | – |
| PEHaplo | 31.845 | 87.0 | 99.937 | 0.062 | 0.047 | 1.173 | 132.333 | 305.0 | 249689.0 | 380.0 | 3353.0 | 818.7 |
| QuRe | 448.215 | 0.0 | 100.0 | 0.998 | 0.989 | 19.991 | 573.0 | 5.0 | 911.333 | 97.0 | 354.0 | 184.133 |
| QVG | 24.215 | 157.0 | 99.892 | 0.032 | 0.024 | 1.409 | 156.667 | 5.0 | 268901.0 | 5387.0 | 125041.0 | 53780.2 |
| SPAdes | 7.09 | 36.0 | 99.906 | 0.036 | 0.025 | 0.162 | 374.0 | 21.0 | 211829.0 | 131.0 | 105080.0 | 10087.1 |
| SSAKE | 5.59 | 21.0 | 99.968 | 0.511 | 0.477 | 0.079 | 100.0 | 364.667 | 116251.0 | 101.333 | 1365.0 | 318.7 |
| TRACESPipe | 177.0 | 143.0 | 99.902 | 0.032 | 0.022 | 9.768 | 371.0 | 4.0 | 143860.0 | 5387.0 | 125041.0 | 35965.0 |
| TRACESPipeLite | 45.255 | 124.0 | 99.9 | 0.033 | 0.023 | 2.82 | 322.333 | 6.0 | 168262.0 | 5387.0 | 125041.0 | 28043.7 |
| VirGenA | 811.215 | 45.0 | 99.882 | 0.647 | 0.614 | 4.22 | 740.0 | 66.0 | 57997.0 | 51.0 | 3006.0 | 878.7 |
| ViSpA | 138.105 | 153.0 | 99.893 | 0.034 | 0.025 | 6.249 | 111.0 | 5.0 | 160089.0 | 5343.0 | 124970.0 | 32017.8 |
| V-pipe | 46.535 | 128.0 | 99.765 | 0.043 | 0.032 | 0.104 | 114.0 | 5.0 | 160429.0 | 5387.0 | 125041.0 | 32085.8 |

Table S16: Results obtained for DS14 using the benchmark proposed. The execution time was measured in seconds, the RAM usage was measured in GB and the CPU usage is presented as a percentage. The executions were, when possible, capped at 6 threads and 48 GB of RAM.

| Reconstruction tool | Execution time | SNPs | Identity | NCSD | NRC | RAM usage | CPU usage | Number of scaffolds | Reconstructed bases | Minimum scaffold length | Maximum scaffold length | Average scaffold length |
|---|---|---|---|---|---|---|---|---|---|---|---|---|
| coronaSPAdes | 3.45 | 63.0 | 99.91 | 0.036 | 0.025 | 0.149 | 431.0 | 10.0 | 203804.0 | 81.0 | 112188.0 | 20380.4 |
| Haploflow | 4.24 | 80.0 | 99.92 | 0.044 | 0.034 | 0.112 | 99.0 | 30.0 | 213645.0 | 752.0 | 23263.0 | 7121.5 |
| IRMA | 201.46 | 0.0 | 100.0 | 0.095 | 0.128 | 1.069 | 562.0 | 5.0 | 144874.0 | 4878.0 | 110250.0 | 28974.8 |
| LAZYPIPE | 7.56 | 61.0 | 99.915 | 0.036 | 0.025 | 1.529 | 268.0 | 10.0 | 203315.0 | 425.0 | 92634.0 | 20331.5 |
| metaSPAdes | 6.365 | 31.0 | 99.823 | 0.037 | 0.026 | 0.23 | 420.333 | 12.0 | 203510.0 | 71.0 | 105136.0 | 16959.2 |
| metaviralSPAdes | – | – | – | – | – | – | – | – | – | – | – | – |
| PEHaplo | 48.675 | 61.0 | 99.954 | 0.072 | 0.057 | 1.594 | 157.667 | 289.0 | 258898.0 | 376.0 | 3938.0 | 895.8 |
| QuRe | 520.955 | 4.333 | 99.743 | 0.993 | 0.979 | 17.191 | 580.0 | 4.333 | 2053.333 | 215.667 | 888.0 | 478.7 |
| QVG | 26.675 | 150.0 | 99.893 | 0.034 | 0.024 | 1.412 | 158.0 | 4.0 | 143860.0 | 5387.0 | 125041.0 | 35965.0 |
| SPAdes | 7.39 | 24.0 | 99.938 | 0.035 | 0.024 | 0.166 | 392.0 | 25.0 | 213118.0 | 167.0 | 105954.0 | 8524.7 |
| SSAKE | 8.7 | 26.333 | 99.973 | 0.193 | 0.167 | 0.096 | 100.0 | 475.0 | 191768.333 | 103.0 | 1626.0 | 403.833 |
| TRACESPipe | 177.085 | 136.0 | 99.903 | 0.033 | 0.023 | 9.768 | 358.0 | 4.0 | 143861.0 | 5387.0 | 125042.0 | 35965.3 |
| TRACESPipeLite | 47.055 | 159.333 | 99.888 | 0.033 | 0.023 | 2.822 | 316.0 | 6.0 | 168262.0 | 5387.0 | 125041.0 | 28043.7 |
| VirGenA | 924.645 | 44.0 | 99.867 | 0.635 | 0.605 | 4.426 | 729.0 | 57.667 | 57130.333 | 159.333 | 3971.0 | 1034.4 |
| ViSpA | 227.875 | 142.0 | 99.901 | 0.034 | 0.024 | 6.886 | 117.667 | 5.0 | 160249.0 | 5369.0 | 125022.0 | 32049.8 |
| V-pipe | 51.565 | 134.0 | 99.89 | 0.035 | 0.025 | 0.104 | 113.667 | 5.0 | 160429.0 | 5387.0 | 125041.0 | 32085.8 |

Table S17: Results obtained for DS15 using the benchmark proposed. The execution time was measured in seconds, the RAM usage was measured in GB and the CPU usage is presented as a percentage. The executions were, when possible, capped at 6 threads and 48 GB of RAM.

| Reconstruction tool | Execution time | SNPs | Identity | NCSD | NRC | RAM usage | CPU usage | Number of scaffolds | Reconstructed bases | Minimum scaffold length | Maximum scaffold length | Average scaffold length |
|---|---|---|---|---|---|---|---|---|---|---|---|---|
| coronaSPAdes | 3.95 | 52.0 | 99.916 | 0.036 | 0.025 | 0.156 | 434.333 | 12.0 | 204047.0 | 81.0 | 105120.0 | 17003.9 |
| Haploflow | 5.015 | 103.0 | 99.908 | 0.035 | 0.026 | 0.125 | 99.333 | 20.0 | 214389.0 | 1571.0 | 64331.0 | 10719.5 |
| IRMA | 237.415 | 0.0 | 100.0 | 0.095 | 0.127 | 1.265 | 581.667 | 5.0 | 144895.0 | 4888.0 | 110259.0 | 28979.0 |
| LAZYPIPE | 7.785 | 66.0 | 99.929 | 0.036 | 0.025 | 1.533 | 254.0 | 8.0 | 202788.0 | 5330.0 | 95777.0 | 25348.5 |
| metaSPAdes | 7.14 | 40.0 | 99.836 | 0.037 | 0.026 | 0.229 | 424.0 | 13.0 | 203610.0 | 71.0 | 105136.0 | 15662.3 |
| metaviralSPAdes | – | – | – | – | – | – | – | – | – | – | – | – |
| PEHaplo | 59.915 | 66.0 | 99.952 | 0.057 | 0.045 | 2.0 | 155.0 | 313.0 | 292885.0 | 382.0 | 2814.0 | 935.7 |
| QuRe | 931.685 | 21.333 | 99.609 | 0.988 | 0.973 | 20.736 | 605.333 | 5.0 | 4849.667 | 139.0 | 1503.667 | 851.733 |
| QVG | 29.835 | 148.0 | 99.893 | 0.034 | 0.024 | 1.413 | 160.333 | 4.0 | 143860.0 | 5387.0 | 125041.0 | 35965.0 |
| SPAdes | 8.485 | 13.0 | 99.947 | 0.033 | 0.023 | 0.175 | 397.333 | 27.0 | 208155.0 | 131.0 | 105954.0 | 7709.4 |
| SSAKE | 10.43 | 28.333 | 99.926 | 0.084 | 0.067 | 0.109 | 100.0 | 405.667 | 219691.667 | 100.333 | 3179.0 | 541.6 |
| TRACESPipe | 184.94 | 140.0 | 99.904 | 0.033 | 0.024 | 9.768 | 362.0 | 4.0 | 143860.0 | 5387.0 | 125041.0 | 35965.0 |
| TRACESPipeLite | 48.405 | 130.333 | 99.908 | 0.032 | 0.023 | 2.824 | 307.333 | 6.0 | 168262.0 | 5387.0 | 125041.0 | 28043.7 |
| VirGenA | 923.85 | 43.0 | 99.888 | 0.647 | 0.62 | 4.896 | 741.0 | 69.0 | 56205.0 | 54.0 | 3365.0 | 814.6 |
| ViSpA | 292.765 | 166.0 | 99.884 | 0.035 | 0.024 | 6.876 | 123.333 | 5.0 | 160211.0 | 5368.0 | 125006.0 | 32042.2 |
| V-pipe | 55.025 | 165.0 | 99.887 | 0.035 | 0.025 | 0.104 | 113.667 | 5.0 | 160429.0 | 5387.0 | 125041.0 | 32085.8 |

Table S18: Results obtained for DS16 using the benchmark proposed. The execution time was measured in seconds, the RAM usage was measured in GB and the CPU usage is presented as a percentage. The executions were, when possible, capped at 6 threads and 48 GB of RAM.

| **Reconstruction tool** | **Execution time** | **SNPs** | **Identity** | **NCSD** | **NRC** | **RAM usage** | **CPU usage** | **Number of scaffolds** | **Reconstructed bases** | **Minimum scaffold length** | **Maximum scaffold length** | **Average scaffold length** |
|---|---|---|---|---|---|---|---|---|---|---|---|---|
| coronaSPAdes | 4.66 | 47.0 | 99.923 | 0.036 | 0.025 | 0.153 | 444.667 | 10.0 | 203843.0 | 81.0 | 112188.0 | 20384.3 |
| Haploflow | 6.52 | 113.0 | 99.876 | 0.033 | 0.024 | 0.151 | 99.667 | 12.0 | 211760.0 | 2592.0 | 88805.0 | 17646.7 |
| IRMA | 308.225 | 0.0 | 100.0 | 0.095 | 0.127 | 1.651 | 598.0 | 5.0 | 144906.0 | 4885.0 | 110253.0 | 28981.2 |
| LAZYPIPE | 9.555 | 53.0 | 99.942 | 0.036 | 0.025 | 1.54 | 270.667 | 9.0 | 203432.0 | 5371.0 | 88951.0 | 22603.6 |
| metaSPAdes | 8.68 | 24.0 | 99.828 | 0.037 | 0.026 | 0.227 | 432.0 | 12.0 | 203549.0 | 71.0 | 105136.0 | 16962.4 |
| metaviralSPAdes | 10.065 | 0.0 | 0.0 | 1.0 | 0.994 | 0.226 | 453.0 | 2.0 | 66553.0 | 16561.0 | 49992.0 | 33276.5 |
| PEHaplo | 81.155 | 64.0 | 99.938 | 0.038 | 0.029 | 2.988 | 149.667 | 220.0 | 321707.0 | 395.0 | 5006.0 | 1462.3 |
| QuRe | 1980.37 | 31.333 | 99.689 | 0.936 | 0.898 | 21.231 | 638.667 | 8.667 | 26562.667 | 410.667 | 5081.0 | 3012.8 |
| QVG | 33.705 | 147.0 | 99.894 | 0.034 | 0.024 | 1.418 | 174.0 | 4.0 | 143860.0 | 5387.0 | 125041.0 | 35965.0 |
| SPAdes | 9.73 | 22.0 | 99.94 | 0.034 | 0.023 | 0.185 | 410.333 | 27.0 | 213297.0 | 131.0 | 105954.0 | 7899.9 |
| SSAKE | 16.85 | 11.667 | 99.989 | 0.04 | 0.03 | 0.132 | 99.667 | 214.0 | 225447.333 | 107.0 | 8185.0 | 1055.367 |
| TRACESPipe | 187.29 | 151.0 | 99.893 | 0.032 | 0.023 | 9.768 | 365.667 | 4.0 | 143860.0 | 5387.0 | 125041.0 | 35965.0 |
| TRACESPipeLite | 53.835 | 142.667 | 99.9 | 0.032 | 0.023 | 2.827 | 293.0 | 6.0 | 168262.0 | 5387.0 | 125041.0 | 28043.7 |
| VirGenA | 1174.83 | 40.0 | 99.844 | 0.629 | 0.596 | 5.112 | 745.667 | 55.0 | 59033.0 | 117.0 | 2741.0 | 1073.3 |
| ViSpA | 478.625 | 164.0 | 99.884 | 0.034 | 0.024 | 6.95 | 128.333 | 5.0 | 160258.0 | 5371.0 | 125015.0 | 32051.6 |
| V-pipe | 63.495 | 132.0 | 99.904 | 0.034 | 0.024 | 0.104 | 113.0 | 5.0 | 160429.0 | 5387.0 | 125041.0 | 32085.8 |

Table S19: Results obtained for DS17 using the benchmark proposed. The execution time was measured in seconds, the RAM usage was measured in GB and the CPU usage is presented as a percentage. The executions were, when possible, capped at 6 threads and 48 GB of RAM.

| **Reconstruction tool** | **Execution time** | **SNPs** | **Identity** | **NCSD** | **NRC** | **RAM usage** | **CPU usage** | **Number of scaffolds** | **Reconstructed bases** | **Minimum scaffold length** | **Maximum scaffold length** | **Average scaffold length** |
|---|---|---|---|---|---|---|---|---|---|---|---|---|
| coronaSPAdes | 1.25 | 95.0 | 99.886 | 0.448 | 0.413 | 0.134 | 355.333 | 259.0 | 122290.0 | 100.0 | 1626.0 | 472.2 |
| Haploflow | – | – | – | – | – | – | – | – | – | – | – | – |
| IRMA | 38.91 | 108.333 | 95.83 | 0.371 | 0.379 | 0.129 | 326.667 | 5.0 | 108214.0 | 3983.0 | 82349.0 | 21642.8 |
| LAZYPIPE | 4.09 | 29.0 | 99.949 | 0.62 | 0.583 | 1.495 | 178.333 | 159.0 | 82856.0 | 306.0 | 1572.0 | 521.1 |
| metaSPAdes | 2.075 | 83.0 | 99.909 | 0.393 | 0.357 | 0.225 | 326.667 | 306.0 | 132688.0 | 56.0 | 1626.0 | 433.6 |
| metaviralSPAdes | – | – | – | – | – | – | – | – | – | – | – | – |
| PEHaplo | 9.67 | 654.333 | 99.878 | 0.603 | 0.566 | 0.091 | 62.0 | 137.0 | 86539.0 | 375.0 | 1552.0 | 631.7 |
| QuRe | – | – | – | – | – | – | – | – | – | – | – | – |
| QVG | – | – | – | – | – | – | – | – | – | – | – | – |
| SPAdes | – | – | – | – | – | – | – | – | – | – | – | – |
| SSAKE | 2.62 | 0.0 | 100.0 | 0.999 | 0.993 | 0.049 | 100.0 | 2.667 | 410.0 | 141.333 | 161.667 | 150.167 |
| TRACESPipe | 164.61 | 15.0 | 97.482 | 0.324 | 0.287 | 9.768 | 361.0 | 4.0 | 143860.0 | 5387.0 | 125041.0 | 35965.0 |
| TRACESPipeLite | 37.01 | 14.333 | 96.444 | 0.317 | 0.286 | 2.776 | 362.0 | 6.0 | 168262.0 | 5387.0 | 125041.0 | 28043.7 |
| VirGenA | 263.35 | 55.0 | 99.867 | 0.576 | 0.543 | 3.824 | 608.333 | 214.0 | 68240.0 | 52.0 | 1205.0 | 318.9 |
| ViSpA | 43.22 | 54.0 | 96.054 | 0.501 | 0.482 | 5.525 | 56.333 | 86.0 | 2411952.0 | 450.0 | 70566.0 | 28046.0 |
| V-pipe | 30.85 | 0.0 | 95.912 | 0.975 | 0.961 | 0.104 | 115.333 | 5.0 | 160429.0 | 5387.0 | 125041.0 | 32085.8 |

Table S20: Results obtained for DS18 using the benchmark proposed. The execution time was measured in seconds, the RAM usage was measured in GB and the CPU usage is presented as a percentage. The executions were, when possible, capped at 6 threads and 48 GB of RAM.

| **Reconstruction tool** | **Execution time** | **SNPs** | **Identity** | **NCSD** | **NRC** | **RAM usage** | **CPU usage** | **Number of scaffolds** | **Reconstructed bases** | **Minimum scaffold length** | **Maximum scaffold length** | **Average scaffold length** |
|---|---|---|---|---|---|---|---|---|---|---|---|---|
| coronaSPAdes | 1.27 | 215.0 | 99.73 | 0.492 | 0.456 | 0.135 | 357.0 | 272.0 | 120288.0 | 224.0 | 1540.0 | 442.2 |
| Haploflow | – | – | – | – | – | – | – | – | – | – | – | – |
| IRMA | 42.26 | 115.0 | 95.586 | 0.412 | 0.409 | 0.122 | 326.667 | 5.0 | 104407.0 | 3598.0 | 79183.0 | 20881.4 |
| LAZYPIPE | 4.225 | 119.0 | 99.741 | 0.668 | 0.631 | 1.495 | 188.333 | 157.0 | 75760.0 | 301.0 | 1464.0 | 482.5 |
| metaSPAdes | 2.095 | 152.0 | 99.818 | 0.447 | 0.409 | 0.225 | 330.333 | 327.0 | 130710.0 | 97.0 | 1540.0 | 399.7 |
| metaviralSPAdes | – | – | – | – | – | – | – | – | – | – | – | – |
| PEHaplo | 9.575 | 2506.667 | 99.631 | 0.635 | 0.6 | 0.091 | 62.333 | 132.0 | 80018.0 | 375.0 | 2641.0 | 606.2 |
| QuRe | – | – | – | – | – | – | – | – | – | – | – | – |
| QVG | 14.94 | 4037.0 | 96.924 | 0.183 | 0.16 | 1.389 | 103.0 | 3.0 | 138473.0 | 5596.0 | 125041.0 | 46157.7 |
| SPAdes | – | – | – | – | – | – | – | – | – | – | – | – |
| SSAKE | 2.695 | 0.0 | 100.0 | 1.0 | 0.995 | 0.049 | 100.0 | 1.667 | 224.667 | 134.667 | 140.333 | 137.333 |
| TRACESPipe | 159.8 | 264.0 | 96.898 | 0.344 | 0.311 | 9.768 | 362.0 | 4.0 | 143859.0 | 5387.0 | 125040.0 | 35964.8 |
| TRACESPipeLite | 37.09 | 116.333 | 97.189 | 0.435 | 0.403 | 2.779 | 366.0 | 6.0 | 168076.0 | 5201.0 | 125041.0 | 28012.7 |
| VirGenA | 239.27 | 112.0 | 99.808 | 0.589 | 0.556 | 3.906 | 593.333 | 211.0 | 68194.0 | 62.0 | 1072.0 | 323.2 |
| ViSpA | 42.02 | 158.333 | 94.834 | 0.542 | 0.519 | 5.458 | 59.333 | 81.0 | 2419083.0 | 390.0 | 69277.0 | 29865.2 |
| V-pipe | 30.93 | 6.0 | 95.968 | 0.97 | 0.955 | 0.104 | 115.0 | 5.0 | 160429.0 | 5387.0 | 125041.0 | 32085.8 |

Table S21: Results obtained for DS19 using the benchmark proposed. The execution time was measured in seconds, the RAM usage was measured in GB and the CPU usage is presented as a percentage. The executions were, when possible, capped at 6 threads and 48 GB of RAM.

| **Reconstruction tool** | **Execution time** | **SNPs** | **Identity** | **NCSD** | **NRC** | **RAM usage** | **CPU usage** | **Number of scaffolds** | **Reconstructed bases** | **Minimum scaffold length** | **Maximum scaffold length** | **Average scaffold length** |
|---|---|---|---|---|---|---|---|---|---|---|---|---|
| coronaSPAdes | 1.27 | 164.0 | 99.656 | 0.48 | 0.446 | 0.134 | 357.667 | 274.0 | 124875.0 | 226.0 | 1302.0 | 455.7 |
| Haploflow | – | – | – | – | – | – | – | – | – | – | – | – |
| IRMA | 45.655 | 120.667 | 96.229 | 0.4 | 0.395 | 0.126 | 328.333 | 5.0 | 106677.0 | 3844.0 | 80453.0 | 21335.4 |
| LAZYPIPE | 3.935 | 60.0 | 99.773 | 0.633 | 0.603 | 1.495 | 174.667 | 182.0 | 86241.0 | 297.0 | 1144.0 | 473.9 |
| metaSPAdes | 2.105 | 126.0 | 99.747 | 0.439 | 0.407 | 0.224 | 328.333 | 323.0 | 134539.0 | 206.0 | 1235.0 | 416.5 |
| metaviralSPAdes | – | – | – | – | – | – | – | – | – | – | – | – |
| PEHaplo | 9.625 | 1272.0 | 99.598 | 0.632 | 0.6 | 0.091 | 62.333 | 147.0 | 86902.0 | 377.0 | 1507.0 | 591.2 |
| QuRe | – | – | – | – | – | – | – | – | – | – | – | – |
| QVG | 15.81 | 6858.0 | 95.231 | 0.207 | 0.186 | 1.389 | 102.333 | 4.0 | 143860.0 | 5387.0 | 125041.0 | 35965.0 |
| SPAdes | – | – | – | – | – | – | – | – | – | – | – | – |
| SSAKE | 2.62 | 0.0 | 100.0 | 1.0 | 0.996 | 0.049 | 100.0 | 4.5 | 672.5 | 123.5 | 173.5 | 149.7 |
| TRACESPipe | 159.595 | 555.0 | 96.286 | 0.324 | 0.295 | 9.768 | 361.667 | 4.0 | 143859.0 | 5387.0 | 125040.0 | 35964.8 |
| TRACESPipeLite | 36.79 | 71.0 | 98.034 | 0.693 | 0.667 | 2.782 | 365.0 | 6.0 | 168246.0 | 5387.0 | 125041.0 | 28041.0 |
| VirGenA | 241.31 | 85.0 | 99.79 | 0.608 | 0.58 | 3.738 | 600.667 | 195.0 | 64953.0 | 54.0 | 1673.0 | 333.1 |
| ViSpA | 41.965 | 299.0 | 94.594 | 0.536 | 0.518 | 5.581 | 57.667 | 86.0 | 2370844.0 | 744.0 | 69927.0 | 27568.0 |
| V-pipe | 30.795 | 1.0 | 95.875 | 0.972 | 0.959 | 0.104 | 115.333 | 5.0 | 160429.0 | 5387.0 | 125041.0 | 32085.8 |

Table S22: Results obtained for DS20 using the benchmark proposed. The execution time was measured in seconds, the RAM usage was measured in GB and the CPU usage is presented as a percentage. The executions were, when possible, capped at 6 threads and 48 GB of RAM.

| **Reconstruction tool** | **Execution time** | **SNPs** | **Identity** | **NCSD** | **NRC** | **RAM usage** | **CPU usage** | **Number of scaffolds** | **Reconstructed bases** | **Minimum scaffold length** | **Maximum scaffold length** | **Average scaffold length** |
|---|---|---|---|---|---|---|---|---|---|---|---|---|
| coronaSPAdes | 1.25 | 245.0 | 99.694 | 0.501 | 0.473 | 0.134 | 355.333 | 265.0 | 117128.0 | 224.0 | 1400.0 | 442.0 |
| Haploflow | – | – | – | – | – | – | – | – | – | – | – | – |
| IRMA | 44.69 | 120.0 | 96.585 | 0.41 | 0.401 | 0.124 | 332.333 | 5.0 | 105256.0 | 3636.0 | 79641.0 | 21051.2 |
| LAZYPIPE | 3.95 | 48.0 | 99.831 | 0.679 | 0.659 | 1.495 | 180.667 | 154.0 | 72806.0 | 301.0 | 1331.0 | 472.8 |
| metaSPAdes | 2.09 | 163.0 | 99.805 | 0.442 | 0.415 | 0.225 | 328.333 | 327.0 | 130602.0 | 206.0 | 1400.0 | 399.4 |
| metaviralSPAdes | – | – | – | – | – | – | – | – | – | – | – | – |
| PEHaplo | 9.63 | 2558.0 | 99.72 | 0.639 | 0.613 | 0.091 | 62.333 | 138.0 | 81582.0 | 377.0 | 1409.0 | 591.2 |
| QuRe | – | – | – | – | – | – | – | – | – | – | – | – |
| QVG | 15.93 | 9676.0 | 93.217 | 0.264 | 0.246 | 1.389 | 103.0 | 4.0 | 143860.0 | 5387.0 | 125041.0 | 35965.0 |
| SPAdes | – | – | – | – | – | – | – | – | – | – | – | – |
| SSAKE | 2.71 | 0.0 | 100.0 | 0.999 | 0.994 | 0.049 | 99.667 | 4.333 | 674.333 | 129.667 | 182.0 | 156.067 |
| TRACESPipe | 164.135 | 580.0 | 96.547 | 0.368 | 0.34 | 9.768 | 346.0 | 4.0 | 143658.0 | 5185.0 | 125041.0 | 35914.5 |
| TRACESPipeLite | 36.01 | 3.0 | 98.918 | 0.951 | 0.943 | 2.783 | 369.333 | 6.0 | 168060.0 | 5201.0 | 125041.0 | 28010.0 |
| VirGenA | 243.54 | 75.333 | 99.82 | 0.601 | 0.578 | 3.817 | 603.667 | 201.667 | 66410.667 | 72.0 | 1400.0 | 329.967 |
| ViSpA | 39.75 | 231.333 | 95.15 | 0.553 | 0.539 | 5.384 | 58.0 | 77.0 | 2157354.0 | 412.0 | 66879.0 | 28017.6 |
| V-pipe | 30.835 | 7.0 | 96.532 | 0.967 | 0.956 | 0.104 | 115.667 | 5.0 | 160429.0 | 5387.0 | 125041.0 | 32085.8 |

Table S23: Results obtained for DS21 using the benchmark proposed. The execution time was measured in seconds, the RAM usage was measured in GB and the CPU usage is presented as a percentage. The executions were, when possible, capped at 6 threads and 48 GB of RAM.

| **Reconstruction tool** | **Execution time** | **SNPs** | **Identity** | **NCSD** | **NRC** | **RAM usage** | **CPU usage** | **Number of scaffolds** | **Reconstructed bases** | **Minimum scaffold length** | **Maximum scaffold length** | **Average scaffold length** |
|---|---|---|---|---|---|---|---|---|---|---|---|---|
| coronaSPAdes | 1.25 | 143.0 | 99.663 | 0.473 | 0.446 | 0.134 | 356.0 | 279.0 | 124323.0 | 224.0 | 1491.0 | 445.6 |
| Haploflow | – | – | – | – | – | – | – | – | – | – | – | – |
| IRMA | 52.525 | 79.667 | 96.301 | 0.413 | 0.402 | 0.126 | 333.667 | 5.0 | 105334.0 | 3432.0 | 80110.0 | 21066.8 |
| LAZYPIPE | 3.93 | 75.0 | 99.864 | 0.663 | 0.64 | 1.496 | 172.667 | 169.0 | 80739.0 | 298.0 | 1388.0 | 477.7 |
| metaSPAdes | 2.105 | 104.0 | 99.743 | 0.43 | 0.404 | 0.225 | 326.667 | 329.0 | 135092.0 | 206.0 | 1491.0 | 410.6 |
| metaviralSPAdes | – | – | – | – | – | – | – | – | – | – | – | – |
| PEHaplo | 9.76 | 2053.0 | 99.933 | 0.638 | 0.618 | 0.091 | 63.0 | 147.0 | 88046.0 | 376.0 | 1768.0 | 599.0 |
| QuRe | – | – | – | – | – | – | – | – | – | – | – | – |
| QVG | 16.015 | 12401.0 | 91.349 | 0.322 | 0.304 | 1.389 | 102.0 | 4.0 | 143860.0 | 5387.0 | 125041.0 | 35965.0 |
| SPAdes | – | – | – | – | – | – | – | – | – | – | – | – |
| SSAKE | 2.755 | 0.0 | 100.0 | 1.0 | 0.997 | 0.05 | 100.0 | 3.0 | 456.5 | 141.5 | 166.0 | 152.2 |
| TRACESPipe | 154.63 | 1200.0 | 95.608 | 0.468 | 0.441 | 9.768 | 366.667 | 4.0 | 143862.0 | 5387.0 | 125043.0 | 35965.5 |
| TRACESPipeLite | 36.23 | 0.0 | 99.886 | 0.989 | 0.984 | 2.786 | 367.333 | 6.0 | 168251.0 | 5387.0 | 125030.0 | 28041.8 |
| VirGenA | 246.785 | 114.667 | 99.715 | 0.602 | 0.58 | 3.823 | 600.0 | 194.333 | 66372.667 | 80.333 | 1257.0 | 342.0 |
| ViSpA | 32.7 | 248.0 | 94.797 | 0.6 | 0.59 | 5.221 | 58.667 | 69.0 | 1578880.0 | 282.0 | 61010.0 | 22882.3 |
| V-pipe | 30.975 | 5.0 | 94.181 | 0.974 | 0.965 | 0.104 | 115.0 | 5.0 | 160429.0 | 5387.0 | 125041.0 | 32085.8 |

Table S24: Results obtained for DS22 using the benchmark proposed. The execution time was measured in seconds, the RAM usage was measured in GB and the CPU usage is presented as a percentage. The executions were, when possible, capped at 6 threads and 48 GB of RAM.

| Reconstruction tool | Execution time | SNPs | Identity | NCSD | NRC | RAM usage | CPU usage | Number of scaffolds | Reconstructed bases | Minimum scaffold length | Maximum scaffold length | Average scaffold length |
|---|---|---|---|---|---|---|---|---|---|---|---|---|
| coronaSPAdes | 1.26 | 126.0 | 99.848 | 0.488 | 0.467 | 0.134 | 355.667 | 263.0 | 121202.0 | 224.0 | 2079.0 | 460.8 |
| Haploflow | – | – | – | – | – | – | – | – | – | – | – | – |
| IRMA | 55.205 | 118.0 | 96.011 | 0.429 | 0.411 | 0.125 | 334.667 | 5.0 | 103635.667 | 3378.0 | 79458.0 | 20727.133 |
| LAZYPIPE | 4.23 | 36.0 | 99.934 | 0.666 | 0.649 | 1.496 | 183.667 | 163.0 | 81213.0 | 302.0 | 1958.0 | 498.2 |
| metaSPAdes | 2.085 | 85.0 | 99.902 | 0.437 | 0.414 | 0.225 | 326.0 | 314.0 | 132017.0 | 206.0 | 2079.0 | 420.4 |
| metaviralSPAdes | – | – | – | – | – | – | – | – | – | – | – | – |
| PEHaplo | 9.715 | 1950.0 | 99.939 | 0.641 | 0.624 | 0.091 | 62.667 | 142.0 | 85791.0 | 375.0 | 2079.0 | 604.2 |
| QuRe | – | – | – | – | – | – | – | – | – | – | – | – |
| QVG | 15.49 | 15309.0 | 89.322 | 0.402 | 0.389 | 1.389 | 102.0 | 4.0 | 143860.0 | 5387.0 | 125041.0 | 35965.0 |
| SPAdes | – | – | – | – | – | – | – | – | – | – | – | – |
| SSAKE | 2.715 | 0.0 | 100.0 | 1.0 | 0.997 | 0.05 | 100.0 | 4.0 | 613.333 | 123.667 | 203.333 | 156.2 |
| TRACESPipe | 176.48 | 1294.0 | 96.259 | 0.585 | 0.559 | 9.768 | 336.0 | 4.0 | 143658.0 | 5185.0 | 125041.0 | 35914.5 |
| TRACESPipeLite | 34.65 | 0.0 | 100.0 | 1.0 | 0.998 | 2.787 | 368.333 | 6.0 | 168262.0 | 5387.0 | 125041.0 | 28043.7 |
| VirGenA | 239.175 | 99.0 | 99.833 | 0.604 | 0.584 | 3.876 | 598.0 | 208.0 | 66126.0 | 52.0 | 1445.0 | 317.9 |
| ViSpA | 26.6 | 175.0 | 95.432 | 0.685 | 0.675 | 4.889 | 58.667 | 49.0 | 962393.0 | 0.0 | 50461.0 | 19640.7 |
| V-pipe | 30.695 | 5.0 | 97.153 | 0.978 | 0.969 | 0.104 | 116.0 | 5.0 | 160429.0 | 5387.0 | 125041.0 | 32085.8 |

Table S25: Results obtained for DS23 using the benchmark proposed. The execution time was measured in seconds, the RAM usage was measured in GB and the CPU usage is presented as a percentage. The executions were, when possible, capped at 6 threads and 48 GB of RAM.

| Reconstruction tool | Execution time | SNPs | Identity | NCSD | NRC | RAM usage | CPU usage | Number of scaffolds | Reconstructed bases | Minimum scaffold length | Maximum scaffold length | Average scaffold length |
|---|---|---|---|---|---|---|---|---|---|---|---|---|
| coronaSPAdes | 1.275 | 149.0 | 99.794 | 0.47 | 0.446 | 0.134 | 356.0 | 265.0 | 124902.0 | 225.0 | 1537.0 | 471.3 |
| Haploflow | 0.63 | 1.0 | 99.84 | 0.999 | 0.996 | 0.034 | 99.0 | 1.0 | 611.0 | 611.0 | 611.0 | 611.0 |
| IRMA | 57.29 | 101.667 | 96.919 | 0.452 | 0.431 | 0.122 | 344.0 | 5.0 | 100766.667 | 3252.0 | 77451.0 | 20153.333 |
| LAZYPIPE | 3.87 | 67.0 | 99.885 | 0.646 | 0.626 | 1.495 | 176.0 | 178.0 | 87278.0 | 297.0 | 1496.0 | 490.3 |
| metaSPAdes | 2.095 | 96.0 | 99.895 | 0.43 | 0.406 | 0.224 | 327.667 | 307.0 | 133617.0 | 206.0 | 1537.0 | 435.2 |
| metaviralSPAdes | – | – | – | – | – | – | – | – | – | – | – | – |
| PEHaplo | 9.69 | 1047.0 | 99.912 | 0.662 | 0.643 | 0.091 | 62.333 | 143.0 | 86513.0 | 375.0 | 1566.0 | 605.0 |
| QuRe | – | – | – | – | – | – | – | – | – | – | – | – |
| QVG | 15.89 | 18093.0 | 87.202 | 0.504 | 0.489 | 1.389 | 102.667 | 4.0 | 143860.0 | 5387.0 | 125041.0 | 35965.0 |
| SPAdes | – | – | – | – | – | – | – | – | – | – | – | – |
| SSAKE | 2.635 | 0.0 | 100.0 | 1.0 | 0.998 | 0.049 | 100.0 | 4.667 | 621.333 | 113.333 | 149.333 | 132.967 |
| TRACESPipe | 173.885 | 1082.0 | 96.453 | 0.632 | 0.609 | 9.768 | 337.0 | 4.0 | 143814.0 | 5341.0 | 125041.0 | 35953.5 |
| TRACESPipeLite | 33.535 | 0.0 | 0.0 | 1.0 | 0.998 | 2.788 | 380.667 | 5.0 | 160429.0 | 5387.0 | 125041.0 | 32085.8 |
| VirGenA | 226.8 | 71.0 | 99.849 | 0.597 | 0.576 | 3.826 | 594.333 | 210.0 | 67346.0 | 51.0 | 971.0 | 320.7 |
| ViSpA | 21.325 | 100.0 | 95.819 | 0.813 | 0.799 | 4.838 | 53.667 | 24.0 | 216346.0 | 0.0 | 32011.0 | 9014.4 |
| V-pipe | 30.995 | 0.0 | 92.188 | 0.991 | 0.985 | 0.103 | 115.0 | 5.0 | 160429.0 | 5387.0 | 125041.0 | 32085.8 |

Table S26: Results obtained for DS24 using the benchmark proposed. The execution time was measured in seconds, the RAM usage was measured in GB and the CPU usage is presented as a percentage. The executions were, when possible, capped at 6 threads and 48 GB of RAM.

| Reconstruction tool | Execution time | SNPs | Identity | NCSD | NRC | RAM usage | CPU usage | Number of scaffolds | Reconstructed bases | Minimum scaffold length | Maximum scaffold length | Average scaffold length |
|---|---|---|---|---|---|---|---|---|---|---|---|---|
| coronaSPAdes | 1.275 | 108.0 | 99.874 | 0.467 | 0.443 | 0.134 | 358.0 | 280.0 | 124665.0 | 225.0 | 1850.0 | 445.2 |
| Haploflow | 0.64 | 1.0 | 99.84 | 0.999 | 0.997 | 0.035 | 99.333 | 1.0 | 630.0 | 630.0 | 630.0 | 630.0 |
| IRMA | 55.53 | 124.0 | 97.017 | 0.471 | 0.446 | 0.118 | 340.0 | 5.0 | 98844.667 | 3403.0 | 75189.667 | 19768.933 |
| LAZYPIPE | 3.715 | 32.0 | 99.952 | 0.593 | 0.571 | 1.496 | 153.333 | 228.0 | 96038.0 | 215.0 | 1807.0 | 421.2 |
| metaSPAdes | 2.115 | 108.0 | 99.88 | 0.435 | 0.411 | 0.224 | 328.333 | 321.0 | 133395.0 | 206.0 | 1850.0 | 415.6 |
| metaviralSPAdes | – | – | – | – | – | – | – | – | – | – | – | – |
| PEHaplo | 9.7 | 2425.0 | 99.915 | 0.638 | 0.617 | 0.091 | 62.667 | 156.0 | 89145.0 | 375.0 | 1281.0 | 571.4 |
| QuRe | – | – | – | – | – | – | – | – | – | – | – | – |
| QVG | 16.91 | 20734.0 | 85.427 | 0.632 | 0.62 | 1.389 | 102.333 | 5.0 | 268901.0 | 5387.0 | 125041.0 | 53780.2 |
| SPAdes | – | – | – | – | – | – | – | – | – | – | – | – |
| SSAKE | 2.7 | 0.0 | 100.0 | 1.0 | 0.998 | 0.049 | 100.0 | 6.0 | 977.667 | 140.667 | 205.333 | 170.367 |
| TRACESPipe | 170.865 | 1107.0 | 96.089 | 0.703 | 0.681 | 9.768 | 339.0 | 4.0 | 143852.0 | 5379.0 | 125041.0 | 35963.0 |
| TRACESPipeLite | 33.675 | 0.0 | 0.0 | 1.0 | 0.999 | 2.791 | 378.333 | 5.0 | 160227.0 | 5185.0 | 125041.0 | 32045.4 |
| VirGenA | 250.375 | 80.667 | 99.849 | 0.59 | 0.568 | 4.029 | 602.0 | 223.333 | 69013.667 | 48.333 | 1190.667 | 309.0 |
| ViSpA | 18.21 | 77.0 | 97.678 | 0.909 | 0.9 | 1.675 | 48.0 | 15.0 | 88580.0 | 0.0 | 18761.0 | 5905.3 |
| V-pipe | 30.66 | 0.0 | 96.533 | 0.993 | 0.986 | 0.103 | 115.333 | 5.0 | 160429.0 | 5387.0 | 125041.0 | 32085.8 |

Table S27: Results obtained for DS25 using the benchmark proposed. The execution time was measured in seconds, the RAM usage was measured in GB and the CPU usage is presented as a percentage. The executions were, when possible, capped at 6 threads and 48 GB of RAM.

| Reconstruction tool | Execution time | SNPs | Identity | NCSD | NRC | RAM usage | CPU usage | Number of scaffolds | Reconstructed bases | Minimum scaffold length | Maximum scaffold length | Average scaffold length |
|---|---|---|---|---|---|---|---|---|---|---|---|---|
| coronaSPAdes | 1.65 | 84.0 | 99.832 | 0.101 | 0.08 | 0.135 | 395.667 | 162.0 | 194144.0 | 229.0 | 5668.0 | 1198.4 |
| Haploflow | 1.15 | 0.0 | 100.0 | 0.973 | 0.942 | 0.052 | 99.333 | 5.0 | 4928.0 | 527.0 | 1589.0 | 985.6 |
| IRMA | 74.755 | 48.667 | 98.435 | 0.141 | 0.17 | 0.276 | 388.667 | 5.0 | 140079.0 | 4702.0 | 106678.0 | 28015.8 |
| LAZYPIPE | 4.895 | 20.0 | 99.984 | 0.177 | 0.149 | 1.502 | 217.667 | 150.0 | 178673.0 | 310.0 | 3436.0 | 1191.2 |
| metaSPAdes | 61.665 | 89.667 | 99.912 | 0.188 | 0.159 | 0.227 | 587.667 | 260.667 | 180076.667 | 71.0 | 3641.333 | 691.067 |
| metaviralSPAdes | – | – | – | – | – | – | – | – | – | – | – | – |
| PEHaplo | 11.92 | 41.0 | 99.951 | 0.127 | 0.103 | 0.172 | 83.0 | 161.0 | 193494.0 | 383.0 | 5108.0 | 1201.8 |
| QuRe | – | – | – | – | – | – | – | – | – | – | – | – |
| QVG | – | – | – | – | – | – | – | – | – | – | – | – |
| SPAdes | 6.455 | 55.0 | 99.851 | 0.1 | 0.079 | 0.151 | 239.0 | 163.0 | 194317.0 | 229.0 | 4121.0 | 1192.1 |
| SSAKE | 5.995 | 0.667 | 99.99 | 0.953 | 0.926 | 0.067 | 100.0 | 82.0 | 12700.333 | 101.333 | 266.0 | 155.033 |
| TRACESPipe | 171.67 | 7.0 | 98.92 | 0.092 | 0.073 | 9.768 | 357.0 | 4.0 | 143860.0 | 5387.0 | 125041.0 | 35965.0 |
| TRACESPipeLite | 38.145 | 2.0 | 98.858 | 0.08 | 0.062 | 2.81 | 352.333 | 6.0 | 168262.0 | 5387.0 | 125041.0 | 28043.7 |
| VirGenA | 419.745 | 41.333 | 99.914 | 0.504 | 0.472 | 4.72 | 686.0 | 110.667 | 78565.667 | 141.0 | 2993.333 | 717.867 |
| ViSpA | 74.405 | 34.0 | 96.982 | 0.159 | 0.14 | 6.814 | 74.667 | 89.0 | 3712310.0 | 339.0 | 113271.0 | 41711.3 |
| V-pipe | 34.1 | 5.0 | 95.046 | 0.64 | 0.597 | 0.104 | 115.0 | 5.0 | 160429.0 | 5387.0 | 125041.0 | 32085.8 |

Table S28: Results obtained for DS26 using the benchmark proposed. The execution time was measured in seconds, the RAM usage was measured in GB and the CPU usage is presented as a percentage. The executions were, when possible, capped at 6 threads and 48 GB of RAM.

| Reconstruction tool | Execution time | SNPs | Identity | NCSD | NRC | RAM usage | CPU usage | Number of scaffolds | Reconstructed bases | Minimum scaffold length | Maximum scaffold length | Average scaffold length |
|---|---|---|---|---|---|---|---|---|---|---|---|---|
| coronaSPAdes | 1.67 | 186.0 | 99.737 | 0.116 | 0.093 | 0.14 | 397.667 | 162.0 | 197605.0 | 239.0 | 4192.0 | 1219.8 |
| Haploflow | 1.15 | 0.0 | 0.0 | 1.0 | 0.996 | 0.053 | 99.333 | 1.0 | 520.0 | 520.0 | 520.0 | 520.0 |
| IRMA | 67.125 | 60.333 | 98.239 | 0.163 | 0.177 | 0.277 | 389.333 | 5.0 | 139295.0 | 4696.0 | 106132.0 | 27859.0 |
| LAZYPIPE | 4.97 | 56.0 | 99.847 | 0.181 | 0.155 | 1.502 | 221.333 | 154.0 | 184203.0 | 302.0 | 4152.0 | 1196.1 |
| metaSPAdes | 83.69 | 127.333 | 99.861 | 0.298 | 0.264 | 0.227 | 590.667 | 305.667 | 163554.667 | 82.0 | 2060.333 | 535.067 |
| metaviralSPAdes | – | – | – | – | – | – | – | – | – | – | – | – |
| PEHaplo | 11.835 | 336.0 | 99.73 | 0.146 | 0.122 | 0.167 | 82.667 | 162.0 | 205525.0 | 375.0 | 3959.0 | 1268.7 |
| QuRe | – | – | – | – | – | – | – | – | – | – | – | – |
| QVG | 18.86 | 3136.0 | 97.748 | 0.126 | 0.106 | 1.395 | 116.0 | 5.0 | 268901.0 | 5387.0 | 125041.0 | 53780.2 |
| SPAdes | 3.44 | 247.0 | 99.807 | 0.118 | 0.095 | 0.151 | 361.667 | 158.0 | 194868.0 | 104.0 | 4192.0 | 1233.3 |
| SSAKE | 5.965 | 0.333 | 99.996 | 0.959 | 0.946 | 0.067 | 100.0 | 75.667 | 11887.0 | 101.667 | 244.333 | 157.267 |
| TRACESPipe | 171.055 | 502.0 | 98.527 | 0.102 | 0.083 | 9.768 | 354.667 | 4.0 | 143861.0 | 5387.0 | 125042.0 | 35965.3 |
| TRACESPipeLite | 38.94 | 226.333 | 98.125 | 0.149 | 0.126 | 2.813 | 358.667 | 6.0 | 168262.0 | 5387.0 | 125041.0 | 28043.7 |
| VirGenA | 377.075 | 73.333 | 99.773 | 0.587 | 0.558 | 4.269 | 686.667 | 125.667 | 66339.0 | 64.0 | 1416.667 | 527.867 |
| ViSpA | 77.01 | 128.0 | 97.005 | 0.165 | 0.148 | 6.568 | 75.333 | 86.0 | 3891818.0 | 773.0 | 113335.0 | 45253.7 |
| V-pipe | 34.115 | 58.0 | 94.635 | 0.662 | 0.632 | 0.104 | 115.0 | 5.0 | 160429.0 | 5387.0 | 125041.0 | 32085.8 |

Table S29: Results obtained for DS27 using the benchmark proposed. The execution time was measured in seconds, the RAM usage was measured in GB and the CPU usage is presented as a percentage. The executions were, when possible, capped at 6 threads and 48 GB of RAM.

| Reconstruction tool | Execution time | SNPs | Identity | NCSD | NRC | RAM usage | CPU usage | Number of scaffolds | Reconstructed bases | Minimum scaffold length | Maximum scaffold length | Average scaffold length |
|---|---|---|---|---|---|---|---|---|---|---|---|---|
| coronaSPAdes | 1.66 | 176.0 | 99.79 | 0.099 | 0.08 | 0.138 | 398.333 | 165.0 | 202532.0 | 233.0 | 5535.0 | 1227.5 |
| Haploflow | – | – | – | – | – | – | – | – | – | – | – | – |
| IRMA | 83.155 | 46.667 | 98.697 | 0.163 | 0.174 | 0.277 | 391.0 | 5.0 | 139708.0 | 4808.0 | 106434.0 | 27941.6 |
| LAZYPIPE | 4.97 | 60.0 | 99.934 | 0.176 | 0.152 | 1.501 | 222.667 | 150.0 | 186736.0 | 310.0 | 5458.0 | 1244.9 |
| metaSPAdes | 94.425 | 112.0 | 99.837 | 0.248 | 0.22 | 0.227 | 591.667 | 315.0 | 174803.0 | 206.0 | 2200.667 | 554.933 |
| metaviralSPAdes | – | – | – | – | – | – | – | – | – | – | – | – |
| PEHaplo | 11.64 | 348.0 | 99.603 | 0.14 | 0.119 | 0.169 | 82.333 | 171.0 | 201993.0 | 387.0 | 3882.0 | 1181.2 |
| QuRe | – | – | – | – | – | – | – | – | – | – | – | – |
| QVG | 18.35 | 5172.0 | 96.4 | 0.169 | 0.149 | 1.394 | 114.0 | 4.0 | 143860.0 | 5387.0 | 125041.0 | 35965.0 |
| SPAdes | 4.46 | 188.0 | 99.772 | 0.102 | 0.083 | 0.152 | 306.0 | 160.0 | 199571.0 | 104.0 | 5535.0 | 1247.3 |
| SSAKE | 6.13 | 0.333 | 99.996 | 0.963 | 0.952 | 0.068 | 100.0 | 70.0 | 11021.0 | 104.0 | 286.667 | 157.333 |
| TRACESPipe | 171.06 | 670.0 | 98.687 | 0.088 | 0.072 | 9.768 | 360.667 | 4.0 | 143861.0 | 5387.0 | 125042.0 | 35965.3 |
| TRACESPipeLite | 36.805 | 171.333 | 97.728 | 0.501 | 0.471 | 2.815 | 361.333 | 6.0 | 168262.0 | 5387.0 | 125041.0 | 28043.7 |
| VirGenA | 401.61 | 153.0 | 99.605 | 0.527 | 0.504 | 4.774 | 684.333 | 114.0 | 76231.667 | 93.0 | 2432.667 | 674.5 |
| ViSpA | 63.97 | 620.0 | 96.542 | 0.171 | 0.154 | 6.704 | 75.0 | 78.0 | 3366127.0 | 1456.0 | 113769.0 | 43155.5 |
| V-pipe | 34.535 | 61.0 | 95.529 | 0.67 | 0.642 | 0.104 | 115.0 | 5.0 | 160429.0 | 5387.0 | 125041.0 | 32085.8 |

Table S30: Results obtained for DS28 using the benchmark proposed. The execution time was measured in seconds, the RAM usage was measured in GB and the CPU usage is presented as a percentage. The executions were, when possible, capped at 6 threads and 48 GB of RAM.

| Reconstruction tool | Execution time | SNPs | Identity | NCSD | NRC | RAM usage | CPU usage | Number of scaffolds | Reconstructed bases | Minimum scaffold length | Maximum scaffold length | Average scaffold length |
|---|---|---|---|---|---|---|---|---|---|---|---|---|
| coronaSPAdes | 1.64 | 97.0 | 99.84 | 0.109 | 0.088 | 0.138 | 399.0 | 156.0 | 201020.0 | 225.0 | 4774.0 | 1288.6 |
| Haploflow | – | – | – | – | – | – | – | – | – | – | – | – |
| IRMA | 101.42 | 66.0 | 98.533 | 0.178 | 0.182 | 0.272 | 392.333 | 5.0 | 138542.0 | 4401.0 | 105504.0 | 27708.4 |
| LAZYPIPE | 4.97 | 24.0 | 99.94 | 0.171 | 0.146 | 1.502 | 218.0 | 149.0 | 187973.0 | 303.0 | 4720.0 | 1261.6 |
| metaSPAdes | 95.205 | 105.0 | 99.9 | 0.317 | 0.288 | 0.227 | 592.333 | 329.333 | 160981.0 | 206.0 | 2418.333 | 489.133 |
| metaviralSPAdes | – | – | – | – | – | – | – | – | – | – | – | – |
| PEHaplo | 12.045 | 323.0 | 99.525 | 0.137 | 0.117 | 0.17 | 118.667 | 166.0 | 221032.0 | 378.0 | 5395.0 | 1331.5 |
| QuRe | – | – | – | – | – | – | – | – | – | – | – | – |
| QVG | 19.625 | 7602.0 | 94.671 | 0.233 | 0.215 | 1.394 | 117.0 | 5.0 | 268901.0 | 5387.0 | 125041.0 | 53780.2 |
| SPAdes | 3.205 | 61.0 | 99.85 | 0.108 | 0.087 | 0.151 | 380.0 | 156.0 | 201005.0 | 230.0 | 4774.0 | 1288.5 |
| SSAKE | 6.0 | 0.333 | 99.996 | 0.963 | 0.954 | 0.068 | 100.0 | 68.667 | 10430.667 | 100.667 | 254.667 | 151.9 |
| TRACESPipe | 178.92 | 1322.0 | 98.045 | 0.116 | 0.097 | 9.768 | 344.667 | 4.0 | 143858.0 | 5387.0 | 125039.0 | 35964.5 |
| TRACESPipeLite | 37.02 | 7.0 | 98.161 | 0.9 | 0.886 | 2.816 | 362.333 | 6.0 | 168262.0 | 5387.0 | 125041.0 | 28043.7 |
| VirGenA | 384.205 | 126.0 | 99.474 | 0.561 | 0.534 | 4.382 | 679.667 | 126.0 | 71738.0 | 62.0 | 1745.0 | 569.3 |
| ViSpA | 62.585 | 911.0 | 96.58 | 0.189 | 0.173 | 7.765 | 78.667 | 90.0 | 3324538.0 | 872.0 | 112463.0 | 36939.3 |
| V-pipe | 34.81 | 182.0 | 94.928 | 0.678 | 0.648 | 0.104 | 115.0 | 5.0 | 160429.0 | 5387.0 | 125041.0 | 32085.8 |

Table S31: Results obtained for DS29 using the benchmark proposed. The execution time was measured in seconds, the RAM usage was measured in GB and the CPU usage is presented as a percentage. The executions were, when possible, capped at 6 threads and 48 GB of RAM.

| Reconstruction tool | Execution time | SNPs | Identity | NCSD | NRC | RAM usage | CPU usage | Number of scaffolds | Reconstructed bases | Minimum scaffold length | Maximum scaffold length | Average scaffold length |
|---|---|---|---|---|---|---|---|---|---|---|---|---|
| coronaSPAdes | 1.65 | 88.0 | 99.934 | 0.124 | 0.106 | 0.138 | 393.0 | 161.0 | 198559.0 | 231.0 | 6121.0 | 1233.3 |
| Haploflow | 1.15 | 0.0 | 0.0 | 1.0 | 0.998 | 0.053 | 99.667 | 1.0 | 511.0 | 511.0 | 511.0 | 511.0 |
| IRMA | 96.65 | 47.333 | 98.492 | 0.183 | 0.182 | 0.276 | 399.0 | 5.0 | 138702.0 | 4592.0 | 105718.0 | 27740.4 |
| LAZYPIPE | 4.195 | 17.0 | 99.987 | 0.169 | 0.148 | 1.502 | 194.667 | 160.0 | 188434.0 | 239.0 | 6018.0 | 1177.7 |
| metaSPAdes | 82.22 | 88.667 | 99.9 | 0.356 | 0.333 | 0.227 | 591.0 | 320.333 | 153235.333 | 207.0 | 2223.667 | 478.333 |
| metaviralSPAdes | – | – | – | – | – | – | – | – | – | – | – | – |
| PEHaplo | 11.695 | 212.0 | 99.763 | 0.157 | 0.137 | 0.169 | 82.0 | 170.0 | 197190.0 | 379.0 | 6615.0 | 1159.9 |
| QuRe | – | – | – | – | – | – | – | – | – | – | – | – |
| QVG | 21.93 | 9748.0 | 93.194 | 0.285 | 0.269 | 1.395 | 117.333 | 7.0 | 282124.0 | 5387.0 | 125041.0 | 40303.4 |
| SPAdes | 3.05 | 45.0 | 99.965 | 0.118 | 0.1 | 0.151 | 394.0 | 163.0 | 199677.0 | 231.0 | 6121.0 | 1225.0 |
| SSAKE | 6.12 | 0.0 | 100.0 | 0.96 | 0.951 | 0.068 | 100.0 | 75.667 | 11525.0 | 102.667 | 220.667 | 152.333 |
| TRACESPipe | 182.195 | 1149.0 | 97.348 | 0.177 | 0.156 | 9.768 | 345.333 | 4.0 | 143860.0 | 5387.0 | 125041.0 | 35965.0 |
| TRACESPipeLite | 37.595 | 0.0 | 98.534 | 0.99 | 0.985 | 2.818 | 362.333 | 6.0 | 168262.0 | 5387.0 | 125041.0 | 28043.7 |
| VirGenA | 379.705 | 94.0 | 99.752 | 0.615 | 0.593 | 4.223 | 685.667 | 121.0 | 63587.0 | 92.0 | 2157.0 | 525.5 |
| ViSpA | 80.93 | 807.0 | 95.473 | 0.248 | 0.236 | 6.841 | 75.333 | 107.0 | 4543976.0 | 804.0 | 105356.0 | 42467.1 |
| V-pipe | 35.18 | 187.0 | 94.365 | 0.684 | 0.66 | 0.104 | 114.333 | 5.0 | 160429.0 | 5387.0 | 125041.0 | 32085.8 |

Table S32: Results obtained for DS30 using the benchmark proposed. The execution time was measured in seconds, the RAM usage was measured in GB and the CPU usage is presented as a percentage. The executions were, when possible, capped at 6 threads and 48 GB of RAM.

| Reconstruction tool | Execution time | SNPs | Identity | NCSD | NRC | RAM usage | CPU usage | Number of scaffolds | Reconstructed bases | Minimum scaffold length | Maximum scaffold length | Average scaffold length |
|---|---|---|---|---|---|---|---|---|---|---|---|---|
| coronaSPAdes | 1.64 | 72.0 | 99.946 | 0.115 | 0.098 | 0.137 | 396.0 | 167.0 | 198003.0 | 231.0 | 4439.0 | 1185.6 |
| Haploflow | 1.14 | 0.0 | 100.0 | 0.999 | 0.997 | 0.053 | 99.0 | 1.0 | 540.0 | 540.0 | 540.0 | 540.0 |
| IRMA | 92.51 | 41.333 | 98.577 | 0.186 | 0.176 | 0.278 | 406.667 | 5.0 | 139538.0 | 4547.0 | 106270.0 | 27907.6 |
| LAZYPIPE | 4.425 | 24.0 | 99.981 | 0.176 | 0.156 | 1.502 | 206.0 | 155.0 | 183551.0 | 300.0 | 4357.0 | 1184.2 |
| metaSPAdes | 85.915 | 87.5 | 99.92 | 0.264 | 0.241 | 0.227 | 591.0 | 314.5 | 170446.0 | 206.0 | 2317.0 | 541.95 |
| metaviralSPAdes | – | – | – | – | – | – | – | – | – | – | – | – |
| PEHaplo | 11.7 | 36.0 | 99.943 | 0.14 | 0.122 | 0.156 | 79.0 | 168.0 | 200107.0 | 381.0 | 5666.0 | 1191.1 |
| QuRe | – | – | – | – | – | – | – | – | – | – | – | – |
| QVG | 22.13 | 12704.0 | 91.129 | 0.371 | 0.36 | 1.395 | 115.667 | 7.0 | 282124.0 | 5387.0 | 125041.0 | 40303.4 |
| SPAdes | 3.115 | 89.0 | 99.931 | 0.113 | 0.096 | 0.151 | 385.0 | 167.0 | 198125.0 | 231.0 | 4439.0 | 1186.4 |
| SSAKE | 6.065 | 1.333 | 99.983 | 0.965 | 0.956 | 0.069 | 100.0 | 77.0 | 12252.667 | 105.667 | 297.0 | 159.1 |
| TRACESPipe | 165.765 | 1352.0 | 97.63 | 0.203 | 0.184 | 9.768 | 351.0 | 4.0 | 143859.0 | 5386.0 | 125041.0 | 35964.8 |
| TRACESPipeLite | 36.75 | 0.0 | 0.0 | 1.001 | 0.998 | 2.818 | 365.0 | 6.0 | 168262.0 | 5387.0 | 125041.0 | 28043.7 |
| VirGenA | 389.84 | 32.0 | 99.755 | 0.62 | 0.602 | 4.315 | 684.667 | 112.0 | 62075.0 | 52.0 | 2320.0 | 554.2 |
| ViSpA | 84.435 | 688.0 | 94.351 | 0.342 | 0.332 | 6.119 | 70.667 | 133.0 | 5417908.0 | 500.0 | 94119.0 | 40736.2 |
| V-pipe | 34.785 | 173.0 | 94.003 | 0.728 | 0.706 | 0.104 | 114.0 | 5.0 | 160429.0 | 5387.0 | 125041.0 | 32085.8 |

Table S33: Results obtained for DS31 using the benchmark proposed. The execution time was measured in seconds, the RAM usage was measured in GB and the CPU usage is presented as a percentage. The executions were, when possible, capped at 6 threads and 48 GB of RAM.

| **Reconstruction tool** | **Execution time** | **SNPs** | **Identity** | **NCSD** | **NRC** | **RAM usage** | **CPU usage** | **Number of scaffolds** | **Reconstructed bases** | **Minimum scaffold length** | **Maximum scaffold length** | **Average scaffold length** |
|---|---|---|---|---|---|---|---|---|---|---|---|---|
| coronaSPAdes | 1.66 | 75.0 | 99.943 | 0.107 | 0.09 | 0.138 | 399.0 | 171.0 | 200252.0 | 230.0 | 6031.0 | 1171.1 |
| Haploflow | 1.15 | 0.0 | 100.0 | 0.998 | 0.997 | 0.053 | 99.0 | 1.0 | 533.0 | 533.0 | 533.0 | 533.0 |
| IRMA | 99.68 | 43.0 | 98.459 | 0.202 | 0.188 | 0.274 | 407.667 | 5.0 | 137704.0 | 4718.0 | 105011.0 | 27540.8 |
| LAZYPIPE | 4.2 | 20.0 | 99.984 | 0.173 | 0.153 | 1.501 | 195.333 | 164.0 | 186700.0 | 249.0 | 4643.0 | 1138.4 |
| metaSPAdes | 90.38 | 90.667 | 99.906 | 0.359 | 0.335 | 0.227 | 592.333 | 325.667 | 152286.0 | 206.0 | 1763.667 | 467.633 |
| metaviralSPAdes | – | – | – | – | – | – | – | – | – | – | – | – |
| PEHaplo | 11.62 | 364.0 | 99.956 | 0.147 | 0.128 | 0.166 | 82.333 | 163.0 | 194113.0 | 377.0 | 4461.0 | 1190.9 |
| QuRe | – | – | – | – | – | – | – | – | – | – | – | – |
| QVG | 20.56 | 16044.0 | 88.634 | 0.489 | 0.476 | 1.395 | 115.667 | 6.0 | 274501.0 | 5387.0 | 125043.0 | 45750.2 |
| SPAdes | 4.28 | 44.0 | 99.966 | 0.105 | 0.088 | 0.151 | 312.667 | 171.0 | 200309.0 | 230.0 | 6031.0 | 1171.4 |
| SSAKE | 6.145 | 0.0 | 100.0 | 0.969 | 0.959 | 0.068 | 100.0 | 62.667 | 10026.333 | 106.667 | 235.333 | 159.933 |
| TRACESPipe | 186.325 | 1812.0 | 97.055 | 0.237 | 0.218 | 9.768 | 333.333 | 4.0 | 143860.0 | 5387.0 | 125041.0 | 35965.0 |
| TRACESPipeLite | 37.58 | 0.0 | 0.0 | 1.0 | 0.998 | 2.819 | 362.333 | 6.0 | 168262.0 | 5387.0 | 125041.0 | 28043.7 |
| VirGenA | 390.005 | 34.0 | 99.907 | 0.591 | 0.572 | 4.396 | 678.333 | 112.333 | 66470.333 | 97.333 | 1887.0 | 594.333 |
| ViSpA | 50.215 | 292.667 | 94.943 | 0.526 | 0.515 | 7.026 | 71.333 | 107.0 | 3041834.0 | 450.0 | 71401.0 | 28428.4 |
| V-pipe | 34.28 | 162.0 | 93.088 | 0.792 | 0.771 | 0.104 | 114.667 | 5.0 | 160429.0 | 5387.0 | 125041.0 | 32085.8 |

Table S34: Results obtained for DS32 using the benchmark proposed. The execution time was measured in seconds, the RAM usage was measured in GB and the CPU usage is presented as a percentage. The executions were, when possible, capped at 6 threads and 48 GB of RAM.

| **Reconstruction tool** | **Execution time** | **SNPs** | **Identity** | **NCSD** | **NRC** | **RAM usage** | **CPU usage** | **Number of scaffolds** | **Reconstructed bases** | **Minimum scaffold length** | **Maximum scaffold length** | **Average scaffold length** |
|---|---|---|---|---|---|---|---|---|---|---|---|---|
| coronaSPAdes | 1.665 | 66.0 | 99.951 | 0.099 | 0.082 | 0.137 | 394.0 | 173.0 | 201373.0 | 230.0 | 4977.0 | 1164.0 |
| Haploflow | – | – | – | – | – | – | – | – | – | – | – | – |
| IRMA | 128.915 | 63.667 | 98.527 | 0.21 | 0.191 | 0.281 | 413.333 | 5.0 | 138139.333 | 4874.0 | 105461.333 | 27627.867 |
| LAZYPIPE | 3.975 | 19.0 | 99.986 | 0.151 | 0.133 | 1.502 | 180.667 | 172.0 | 189101.0 | 239.0 | 4894.0 | 1099.4 |
| metaSPAdes | 86.175 | 79.333 | 99.932 | 0.237 | 0.214 | 0.227 | 591.333 | 313.667 | 173362.0 | 206.0 | 2234.0 | 552.967 |
| metaviralSPAdes | – | – | – | – | – | – | – | – | – | – | – | – |
| PEHaplo | 11.685 | 34.0 | 99.963 | 0.128 | 0.11 | 0.156 | 82.0 | 163.0 | 197234.0 | 379.0 | 4460.0 | 1210.0 |
| QuRe | – | – | – | – | – | – | – | – | – | – | – | – |
| QVG | 21.015 | 19218.0 | 86.483 | 0.622 | 0.612 | 1.394 | 117.0 | 7.0 | 282329.0 | 5387.0 | 125039.0 | 40332.7 |
| SPAdes | 3.12 | 44.0 | 99.967 | 0.096 | 0.08 | 0.151 | 384.0 | 173.0 | 201591.0 | 230.0 | 4977.0 | 1165.3 |
| SSAKE | 6.015 | 0.0 | 100.0 | 0.971 | 0.96 | 0.068 | 100.0 | 73.0 | 11525.333 | 102.0 | 296.0 | 157.667 |
| TRACESPipe | 183.67 | 1406.0 | 97.524 | 0.303 | 0.283 | 9.768 | 339.0 | 4.0 | 143814.0 | 5341.0 | 125041.0 | 35953.5 |
| TRACESPipeLite | 37.17 | 0.0 | 0.0 | 1.0 | 0.999 | 2.82 | 364.0 | 6.0 | 168289.0 | 5387.0 | 125041.0 | 28048.2 |
| VirGenA | 400.325 | 56.667 | 99.747 | 0.604 | 0.584 | 4.473 | 686.0 | 124.333 | 65622.667 | 85.667 | 1602.333 | 533.633 |
| ViSpA | 28.805 | 248.667 | 95.949 | 0.74 | 0.729 | 6.256 | 67.333 | 54.0 | 1043140.0 | 404.0 | 44347.0 | 19317.4 |
| V-pipe | 34.075 | 72.0 | 93.219 | 0.856 | 0.837 | 0.104 | 114.667 | 5.0 | 160429.0 | 5387.0 | 125041.0 | 32085.8 |

Table S35: Results obtained for DS33 using the benchmark proposed. The execution time was measured in seconds, the RAM usage was measured in GB and the CPU usage is presented as a percentage. The executions were, when possible, capped at 6 threads and 48 GB of RAM.

| **Reconstruction tool** | **Execution time** | **SNPs** | **Identity** | **NCSD** | **NRC** | **RAM usage** | **CPU usage** | **Number of scaffolds** | **Reconstructed bases** | **Minimum scaffold length** | **Maximum scaffold length** | **Average scaffold length** |
|---|---|---|---|---|---|---|---|---|---|---|---|---|
| coronaSPAdes | 2.125 | 10.0 | 99.908 | 0.035 | 0.026 | 0.138 | 411.333 | 27.0 | 202192.0 | 165.0 | 41011.0 | 7488.6 |
| Haploflow | 2.04 | 0.0 | 100.0 | 0.621 | 0.576 | 0.069 | 99.333 | 106.0 | 84303.0 | 504.0 | 2747.0 | 795.3 |
| IRMA | 95.06 | 7.333 | 99.952 | 0.092 | 0.131 | 0.488 | 456.667 | 5.0 | 144638.0 | 4880.0 | 110164.0 | 28927.6 |
| LAZYPIPE | 5.47 | 4.0 | 99.995 | 0.041 | 0.033 | 1.509 | 248.333 | 24.0 | 201014.0 | 302.0 | 40981.0 | 8375.6 |
| metaSPAdes | 3.74 | 8.0 | 99.858 | 0.034 | 0.025 | 0.227 | 395.0 | 27.0 | 202264.0 | 71.0 | 41011.0 | 7491.3 |
| metaviralSPAdes | – | – | – | – | – | – | – | – | – | – | – | – |
| PEHaplo | 16.93 | 16.0 | 99.983 | 0.053 | 0.044 | 0.428 | 102.0 | 208.0 | 226110.0 | 379.0 | 5296.0 | 1087.1 |
| QuRe | 720.04 | 0.0 | 100.0 | 1.0 | 0.995 | 55.254 | 611.5 | 3.0 | 151.0 | 40.0 | 71.0 | 50.3 |
| QVG | – | – | – | – | – | – | – | – | – | – | – | – |
| SPAdes | 12.25 | 5.0 | 99.863 | 0.035 | 0.026 | 0.158 | 206.667 | 26.0 | 202307.0 | 165.0 | 41011.0 | 7781.0 |
| SSAKE | 1.665 | 1.0 | 99.994 | 0.916 | 0.889 | 0.049 | 100.0 | 115.667 | 21545.333 | 100.333 | 473.667 | 186.267 |
| TRACESPipe | 165.305 | 0.0 | 99.957 | 0.031 | 0.023 | 9.768 | 364.667 | 4.0 | 143860.0 | 5387.0 | 125041.0 | 35965.0 |
| TRACESPipeLite | 40.27 | 0.0 | 99.909 | 0.031 | 0.023 | 2.816 | 339.0 | 6.0 | 168262.0 | 5387.0 | 125041.0 | 28043.7 |
| VirGenA | 586.465 | 21.0 | 99.894 | 0.563 | 0.533 | 4.498 | 734.333 | 85.0 | 69043.0 | 226.0 | 2345.0 | 812.3 |
| ViSpA | 40.11 | 4.0 | 99.635 | 0.042 | 0.033 | 5.94 | 91.0 | 11.0 | 454253.0 | 2029.0 | 124437.0 | 41295.7 |
| V-pipe | 38.28 | 5.0 | 96.583 | 0.223 | 0.194 | 0.104 | 114.0 | 5.0 | 160429.0 | 5387.0 | 125041.0 | 32085.8 |

Table S36: Results obtained for DS34 using the benchmark proposed. The execution time was measured in seconds, the RAM usage was measured in GB and the CPU usage is presented as a percentage. The executions were, when possible, capped at 6 threads and 48 GB of RAM.

| Reconstruction tool | Execution time | SNPs | Identity | NCSD | NRC | RAM usage | CPU usage | Number of scaffolds | Reconstructed bases | Minimum scaffold length | Maximum scaffold length | Average scaffold length |
|---|---|---|---|---|---|---|---|---|---|---|---|---|
| coronaSPAdes | 2.19 | 55.0 | 99.96 | 0.04 | 0.028 | 0.154 | 409.0 | 33.0 | 207952.0 | 242.0 | 26757.0 | 6301.6 |
| Haploflow | 2.07 | 44.0 | 99.921 | 0.641 | 0.612 | 0.07 | 99.667 | 103.0 | 80047.0 | 502.0 | 2199.0 | 777.2 |
| IRMA | 110.605 | 3.0 | 99.946 | 0.106 | 0.13 | 0.484 | 457.333 | 5.0 | 144544.0 | 4878.0 | 110152.0 | 28908.8 |
| LAZYPIPE | 5.59 | 148.0 | 99.856 | 0.043 | 0.032 | 1.51 | 252.0 | 32.0 | 208755.0 | 328.0 | 26713.0 | 6523.6 |
| metaSPAdes | 3.815 | 40.0 | 99.924 | 0.04 | 0.029 | 0.227 | 397.667 | 39.0 | 208108.0 | 69.0 | 26757.0 | 5336.1 |
| metaviralSPAdes | – | – | – | – | – | – | – | – | – | – | – | – |
| PEHaplo | 17.035 | 232.0 | 99.753 | 0.059 | 0.047 | 0.418 | 107.0 | 240.0 | 243109.0 | 380.0 | 4924.0 | 1013.0 |
| QuRe | 619.59 | 0.0 | 100.0 | 1.0 | 0.995 | 52.658 | 577.333 | 1.0 | 94.0 | 94.0 | 94.0 | 94.0 |
| QVG | 21.43 | 1388.0 | 98.963 | 0.077 | 0.061 | 1.399 | 133.667 | 5.0 | 268901.0 | 5387.0 | 125041.0 | 53780.2 |
| SPAdes | 7.28 | 20.0 | 99.918 | 0.037 | 0.026 | 0.159 | 277.0 | 34.0 | 209307.0 | 85.0 | 26757.0 | 6156.1 |
| SSAKE | 1.645 | 0.333 | 99.998 | 0.932 | 0.919 | 0.049 | 100.0 | 118.0 | 20774.0 | 102.333 | 412.0 | 176.033 |
| TRACESPipe | 168.35 | 511.0 | 99.591 | 0.042 | 0.032 | 9.768 | 363.0 | 4.0 | 143862.0 | 5387.0 | 125043.0 | 35965.5 |
| TRACESPipeLite | 42.105 | 457.667 | 99.12 | 0.075 | 0.061 | 2.818 | 345.333 | 6.0 | 168262.0 | 5387.0 | 125041.0 | 28043.7 |
| VirGenA | 561.335 | 80.0 | 99.75 | 0.627 | 0.6 | 4.497 | 739.0 | 86.0 | 60061.0 | 57.0 | 2560.0 | 698.4 |
| ViSpA | 39.015 | 470.0 | 99.477 | 0.054 | 0.043 | 6.074 | 94.667 | 9.0 | 314557.0 | 2166.333 | 173078.333 | 40034.533 |
| V-pipe | 38.545 | 297.0 | 96.547 | 0.257 | 0.225 | 0.104 | 114.0 | 5.0 | 160429.0 | 5387.0 | 125041.0 | 32085.8 |

Table S37: Results obtained for DS35 using the benchmark proposed. The execution time was measured in seconds, the RAM usage was measured in GB and the CPU usage is presented as a percentage. The executions were, when possible, capped at 6 threads and 48 GB of RAM.

| Reconstruction tool | Execution time | SNPs | Identity | NCSD | NRC | RAM usage | CPU usage | Number of scaffolds | Reconstructed bases | Minimum scaffold length | Maximum scaffold length | Average scaffold length |
|---|---|---|---|---|---|---|---|---|---|---|---|---|
| coronaSPAdes | 2.1 | 28.0 | 99.976 | 0.038 | 0.028 | 0.136 | 414.667 | 25.0 | 209914.0 | 298.0 | 34355.0 | 8396.6 |
| Haploflow | 2.08 | 34.0 | 99.907 | 0.672 | 0.642 | 0.07 | 99.333 | 101.0 | 76112.0 | 502.0 | 1781.0 | 753.6 |
| IRMA | 144.775 | 6.667 | 99.851 | 0.114 | 0.133 | 0.478 | 459.0 | 5.0 | 144500.0 | 4887.0 | 110002.0 | 28900.0 |
| LAZYPIPE | 4.98 | 9.0 | 99.876 | 0.045 | 0.035 | 1.509 | 238.667 | 27.0 | 208600.0 | 363.0 | 26610.0 | 7725.9 |
| metaSPAdes | 3.845 | 29.0 | 99.974 | 0.039 | 0.029 | 0.227 | 398.667 | 30.0 | 209653.0 | 184.0 | 34355.0 | 6988.4 |
| metaviralSPAdes | – | – | – | – | – | – | – | – | – | – | – | – |
| PEHaplo | 16.905 | 114.0 | 99.787 | 0.067 | 0.053 | 0.419 | 102.0 | 221.0 | 235361.0 | 377.0 | 3273.0 | 1065.0 |
| QuRe | – | – | – | – | – | – | – | – | – | – | – | – |
| QVG | 23.075 | 2184.0 | 98.482 | 0.098 | 0.083 | 1.399 | 133.333 | 5.0 | 268901.0 | 5387.0 | 125041.0 | 53780.2 |
| SPAdes | 7.905 | 2.0 | 99.508 | 0.038 | 0.028 | 0.159 | 267.667 | 28.0 | 209843.0 | 83.0 | 34355.0 | 7494.4 |
| SSAKE | 1.7 | 0.333 | 99.998 | 0.924 | 0.908 | 0.049 | 100.0 | 116.667 | 21313.0 | 102.0 | 505.333 | 182.667 |
| TRACESPipe | 171.06 | 665.0 | 99.454 | 0.047 | 0.037 | 9.768 | 349.333 | 4.0 | 143860.0 | 5387.0 | 125042.0 | 35965.0 |
| TRACESPipeLite | 41.075 | 249.0 | 97.658 | 0.393 | 0.363 | 2.82 | 347.0 | 6.0 | 168262.0 | 5387.0 | 125041.0 | 28043.7 |
| VirGenA | 523.695 | 140.667 | 99.63 | 0.576 | 0.55 | 4.807 | 733.0 | 79.333 | 66812.333 | 149.0 | 2257.667 | 877.0 |
| ViSpA | 40.73 | 647.0 | 99.377 | 0.058 | 0.047 | 5.922 | 92.0 | 11.0 | 506252.0 | 5265.0 | 124298.0 | 46022.9 |
| V-pipe | 38.685 | 271.0 | 96.769 | 0.254 | 0.228 | 0.104 | 114.0 | 5.0 | 160429.0 | 5387.0 | 125041.0 | 32085.8 |

Table S38: Results obtained for DS36 using the benchmark proposed. The execution time was measured in seconds, the RAM usage was measured in GB and the CPU usage is presented as a percentage. The executions were, when possible, capped at 6 threads and 48 GB of RAM.

| Reconstruction tool | Execution time | SNPs | Identity | NCSD | NRC | RAM usage | CPU usage | Number of scaffolds | Reconstructed bases | Minimum scaffold length | Maximum scaffold length | Average scaffold length |
|---|---|---|---|---|---|---|---|---|---|---|---|---|
| coronaSPAdes | 2.12 | 19.0 | 99.986 | 0.034 | 0.024 | 0.135 | 414.667 | 16.0 | 210030.0 | 481.0 | 76427.0 | 13126.9 |
| Haploflow | 2.08 | 21.0 | 99.886 | 0.657 | 0.634 | 0.07 | 99.333 | 109.0 | 82555.0 | 501.0 | 2041.0 | 757.4 |
| IRMA | 148.69 | 4.667 | 99.975 | 0.117 | 0.13 | 0.487 | 468.667 | 5.0 | 144632.0 | 4794.0 | 110222.0 | 28926.4 |
| LAZYPIPE | 4.81 | 8.0 | 99.953 | 0.036 | 0.027 | 1.51 | 228.333 | 17.0 | 209331.0 | 447.0 | 76392.0 | 12313.6 |
| metaSPAdes | 3.855 | 16.0 | 99.986 | 0.033 | 0.024 | 0.227 | 394.333 | 17.0 | 210105.0 | 481.0 | 76427.0 | 12359.1 |
| metaviralSPAdes | – | – | – | – | – | – | – | – | – | – | – | – |
| PEHaplo | 16.81 | 157.0 | 99.806 | 0.064 | 0.053 | 0.419 | 102.0 | 200.0 | 229677.0 | 379.0 | 4327.0 | 1148.4 |
| QuRe | 602.145 | 0.0 | 100.0 | 1.0 | 0.996 | 52.768 | 621.667 | 2.0 | 117.0 | 47.0 | 70.0 | 58.5 |
| QVG | 24.935 | 3358.0 | 97.624 | 0.13 | 0.117 | 1.398 | 134.0 | 6.0 | 274288.0 | 5387.0 | 125041.0 | 45714.7 |
| SPAdes | 6.585 | 9.0 | 99.989 | 0.033 | 0.024 | 0.159 | 301.0 | 17.0 | 210126.0 | 481.0 | 76427.0 | 12360.4 |
| SSAKE | 1.71 | 0.333 | 99.998 | 0.921 | 0.908 | 0.049 | 100.0 | 116.0 | 22004.333 | 102.333 | 464.667 | 189.6 |
| TRACESPipe | 172.975 | 793.0 | 99.225 | 0.06 | 0.049 | 9.768 | 356.333 | 4.0 | 143869.0 | 5387.0 | 125050.0 | 35967.3 |
| TRACESPipeLite | 39.28 | 65.667 | 98.668 | 0.841 | 0.823 | 2.821 | 352.0 | 6.0 | 168262.0 | 5387.0 | 125041.0 | 28043.7 |
| VirGenA | 560.25 | 164.0 | 99.644 | 0.597 | 0.575 | 4.641 | 736.333 | 81.667 | 64253.667 | 144.667 | 2783.333 | 804.9 |
| ViSpA | 37.865 | 632.0 | 98.667 | 0.069 | 0.058 | 6.166 | 92.333 | 15.0 | 457100.0 | 4009.0 | 124187.0 | 30473.3 |
| V-pipe | 38.605 | 424.0 | 95.936 | 0.28 | 0.254 | 0.104 | 114.333 | 5.0 | 160429.0 | 5387.0 | 125041.0 | 32085.8 |

Table S39: Results obtained for DS37 using the benchmark proposed. The execution time was measured in seconds, the RAM usage was measured in GB and the CPU usage is presented as a percentage. The executions were, when possible, capped at 6 threads and 48 GB of RAM.

| Reconstruction tool | Execution time | SNPs | Identity | NCSD | NRC | RAM usage | CPU usage | Number of scaffolds | Reconstructed bases | Minimum scaffold length | Maximum scaffold length | Average scaffold length |
|---|---|---|---|---|---|---|---|---|---|---|---|---|
| coronaSPAdes | 2.11 | 12.0 | 99.99 | 0.033 | 0.025 | 0.134 | 415.333 | 24.0 | 210176.0 | 560.0 | 29993.0 | 8757.3 |
| Haploflow | 2.08 | 134.0 | 99.741 | 0.673 | 0.652 | 0.07 | 99.0 | 98.0 | 78659.0 | 501.0 | 2497.0 | 802.6 |
| IRMA | 151.255 | 12.333 | 99.925 | 0.119 | 0.127 | 0.482 | 476.333 | 5.0 | 145406.333 | 4867.0 | 110886.333 | 29081.267 |
| LAZYPIPE | 4.565 | 4.0 | 99.995 | 0.041 | 0.032 | 1.509 | 204.667 | 26.0 | 209151.0 | 526.0 | 29973.0 | 8044.3 |
| metaSPAdes | 3.765 | 7.0 | 99.994 | 0.032 | 0.024 | 0.227 | 393.667 | 24.0 | 210211.0 | 560.0 | 29993.0 | 8758.8 |
| metaviralSPAdes | – | – | – | – | – | – | – | – | – | – | – | – |
| PEHaplo | 16.755 | 109.0 | 99.925 | 0.063 | 0.051 | 0.418 | 101.667 | 202.0 | 230677.0 | 380.0 | 5122.0 | 1142.0 |
| QuRe | 168.765 | 0.0 | 100.0 | 1.0 | 0.997 | 12.763 | 629.0 | 3.0 | 334.0 | 98.0 | 118.0 | 111.3 |
| QVG | 26.29 | 4998.0 | 96.505 | 0.171 | 0.163 | 1.399 | 133.0 | 6.0 | 274497.0 | 5387.0 | 125041.0 | 45749.5 |
| SPAdes | 5.305 | 9.0 | 99.991 | 0.032 | 0.024 | 0.159 | 342.333 | 24.0 | 210211.0 | 560.0 | 29993.0 | 8758.8 |
| SSAKE | 1.82 | 1.333 | 99.991 | 0.927 | 0.913 | 0.05 | 100.0 | 114.333 | 21220.0 | 101.667 | 363.667 | 185.667 |
| TRACESPipe | 179.625 | 530.0 | 99.5 | 0.073 | 0.061 | 9.768 | 354.0 | 4.0 | 143860.0 | 5387.0 | 125041.0 | 35965.0 |
| TRACESPipeLite | 38.6 | 1.0 | 99.976 | 0.983 | 0.975 | 2.824 | 354.667 | 6.0 | 168262.0 | 5387.0 | 125041.0 | 28043.7 |
| VirGenA | 559.23 | 66.0 | 99.727 | 0.602 | 0.583 | 4.454 | 736.667 | 77.0 | 63709.333 | 229.667 | 2382.0 | 834.2 |
| ViSpA | 58.445 | 1051.333 | 97.847 | 0.097 | 0.085 | 5.949 | 91.0 | 41.0 | 2128909.0 | 1839.0 | 121179.0 | 51924.6 |
| V-pipe | 39.24 | 689.0 | 95.647 | 0.291 | 0.267 | 0.104 | 113.667 | 5.0 | 160429.0 | 5387.0 | 125041.0 | 32085.8 |

Table S40: Results obtained for DS38 using the benchmark proposed. The execution time was measured in seconds, the RAM usage was measured in GB and the CPU usage is presented as a percentage. The executions were, when possible, capped at 6 threads and 48 GB of RAM.

| Reconstruction tool | Execution time | SNPs | Identity | NCSD | NRC | RAM usage | CPU usage | Number of scaffolds | Reconstructed bases | Minimum scaffold length | Maximum scaffold length | Average scaffold length |
|---|---|---|---|---|---|---|---|---|---|---|---|---|
| coronaSPAdes | 2.08 | 13.0 | 99.989 | 0.034 | 0.026 | 0.136 | 413.333 | 19.0 | 210132.0 | 696.0 | 49969.0 | 11059.6 |
| Haploflow | 2.05 | 0.0 | 100.0 | 0.619 | 0.601 | 0.069 | 99.0 | 104.0 | 86123.0 | 503.0 | 2164.0 | 828.1 |
| IRMA | 168.96 | 9.333 | 99.847 | 0.13 | 0.128 | 0.486 | 474.0 | 5.0 | 145534.0 | 4871.0 | 110982.0 | 29106.8 |
| LAZYPIPE | 4.515 | 4.0 | 99.995 | 0.039 | 0.032 | 1.51 | 210.667 | 22.0 | 209468.0 | 262.0 | 48758.0 | 9521.3 |
| metaSPAdes | 3.8 | 7.0 | 99.992 | 0.033 | 0.026 | 0.227 | 394.333 | 20.0 | 210228.0 | 341.0 | 49969.0 | 10511.4 |
| metaviralSPAdes | – | – | – | – | – | – | – | – | – | – | – | – |
| PEHaplo | 16.915 | 4.0 | 99.615 | 0.056 | 0.046 | 0.424 | 101.667 | 192.0 | 229893.0 | 382.0 | 4797.0 | 1197.4 |
| QuRe | 1250.83 | 0.0 | 100.0 | 1.0 | 0.998 | 53.058 | 508.333 | 2.0 | 256.0 | 128.0 | 128.0 | 128.0 |
| QVG | 27.45 | 7540.0 | 94.73 | 0.246 | 0.241 | 1.4 | 133.333 | 7.0 | 279890.0 | 5387.0 | 125044.0 | 39984.3 |
| SPAdes | 5.92 | 5.0 | 99.993 | 0.033 | 0.026 | 0.159 | 316.333 | 18.0 | 210085.0 | 2166.0 | 49969.0 | 11671.4 |
| SSAKE | 1.7 | 0.667 | 99.996 | 0.928 | 0.916 | 0.05 | 100.0 | 113.667 | 20939.333 | 102.0 | 451.667 | 184.233 |
| TRACESPipe | 175.74 | 758.0 | 99.217 | 0.08 | 0.07 | 9.768 | 357.0 | 4.0 | 143860.0 | 5387.0 | 125041.0 | 35965.0 |
| TRACESPipeLite | 39.02 | 0.0 | 100.0 | 0.999 | 0.997 | 2.824 | 351.667 | 6.0 | 168262.0 | 5387.0 | 125041.0 | 28043.7 |
| VirGenA | 544.415 | 74.667 | 99.727 | 0.663 | 0.645 | 4.466 | 737.0 | 73.0 | 55130.0 | 105.667 | 2842.667 | 772.867 |
| ViSpA | 76.595 | 915.333 | 97.045 | 0.168 | 0.158 | 6.355 | 85.667 | 88.0 | 3718653.0 | 412.0 | 113601.0 | 42257.4 |
| V-pipe | 38.6 | 78.0 | 94.147 | 0.357 | 0.334 | 0.104 | 114.667 | 5.0 | 160429.0 | 5387.0 | 125041.0 | 32085.8 |

Table S41: Results obtained for DS39 using the benchmark proposed. The execution time was measured in seconds, the RAM usage was measured in GB and the CPU usage is presented as a percentage. The executions were, when possible, capped at 6 threads and 48 GB of RAM.

| Reconstruction tool | Execution time | SNPs | Identity | NCSD | NRC | RAM usage | CPU usage | Number of scaffolds | Reconstructed bases | Minimum scaffold length | Maximum scaffold length | Average scaffold length |
|---|---|---|---|---|---|---|---|---|---|---|---|---|
| coronaSPAdes | 2.095 | 4.0 | 99.997 | 0.033 | 0.026 | 0.134 | 413.333 | 26.0 | 209877.0 | 392.0 | 28454.0 | 8072.2 |
| Haploflow | 2.06 | 0.0 | 100.0 | 0.631 | 0.613 | 0.069 | 99.667 | 112.0 | 87072.0 | 508.0 | 1820.0 | 777.4 |
| IRMA | 205.48 | 49.667 | 99.813 | 0.134 | 0.126 | 0.497 | 475.0 | 5.0 | 146048.0 | 4892.0 | 111889.0 | 29209.6 |
| LAZYPIPE | 4.105 | 3.0 | 99.998 | 0.043 | 0.035 | 1.509 | 197.333 | 27.0 | 208705.0 | 362.0 | 28385.0 | 7729.8 |
| metaSPAdes | 3.71 | 2.0 | 99.998 | 0.033 | 0.026 | 0.228 | 394.333 | 26.0 | 209877.0 | 392.0 | 28454.0 | 8072.2 |
| metaviralSPAdes | – | – | – | – | – | – | – | – | – | – | – | – |
| PEHaplo | 16.795 | 3.0 | 99.997 | 0.049 | 0.04 | 0.423 | 101.667 | 207.0 | 233449.0 | 385.0 | 5356.0 | 1127.8 |
| QuRe | 156.305 | 0.0 | 100.0 | 0.999 | 0.997 | 29.787 | 567.667 | 3.333 | 570.667 | 152.0 | 200.0 | 170.667 |
| QVG | 26.205 | 11690.0 | 91.79 | 0.395 | 0.387 | 1.399 | 135.667 | 7.0 | 279880.0 | 5385.0 | 125041.0 | 39982.9 |
| SPAdes | 7.28 | 2.0 | 99.998 | 0.033 | 0.026 | 0.158 | 278.0 | 26.0 | 209877.0 | 392.0 | 28454.0 | 8072.2 |
| SSAKE | 1.8 | 1.333 | 99.992 | 0.917 | 0.903 | 0.05 | 100.0 | 118.0 | 22427.0 | 104.333 | 426.333 | 190.067 |
| TRACESPipe | 195.91 | 345.0 | 98.7 | 0.093 | 0.08 | 9.768 | 329.667 | 4.0 | 143860.0 | 5387.0 | 125041.0 | 35965.0 |
| TRACESPipeLite | 36.545 | 0.0 | 0.0 | 1.0 | 0.998 | 2.824 | 365.333 | 5.0 | 160429.0 | 5387.0 | 125041.0 | 32085.8 |
| VirGenA | 554.24 | 59.667 | 99.8 | 0.626 | 0.608 | 4.352 | 733.333 | 80.667 | 60899.333 | 113.333 | 2620.0 | 762.467 |
| ViSpA | 80.615 | 489.667 | 95.815 | 0.361 | 0.351 | 6.056 | 85.333 | 139.0 | 5120204.0 | 464.0 | 91748.0 | 36836.0 |
| V-pipe | 38.49 | 457.0 | 93.037 | 0.478 | 0.453 | 0.104 | 114.667 | 5.0 | 160429.0 | 5387.0 | 125041.0 | 32085.8 |

Table S42: Results obtained for DS40 using the benchmark proposed. The execution time was measured in seconds, the RAM usage was measured in GB and the CPU usage is presented as a percentage. The executions were, when possible, capped at 6 threads and 48 GB of RAM.

| Reconstruction tool | Execution time | SNPs | Identity | NCSD | NRC | RAM usage | CPU usage | Number of scaffolds | Reconstructed bases | Minimum scaffold length | Maximum scaffold length | Average scaffold length |
|---|---|---|---|---|---|---|---|---|---|---|---|---|
| coronaSPAdes | 2.09 | 12.0 | 99.991 | 0.033 | 0.025 | 0.134 | 415.667 | 19.0 | 209941.0 | 747.0 | 43152.0 | 11049.5 |
| Haploflow | 2.08 | 0.0 | 100.0 | 0.676 | 0.657 | 0.07 | 99.667 | 100.0 | 75548.0 | 502.0 | 1689.0 | 755.5 |
| IRMA | 223.015 | 36.667 | 99.801 | 0.144 | 0.131 | 0.506 | 475.0 | 5.0 | 145808.0 | 4892.0 | 111955.0 | 29161.6 |
| LAZYPIPE | 4.055 | 4.0 | 99.996 | 0.04 | 0.032 | 1.509 | 194.667 | 19.0 | 209013.0 | 740.0 | 43095.0 | 11000.7 |
| metaSPAdes | 3.83 | 7.0 | 99.993 | 0.033 | 0.025 | 0.227 | 396.0 | 19.0 | 209941.0 | 747.0 | 43152.0 | 11049.5 |
| metaviralSPAdes | – | – | – | – | – | – | – | – | – | – | – | – |
| PEHaplo | 16.88 | 3.0 | 99.997 | 0.044 | 0.037 | 0.42 | 101.333 | 201.0 | 238048.0 | 378.0 | 5518.0 | 1184.3 |
| QuRe | 169.19 | 0.0 | 100.0 | 1.0 | 0.999 | 27.323 | 588.333 | 1.0 | 164.0 | 164.0 | 164.0 | 164.0 |
| QVG | 24.015 | 15561.0 | 89.044 | 0.521 | 0.516 | 1.398 | 134.333 | 6.0 | 274495.0 | 5387.0 | 125041.0 | 45749.2 |
| SPAdes | 6.76 | 5.0 | 99.996 | 0.032 | 0.025 | 0.158 | 290.333 | 19.0 | 209941.0 | 747.0 | 43152.0 | 11049.5 |
| SSAKE | 1.745 | 0.667 | 99.996 | 0.927 | 0.913 | 0.05 | 100.0 | 117.667 | 21592.0 | 103.333 | 377.667 | 183.5 |
| TRACESPipe | 194.315 | 1647.0 | 98.52 | 0.145 | 0.134 | 9.768 | 322.0 | 4.0 | 143860.0 | 5387.0 | 125041.0 | 35965.0 |
| TRACESPipeLite | 36.535 | 0.0 | 0.0 | 1.0 | 0.999 | 2.825 | 366.0 | 5.0 | 160429.0 | 5387.0 | 125041.0 | 32085.8 |
| VirGenA | 529.36 | 72.0 | 99.819 | 0.614 | 0.596 | 4.517 | 732.0 | 82.333 | 62748.333 | 155.0 | 2665.667 | 773.8 |
| ViSpA | 53.24 | 228.0 | 95.858 | 0.586 | 0.576 | 5.694 | 79.0 | 115.0 | 3300714.0 | 411.0 | 63874.0 | 28701.9 |
| V-pipe | 38.25 | 212.0 | 91.872 | 0.621 | 0.596 | 0.104 | 115.0 | 5.0 | 160429.0 | 5387.0 | 125041.0 | 32085.8 |

Table S43: Results obtained for DS41 using the benchmark proposed. The execution time was measured in seconds, the RAM usage was measured in GB and the CPU usage is presented as a percentage. The executions were, when possible, capped at 6 threads and 48 GB of RAM.

| Reconstruction tool | Execution time | SNPs | Identity | NCSD | NRC | RAM usage | CPU usage | Number of scaffolds | Reconstructed bases | Minimum scaffold length | Maximum scaffold length | Average scaffold length |
|---|---|---|---|---|---|---|---|---|---|---|---|---|
| coronaSPAdes | 3.135 | 1.0 | 99.9 | 0.03 | 0.022 | 0.142 | 410.0 | 9.0 | 202464.0 | 127.0 | 117531.0 | 22496.0 |
| Haploflow | 3.565 | 13.0 | 99.976 | 0.061 | 0.048 | 0.1 | 99.0 | 53.0 | 201565.0 | 556.0 | 11370.0 | 3803.1 |
| IRMA | 167.785 | 0.0 | 100.0 | 0.088 | 0.128 | 0.879 | 537.0 | 5.0 | 144829.0 | 4887.0 | 110250.0 | 28965.8 |
| LAZYPIPE | 7.115 | 0.0 | 99.971 | 0.03 | 0.022 | 1.525 | 254.0 | 7.0 | 202223.0 | 5322.0 | 96482.0 | 28889.0 |
| metaSPAdes | 5.59 | 0.0 | 99.952 | 0.029 | 0.021 | 0.229 | 410.667 | 9.0 | 202467.0 | 71.0 | 105065.0 | 22496.3 |
| metaviralSPAdes | – | – | – | – | – | – | – | – | – | – | – | – |
| PEHaplo | 31.735 | 0.0 | 100.0 | 0.073 | 0.061 | 1.162 | 133.333 | 312.0 | 252737.0 | 375.0 | 2818.0 | 810.1 |
| QuRe | 438.51 | 0.0 | 100.0 | 0.998 | 0.992 | 19.134 | 566.667 | 4.333 | 764.333 | 124.0 | 271.0 | 179.333 |
| QVG | – | – | – | – | – | – | – | – | – | – | – | – |
| SPAdes | 9.97 | 0.0 | 99.917 | 0.029 | 0.021 | 0.161 | 285.667 | 8.0 | 202590.0 | 401.0 | 112460.0 | 25323.8 |
| SSAKE | 5.595 | 3.667 | 99.977 | 0.475 | 0.431 | 0.08 | 100.0 | 371.333 | 121382.0 | 102.0 | 2740.0 | 326.9 |
| TRACESPipe | 174.335 | 0.0 | 100.0 | 0.029 | 0.021 | 9.768 | 359.333 | 4.0 | 143860.0 | 5387.0 | 125041.0 | 35965.0 |
| TRACESPipeLite | 44.705 | 1.0 | 99.991 | 0.028 | 0.02 | 2.819 | 319.333 | 6.0 | 168262.0 | 5387.0 | 125041.0 | 28043.7 |
| VirGenA | – | – | – | – | – | – | – | – | – | – | – | – |
| ViSpA | 154.7 | 0.0 | 100.0 | 0.029 | 0.021 | 5.972 | 112.333 | 5.0 | 160119.0 | 5322.0 | 124995.0 | 32023.8 |
| V-pipe | 45.975 | 0.0 | 99.837 | 0.042 | 0.032 | 0.104 | 114.333 | 5.0 | 160429.0 | 5387.0 | 125041.0 | 32085.8 |

Table S44: Results obtained for DS42 using the benchmark proposed. The execution time was measured in seconds, the RAM usage was measured in GB and the CPU usage is presented as a percentage. The executions were, when possible, capped at 6 threads and 48 GB of RAM.

| Reconstruction tool | Execution time | SNPs | Identity | NCSD | NRC | RAM usage | CPU usage | Number of scaffolds | Reconstructed bases | Minimum scaffold length | Maximum scaffold length | Average scaffold length |
|---|---|---|---|---|---|---|---|---|---|---|---|---|
| coronaSPAdes | 3.07 | 7.0 | 99.995 | 0.036 | 0.026 | 0.165 | 418.667 | 13.0 | 207930.0 | 262.0 | 105627.0 | 15994.6 |
| Haploflow | 3.58 | 142.0 | 99.844 | 0.052 | 0.04 | 0.1 | 99.0 | 57.0 | 206813.0 | 507.0 | 16242.0 | 3628.3 |
| IRMA | 195.99 | 0.0 | 100.0 | 0.104 | 0.128 | 0.877 | 540.0 | 5.0 | 144804.0 | 4868.0 | 110235.0 | 28960.8 |
| LAZYPIPE | 6.86 | 291.0 | 99.786 | 0.032 | 0.022 | 1.525 | 267.0 | 10.0 | 210000.0 | 392.0 | 97589.0 | 21000.0 |
| metaSPAdes | 5.62 | 84.0 | 99.944 | 0.035 | 0.024 | 0.229 | 412.0 | 7.0 | 208833.0 | 1027.0 | 122573.0 | 29833.3 |
| metaviralSPAdes | – | – | – | – | – | – | – | – | – | – | – | – |
| PEHaplo | 31.74 | 108.0 | 99.888 | 0.084 | 0.068 | 1.142 | 132.333 | 325.0 | 263012.0 | 375.0 | 2428.0 | 809.3 |
| QuRe | 939.94 | 16.667 | 99.476 | 0.989 | 0.977 | 42.47 | 546.667 | 6.0 | 4321.0 | 118.667 | 1437.0 | 707.933 |
| QVG | 27.39 | 551.0 | 99.537 | 0.046 | 0.037 | 1.409 | 156.333 | 5.0 | 268901.0 | 5387.0 | 125041.0 | 53780.2 |
| SPAdes | 6.025 | 2.0 | 99.999 | 0.031 | 0.021 | 0.161 | 399.333 | 7.0 | 210263.0 | 5370.0 | 107709.0 | 30037.6 |
| SSAKE | 5.59 | 6.667 | 99.985 | 0.537 | 0.506 | 0.079 | 100.0 | 359.0 | 113578.333 | 101.333 | 1099.0 | 316.433 |
| TRACESPipe | 170.04 | 438.0 | 99.663 | 0.039 | 0.029 | 9.768 | 360.667 | 4.0 | 143860.0 | 5387.0 | 125041.0 | 35965.0 |
| TRACESPipeLite | 46.425 | 457.333 | 99.353 | 0.06 | 0.048 | 2.821 | 328.333 | 6.0 | 168262.0 | 5387.0 | 125041.0 | 28043.7 |
| VirGenA | 830.625 | 127.667 | 99.713 | 0.65 | 0.623 | 4.427 | 743.0 | 55.333 | 54882.333 | 250.333 | 3061.0 | 1010.967 |
| ViSpA | 148.715 | 444.0 | 99.676 | 0.043 | 0.032 | 6.12 | 110.0 | 5.0 | 160099.0 | 5336.0 | 124985.0 | 32019.8 |
| V-pipe | 47.45 | 362.0 | 99.398 | 0.051 | 0.04 | 0.104 | 112.667 | 5.0 | 160429.0 | 5387.0 | 125041.0 | 32085.8 |

Table S45: Results obtained for DS43 using the benchmark proposed. The execution time was measured in seconds, the RAM usage was measured in GB and the CPU usage is presented as a percentage. The executions were, when possible, capped at 6 threads and 48 GB of RAM.

| Reconstruction tool | Execution time | SNPs | Identity | NCSD | NRC | RAM usage | CPU usage | Number of scaffolds | Reconstructed bases | Minimum scaffold length | Maximum scaffold length | Average scaffold length |
|---|---|---|---|---|---|---|---|---|---|---|---|---|
| coronaSPAdes | 3.1 | 2.0 | 99.991 | 0.031 | 0.022 | 0.142 | 408.667 | 6.0 | 210231.0 | 5328.0 | 124992.0 | 35038.5 |
| Haploflow | 3.56 | 616.0 | 99.537 | 0.058 | 0.045 | 0.1 | 99.0 | 44.0 | 206813.0 | 600.0 | 15451.0 | 4700.3 |
| IRMA | 262.66 | 0.0 | 99.999 | 0.109 | 0.128 | 0.88 | 537.667 | 5.0 | 144812.0 | 4878.0 | 110278.0 | 28962.4 |
| LAZYPIPE | 6.155 | 6.0 | 99.981 | 0.032 | 0.022 | 1.525 | 254.0 | 8.0 | 210138.0 | 351.0 | 109533.0 | 26267.3 |
| metaSPAdes | 5.63 | 2.0 | 99.998 | 0.031 | 0.022 | 0.229 | 413.0 | 6.0 | 210231.0 | 5328.0 | 124992.0 | 35038.5 |
| metaviralSPAdes | – | – | – | – | – | – | – | – | – | – | – | – |
| PEHaplo | 31.79 | 145.0 | 99.839 | 0.075 | 0.062 | 1.144 | 133.667 | 358.0 | 273251.0 | 375.0 | 3051.0 | 763.3 |
| QuRe | 410.405 | 1.333 | 99.85 | 0.995 | 0.988 | 16.62 | 562.0 | 7.0 | 1825.333 | 105.0 | 531.0 | 262.167 |
| QVG | 28.615 | 812.0 | 99.432 | 0.053 | 0.043 | 1.409 | 154.333 | 5.0 | 268901.0 | 5387.0 | 125041.0 | 53780.2 |
| SPAdes | 6.31 | 0.0 | 100.0 | 0.031 | 0.022 | 0.162 | 389.0 | 6.0 | 210231.0 | 5328.0 | 124992.0 | 35038.5 |
| SSAKE | 5.385 | 7.333 | 99.988 | 0.522 | 0.493 | 0.079 | 100.0 | 377.667 | 119055.667 | 102.667 | 1108.333 | 315.333 |
| TRACESPipe | 176.49 | 741.0 | 99.481 | 0.041 | 0.031 | 9.768 | 360.333 | 4.0 | 143864.0 | 5387.0 | 125045.0 | 35966.0 |
| TRACESPipeLite | 44.145 | 271.333 | 97.621 | 0.315 | 0.287 | 2.824 | 329.0 | 6.0 | 168264.0 | 5387.0 | 125043.0 | 28044.0 |
| VirGenA | 729.97 | 93.0 | 99.788 | 0.666 | 0.642 | 4.277 | 736.5 | 68.0 | 54227.0 | 65.0 | 2055.0 | 797.5 |
| ViSpA | 149.68 | 801.0 | 99.41 | 0.049 | 0.038 | 6.123 | 111.667 | 5.0 | 160128.0 | 5317.0 | 124991.0 | 32025.6 |
| V-pipe | 47.43 | 693.0 | 99.348 | 0.055 | 0.045 | 0.104 | 113.667 | 5.0 | 160429.0 | 5387.0 | 125041.0 | 32085.8 |

Table S46: Results obtained for DS44 using the benchmark proposed. The execution time was measured in seconds, the RAM usage was measured in GB and the CPU usage is presented as a percentage. The executions were, when possible, capped at 6 threads and 48 GB of RAM.

| Reconstruction tool | Execution time | SNPs | Identity | NCSD | NRC | RAM usage | CPU usage | Number of scaffolds | Reconstructed bases | Minimum scaffold length | Maximum scaffold length | Average scaffold length |
|---|---|---|---|---|---|---|---|---|---|---|---|---|
| coronaSPAdes | 2.89 | 0.0 | 100.0 | 0.03 | 0.021 | 0.144 | 431.0 | 6.0 | 210201.0 | 5363.0 | 125004.0 | 35033.5 |
| Haploflow | 3.56 | 69.0 | 99.933 | 0.058 | 0.047 | 0.1 | 99.0 | 61.0 | 206286.0 | 534.0 | 11587.0 | 3381.7 |
| IRMA | 269.055 | 0.0 | 100.0 | 0.114 | 0.127 | 0.881 | 551.0 | 5.0 | 145000.0 | 4908.0 | 110436.0 | 29000.0 |
| LAZYPIPE | 5.315 | 0.0 | 99.983 | 0.031 | 0.022 | 1.525 | 235.333 | 6.0 | 210024.0 | 5318.0 | 124924.0 | 35004.0 |
| metaSPAdes | 5.54 | 0.0 | 100.0 | 0.03 | 0.021 | 0.23 | 412.667 | 6.0 | 210201.0 | 5363.0 | 125004.0 | 35033.5 |
| metaviralSPAdes | – | – | – | – | – | – | – | – | – | – | – | – |
| PEHaplo | 31.285 | 44.0 | 99.949 | 0.078 | 0.065 | 1.147 | 133.667 | 328.0 | 268990.0 | 377.0 | 2897.0 | 820.1 |
| QuRe | 436.055 | 1.0 | 99.766 | 0.999 | 0.994 | 19.368 | 579.0 | 5.0 | 893.333 | 141.667 | 220.333 | 178.7 |
| QVG | 32.375 | 1293.0 | 99.056 | 0.068 | 0.058 | 1.409 | 155.333 | 6.0 | 274497.0 | 5387.0 | 125041.0 | 45749.5 |
| SPAdes | 6.33 | 0.0 | 100.0 | 0.03 | 0.021 | 0.162 | 389.333 | 6.0 | 210201.0 | 5363.0 | 125004.0 | 35033.5 |
| SSAKE | 5.815 | 3.0 | 99.994 | 0.506 | 0.48 | 0.08 | 100.0 | 377.667 | 119891.0 | 101.0 | 1146.0 | 317.433 |
| TRACESPipe | 189.22 | 893.0 | 99.294 | 0.046 | 0.036 | 9.768 | 338.667 | 4.0 | 143863.0 | 5387.0 | 125043.0 | 35965.8 |
| TRACESPipeLite | 41.835 | 33.333 | 98.268 | 0.801 | 0.781 | 2.825 | 337.0 | 6.0 | 168262.0 | 5387.0 | 125041.0 | 28043.7 |
| VirGenA | 781.02 | 87.0 | 99.827 | 0.649 | 0.631 | 4.476 | 737.667 | 63.0 | 56079.0 | 80.0 | 2749.0 | 890.1 |
| ViSpA | 72.62 | 1069.0 | 99.064 | 0.057 | 0.047 | 6.0 | 111.333 | 10.0 | 304898.0 | 4090.0 | 124891.0 | 30489.8 |
| V-pipe | 47.4 | 840.0 | 99.045 | 0.065 | 0.055 | 0.104 | 113.333 | 5.0 | 160429.0 | 5387.0 | 125041.0 | 32085.8 |

Table S47: Results obtained for DS45 using the benchmark proposed. The execution time was measured in seconds, the RAM usage was measured in GB and the CPU usage is presented as a percentage. The executions were, when possible, capped at 6 threads and 48 GB of RAM.

| Reconstruction tool | Execution time | SNPs | Identity | NCSD | NRC | RAM usage | CPU usage | Number of scaffolds | Reconstructed bases | Minimum scaffold length | Maximum scaffold length | Average scaffold length |
|---|---|---|---|---|---|---|---|---|---|---|---|---|
| coronaSPAdes | 2.9 | 1.0 | 99.991 | 0.029 | 0.022 | 0.142 | 428.667 | 6.0 | 210140.0 | 5364.0 | 124966.0 | 35023.3 |
| Haploflow | 3.57 | 0.0 | 100.0 | 0.059 | 0.048 | 0.1 | 99.333 | 45.0 | 206927.0 | 647.0 | 19632.0 | 4598.4 |
| IRMA | 316.405 | 8.333 | 99.994 | 0.115 | 0.123 | 0.888 | 560.0 | 5.0 | 145734.333 | 4894.0 | 111130.333 | 29146.867 |
| LAZYPIPE | 5.325 | 0.0 | 100.0 | 0.03 | 0.023 | 1.525 | 236.667 | 6.0 | 210022.0 | 5316.0 | 124919.0 | 35003.7 |
| metaSPAdes | 5.565 | 1.0 | 99.991 | 0.029 | 0.022 | 0.229 | 412.333 | 6.0 | 210147.0 | 5364.0 | 124966.0 | 35024.5 |
| metaviralSPAdes | – | – | – | – | – | – | – | – | – | – | – | – |
| PEHaplo | 31.145 | 39.0 | 99.935 | 0.077 | 0.064 | 1.134 | 132.333 | 317.0 | 255235.0 | 377.0 | 3917.0 | 805.2 |
| QuRe | 701.145 | 1.667 | 99.872 | 0.995 | 0.99 | 17.776 | 596.333 | 5.0 | 1501.333 | 139.0 | 487.0 | 303.367 |
| QVG | 35.155 | 1950.0 | 98.633 | 0.085 | 0.078 | 1.409 | 156.667 | 7.0 | 279882.0 | 5387.0 | 125040.0 | 39983.1 |
| SPAdes | 6.86 | 1.0 | 99.991 | 0.029 | 0.022 | 0.161 | 368.0 | 6.0 | 210147.0 | 5364.0 | 124966.0 | 35024.5 |
| SSAKE | 5.52 | 4.0 | 99.993 | 0.536 | 0.514 | 0.079 | 100.0 | 364.333 | 113342.667 | 102.667 | 1526.333 | 311.067 |
| TRACESPipe | 181.755 | 1516.0 | 98.594 | 0.088 | 0.077 | 9.768 | 357.0 | 4.0 | 143847.0 | 5387.0 | 125052.0 | 35961.8 |
| TRACESPipeLite | 41.105 | 2.0 | 99.509 | 0.971 | 0.962 | 2.827 | 340.667 | 6.0 | 168262.0 | 5387.0 | 125041.0 | 28043.7 |
| VirGenA | – | – | – | – | – | – | – | – | – | – | – | – |
| ViSpA | 65.22 | 1287.667 | 98.429 | 0.074 | 0.063 | 5.994 | 113.667 | 5.0 | 1501657.0 | 5314.0 | 1460372.0 | 300331.4 |
| V-pipe | 47.365 | 908.0 | 98.817 | 0.066 | 0.056 | 0.104 | 114.0 | 5.0 | 160429.0 | 5387.0 | 125041.0 | 32085.8 |

Table S48: Results obtained for DS46 using the benchmark proposed. The execution time was measured in seconds, the RAM usage was measured in GB and the CPU usage is presented as a percentage. The executions were, when possible, capped at 6 threads and 48 GB of RAM.

| Reconstruction tool | Execution time | SNPs | Identity | NCSD | NRC | RAM usage | CPU usage | Number of scaffolds | Reconstructed bases | Minimum scaffold length | Maximum scaffold length | Average scaffold length |
|---|---|---|---|---|---|---|---|---|---|---|---|---|
| coronaSPAdes | 2.88 | 3.0 | 99.99 | 0.028 | 0.021 | 0.142 | 430.333 | 6.0 | 210135.0 | 5381.0 | 124977.0 | 35022.5 |
| Haploflow | 3.58 | 0.0 | 100.0 | 0.057 | 0.046 | 0.1 | 99.667 | 49.0 | 206675.0 | 560.0 | 24401.0 | 4217.9 |
| IRMA | 275.825 | 3.333 | 99.989 | 0.122 | 0.12 | 0.892 | 554.333 | 5.0 | 146285.0 | 4890.0 | 111769.0 | 29257.0 |
| LAZYPIPE | 4.945 | 0.0 | 100.0 | 0.029 | 0.022 | 1.526 | 224.667 | 6.0 | 209964.0 | 5368.0 | 124939.0 | 34994.0 |
| metaSPAdes | 5.545 | 0.0 | 100.0 | 0.028 | 0.021 | 0.229 | 411.0 | 6.0 | 210135.0 | 5381.0 | 124977.0 | 35022.5 |
| metaviralSPAdes | – | – | – | – | – | – | – | – | – | – | – | – |
| PEHaplo | 31.31 | 33.0 | 99.944 | 0.07 | 0.058 | 1.17 | 132.333 | 299.0 | 245592.0 | 375.0 | 2712.0 | 821.4 |
| QuRe | 488.97 | 0.333 | 99.971 | 0.997 | 0.993 | 17.356 | 564.667 | 8.333 | 1372.333 | 73.0 | 259.0 | 164.733 |
| QVG | 34.725 | 3754.0 | 97.368 | 0.139 | 0.134 | 1.409 | 153.333 | 6.0 | 274272.0 | 5387.0 | 125033.0 | 45712.0 |
| SPAdes | 7.055 | 0.0 | 100.0 | 0.028 | 0.021 | 0.161 | 362.667 | 6.0 | 210135.0 | 5381.0 | 124977.0 | 35022.5 |
| SSAKE | 5.49 | 4.0 | 99.994 | 0.524 | 0.502 | 0.079 | 100.0 | 357.0 | 115824.333 | 102.0 | 1312.333 | 324.5 |
| TRACESPipe | 189.475 | 239.0 | 99.809 | 0.039 | 0.031 | 9.768 | 346.0 | 4.0 | 143831.0 | 5358.0 | 125041.0 | 35957.8 |
| TRACESPipeLite | 40.565 | 0.0 | 100.0 | 1.0 | 0.997 | 2.827 | 340.333 | 6.0 | 168262.0 | 5387.0 | 125041.0 | 28043.7 |
| VirGenA | 736.52 | 30.0 | 99.694 | 0.699 | 0.682 | 4.071 | 740.667 | 56.0 | 48874.0 | 116.0 | 2225.0 | 872.8 |
| ViSpA | 91.885 | 1196.333 | 97.355 | 0.121 | 0.109 | 5.956 | 106.0 | 69.0 | 3266869.0 | 403.0 | 118958.0 | 47345.9 |
| V-pipe | 47.595 | 1120.0 | 97.644 | 0.123 | 0.108 | 0.104 | 113.333 | 5.0 | 160429.0 | 5387.0 | 125041.0 | 32085.8 |

Table S49: Results obtained for DS47 using the benchmark proposed. The execution time was measured in seconds, the RAM usage was measured in GB and the CPU usage is presented as a percentage. The executions were, when possible, capped at 6 threads and 48 GB of RAM.

| Reconstruction tool | Execution time | SNPs | Identity | NCSD | NRC | RAM usage | CPU usage | Number of scaffolds | Reconstructed bases | Minimum scaffold length | Maximum scaffold length | Average scaffold length |
|---|---|---|---|---|---|---|---|---|---|---|---|---|
| coronaSPAdes | 2.93 | 0.0 | 100.0 | 0.027 | 0.02 | 0.145 | 429.333 | 6.0 | 210293.0 | 5374.0 | 124987.0 | 35048.8 |
| Haploflow | 3.605 | 0.0 | 100.0 | 0.047 | 0.039 | 0.101 | 99.333 | 61.0 | 208117.0 | 504.0 | 12951.0 | 3411.8 |
| IRMA | 303.18 | 11.333 | 99.99 | 0.128 | 0.122 | 0.898 | 553.0 | 5.0 | 146154.667 | 4886.0 | 111820.667 | 29230.933 |
| LAZYPIPE | 6.365 | 2.0 | 99.997 | 0.033 | 0.026 | 1.525 | 269.0 | 24.0 | 210020.0 | 589.0 | 39288.0 | 8750.8 |
| metaSPAdes | 5.56 | 1.0 | 99.991 | 0.027 | 0.021 | 0.229 | 410.333 | 6.0 | 210293.0 | 5374.0 | 124987.0 | 35048.8 |
| metaviralSPAdes | – | – | – | – | – | – | – | – | – | – | – | – |
| PEHaplo | 31.27 | 0.0 | 100.0 | 0.072 | 0.061 | 1.155 | 131.667 | 310.0 | 252286.0 | 377.0 | 2966.0 | 813.8 |
| QuRe | 372.765 | 3.0 | 99.751 | 0.997 | 0.993 | 16.72 | 559.333 | 4.333 | 1143.333 | 192.0 | 396.333 | 262.8 |
| QVG | 35.275 | 6950.0 | 94.724 | 0.256 | 0.252 | 1.409 | 155.333 | 7.0 | 282112.0 | 5387.0 | 125033.0 | 40301.7 |
| SPAdes | 6.135 | 1.0 | 99.991 | 0.027 | 0.021 | 0.161 | 397.333 | 6.0 | 210293.0 | 5374.0 | 124987.0 | 35048.8 |
| SSAKE | 5.145 | 3.667 | 99.992 | 0.533 | 0.512 | 0.08 | 100.0 | 360.0 | 115435.0 | 102.0 | 1227.667 | 320.667 |
| TRACESPipe | 202.665 | 3.0 | 99.991 | 0.028 | 0.022 | 9.768 | 326.0 | 4.0 | 143701.0 | 5298.0 | 125007.0 | 35925.3 |
| TRACESPipeLite | 37.64 | 0.0 | 0.0 | 1.0 | 0.998 | 2.828 | 353.333 | 5.0 | 160429.0 | 5387.0 | 125041.0 | 32085.8 |
| VirGenA | 764.02 | 96.0 | 99.753 | 0.686 | 0.67 | 4.195 | 737.333 | 57.0 | 51406.0 | 94.0 | 2356.0 | 901.9 |
| ViSpA | 115.685 | 1109.0 | 96.25 | 0.283 | 0.273 | 6.334 | 99.0 | 162.0 | 6569618.0 | 500.0 | 101012.0 | 40553.2 |
| V-pipe | 47.425 | 1569.0 | 95.842 | 0.23 | 0.21 | 0.104 | 113.333 | 5.0 | 160429.0 | 5387.0 | 125041.0 | 32085.8 |

Table S50: Results obtained for DS48 using the benchmark proposed. The execution time was measured in seconds, the RAM usage was measured in GB and the CPU usage is presented as a percentage. The executions were, when possible, capped at 6 threads and 48 GB of RAM.

| Reconstruction tool | Execution time | SNPs | Identity | NCSD | NRC | RAM usage | CPU usage | Number of scaffolds | Reconstructed bases | Minimum scaffold length | Maximum scaffold length | Average scaffold length |
|---|---|---|---|---|---|---|---|---|---|---|---|---|
| coronaSPAdes | 3.005 | 0.0 | 100.0 | 0.027 | 0.021 | 0.146 | 421.333 | 6.0 | 210218.0 | 5341.0 | 125035.0 | 35036.3 |
| Haploflow | 3.58 | 0.0 | 100.0 | 0.046 | 0.038 | 0.1 | 99.667 | 50.0 | 208209.0 | 554.0 | 15846.0 | 4164.2 |
| IRMA | 341.925 | 55.667 | 99.956 | 0.128 | 0.117 | 0.919 | 555.333 | 5.0 | 147253.333 | 5033.0 | 112899.333 | 29450.667 |
| LAZYPIPE | 4.59 | 0.0 | 100.0 | 0.029 | 0.023 | 1.525 | 210.0 | 6.0 | 209997.0 | 5291.0 | 124999.0 | 34999.5 |
| metaSPAdes | 5.595 | 0.0 | 100.0 | 0.027 | 0.021 | 0.229 | 410.667 | 6.0 | 210218.0 | 5341.0 | 125035.0 | 35036.3 |
| metaviralSPAdes | – | – | – | – | – | – | – | – | – | – | – | – |
| PEHaplo | 31.275 | 9.0 | 99.986 | 0.09 | 0.076 | 1.157 | 131.667 | 305.0 | 247198.0 | 377.0 | 3432.0 | 810.5 |
| QuRe | 725.785 | 8.667 | 99.666 | 0.99 | 0.984 | 27.461 | 599.667 | 7.333 | 2964.0 | 109.333 | 1368.0 | 399.9 |
| QVG | 34.345 | 11569.0 | 91.836 | 0.398 | 0.395 | 1.409 | 160.0 | 8.0 | 287720.0 | 5387.0 | 125041.0 | 35965.0 |
| SPAdes | 5.935 | 0.0 | 100.0 | 0.027 | 0.021 | 0.161 | 405.667 | 6.0 | 210218.0 | 5341.0 | 125035.0 | 35036.3 |
| SSAKE | 5.795 | 3.333 | 99.995 | 0.499 | 0.476 | 0.081 | 100.0 | 384.667 | 124712.333 | 102.667 | 1256.0 | 324.133 |
| TRACESPipe | 202.775 | 52.0 | 99.851 | 0.039 | 0.032 | 9.768 | 323.333 | 4.0 | 143811.0 | 5361.0 | 125041.0 | 35952.8 |
| TRACESPipeLite | 37.35 | 0.0 | 0.0 | 1.0 | 0.999 | 2.828 | 354.0 | 5.0 | 160429.0 | 5387.0 | 125041.0 | 32085.8 |
| VirGenA | 790.66 | 10.0 | 99.761 | 0.734 | 0.717 | 3.95 | 744.0 | 60.0 | 44696.0 | 187.0 | 2269.0 | 744.9 |
| ViSpA | 89.27 | 514.667 | 95.995 | 0.482 | 0.473 | 5.942 | 97.333 | 168.0 | 5743888.0 | 433.0 | 77451.0 | 34189.8 |
| V-pipe | 47.02 | 566.0 | 94.245 | 0.386 | 0.364 | 0.104 | 113.333 | 5.0 | 160429.0 | 5387.0 | 125041.0 | 32085.8 |

Table S51: Results obtained for DS49 using the benchmark proposed. The execution time was measured in seconds, the RAM usage was measured in GB and the CPU usage is presented as a percentage. The executions were, when possible, capped at 6 threads and 48 GB of RAM.

| **Reconstruction tool** | **Execution time** | **SNPs** | **Identity** | **NCSD** | **NRC** | **RAM usage** | **CPU usage** | **Number of scaffolds** | **Reconstructed bases** | **Minimum scaffold length** | **Maximum scaffold length** | **Average scaffold length** |
|---|---|---|---|---|---|---|---|---|---|---|---|---|
| coronaSPAdes | 4.665 | 0.0 | 99.949 | 0.029 | 0.022 | 0.149 | 438.333 | 9.0 | 202617.0 | 147.0 | 112509.0 | 22513.0 |
| Haploflow | 6.5 | 8.0 | 99.989 | 0.03 | 0.022 | 0.151 | 99.0 | 14.0 | 203336.0 | 1385.0 | 54636.0 | 14524.0 |
| IRMA | 307.225 | 0.0 | 100.0 | 0.088 | 0.127 | 1.648 | 597.333 | 5.0 | 144862.0 | 4876.0 | 110239.0 | 28972.4 |
| LAZYPIPE | 9.515 | 0.0 | 99.994 | 0.028 | 0.021 | 1.541 | 269.0 | 8.0 | 202542.0 | 5373.0 | 97853.0 | 25317.8 |
| metaSPAdes | 8.75 | 15.0 | 99.8 | 0.029 | 0.021 | 0.226 | 429.667 | 10.0 | 202605.0 | 71.0 | 112462.0 | 20260.5 |
| metaviralSPAdes | 9.955 | 0.0 | 0.0 | 1.0 | 0.994 | 0.227 | 450.333 | 2.0 | 66537.0 | 16548.0 | 49989.0 | 33268.5 |
| PEHaplo | 83.755 | 0.0 | 99.999 | 0.031 | 0.024 | 3.052 | 142.0 | 215.0 | 311054.0 | 382.0 | 6699.0 | 1446.8 |
| QuRe | 3261.255 | 0.333 | 99.996 | 0.946 | 0.905 | 21.594 | 627.333 | 5.0 | 10109.0 | 312.0 | 6698.333 | 2021.8 |
| QVG | – | – | – | – | – | – | – | – | – | – | – | – |
| SPAdes | 10.155 | 0.0 | 99.9 | 0.029 | 0.02 | 0.185 | 393.0 | 9.0 | 202724.0 | 401.0 | 105060.0 | 22524.9 |
| SSAKE | 16.44 | 5.333 | 99.992 | 0.041 | 0.032 | 0.132 | 99.667 | 170.0 | 216031.667 | 103.0 | 11406.667 | 1271.9 |
| TRACESPipe | 188.11 | 0.0 | 100.0 | 0.028 | 0.02 | 9.768 | 360.667 | 4.0 | 143860.0 | 5387.0 | 125041.0 | 35965.0 |
| TRACESPipeLite | 52.26 | 0.0 | 100.0 | 0.028 | 0.02 | 2.826 | 286.333 | 6.0 | 168262.0 | 5387.0 | 125041.0 | 28043.7 |
| VirGenA | 1120.54 | 9.0 | 99.932 | 0.673 | 0.64 | 4.812 | 744.667 | 62.0 | 52819.333 | 68.0 | 2550.333 | 852.467 |
| ViSpA | 483.085 | 0.0 | 100.0 | 0.028 | 0.02 | 7.009 | 133.0 | 5.0 | 160254.0 | 5373.0 | 125010.0 | 32050.8 |
| V-pipe | 62.42 | 0.0 | 100.0 | 0.029 | 0.022 | 0.104 | 113.333 | 5.0 | 160429.0 | 5387.0 | 125041.0 | 32085.8 |

Table S52: Results obtained for DS50 using the benchmark proposed. The execution time was measured in seconds, the RAM usage was measured in GB and the CPU usage is presented as a percentage. The executions were, when possible, capped at 6 threads and 48 GB of RAM.

| **Reconstruction tool** | **Execution time** | **SNPs** | **Identity** | **NCSD** | **NRC** | **RAM usage** | **CPU usage** | **Number of scaffolds** | **Reconstructed bases** | **Minimum scaffold length** | **Maximum scaffold length** | **Average scaffold length** |
|---|---|---|---|---|---|---|---|---|---|---|---|---|
| coronaSPAdes | 4.65 | 6.0 | 99.996 | 0.036 | 0.025 | 0.171 | 445.667 | 12.0 | 207942.0 | 530.0 | 105623.0 | 17328.5 |
| Haploflow | 6.57 | 315.0 | 99.704 | 0.034 | 0.023 | 0.152 | 99.333 | 11.0 | 209651.0 | 1817.0 | 88819.0 | 19059.2 |
| IRMA | 363.795 | 0.0 | 100.0 | 0.103 | 0.127 | 1.646 | 598.667 | 5.0 | 144911.0 | 4884.0 | 110274.0 | 28982.2 |
| LAZYPIPE | 9.715 | 1.0 | 99.963 | 0.031 | 0.022 | 1.542 | 273.333 | 8.0 | 210172.0 | 5362.0 | 105627.0 | 26271.5 |
| metaSPAdes | 8.655 | 78.0 | 99.947 | 0.033 | 0.023 | 0.226 | 431.333 | 6.0 | 210344.0 | 5369.0 | 125025.0 | 35057.3 |
| metaviralSPAdes | 9.85 | 0.0 | 100.0 | 0.163 | 0.149 | 0.227 | 452.333 | 3.0 | 191573.0 | 16560.0 | 125025.0 | 63857.7 |
| PEHaplo | 79.955 | 114.0 | 99.917 | 0.036 | 0.027 | 2.966 | 145.0 | 243.0 | 344843.0 | 387.0 | 4514.0 | 1419.1 |
| QuRe | 4168.455 | 130.333 | 98.838 | 0.954 | 0.922 | 20.104 | 636.0 | 5.333 | 16396.333 | 627.333 | 5111.0 | 2704.367 |
| QVG | 39.435 | 498.0 | 99.577 | 0.041 | 0.034 | 1.419 | 175.333 | 5.0 | 268901.0 | 5387.0 | 125041.0 | 53780.2 |
| SPAdes | 9.78 | 1.0 | 99.999 | 0.031 | 0.021 | 0.186 | 415.333 | 6.0 | 210358.0 | 5369.0 | 125025.0 | 35059.7 |
| SSAKE | 16.61 | 34.667 | 99.943 | 0.042 | 0.031 | 0.132 | 100.0 | 167.667 | 222211.0 | 105.333 | 10487.0 | 1325.567 |
| TRACESPipe | 189.17 | 404.0 | 99.719 | 0.038 | 0.028 | 9.768 | 362.0 | 4.0 | 143861.0 | 5387.0 | 125042.0 | 35965.3 |
| TRACESPipeLite | 53.82 | 473.333 | 99.528 | 0.053 | 0.042 | 2.828 | 304.0 | 6.0 | 168261.0 | 5387.0 | 125040.0 | 28043.5 |
| VirGenA | 1118.28 | 136.0 | 99.536 | 0.728 | 0.696 | 4.464 | 748.667 | 42.333 | 44162.0 | 239.667 | 3002.667 | 1056.5 |
| ViSpA | 454.14 | 551.0 | 99.569 | 0.041 | 0.031 | 6.934 | 130.0 | 5.0 | 160260.0 | 5362.0 | 125030.0 | 32052.0 |
| V-pipe | 64.33 | 476.0 | 99.664 | 0.041 | 0.031 | 0.104 | 113.0 | 5.0 | 160429.0 | 5387.0 | 125041.0 | 32085.8 |

Table S53: Results obtained for DS51 using the benchmark proposed. The execution time was measured in seconds, the RAM usage was measured in GB and the CPU usage is presented as a percentage. The executions were, when possible, capped at 6 threads and 48 GB of RAM.

| **Reconstruction tool** | **Execution time** | **SNPs** | **Identity** | **NCSD** | **NRC** | **RAM usage** | **CPU usage** | **Number of scaffolds** | **Reconstructed bases** | **Minimum scaffold length** | **Maximum scaffold length** | **Average scaffold length** |
|---|---|---|---|---|---|---|---|---|---|---|---|---|
| coronaSPAdes | 4.555 | 2.0 | 99.999 | 0.03 | 0.021 | 0.153 | 436.333 | 6.0 | 210359.0 | 5371.0 | 125029.0 | 35059.8 |
| Haploflow | 6.505 | 245.0 | 99.792 | 0.033 | 0.024 | 0.15 | 99.667 | 11.0 | 216210.0 | 2101.0 | 110024.0 | 19655.5 |
| IRMA | 366.775 | 0.0 | 100.0 | 0.108 | 0.128 | 1.651 | 597.333 | 5.0 | 144931.0 | 4892.0 | 110278.0 | 28986.2 |
| LAZYPIPE | 9.62 | 1.0 | 99.478 | 0.03 | 0.021 | 1.541 | 271.0 | 8.0 | 210589.0 | 393.0 | 109777.0 | 26323.6 |
| metaSPAdes | 8.5 | 1.0 | 99.999 | 0.03 | 0.021 | 0.227 | 431.0 | 6.0 | 210359.0 | 5371.0 | 125029.0 | 35059.8 |
| metaviralSPAdes | 9.73 | 0.0 | 100.0 | 0.162 | 0.149 | 0.227 | 452.667 | 3.0 | 191575.0 | 16567.0 | 125029.0 | 63858.3 |
| PEHaplo | 76.86 | 0.0 | 99.944 | 0.046 | 0.035 | 2.917 | 146.0 | 242.0 | 324762.0 | 377.0 | 4784.0 | 1342.0 |
| QuRe | 2744.455 | 4.333 | 99.567 | 0.961 | 0.951 | 19.926 | 640.667 | 4.667 | 8286.0 | 543.333 | 3248.0 | 1778.933 |
| QVG | 42.745 | 741.0 | 99.488 | 0.047 | 0.04 | 1.418 | 174.333 | 5.0 | 268901.0 | 5387.0 | 125041.0 | 53780.2 |
| SPAdes | 9.16 | 1.0 | 99.999 | 0.03 | 0.021 | 0.187 | 423.667 | 6.0 | 210359.0 | 5371.0 | 125029.0 | 35059.8 |
| SSAKE | 16.565 | 0.333 | 99.999 | 0.041 | 0.031 | 0.133 | 100.0 | 170.667 | 222789.0 | 113.667 | 10299.0 | 1310.0 |
| TRACESPipe | 199.645 | 609.0 | 99.579 | 0.042 | 0.033 | 9.768 | 347.0 | 4.0 | 143862.0 | 5388.0 | 125041.0 | 35965.5 |
| TRACESPipeLite | 49.92 | 304.0 | 97.736 | 0.305 | 0.275 | 2.829 | 307.667 | 6.0 | 156261.0 | 5387.0 | 113040.0 | 26043.5 |
| VirGenA | 1071.385 | 62.0 | 99.659 | 0.694 | 0.674 | 4.739 | 747.667 | 47.0 | 48702.0 | 86.0 | 3067.0 | 1036.2 |
| ViSpA | 420.03 | 638.0 | 99.522 | 0.05 | 0.039 | 6.923 | 134.667 | 5.0 | 160284.0 | 5344.0 | 125026.0 | 32056.8 |
| V-pipe | 65.21 | 590.0 | 99.591 | 0.042 | 0.033 | 0.104 | 112.333 | 5.0 | 160429.0 | 5387.0 | 125041.0 | 32085.8 |

Table S54: Results obtained for DS52 using the benchmark proposed. The execution time was measured in seconds, the RAM usage was measured in GB and the CPU usage is presented as a percentage. The executions were, when possible, capped at 6 threads and 48 GB of RAM.

| Reconstruction tool | Execution time | SNPs | Identity | NCSD | NRC | RAM usage | CPU usage | Number of scaffolds | Reconstructed bases | Minimum scaffold length | Maximum scaffold length | Average scaffold length |
|---|---|---|---|---|---|---|---|---|---|---|---|---|
| coronaSPAdes | 4.485 | 0.0 | 100.0 | 0.029 | 0.021 | 0.156 | 450.333 | 6.0 | 210345.0 | 5354.0 | 125021.0 | 35057.5 |
| Haploflow | 6.705 | 499.0 | 99.371 | 0.033 | 0.024 | 0.158 | 99.0 | 9.0 | 209870.0 | 4765.0 | 94245.0 | 23318.9 |
| IRMA | 502.225 | 0.333 | 100.0 | 0.114 | 0.127 | 1.648 | 609.333 | 5.0 | 144956.0 | 4910.0 | 110307.0 | 28991.2 |
| LAZYPIPE | 7.88 | 0.0 | 99.985 | 0.029 | 0.021 | 1.54 | 244.0 | 6.0 | 210232.0 | 5332.0 | 124944.0 | 35038.7 |
| metaSPAdes | 8.51 | 0.0 | 100.0 | 0.029 | 0.021 | 0.226 | 431.333 | 6.0 | 210346.0 | 5355.0 | 125021.0 | 35057.7 |
| metaviralSPAdes | 9.695 | 0.0 | 100.0 | 0.161 | 0.149 | 0.227 | 452.0 | 3.0 | 191576.0 | 16565.0 | 125021.0 | 63858.7 |
| PEHaplo | 77.47 | 18.0 | 99.983 | 0.036 | 0.028 | 2.949 | 145.667 | 217.0 | 302159.0 | 393.0 | 5663.0 | 1392.4 |
| QuRe | 2009.37 | 7.333 | 99.882 | 0.966 | 0.958 | 21.751 | 638.667 | 5.0 | 7871.667 | 755.667 | 2247.333 | 1574.333 |
| QVG | 50.08 | 1058.0 | 99.221 | 0.056 | 0.049 | 1.418 | 177.667 | 7.0 | 282333.0 | 5387.0 | 125041.0 | 40333.3 |
| SPAdes | 9.075 | 0.0 | 100.0 | 0.029 | 0.021 | 0.186 | 424.333 | 6.0 | 210346.0 | 5355.0 | 125021.0 | 35057.7 |
| SSAKE | 16.49 | 0.0 | 99.998 | 0.038 | 0.028 | 0.134 | 99.667 | 157.333 | 222399.333 | 108.667 | 10223.333 | 1417.333 |
| TRACESPipe | 202.6 | 6.0 | 99.986 | 0.029 | 0.021 | 9.768 | 342.667 | 4.0 | 143860.0 | 5387.0 | 125041.0 | 35965.0 |
| TRACESPipeLite | 44.915 | 94.667 | 98.341 | 0.76 | 0.737 | 2.83 | 316.333 | 6.0 | 168263.0 | 5387.0 | 125042.0 | 28043.8 |
| VirGenA | 1129.92 | 100.0 | 99.732 | 0.716 | 0.697 | 4.641 | 749.0 | 43.333 | 45948.667 | 337.667 | 2685.0 | 1077.833 |
| ViSpA | 212.14 | 1025.667 | 99.072 | 0.059 | 0.051 | 7.054 | 133.333 | 11.0 | 492337.0 | 4092.0 | 125027.0 | 44757.9 |
| V-pipe | 65.87 | 556.0 | 99.201 | 0.048 | 0.039 | 0.104 | 111.667 | 5.0 | 160429.0 | 5387.0 | 125041.0 | 32085.8 |

Table S55: Results obtained for DS53 using the benchmark proposed. The execution time was measured in seconds, the RAM usage was measured in GB and the CPU usage is presented as a percentage. The executions were, when possible, capped at 6 threads and 48 GB of RAM.

| Reconstruction tool | Execution time | SNPs | Identity | NCSD | NRC | RAM usage | CPU usage | Number of scaffolds | Reconstructed bases | Minimum scaffold length | Maximum scaffold length | Average scaffold length |
|---|---|---|---|---|---|---|---|---|---|---|---|---|
| coronaSPAdes | 4.51 | 1.0 | 99.999 | 0.028 | 0.021 | 0.153 | 449.333 | 6.0 | 210316.0 | 5363.0 | 125016.0 | 35052.7 |
| Haploflow | 6.56 | 0.0 | 99.862 | 0.031 | 0.024 | 0.152 | 99.333 | 8.0 | 211036.0 | 5267.0 | 106254.0 | 26379.5 |
| IRMA | 509.245 | 7.333 | 99.976 | 0.114 | 0.121 | 1.67 | 615.667 | 5.0 | 146083.667 | 4898.0 | 111451.667 | 29216.733 |
| LAZYPIPE | 7.87 | 1.0 | 99.999 | 0.029 | 0.021 | 1.541 | 243.0 | 6.0 | 210236.0 | 5362.0 | 124985.0 | 35039.3 |
| metaSPAdes | 8.775 | 1.0 | 99.999 | 0.028 | 0.021 | 0.227 | 432.667 | 6.0 | 210317.0 | 5364.0 | 125016.0 | 35052.8 |
| metaviralSPAdes | 9.735 | 0.0 | 100.0 | 0.159 | 0.149 | 0.228 | 450.667 | 3.0 | 191566.0 | 16562.0 | 125016.0 | 63855.3 |
| PEHaplo | 77.42 | 148.0 | 99.874 | 0.039 | 0.032 | 2.949 | 145.667 | 230.0 | 312923.0 | 385.0 | 5712.0 | 1360.5 |
| QuRe | 4281.52 | 11.333 | 97.628 | 0.955 | 0.94 | 21.561 | 661.667 | 4.667 | 9349.667 | 327.667 | 5612.0 | 1921.567 |
| QVG | 52.19 | 1495.0 | 98.959 | 0.067 | 0.061 | 1.418 | 176.333 | 6.0 | 274497.0 | 5387.0 | 125041.0 | 45749.5 |
| SPAdes | 8.805 | 1.0 | 99.999 | 0.028 | 0.021 | 0.186 | 434.0 | 6.0 | 210317.0 | 5364.0 | 125016.0 | 35052.8 |
| SSAKE | 16.84 | 0.333 | 99.998 | 0.04 | 0.032 | 0.134 | 100.0 | 177.333 | 223227.0 | 109.0 | 9255.0 | 1258.9 |
| TRACESPipe | 191.53 | 1.0 | 99.999 | 0.028 | 0.021 | 9.768 | 347.333 | 4.0 | 143826.0 | 5387.0 | 125015.0 | 35956.5 |
| TRACESPipeLite | 44.375 | 0.0 | 99.651 | 0.963 | 0.953 | 2.832 | 316.667 | 6.0 | 168262.0 | 5387.0 | 125041.0 | 28043.7 |
| VirGenA | 1048.67 | 117.0 | 99.608 | 0.772 | 0.755 | 4.34 | 751.0 | 45.0 | 38219.0 | 49.0 | 2393.0 | 849.3 |
| ViSpA | 117.795 | 1309.667 | 98.537 | 0.081 | 0.07 | 6.876 | 132.667 | 11.333 | 1070682.0 | 4259.667 | 731683.333 | 157628.2 |
| V-pipe | 65.76 | 1157.0 | 99.096 | 0.054 | 0.046 | 0.104 | 112.333 | 5.0 | 160429.0 | 5387.0 | 125041.0 | 32085.8 |

Table S56: Results obtained for DS54 using the benchmark proposed. The execution time was measured in seconds, the RAM usage was measured in GB and the CPU usage is presented as a percentage. The executions were, when possible, capped at 6 threads and 48 GB of RAM.

| Reconstruction tool | Execution time | SNPs | Identity | NCSD | NRC | RAM usage | CPU usage | Number of scaffolds | Reconstructed bases | Minimum scaffold length | Maximum scaffold length | Average scaffold length |
|---|---|---|---|---|---|---|---|---|---|---|---|---|
| coronaSPAdes | 4.285 | 0.0 | 100.0 | 0.027 | 0.02 | 0.147 | 450.0 | 6.0 | 210356.0 | 5377.0 | 125033.0 | 35059.3 |
| Haploflow | 6.52 | 0.0 | 100.0 | 0.031 | 0.024 | 0.15 | 99.0 | 7.0 | 209821.0 | 5246.0 | 97636.0 | 29974.4 |
| IRMA | 511.585 | 6.0 | 99.993 | 0.121 | 0.119 | 1.681 | 615.0 | 5.0 | 146449.0 | 4911.0 | 111796.0 | 29289.8 |
| LAZYPIPE | 6.73 | 0.0 | 100.0 | 0.027 | 0.021 | 1.54 | 247.667 | 6.0 | 210294.0 | 5371.0 | 125028.0 | 35049.0 |
| metaSPAdes | 8.68 | 0.0 | 100.0 | 0.027 | 0.02 | 0.227 | 430.333 | 6.0 | 210360.0 | 5377.0 | 125037.0 | 35060.0 |
| metaviralSPAdes | 9.805 | 0.0 | 100.0 | 0.157 | 0.149 | 0.227 | 451.667 | 3.0 | 191590.0 | 16563.0 | 125037.0 | 63863.3 |
| PEHaplo | 77.29 | 14.0 | 99.99 | 0.033 | 0.027 | 2.934 | 146.667 | 236.0 | 317531.0 | 386.0 | 5125.0 | 1345.5 |
| QuRe | 2940.205 | 9.333 | 98.734 | 0.963 | 0.956 | 21.97 | 653.333 | 4.667 | 9153.667 | 288.0 | 3775.667 | 1847.5 |
| QVG | 51.87 | 2719.0 | 98.103 | 0.102 | 0.099 | 1.418 | 180.0 | 7.0 | 282128.0 | 5387.0 | 125043.0 | 40304.0 |
| SPAdes | 9.225 | 0.0 | 100.0 | 0.027 | 0.02 | 0.184 | 431.333 | 6.0 | 210360.0 | 5377.0 | 125037.0 | 35060.0 |
| SSAKE | 16.785 | 0.0 | 99.999 | 0.04 | 0.032 | 0.134 | 100.0 | 168.667 | 222414.0 | 107.667 | 10779.333 | 1320.9 |
| TRACESPipe | 206.685 | 0.0 | 100.0 | 0.027 | 0.02 | 9.768 | 332.0 | 4.0 | 143844.0 | 5387.0 | 125034.0 | 35961.0 |
| TRACESPipeLite | 43.725 | 0.0 | 100.0 | 0.998 | 0.995 | 2.832 | 318.0 | 6.0 | 168262.0 | 5387.0 | 125041.0 | 28043.7 |
| VirGenA | 1104.195 | 106.0 | 99.366 | 0.755 | 0.737 | 4.453 | 748.333 | 47.0 | 40533.0 | 70.0 | 2131.0 | 862.4 |
| ViSpA | 123.16 | 723.333 | 98.415 | 0.113 | 0.101 | 7.22 | 131.0 | 57.0 | 2607820.0 | 446.0 | 120733.0 | 45751.2 |
| V-pipe | 65.31 | 1248.0 | 98.469 | 0.088 | 0.077 | 0.104 | 112.333 | 5.0 | 160429.0 | 5387.0 | 125041.0 | 32085.8 |

Table S57: Results obtained for DS55 using the benchmark proposed. The execution time was measured in seconds, the RAM usage was measured in GB and the CPU usage is presented as a percentage. The executions were, when possible, capped at 6 threads and 48 GB of RAM.

| Reconstruction tool | Execution time | SNPs | Identity | NCSD | NRC | RAM usage | CPU usage | Number of scaffolds | Reconstructed bases | Minimum scaffold length | Maximum scaffold length | Average scaffold length |
|---|---|---|---|---|---|---|---|---|---|---|---|---|
| coronaSPAdes | 4.41 | 1.0 | 99.991 | 0.027 | 0.02 | 0.152 | 452.0 | 6.0 | 210365.0 | 5352.0 | 125038.0 | 35060.8 |
| Haploflow | 6.73 | 0.0 | 100.0 | 0.03 | 0.023 | 0.158 | 99.0 | 6.0 | 209884.0 | 5215.0 | 125010.0 | 34980.7 |
| IRMA | 625.51 | 30.333 | 99.969 | 0.126 | 0.121 | 1.686 | 614.667 | 5.0 | 146254.0 | 4919.0 | 111752.0 | 29250.8 |
| LAZYPIPE | 6.975 | 0.0 | 100.0 | 0.028 | 0.021 | 1.54 | 239.667 | 6.0 | 210254.0 | 5328.0 | 125032.0 | 35042.3 |
| metaSPAdes | 8.71 | 0.0 | 100.0 | 0.027 | 0.02 | 0.228 | 433.333 | 6.0 | 210365.0 | 5352.0 | 125038.0 | 35060.8 |
| metaviralSPAdes | 9.81 | 0.0 | 100.0 | 0.157 | 0.149 | 0.227 | 452.333 | 3.0 | 191599.0 | 16563.0 | 125038.0 | 63866.3 |
| PEHaplo | 77.335 | 1.0 | 99.999 | 0.03 | 0.023 | 2.952 | 146.667 | 231.0 | 311157.0 | 381.0 | 4353.0 | 1347.0 |
| QuRe | 4987.555 | 10.333 | 99.756 | 0.981 | 0.973 | 20.421 | 644.333 | 4.667 | 5779.667 | 257.333 | 2589.333 | 1370.1 |
| QVG | 51.48 | 5078.0 | 96.251 | 0.186 | 0.183 | 1.418 | 175.333 | 7.0 | 282124.0 | 5387.0 | 125039.0 | 40303.4 |
| SPAdes | 8.875 | 0.0 | 100.0 | 0.027 | 0.02 | 0.186 | 432.333 | 6.0 | 210365.0 | 5352.0 | 125038.0 | 35060.8 |
| SSAKE | 16.715 | 1.0 | 99.998 | 0.037 | 0.03 | 0.133 | 100.0 | 172.333 | 221907.0 | 111.333 | 9392.667 | 1287.967 |
| TRACESPipe | 214.09 | 1.0 | 99.999 | 0.027 | 0.021 | 9.768 | 328.0 | 4.0 | 143780.0 | 5369.0 | 125030.0 | 35945.0 |
| TRACESPipeLite | 41.18 | 0.0 | 0.0 | 1.0 | 0.998 | 2.832 | 329.333 | 5.0 | 160429.0 | 5387.0 | 125041.0 | 32085.8 |
| VirGenA | 1131.14 | 47.0 | 99.869 | 0.705 | 0.689 | 5.079 | 746.0 | 50.0 | 48048.0 | 64.0 | 2789.0 | 961.0 |
| ViSpA | 143.375 | 885.667 | 96.128 | 0.258 | 0.247 | 6.886 | 128.667 | 147.0 | 6180917.0 | 547.0 | 104055.0 | 42047.1 |
| V-pipe | 64.945 | 1529.0 | 97.021 | 0.162 | 0.146 | 0.104 | 112.333 | 5.0 | 160429.0 | 5387.0 | 125041.0 | 32085.8 |

Table S58: Results obtained for DS56 using the benchmark proposed. The execution time was measured in seconds, the RAM usage was measured in GB and the CPU usage is presented as a percentage. The executions were, when possible, capped at 6 threads and 48 GB of RAM.

| Reconstruction tool | Execution time | SNPs | Identity | NCSD | NRC | RAM usage | CPU usage | Number of scaffolds | Reconstructed bases | Minimum scaffold length | Maximum scaffold length | Average scaffold length |
|---|---|---|---|---|---|---|---|---|---|---|---|---|
| coronaSPAdes | 4.425 | 0.0 | 100.0 | 0.027 | 0.02 | 0.145 | 446.333 | 6.0 | 210347.0 | 5383.0 | 125013.0 | 35057.8 |
| Haploflow | 6.58 | 0.0 | 100.0 | 0.029 | 0.023 | 0.151 | 99.0 | 6.0 | 209839.0 | 5266.0 | 125013.0 | 34973.2 |
| IRMA | 760.465 | 131.667 | 99.832 | 0.118 | 0.109 | 1.747 | 614.333 | 5.0 | 148365.667 | 4898.0 | 113863.667 | 29673.133 |
| LAZYPIPE | 6.265 | 0.0 | 100.0 | 0.027 | 0.021 | 1.541 | 233.0 | 6.0 | 210280.0 | 5361.0 | 125011.0 | 35046.7 |
| metaSPAdes | 8.685 | 0.0 | 100.0 | 0.027 | 0.02 | 0.227 | 431.667 | 6.0 | 210347.0 | 5383.0 | 125013.0 | 35057.8 |
| metaviralSPAdes | 9.62 | 0.0 | 100.0 | 0.157 | 0.149 | 0.227 | 448.667 | 3.0 | 191553.0 | 16553.0 | 125013.0 | 63851.0 |
| PEHaplo | 76.98 | 0.0 | 99.982 | 0.03 | 0.023 | 2.929 | 145.667 | 232.0 | 319486.0 | 378.0 | 6558.0 | 1377.1 |
| QuRe | 3719.365 | 19.333 | 99.452 | 0.969 | 0.961 | 21.975 | 644.333 | 3.667 | 6776.0 | 777.333 | 3287.0 | 1907.833 |
| QVG | 45.865 | 9454.0 | 93.317 | 0.322 | 0.319 | 1.419 | 181.333 | 6.0 | 276745.0 | 5387.0 | 125045.0 | 46124.2 |
| SPAdes | 8.895 | 0.0 | 100.0 | 0.027 | 0.02 | 0.185 | 430.0 | 6.0 | 210347.0 | 5383.0 | 125013.0 | 35057.8 |
| SSAKE | 16.835 | 0.0 | 99.999 | 0.043 | 0.035 | 0.134 | 100.0 | 188.0 | 223020.0 | 103.0 | 10877.333 | 1186.367 |
| TRACESPipe | 214.57 | 0.0 | 100.0 | 0.027 | 0.021 | 9.768 | 327.333 | 4.0 | 143754.0 | 5381.0 | 124971.0 | 35938.5 |
| TRACESPipeLite | 41.935 | 0.0 | 0.0 | 1.0 | 0.999 | 2.833 | 328.0 | 5.0 | 160429.0 | 5387.0 | 125041.0 | 32085.8 |
| VirGenA | 1090.135 | 41.0 | 99.859 | 0.712 | 0.694 | 4.887 | 747.333 | 46.0 | 47781.0 | 301.0 | 3604.0 | 1038.7 |
| ViSpA | 121.98 | 669.667 | 96.097 | 0.434 | 0.426 | 6.946 | 119.0 | 200.0 | 6887728.0 | 461.0 | 83474.0 | 34438.6 |
| V-pipe | 64.435 | 1088.0 | 94.938 | 0.301 | 0.28 | 0.104 | 112.0 | 5.0 | 160429.0 | 5387.0 | 125041.0 | 32085.8 |

Table S59: Results obtained for DS57 using the benchmark proposed. The execution time was measured in seconds, the RAM usage was measured in GB and the CPU usage is presented as a percentage. The executions were, when possible, capped at 6 threads and 48 GB of RAM.

| Reconstruction tool | Execution time | SNPs | Identity | NCSD | NRC | RAM usage | CPU usage | Number of scaffolds | Reconstructed bases | Minimum scaffold length | Maximum scaffold length | Average scaffold length |
|---|---|---|---|---|---|---|---|---|---|---|---|---|
| coronaSPAdes | 2.935 | 67.0 | 99.901 | 0.036 | 0.026 | 0.148 | 414.667 | 8.0 | 187175.0 | 81.0 | 118363.0 | 23396.9 |
| Haploflow | 3.305 | 97.0 | 99.926 | 0.052 | 0.039 | 0.092 | 99.333 | 50.0 | 196302.0 | 513.0 | 15318.0 | 3926.0 |
| IRMA | 157.865 | 0.0 | 100.0 | 0.096 | 0.128 | 0.874 | 540.667 | 4.0 | 128277.0 | 4871.0 | 110232.0 | 32069.3 |
| LAZYPIPE | 6.71 | 16.0 | 99.874 | 0.036 | 0.025 | 1.523 | 260.333 | 8.0 | 186605.0 | 330.0 | 95821.0 | 23325.6 |
| metaSPAdes | 5.36 | 54.0 | 99.867 | 0.037 | 0.026 | 0.229 | 409.667 | 11.0 | 186838.0 | 71.0 | 105136.0 | 16985.3 |
| metaviralSPAdes | – | – | – | – | – | – | – | – | – | – | – | – |
| PEHaplo | 31.375 | 56.0 | 99.94 | 0.075 | 0.06 | 1.091 | 140.333 | 291.0 | 235200.0 | 376.0 | 2611.0 | 808.2 |
| QuRe | 571.335 | 17.667 | 99.514 | 0.986 | 0.969 | 14.26 | 592.333 | 8.333 | 6253.0 | 77.0 | 1723.0 | 716.933 |
| QVG | 21.51 | 160.0 | 99.893 | 0.035 | 0.025 | 1.408 | 153.333 | 4.0 | 143860.0 | 5387.0 | 125041.0 | 35965.0 |
| SPAdes | 6.93 | 38.0 | 99.928 | 0.035 | 0.024 | 0.162 | 355.0 | 22.0 | 195346.0 | 131.0 | 105523.0 | 8879.4 |
| SSAKE | 4.97 | 12.333 | 99.981 | 0.524 | 0.489 | 0.075 | 100.0 | 347.667 | 106998.0 | 103.333 | 1467.0 | 307.7 |
| TRACESPipe | 197.17 | 131.0 | 99.91 | 0.034 | 0.025 | 9.768 | 332.333 | 4.0 | 143861.0 | 5387.0 | 125042.0 | 35965.3 |
| TRACESPipeLite | 41.355 | 132.0 | 99.911 | 0.033 | 0.023 | 2.813 | 336.333 | 5.0 | 151693.0 | 5387.0 | 125041.0 | 30338.6 |
| VirGenA | 764.6 | 32.0 | 99.901 | 0.549 | 0.518 | 4.891 | 727.667 | 72.0 | 70380.0 | 115.0 | 2888.0 | 977.5 |
| ViSpA | 138.655 | 168.0 | 99.883 | 0.034 | 0.024 | 5.0 | 99.0 | 5.0 | 143673.0 | 0.0 | 124983.0 | 28734.6 |
| V-pipe | 45.1 | 141.0 | 99.684 | 0.043 | 0.032 | 0.104 | 114.0 | 5.0 | 160429.0 | 5387.0 | 125041.0 | 32085.8 |

Table S60: Results obtained for DS58 using the benchmark proposed. The execution time was measured in seconds, the RAM usage was measured in GB and the CPU usage is presented as a percentage. The executions were, when possible, capped at 6 threads and 48 GB of RAM.

| Reconstruction tool | Execution time | SNPs | Identity | NCSD | NRC | RAM usage | CPU usage | Number of scaffolds | Reconstructed bases | Minimum scaffold length | Maximum scaffold length | Average scaffold length |
|---|---|---|---|---|---|---|---|---|---|---|---|---|
| coronaSPAdes | 2.63 | 62.0 | 99.908 | 0.036 | 0.025 | 0.147 | 407.333 | 10.0 | 153918.0 | 81.0 | 105120.0 | 15391.8 |
| Haploflow | 2.67 | 94.0 | 99.914 | 0.069 | 0.053 | 0.076 | 99.333 | 40.0 | 160635.0 | 542.0 | 18358.0 | 4015.9 |
| IRMA | 162.62 | 0.667 | 99.999 | 0.095 | 0.127 | 0.874 | 508.333 | 5.0 | 144910.0 | 4882.0 | 110266.0 | 28982.0 |
| LAZYPIPE | 6.105 | 71.333 | 99.939 | 0.037 | 0.027 | 1.519 | 247.333 | 7.0 | 152818.0 | 5318.0 | 96717.0 | 21831.1 |
| metaSPAdes | 4.695 | 33.0 | 99.84 | 0.037 | 0.026 | 0.226 | 401.333 | 11.0 | 153524.0 | 71.0 | 105136.0 | 13956.7 |
| metaviralSPAdes | – | – | – | – | – | – | – | – | – | – | – | – |
| PEHaplo | 26.01 | 58.0 | 99.958 | 0.094 | 0.077 | 0.882 | 128.333 | 224.0 | 186980.0 | 378.0 | 3000.0 | 834.7 |
| QuRe | 427.43 | 3.333 | 99.789 | 0.995 | 0.983 | 21.496 | 540.333 | 7.0 | 2176.667 | 55.0 | 617.667 | 310.933 |
| QVG | 23.075 | 168.0 | 99.881 | 0.032 | 0.024 | 1.406 | 143.667 | 5.0 | 268901.0 | 5387.0 | 125041.0 | 53780.2 |
| SPAdes | 6.175 | 50.0 | 99.896 | 0.036 | 0.025 | 0.161 | 343.0 | 22.0 | 161709.0 | 131.0 | 105080.0 | 7350.4 |
| SSAKE | 4.31 | 12.0 | 99.98 | 0.507 | 0.468 | 0.067 | 100.0 | 309.0 | 92565.0 | 102.333 | 1256.667 | 299.6 |
| TRACESPipe | 168.845 | 174.0 | 99.877 | 0.033 | 0.023 | 9.768 | 366.0 | 4.0 | 143860.0 | 5387.0 | 125041.0 | 35965.0 |
| TRACESPipeLite | 42.845 | 141.333 | 99.903 | 0.033 | 0.023 | 2.782 | 327.0 | 6.0 | 168262.0 | 5387.0 | 125041.0 | 28043.7 |
| VirGenA | 674.175 | 105.333 | 99.88 | 0.301 | 0.277 | 4.426 | 718.667 | 59.0 | 107721.667 | 233.0 | 7460.667 | 2014.267 |
| ViSpA | 155.465 | 188.0 | 99.867 | 0.035 | 0.025 | 6.154 | 107.333 | 5.0 | 160133.0 | 5318.0 | 125016.0 | 32026.6 |
| V-pipe | 42.7 | 148.0 | 99.74 | 0.044 | 0.033 | 0.104 | 114.667 | 5.0 | 160429.0 | 5387.0 | 125041.0 | 32085.8 |

Table S61: Results obtained for DS59 using the benchmark proposed. The execution time was measured in seconds, the RAM usage was measured in GB and the CPU usage is presented as a percentage. The executions were, when possible, capped at 6 threads and 48 GB of RAM.

| Reconstruction tool | Execution time | SNPs | Identity | NCSD | NRC | RAM usage | CPU usage | Number of scaffolds | Reconstructed bases | Minimum scaffold length | Maximum scaffold length | Average scaffold length |
|---|---|---|---|---|---|---|---|---|---|---|---|---|
| coronaSPAdes | 3.875 | 46.0 | 99.967 | 0.037 | 0.026 | 0.164 | 435.0 | 12.0 | 207916.0 | 531.0 | 105629.0 | 17326.3 |
| Haploflow | 4.99 | 457.0 | 99.66 | 0.052 | 0.04 | 0.126 | 99.333 | 16.0 | 211700.0 | 1680.0 | 45394.0 | 13231.3 |
| IRMA | 279.805 | 1.0 | 99.999 | 0.103 | 0.128 | 1.261 | 579.0 | 5.0 | 144874.0 | 4888.0 | 110252.0 | 28974.8 |
| LAZYPIPE | 8.545 | 1.0 | 99.963 | 0.032 | 0.022 | 1.533 | 265.667 | 8.0 | 210190.0 | 4291.0 | 105465.0 | 26273.8 |
| metaSPAdes | 7.23 | 76.0 | 99.946 | 0.033 | 0.023 | 0.229 | 429.333 | 7.0 | 210082.0 | 328.0 | 124464.0 | 30011.7 |
| metaviralSPAdes | – | – | – | – | – | – | – | – | – | – | – | – |
| PEHaplo | 53.755 | 132.0 | 99.886 | 0.055 | 0.043 | 1.981 | 145.333 | 301.0 | 314300.0 | 381.0 | 3694.0 | 1044.2 |
| QuRe | 1377.655 | 21.0 | 99.51 | 0.976 | 0.954 | 21.302 | 610.667 | 5.0 | 6075.0 | 212.667 | 3038.667 | 1163.5 |
| QVG | 32.245 | 503.0 | 99.573 | 0.045 | 0.035 | 1.413 | 162.0 | 4.0 | 143860.0 | 5387.0 | 125041.0 | 35965.0 |
| SPAdes | 7.705 | 2.0 | 99.998 | 0.031 | 0.021 | 0.174 | 411.667 | 6.0 | 210332.0 | 5371.0 | 125032.0 | 35055.3 |
| SSAKE | 14.73 | 1.333 | 99.985 | 0.082 | 0.064 | 0.116 | 100.0 | 409.667 | 220053.333 | 101.333 | 3104.333 | 537.167 |
| TRACESPipe | 180.345 | 498.0 | 99.653 | 0.039 | 0.028 | 9.768 | 363.0 | 4.0 | 143859.0 | 5387.0 | 125040.0 | 35964.8 |
| TRACESPipeLite | 50.66 | 473.0 | 99.503 | 0.055 | 0.044 | 2.825 | 315.0 | 6.0 | 168262.0 | 5387.0 | 125041.0 | 28043.7 |
| VirGenA | 976.155 | 96.0 | 99.713 | 0.706 | 0.688 | 4.682 | 744.0 | 53.0 | 46445.0 | 170.0 | 2247.0 | 876.3 |
| ViSpA | 276.105 | 610.0 | 99.564 | 0.042 | 0.032 | 6.875 | 123.0 | 5.0 | 160186.0 | 5342.0 | 125007.0 | 32037.2 |
| V-pipe | 55.975 | 454.0 | 99.683 | 0.042 | 0.031 | 0.104 | 113.0 | 5.0 | 160429.0 | 5387.0 | 125041.0 | 32085.8 |

Table S62: Results obtained for DS60 using the benchmark proposed. The execution time was measured in seconds, the RAM usage was measured in GB and the CPU usage is presented as a percentage. The executions were, when possible, capped at 6 threads and 48 GB of RAM.

| Reconstruction tool | Execution time | SNPs | Identity | NCSD | NRC | RAM usage | CPU usage | Number of scaffolds | Reconstructed bases | Minimum scaffold length | Maximum scaffold length | Average scaffold length |
|---|---|---|---|---|---|---|---|---|---|---|---|---|
| coronaSPAdes | 4.54 | 0.0 | 100.0 | 0.03 | 0.021 | 0.153 | 437.333 | 6.0 | 210367.0 | 5374.0 | 125037.0 | 35061.2 |
| Haploflow | 6.56 | 206.0 | 99.844 | 0.032 | 0.023 | 0.152 | 99.0 | 8.0 | 211720.0 | 5095.0 | 115022.0 | 26465.0 |
| IRMA | 364.3 | 0.0 | 100.0 | 0.108 | 0.128 | 1.639 | 594.667 | 5.0 | 144923.0 | 4892.0 | 110283.0 | 28984.6 |
| LAZYPIPE | 8.9 | 0.0 | 99.976 | 0.031 | 0.022 | 1.54 | 256.0 | 7.0 | 210287.0 | 351.0 | 124743.0 | 30041.0 |
| metaSPAdes | 8.62 | 0.0 | 100.0 | 0.03 | 0.021 | 0.228 | 433.0 | 6.0 | 210367.0 | 5374.0 | 125037.0 | 35061.2 |
| metaviralSPAdes | 9.695 | 0.0 | 100.0 | 0.162 | 0.149 | 0.226 | 450.667 | 3.0 | 191597.0 | 16565.0 | 125037.0 | 63865.7 |
| PEHaplo | 78.54 | 146.0 | 99.894 | 0.032 | 0.024 | 2.94 | 149.667 | 218.0 | 334264.0 | 411.0 | 4869.0 | 1533.3 |
| QuRe | 2492.145 | 6.333 | 99.863 | 0.978 | 0.97 | 22.494 | 632.0 | 4.333 | 5290.667 | 365.0 | 2446.0 | 1215.2 |
| QVG | 42.885 | 736.0 | 99.488 | 0.047 | 0.04 | 1.419 | 173.333 | 5.0 | 268901.0 | 5387.0 | 125041.0 | 53780.2 |
| SPAdes | 9.33 | 0.0 | 100.0 | 0.03 | 0.021 | 0.185 | 427.333 | 6.0 | 210367.0 | 5374.0 | 125037.0 | 35061.2 |
| SSAKE | 17.125 | 0.333 | 99.998 | 0.042 | 0.032 | 0.134 | 100.0 | 175.0 | 224017.667 | 115.333 | 8673.0 | 1280.467 |
| TRACESPipe | 191.88 | 689.0 | 99.51 | 0.044 | 0.035 | 9.768 | 361.333 | 4.0 | 143867.0 | 5387.0 | 125046.0 | 35966.8 |
| TRACESPipeLite | 50.12 | 218.667 | 97.556 | 0.273 | 0.246 | 2.829 | 309.0 | 6.0 | 168263.0 | 5387.0 | 125041.0 | 28043.8 |
| VirGenA | 1022.68 | 49.0 | 99.685 | 0.711 | 0.69 | 5.317 | 746.0 | 52.0 | 46643.0 | 61.0 | 2554.0 | 897.0 |
| ViSpA | 438.595 | 784.0 | 99.413 | 0.051 | 0.04 | 6.925 | 128.0 | 5.0 | 160360.0 | 5360.0 | 125072.0 | 32072.0 |
| V-pipe | 65.25 | 586.0 | 99.59 | 0.042 | 0.033 | 0.104 | 112.667 | 5.0 | 160429.0 | 5387.0 | 125041.0 | 32085.8 |

Table S63: Results obtained for DS61 using the benchmark proposed. The execution time was measured in seconds, the RAM usage was measured in GB and the CPU usage is presented as a percentage. The executions were, when possible, capped at 6 threads and 48 GB of RAM.

| Reconstruction tool | Execution time | SNPs | Identity | NCSD | NRC | RAM usage | CPU usage | Number of scaffolds | Reconstructed bases | Minimum scaffold length | Maximum scaffold length | Average scaffold length |
|---|---|---|---|---|---|---|---|---|---|---|---|---|
| coronaSPAdes | 6.255 | 219.0 | 99.732 | 0.047 | 0.031 | 0.181 | 466.0 | 18.0 | 202859.0 | 51.0 | 84751.0 | 11269.9 |
| Haploflow | – | – | – | – | – | – | – | – | – | – | – | – |
| IRMA | – | – | – | – | – | – | – | – | – | – | – | – |
| LAZYPIPE | 5.09 | 504.0 | 99.384 | 0.406 | 0.367 | 1.521 | 236.667 | 236.0 | 139792.0 | 237.0 | 2358.0 | 592.3 |
| metaSPAdes | 6.09 | 266.0 | 99.773 | 0.066 | 0.049 | 0.233 | 441.0 | 72.0 | 204176.0 | 97.0 | 34311.0 | 2835.8 |
| metaviralSPAdes | – | – | – | – | – | – | – | – | – | – | – | – |
| PEHaplo | 42.93 | 51.0 | 99.931 | 0.349 | 0.322 | 1.725 | 124.667 | 273.0 | 160072.0 | 375.0 | 1933.0 | 586.3 |
| QuRe | 601.72 | 0.667 | 99.888 | 0.999 | 0.992 | 15.131 | 585.333 | 5.0 | 507.667 | 85.0 | 124.0 | 104.9 |
| QVG | 21.305 | 1203.0 | 99.166 | 0.074 | 0.054 | 1.406 | 134.667 | 4.0 | 143860.0 | 5387.0 | 125041.0 | 35965.0 |
| SPAdes | 10.175 | 133.0 | 99.844 | 0.042 | 0.028 | 0.337 | 442.333 | 17.0 | 207875.0 | 64.0 | 64084.0 | 12227.9 |
| SSAKE | – | – | – | – | – | – | – | – | – | – | – | – |
| TRACESPipe | 167.105 | 153.0 | 99.896 | 0.033 | 0.023 | 9.768 | 365.333 | 4.0 | 143860.0 | 5387.0 | 125041.0 | 35965.0 |
| TRACESPipeLite | 31.71 | 8.667 | 98.183 | 0.998 | 0.987 | 2.532 | 381.333 | 5.0 | 160560.0 | 5185.0 | 125374.0 | 32112.0 |
| VirGenA | – | – | – | – | – | – | – | – | – | – | – | – |
| ViSpA | 290.76 | 70.333 | 99.565 | 0.05 | 0.035 | 7.382 | 115.667 | 5.0 | 431876.0 | 5375.0 | 396401.0 | 86375.2 |
| V-pipe | – | – | – | – | – | – | – | – | – | – | – | – |

Table S64: Results obtained for DS62 using the benchmark proposed. The execution time was measured in seconds, the RAM usage was measured in GB and the CPU usage is presented as a percentage. The executions were, when possible, capped at 6 threads and 48 GB of RAM.

| Reconstruction tool | Execution time | SNPs | Identity | NCSD | NRC | RAM usage | CPU usage | Number of scaffolds | Reconstructed bases | Minimum scaffold length | Maximum scaffold length | Average scaffold length |
|---|---|---|---|---|---|---|---|---|---|---|---|---|
| coronaSPAdes | 7.12 | 35.0 | 99.969 | 0.036 | 0.025 | 0.155 | 443.333 | 12.0 | 205320.0 | 468.0 | 107424.0 | 17110.0 |
| Haploflow | 6.84 | 65.0 | 99.91 | 0.19 | 0.168 | 0.251 | 99.0 | 111.0 | 180647.0 | 513.0 | 5061.0 | 1627.5 |
| IRMA | 136.635 | 1.0 | 99.999 | 0.096 | 0.128 | 0.581 | 534.0 | 5.0 | 144840.0 | 4888.0 | 110225.0 | 28968.0 |
| LAZYPIPE | 7.12 | 53.0 | 99.901 | 0.036 | 0.025 | 1.524 | 288.667 | 9.0 | 204459.0 | 1010.0 | 109941.0 | 22717.7 |
| metaSPAdes | 5.885 | 50.0 | 99.855 | 0.037 | 0.026 | 0.227 | 422.0 | 14.0 | 203516.0 | 71.0 | 105176.0 | 14536.9 |
| metaviralSPAdes | 7.26 | 56.0 | 99.953 | 0.179 | 0.159 | 0.228 | 444.667 | 4.0 | 183652.0 | 5358.0 | 111885.0 | 45913.0 |
| PEHaplo | 27.67 | 49.0 | 99.956 | 0.054 | 0.042 | 0.806 | 152.0 | 214.0 | 271186.0 | 394.0 | 6577.0 | 1267.2 |
| QuRe | 792.355 | 0.0 | 100.0 | 0.997 | 0.987 | 42.713 | 534.667 | 2.333 | 982.333 | 249.0 | 523.0 | 404.0 |
| QVG | 23.81 | 151.0 | 99.891 | 0.032 | 0.024 | 1.406 | 154.667 | 5.0 | 268901.0 | 5387.0 | 125041.0 | 53780.2 |
| SPAdes | 10.295 | 33.0 | 99.98 | 0.032 | 0.022 | 0.196 | 393.333 | 15.0 | 209243.0 | 355.0 | 106624.0 | 13949.5 |
| SSAKE | 2.175 | 2.0 | 99.987 | 0.918 | 0.898 | 0.045 | 100.0 | 108.0 | 22069.333 | 103.0 | 518.333 | 204.333 |
| TRACESPipe | 172.825 | 154.0 | 99.894 | 0.034 | 0.024 | 9.768 | 364.667 | 4.0 | 143860.0 | 5387.0 | 125041.0 | 35965.0 |
| TRACESPipeLite | 59.39 | 127.667 | 99.91 | 0.033 | 0.024 | 2.84 | 350.0 | 6.0 | 168262.0 | 5387.0 | 125041.0 | 28043.7 |
| VirGenA | 919.355 | 55.0 | 99.844 | 0.756 | 0.734 | 3.695 | 727.0 | 28.0 | 37394.0 | 682.0 | 2955.0 | 1335.5 |
| ViSpA | 73.055 | 256.0 | 99.815 | 0.04 | 0.027 | 5.777 | 108.0 | 5.0 | 160120.0 | 5355.0 | 124967.0 | 32024.0 |
| V-pipe | 47.08 | 134.0 | 99.715 | 0.052 | 0.038 | 0.104 | 112.333 | 5.0 | 160429.0 | 5387.0 | 125041.0 | 32085.8 |

Table S65: Results obtained for DS63 using the benchmark proposed. The execution time was measured in seconds, the RAM usage was measured in GB and the CPU usage is presented as a percentage. The executions were, when possible, capped at 6 threads and 48 GB of RAM.

| Reconstruction tool | Execution time | SNPs | Identity | NCSD | NRC | RAM usage | CPU usage | Number of scaffolds | Reconstructed bases | Minimum scaffold length | Maximum scaffold length | Average scaffold length |
|---|---|---|---|---|---|---|---|---|---|---|---|---|
| coronaSPAdes | 5.115 | 126.0 | 99.957 | 0.027 | 0.018 | 0.177 | 437.333 | 17.0 | 358134.0 | 79.0 | 125432.0 | 21066.7 |
| Haploflow | 6.62 | 217.0 | 99.888 | 0.04 | 0.031 | 0.174 | 99.0 | 89.0 | 373224.0 | 521.0 | 21945.0 | 4193.5 |
| IRMA | 824.055 | 97.0 | 99.921 | 0.056 | 0.068 | 1.715 | 610.0 | 6.0 | 298596.333 | 4877.0 | 153774.333 | 49766.067 |
| LAZYPIPE | 10.14 | 152.0 | 99.854 | 0.027 | 0.018 | 1.539 | 281.0 | 14.0 | 356563.0 | 350.0 | 131634.0 | 25468.8 |
| metaSPAdes | 9.305 | 113.0 | 99.868 | 0.027 | 0.019 | 0.233 | 437.667 | 29.0 | 358115.0 | 57.0 | 145112.0 | 12348.8 |
| metaviralSPAdes | – | – | – | – | – | – | – | – | – | – | – | – |
| PEHaplo | 192.015 | 242.0 | 99.895 | 0.045 | 0.037 | 1.867 | 120.0 | 535.0 | 486718.0 | 376.0 | 4058.0 | 909.8 |
| QuRe | 2122.53 | 11.0 | 99.367 | 0.996 | 0.975 | 20.128 | 623.667 | 14.333 | 4255.333 | 71.0 | 907.0 | 296.467 |
| QVG | 45.55 | 335.667 | 99.881 | 0.024 | 0.019 | 1.419 | 168.0 | 7.0 | 593129.0 | 5387.0 | 162114.0 | 84732.7 |
| SPAdes | 13.14 | 54.0 | 99.939 | 0.025 | 0.017 | 0.188 | 321.667 | 53.0 | 370365.0 | 131.0 | 121039.0 | 6988.0 |
| SSAKE | 7.265 | 14.667 | 99.968 | 0.671 | 0.641 | 0.109 | 99.333 | 416.0 | 134437.0 | 100.667 | 1270.667 | 322.9 |
| TRACESPipe | 255.81 | 336.0 | 99.89 | 0.022 | 0.016 | 9.768 | 300.333 | 7.0 | 624802.0 | 5387.0 | 162114.0 | 89257.4 |
| TRACESPipeLite | 55.815 | 303.333 | 99.894 | 0.024 | 0.016 | 2.844 | 289.0 | 7.0 | 330376.0 | 5387.0 | 162114.0 | 47196.6 |
| VirGenA | 2002.26 | 52.333 | 99.877 | 0.763 | 0.742 | 5.45 | 739.0 | 76.667 | 76151.0 | 135.333 | 3894.0 | 996.9 |
| ViSpA | 321.31 | 275.0 | 99.873 | 0.023 | 0.017 | 6.139 | 124.0 | 12.0 | 747730.0 | 346.0 | 153863.0 | 62310.8 |
| V-pipe | 74.32 | 321.0 | 99.756 | 0.033 | 0.025 | 0.111 | 113.667 | 6.0 | 322543.0 | 5387.0 | 162114.0 | 53757.2 |

Table S66: Results obtained for DS64 using the benchmark proposed. The execution time was measured in seconds, the RAM usage was measured in GB and the CPU usage is presented as a percentage. The executions were, when possible, capped at 6 threads and 48 GB of RAM.

| Reconstruction tool | Execution time | SNPs | Identity | NCSD | NRC | RAM usage | CPU usage | Number of scaffolds | Reconstructed bases | Minimum scaffold length | Maximum scaffold length | Average scaffold length |
|---|---|---|---|---|---|---|---|---|---|---|---|---|
| coronaSPAdes | 5.185 | 81.0 | 99.821 | 0.03 | 0.02 | 0.17 | 446.0 | 22.0 | 351746.0 | 81.0 | 105120.0 | 15988.5 |
| Haploflow | 6.725 | 203.0 | 99.897 | 0.048 | 0.037 | 0.177 | 99.0 | 89.0 | 368848.0 | 524.0 | 19368.0 | 4144.4 |
| IRMA | 924.655 | 61.0 | 99.786 | 0.063 | 0.073 | 1.79 | 616.667 | 6.0 | 297266.333 | 4863.0 | 152903.333 | 49544.4 |
| LAZYPIPE | 10.56 | 131.0 | 99.929 | 0.029 | 0.023 | 1.539 | 293.667 | 32.0 | 354981.0 | 273.0 | 92612.0 | 11093.2 |
| metaSPAdes | 9.575 | 92.0 | 99.893 | 0.031 | 0.021 | 0.233 | 444.0 | 17.0 | 350428.0 | 71.0 | 105234.0 | 20613.4 |
| metaviralSPAdes | 11.27 | 2.0 | 99.96 | 0.985 | 0.968 | 0.232 | 457.333 | 1.0 | 5358.0 | 5358.0 | 5358.0 | 5358.0 |
| PEHaplo | 70.365 | 142.0 | 99.952 | 0.053 | 0.041 | 2.036 | 138.333 | 524.0 | 472293.0 | 376.0 | 4078.0 | 901.3 |
| QuRe | 1708.795 | 8.0 | 99.632 | 0.997 | 0.971 | 16.215 | 632.0 | 10.0 | 2621.667 | 131.667 | 460.333 | 260.267 |
| QVG | 43.555 | 429.667 | 99.857 | 0.029 | 0.02 | 1.417 | 167.667 | 6.0 | 441230.0 | 4952.0 | 172764.0 | 73538.3 |
| SPAdes | 15.89 | 48.0 | 99.846 | 0.029 | 0.02 | 0.187 | 312.667 | 62.0 | 364614.0 | 80.0 | 105128.0 | 5880.9 |
| SSAKE | 6.665 | 29.667 | 99.967 | 0.692 | 0.619 | 0.109 | 100.0 | 373.333 | 122899.0 | 102.333 | 1390.0 | 328.833 |
| TRACESPipe | 217.615 | 398.0 | 99.866 | 0.027 | 0.018 | 9.768 | 331.667 | 6.0 | 321125.0 | 4935.0 | 172764.0 | 53520.8 |
| TRACESPipeLite | 59.8 | 391.667 | 99.868 | 0.027 | 0.019 | 2.843 | 283.0 | 7.0 | 340591.0 | 4952.0 | 172764.0 | 48655.9 |
| VirGenA | 2028.57 | 32.333 | 99.85 | 0.875 | 0.845 | 5.204 | 719.667 | 39.0 | 39978.333 | 230.0 | 2755.0 | 944.7 |
| ViSpA | 241.025 | 231.667 | 99.901 | 0.025 | 0.019 | 5.383 | 121.667 | 13.0 | 851902.0 | 4935.0 | 148680.0 | 65530.9 |
| V-pipe | 77.81 | 434.0 | 99.686 | 0.038 | 0.027 | 0.113 | 112.667 | 6.0 | 332758.0 | 4952.0 | 172764.0 | 55459.7 |

Table S67: Results obtained for DS65 using the benchmark proposed. The execution time was measured in seconds, the RAM usage was measured in GB and the CPU usage is presented as a percentage. The executions were, when possible, capped at 6 threads and 48 GB of RAM.

| Reconstruction tool | Execution time | SNPs | Identity | NCSD | NRC | RAM usage | CPU usage | Number of scaffolds | Reconstructed bases | Minimum scaffold length | Maximum scaffold length | Average scaffold length |
|---|---|---|---|---|---|---|---|---|---|---|---|---|
| coronaSPAdes | 5.685 | 61.0 | 99.963 | 0.02 | 0.014 | 0.173 | 452.667 | 16.0 | 435772.0 | 81.0 | 193790.0 | 27235.8 |
| Haploflow | 7.96 | 63.0 | 99.958 | 0.039 | 0.031 | 0.208 | 99.333 | 113.0 | 448894.0 | 544.0 | 18022.0 | 3972.5 |
| IRMA | 690.105 | 0.667 | 100.0 | 0.051 | 0.071 | 3.483 | 613.667 | 6.0 | 373222.0 | 4881.0 | 228347.0 | 62203.7 |
| LAZYPIPE | 11.34 | 70.0 | 99.968 | 0.02 | 0.014 | 1.543 | 285.333 | 10.0 | 435025.0 | 5372.0 | 196252.0 | 43502.5 |
| metaSPAdes | 10.91 | 40.0 | 99.919 | 0.02 | 0.014 | 0.234 | 442.333 | 15.0 | 434957.0 | 71.0 | 192881.0 | 28997.1 |
| metaviralSPAdes | – | – | – | – | – | – | – | – | – | – | – | – |
| PEHaplo | 60.06 | 86.0 | 99.974 | 0.039 | 0.033 | 2.059 | 154.333 | 592.0 | 587990.0 | 378.0 | 4705.0 | 993.2 |
| QuRe | 1598.62 | 3.667 | 99.636 | 0.999 | 0.984 | 15.896 | 634.667 | 8.0 | 1648.667 | 105.333 | 341.0 | 217.2 |
| QVG | 48.065 | 221.667 | 99.941 | 0.02 | 0.014 | 1.42 | 172.0 | 6.0 | 504230.0 | 5387.0 | 235329.0 | 84038.3 |
| SPAdes | 11.665 | 33.0 | 99.97 | 0.019 | 0.014 | 0.202 | 421.667 | 34.0 | 445057.0 | 167.0 | 192883.0 | 13089.9 |
| SSAKE | 8.3 | 17.667 | 99.981 | 0.688 | 0.665 | 0.124 | 100.0 | 454.0 | 153953.667 | 102.0 | 1281.0 | 338.967 |
| TRACESPipe | 307.15 | 196.0 | 99.951 | 0.018 | 0.013 | 9.768 | 262.0 | 5.0 | 379189.0 | 5387.0 | 235329.0 | 75837.8 |
| TRACESPipeLite | 63.295 | 189.667 | 99.951 | 0.018 | 0.013 | 2.847 | 280.333 | 7.0 | 403590.0 | 5387.0 | 235329.0 | 57655.7 |
| VirGenA | 2023.705 | 2.0 | 99.964 | 0.989 | 0.975 | 6.078 | 747.0 | 6.0 | 5797.0 | 617.0 | 1716.0 | 966.2 |
| ViSpA | 670.29 | 232.0 | 99.94 | 0.019 | 0.013 | 4.608 | 120.0 | 6.0 | 395003.0 | 5347.0 | 234813.0 | 65833.8 |
| V-pipe | 82.375 | 195.0 | 99.805 | 0.029 | 0.022 | 0.128 | 113.333 | 6.0 | 395758.0 | 5387.0 | 235329.0 | 65959.7 |

Table S68: Results obtained by dnadiff [4] when analysing the reconstruction of DS1. Only one execution of HVRS is represented.

| Reconstruction tool | Total seqs | Aligned seqs | Aligned bases | Aligned bases (%) | Unaligned bases | Unaligned bases (%) | Average length | SNPs | Identity |
|---|---|---|---|---|---|---|---|---|---|
| coronaSPAdes | 182 | 182 | 83085 | 99.9976 | 2 | 0.0024 | 455.1530 | 131 | 99,824 |
| Haploflow | – | – | – | – | – | – | – | – | – |
| IRMA | 4 | 4 | 95726 | 99.9958 | 4 | 0.0042 | 712.6889 | 99 | 96,085 |
| LAZYPIPE | 152 | 152 | 65304 | 99.9954 | 3 | 0.0046 | 425.7403 | 64 | 99,881 |
| metaSPAdes | 210 | 210 | 88861 | 99.9977 | 2 | 0.0023 | 423.1476 | 128 | 99,854 |
| metaviralSPAdes | – | – | – | – | – | – | – | – | – |
| PEHaplo | 98 | 98 | 58459 | 99.9949 | 3 | 0.0051 | 590.3500 | 672 | 99,749 |
| QuRe | – | – | – | – | – | – | – | – | – |
| QVG | 3 | 3 | 138473 | 100.0000 | 0 | 0.0000 | 46157.6667 | 1301 | 99,064 |
| SPAdes | 179 | 179 | 82066 | 99.9976 | 2 | 0.0024 | 457.0778 | 104 | 99,858 |
| SSAKE | 2 | 2 | 248 | 100.0000 | 0 | 0.0000 | 124.0000 | 0 | 100,000 |
| TRACESPipe | 4 | 4 | 113776 | 79.0880 | 30084 | 20.9120 | 904.0250 | 100 | 97,349 |
| TRACESPipeLite | 5 | 4 | 112549 | 74.1952 | 39144 | 25.8048 | 848.6031 | 66 | 97,028 |
| V-pipe | 5 | 3 | 5704 | 3.5555 | 154725 | 96.4445 | 139.1220 | 0 | 95,791 |
| VirGenA | 238 | 237 | 85972 | 99.9314 | 59 | 0.0686 | 367.6026 | 88 | 99,884 |
| ViSpA | 75 | 74 | 2289898 | 99.8929 | 2454 | 0.1071 | 326.3165 | 94 | 95,123 |

Table S69: Results obtained by dnadiff [4] when analysing the reconstruction of DS2. Only one execution of HVRS is represented.

| Reconstruction tool | Total seqs | Aligned seqs | Aligned bases | Aligned bases (%) | Unaligned bases | Unaligned bases (%) | Average length | SNPs | Identity |
|---|---|---|---|---|---|---|---|---|---|
| coronaSPAdes | 100 | 100 | 129534 | 99.9954 | 6 | 0.0046 | 1295.2647 | 120 | 99,775 |
| Haploflow | – | – | – | – | – | – | – | – | – |
| IRMA | 4 | 4 | 122405 | 100.0000 | 0 | 0.0000 | 4372.4643 | 49 | 98,394 |
| LAZYPIPE | 92 | 92 | 121346 | 99.9959 | 5 | 0.0041 | 1291.8723 | 114 | 99,896 |
| metaSPAdes | 104 | 104 | 130237 | 99.9954 | 6 | 0.0046 | 1244.3883 | 156 | 99,721 |
| metaviralSPAdes | – | – | – | – | – | – | – | – | – |
| PEHaplo | 102 | 102 | 127508 | 99.9969 | 4 | 0.0031 | 1250.0784 | 114 | 99,877 |
| QuRe | – | – | – | – | – | – | – | – | – |
| QVG | 4 | 4 | 143858 | 99.9986 | 2 | 0.0014 | 35964.5000 | 955 | 99,333 |
| SPAdes | 101 | 101 | 129739 | 99.9214 | 102 | 0.0786 | 1292.6600 | 109 | 99,909 |
| SSAKE | 55 | 55 | 8469 | 100.0000 | 0 | 0.0000 | 153.9818 | 3 | 99,965 |
| TRACESPipe | 4 | 4 | 141490 | 98.3526 | 2370 | 1.6474 | 7860.5556 | 135 | 99,060 |
| TRACESPipeLite | 5 | 4 | 140020 | 92.3049 | 11673 | 7.6951 | 7000.8500 | 153 | 98,787 |
| V-pipe | 5 | 4 | 68492 | 42.6930 | 91937 | 57.3070 | 317.7523 | 24 | 94,134 |
| VirGenA | 114 | 113 | 108795 | 99.9458 | 59 | 0.0542 | 928.0348 | 97 | 99,881 |
| ViSpA | 80 | 79 | 3611375 | 99.9794 | 745 | 0.0206 | 1894.4265 | 137 | 97,596 |

Table S70: Results obtained by dnadiff [4] when analysing the reconstruction of DS3. Only one execution of HVRS is represented.

| Reconstruction tool | Total seqs | Aligned seqs | Aligned bases | Aligned bases (%) | Unaligned bases | Unaligned bases (%) | Average length | SNPs | Identity |
|---|---|---|---|---|---|---|---|---|---|
| coronaSPAdes | 21 | 21 | 138037 | 100.0000 | 0 | 0.0000 | 7628.9444 | 39 | 99,868 |
| Haploflow | 65 | 65 | 52460 | 100.0000 | 0 | 0.0000 | 802.8254 | 34 | 99,882 |
| IRMA | 4 | 4 | 127894 | 100.0000 | 0 | 0.0000 | 25579.6000 | 4 | 99,921 |
| LAZYPIPE | 19 | 19 | 135584 | 100.0000 | 0 | 0.0000 | 6793.4000 | 60 | 99,950 |
| metaSPAdes | 17 | 17 | 136529 | 99.9268 | 100 | 0.0732 | 8005.9412 | 60 | 99,813 |
| metaviralSPAdes | – | – | – | – | – | – | – | – | – |
| PEHaplo | 116 | 116 | 146720 | 100.0000 | 0 | 0.0000 | 1260.0345 | 55 | 99,931 |
| QuRe | 3 | 2 | 192 | 66.8990 | 95 | 33.1010 | 96.0000 | 0 | 100,000 |
| QVG | 4 | 4 | 143858 | 99.9986 | 2 | 0.0014 | 35964.5000 | 452 | 99,685 |
| SPAdes | 27 | 27 | 143582 | 99.9304 | 100 | 0.0696 | 5126.0000 | 37 | 99,926 |
| SSAKE | 114 | 114 | 19911 | 100.0000 | 0 | 0.0000 | 174.6579 | 4 | 99,980 |
| TRACESPipe | 4 | 4 | 143518 | 99.7623 | 342 | 0.2377 | 35879.5000 | 152 | 99,850 |
| TRACESPipeLite | 5 | 5 | 143881 | 94.8501 | 7812 | 5.1499 | 35879.5000 | 126 | 99,869 |
| V-pipe | 5 | 4 | 126205 | 78.6672 | 34224 | 21.3328 | 1225.0101 | 46 | 97,085 |
| VirGenA | 64 | 63 | 114363 | 99.8952 | 120 | 0.1048 | 1549.3243 | 66 | 99,864 |
| ViSpA | 9 | 8 | 483660 | 100.0000 | 0 | 0.0000 | 25607.0000 | 132 | 99,514 |

Table S71: Results obtained by dnadiff [4] when analysing the reconstruction of DS4. Only one execution of HVRS is represented.

| Reconstruction tool | Total seqs | Aligned seqs | Aligned bases | Aligned bases (%) | Unaligned bases | Unaligned bases (%) | Average length | SNPs | Identity |
|---|---|---|---|---|---|---|---|---|---|
| coronaSPAdes | 14 | 14 | 138001 | 99.9276 | 100 | 0.0724 | 9851.4286 | 74 | 99,837 |
| Haploflow | 82 | 82 | 126008 | 100.0000 | 0 | 0.0000 | 1503.9500 | 68 | 99,908 |
| IRMA | 4 | 4 | 128279 | 100.0000 | 0 | 0.0000 | 25655.8000 | 1 | 99,992 |
| LAZYPIPE | 8 | 8 | 136217 | 100.0000 | 0 | 0.0000 | 17011.5000 | 72 | 99,889 |
| metaSPAdes | 8 | 8 | 136315 | 99.9282 | 98 | 0.0718 | 12383.2727 | 73 | 99,899 |
| metaviralSPAdes | 1 | 1 | 5358 | 100.0000 | 0 | 0.0000 | 5358.0000 | 1 | 99,940 |
| PEHaplo | 180 | 180 | 165859 | 99.9988 | 2 | 0.0012 | 931.3077 | 86 | 99,935 |
| QuRe | 7 | 6 | 620 | 87.2011 | 91 | 12.7989 | 108.2000 | 0 | 100,000 |
| QVG | 4 | 4 | 143858 | 99.9986 | 2 | 0.0014 | 35964.5000 | 197 | 99,859 |
| SPAdes | 7 | 7 | 137172 | 99.8559 | 198 | 0.1441 | 15257.6667 | 68 | 99,791 |
| SSAKE | 174 | 174 | 37612 | 99.9973 | 1 | 0.0027 | 216.1609 | 6 | 99,984 |
| TRACESPipe | 4 | 4 | 143701 | 99.8895 | 159 | 0.1105 | 35925.2500 | 111 | 99,920 |
| TRACESPipeLite | 5 | 5 | 143953 | 94.8976 | 7740 | 5.1024 | 35936.2500 | 128 | 99,910 |
| V-pipe | 5 | 4 | 142613 | 88.8948 | 17816 | 11.1052 | 15845.8889 | 151 | 99,490 |
| VirGenA | 50 | 50 | 114754 | 99.9573 | 49 | 0.0427 | 2028.7544 | 98 | 99,831 |
| ViSpA | 5 | 4 | 143528 | 100.0000 | 0 | 0.0000 | 30096.6000 | 162 | 99,782 |

Table S72: Results obtained by dnadiff [4] when analysing the reconstruction of DS5. Only one execution of HVRS is represented.

| Reconstruction tool | Total seqs | Aligned seqs | Aligned bases | Aligned bases (%) | Unaligned bases | Unaligned bases (%) | Average length | SNPs | Identity |
|---|---|---|---|---|---|---|---|---|---|
| coronaSPAdes | 9 | 9 | 137420 | 100.0000 | 0 | 0.0000 | 17150.6250 | 36 | 99,884 |
| Haploflow | 42 | 42 | 143037 | 100.0000 | 0 | 0.0000 | 3533.0526 | 60 | 99,949 |
| IRMA | 4 | 4 | 128314 | 100.0000 | 0 | 0.0000 | 25663.2000 | 0 | 100,000 |
| LAZYPIPE | 10 | 10 | 137370 | 100.0000 | 0 | 0.0000 | 13677.4000 | 53 | 99,947 |
| metaSPAdes | 7 | 7 | 136233 | 99.9267 | 100 | 0.0733 | 15670.8889 | 37 | 99,824 |
| metaviralSPAdes | – | – | – | – | – | – | – | – | – |
| PEHaplo | 204 | 204 | 167437 | 99.9994 | 1 | 0.0006 | 810.4350 | 80 | 99,919 |
| QuRe | 6 | 5 | 1328 | 90.7724 | 135 | 9.2276 | 265.6000 | 6 | 99,471 |
| QVG | 4 | 4 | 143858 | 99.9986 | 2 | 0.0014 | 35964.5000 | 157 | 99,891 |
| SPAdes | 19 | 19 | 145008 | 99.9318 | 99 | 0.0682 | 8170.3529 | 32 | 99,931 |
| SSAKE | 313 | 313 | 91113 | 100.0000 | 0 | 0.0000 | 294.6271 | 7 | 99,982 |
| TRACESPipe | 4 | 4 | 143762 | 99.9319 | 98 | 0.0681 | 35940.5000 | 153 | 99,893 |
| TRACESPipeLite | 5 | 5 | 143991 | 94.9226 | 7702 | 5.0774 | 35948.0000 | 148 | 99,885 |
| V-pipe | 5 | 4 | 143412 | 89.3928 | 17017 | 10.6072 | 34271.2500 | 64 | 99,627 |
| VirGenA | 40 | 40 | 114988 | 99.9748 | 29 | 0.0252 | 2570.5349 | 67 | 99,866 |
| ViSpA | 5 | 4 | 143678 | 100.0000 | 0 | 0.0000 | 35919.5000 | 158 | 99,884 |

Table S73: Results obtained by dnadiff [4] when analysing the reconstruction of DS6. Only one execution of HVRS is represented.

| Reconstruction tool | Total seqs | Aligned seqs | Aligned bases | Aligned bases (%) | Unaligned bases | Unaligned bases (%) | Average length | SNPs | Identity |
|---|---|---|---|---|---|---|---|---|---|
| coronaSPAdes | 10 | 10 | 137343 | 99.9287 | 98 | 0.0713 | 17109.7500 | 58 | 99,912 |
| Haploflow | 21 | 21 | 165922 | 99.9711 | 48 | 0.0289 | 6905.3500 | 98 | 99,923 |
| IRMA | 4 | 4 | 128304 | 100.0000 | 0 | 0.0000 | 25660.8000 | 0 | 100,000 |
| LAZYPIPE | 8 | 8 | 136723 | 100.0000 | 0 | 0.0000 | 17025.1250 | 62 | 99,954 |
| metaSPAdes | 9 | 9 | 136739 | 99.8539 | 200 | 0.1461 | 12391.5455 | 68 | 99,904 |
| metaviralSPAdes | 1 | 1 | 5358 | 100.0000 | 0 | 0.0000 | 5358.0000 | 1 | 99,940 |
| PEHaplo | 203 | 203 | 177772 | 100.0000 | 0 | 0.0000 | 851.4518 | 41 | 99,962 |
| QuRe | 2 | 2 | 1429 | 100.0000 | 0 | 0.0000 | 714.5000 | 9 | 99,367 |
| QVG | 4 | 4 | 143858 | 99.9986 | 2 | 0.0014 | 35964.5000 | 149 | 99,893 |
| SPAdes | 25 | 25 | 146732 | 99.9980 | 3 | 0.0020 | 5662.7200 | 16 | 99,944 |
| SSAKE | 366 | 366 | 138013 | 99.9978 | 3 | 0.0022 | 387.2277 | 10 | 99,988 |
| TRACESPipe | 4 | 4 | 143780 | 99.9444 | 80 | 0.0556 | 35945.0000 | 148 | 99,894 |
| TRACESPipeLite | 5 | 4 | 143724 | 94.7466 | 7969 | 5.2534 | 35931.0000 | 163 | 99,885 |
| V-pipe | 5 | 4 | 143379 | 89.3722 | 17050 | 10.6278 | 35844.7500 | 148 | 99,886 |
| VirGenA | 40 | 40 | 116127 | 99.9466 | 62 | 0.0534 | 2425.4783 | 63 | 99,851 |
| ViSpA | 5 | 4 | 143613 | 100.0000 | 0 | 0.0000 | 35903.2500 | 156 | 99,891 |

Table S74: Results obtained by dnadiff [4] when analysing the reconstruction of DS7. Only one execution of HVRS is represented.

| Reconstruction tool | Total seqs | Aligned seqs | Aligned bases | Aligned bases (%) | Unaligned bases | Unaligned bases (%) | Average length | SNPs | Identity |
|---|---|---|---|---|---|---|---|---|---|
| coronaSPAdes | 11 | 11 | 137556 | 99.9985 | 2 | 0.0015 | 17125.1250 | 60 | 99,864 |
| Haploflow | 17 | 17 | 147874 | 99.9459 | 80 | 0.0541 | 8398.6471 | 42 | 99,862 |
| IRMA | 4 | 4 | 128337 | 100.0000 | 0 | 0.0000 | 25667.4000 | 0 | 100,000 |
| LAZYPIPE | 5 | 5 | 136271 | 100.0000 | 0 | 0.0000 | 20978.2857 | 95 | 99,733 |
| metaSPAdes | 10 | 10 | 136839 | 99.9270 | 100 | 0.0730 | 11394.9167 | 52 | 99,869 |
| metaviralSPAdes | – | – | – | – | – | – | – | – | – |
| PEHaplo | 208 | 208 | 192610 | 99.9964 | 7 | 0.0036 | 944.5989 | 55 | 99,957 |
| QuRe | 5 | 5 | 3694 | 99.9729 | 1 | 0.0271 | 586.7500 | 0 | 99,706 |
| QVG | 4 | 4 | 143858 | 99.9986 | 2 | 0.0014 | 35964.5000 | 149 | 99,893 |
| SPAdes | 25 | 25 | 146714 | 99.9980 | 3 | 0.0020 | 6129.3478 | 20 | 99,941 |
| SSAKE | 304 | 304 | 152481 | 100.0000 | 0 | 0.0000 | 534.7591 | 24 | 99,976 |
| TRACESPipe | 4 | 4 | 143834 | 99.9819 | 26 | 0.0181 | 35958.5000 | 162 | 99,885 |
| TRACESPipeLite | 5 | 5 | 144200 | 95.0604 | 7493 | 4.9396 | 35952.5000 | 144 | 99,896 |
| V-pipe | 5 | 4 | 143447 | 89.4146 | 16982 | 10.5854 | 35861.7500 | 147 | 99,886 |
| VirGenA | 15 | 15 | 117822 | 99.9847 | 18 | 0.0153 | 5139.9565 | 93 | 99,881 |
| ViSpA | 5 | 4 | 143678 | 100.0000 | 0 | 0.0000 | 30127.2000 | 165 | 99,840 |

Table S75: Results obtained by dnadiff [4] when analysing the reconstruction of DS8. Only one execution of HVRS is represented.

| Reconstruction tool | Total seqs | Aligned seqs | Aligned bases | Aligned bases (%) | Unaligned bases | Unaligned bases (%) | Average length | SNPs | Identity |
|---|---|---|---|---|---|---|---|---|---|
| coronaSPAdes | 10 | 10 | 137518 | 100.0000 | 0 | 0.0000 | 19572.5714 | 46 | 99,876 |
| Haploflow | 12 | 12 | 147403 | 99.9722 | 41 | 0.0278 | 11911.0000 | 119 | 99,870 |
| IRMA | 4 | 4 | 128349 | 100.0000 | 0 | 0.0000 | 25669.8000 | 0 | 100,000 |
| LAZYPIPE | 9 | 9 | 137996 | 100.0000 | 0 | 0.0000 | 16365.4444 | 52 | 99,879 |
| metaSPAdes | 10 | 10 | 136896 | 99.9263 | 101 | 0.0737 | 9973.8571 | 21 | 99,829 |
| metaviralSPAdes | – | – | – | – | – | – | – | – | – |
| PEHaplo | 186 | 186 | 208290 | 99.9990 | 2 | 0.0010 | 1138.3012 | 51 | 99,951 |
| QuRe | 3 | 3 | 9628 | 100.0000 | 0 | 0.0000 | 3209.3333 | 33 | 99,378 |
| QVG | 4 | 4 | 143858 | 99.9986 | 2 | 0.0014 | 35964.5000 | 147 | 99,894 |
| SPAdes | 25 | 25 | 146751 | 100.0000 | 0 | 0.0000 | 5894.0000 | 22 | 99,940 |
| SSAKE | 132 | 132 | 153157 | 100.0000 | 0 | 0.0000 | 1433.9216 | 5 | 99,994 |
| TRACESPipe | 4 | 4 | 143827 | 99.9771 | 33 | 0.0229 | 35956.7500 | 126 | 99,912 |
| TRACESPipeLite | 5 | 5 | 144026 | 94.9457 | 7667 | 5.0543 | 35956.0000 | 129 | 99,911 |
| V-pipe | 5 | 4 | 143517 | 89.4583 | 16912 | 10.5417 | 35879.2500 | 160 | 99,887 |
| VirGenA | 41 | 41 | 116653 | 99.9186 | 95 | 0.0814 | 2401.8723 | 96 | 99,869 |
| ViSpA | 5 | 4 | 143719 | 99.9993 | 1 | 0.0007 | 35929.7500 | 158 | 99,884 |

Table S76: Results obtained by dnadiff [4] when analysing the reconstruction of DS9. Only one execution of HVRS is represented.

| Reconstruction tool | Total seqs | Aligned seqs | Aligned bases | Aligned bases (%) | Unaligned bases | Unaligned bases (%) | Average length | SNPs | Identity |
|---|---|---|---|---|---|---|---|---|---|
| coronaSPAdes | 267 | 183 | 81775 | 67.8406 | 38765 | 32.1594 | 446.8579 | 117 | 99,856 |
| Haploflow | – | – | – | – | – | – | – | – | – |
| IRMA | 5 | 4 | 92865 | 88.3301 | 12269 | 11.6699 | 701.2406 | 106 | 96,424 |
| LAZYPIPE | 162 | 116 | 55407 | 69.2865 | 24561 | 30.7135 | 477.6466 | 61 | 99,890 |
| metaSPAdes | 305 | 209 | 86582 | 67.7057 | 41298 | 32.2943 | 414.2679 | 114 | 99,866 |
| metaviralSPAdes | – | – | – | – | – | – | – | – | – |
| PEHaplo | 125 | 87 | 53115 | 68.0665 | 24919 | 31.9335 | 582.2529 | 1616 | 99,819 |
| QuRe | – | – | – | – | – | – | – | – | – |
| QVG | 3 | 3 | 138262 | 99.9986 | 2 | 0.0014 | 46087.3333 | 1300 | 99,059 |
| SPAdes | 258 | 176 | 79767 | 67.5528 | 38314 | 32.4472 | 457.1494 | 94 | 99,880 |
| SSAKE | 4 | 0 | 0 | 0.0000 | 603 | 100.0000 | 0.0000 | 0 | 0,000 |
| TRACESPipe | 4 | 4 | 114586 | 79.6510 | 29274 | 20.3490 | 848.9380 | 127 | 96,979 |
| TRACESPipeLite | 6 | 4 | 109703 | 65.1873 | 58586 | 34.8127 | 893.3967 | 62 | 97,105 |
| V-pipe | 5 | 4 | 6397 | 3.9874 | 154032 | 96.0126 | 116.3091 | 0 | 98,428 |
| VirGenA | 200 | 196 | 66371 | 98.9254 | 721 | 1.0746 | 341.8842 | 79 | 99,849 |
| ViSpA | 85 | 77 | 2498349 | 98.2043 | 45683 | 1.7957 | 340.7778 | 96 | 94,399 |

Table S77: Results obtained by dnadiff [4] when analysing the reconstruction of DS10. Only one execution of HVRS is represented.

| Reconstruction tool | Total seqs | Aligned seqs | Aligned bases | Aligned bases (%) | Unaligned bases | Unaligned bases (%) | Average length | SNPs | Identity |
|---|---|---|---|---|---|---|---|---|---|
| coronaSPAdes | 174 | 121 | 131458 | 67.6391 | 62894 | 32.3609 | 1094.1000 | 132 | 99,856 |
| Haploflow | – | – | – | – | – | – | – | – | – |
| IRMA | 5 | 4 | 123283 | 88.6596 | 15769 | 11.3404 | 4252.1034 | 39 | 98,685 |
| LAZYPIPE | 145 | 102 | 118542 | 67.6687 | 56638 | 32.3313 | 1130.3905 | 109 | 99,867 |
| metaSPAdes | 273 | 182 | 121872 | 67.5060 | 58663 | 32.4940 | 665.9672 | 154 | 99,867 |
| metaviralSPAdes | – | – | – | – | – | – | – | – | – |
| PEHaplo | 155 | 107 | 123302 | 67.1550 | 60306 | 32.8450 | 1134.5741 | 88 | 99,917 |
| QuRe | – | – | – | – | – | – | – | – | – |
| QVG | 4 | 4 | 143858 | 99.9986 | 2 | 0.0014 | 35964.5000 | 989 | 99,317 |
| SPAdes | 167 | 114 | 129567 | 67.3212 | 62894 | 32.6788 | 1147.7130 | 118 | 99,716 |
| SSAKE | 76 | 56 | 8350 | 71.9952 | 3248 | 28.0048 | 149.1071 | 1 | 99,988 |
| TRACESPipe | 4 | 4 | 140451 | 97.6303 | 3409 | 2.3697 | 6384.1364 | 177 | 98,896 |
| TRACESPipeLite | 6 | 4 | 140336 | 83.4033 | 27926 | 16.5967 | 6973.5000 | 149 | 98,883 |
| V-pipe | 5 | 4 | 68824 | 42.9000 | 91605 | 57.1000 | 332.6683 | 22 | 94,328 |
| VirGenA | 124 | 123 | 73477 | 99.4168 | 431 | 0.5832 | 570.2913 | 65 | 99,830 |
| ViSpA | 92 | 82 | 4391811 | 98.1675 | 81984 | 1.8325 | 1988.2969 | 125 | 96,964 |

Table S78: Results obtained by dnadiff [4] when analysing the reconstruction of DS11. Only one execution of HVRS is represented.

| Reconstruction tool | Total seqs | Aligned seqs | Aligned bases | Aligned bases (%) | Unaligned bases | Unaligned bases (%) | Average length | SNPs | Identity |
|---|---|---|---|---|---|---|---|---|---|
| coronaSPAdes | 24 | 18 | 137432 | 67.3709 | 66561 | 32.6291 | 10514.2308 | 83 | 99,825 |
| Haploflow | 107 | 72 | 60173 | 69.9458 | 25855 | 30.0542 | 823.7361 | 38 | 99,933 |
| IRMA | 5 | 4 | 128055 | 88.5624 | 16538 | 11.4376 | 21342.8333 | 4 | 99,974 |
| LAZYPIPE | 20 | 14 | 135935 | 67.2014 | 66345 | 32.7986 | 9703.9286 | 89 | 99,911 |
| metaSPAdes | 25 | 19 | 136999 | 67.2715 | 66652 | 32.7285 | 6834.5000 | 47 | 99,853 |
| metaviralSPAdes | – | – | – | – | – | – | – | – | – |
| PEHaplo | 202 | 139 | 150703 | 67.6733 | 71989 | 32.3267 | 1061.7338 | 58 | 99,952 |
| QuRe | – | – | – | – | – | – | – | – | – |
| QVG | 4 | 4 | 143858 | 99.9986 | 2 | 0.0014 | 35964.5000 | 408 | 99,715 |
| SPAdes | 26 | 20 | 137920 | 67.3839 | 66758 | 32.6161 | 7022.8000 | 27 | 99,862 |
| SSAKE | 114 | 80 | 15281 | 72.0428 | 5930 | 27.9572 | 191.0125 | 4 | 99,974 |
| TRACESPipe | 4 | 4 | 143756 | 99.9277 | 104 | 0.0723 | 35939.0000 | 135 | 99,861 |
| TRACESPipeLite | 6 | 5 | 143842 | 85.4869 | 24420 | 14.5131 | 35911.2500 | 148 | 99,861 |
| V-pipe | 5 | 4 | 126788 | 79.0306 | 33641 | 20.9694 | 1333.6458 | 54 | 96,844 |
| VirGenA | 93 | 92 | 60436 | 99.3001 | 426 | 0.6999 | 583.4216 | 42 | 99,886 |
| ViSpA | 8 | 7 | 313106 | 95.0021 | 16472 | 4.9979 | 29090.5000 | 3 | 99,732 |

Table S79: Results obtained by dnadiff [4] when analysing the reconstruction of DS12. Only one execution of HVRS is represented.

| Reconstruction tool | Total seqs | Aligned seqs | Aligned bases | Aligned bases (%) | Unaligned bases | Unaligned bases (%) | Average length | SNPs | Identity |
|---|---|---|---|---|---|---|---|---|---|
| coronaSPAdes | 12 | 10 | 137312 | 67.3521 | 66560 | 32.6479 | 13723.1000 | 53 | 99,867 |
| Haploflow | 118 | 80 | 123050 | 68.3232 | 57050 | 31.6768 | 1523.6579 | 75 | 99,932 |
| IRMA | 5 | 4 | 128255 | 88.5916 | 16516 | 11.4084 | 25651.0000 | 0 | 99,984 |
| LAZYPIPE | 10 | 8 | 136253 | 67.2695 | 66295 | 32.7305 | 15137.3333 | 74 | 99,938 |
| metaSPAdes | 13 | 11 | 136713 | 67.2555 | 66561 | 32.7445 | 10526.3846 | 55 | 99,823 |
| metaviralSPAdes | – | – | – | – | – | – | – | – | – |
| PEHaplo | 276 | 189 | 159825 | 67.6522 | 76420 | 32.3478 | 833.5574 | 38 | 99,969 |
| QuRe | 4 | 2 | 224 | 56.2814 | 174 | 43.7186 | 112.0000 | 0 | 100,000 |
| QVG | 4 | 4 | 143858 | 99.9986 | 2 | 0.0014 | 35964.5000 | 225 | 99,843 |
| SPAdes | 19 | 17 | 138979 | 67.5511 | 66760 | 32.4489 | 7294.8947 | 47 | 99,918 |
| SSAKE | 214 | 148 | 34855 | 69.3190 | 15427 | 30.6810 | 235.5068 | 8 | 99,968 |
| TRACESPipe | 4 | 4 | 143717 | 99.9006 | 143 | 0.0994 | 35929.2500 | 147 | 99,896 |
| TRACESPipeLite | 6 | 5 | 144132 | 85.6593 | 24130 | 14.3407 | 35933.2500 | 128 | 99,903 |
| V-pipe | 5 | 4 | 142389 | 88.7552 | 18040 | 11.2448 | 15821.0000 | 145 | 99,134 |
| VirGenA | 72 | 71 | 59108 | 99.2894 | 423 | 0.7106 | 741.7722 | 46 | 99,864 |
| ViSpA | 6 | 5 | 180530 | 91.6373 | 16475 | 8.3627 | 28693.4000 | 150 | 99,889 |

Table S80: Results obtained by dnadiff [4] when analysing the reconstruction of DS13. Only one execution of HVRS is represented.

| Reconstruction tool | Total seqs | Aligned seqs | Aligned bases | Aligned bases (%) | Unaligned bases | Unaligned bases (%) | Average length | SNPs | Identity |
|---|---|---|---|---|---|---|---|---|---|
| coronaSPAdes | 14 | 12 | 137731 | 67.4332 | 66517 | 32.5668 | 12662.9091 | 44 | 99,821 |
| Haploflow | 42 | 32 | 142611 | 68.5547 | 65414 | 31.4453 | 4715.4828 | 92 | 99,913 |
| IRMA | 5 | 4 | 128300 | 88.5995 | 16509 | 11.4005 | 25660.2000 | 0 | 100,000 |
| LAZYPIPE | 7 | 5 | 136414 | 67.2616 | 66397 | 32.7384 | 17129.5000 | 56 | 99,932 |
| metaSPAdes | 13 | 11 | 136759 | 67.2772 | 66518 | 32.7228 | 11396.5833 | 59 | 99,861 |
| metaviralSPAdes | – | – | – | – | – | – | – | – | – |
| PEHaplo | 305 | 211 | 173528 | 69.4977 | 76161 | 30.5023 | 834.8408 | 87 | 99,937 |
| QuRe | 4 | 3 | 695 | 87.8635 | 96 | 12.1365 | 231.6667 | 0 | 100,000 |
| QVG | 4 | 4 | 143858 | 99.9986 | 2 | 0.0014 | 35964.5000 | 157 | 99,892 |
| SPAdes | 21 | 19 | 145210 | 68.5506 | 66619 | 31.4494 | 6673.6190 | 36 | 99,906 |
| SSAKE | 373 | 266 | 84777 | 69.9457 | 36427 | 30.0543 | 320.4580 | 28 | 99,964 |
| TRACESPipe | 4 | 4 | 143825 | 99.9757 | 35 | 0.0243 | 35956.2500 | 143 | 99,902 |
| TRACESPipeLite | 6 | 4 | 143716 | 85.4120 | 24546 | 14.5880 | 28808.8000 | 123 | 99,900 |
| V-pipe | 5 | 4 | 143294 | 89.3193 | 17135 | 10.6807 | 35823.5000 | 128 | 99,765 |
| VirGenA | 66 | 65 | 57306 | 98.8086 | 691 | 1.1914 | 717.4805 | 45 | 99,882 |
| ViSpA | 5 | 4 | 143592 | 89.6951 | 16497 | 10.3049 | 35898.0000 | 153 | 99,893 |

Table S81: Results obtained by dnadiff [4] when analysing the reconstruction of DS14. Only one execution of HVRS is represented.

| Reconstruction tool | Total seqs | Aligned seqs | Aligned bases | Aligned bases (%) | Unaligned bases | Unaligned bases (%) | Average length | SNPs | Identity |
|---|---|---|---|---|---|---|---|---|---|
| coronaSPAdes | 10 | 8 | 137186 | 67.3127 | 66618 | 32.6873 | 19568.8571 | 63 | 99,910 |
| Haploflow | 30 | 23 | 147275 | 68.9344 | 66370 | 31.0656 | 6243.7273 | 80 | 99,920 |
| IRMA | 5 | 4 | 128330 | 88.5804 | 16544 | 11.4196 | 25666.2000 | 0 | 100,000 |
| LAZYPIPE | 10 | 8 | 136874 | 67.3212 | 66441 | 32.6788 | 15232.7778 | 61 | 99,915 |
| metaSPAdes | 12 | 10 | 136892 | 67.2655 | 66618 | 32.7345 | 10740.7692 | 31 | 99,823 |
| metaviralSPAdes | – | – | – | – | – | – | – | – | – |
| PEHaplo | 289 | 184 | 173544 | 67.0318 | 85354 | 32.9682 | 942.4171 | 61 | 99,954 |
| QuRe | 4 | 3 | 1460 | 82.7195 | 305 | 17.2805 | 486.6667 | 4 | 99,722 |
| QVG | 4 | 4 | 143858 | 99.9986 | 2 | 0.0014 | 35964.5000 | 150 | 99,893 |
| SPAdes | 25 | 23 | 146599 | 68.7877 | 66519 | 31.2123 | 6707.6190 | 24 | 99,938 |
| SSAKE | 489 | 345 | 134050 | 69.4322 | 59016 | 30.5678 | 396.7190 | 25 | 99,969 |
| TRACESPipe | 4 | 4 | 143798 | 99.9562 | 63 | 0.0438 | 35949.5000 | 136 | 99,903 |
| TRACESPipeLite | 6 | 5 | 144010 | 85.5868 | 24252 | 14.4132 | 35949.0000 | 161 | 99,885 |
| V-pipe | 5 | 4 | 143451 | 89.4171 | 16978 | 10.5829 | 35862.7500 | 134 | 99,890 |
| VirGenA | 68 | 67 | 57858 | 99.7036 | 172 | 0.2964 | 716.9630 | 33 | 99,899 |
| ViSpA | 5 | 4 | 143728 | 89.6904 | 16521 | 10.3096 | 35932.0000 | 142 | 99,901 |

Table S82: Results obtained by dnadiff [4] when analysing the reconstruction of DS15. Only one execution of HVRS is represented.

| Reconstruction tool | Total seqs | Aligned seqs | Aligned bases | Aligned bases (%) | Unaligned bases | Unaligned bases (%) | Average length | SNPs | Identity |
|---|---|---|---|---|---|---|---|---|---|
| coronaSPAdes | 12 | 10 | 137407 | 67.3409 | 66640 | 32.6591 | 17112.1250 | 52 | 99,916 |
| Haploflow | 20 | 16 | 147883 | 68.9788 | 66506 | 31.0212 | 9187.7333 | 103 | 99,908 |
| IRMA | 5 | 4 | 128341 | 88.5752 | 16554 | 11.4248 | 25668.4000 | 0 | 100,000 |
| LAZYPIPE | 8 | 6 | 136283 | 67.2047 | 66505 | 32.7953 | 17121.3750 | 66 | 99,929 |
| metaSPAdes | 13 | 11 | 136969 | 67.2703 | 66641 | 32.7297 | 10737.5385 | 40 | 99,836 |
| metaviralSPAdes | – | – | – | – | – | – | – | – | – |
| PEHaplo | 313 | 207 | 197805 | 67.5367 | 95080 | 32.4633 | 972.9206 | 66 | 99,952 |
| QuRe | 5 | 4 | 1017 | 82.4818 | 216 | 17.5182 | 254.2500 | 2 | 99,804 |
| QVG | 4 | 4 | 143858 | 99.9986 | 2 | 0.0014 | 35964.5000 | 148 | 99,893 |
| SPAdes | 27 | 25 | 141613 | 68.0325 | 66542 | 31.9675 | 5893.5833 | 13 | 99,947 |
| SSAKE | 404 | 288 | 149793 | 68.6158 | 68514 | 31.3842 | 557.5290 | 39 | 99,915 |
| TRACESPipe | 4 | 4 | 143764 | 99.9333 | 96 | 0.0667 | 35941.0000 | 140 | 99,904 |
| TRACESPipeLite | 6 | 5 | 144130 | 85.6581 | 24132 | 14.3419 | 35941.7500 | 111 | 99,920 |
| V-pipe | 5 | 4 | 143381 | 89.3735 | 17048 | 10.6265 | 35845.2500 | 165 | 99,887 |
| VirGenA | 69 | 68 | 55640 | 98.9948 | 565 | 1.0052 | 713.9333 | 43 | 99,888 |
| ViSpA | 5 | 4 | 143681 | 89.6824 | 16530 | 10.3176 | 35920.2500 | 166 | 99,884 |

Table S83: Results obtained by dnadiff [4] when analysing the reconstruction of DS16. Only one execution of HVRS is represented.

| Reconstruction tool | Total seqs | Aligned seqs | Aligned bases | Aligned bases (%) | Unaligned bases | Unaligned bases (%) | Average length | SNPs | Identity |
|---|---|---|---|---|---|---|---|---|---|
| coronaSPAdes | 10 | 8 | 137191 | 67.3023 | 66652 | 32.6977 | 19569.5714 | 47 | 99,923 |
| Haploflow | 12 | 10 | 145468 | 68.6947 | 66292 | 31.3053 | 13811.4000 | 113 | 99,876 |
| IRMA | 5 | 4 | 128345 | 88.5712 | 16561 | 11.4288 | 25669.0000 | 0 | 100,000 |
| LAZYPIPE | 9 | 7 | 136903 | 67.2967 | 66529 | 32.7033 | 19557.5714 | 53 | 99,942 |
| metaSPAdes | 12 | 10 | 136897 | 67.2551 | 66652 | 32.7449 | 10741.1538 | 24 | 99,828 |
| metaviralSPAdes | 2 | 0 | 0 | 0.0000 | 66553 | 100.0000 | 0.0000 | 0 | 0,000 |
| PEHaplo | 220 | 149 | 222518 | 69.1679 | 99189 | 30.8321 | 1489.3525 | 64 | 99,938 |
| QuRe | 8 | 7 | 25124 | 97.6941 | 593 | 2.3059 | 2470.2500 | 32 | 99,676 |
| QVG | 4 | 4 | 143858 | 99.9986 | 2 | 0.0014 | 35964.5000 | 147 | 99,894 |
| SPAdes | 27 | 25 | 146744 | 68.7980 | 66553 | 31.2020 | 5893.7083 | 22 | 99,940 |
| SSAKE | 223 | 164 | 155802 | 68.7576 | 70794 | 31.2424 | 1224.4667 | 10 | 99,993 |
| TRACESPipe | 4 | 4 | 143813 | 99.9673 | 47 | 0.0327 | 35953.2500 | 151 | 99,893 |
| TRACESPipeLite | 6 | 4 | 143814 | 85.4703 | 24448 | 14.5297 | 35953.5000 | 138 | 99,903 |
| V-pipe | 5 | 4 | 143515 | 89.4570 | 16914 | 10.5430 | 35878.7500 | 132 | 99,904 |
| VirGenA | 55 | 55 | 58585 | 99.2411 | 448 | 0.7589 | 804.4429 | 40 | 99,844 |
| ViSpA | 5 | 4 | 143700 | 89.6679 | 16558 | 10.3321 | 35925.0000 | 164 | 99,884 |

Table S84: Results obtained by dnadiff [4] when analysing the reconstruction of DS17. Only one execution of HVRS is represented.

| Reconstruction tool | Total seqs | Aligned seqs | Aligned bases | Aligned bases (%) | Unaligned bases | Unaligned bases (%) | Average length | SNPs | Identity |
|---|---|---|---|---|---|---|---|---|---|
| coronaSPAdes | 259 | 174 | 84390 | 69.0081 | 37900 | 30.9919 | 485.0000 | 95 | 99,886 |
| Haploflow | – | – | – | – | – | – | – | – | – |
| IRMA | 5 | 4 | 96234 | 88.9293 | 11980 | 11.0707 | 730.7576 | 112 | 95,775 |
| LAZYPIPE | 159 | 109 | 58380 | 70.4596 | 24476 | 29.5404 | 526.9550 | 29 | 99,949 |
| metaSPAdes | 306 | 212 | 92930 | 70.0365 | 39758 | 29.9635 | 436.3726 | 83 | 99,909 |
| metaviralSPAdes | – | – | – | – | – | – | – | – | – |
| PEHaplo | 131 | 94 | 60152 | 71.9005 | 23508 | 28.0995 | 627.6632 | 522 | 99,878 |
| QuRe | – | – | – | – | – | – | – | – | – |
| QVG | – | – | – | – | – | – | – | – | – |
| SPAdes | – | – | – | – | – | – | – | – | – |
| SSAKE | 2 | 2 | 251 | 100.0000 | 0 | 0.0000 | 125.5000 | 0 | 100,000 |
| TRACESPipe | 4 | 4 | 114023 | 79.2597 | 29837 | 20.7403 | 857.2960 | 15 | 97,482 |
| TRACESPipeLite | 6 | 4 | 113527 | 67.4704 | 54735 | 32.5296 | 930.9835 | 15 | 96,411 |
| V-pipe | 5 | 3 | 5686 | 3.5442 | 154743 | 96.4558 | 132.2326 | 0 | 95,912 |
| VirGenA | 214 | 210 | 67931 | 99.5472 | 309 | 0.4528 | 321.8883 | 55 | 99,867 |
| ViSpA | 86 | 75 | 2357091 | 97.7255 | 54861 | 2.2745 | 338.8186 | 54 | 96,054 |

Table S85: Results obtained by dnadiff [4] when analysing the reconstruction of DS18. Only one execution of HVRS is represented.

| Reconstruction tool | Total seqs | Aligned seqs | Aligned bases | Aligned bases (%) | Unaligned bases | Unaligned bases (%) | Average length | SNPs | Identity |
|---|---|---|---|---|---|---|---|---|---|
| coronaSPAdes | 272 | 182 | 80543 | 66.9585 | 39745 | 33.0415 | 444.3128 | 215 | 99,730 |
| Haploflow | – | – | – | – | – | – | – | – | – |
| IRMA | 5 | 4 | 91913 | 88.0334 | 12494 | 11.9666 | 720.5312 | 109 | 95,614 |
| LAZYPIPE | 157 | 113 | 52867 | 69.7822 | 22893 | 30.2178 | 467.8496 | 119 | 99,741 |
| metaSPAdes | 327 | 219 | 87274 | 66.7692 | 43436 | 33.2308 | 399.8945 | 152 | 99,818 |
| metaviralSPAdes | – | – | – | – | – | – | – | – | – |
| PEHaplo | 118 | 92 | 53767 | 73.0381 | 19848 | 26.9619 | 591.6413 | 2702 | 99,631 |
| QuRe | – | – | – | – | – | – | – | – | – |
| QVG | 3 | 3 | 138473 | 100.0000 | 0 | 0.0000 | 34647.7500 | 4037 | 96,924 |
| SPAdes | – | – | – | – | – | – | – | – | – |
| SSAKE | 1 | 1 | 167 | 100.0000 | 0 | 0.0000 | 167.0000 | 0 | 100,000 |
| TRACESPipe | 4 | 4 | 112911 | 78.4873 | 30948 | 21.5127 | 899.5920 | 264 | 96,898 |
| TRACESPipeLite | 6 | 4 | 95121 | 56.5940 | 72955 | 43.4060 | 617.0987 | 120 | 97,176 |
| V-pipe | 5 | 3 | 6746 | 4.2050 | 153683 | 95.7950 | 140.5417 | 6 | 95,968 |
| VirGenA | 211 | 210 | 67948 | 99.6393 | 246 | 0.3607 | 323.3269 | 112 | 99,808 |
| ViSpA | 81 | 72 | 2367542 | 97.8694 | 51541 | 2.1306 | 328.8703 | 174 | 94,834 |

Table S86: Results obtained by dnadiff [4] when analysing the reconstruction of DS19. Only one execution of HVRS is represented.

| Reconstruction tool | Total seqs | Aligned seqs | Aligned bases | Aligned bases (%) | Unaligned bases | Unaligned bases (%) | Average length | SNPs | Identity |
|---|---|---|---|---|---|---|---|---|---|
| coronaSPAdes | 274 | 184 | 83666 | 66.9998 | 41209 | 33.0002 | 460.4348 | 164 | 99,656 |
| Haploflow | – | – | – | – | – | – | – | – | – |
| IRMA | 5 | 4 | 94356 | 88.4502 | 12321 | 11.5498 | 674.8500 | 126 | 96,223 |
| LAZYPIPE | 182 | 128 | 59305 | 68.7666 | 26936 | 31.2334 | 464.5349 | 60 | 99,773 |
| metaSPAdes | 323 | 219 | 90359 | 67.1619 | 44180 | 32.8381 | 415.5000 | 126 | 99,747 |
| metaviralSPAdes | – | – | – | – | – | – | – | – | – |
| PEHaplo | 138 | 97 | 56919 | 69.0530 | 25509 | 30.9470 | 594.6146 | 1272 | 99,598 |
| QuRe | – | – | – | – | – | – | – | – | – |
| QVG | 4 | 4 | 143858 | 99.9986 | 2 | 0.0014 | 35964.5000 | 6858 | 95,231 |
| SPAdes | – | – | – | – | – | – | – | – | – |
| SSAKE | – | – | – | – | – | – | – | – | – |
| TRACESPipe | 4 | 4 | 117131 | 81.4207 | 26728 | 18.5793 | 974.2667 | 555 | 96,286 |
| TRACESPipeLite | 6 | 4 | 53684 | 31.9080 | 114562 | 68.0920 | 331.5000 | 73 | 98,109 |
| V-pipe | 5 | 4 | 6608 | 4.1190 | 153821 | 95.8810 | 132.1600 | 1 | 95,875 |
| VirGenA | 195 | 195 | 64904 | 99.9246 | 49 | 0.0754 | 321.3941 | 85 | 99,790 |
| ViSpA | 86 | 79 | 2326634 | 98.1353 | 44210 | 1.8647 | 331.6303 | 299 | 94,594 |

Table S87: Results obtained by dnadiff [4] when analysing the reconstruction of DS20. Only one execution of HVRS is represented.

| Reconstruction tool | Total seqs | Aligned seqs | Aligned bases | Aligned bases (%) | Unaligned bases | Unaligned bases (%) | Average length | SNPs | Identity |
|---|---|---|---|---|---|---|---|---|---|
| coronaSPAdes | 265 | 176 | 80299 | 68.5566 | 36829 | 31.4434 | 457.4971 | 245 | 99,694 |
| Haploflow | – | – | – | – | – | – | – | – | – |
| IRMA | 5 | 4 | 93433 | 88.7598 | 11832 | 11.2402 | 628.2013 | 120 | 96,533 |
| LAZYPIPE | 154 | 111 | 52118 | 71.5848 | 20688 | 28.4152 | 468.6667 | 48 | 99,831 |
| metaSPAdes | 327 | 221 | 90222 | 69.0816 | 40380 | 30.9184 | 409.6575 | 163 | 99,805 |
| metaviralSPAdes | – | – | – | – | – | – | – | – | – |
| PEHaplo | 125 | 93 | 54594 | 72.7532 | 20446 | 27.2468 | 583.7684 | 2658 | 99,720 |
| QuRe | – | – | – | – | – | – | – | – | – |
| QVG | 4 | 4 | 143860 | 100.0000 | 0 | 0.0000 | 35965.0000 | 9676 | 93,217 |
| SPAdes | – | – | – | – | – | – | – | – | – |
| SSAKE | 5 | 4 | 609 | 78.5806 | 166 | 21.4194 | 152.2500 | 0 | 100,000 |
| TRACESPipe | 4 | 4 | 110655 | 77.0267 | 33003 | 22.9733 | 791.9023 | 580 | 96,547 |
| TRACESPipeLite | 6 | 2 | 9730 | 5.7896 | 158330 | 94.2104 | 237.3171 | 3 | 98,918 |
| V-pipe | 5 | 4 | 7649 | 4.7678 | 152780 | 95.2322 | 141.6481 | 7 | 96,532 |
| VirGenA | 206 | 201 | 64285 | 99.3755 | 404 | 0.6245 | 319.2150 | 70 | 99,812 |
| ViSpA | 77 | 69 | 2115952 | 98.0809 | 41402 | 1.9191 | 305.1036 | 224 | 95,150 |

Table S88: Results obtained by dnadiff [4] when analysing the reconstruction of DS21. Only one execution of HVRS is represented.

| Reconstruction tool | Total seqs | Aligned seqs | Aligned bases | Aligned bases (%) | Unaligned bases | Unaligned bases (%) | Average length | SNPs | Identity |
|---|---|---|---|---|---|---|---|---|---|
| coronaSPAdes | 279 | 190 | 85483 | 68.7588 | 38840 | 31.2412 | 450.5550 | 143 | 99,663 |
| Haploflow | – | – | – | – | – | – | – | – | – |
| IRMA | 5 | 4 | 92996 | 88.2868 | 12338 | 11.7132 | 656.0352 | 80 | 96,255 |
| LAZYPIPE | 169 | 111 | 54996 | 68.1158 | 25743 | 31.8842 | 495.4595 | 75 | 99,864 |
| metaSPAdes | 329 | 221 | 92276 | 68.3060 | 42816 | 31.6940 | 418.2387 | 104 | 99,743 |
| metaviralSPAdes | – | – | – | – | – | – | – | – | – |
| PEHaplo | 138 | 94 | 57468 | 68.6620 | 26229 | 31.3380 | 613.8602 | 2053 | 99,934 |
| QuRe | – | – | – | – | – | – | – | – | – |
| QVG | 4 | 4 | 143816 | 99.9694 | 44 | 0.0306 | 35953.2500 | 12401 | 91,349 |
| SPAdes | – | – | – | – | – | – | – | – | – |
| SSAKE | – | – | – | – | – | – | – | – | – |
| TRACESPipe | 4 | 4 | 98066 | 68.1667 | 45796 | 31.8333 | 694.0821 | 1200 | 95,608 |
| TRACESPipeLite | 6 | 2 | 2614 | 1.5536 | 165637 | 98.4464 | 201.0769 | 0 | 99,886 |
| V-pipe | 5 | 4 | 6352 | 3.9594 | 154077 | 96.0406 | 129.6327 | 5 | 94,181 |
| VirGenA | 198 | 193 | 64608 | 99.0814 | 599 | 0.9186 | 330.9643 | 95 | 99,732 |
| ViSpA | 69 | 58 | 1514235 | 95.9056 | 64645 | 4.0944 | 278.6948 | 248 | 94,797 |

Table S89: Results obtained by dnadiff [4] when analysing the reconstruction of DS22. Only one execution of HVRS is represented.

| Reconstruction tool | Total seqs | Aligned seqs | Aligned bases | Aligned bases (%) | Unaligned bases | Unaligned bases (%) | Average length | SNPs | Identity |
|---|---|---|---|---|---|---|---|---|---|
| coronaSPAdes | 263 | 184 | 82843 | 68.3512 | 38359 | 31.6488 | 450.2337 | 126 | 99,848 |
| Haploflow | – | – | – | – | – | – | – | – | – |
| IRMA | 5 | 4 | 91779 | 88.5675 | 11847 | 11.4325 | 651.7376 | 114 | 96,015 |
| LAZYPIPE | 163 | 111 | 54566 | 67.1888 | 26647 | 32.8112 | 491.5856 | 36 | 99,934 |
| metaSPAdes | 314 | 224 | 91343 | 69.1903 | 40674 | 30.8097 | 407.7812 | 85 | 99,902 |
| metaviralSPAdes | – | – | – | – | – | – | – | – | – |
| PEHaplo | 133 | 93 | 55541 | 68.0008 | 26136 | 31.9992 | 578.8958 | 1950 | 99,939 |
| QuRe | – | – | – | – | – | – | – | – | – |
| QVG | 4 | 4 | 143860 | 100.0000 | 0 | 0.0000 | 35965.0000 | 15309 | 89,322 |
| SPAdes | – | – | – | – | – | – | – | – | – |
| SSAKE | 3 | 2 | 369 | 70.8253 | 152 | 29.1747 | 184.5000 | 0 | 100,000 |
| TRACESPipe | 4 | 4 | 74753 | 52.0354 | 68905 | 47.9646 | 528.3147 | 1294 | 96,259 |
| TRACESPipeLite | 6 | 1 | 221 | 0.1313 | 168041 | 99.8687 | 221.0000 | 0 | 100,000 |
| V-pipe | 5 | 4 | 5301 | 3.3043 | 155128 | 96.6957 | 117.8000 | 5 | 97,153 |
| VirGenA | 208 | 203 | 65807 | 99.5176 | 319 | 0.4824 | 318.0628 | 99 | 99,833 |
| ViSpA | 49 | 39 | 906889 | 94.2327 | 55504 | 5.7673 | 249.4409 | 175 | 95,431 |

Table S90: Results obtained by dnadiff [4] when analysing the reconstruction of DS23. Only one execution of HVRS is represented.

| Reconstruction tool | Total seqs | Aligned seqs | Aligned bases | Aligned bases (%) | Unaligned bases | Unaligned bases (%) | Average length | SNPs | Identity |
|---|---|---|---|---|---|---|---|---|---|
| coronaSPAdes | 265 | 188 | 85953 | 68.8164 | 38949 | 31.1836 | 455.6402 | 149 | 99,794 |
| Haploflow | 1 | 1 | 611 | 100.0000 | 0 | 0.0000 | 611.0000 | 1 | 99,840 |
| IRMA | 5 | 4 | 89102 | 88.4492 | 11636 | 11.5508 | 507.0511 | 97 | 96,971 |
| LAZYPIPE | 178 | 121 | 58166 | 66.6445 | 29112 | 33.3555 | 480.7107 | 67 | 99,885 |
| metaSPAdes | 307 | 219 | 92416 | 69.1649 | 41201 | 30.8351 | 421.9909 | 96 | 99,895 |
| metaviralSPAdes | – | – | – | – | – | – | – | – | – |
| PEHaplo | 134 | 89 | 53334 | 64.8903 | 28857 | 35.1097 | 599.2584 | 1047 | 99,912 |
| QuRe | – | – | – | – | – | – | – | – | – |
| QVG | 4 | 4 | 143818 | 99.9708 | 42 | 0.0292 | 35954.5000 | 18093 | 87,202 |
| SPAdes | – | – | – | – | – | – | – | – | – |
| SSAKE | 3 | 1 | 124 | 32.7177 | 255 | 67.2823 | 124.0000 | 0 | 100,000 |
| TRACESPipe | 4 | 4 | 66560 | 46.2820 | 77254 | 53.7180 | 469.6993 | 1082 | 96,453 |
| TRACESPipeLite | 5 | 0 | 0 | 0.0000 | 160429 | 100.0000 | 0.0000 | 0 | 0,000 |
| V-pipe | 5 | 3 | 2539 | 1.5826 | 157890 | 98.4174 | 133.6316 | 0 | 92,188 |
| VirGenA | 210 | 205 | 66928 | 99.3793 | 418 | 0.6207 | 320.5311 | 71 | 99,849 |
| ViSpA | 24 | 15 | 173066 | 79.9950 | 43280 | 20.0050 | 205.2892 | 100 | 95,819 |

Table S91: Results obtained by dnadiff [4] when analysing the reconstruction of DS24. Only one execution of HVRS is represented.

| Reconstruction tool | Total seqs | Aligned seqs | Aligned bases | Aligned bases (%) | Unaligned bases | Unaligned bases (%) | Average length | SNPs | Identity |
|---|---|---|---|---|---|---|---|---|---|
| coronaSPAdes | 280 | 190 | 86772 | 69.6041 | 37893 | 30.3959 | 456.6947 | 108 | 99,874 |
| Haploflow | 1 | 1 | 630 | 100.0000 | 0 | 0.0000 | 630.0000 | 1 | 99,840 |
| IRMA | 5 | 4 | 86789 | 87.7667 | 12097 | 12.2333 | 489.2697 | 130 | 97,010 |
| LAZYPIPE | 228 | 154 | 67093 | 69.8609 | 28945 | 30.1391 | 435.6688 | 32 | 99,952 |
| metaSPAdes | 321 | 214 | 91855 | 68.8594 | 41540 | 31.1406 | 429.2290 | 108 | 99,880 |
| metaviralSPAdes | – | – | – | – | – | – | – | – | – |
| PEHaplo | 139 | 98 | 57283 | 71.1661 | 23209 | 28.8339 | 567.5248 | 2425 | 99,915 |
| QuRe | – | – | – | – | – | – | – | – | – |
| QVG | 4 | 4 | 143851 | 99.9937 | 9 | 0.0063 | 35962.7500 | 20734 | 85,427 |
| SPAdes | – | – | – | – | – | – | – | – | – |
| SSAKE | 7 | 3 | 507 | 42.3559 | 690 | 57.6441 | 169.0000 | 0 | 100,000 |
| TRACESPipe | 4 | 4 | 56696 | 39.4127 | 87156 | 60.5873 | 443.2171 | 1107 | 96,089 |
| TRACESPipeLite | 5 | 0 | 0 | 0.0000 | 160227 | 100.0000 | 0.0000 | 0 | 0,000 |
| V-pipe | 5 | 4 | 2595 | 1.6175 | 157834 | 98.3825 | 108.1250 | 0 | 96,533 |
| VirGenA | 222 | 214 | 68076 | 99.2376 | 523 | 0.7624 | 311.0548 | 79 | 99,853 |
| ViSpA | 15 | 4 | 47037 | 53.1012 | 41543 | 46.8988 | 157.9459 | 77 | 97,678 |

Table S92: Results obtained by dnadiff [4] when analysing the reconstruction of DS25. Only one execution of HVRS is represented.

| Reconstruction tool | Total seqs | Aligned seqs | Aligned bases | Aligned bases (%) | Unaligned bases | Unaligned bases (%) | Average length | SNPs | Identity |
|---|---|---|---|---|---|---|---|---|---|
| coronaSPAdes | 162 | 105 | 130031 | 66.9766 | 64113 | 33.0234 | 1227.5377 | 84 | 99,832 |
| Haploflow | 5 | 4 | 4374 | 88.7581 | 554 | 11.2419 | 1093.5000 | 0 | 100,000 |
| IRMA | 5 | 4 | 124223 | 88.6807 | 15856 | 11.3193 | 5916.0952 | 54 | 98,431 |
| LAZYPIPE | 150 | 97 | 119840 | 67.0722 | 58833 | 32.9278 | 1235.4639 | 20 | 99,984 |
| metaSPAdes | 265 | 171 | 121006 | 66.7513 | 60273 | 33.2487 | 703.5233 | 96 | 99,888 |
| metaviralSPAdes | – | – | – | – | – | – | – | – | – |
| PEHaplo | 159 | 106 | 129864 | 67.4051 | 62798 | 32.5949 | 1201.2991 | 41 | 99,951 |
| QuRe | – | – | – | – | – | – | – | – | – |
| QVG | – | – | – | – | – | – | – | – | – |
| SPAdes | 163 | 106 | 130203 | 67.0055 | 64114 | 32.9945 | 1217.6729 | 55 | 99,851 |
| SSAKE | 93 | 71 | 10866 | 75.8322 | 3463 | 24.1678 | 152.8028 | 1 | 99,982 |
| TRACESPipe | 4 | 4 | 140919 | 97.9557 | 2941 | 2.0443 | 7045.9500 | 7 | 98,920 |
| TRACESPipeLite | 6 | 5 | 142456 | 84.6632 | 25806 | 15.3368 | 12932.5455 | 2 | 98,858 |
| V-pipe | 5 | 4 | 68320 | 42.5858 | 92109 | 57.4142 | 321.9183 | 5 | 95,046 |
| VirGenA | 104 | 104 | 81489 | 99.9375 | 51 | 0.0625 | 741.0472 | 44 | 99,916 |
| ViSpA | 89 | 76 | 3628951 | 97.7545 | 83359 | 2.2455 | 1843.3582 | 34 | 96,987 |

Table S93: Results obtained by dnadiff [4] when analysing the reconstruction of DS26. Only one execution of HVRS is represented.

| Reconstruction tool | Total seqs | Aligned seqs | Aligned bases | Aligned bases (%) | Unaligned bases | Unaligned bases (%) | Average length | SNPs | Identity |
|---|---|---|---|---|---|---|---|---|---|
| coronaSPAdes | 162 | 114 | 133959 | 67.7913 | 63646 | 32.2087 | 1219.8532 | 186 | 99,737 |
| Haploflow | 1 | 0 | 0 | 0.0000 | 520 | 100.0000 | 0.0000 | 0 | 0,000 |
| IRMA | 5 | 4 | 123337 | 88.5437 | 15958 | 11.4563 | 5377.6087 | 48 | 98,297 |
| LAZYPIPE | 154 | 107 | 124737 | 67.7171 | 59466 | 32.2829 | 1202.6505 | 56 | 99,847 |
| metaSPAdes | 308 | 217 | 110049 | 67.5524 | 52860 | 32.4476 | 515.9275 | 110 | 99,891 |
| metaviralSPAdes | – | – | – | – | – | – | – | – | – |
| PEHaplo | 162 | 112 | 140459 | 68.3416 | 65066 | 31.6584 | 1214.0741 | 336 | 99,730 |
| QuRe | – | – | – | – | – | – | – | – | – |
| QVG | 4 | 4 | 143860 | 100.0000 | 0 | 0.0000 | 35965.0000 | 3136 | 97,748 |
| SPAdes | 158 | 110 | 131221 | 67.3384 | 63647 | 32.6616 | 1215.7477 | 247 | 99,807 |
| SSAKE | 78 | 54 | 8741 | 70.7544 | 3613 | 29.2456 | 161.8704 | 0 | 100,000 |
| TRACESPipe | 4 | 4 | 140829 | 97.8924 | 3032 | 2.1076 | 7041.1500 | 502 | 98,527 |
| TRACESPipeLite | 6 | 4 | 135071 | 80.2742 | 33191 | 19.7258 | 2811.5532 | 215 | 98,103 |
| V-pipe | 5 | 4 | 65656 | 40.9253 | 94773 | 59.0747 | 319.8768 | 58 | 94,635 |
| VirGenA | 126 | 124 | 66483 | 99.2891 | 476 | 0.7109 | 503.2308 | 62 | 99,783 |
| ViSpA | 86 | 78 | 3828546 | 98.3742 | 63272 | 1.6258 | 1925.1970 | 128 | 97,005 |

Table S94: Results obtained by dnadiff [4] when analysing the reconstruction of DS27. Only one execution of HVRS is represented.

| Reconstruction tool | Total seqs | Aligned seqs | Aligned bases | Aligned bases (%) | Unaligned bases | Unaligned bases (%) | Average length | SNPs | Identity |
|---|---|---|---|---|---|---|---|---|---|
| coronaSPAdes | 165 | 121 | 138722 | 68.4939 | 63810 | 31.5061 | 1157.6218 | 176 | 99,790 |
| Haploflow | – | – | – | – | – | – | – | – | – |
| IRMA | 5 | 4 | 123765 | 88.5883 | 15943 | 11.4117 | 4760.9615 | 50 | 98,694 |
| LAZYPIPE | 150 | 107 | 126613 | 67.8032 | 60123 | 32.1968 | 1175.1321 | 60 | 99,934 |
| metaSPAdes | 315 | 220 | 120372 | 68.1122 | 56354 | 31.8878 | 547.9816 | 142 | 99,854 |
| metaviralSPAdes | – | – | – | – | – | – | – | – | – |
| PEHaplo | 170 | 120 | 137977 | 68.6006 | 63154 | 31.3994 | 1145.8017 | 348 | 99,603 |
| QuRe | – | – | – | – | – | – | – | – | – |
| QVG | 4 | 4 | 143858 | 99.9986 | 2 | 0.0014 | 35964.5000 | 5172 | 96,400 |
| SPAdes | 160 | 116 | 135761 | 68.0264 | 63810 | 31.9736 | 1182.3304 | 188 | 99,772 |
| SSAKE | 75 | 51 | 8352 | 69.7104 | 3629 | 30.2896 | 163.7647 | 1 | 99,988 |
| TRACESPipe | 4 | 4 | 142304 | 98.9177 | 1557 | 1.0823 | 10164.5714 | 670 | 98,687 |
| TRACESPipeLite | 6 | 4 | 85311 | 50.7013 | 82951 | 49.2987 | 543.8258 | 111 | 97,683 |
| V-pipe | 5 | 4 | 63188 | 39.3869 | 97241 | 60.6131 | 284.2133 | 61 | 95,529 |
| VirGenA | 120 | 120 | 74055 | 99.6622 | 251 | 0.3378 | 563.0385 | 158 | 99,568 |
| ViSpA | 78 | 70 | 3285350 | 97.6003 | 80777 | 2.3997 | 2079.4355 | 620 | 96,542 |

Table S95: Results obtained by dnadiff [4] when analysing the reconstruction of DS28. Only one execution of HVRS is represented.

| Reconstruction tool | Total seqs | Aligned seqs | Aligned bases | Aligned bases (%) | Unaligned bases | Unaligned bases (%) | Average length | SNPs | Identity |
|---|---|---|---|---|---|---|---|---|---|
| coronaSPAdes | 156 | 103 | 136938 | 68.1216 | 64082 | 31.8784 | 1340.0294 | 97 | 99,840 |
| Haploflow | – | – | – | – | – | – | – | – | – |
| IRMA | 5 | 4 | 122469 | 88.3985 | 16073 | 11.6015 | 4224.1724 | 78 | 98,519 |
| LAZYPIPE | 149 | 97 | 127856 | 68.0183 | 60117 | 31.9817 | 1318.1031 | 24 | 99,940 |
| metaSPAdes | 338 | 230 | 107948 | 67.5093 | 51953 | 32.4907 | 469.6594 | 105 | 99,901 |
| metaviralSPAdes | – | – | – | – | – | – | – | – | – |
| PEHaplo | 166 | 115 | 158080 | 71.5191 | 62952 | 28.4809 | 1281.1509 | 323 | 99,525 |
| QuRe | – | – | – | – | – | – | – | – | – |
| QVG | 4 | 4 | 143860 | 100.0000 | 0 | 0.0000 | 35965.0000 | 7602 | 94,671 |
| SPAdes | 156 | 103 | 136919 | 68.1172 | 64086 | 31.8828 | 1337.9903 | 61 | 99,850 |
| SSAKE | 65 | 45 | 6832 | 69.8569 | 2948 | 30.1431 | 151.8222 | 0 | 100,000 |
| TRACESPipe | 4 | 4 | 141043 | 98.0432 | 2815 | 1.9568 | 7835.5556 | 1322 | 98,045 |
| TRACESPipeLite | 6 | 3 | 19599 | 11.6479 | 148663 | 88.3521 | 283.7391 | 9 | 98,176 |
| V-pipe | 5 | 4 | 63775 | 39.7528 | 96654 | 60.2472 | 302.8990 | 182 | 94,928 |
| VirGenA | 126 | 125 | 71184 | 99.2277 | 554 | 0.7723 | 544.1439 | 126 | 99,474 |
| ViSpA | 90 | 76 | 3190744 | 95.9756 | 133794 | 4.0244 | 1778.5270 | 911 | 96,581 |

Table S96: Results obtained by dnadiff [4] when analysing the reconstruction of DS29. Only one execution of HVRS is represented.

| Reconstruction tool | Total seqs | Aligned seqs | Aligned bases | Aligned bases (%) | Unaligned bases | Unaligned bases (%) | Average length | SNPs | Identity |
|---|---|---|---|---|---|---|---|---|---|
| coronaSPAdes | 161 | 111 | 134153 | 67.5633 | 64406 | 32.4367 | 1208.5856 | 88 | 99,934 |
| Haploflow | 1 | 0 | 0 | 0.0000 | 511 | 100.0000 | 0.0000 | 0 | 0,000 |
| IRMA | 5 | 4 | 122742 | 88.4933 | 15960 | 11.5067 | 3610.6765 | 47 | 98,493 |
| LAZYPIPE | 160 | 111 | 127975 | 67.9150 | 60459 | 32.0850 | 1152.9279 | 17 | 99,987 |
| metaSPAdes | 317 | 211 | 100064 | 66.7129 | 49928 | 33.2871 | 473.7048 | 90 | 99,908 |
| metaviralSPAdes | – | – | – | – | – | – | – | – | – |
| PEHaplo | 170 | 120 | 133304 | 67.6018 | 63886 | 32.3982 | 1115.9496 | 212 | 99,763 |
| QuRe | – | – | – | – | – | – | – | – | – |
| QVG | 4 | 4 | 143813 | 99.9673 | 47 | 0.0327 | 35953.2500 | 9748 | 93,194 |
| SPAdes | 163 | 113 | 135271 | 67.7449 | 64406 | 32.2551 | 1197.0885 | 45 | 99,965 |
| SSAKE | 68 | 50 | 7572 | 72.9550 | 2807 | 27.0450 | 151.4400 | 0 | 100,000 |
| TRACESPipe | 4 | 4 | 136260 | 94.7171 | 7600 | 5.2829 | 3696.7222 | 1149 | 97,348 |
| TRACESPipeLite | 6 | 2 | 2485 | 1.4769 | 165777 | 98.5231 | 225.9091 | 0 | 98,534 |
| V-pipe | 5 | 4 | 63104 | 39.3345 | 97325 | 60.6655 | 299.6019 | 187 | 94,365 |
| VirGenA | 121 | 121 | 63149 | 99.3112 | 438 | 0.6888 | 490.2266 | 94 | 99,752 |
| ViSpA | 107 | 97 | 4450371 | 97.9400 | 93605 | 2.0600 | 1215.4554 | 807 | 95,473 |

Table S97: Results obtained by dnadiff [4] when analysing the reconstruction of DS30. Only one execution of HVRS is represented.

| Reconstruction tool | Total seqs | Aligned seqs | Aligned bases | Aligned bases (%) | Unaligned bases | Unaligned bases (%) | Average length | SNPs | Identity |
|---|---|---|---|---|---|---|---|---|---|
| coronaSPAdes | 167 | 114 | 135762 | 68.5656 | 62241 | 31.4344 | 1190.8947 | 72 | 99,946 |
| Haploflow | 1 | 1 | 540 | 100.0000 | 0 | 0.0000 | 540.0000 | 0 | 100,000 |
| IRMA | 5 | 4 | 123633 | 88.6017 | 15905 | 11.3983 | 3254.7895 | 41 | 98,577 |
| LAZYPIPE | 155 | 110 | 126982 | 69.1808 | 56569 | 30.8192 | 1154.3818 | 24 | 99,981 |
| metaSPAdes | 313 | 210 | 116656 | 68.2822 | 54188 | 31.7178 | 555.5048 | 90 | 99,918 |
| metaviralSPAdes | – | – | – | – | – | – | – | – | – |
| PEHaplo | 168 | 112 | 135846 | 67.8867 | 64261 | 32.1133 | 1205.4414 | 36 | 99,943 |
| QuRe | – | – | – | – | – | – | – | – | – |
| QVG | 4 | 4 | 143860 | 100.0000 | 0 | 0.0000 | 35965.0000 | 12704 | 91,129 |
| SPAdes | 167 | 114 | 135875 | 68.5804 | 62250 | 31.4196 | 1198.9027 | 89 | 99,931 |
| SSAKE | 81 | 52 | 8211 | 62.9581 | 4831 | 37.0419 | 157.9038 | 1 | 99,988 |
| TRACESPipe | 4 | 4 | 131760 | 91.5897 | 12099 | 8.4103 | 2787.9348 | 1352 | 97,630 |
| TRACESPipeLite | 6 | 0 | 0 | 0.0000 | 168262 | 100.0000 | 0.0000 | 0 | 0,000 |
| V-pipe | 5 | 4 | 55920 | 34.8565 | 104509 | 65.1435 | 273.0493 | 173 | 94,003 |
| VirGenA | 112 | 109 | 61705 | 99.4039 | 370 | 0.5961 | 524.6864 | 32 | 99,755 |
| ViSpA | 133 | 128 | 5342677 | 98.6114 | 75231 | 1.3886 | 724.9864 | 688 | 94,350 |

Table S98: Results obtained by dnadiff [4] when analysing the reconstruction of DS31. Only one execution of HVRS is represented.

| Reconstruction tool | Total seqs | Aligned seqs | Aligned bases | Aligned bases (%) | Unaligned bases | Unaligned bases (%) | Average length | SNPs | Identity |
|---|---|---|---|---|---|---|---|---|---|
| coronaSPAdes | 171 | 117 | 136962 | 68.3948 | 63290 | 31.6052 | 1170.6154 | 75 | 99,943 |
| Haploflow | 1 | 1 | 533 | 100.0000 | 0 | 0.0000 | 533.0000 | 0 | 100,000 |
| IRMA | 5 | 4 | 122039 | 88.6242 | 15665 | 11.3758 | 2491.7143 | 40 | 98,481 |
| LAZYPIPE | 164 | 112 | 127384 | 68.2292 | 59316 | 31.7708 | 1137.3571 | 20 | 99,984 |
| metaSPAdes | 322 | 227 | 104046 | 68.4747 | 47902 | 31.5253 | 458.3524 | 84 | 99,913 |
| metaviralSPAdes | – | – | – | – | – | – | – | – | – |
| PEHaplo | 162 | 112 | 133603 | 68.9795 | 60082 | 31.0205 | 1192.8839 | 364 | 99,956 |
| QuRe | – | – | – | – | – | – | – | – | – |
| QVG | 4 | 4 | 143820 | 99.9708 | 42 | 0.0292 | 35955.0000 | 16044 | 88,634 |
| SPAdes | 171 | 117 | 137018 | 68.4033 | 63291 | 31.5967 | 1171.0940 | 44 | 99,966 |
| SSAKE | 61 | 43 | 6817 | 70.8628 | 2803 | 29.1372 | 158.5349 | 0 | 100,000 |
| TRACESPipe | 4 | 4 | 128565 | 89.3681 | 15295 | 10.6319 | 2272.0714 | 1812 | 97,055 |
| TRACESPipeLite | 6 | 0 | 0 | 0.0000 | 168262 | 100.0000 | 0.0000 | 0 | 0,000 |
| V-pipe | 5 | 4 | 45781 | 28.5366 | 114648 | 71.4634 | 235.2092 | 162 | 93,088 |
| VirGenA | 105 | 105 | 70354 | 99.2929 | 501 | 0.7071 | 602.2564 | 34 | 99,883 |
| ViSpA | 107 | 96 | 2925222 | 96.1664 | 116612 | 3.8336 | 419.7128 | 292 | 94,943 |

Table S99: Results obtained by dnadiff [4] when analysing the reconstruction of DS32. Only one execution of HVRS is represented.

| Reconstruction tool | Total seqs | Aligned seqs | Aligned bases | Aligned bases (%) | Unaligned bases | Unaligned bases (%) | Average length | SNPs | Identity |
|---|---|---|---|---|---|---|---|---|---|
| coronaSPAdes | 173 | 116 | 137962 | 68.5107 | 63411 | 31.4893 | 1189.3276 | 66 | 99,951 |
| Haploflow | – | – | – | – | – | – | – | – | – |
| IRMA | 5 | 4 | 121836 | 88.1662 | 16353 | 11.8338 | 1792.6618 | 58 | 98,531 |
| LAZYPIPE | 172 | 118 | 130397 | 68.9563 | 58704 | 31.0437 | 1105.0593 | 19 | 99,986 |
| metaSPAdes | 319 | 218 | 121490 | 69.5110 | 53288 | 30.4890 | 557.2936 | 75 | 99,936 |
| metaviralSPAdes | – | – | – | – | – | – | – | – | – |
| PEHaplo | 162 | 112 | 134948 | 68.6758 | 61552 | 31.3242 | 1204.8929 | 34 | 99,963 |
| QuRe | – | – | – | – | – | – | – | – | – |
| QVG | 4 | 4 | 143849 | 99.9937 | 9 | 0.0063 | 35962.2500 | 19218 | 86,483 |
| SPAdes | 173 | 116 | 138178 | 68.5437 | 63413 | 31.4563 | 1191.1897 | 44 | 99,967 |
| SSAKE | 74 | 46 | 6815 | 60.2191 | 4502 | 39.7809 | 148.1522 | 0 | 100,000 |
| TRACESPipe | 4 | 4 | 116877 | 81.2696 | 26937 | 18.7304 | 1523.8816 | 1406 | 97,524 |
| TRACESPipeLite | 6 | 0 | 0 | 0.0000 | 168289 | 100.0000 | 0.0000 | 0 | 0,000 |
| V-pipe | 5 | 4 | 32649 | 20.3511 | 127780 | 79.6489 | 195.5030 | 72 | 93,219 |
| VirGenA | 129 | 127 | 62578 | 98.8204 | 747 | 1.1796 | 453.0647 | 63 | 99,640 |
| ViSpA | 54 | 46 | 952339 | 91.2954 | 90801 | 8.7046 | 263.7931 | 249 | 95,949 |

Table S100: Results obtained by dnadiff [4] when analysing the reconstruction of DS33. Only one execution of HVRS is represented.

| Reconstruction tool | Total seqs | Aligned seqs | Aligned bases | Aligned bases (%) | Unaligned bases | Unaligned bases (%) | Average length | SNPs | Identity |
|---|---|---|---|---|---|---|---|---|---|
| coronaSPAdes | 27 | 20 | 135645 | 67.0872 | 66547 | 32.9128 | 6464.4762 | 10 | 99,908 |
| Haploflow | 106 | 72 | 59512 | 70.5930 | 24791 | 29.4070 | 826.5556 | 0 | 100,000 |
| IRMA | 5 | 4 | 128071 | 88.5459 | 16567 | 11.4541 | 30076.5000 | 8 | 99,947 |
| LAZYPIPE | 24 | 17 | 134839 | 67.0794 | 66175 | 32.9206 | 6749.3000 | 4 | 99,995 |
| metaSPAdes | 27 | 20 | 135717 | 67.0989 | 66547 | 32.9011 | 6467.3333 | 8 | 99,858 |
| metaviralSPAdes | – | – | – | – | – | – | – | – | – |
| PEHaplo | 208 | 144 | 152091 | 67.2642 | 74019 | 32.7358 | 1041.5972 | 16 | 99,983 |
| QuRe | 3 | 1 | 71 | 47.0199 | 80 | 52.9801 | 71.0000 | 0 | 100,000 |
| QVG | – | – | – | – | – | – | – | – | – |
| SPAdes | 26 | 19 | 135760 | 67.1059 | 66547 | 32.8941 | 6473.8571 | 5 | 99,863 |
| SSAKE | 120 | 83 | 15779 | 69.9052 | 6793 | 30.0948 | 190.1084 | 1 | 99,994 |
| TRACESPipe | 4 | 4 | 143815 | 99.9687 | 45 | 0.0313 | 35953.7500 | 0 | 99,957 |
| TRACESPipeLite | 6 | 5 | 143992 | 85.5761 | 24270 | 14.4239 | 35899.5000 | 0 | 99,909 |
| V-pipe | 5 | 4 | 129232 | 80.5540 | 31197 | 19.4460 | 1621.7564 | 5 | 96,583 |
| VirGenA | 85 | 85 | 68280 | 98.8949 | 763 | 1.1051 | 727.4362 | 21 | 99,894 |
| ViSpA | 11 | 10 | 437940 | 96.4088 | 16313 | 3.5912 | 21688.2500 | 4 | 99,630 |

Table S101: Results obtained by dnadiff [4] when analysing the reconstruction of DS34. Only one execution of HVRS is represented.

| Reconstruction tool | Total seqs | Aligned seqs | Aligned bases | Aligned bases (%) | Unaligned bases | Unaligned bases (%) | Average length | SNPs | Identity |
|---|---|---|---|---|---|---|---|---|---|
| coronaSPAdes | 33 | 23 | 141458 | 68.0244 | 66494 | 31.9756 | 6417.0909 | 55 | 99,960 |
| Haploflow | 103 | 69 | 56298 | 70.3312 | 23749 | 29.6688 | 819.5735 | 44 | 99,921 |
| IRMA | 5 | 4 | 128125 | 88.6408 | 16419 | 11.3592 | 25625.2000 | 1 | 99,951 |
| LAZYPIPE | 32 | 22 | 142705 | 68.3600 | 66050 | 31.6400 | 8057.4706 | 148 | 99,856 |
| metaSPAdes | 39 | 29 | 141614 | 68.0483 | 66494 | 31.9517 | 5048.4643 | 40 | 99,924 |
| metaviralSPAdes | – | – | – | – | – | – | – | – | – |
| PEHaplo | 240 | 170 | 170017 | 69.9345 | 73092 | 30.0655 | 984.4410 | 232 | 99,753 |
| QuRe | 1 | 1 | 94 | 100.0000 | 0 | 0.0000 | 94.0000 | 0 | 100,000 |
| QVG | 4 | 4 | 143860 | 100.0000 | 0 | 0.0000 | 35965.0000 | 1388 | 98,963 |
| SPAdes | 34 | 24 | 142811 | 68.2304 | 66496 | 31.7696 | 6772.8571 | 20 | 99,918 |
| SSAKE | 118 | 76 | 13291 | 62.4548 | 7990 | 37.5452 | 174.8816 | 1 | 99,993 |
| TRACESPipe | 4 | 4 | 143608 | 99.8234 | 254 | 0.1766 | 28793.8000 | 511 | 99,591 |
| TRACESPipeLite | 6 | 4 | 142274 | 84.5550 | 25988 | 15.4450 | 11856.1667 | 426 | 99,119 |
| V-pipe | 5 | 4 | 126584 | 78.9034 | 33845 | 21.0966 | 1301.9574 | 297 | 96,547 |
| VirGenA | 86 | 85 | 59524 | 99.1059 | 537 | 0.8941 | 643.8000 | 80 | 99,750 |
| ViSpA | 11 | 9 | 289066 | 91.8962 | 25491 | 8.1038 | 17846.3750 | 470 | 99,476 |

Table S102: Results obtained by dnadiff [4] when analysing the reconstruction of DS35. Only one execution of HVRS is represented.

| Reconstruction tool | Total seqs | Aligned seqs | Aligned bases | Aligned bases (%) | Unaligned bases | Unaligned bases (%) | Average length | SNPs | Identity |
|---|---|---|---|---|---|---|---|---|---|
| coronaSPAdes | 25 | 17 | 143450 | 68.3375 | 66464 | 31.6625 | 8438.2353 | 28 | 99,976 |
| Haploflow | 101 | 69 | 52541 | 69.0312 | 23571 | 30.9688 | 752.6714 | 34 | 99,907 |
| IRMA | 5 | 4 | 127959 | 88.5529 | 16541 | 11.4471 | 25591.8000 | 7 | 99,851 |
| LAZYPIPE | 27 | 18 | 142502 | 68.3135 | 66098 | 31.6865 | 7518.6842 | 9 | 99,876 |
| metaSPAdes | 30 | 21 | 143105 | 68.2580 | 66548 | 31.7420 | 6814.5238 | 29 | 99,974 |
| metaviralSPAdes | – | – | – | – | – | – | – | – | – |
| PEHaplo | 221 | 160 | 162054 | 68.8534 | 73307 | 31.1466 | 1020.8312 | 114 | 99,787 |
| QuRe | – | – | – | – | – | – | – | – | – |
| QVG | 4 | 4 | 143858 | 99.9986 | 2 | 0.0014 | 35964.5000 | 2184 | 98,482 |
| SPAdes | 28 | 20 | 143379 | 68.3268 | 66464 | 31.6732 | 9595.7333 | 2 | 99,508 |
| SSAKE | 116 | 82 | 14497 | 69.6971 | 6303 | 30.3029 | 176.7927 | 1 | 99,993 |
| TRACESPipe | 4 | 4 | 143550 | 99.7845 | 310 | 0.2155 | 35887.5000 | 665 | 99,454 |
| TRACESPipeLite | 6 | 4 | 102550 | 60.9466 | 65712 | 39.0534 | 779.6562 | 308 | 97,694 |
| V-pipe | 5 | 4 | 126469 | 78.8318 | 33960 | 21.1682 | 1356.4556 | 271 | 96,769 |
| VirGenA | 60 | 60 | 69319 | 99.3579 | 448 | 0.6421 | 875.3462 | 164 | 99,505 |
| ViSpA | 11 | 10 | 489793 | 96.7489 | 16459 | 3.2511 | 15837.8889 | 647 | 99,377 |

Table S103: Results obtained by dnadiff [4] when analysing the reconstruction of DS36. Only one execution of HVRS is represented.

| Reconstruction tool | Total seqs | Aligned seqs | Aligned bases | Aligned bases (%) | Unaligned bases | Unaligned bases (%) | Average length | SNPs | Identity |
|---|---|---|---|---|---|---|---|---|---|
| coronaSPAdes | 16 | 7 | 143569 | 68.3564 | 66461 | 31.6436 | 20509.8571 | 19 | 99,986 |
| Haploflow | 109 | 74 | 54771 | 66.3449 | 27784 | 33.6551 | 740.1486 | 21 | 99,886 |
| IRMA | 5 | 4 | 128093 | 88.5648 | 16539 | 11.4352 | 25618.6000 | 4 | 99,975 |
| LAZYPIPE | 17 | 8 | 143217 | 68.4165 | 66114 | 31.5835 | 17902.1250 | 8 | 99,953 |
| metaSPAdes | 17 | 8 | 143644 | 68.3677 | 66461 | 31.6323 | 17955.5000 | 16 | 99,986 |
| metaviralSPAdes | – | – | – | – | – | – | – | – | – |
| PEHaplo | 200 | 143 | 157637 | 68.6342 | 72040 | 31.3658 | 1096.0634 | 157 | 99,806 |
| QuRe | 2 | 1 | 70 | 59.8291 | 47 | 40.1709 | 70.0000 | 0 | 100,000 |
| QVG | 4 | 4 | 143860 | 100.0000 | 0 | 0.0000 | 35965.0000 | 3358 | 97,624 |
| SPAdes | 17 | 8 | 143665 | 68.3709 | 66461 | 31.6291 | 17958.1250 | 9 | 99,989 |
| SSAKE | 112 | 74 | 13967 | 67.5485 | 6710 | 32.4515 | 188.7432 | 0 | 100,000 |
| TRACESPipe | 4 | 4 | 143539 | 99.7706 | 330 | 0.2294 | 28707.8000 | 793 | 99,225 |
| TRACESPipeLite | 6 | 4 | 29276 | 17.3991 | 138986 | 82.6009 | 312.7368 | 63 | 98,598 |
| V-pipe | 5 | 4 | 125828 | 78.4322 | 34601 | 21.5678 | 1312.6809 | 424 | 95,936 |
| VirGenA | 89 | 89 | 62324 | 99.4939 | 317 | 0.5061 | 645.8211 | 135 | 99,656 |
| ViSpA | 15 | 13 | 429646 | 93.9939 | 27454 | 6.0061 | 19072.7500 | 632 | 98,667 |

Table S104: Results obtained by dnadiff [4] when analysing the reconstruction of DS37. Only one execution of HVRS is represented.

| Reconstruction tool | Total seqs | Aligned seqs | Aligned bases | Aligned bases (%) | Unaligned bases | Unaligned bases (%) | Average length | SNPs | Identity |
|---|---|---|---|---|---|---|---|---|---|
| coronaSPAdes | 24 | 16 | 143707 | 68.3746 | 66469 | 31.6254 | 8981.6875 | 12 | 99,990 |
| Haploflow | 98 | 66 | 52403 | 66.6205 | 26256 | 33.3795 | 796.9692 | 134 | 99,741 |
| IRMA | 5 | 4 | 128872 | 88.6291 | 16534 | 11.3709 | 14321.2222 | 13 | 99,905 |
| LAZYPIPE | 26 | 18 | 143054 | 68.3975 | 66097 | 31.6025 | 7947.4444 | 4 | 99,995 |
| metaSPAdes | 24 | 16 | 143742 | 68.3799 | 66469 | 31.6201 | 8983.8750 | 7 | 99,994 |
| metaviralSPAdes | – | – | – | – | – | – | – | – | – |
| PEHaplo | 202 | 137 | 158122 | 68.5469 | 72555 | 31.4531 | 1151.0224 | 109 | 99,925 |
| QuRe | 3 | 2 | 236 | 70.6587 | 98 | 29.3413 | 118.0000 | 0 | 100,000 |
| QVG | 4 | 4 | 143813 | 99.9673 | 47 | 0.0327 | 35953.2500 | 4998 | 96,505 |
| SPAdes | 24 | 16 | 143742 | 68.3799 | 66469 | 31.6201 | 8983.8750 | 9 | 99,991 |
| SSAKE | 113 | 78 | 14434 | 68.9995 | 6485 | 31.0005 | 185.0513 | 2 | 99,986 |
| TRACESPipe | 4 | 4 | 141961 | 98.6800 | 1899 | 1.3200 | 19776.8571 | 530 | 99,500 |
| TRACESPipeLite | 6 | 2 | 4163 | 2.4741 | 164099 | 97.5259 | 219.1053 | 1 | 99,976 |
| V-pipe | 5 | 4 | 125968 | 78.5195 | 34461 | 21.4805 | 1258.7938 | 689 | 95,647 |
| VirGenA | 81 | 81 | 62280 | 99.9904 | 6 | 0.0096 | 655.4105 | 61 | 99,705 |
| ViSpA | 41 | 40 | 2112414 | 99.2252 | 16495 | 0.7748 | 6898.8500 | 1051 | 97,848 |

Table S105: Results obtained by dnadiff [4] when analysing the reconstruction of DS38. Only one execution of HVRS is represented.

| Reconstruction tool | Total seqs | Aligned seqs | Aligned bases | Aligned bases (%) | Unaligned bases | Unaligned bases (%) | Average length | SNPs | Identity |
|---|---|---|---|---|---|---|---|---|---|
| coronaSPAdes | 19 | 16 | 143610 | 68.3428 | 66522 | 31.6572 | 8975.6250 | 13 | 99,989 |
| Haploflow | 104 | 73 | 60641 | 70.4121 | 25482 | 29.5879 | 830.6986 | 0 | 100,000 |
| IRMA | 5 | 4 | 128992 | 88.6336 | 16542 | 11.3664 | 9214.3571 | 8 | 99,852 |
| LAZYPIPE | 22 | 18 | 143056 | 68.2949 | 66412 | 31.7051 | 7947.5556 | 4 | 99,995 |
| metaSPAdes | 20 | 17 | 143706 | 68.3572 | 66522 | 31.6428 | 8453.2941 | 7 | 99,992 |
| metaviralSPAdes | – | – | – | – | – | – | – | – | – |
| PEHaplo | 192 | 134 | 157837 | 68.6567 | 72056 | 31.3433 | 1193.7727 | 4 | 99,615 |
| QuRe | 2 | 2 | 256 | 100.0000 | 0 | 0.0000 | 128.0000 | 0 | 100,000 |
| QVG | 4 | 4 | 143863 | 100.0000 | 0 | 0.0000 | 35965.7500 | 7540 | 94,730 |
| SPAdes | 18 | 15 | 143563 | 68.3357 | 66522 | 31.6643 | 9570.8667 | 5 | 99,993 |
| SSAKE | 112 | 79 | 14076 | 68.8077 | 6381 | 31.1923 | 178.1772 | 0 | 100,000 |
| TRACESPipe | 4 | 4 | 141458 | 98.3303 | 2402 | 1.6697 | 23576.3333 | 758 | 99,217 |
| TRACESPipeLite | 6 | 1 | 434 | 0.2579 | 167828 | 99.7421 | 217.0000 | 0 | 100,000 |
| V-pipe | 5 | 4 | 118868 | 74.0938 | 41561 | 25.9062 | 996.4262 | 78 | 94,147 |
| VirGenA | 79 | 78 | 52175 | 99.2788 | 379 | 0.7212 | 544.9688 | 77 | 99,786 |
| ViSpA | 88 | 87 | 3699971 | 99.4976 | 18682 | 0.5024 | 2102.8525 | 915 | 97,046 |

Table S106: Results obtained by dnadiff [4] when analysing the reconstruction of DS39. Only one execution of HVRS is represented.

| Reconstruction tool | Total seqs | Aligned seqs | Aligned bases | Aligned bases (%) | Unaligned bases | Unaligned bases (%) | Average length | SNPs | Identity |
|---|---|---|---|---|---|---|---|---|---|
| coronaSPAdes | 26 | 22 | 143612 | 68.4267 | 66265 | 31.5733 | 6527.8182 | 4 | 99,997 |
| Haploflow | 112 | 76 | 59297 | 68.1011 | 27775 | 31.8989 | 780.2237 | 0 | 100,000 |
| IRMA | 5 | 4 | 129301 | 88.6127 | 16616 | 11.3873 | 5622.6522 | 58 | 99,818 |
| LAZYPIPE | 27 | 23 | 142691 | 68.3697 | 66014 | 31.6303 | 6203.9565 | 3 | 99,998 |
| metaSPAdes | 26 | 22 | 143612 | 68.4267 | 66265 | 31.5733 | 6527.8182 | 2 | 99,998 |
| metaviralSPAdes | – | – | – | – | – | – | – | – | – |
| PEHaplo | 207 | 137 | 158836 | 68.0388 | 74613 | 31.9612 | 1177.6692 | 3 | 99,997 |
| QuRe | 3 | 3 | 504 | 100.0000 | 0 | 0.0000 | 168.0000 | 0 | 100,000 |
| QVG | 4 | 4 | 143816 | 99.9708 | 42 | 0.0292 | 35954.0000 | 11690 | 91,790 |
| SPAdes | 26 | 22 | 143612 | 68.4267 | 66265 | 31.5733 | 6527.8182 | 2 | 99,998 |
| SSAKE | 119 | 81 | 16045 | 69.1386 | 7162 | 30.8614 | 198.0864 | 0 | 100,000 |
| TRACESPipe | 4 | 4 | 141230 | 98.1718 | 2630 | 1.8282 | 12042.0833 | 345 | 98,700 |
| TRACESPipeLite | 5 | 0 | 0 | 0.0000 | 160429 | 100.0000 | 0.0000 | 0 | 0,000 |
| V-pipe | 5 | 4 | 103260 | 64.3649 | 57169 | 35.6351 | 603.4588 | 457 | 93,037 |
| VirGenA | 85 | 85 | 58507 | 99.3699 | 371 | 0.6301 | 637.5714 | 66 | 99,810 |
| ViSpA | 139 | 138 | 5096959 | 99.5460 | 23245 | 0.4540 | 687.4257 | 490 | 95,815 |

Table S107: Results obtained by dnadiff [4] when analysing the reconstruction of DS40. Only one execution of HVRS is represented.

| Reconstruction tool | Total seqs | Aligned seqs | Aligned bases | Aligned bases (%) | Unaligned bases | Unaligned bases (%) | Average length | SNPs | Identity |
|---|---|---|---|---|---|---|---|---|---|
| coronaSPAdes | 19 | 13 | 143535 | 68.3692 | 66406 | 31.6308 | 11041.1538 | 12 | 99,991 |
| Haploflow | 100 | 70 | 52778 | 69.8602 | 22770 | 30.1398 | 753.9714 | 0 | 100,000 |
| IRMA | 5 | 4 | 129116 | 88.4671 | 16832 | 11.5329 | 4310.2000 | 8 | 99,770 |
| LAZYPIPE | 19 | 13 | 142865 | 68.3522 | 66148 | 31.6478 | 10989.6154 | 4 | 99,996 |
| metaSPAdes | 19 | 13 | 143535 | 68.3692 | 66406 | 31.6308 | 11041.1538 | 7 | 99,993 |
| metaviralSPAdes | – | – | – | – | – | – | – | – | – |
| PEHaplo | 201 | 148 | 163127 | 68.5269 | 74921 | 31.4731 | 1080.3878 | 3 | 99,997 |
| QuRe | 1 | 1 | 164 | 100.0000 | 0 | 0.0000 | 164.0000 | 0 | 100,000 |
| QVG | 4 | 4 | 143849 | 99.9937 | 9 | 0.0063 | 35962.2500 | 15561 | 89,044 |
| SPAdes | 19 | 13 | 143535 | 68.3692 | 66406 | 31.6308 | 11041.1538 | 5 | 99,996 |
| SSAKE | 118 | 78 | 14256 | 65.7201 | 7436 | 34.2799 | 182.7692 | 0 | 100,000 |
| TRACESPipe | 4 | 4 | 136021 | 94.5510 | 7839 | 5.4490 | 7159.0000 | 1647 | 98,520 |
| TRACESPipeLite | 5 | 0 | 0 | 0.0000 | 160429 | 100.0000 | 0.0000 | 0 | 0,000 |
| V-pipe | 5 | 4 | 80756 | 50.3375 | 79673 | 49.6625 | 412.8918 | 212 | 91,872 |
| VirGenA | 71 | 71 | 66652 | 99.5534 | 299 | 0.4466 | 806.4268 | 142 | 99,743 |
| ViSpA | 115 | 114 | 3266451 | 98.9620 | 34263 | 1.0380 | 360.3264 | 228 | 95,858 |

Table S108: Results obtained by dnadiff [4] when analysing the reconstruction of DS41. Only one execution of HVRS is represented.

| Reconstruction tool | Total seqs | Aligned seqs | Aligned bases | Aligned bases (%) | Unaligned bases | Unaligned bases (%) | Average length | SNPs | Identity |
|---|---|---|---|---|---|---|---|---|---|
| coronaSPAdes | 9 | 7 | 135952 | 67.1487 | 66512 | 32.8513 | 22627.6667 | 1 | 99,900 |
| Haploflow | 53 | 37 | 136232 | 67.5871 | 65333 | 32.4129 | 3699.4167 | 13 | 99,976 |
| IRMA | 5 | 4 | 128291 | 88.5810 | 16538 | 11.4190 | 25658.2000 | 0 | 100,000 |
| LAZYPIPE | 7 | 5 | 135740 | 67.1239 | 66483 | 32.8761 | 17046.7500 | 0 | 99,971 |
| metaSPAdes | 9 | 7 | 135754 | 67.0499 | 66713 | 32.9501 | 15083.7778 | 0 | 99,952 |
| metaviralSPAdes | – | – | – | – | – | – | – | – | – |
| PEHaplo | 312 | 211 | 172528 | 68.2638 | 80209 | 31.7362 | 811.3268 | 0 | 100,000 |
| QuRe | 6 | 5 | 786 | 79.0744 | 208 | 20.9256 | 157.2000 | 0 | 100,000 |
| QVG | – | – | – | – | – | – | – | – | – |
| SPAdes | 8 | 6 | 136077 | 67.1687 | 66513 | 32.8313 | 22676.5000 | 0 | 99,917 |
| SSAKE | 375 | 254 | 83659 | 69.1791 | 37272 | 30.8209 | 328.5882 | 6 | 99,940 |
| TRACESPipe | 4 | 4 | 143743 | 99.9187 | 117 | 0.0813 | 35935.7500 | 0 | 100,000 |
| TRACESPipeLite | 6 | 5 | 144112 | 85.6474 | 24150 | 14.3526 | 35932.2500 | 1 | 99,991 |
| V-pipe | 5 | 4 | 143178 | 89.2470 | 17251 | 10.7530 | 28635.6000 | 0 | 99,837 |
| VirGenA | – | – | – | – | – | – | – | – | – |
| ViSpA | 5 | 4 | 143596 | 89.6808 | 16523 | 10.3192 | 35899.0000 | 0 | 100,000 |

Table S109: Results obtained by dnadiff [4] when analysing the reconstruction of DS42. Only one execution of HVRS is represented.

| Reconstruction tool | Total seqs | Aligned seqs | Aligned bases | Aligned bases (%) | Unaligned bases | Unaligned bases (%) | Average length | SNPs | Identity |
|---|---|---|---|---|---|---|---|---|---|
| coronaSPAdes | 13 | 11 | 141427 | 68.0166 | 66503 | 31.9834 | 12857.0000 | 7 | 99,995 |
| Haploflow | 57 | 40 | 141863 | 68.5948 | 64950 | 31.4052 | 3563.6410 | 142 | 99,844 |
| IRMA | 5 | 4 | 128272 | 88.5832 | 16532 | 11.4168 | 25654.4000 | 0 | 100,000 |
| LAZYPIPE | 10 | 8 | 143546 | 68.3552 | 66454 | 31.6448 | 27438.6000 | 291 | 99,786 |
| metaSPAdes | 7 | 5 | 142230 | 68.1071 | 66603 | 31.8929 | 28240.6000 | 84 | 99,944 |
| metaviralSPAdes | – | – | – | – | – | – | – | – | – |
| PEHaplo | 325 | 227 | 185352 | 70.4728 | 77660 | 29.5272 | 800.9231 | 108 | 99,888 |
| QuRe | 5 | 5 | 3332 | 99.9100 | 3 | 0.0900 | 666.4000 | 17 | 99,457 |
| QVG | 4 | 4 | 143860 | 100.0000 | 0 | 0.0000 | 35965.0000 | 551 | 99,537 |
| SPAdes | 7 | 5 | 143760 | 68.3715 | 66503 | 31.6285 | 28752.0000 | 2 | 99,999 |
| SSAKE | 341 | 230 | 73715 | 68.6519 | 33660 | 31.3481 | 323.0177 | 9 | 99,979 |
| TRACESPipe | 4 | 4 | 143774 | 99.9402 | 86 | 0.0598 | 28827.0000 | 438 | 99,663 |
| TRACESPipeLite | 6 | 4 | 143111 | 85.0525 | 25151 | 14.9475 | 17888.6250 | 479 | 99,338 |
| V-pipe | 5 | 4 | 142834 | 89.0325 | 17595 | 10.9675 | 25481.8333 | 362 | 99,398 |
| VirGenA | 46 | 46 | 56111 | 98.5527 | 824 | 1.4473 | 971.4386 | 155 | 99,655 |
| ViSpA | 5 | 4 | 143596 | 89.6920 | 16503 | 10.3080 | 35899.0000 | 444 | 99,676 |

Table S110: Results obtained by dnadiff [4] when analysing the reconstruction of DS43. Only one execution of HVRS is represented.

| Reconstruction tool | Total seqs | Aligned seqs | Aligned bases | Aligned bases (%) | Unaligned bases | Unaligned bases (%) | Average length | SNPs | Identity |
|---|---|---|---|---|---|---|---|---|---|
| coronaSPAdes | 6 | 4 | 143694 | 68.3505 | 66537 | 31.6495 | 35923.5000 | 2 | 99,991 |
| Haploflow | 44 | 30 | 141462 | 68.4009 | 65351 | 31.5991 | 4426.0312 | 616 | 99,537 |
| IRMA | 5 | 4 | 128264 | 88.5728 | 16548 | 11.4272 | 25654.4000 | 0 | 99,999 |
| LAZYPIPE | 8 | 6 | 143685 | 68.3765 | 66453 | 31.6235 | 20538.1429 | 6 | 99,981 |
| metaSPAdes | 6 | 4 | 143694 | 68.3505 | 66537 | 31.6495 | 35923.5000 | 2 | 99,998 |
| metaviralSPAdes | – | – | – | – | – | – | – | – | – |
| PEHaplo | 358 | 243 | 190191 | 69.6030 | 83060 | 30.3970 | 774.5275 | 145 | 99,839 |
| QuRe | 6 | 5 | 1596 | 91.4613 | 149 | 8.5387 | 319.2000 | 1 | 99,874 |
| QVG | 4 | 4 | 143858 | 99.9986 | 2 | 0.0014 | 35964.5000 | 812 | 99,432 |
| SPAdes | 6 | 4 | 143694 | 68.3505 | 66537 | 31.6495 | 35923.5000 | 0 | 100,000 |
| SSAKE | 374 | 244 | 77929 | 65.7773 | 40545 | 34.2227 | 321.4149 | 6 | 99,991 |
| TRACESPipe | 4 | 4 | 143811 | 99.9632 | 53 | 0.0368 | 35952.7500 | 741 | 99,481 |
| TRACESPipeLite | 6 | 4 | 114628 | 68.1239 | 53636 | 31.8761 | 1097.5192 | 359 | 97,533 |
| V-pipe | 5 | 4 | 143283 | 89.3124 | 17146 | 10.6876 | 35820.7500 | 693 | 99,348 |
| VirGenA | 68 | 68 | 53807 | 99.2255 | 420 | 0.7745 | 662.0506 | 93 | 99,788 |
| ViSpA | 5 | 4 | 143614 | 89.6870 | 16514 | 10.3130 | 35903.5000 | 801 | 99,410 |

Table S111: Results obtained by dnadiff [4] when analysing the reconstruction of DS44. Only one execution of HVRS is represented.

| Reconstruction tool | Total seqs | Aligned seqs | Aligned bases | Aligned bases (%) | Unaligned bases | Unaligned bases (%) | Average length | SNPs | Identity |
|---|---|---|---|---|---|---|---|---|---|
| coronaSPAdes | 6 | 4 | 143731 | 68.3779 | 66470 | 31.6221 | 35932.7500 | 0 | 100,000 |
| Haploflow | 61 | 42 | 141410 | 68.5505 | 64876 | 31.4495 | 3356.0476 | 69 | 99,933 |
| IRMA | 5 | 4 | 128494 | 88.6166 | 16506 | 11.3834 | 21416.0000 | 0 | 100,000 |
| LAZYPIPE | 6 | 4 | 143600 | 68.3731 | 66424 | 31.6269 | 35900.0000 | 0 | 99,983 |
| metaSPAdes | 6 | 4 | 143731 | 68.3779 | 66470 | 31.6221 | 35932.7500 | 0 | 100,000 |
| metaviralSPAdes | – | – | – | – | – | – | – | – | – |
| PEHaplo | 328 | 237 | 191181 | 71.0736 | 77809 | 28.9264 | 775.8610 | 44 | 99,949 |
| QuRe | 5 | 4 | 650 | 77.1971 | 192 | 22.8029 | 171.3333 | 0 | 99,806 |
| QVG | 4 | 4 | 143860 | 100.0000 | 0 | 0.0000 | 35965.0000 | 1293 | 99,056 |
| SPAdes | 6 | 4 | 143731 | 68.3779 | 66470 | 31.6221 | 35932.7500 | 0 | 100,000 |
| SSAKE | 348 | 236 | 76303 | 68.7198 | 34732 | 31.2802 | 323.3178 | 2 | 99,997 |
| TRACESPipe | 4 | 4 | 143771 | 99.9361 | 92 | 0.0639 | 35941.2500 | 893 | 99,294 |
| TRACESPipeLite | 6 | 4 | 37385 | 22.2183 | 130877 | 77.7817 | 330.1982 | 38 | 98,367 |
| V-pipe | 5 | 4 | 142897 | 89.0718 | 17532 | 10.9282 | 28579.4000 | 840 | 99,045 |
| VirGenA | 63 | 63 | 55574 | 99.0995 | 505 | 0.9005 | 742.2933 | 87 | 99,827 |
| ViSpA | 10 | 9 | 288411 | 94.5926 | 16487 | 5.4074 | 24618.3333 | 1069 | 99,064 |

Table S112: Results obtained by dnadiff [4] when analysing the reconstruction of DS45. Only one execution of HVRS is represented.

| Reconstruction tool | Total seqs | Aligned seqs | Aligned bases | Aligned bases (%) | Unaligned bases | Unaligned bases (%) | Average length | SNPs | Identity |
|---|---|---|---|---|---|---|---|---|---|
| coronaSPAdes | 6 | 4 | 143666 | 68.3668 | 66474 | 31.6332 | 35916.5000 | 1 | 99,991 |
| Haploflow | 45 | 35 | 140928 | 68.1052 | 65999 | 31.8948 | 4026.5143 | 0 | 100,000 |
| IRMA | 5 | 4 | 129184 | 88.6413 | 16554 | 11.3587 | 11744.2727 | 8 | 99,994 |
| LAZYPIPE | 6 | 4 | 143567 | 68.3581 | 66455 | 31.6419 | 35891.7500 | 0 | 100,000 |
| metaSPAdes | 6 | 4 | 143671 | 68.3669 | 66476 | 31.6331 | 35917.7500 | 1 | 99,991 |
| metaviralSPAdes | – | – | – | – | – | – | – | – | – |
| PEHaplo | 317 | 218 | 177593 | 69.5802 | 77642 | 30.4198 | 803.7321 | 39 | 99,935 |
| QuRe | 5 | 4 | 1545 | 91.7458 | 139 | 8.2542 | 386.2500 | 1 | 99,936 |
| QVG | 4 | 4 | 143812 | 99.9673 | 47 | 0.0327 | 35953.0000 | 1950 | 98,633 |
| SPAdes | 6 | 4 | 143671 | 68.3669 | 66476 | 31.6331 | 35917.7500 | 1 | 99,991 |
| SSAKE | 387 | 276 | 84168 | 69.6876 | 36611 | 30.3124 | 305.6667 | 2 | 99,994 |
| TRACESPipe | 4 | 4 | 143175 | 99.5328 | 672 | 0.4672 | 17896.8750 | 1516 | 98,594 |
| TRACESPipeLite | 6 | 3 | 6474 | 3.8476 | 161788 | 96.1524 | 239.7778 | 2 | 99,509 |
| V-pipe | 5 | 4 | 143231 | 89.2800 | 17198 | 10.7200 | 29026.6000 | 908 | 98,817 |
| VirGenA | – | – | – | – | – | – | – | – | – |
| ViSpA | 5 | 4 | 1485102 | 98.8976 | 16555 | 1.1024 | 12345.7500 | 1289 | 98,403 |

Table S113: Results obtained by dnadiff [4] when analysing the reconstruction of DS46. Only one execution of HVRS is represented.

| Reconstruction tool | Total seqs | Aligned seqs | Aligned bases | Aligned bases (%) | Unaligned bases | Unaligned bases (%) | Average length | SNPs | Identity |
|---|---|---|---|---|---|---|---|---|---|
| coronaSPAdes | 6 | 4 | 143695 | 68.3822 | 66440 | 31.6178 | 35923.7500 | 3 | 99,990 |
| Haploflow | 49 | 30 | 141422 | 68.4272 | 65253 | 31.5728 | 4714.0667 | 0 | 100,000 |
| IRMA | 5 | 4 | 129691 | 88.6564 | 16594 | 11.3436 | 9264.9286 | 3 | 99,989 |
| LAZYPIPE | 6 | 4 | 143602 | 68.3936 | 66362 | 31.6064 | 35900.5000 | 0 | 100,000 |
| metaSPAdes | 6 | 4 | 143695 | 68.3822 | 66440 | 31.6178 | 35923.7500 | 0 | 100,000 |
| metaviralSPAdes | – | – | – | – | – | – | – | – | – |
| PEHaplo | 299 | 201 | 167945 | 68.3837 | 77647 | 31.6163 | 836.8693 | 33 | 99,944 |
| QuRe | 8 | 7 | 1074 | 85.3058 | 185 | 14.6942 | 153.4286 | 0 | 100,000 |
| QVG | 4 | 4 | 143852 | 100.0000 | 0 | 0.0000 | 35963.0000 | 3754 | 97,368 |
| SPAdes | 6 | 4 | 143695 | 68.3822 | 66440 | 31.6178 | 35923.7500 | 0 | 100,000 |
| SSAKE | 377 | 260 | 83650 | 68.4013 | 38643 | 31.5987 | 322.9535 | 6 | 99,992 |
| TRACESPipe | 4 | 4 | 143660 | 99.8811 | 171 | 0.1189 | 35914.2500 | 239 | 99,809 |
| TRACESPipeLite | 6 | 1 | 227 | 0.1349 | 168035 | 99.8651 | 227.0000 | 0 | 100,000 |
| V-pipe | 5 | 4 | 140715 | 87.7117 | 19714 | 12.2883 | 7035.7500 | 1120 | 97,644 |
| VirGenA | 56 | 56 | 48588 | 99.4148 | 286 | 0.5852 | 675.3973 | 30 | 99,694 |
| ViSpA | 69 | 68 | 3250188 | 99.4894 | 16681 | 0.5106 | 3986.9714 | 1195 | 97,355 |

Table S114: Results obtained by dnadiff [4] when analysing the reconstruction of DS47. Only one execution of HVRS is represented.

| Reconstruction tool | Total seqs | Aligned seqs | Aligned bases | Aligned bases (%) | Unaligned bases | Unaligned bases (%) | Average length | SNPs | Identity |
|---|---|---|---|---|---|---|---|---|---|
| coronaSPAdes | 6 | 4 | 143784 | 68.3732 | 66509 | 31.6268 | 35946.0000 | 0 | 100,000 |
| Haploflow | 61 | 40 | 142664 | 68.5499 | 65453 | 31.4501 | 3566.6000 | 0 | 100,000 |
| IRMA | 5 | 4 | 129557 | 88.6618 | 16568 | 11.3382 | 7621.2941 | 13 | 99,988 |
| LAZYPIPE | 24 | 19 | 143592 | 68.3706 | 66428 | 31.6294 | 7557.4737 | 2 | 99,997 |
| metaSPAdes | 6 | 4 | 143784 | 68.3732 | 66509 | 31.6268 | 35946.0000 | 1 | 99,991 |
| metaviralSPAdes | – | – | – | – | – | – | – | – | – |
| PEHaplo | 310 | 207 | 170956 | 67.7628 | 81330 | 32.2372 | 824.4406 | 0 | 100,000 |
| QuRe | 4 | 4 | 1128 | 99.9114 | 1 | 0.0886 | 282.0000 | 4 | 99,648 |
| QVG | 4 | 4 | 143812 | 99.9708 | 42 | 0.0292 | 30786.6000 | 6950 | 94,724 |
| SPAdes | 6 | 4 | 143784 | 68.3732 | 66509 | 31.6268 | 35946.0000 | 1 | 99,991 |
| SSAKE | 357 | 236 | 78994 | 68.3830 | 36523 | 31.6170 | 335.6213 | 3 | 99,996 |
| TRACESPipe | 4 | 4 | 143701 | 100.0000 | 0 | 0.0000 | 35925.2500 | 3 | 99,991 |
| TRACESPipeLite | 5 | 0 | 0 | 0.0000 | 160429 | 100.0000 | 0.0000 | 0 | 0,000 |
| V-pipe | 5 | 4 | 131713 | 82.1005 | 28716 | 17.8995 | 1995.6515 | 1569 | 95,842 |
| VirGenA | 57 | 57 | 50620 | 98.4710 | 786 | 1.5290 | 708.3662 | 96 | 99,753 |
| ViSpA | 162 | 161 | 6551066 | 99.7176 | 18552 | 0.2824 | 1033.9009 | 1108 | 96,251 |

Table S115: Results obtained by dnadiff [4] when analysing the reconstruction of DS48. Only one execution of HVRS is represented.

| Reconstruction tool | Total seqs | Aligned seqs | Aligned bases | Aligned bases (%) | Unaligned bases | Unaligned bases (%) | Average length | SNPs | Identity |
|---|---|---|---|---|---|---|---|---|---|
| coronaSPAdes | 6 | 4 | 143723 | 68.3686 | 66495 | 31.6314 | 35930.7500 | 0 | 100,000 |
| Haploflow | 50 | 29 | 142509 | 68.4452 | 65700 | 31.5548 | 4914.1034 | 0 | 100,000 |
| IRMA | 5 | 4 | 130625 | 88.7302 | 16591 | 11.2698 | 5939.1818 | 55 | 99,958 |
| LAZYPIPE | 6 | 4 | 143565 | 68.3653 | 66432 | 31.6347 | 35891.2500 | 0 | 100,000 |
| metaSPAdes | 6 | 4 | 143723 | 68.3686 | 66495 | 31.6314 | 35930.7500 | 0 | 100,000 |
| metaviralSPAdes | – | – | – | – | – | – | – | – | – |
| PEHaplo | 305 | 207 | 166087 | 67.1878 | 81111 | 32.8122 | 806.0739 | 9 | 99,986 |
| QuRe | 7 | 6 | 2236 | 90.5630 | 233 | 9.4370 | 372.6667 | 7 | 99,639 |
| QVG | 4 | 4 | 143851 | 99.9937 | 9 | 0.0063 | 35962.7500 | 11569 | 91,836 |
| SPAdes | 6 | 4 | 143723 | 68.3686 | 66495 | 31.6314 | 35930.7500 | 0 | 100,000 |
| SSAKE | 360 | 249 | 78812 | 68.0047 | 37080 | 31.9953 | 318.0850 | 3 | 99,994 |
| TRACESPipe | 4 | 4 | 143770 | 99.9715 | 41 | 0.0285 | 35942.5000 | 52 | 99,851 |
| TRACESPipeLite | 5 | 0 | 0 | 0.0000 | 160429 | 100.0000 | 0.0000 | 0 | 0,000 |
| V-pipe | 5 | 4 | 114314 | 71.2552 | 46115 | 28.7448 | 905.5476 | 566 | 94,245 |
| VirGenA | 60 | 60 | 44245 | 98.9910 | 451 | 1.0090 | 638.4143 | 10 | 99,761 |
| ViSpA | 168 | 167 | 5712543 | 99.4543 | 31345 | 0.5457 | 500.4251 | 515 | 95,995 |

Table S116: Results obtained by dnadiff [4] when analysing the reconstruction of DS49. Only one execution of HVRS is represented.

| Reconstruction tool | Total seqs | Aligned seqs | Aligned bases | Aligned bases (%) | Unaligned bases | Unaligned bases (%) | Average length | SNPs | Identity |
|---|---|---|---|---|---|---|---|---|---|
| coronaSPAdes | 9 | 7 | 135981 | 67.1123 | 66636 | 32.8877 | 15117.0000 | 0 | 99,949 |
| Haploflow | 14 | 11 | 136947 | 67.3501 | 66389 | 32.6499 | 13556.2000 | 8 | 99,989 |
| IRMA | 5 | 4 | 128314 | 88.5767 | 16548 | 11.4233 | 25662.8000 | 0 | 100,000 |
| LAZYPIPE | 8 | 6 | 136021 | 67.1569 | 66521 | 32.8431 | 17016.6250 | 0 | 99,994 |
| metaSPAdes | 10 | 8 | 136068 | 67.1593 | 66537 | 32.8407 | 16994.7500 | 15 | 99,800 |
| metaviralSPAdes | 2 | 0 | 0 | 0.0000 | 66537 | 100.0000 | 0.0000 | 0 | 0,000 |
| PEHaplo | 215 | 140 | 210019 | 67.5185 | 101035 | 32.4815 | 1488.6016 | 0 | 99,999 |
| QuRe | 5 | 4 | 8273 | 85.4649 | 1407 | 14.5351 | 2068.2500 | 0 | 100,000 |
| QVG | – | – | – | – | – | – | – | – | – |
| SPAdes | 9 | 7 | 136187 | 67.1785 | 66537 | 32.8215 | 17030.7500 | 0 | 99,900 |
| SSAKE | 169 | 110 | 145530 | 67.1713 | 71125 | 32.8287 | 1871.2162 | 8 | 99,988 |
| TRACESPipe | 4 | 4 | 143829 | 99.9785 | 31 | 0.0215 | 35957.2500 | 0 | 100,000 |
| TRACESPipeLite | 6 | 5 | 144185 | 85.6908 | 24077 | 14.3092 | 35951.0000 | 0 | 100,000 |
| V-pipe | 5 | 4 | 143497 | 89.4458 | 16932 | 10.5542 | 35874.2500 | 0 | 100,000 |
| VirGenA | 61 | 61 | 52214 | 98.5226 | 783 | 1.4774 | 693.5205 | 8 | 99,925 |
| ViSpA | 5 | 4 | 143715 | 89.6795 | 16539 | 10.3205 | 35928.7500 | 0 | 100,000 |

Table S117: Results obtained by dnadiff [4] when analysing the reconstruction of DS50. Only one execution of HVRS is represented.

| Reconstruction tool | Total seqs | Aligned seqs | Aligned bases | Aligned bases (%) | Unaligned bases | Unaligned bases (%) | Average length | SNPs | Identity |
|---|---|---|---|---|---|---|---|---|---|
| coronaSPAdes | 12 | 10 | 141394 | 67.9968 | 66548 | 32.0032 | 14139.4000 | 6 | 99,996 |
| Haploflow | 11 | 9 | 143215 | 68.3111 | 66436 | 31.6889 | 14321.6000 | 315 | 99,704 |
| IRMA | 5 | 4 | 128351 | 88.5723 | 16560 | 11.4277 | 25670.4000 | 0 | 100,000 |
| LAZYPIPE | 8 | 6 | 143661 | 68.3540 | 66511 | 31.6460 | 19257.1250 | 1 | 99,963 |
| metaSPAdes | 6 | 4 | 143796 | 68.3623 | 66548 | 31.6377 | 35949.0000 | 78 | 99,947 |
| metaviralSPAdes | 3 | 1 | 125025 | 65.2623 | 66548 | 34.7377 | 125025.0000 | 0 | 100,000 |
| PEHaplo | 243 | 176 | 249458 | 72.3396 | 95385 | 27.6604 | 1397.3841 | 114 | 99,917 |
| QuRe | 4 | 3 | 4905 | 84.0617 | 930 | 15.9383 | 1635.0000 | 4 | 99,918 |
| QVG | 4 | 4 | 143860 | 100.0000 | 0 | 0.0000 | 35965.0000 | 498 | 99,577 |
| SPAdes | 6 | 4 | 143810 | 68.3644 | 66548 | 31.6356 | 35952.5000 | 1 | 99,999 |
| SSAKE | 164 | 116 | 151144 | 68.2720 | 70241 | 31.7280 | 1594.0769 | 104 | 99,924 |
| TRACESPipe | 4 | 4 | 143833 | 99.9805 | 28 | 0.0195 | 35958.2500 | 404 | 99,719 |
| TRACESPipeLite | 6 | 4 | 143280 | 85.1534 | 24981 | 14.8466 | 20468.5714 | 475 | 99,527 |
| V-pipe | 5 | 4 | 143544 | 89.4751 | 16885 | 10.5249 | 35886.0000 | 476 | 99,664 |
| VirGenA | 37 | 37 | 46433 | 98.9958 | 471 | 1.0042 | 842.7500 | 96 | 99,523 |
| ViSpA | 5 | 4 | 143725 | 89.6824 | 16535 | 10.3176 | 35931.2500 | 551 | 99,569 |

Table S118: Results obtained by dnadiff [4] when analysing the reconstruction of DS51. Only one execution of HVRS is represented.

| Reconstruction tool | Total seqs | Aligned seqs | Aligned bases | Aligned bases (%) | Unaligned bases | Unaligned bases (%) | Average length | SNPs | Identity |
|---|---|---|---|---|---|---|---|---|---|
| coronaSPAdes | 6 | 4 | 143813 | 68.3655 | 66546 | 31.6345 | 35953.2500 | 2 | 99,999 |
| Haploflow | 11 | 9 | 149790 | 69.2799 | 66420 | 30.7201 | 20506.2857 | 245 | 99,792 |
| IRMA | 5 | 4 | 128364 | 88.5690 | 16567 | 11.4310 | 25673.0000 | 0 | 100,000 |
| LAZYPIPE | 8 | 6 | 144054 | 68.4053 | 66535 | 31.5947 | 18936.7500 | 1 | 99,478 |
| metaSPAdes | 6 | 4 | 143813 | 68.3655 | 66546 | 31.6345 | 35953.2500 | 1 | 99,999 |
| metaviralSPAdes | 3 | 1 | 125029 | 65.2637 | 66546 | 34.7363 | 125029.0000 | 0 | 100,000 |
| PEHaplo | 242 | 159 | 223138 | 68.7082 | 101624 | 31.2918 | 1384.3066 | 0 | 99,944 |
| QuRe | 5 | 4 | 6906 | 77.9458 | 1954 | 22.0542 | 1561.6000 | 3 | 98,835 |
| QVG | 4 | 4 | 143858 | 99.9986 | 2 | 0.0014 | 35964.5000 | 741 | 99,488 |
| SPAdes | 6 | 4 | 143813 | 68.3655 | 66546 | 31.6345 | 35953.2500 | 1 | 99,999 |
| SSAKE | 172 | 108 | 151571 | 67.7982 | 71991 | 32.2018 | 1791.6707 | 1 | 99,998 |
| TRACESPipe | 4 | 4 | 143805 | 99.9604 | 57 | 0.0396 | 35951.2500 | 609 | 99,579 |
| TRACESPipeLite | 6 | 4 | 110125 | 70.4750 | 46136 | 29.5250 | 1280.5233 | 315 | 97,753 |
| V-pipe | 5 | 4 | 143518 | 89.4589 | 16911 | 10.5411 | 35879.5000 | 590 | 99,591 |
| VirGenA | 47 | 47 | 48429 | 99.4394 | 273 | 0.5606 | 792.5410 | 62 | 99,659 |
| ViSpA | 5 | 4 | 143735 | 89.6752 | 16549 | 10.3248 | 35933.7500 | 638 | 99,522 |

Table S119: Results obtained by dnadiff [4] when analysing the reconstruction of DS52. Only one execution of HVRS is represented.

| Reconstruction tool | Total seqs | Aligned seqs | Aligned bases | Aligned bases (%) | Unaligned bases | Unaligned bases (%) | Average length | SNPs | Identity |
|---|---|---|---|---|---|---|---|---|---|
| coronaSPAdes | 6 | 4 | 143790 | 68.3591 | 66555 | 31.6409 | 35947.5000 | 0 | 100,000 |
| Haploflow | 9 | 7 | 143379 | 68.3180 | 66491 | 31.6820 | 20162.7143 | 499 | 99,371 |
| IRMA | 5 | 4 | 128391 | 88.5724 | 16565 | 11.4276 | 25678.2000 | 0 | 100,000 |
| LAZYPIPE | 6 | 4 | 143689 | 68.3478 | 66543 | 31.6522 | 28766.4000 | 0 | 99,985 |
| metaSPAdes | 6 | 4 | 143791 | 68.3593 | 66555 | 31.6407 | 35947.7500 | 0 | 100,000 |
| metaviralSPAdes | 3 | 1 | 125021 | 65.2592 | 66555 | 34.7408 | 125021.0000 | 0 | 100,000 |
| PEHaplo | 217 | 155 | 211608 | 70.0320 | 90551 | 29.9680 | 1426.8676 | 18 | 99,983 |
| QuRe | 5 | 4 | 6961 | 81.6444 | 1565 | 18.3556 | 1740.2500 | 4 | 99,942 |
| QVG | 4 | 4 | 143860 | 100.0000 | 0 | 0.0000 | 35965.0000 | 1058 | 99,221 |
| SPAdes | 6 | 4 | 143791 | 68.3593 | 66555 | 31.6407 | 35947.7500 | 0 | 100,000 |
| SSAKE | 147 | 101 | 150862 | 68.2769 | 70094 | 31.7231 | 1793.2073 | 0 | 99,999 |
| TRACESPipe | 4 | 4 | 143804 | 99.9611 | 56 | 0.0389 | 35951.0000 | 6 | 99,986 |
| TRACESPipeLite | 6 | 4 | 43893 | 26.0859 | 124370 | 73.9141 | 359.3025 | 103 | 98,240 |
| V-pipe | 5 | 4 | 143508 | 89.4527 | 16921 | 10.5473 | 30718.4000 | 556 | 99,201 |
| VirGenA | 46 | 46 | 42874 | 99.5704 | 185 | 0.4296 | 661.7833 | 55 | 99,801 |
| ViSpA | 11 | 10 | 475775 | 96.6360 | 16562 | 3.3640 | 35918.5000 | 1028 | 99,078 |

Table S120: Results obtained by dnadiff [4] when analysing the reconstruction of DS53. Only one execution of HVRS is represented.

| Reconstruction tool | Total seqs | Aligned seqs | Aligned bases | Aligned bases (%) | Unaligned bases | Unaligned bases (%) | Average length | SNPs | Identity |
|---|---|---|---|---|---|---|---|---|---|
| coronaSPAdes | 6 | 4 | 143766 | 68.3571 | 66550 | 31.6429 | 35941.5000 | 1 | 99,999 |
| Haploflow | 8 | 6 | 144613 | 68.5253 | 66423 | 31.4747 | 24101.1667 | 0 | 99,862 |
| IRMA | 5 | 4 | 129518 | 88.6606 | 16565 | 11.3394 | 9974.5385 | 7 | 99,976 |
| LAZYPIPE | 6 | 4 | 143712 | 68.3575 | 66524 | 31.6425 | 35928.0000 | 1 | 99,999 |
| metaSPAdes | 6 | 4 | 143767 | 68.3573 | 66550 | 31.6427 | 35941.7500 | 1 | 99,999 |
| metaviralSPAdes | 3 | 1 | 125016 | 65.2600 | 66550 | 34.7400 | 125016.0000 | 0 | 100,000 |
| PEHaplo | 230 | 155 | 215544 | 68.8808 | 97379 | 31.1192 | 1403.2101 | 148 | 99,874 |
| QuRe | 4 | 3 | 1652 | 53.3247 | 1446 | 46.6753 | 550.6667 | 2 | 99,878 |
| QVG | 4 | 4 | 143813 | 99.9673 | 47 | 0.0327 | 35953.2500 | 1495 | 98,959 |
| SPAdes | 6 | 4 | 143767 | 68.3573 | 66550 | 31.6427 | 35941.7500 | 1 | 99,999 |
| SSAKE | 179 | 112 | 151367 | 67.8481 | 71730 | 32.1519 | 1724.0588 | 0 | 99,997 |
| TRACESPipe | 4 | 4 | 143806 | 99.9861 | 20 | 0.0139 | 35951.5000 | 1 | 99,999 |
| TRACESPipeLite | 6 | 3 | 8049 | 4.7836 | 160213 | 95.2164 | 236.5758 | 0 | 99,658 |
| V-pipe | 5 | 4 | 143508 | 89.4527 | 16921 | 10.5473 | 35875.2500 | 1157 | 99,096 |
| VirGenA | – | – | – | – | – | – | – | – | – |
| ViSpA | 5 | 4 | 1054129 | 98.4540 | 16553 | 1.5460 | 19976.6667 | 1293 | 98,564 |

Table S121: Results obtained by dnadiff [4] when analysing the reconstruction of DS54. Only one execution of HVRS is represented.

| Reconstruction tool | Total seqs | Aligned seqs | Aligned bases | Aligned bases (%) | Unaligned bases | Unaligned bases (%) | Average length | SNPs | Identity |
|---|---|---|---|---|---|---|---|---|---|
| coronaSPAdes | 6 | 4 | 143803 | 68.3617 | 66553 | 31.6383 | 35950.7500 | 0 | 100,000 |
| Haploflow | 7 | 5 | 143470 | 68.3773 | 66351 | 31.6227 | 28694.0000 | 0 | 100,000 |
| IRMA | 5 | 4 | 129886 | 88.6903 | 16563 | 11.3097 | 8660.5333 | 6 | 99,993 |
| LAZYPIPE | 6 | 4 | 143771 | 68.3667 | 66523 | 31.6333 | 35942.7500 | 0 | 100,000 |
| metaSPAdes | 6 | 4 | 143807 | 68.3623 | 66553 | 31.6377 | 35951.7500 | 0 | 100,000 |
| metaviralSPAdes | 3 | 1 | 125037 | 65.2628 | 66553 | 34.7372 | 125037.0000 | 0 | 100,000 |
| PEHaplo | 236 | 174 | 227715 | 71.7143 | 89816 | 28.2857 | 1330.0724 | 14 | 99,990 |
| QuRe | 6 | 5 | 14299 | 90.0838 | 1574 | 9.9162 | 2850.4000 | 9 | 96,671 |
| QVG | 4 | 4 | 143862 | 100.0000 | 0 | 0.0000 | 35965.5000 | 2719 | 98,103 |
| SPAdes | 6 | 4 | 143807 | 68.3623 | 66553 | 31.6377 | 35951.7500 | 0 | 100,000 |
| SSAKE | 171 | 113 | 151255 | 68.0070 | 71156 | 31.9930 | 1585.4516 | 0 | 99,998 |
| TRACESPipe | 4 | 4 | 143818 | 99.9819 | 26 | 0.0181 | 35954.5000 | 0 | 100,000 |
| TRACESPipeLite | 6 | 2 | 746 | 0.4434 | 167516 | 99.5566 | 248.6667 | 0 | 100,000 |
| V-pipe | 5 | 4 | 142742 | 88.9752 | 17687 | 11.0248 | 15859.4444 | 1248 | 98,469 |
| VirGenA | 47 | 47 | 40467 | 99.8372 | 66 | 0.1628 | 638.5938 | 106 | 99,366 |
| ViSpA | 57 | 56 | 2591210 | 99.3631 | 16610 | 0.6369 | 4487.0000 | 723 | 98,415 |

Table S122: Results obtained by dnadiff [4] when analysing the reconstruction of DS55. Only one execution of HVRS is represented.

| Reconstruction tool | Total seqs | Aligned seqs | Aligned bases | Aligned bases (%) | Unaligned bases | Unaligned bases (%) | Average length | SNPs | Identity |
|---|---|---|---|---|---|---|---|---|---|
| coronaSPAdes | 6 | 4 | 143804 | 68.3593 | 66561 | 31.6407 | 35951.0000 | 1 | 99,991 |
| Haploflow | 6 | 4 | 143453 | 68.3487 | 66431 | 31.6513 | 35863.2500 | 0 | 100,000 |
| IRMA | 5 | 4 | 129570 | 88.6658 | 16563 | 11.3342 | 8638.4667 | 32 | 99,973 |
| LAZYPIPE | 6 | 4 | 143728 | 68.3592 | 66526 | 31.6408 | 35932.0000 | 0 | 100,000 |
| metaSPAdes | 6 | 4 | 143804 | 68.3593 | 66561 | 31.6407 | 35951.0000 | 0 | 100,000 |
| metaviralSPAdes | 3 | 1 | 125038 | 65.2603 | 66561 | 34.7397 | 125038.0000 | 0 | 100,000 |
| PEHaplo | 231 | 155 | 211923 | 68.1081 | 99234 | 31.8919 | 1394.5214 | 1 | 99,999 |
| QuRe | 5 | 4 | 3165 | 76.8204 | 955 | 23.1796 | 791.2500 | 7 | 99,780 |
| QVG | 4 | 4 | 143818 | 99.9708 | 42 | 0.0292 | 30787.8000 | 5078 | 96,251 |
| SPAdes | 6 | 4 | 143804 | 68.3593 | 66561 | 31.6407 | 35951.0000 | 0 | 100,000 |
| SSAKE | 169 | 111 | 151654 | 68.4257 | 69979 | 31.5743 | 1768.2410 | 1 | 99,998 |
| TRACESPipe | 4 | 4 | 143780 | 100.0000 | 0 | 0.0000 | 35945.0000 | 1 | 99,999 |
| TRACESPipeLite | 5 | 0 | 0 | 0.0000 | 160429 | 100.0000 | 0.0000 | 0 | 0,000 |
| V-pipe | 5 | 4 | 137879 | 85.9439 | 22550 | 14.0561 | 4055.2647 | 1529 | 97,021 |
| VirGenA | 50 | 49 | 47872 | 99.6337 | 176 | 0.3663 | 749.9844 | 47 | 99,869 |
| ViSpA | 147 | 146 | 6162466 | 99.7015 | 18451 | 0.2985 | 1315.8614 | 886 | 96,130 |

Table S123: Results obtained by dnadiff [4] when analysing the reconstruction of DS56. Only one execution of HVRS is represented.

| Reconstruction tool | Total seqs | Aligned seqs | Aligned bases | Aligned bases (%) | Unaligned bases | Unaligned bases (%) | Average length | SNPs | Identity |
|---|---|---|---|---|---|---|---|---|---|
| coronaSPAdes | 6 | 4 | 143807 | 68.3666 | 66540 | 31.6334 | 35951.7500 | 0 | 100,000 |
| Haploflow | 6 | 4 | 143521 | 68.3958 | 66318 | 31.6042 | 35880.2500 | 0 | 100,000 |
| IRMA | 5 | 4 | 131193 | 88.6997 | 16714 | 11.3003 | 8200.5625 | 111 | 99,845 |
| LAZYPIPE | 6 | 4 | 143754 | 68.3631 | 66526 | 31.6369 | 35938.5000 | 0 | 100,000 |
| metaSPAdes | 6 | 4 | 143807 | 68.3666 | 66540 | 31.6334 | 35951.7500 | 0 | 100,000 |
| metaviralSPAdes | 3 | 1 | 125013 | 65.2629 | 66540 | 34.7371 | 125013.0000 | 0 | 100,000 |
| PEHaplo | 232 | 160 | 214186 | 67.0408 | 105300 | 32.9592 | 1313.4211 | 0 | 99,982 |
| QuRe | 4 | 3 | 4482 | 86.2255 | 716 | 13.7745 | 1494.0000 | 13 | 99,710 |
| QVG | 4 | 4 | 143855 | 99.9937 | 9 | 0.0063 | 35963.7500 | 9454 | 93,317 |
| SPAdes | 6 | 4 | 143807 | 68.3666 | 66540 | 31.6334 | 35951.7500 | 0 | 100,000 |
| SSAKE | 189 | 130 | 152803 | 68.5086 | 70239 | 31.4914 | 1499.6429 | 0 | 99,999 |
| TRACESPipe | 4 | 4 | 143754 | 100.0000 | 0 | 0.0000 | 35938.5000 | 0 | 100,000 |
| TRACESPipeLite | 5 | 0 | 0 | 0.0000 | 160429 | 100.0000 | 0.0000 | 0 | 0,000 |
| V-pipe | 5 | 4 | 126236 | 78.6865 | 34193 | 21.3135 | 1491.8675 | 1088 | 94,938 |
| VirGenA | 46 | 46 | 47210 | 98.8050 | 571 | 1.1950 | 728.9692 | 41 | 99,859 |
| ViSpA | 200 | 199 | 6867933 | 99.7126 | 19795 | 0.2874 | 597.7722 | 670 | 96,098 |

Table S124: Results obtained by dnadiff [4] when analysing the reconstruction of DS57. Only one execution of HVRS is represented.

| Reconstruction tool | Total seqs | Aligned seqs | Aligned bases | Aligned bases (%) | Unaligned bases | Unaligned bases (%) | Average length | SNPs | Identity |
|---|---|---|---|---|---|---|---|---|---|
| coronaSPAdes | 8 | 7 | 137147 | 73.2721 | 50028 | 26.7279 | 22823.8333 | 67 | 99,901 |
| Haploflow | 50 | 35 | 146800 | 74.7827 | 49502 | 25.2173 | 4281.9688 | 97 | 99,926 |
| IRMA | 4 | 4 | 128277 | 100.0000 | 0 | 0.0000 | 25655.4000 | 0 | 100,000 |
| LAZYPIPE | 8 | 7 | 136744 | 73.2799 | 49861 | 26.7201 | 14206.8000 | 16 | 99,874 |
| metaSPAdes | 11 | 10 | 136810 | 73.2239 | 50028 | 26.7761 | 11392.5000 | 54 | 99,867 |
| metaviralSPAdes | – | – | – | – | – | – | – | – | – |
| PEHaplo | 291 | 220 | 176357 | 74.9817 | 58843 | 25.0183 | 783.8852 | 56 | 99,940 |
| QuRe | 7 | 7 | 4065 | 99.9754 | 1 | 0.0246 | 459.6000 | 9 | 99,609 |
| QVG | 4 | 4 | 143858 | 99.9986 | 2 | 0.0014 | 35964.5000 | 160 | 99,893 |
| SPAdes | 22 | 21 | 145317 | 74.3895 | 50029 | 25.6105 | 6667.7143 | 38 | 99,928 |
| SSAKE | 371 | 269 | 82929 | 72.1969 | 31936 | 27.8031 | 308.9476 | 10 | 99,983 |
| TRACESPipe | 4 | 4 | 143717 | 99.8999 | 144 | 0.1001 | 35929.2500 | 131 | 99,910 |
| TRACESPipeLite | 5 | 5 | 143976 | 94.9128 | 7717 | 5.0872 | 35945.0000 | 178 | 99,876 |
| V-pipe | 5 | 4 | 143243 | 89.2875 | 17186 | 10.7125 | 28648.6000 | 141 | 99,684 |
| VirGenA | 72 | 72 | 69940 | 99.3748 | 440 | 0.6252 | 802.0714 | 32 | 99,901 |
| ViSpA | 5 | 4 | 143673 | 100.0000 | 0 | 0.0000 | 35918.2500 | 168 | 99,883 |

Table S125: Results obtained by dnadiff [4] when analysing the reconstruction of DS58. Only one execution of HVRS is represented.

| Reconstruction tool | Total seqs | Aligned seqs | Aligned bases | Aligned bases (%) | Unaligned bases | Unaligned bases (%) | Average length | SNPs | Identity |
|---|---|---|---|---|---|---|---|---|---|
| coronaSPAdes | 10 | 9 | 137255 | 89.1741 | 16663 | 10.8259 | 15237.3333 | 62 | 99,908 |
| Haploflow | 40 | 37 | 144560 | 89.9928 | 16075 | 10.0072 | 3653.6486 | 94 | 99,914 |
| IRMA | 5 | 4 | 128346 | 88.5695 | 16564 | 11.4305 | 25669.2000 | 0 | 100,000 |
| LAZYPIPE | 7 | 6 | 136300 | 89.1911 | 16518 | 10.8089 | 19452.5714 | 72 | 99,943 |
| metaSPAdes | 11 | 10 | 136859 | 89.1450 | 16665 | 10.8550 | 9971.2143 | 33 | 99,840 |
| metaviralSPAdes | – | – | – | – | – | – | – | – | – |
| PEHaplo | 224 | 197 | 167559 | 89.6133 | 19421 | 10.3867 | 835.2775 | 58 | 99,958 |
| QuRe | 7 | 5 | 1620 | 90.3010 | 174 | 9.6990 | 307.7500 | 3 | 99,755 |
| QVG | 4 | 4 | 143858 | 99.9986 | 2 | 0.0014 | 35964.5000 | 168 | 99,881 |
| SPAdes | 22 | 21 | 145142 | 89.7551 | 16567 | 10.2449 | 6670.3810 | 50 | 99,896 |
| SSAKE | 296 | 256 | 77442 | 86.9012 | 11673 | 13.0988 | 301.9881 | 16 | 99,970 |
| TRACESPipe | 4 | 4 | 143776 | 99.9416 | 84 | 0.0584 | 35944.0000 | 174 | 99,877 |
| TRACESPipeLite | 6 | 4 | 143790 | 85.4560 | 24472 | 14.5440 | 35947.5000 | 139 | 99,903 |
| V-pipe | 5 | 4 | 143297 | 89.3211 | 17132 | 10.6789 | 35824.2500 | 148 | 99,740 |
| VirGenA | 70 | 70 | 105575 | 99.9650 | 37 | 0.0350 | 1389.0400 | 105 | 99,876 |
| ViSpA | 5 | 4 | 143614 | 89.6842 | 16519 | 10.3158 | 35903.5000 | 188 | 99,867 |

Table S126: Results obtained by dnadiff [4] when analysing the reconstruction of DS59. Only one execution of HVRS is represented.

| Reconstruction tool | Total seqs | Aligned seqs | Aligned bases | Aligned bases (%) | Unaligned bases | Unaligned bases (%) | Average length | SNPs | Identity |
|---|---|---|---|---|---|---|---|---|---|
| coronaSPAdes | 12 | 10 | 141384 | 68.0005 | 66532 | 31.9995 | 14138.4000 | 46 | 99,967 |
| Haploflow | 16 | 13 | 145293 | 68.6316 | 66407 | 31.3684 | 11753.8333 | 457 | 99,660 |
| IRMA | 5 | 4 | 128326 | 88.5777 | 16548 | 11.4223 | 25665.2000 | 1 | 99,999 |
| LAZYPIPE | 8 | 6 | 143705 | 68.3691 | 66485 | 31.6309 | 19259.2500 | 1 | 99,963 |
| metaSPAdes | 7 | 5 | 143550 | 68.3305 | 66532 | 31.6695 | 35805.5000 | 76 | 99,946 |
| metaviralSPAdes | – | – | – | – | – | – | – | – | – |
| PEHaplo | 301 | 215 | 227014 | 72.2284 | 87286 | 27.7716 | 1012.9946 | 132 | 99,886 |
| QuRe | 4 | 3 | 4244 | 95.1143 | 218 | 4.8857 | 1414.6667 | 6 | 99,860 |
| QVG | 4 | 4 | 143860 | 100.0000 | 0 | 0.0000 | 35965.0000 | 503 | 99,573 |
| SPAdes | 6 | 4 | 143800 | 68.3681 | 66532 | 31.6319 | 35950.0000 | 2 | 99,998 |
| SSAKE | 408 | 275 | 150044 | 68.2881 | 69678 | 31.7119 | 558.7424 | 0 | 99,995 |
| TRACESPipe | 4 | 4 | 143806 | 99.9632 | 53 | 0.0368 | 35951.5000 | 498 | 99,653 |
| TRACESPipeLite | 6 | 4 | 143272 | 85.1482 | 24990 | 14.8518 | 20467.4286 | 452 | 99,519 |
| V-pipe | 5 | 4 | 143417 | 89.3959 | 17012 | 10.6041 | 35854.2500 | 454 | 99,683 |
| VirGenA | 53 | 53 | 46333 | 99.7589 | 112 | 0.2411 | 722.7500 | 96 | 99,713 |
| ViSpA | 5 | 4 | 143668 | 89.6882 | 16518 | 10.3118 | 35917.0000 | 610 | 99,564 |

Table S127: Results obtained by dnadiff [4] when analysing the reconstruction of DS60. Only one execution of HVRS is represented.

| Reconstruction tool | Total seqs | Aligned seqs | Aligned bases | Aligned bases (%) | Unaligned bases | Unaligned bases (%) | Average length | SNPs | Identity |
|---|---|---|---|---|---|---|---|---|---|
| coronaSPAdes | 6 | 4 | 143807 | 68.3601 | 66560 | 31.6399 | 35951.7500 | 0 | 100,000 |
| Haploflow | 8 | 6 | 145254 | 68.6067 | 66466 | 31.3933 | 20520.2857 | 206 | 99,844 |
| IRMA | 5 | 4 | 128358 | 88.5698 | 16565 | 11.4302 | 25672.4000 | 0 | 100,000 |
| LAZYPIPE | 7 | 5 | 143750 | 68.3590 | 66537 | 31.6410 | 20574.4286 | 0 | 99,976 |
| metaSPAdes | 6 | 4 | 143807 | 68.3601 | 66560 | 31.6399 | 35951.7500 | 0 | 100,000 |
| metaviralSPAdes | 3 | 1 | 125037 | 65.2604 | 66560 | 34.7396 | 125037.0000 | 0 | 100,000 |
| PEHaplo | 218 | 150 | 238466 | 71.3406 | 95798 | 28.6594 | 1562.7647 | 146 | 99,894 |
| QuRe | 4 | 4 | 4362 | 100.0000 | 0 | 0.0000 | 1090.5000 | 5 | 99,888 |
| QVG | 4 | 4 | 143858 | 99.9986 | 2 | 0.0014 | 35964.5000 | 736 | 99,488 |
| SPAdes | 6 | 4 | 143807 | 68.3601 | 66560 | 31.6399 | 35951.7500 | 0 | 100,000 |
| SSAKE | 171 | 116 | 152862 | 68.4041 | 70607 | 31.5959 | 1784.5000 | 0 | 99,999 |
| TRACESPipe | 4 | 4 | 143816 | 99.9646 | 51 | 0.0354 | 35954.0000 | 689 | 99,510 |
| TRACESPipeLite | 6 | 4 | 121031 | 71.9297 | 47232 | 28.0703 | 1384.5955 | 52 | 97,436 |
| V-pipe | 5 | 4 | 143557 | 89.4832 | 16872 | 10.5168 | 35889.2500 | 586 | 99,590 |
| VirGenA | – | – | – | – | – | – | – | – | – |
| ViSpA | 5 | 4 | 143799 | 89.6726 | 16561 | 10.3274 | 35949.7500 | 784 | 99,413 |

Table S128: Results obtained by dnadiff [4] when analysing the reconstruction of DS61. Only one execution of HVRS is represented.

| Reconstruction tool | Total seqs | Aligned seqs | Aligned bases | Aligned bases (%) | Unaligned bases | Unaligned bases (%) | Average length | SNPs | Identity |
|---|---|---|---|---|---|---|---|---|---|
| coronaSPAdes | 18 | 12 | 136059 | 67.0707 | 66800 | 32.9293 | 10474.0000 | 219 | 99,732 |
| Haploflow | – | – | – | – | – | – | – | – | – |
| IRMA | – | – | – | – | – | – | – | – | – |
| LAZYPIPE | 236 | 157 | 92896 | 66.4530 | 46896 | 33.5470 | 591.6943 | 504 | 99,384 |
| metaSPAdes | 72 | 56 | 137235 | 67.2141 | 66941 | 32.7859 | 3227.3333 | 266 | 99,773 |
| metaviralSPAdes | – | – | – | – | – | – | – | – | – |
| PEHaplo | 273 | 193 | 112848 | 70.4983 | 47224 | 29.5017 | 569.6701 | 51 | 99,931 |
| QuRe | 6 | 2 | 248 | 56.8807 | 188 | 43.1193 | 124.0000 | 0 | 100,000 |
| QVG | 4 | 4 | 143858 | 99.9986 | 2 | 0.0014 | 35964.5000 | 1203 | 99,166 |
| SPAdes | 17 | 15 | 141354 | 67.9995 | 66521 | 32.0005 | 9095.6667 | 133 | 99,844 |
| SSAKE | – | – | – | – | – | – | – | – | – |
| TRACESPipe | 4 | 4 | 143792 | 99.9527 | 68 | 0.0473 | 35948.0000 | 153 | 99,896 |
| TRACESPipeLite | 5 | 1 | 1610 | 1.0027 | 158950 | 98.9973 | 225.5714 | 12 | 98,227 |
| V-pipe | 5 | 0 | 0 | 0.0000 | 160429 | 100.0000 | 0.0000 | 0 | 0,000 |
| VirGenA | – | – | – | – | – | – | – | – | – |
| ViSpA | 5 | 4 | 415333 | 96.1695 | 16543 | 3.8305 | 22690.2857 | 70 | 99,569 |

Table S129: Results obtained by dnadiff [4] when analysing the reconstruction of DS62. Only one execution of HVRS is represented.

| Reconstruction tool | Total seqs | Aligned seqs | Aligned bases | Aligned bases (%) | Unaligned bases | Unaligned bases (%) | Average length | SNPs | Identity |
|---|---|---|---|---|---|---|---|---|---|
| coronaSPAdes | 12 | 10 | 138811 | 67.6071 | 66509 | 32.3929 | 15284.5556 | 35 | 99,969 |
| Haploflow | 111 | 77 | 123729 | 68.4921 | 56918 | 31.5079 | 1605.8919 | 65 | 99,910 |
| IRMA | 5 | 4 | 128289 | 88.5729 | 16551 | 11.4271 | 25657.8000 | 1 | 99,999 |
| LAZYPIPE | 9 | 7 | 138145 | 67.5661 | 66314 | 32.4339 | 17216.7500 | 56 | 99,901 |
| metaSPAdes | 14 | 12 | 136909 | 67.2719 | 66607 | 32.7281 | 11391.5833 | 50 | 99,855 |
| metaviralSPAdes | 4 | 2 | 117242 | 63.8392 | 66410 | 36.1608 | 58621.0000 | 56 | 99,953 |
| PEHaplo | 214 | 142 | 186147 | 68.6418 | 85039 | 31.3582 | 1276.3359 | 49 | 99,956 |
| QuRe | 2 | 2 | 440 | 100.0000 | 0 | 0.0000 | 220.0000 | 0 | 100,000 |
| QVG | 4 | 4 | 143858 | 99.9986 | 2 | 0.0014 | 35964.5000 | 151 | 99,891 |
| SPAdes | 15 | 13 | 142834 | 68.2623 | 66409 | 31.7377 | 11803.7500 | 33 | 99,980 |
| SSAKE | 105 | 69 | 13717 | 64.6876 | 7488 | 35.3124 | 198.7971 | 1 | 99,993 |
| TRACESPipe | 4 | 4 | 143731 | 99.9103 | 129 | 0.0897 | 35932.7500 | 154 | 99,894 |
| TRACESPipeLite | 6 | 4 | 143750 | 85.4322 | 24512 | 14.5678 | 35937.5000 | 113 | 99,922 |
| V-pipe | 5 | 4 | 142702 | 88.9503 | 17727 | 11.0497 | 35675.5000 | 134 | 99,715 |
| VirGenA | – | – | – | – | – | – | – | – | – |
| ViSpA | 5 | 4 | 143636 | 89.7052 | 16484 | 10.2948 | 35909.0000 | 256 | 99,815 |

Table S130: Results obtained by dnadiff [4] when analysing the reconstruction of DS63. Only one execution of HVRS is represented.

| Reconstruction tool | Total seqs | Aligned seqs | Aligned bases | Aligned bases (%) | Unaligned bases | Unaligned bases (%) | Average length | SNPs | Identity |
|---|---|---|---|---|---|---|---|---|---|
| coronaSPAdes | 17 | 15 | 291456 | 81.3818 | 66678 | 18.6182 | 22351.3846 | 126 | 99,957 |
| Haploflow | 89 | 71 | 307366 | 82.3543 | 65858 | 17.6457 | 4463.4030 | 217 | 99,888 |
| IRMA | 6 | 5 | 282079 | 94.4722 | 16505 | 5.5278 | 35306.0000 | 96 | 99,944 |
| LAZYPIPE | 14 | 12 | 290101 | 81.3604 | 66462 | 18.6396 | 20026.4667 | 152 | 99,854 |
| metaSPAdes | 29 | 27 | 291139 | 81.2976 | 66976 | 18.7024 | 12105.9167 | 113 | 99,868 |
| metaviralSPAdes | – | – | – | – | – | – | – | – | – |
| PEHaplo | 535 | 433 | 399761 | 82.1340 | 86957 | 17.8660 | 928.0124 | 242 | 99,895 |
| QuRe | 14 | 11 | 3434 | 88.5965 | 442 | 11.4035 | 238.2222 | 7 | 99,446 |
| QVG | 5 | 5 | 305972 | 99.9993 | 2 | 0.0007 | 61194.4000 | 341 | 99,881 |
| SPAdes | 53 | 51 | 303689 | 81.9972 | 66676 | 18.0028 | 6193.8542 | 54 | 99,939 |
| SSAKE | 417 | 346 | 111014 | 82.5727 | 23430 | 17.4273 | 322.5982 | 18 | 99,980 |
| TRACESPipe | 7 | 7 | 608310 | 97.3604 | 16492 | 2.6396 | 61077.2000 | 336 | 99,890 |
| TRACESPipeLite | 7 | 6 | 306116 | 92.6569 | 24260 | 7.3431 | 61183.8000 | 303 | 99,893 |
| V-pipe | 6 | 5 | 305271 | 94.6451 | 17272 | 5.3549 | 61050.2000 | 321 | 99,756 |
| VirGenA | 83 | 82 | 78143 | 98.7739 | 970 | 1.2261 | 703.3364 | 56 | 99,861 |
| ViSpA | 12 | 11 | 731237 | 97.7943 | 16493 | 2.2057 | 49566.8333 | 275 | 99,871 |

Table S131: Results obtained by dnadiff [4] when analysing the reconstruction of DS64. Only one execution of HVRS is represented.

| Reconstruction tool | Total seqs | Aligned seqs | Aligned bases | Aligned bases (%) | Unaligned bases | Unaligned bases (%) | Average length | SNPs | Identity |
|---|---|---|---|---|---|---|---|---|---|
| coronaSPAdes | 22 | 20 | 285059 | 81.0411 | 66687 | 18.9589 | 15512.4211 | 81 | 99,821 |
| Haploflow | 89 | 72 | 302847 | 82.1062 | 66001 | 17.8938 | 3948.0694 | 203 | 99,897 |
| IRMA | 6 | 5 | 281319 | 94.4423 | 16555 | 5.5577 | 11491.0800 | 44 | 99,726 |
| LAZYPIPE | 32 | 30 | 288596 | 81.2990 | 66385 | 18.7010 | 9284.5161 | 131 | 99,929 |
| metaSPAdes | 17 | 15 | 283200 | 80.8155 | 67228 | 19.1845 | 11828.8750 | 92 | 99,893 |
| metaviralSPAdes | 1 | 1 | 5358 | 100.0000 | 0 | 0.0000 | 5358.0000 | 2 | 99,960 |
| PEHaplo | 524 | 407 | 383606 | 81.2220 | 88687 | 18.7780 | 945.7447 | 142 | 99,952 |
| QuRe | 11 | 10 | 2906 | 93.5608 | 200 | 6.4392 | 284.0000 | 10 | 99,559 |
| QVG | 5 | 5 | 316189 | 100.0000 | 0 | 0.0000 | 54330.5000 | 432 | 99,857 |
| SPAdes | 62 | 60 | 298126 | 81.7648 | 66488 | 18.2352 | 5423.3148 | 48 | 99,846 |
| SSAKE | 376 | 313 | 108791 | 85.6096 | 18287 | 14.3904 | 351.1732 | 28 | 99,971 |
| TRACESPipe | 6 | 6 | 321012 | 99.9648 | 113 | 0.0352 | 52947.0000 | 398 | 99,866 |
| TRACESPipeLite | 7 | 6 | 316226 | 92.8463 | 24365 | 7.1537 | 52938.1667 | 395 | 99,867 |
| V-pipe | 6 | 5 | 315365 | 94.7731 | 17393 | 5.2269 | 52560.8333 | 434 | 99,686 |
| VirGenA | 73 | 73 | 81237 | 98.5611 | 1186 | 1.4389 | 709.0783 | 61 | 99,875 |
| ViSpA | 13 | 12 | 835387 | 98.0614 | 16515 | 1.9386 | 29216.8000 | 235 | 99,894 |

Table S132: Results obtained by dnadiff [4] when analysing the reconstruction of DS65. Only one execution of HVRS is represented.

| Reconstruction tool | Total seqs | Aligned seqs | Aligned bases | Aligned bases (%) | Unaligned bases | Unaligned bases (%) | Average length | SNPs | Identity |
|---|---|---|---|---|---|---|---|---|---|
| coronaSPAdes | 16 | 14 | 369164 | 84.7149 | 66608 | 15.2851 | 33496.1818 | 61 | 99,963 |
| Haploflow | 113 | 96 | 382763 | 85.2680 | 66131 | 14.7320 | 3963.2553 | 63 | 99,958 |
| IRMA | 6 | 5 | 356682 | 95.5683 | 16540 | 4.4317 | 50954.7143 | 0 | 100,000 |
| LAZYPIPE | 10 | 8 | 368598 | 84.7303 | 66427 | 15.2697 | 33603.4545 | 70 | 99,968 |
| metaSPAdes | 15 | 13 | 368348 | 84.6861 | 66609 | 15.3139 | 21879.9412 | 40 | 99,919 |
| metaviralSPAdes | – | – | – | – | – | – | – | – | – |
| PEHaplo | 592 | 510 | 502042 | 85.3827 | 85948 | 14.6173 | 997.5705 | 86 | 99,974 |
| QuRe | 11 | 7 | 1333 | 75.9544 | 422 | 24.0456 | 196.3333 | 2 | 99,746 |
| QVG | 5 | 5 | 379187 | 99.9995 | 2 | 0.0005 | 75837.4000 | 224 | 99,941 |
| SPAdes | 34 | 32 | 378544 | 85.0552 | 66513 | 14.9448 | 12046.7097 | 33 | 99,970 |
| SSAKE | 417 | 349 | 117901 | 84.2156 | 22098 | 15.7844 | 339.9246 | 12 | 99,990 |
| TRACESPipe | 5 | 5 | 379100 | 99.9765 | 89 | 0.0235 | 75820.0000 | 196 | 99,951 |
| TRACESPipeLite | 7 | 6 | 379329 | 93.9887 | 24261 | 6.0113 | 75824.2000 | 190 | 99,951 |
| V-pipe | 6 | 5 | 378196 | 95.5624 | 17562 | 4.4376 | 54028.0000 | 195 | 99,805 |
| VirGenA | 6 | 6 | 5407 | 93.2724 | 390 | 6.7276 | 262.0000 | 2 | 99,964 |
| ViSpA | 6 | 5 | 378485 | 95.8183 | 16518 | 4.1817 | 75697.0000 | 232 | 99,940 |

#### 2.2 Real datasets

Table S133: Viral sequences included in each real dataset according to FALCON-meta [5] (top of similarity value set to 8,000) and the total number of reference genomes found per dataset.

| Virus | SRR23101281 | SRR23101235 | SRR23101259 | SRR23101276 | SRR23101228 | SRR12175231 |
|---|---|---|---|---|---|---|
| B19V | x | x | x | x | x | |
| BKPyV | x | x | x | x | x | x |
| BuV | x | x | x | x | x | x |
| CuV | x | x | x | x | x | x |
| EBV | x | x | x | x | | |
| EV | x | x | x | x | x | x |
| HAV | | x | x | x | x | x |
| HBoV | x | | | | | |
| HCMV | | | | | x | |
| HDV | x | x | x | x | | x |
| HERV | x | x | x | x | x | x |
| HHV6 | x | x | | x | x | |
| HHV7 | x | x | x | x | x | x |
| HPV16 | x | x | x | x | x | x |
| HPV2 | x | | | | | |
| HPV31 | x | | | | x | |
| HPV51 | x | | | | | |
| HPV58 | x | | | | | |
| HPV6 | x | x | | x | x | x |
| HPyV12 | x | x | x | x | x | x |
| HPyV6 | x | x | x | x | | |
| HPyV7 | x | x | x | x | x | x |
| HPyV9 | | x | x | x | x | x |
| HSV-1 | x | x | | x | | |
| HSV-2 | x | x | x | x | | |
| JCPyV | x | x | x | x | x | x |
| KIPyV | x | x | x | | x | x |
| LIPyV | x | x | x | x | x | |
| MCPyV | x | x | x | x | x | x |
| MT | x | x | x | x | x | x |
| MWPyV | x | x | x | x | x | x |
| NJPyV | x | x | x | x | x | x |
| ReDoV | | | | | | x |
| SENV | x | | | | | |
| STLPyV | | x | x | x | x | x |
| SV40 | x | x | x | x | x | x |
| TSPyV | x | x | x | x | x | x |
| TTV | x | x | x | x | x | x |
| TTVmid | x | x | x | x | x | x |
| TTVmin | x | x | x | x | x | x |
| WUPyV | x | | x | | x | |
| Total number of references | 36 | 32 | 30 | 31 | 30 | 26 |

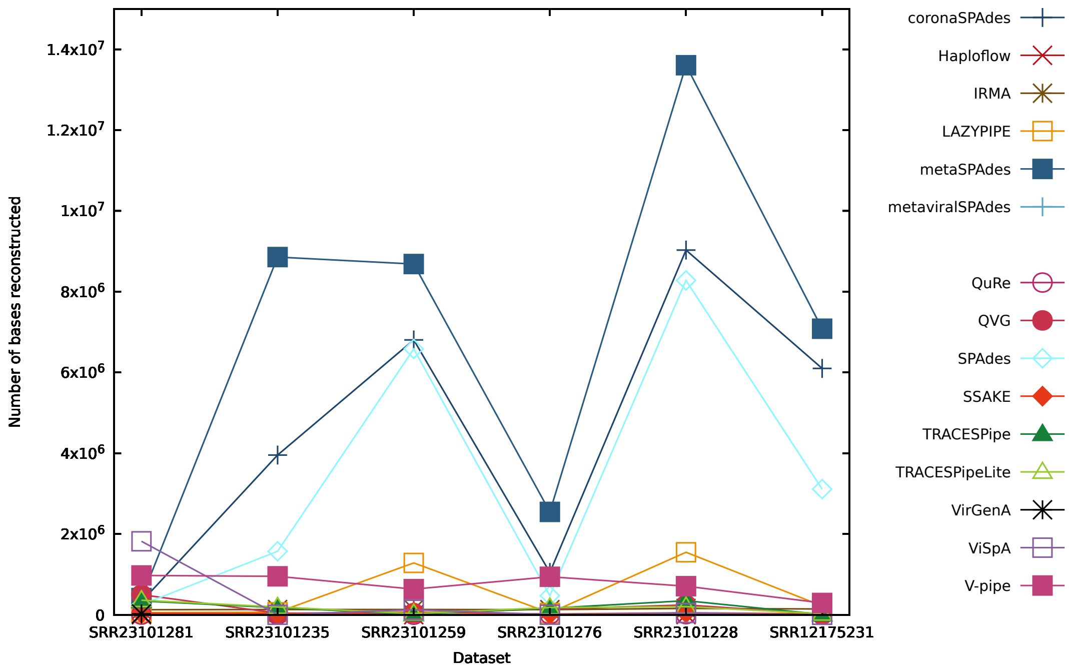


Figure S129: Figure comparing the performance of the reconstruction programs in terms of the number of bases reconstructed for datasets SRR23101281, SRR23101235, SRR23101259, SRR23101276, SRR23101228 and SRR12175231.

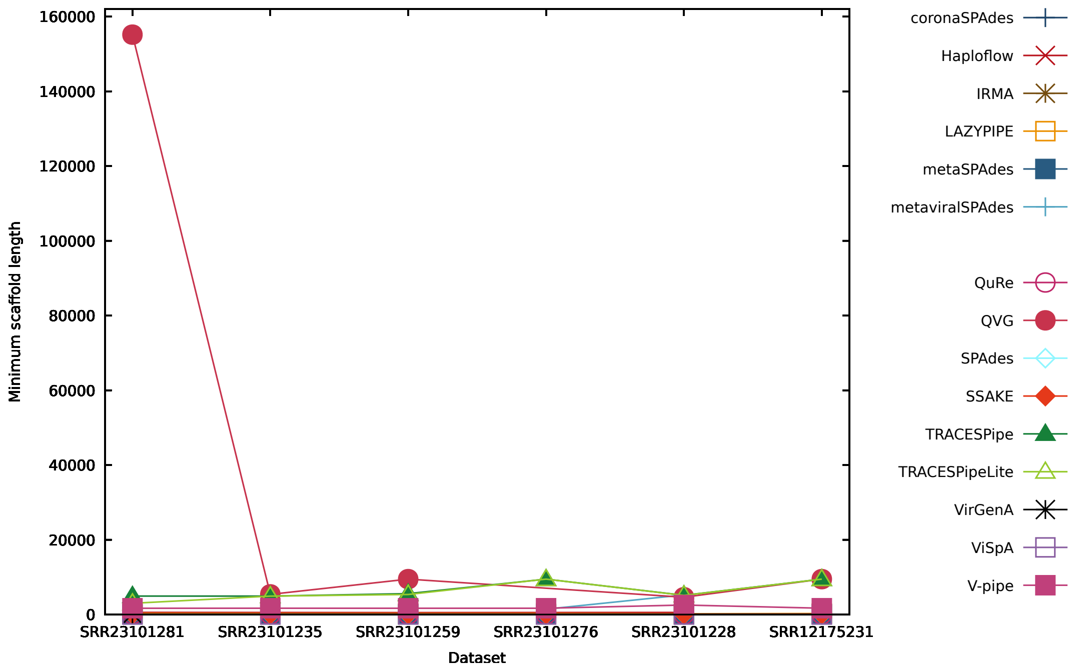


Figure S130: Figure comparing the performance of the reconstruction programs in terms of the minimum number of reconstructed bases per scaffold for datasets SRR23101281, SRR23101235, SRR23101259, SRR23101276, SRR23101228 and SRR12175231.

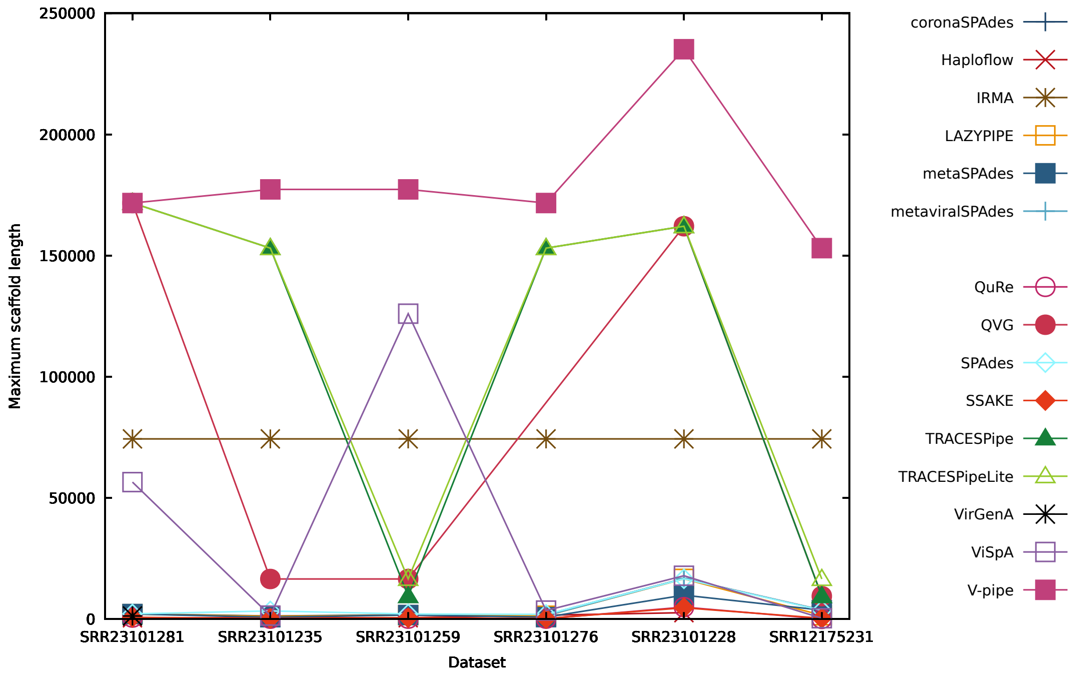


Figure S131: Figure comparing the performance of the reconstruction programs in terms of the maximum number of reconstructed bases per scaffold for datasets SRR23101281, SRR23101235, SRR23101259, SRR23101276, SRR23101228 and SRR12175231.

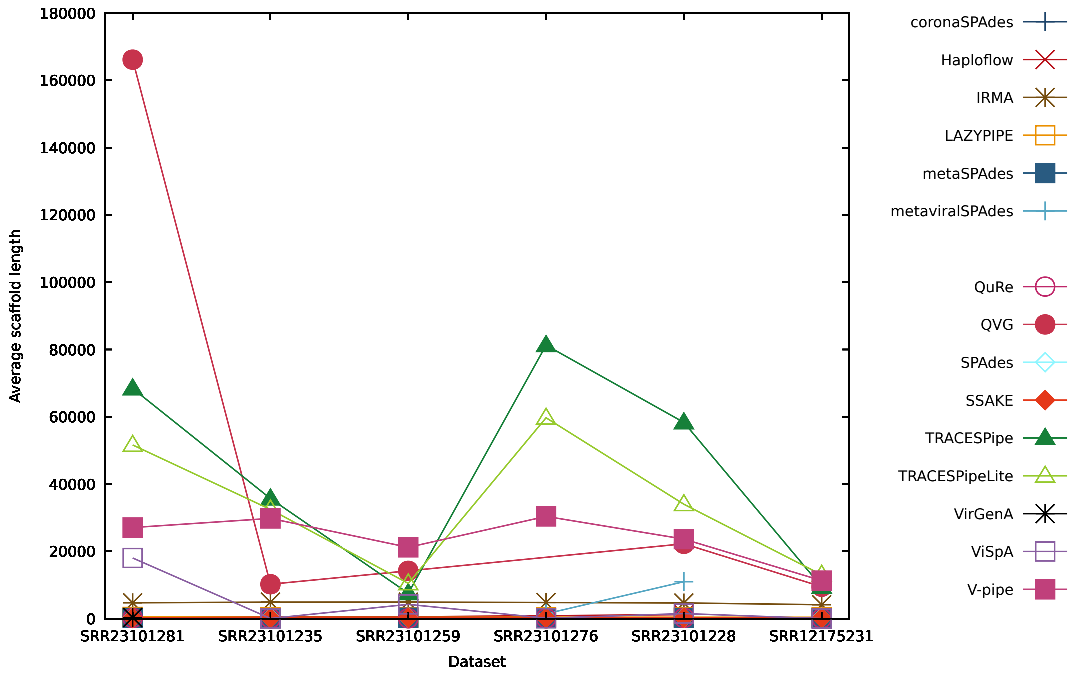


Figure S132: Figure comparing the performance of the reconstruction programs in terms of the average number of reconstructed bases per scaffold for datasets SRR23101281, SRR23101235, SRR23101259, SRR23101276, SRR23101228 and SRR12175231.

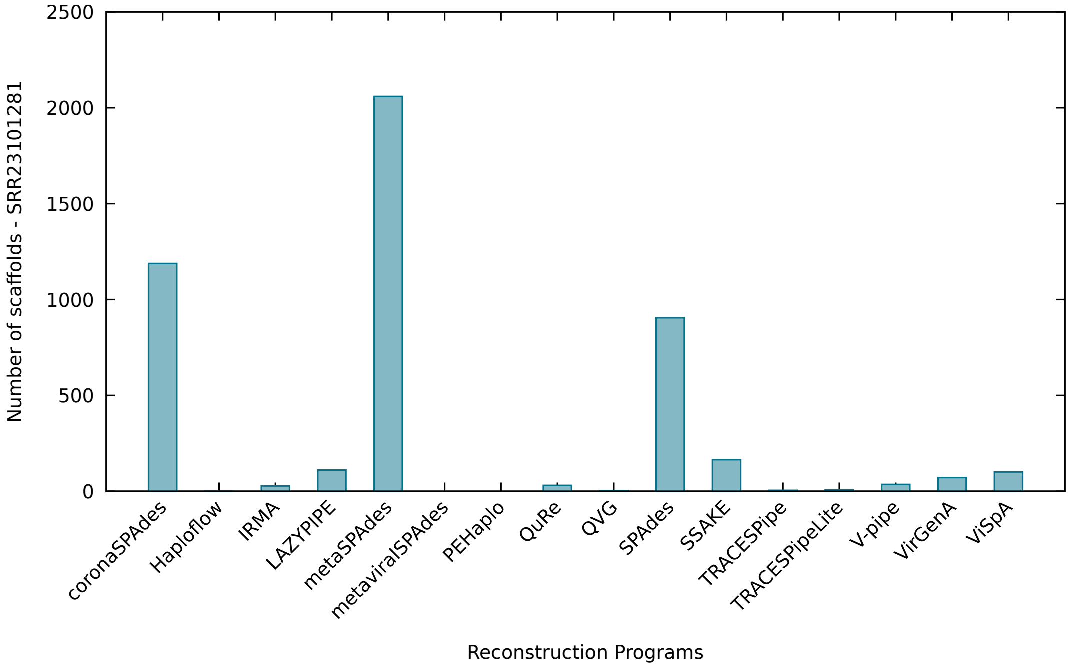


Figure S133: Figure comparing the performance of the reconstruction programs in terms of the number of scaffolds reconstructed in dataset SRR23101281.

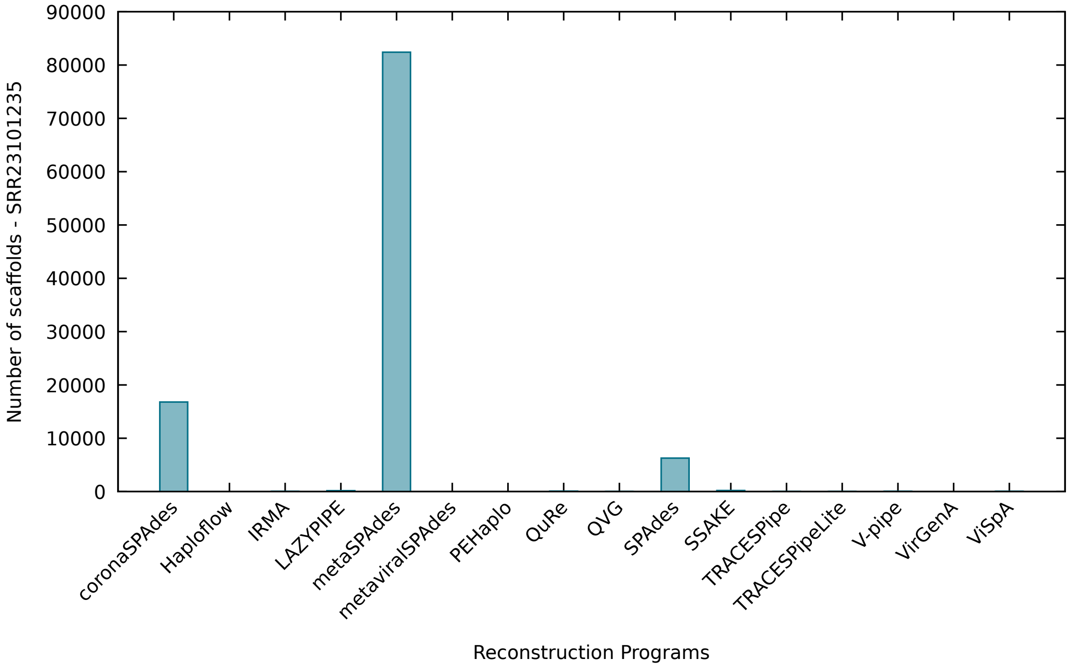


Figure S134: Figure comparing the performance of the reconstruction programs in terms of the number of scaffolds reconstructed in dataset SRR23101235.

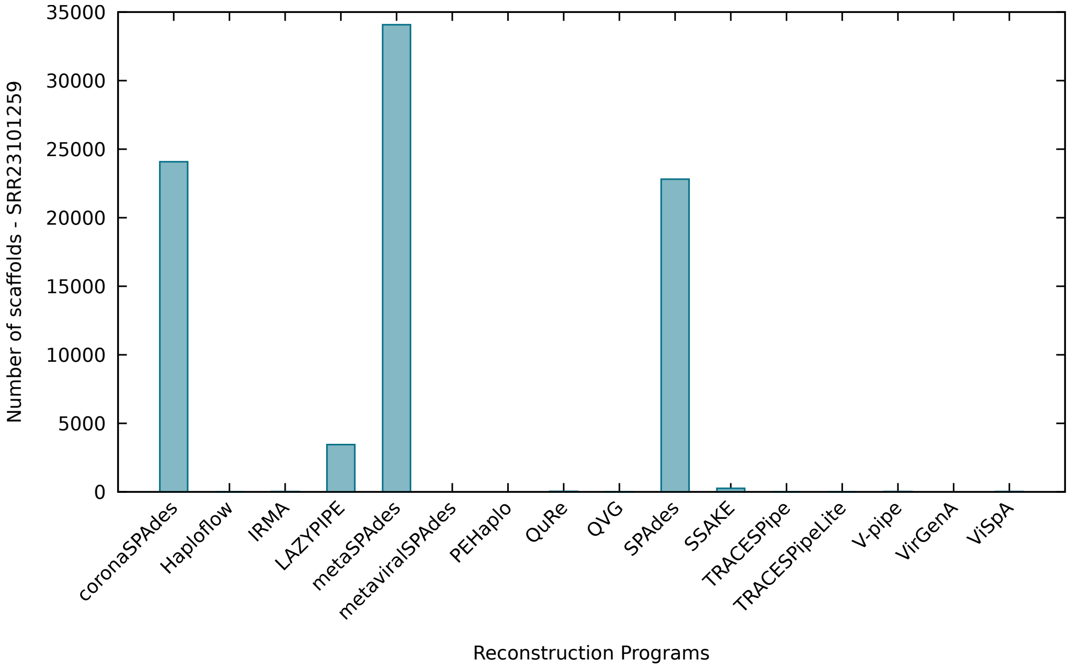


Figure S135: Figure comparing the performance of the reconstruction programs in terms of the number of scaffolds reconstructed in dataset SRR23101259.

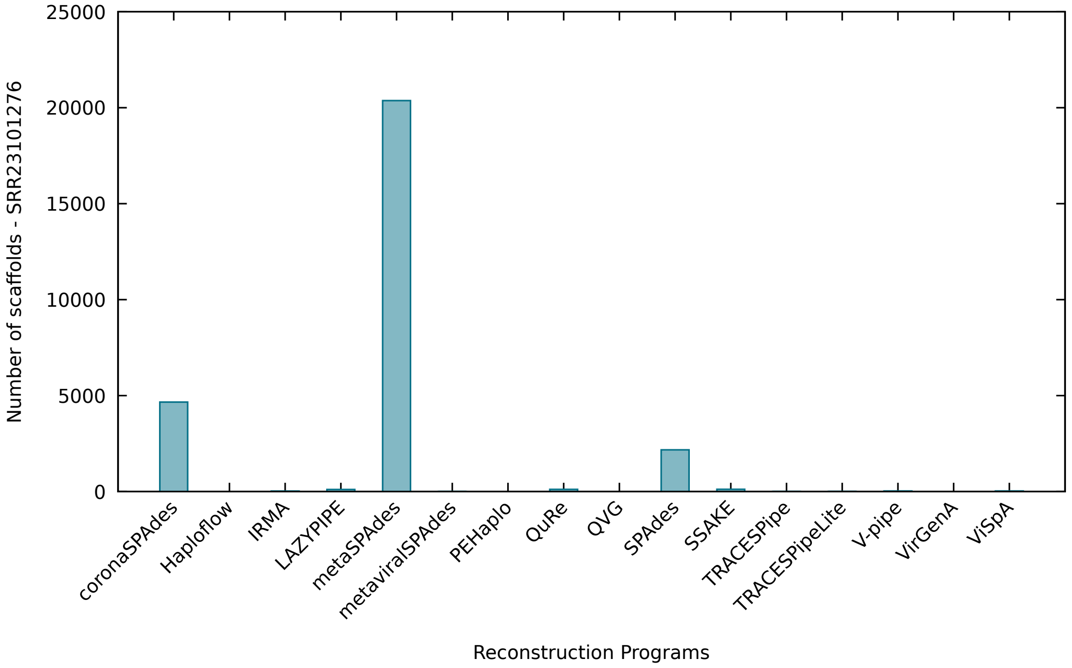


Figure S136: Figure comparing the performance of the reconstruction programs in terms of the number of scaffolds reconstructed in dataset SRR23101276.

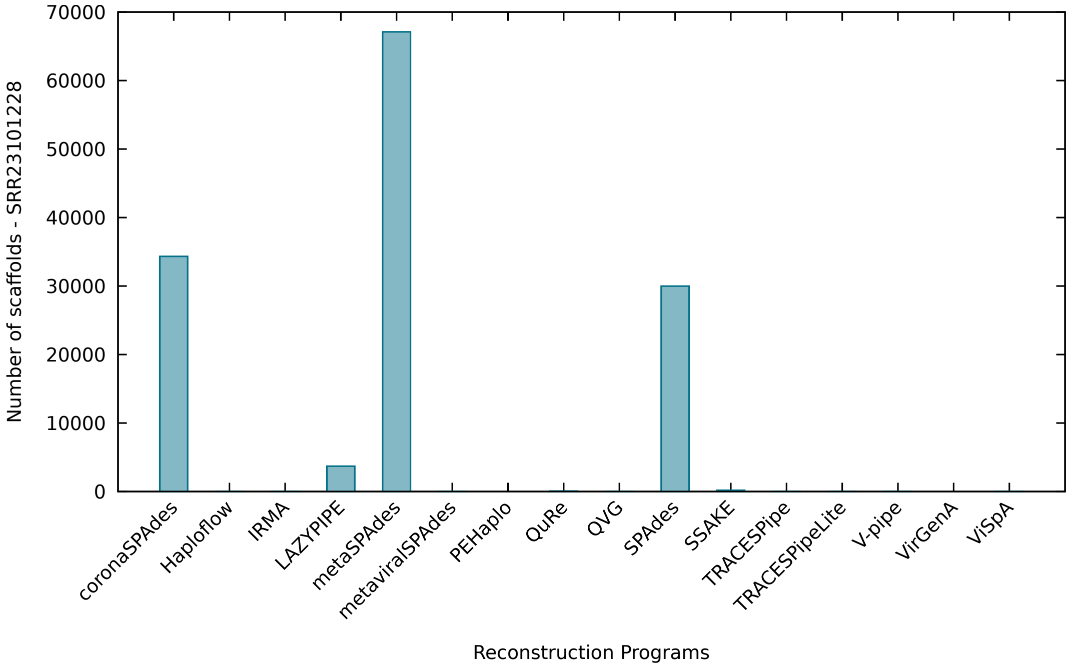


Figure S137: Figure comparing the performance of the reconstruction programs in terms of the number of scaffolds reconstructed in dataset SRR23101228.

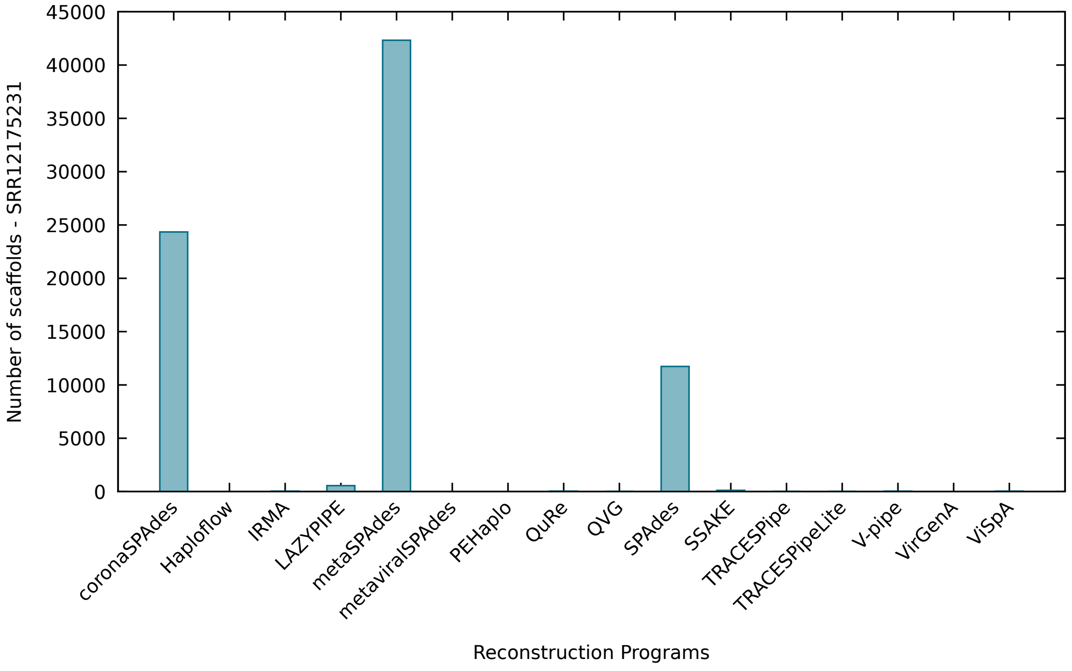


Figure S138: Figure comparing the performance of the reconstruction programs in terms of the number of scaffolds reconstructed in dataset SRR12175231.

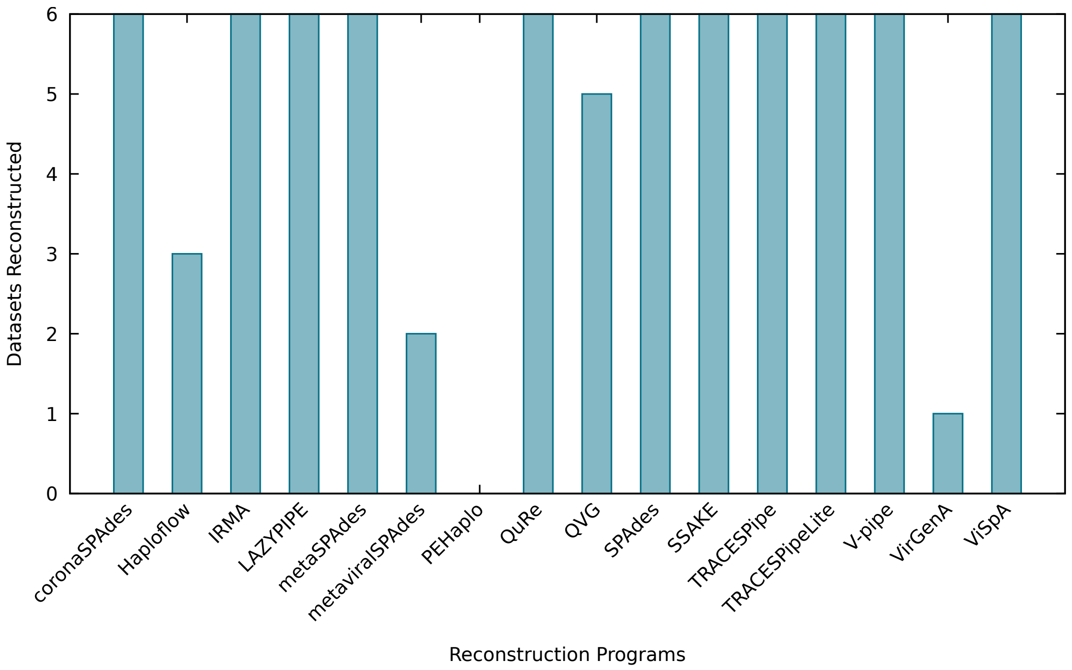


Figure S139: Figure comparing the performance of the reconstruction programs in terms of the number of datasets reconstructed across datasets SRR23101281, SRR23101235, SRR23101259, SRR23101276, SRR23101228 and SRR12175231.

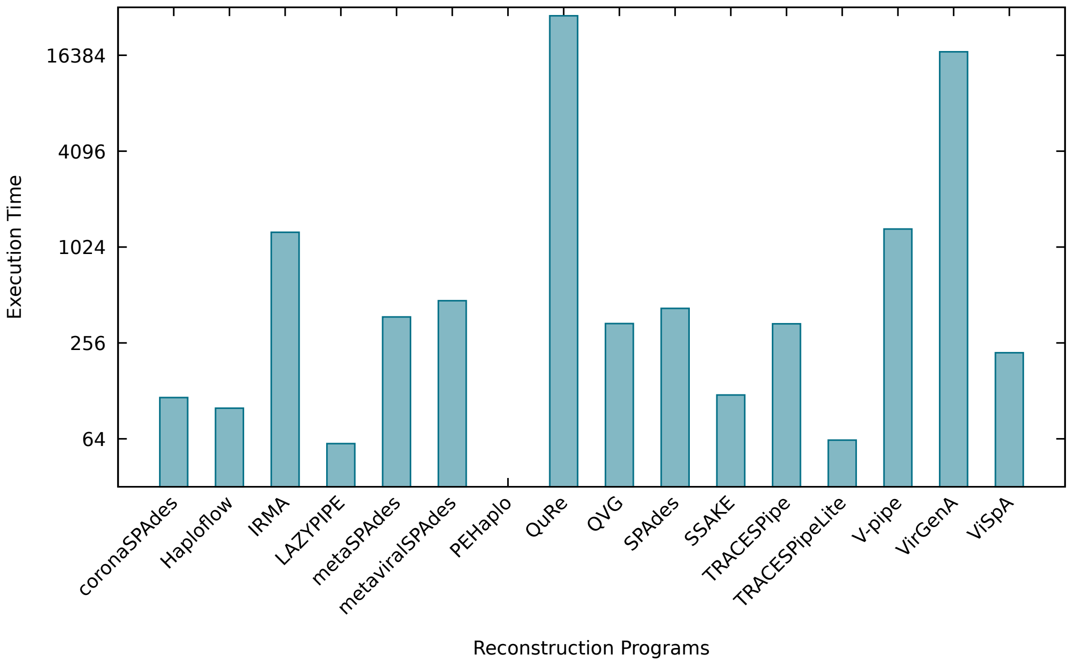


Figure S140: Figure comparing the average performance of the reconstruction programs in terms of the execution time across datasets SRR23101281, SRR23101235, SRR23101259, SRR23101276, SRR23101228 and SRR12175231. The y-axis is presented in a logarithmic scale of base 2.

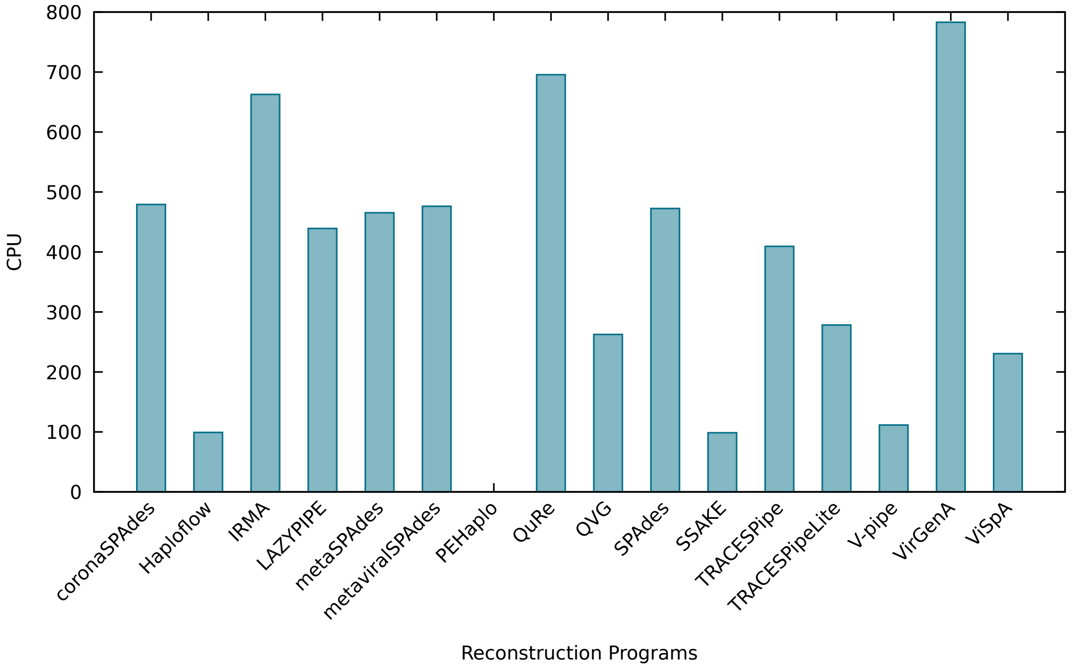


Figure S141: Figure comparing the average performance of the reconstruction programs in terms of the percentage of CPU used across datasets SRR23101281, SRR23101235, SRR23101259, SRR23101276, SRR23101228 and SRR12175231.

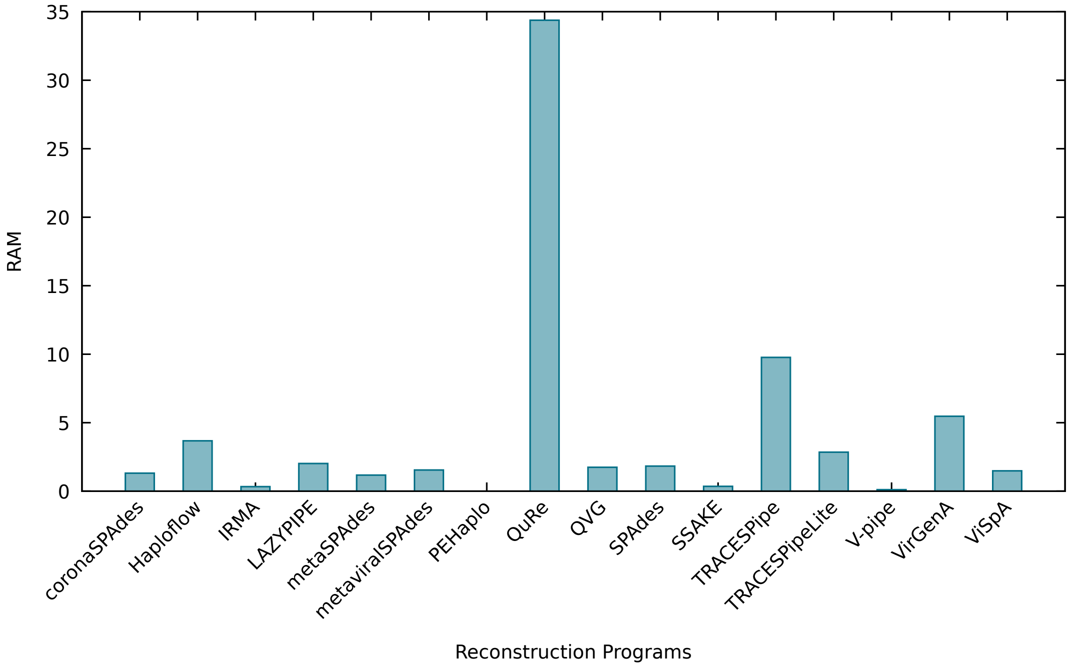


Figure S142: Figure comparing the average performance of the reconstruction programs in terms of the RAM used across datasets SRR23101281, SRR23101235, SRR23101259, SRR23101276, SRR23101228 and SRR12175231.

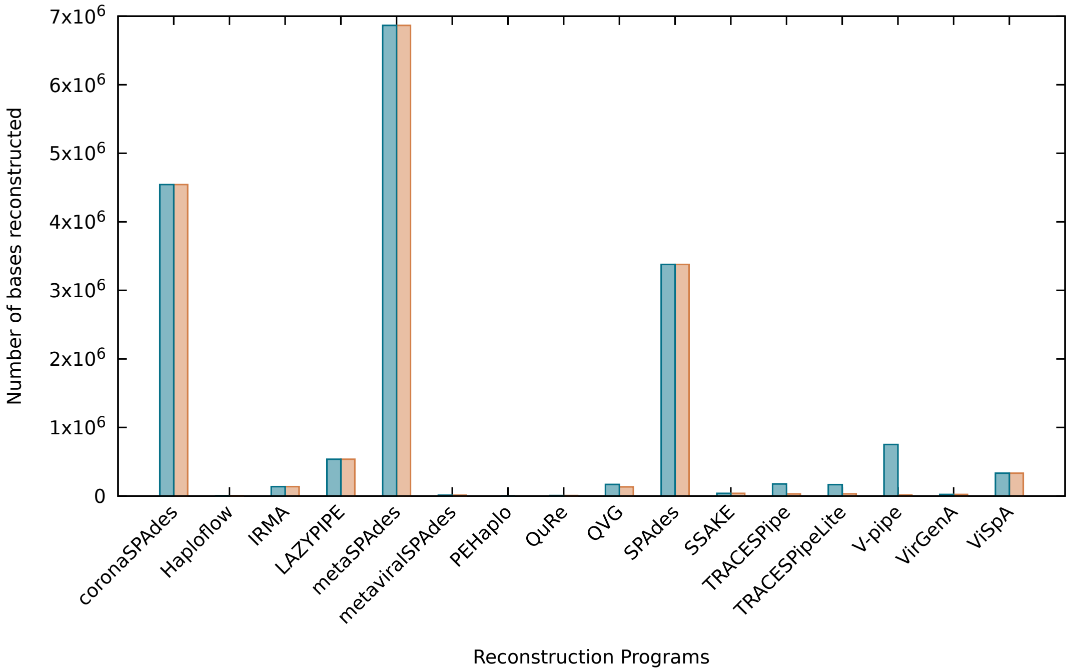


Figure S143: Figure comparing the average performance of the reconstruction programs in terms of the overall number of bases reconstructed across datasets SRR23101281, SRR23101235, SRR23101259, SRR23101276, SRR23101228 and SRR12175231. The blue bars show the performance using all bases present in the reconstructed file, whereas the orange bars show the results excluding non-reconstructed bases (N).

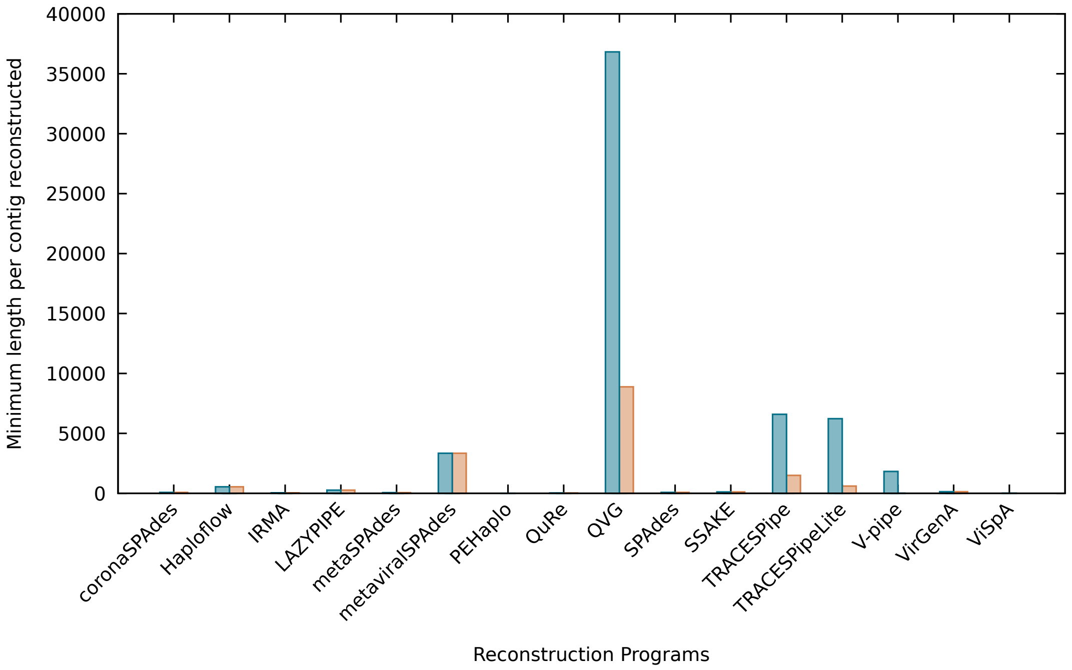


Figure S144: Figure comparing the average performance of the reconstruction programs in terms of the minimum number of bases reconstructed per scaffold across datasets SRR23101281, SRR23101235, SRR23101259, SRR23101276, SRR23101228 and SRR12175231. The blue bars show the performance using all bases present in the reconstructed file, whereas the orange bars show the results excluding non-reconstructed bases (N).

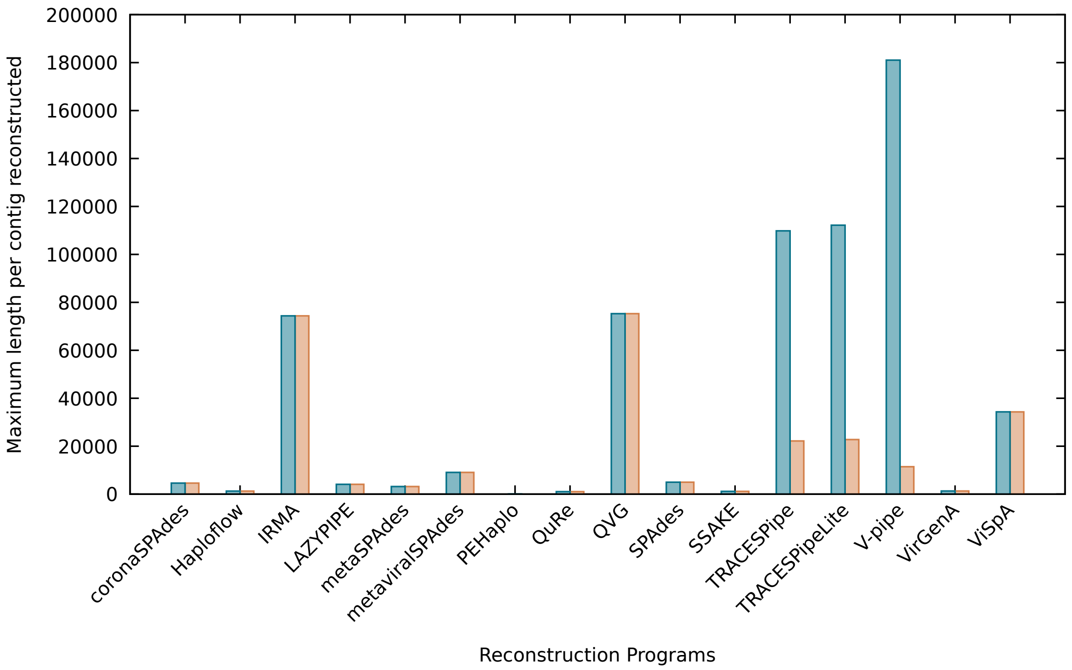


Figure S145: Figure comparing the average performance of the reconstruction programs in terms of the maximum number of bases reconstructed by scaffold across datasets SRR23101281, SRR23101235, SRR23101259, SRR23101276, SRR23101228 and SRR12175231. The blue bars show the performance using all bases present in the reconstructed file, whereas the orange bars show the results excluding non-reconstructed bases (N).

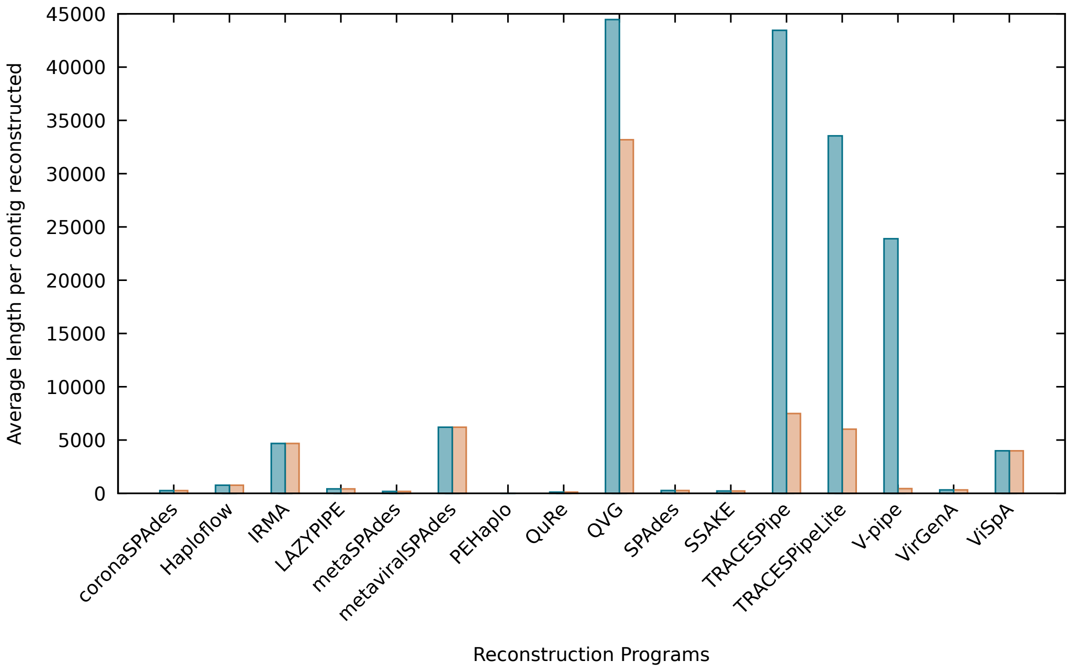


Figure S146: Figure comparing the average performance of the reconstruction programs in terms of the average number of bases reconstructed by scaffold across datasets SRR23101281, SRR23101235, SRR23101259, SRR23101276, SRR23101228 and SRR12175231. The blue bars show the performance using all bases present in the reconstructed file, whereas the orange bars show the results excluding non-reconstructed bases (N).

Table S134: Results obtained for SRR23101281 using the benchmark proposed. The execution time was measured in seconds, the RAM usage was measured in GB and the CPU usage is presented as a percentage. The executions were, when possible, capped at 6 threads and 48 GB of RAM.

| Reconstruction tool | Execution time | RAM usage | CPU usage | Number of scaffolds | Reconstructed bases | Minimum scaffold length | Maximum scaffold length | Average scaffold length | Reconstructed bases (excluding N) | Minimum scaffold length (excluding N) | Maximum scaffold length (excluding N) | Average scaffold length (excluding N) |
|---|---|---|---|---|---|---|---|---|---|---|---|---|
| coronaSPAdes | 8.74 | 0.191 | 471.5 | 1188.0 | 323936.0 | 74.0 | 2171.0 | 272.7 | 323936.0 | 74.0 | 2171.0 | 272.7 |
| Haploflow | 9.705 | 0.456 | 99.0 | 1.0 | 564.0 | 564.0 | 564.0 | 564.0 | 564.0 | 564.0 | 564.0 | 564.0 |
| IRMA | 379.445 | 0.318 | 552.5 | 28.0 | 126512.0 | 44.5 | 74366.0 | 4707.2 | 126510.5 | 44.5 | 74365.0 | 4707.15 |
| LAZYPIPE | 8.175 | 1.792 | 316.5 | 111.0 | 55765.0 | 303.0 | 2065.0 | 502.4 | 55765.0 | 303.0 | 2065.0 | 502.4 |
| metaSPAdes | 22.625 | 0.281 | 444.0 | 2059.0 | 415696.0 | 56.0 | 2111.0 | 201.9 | 415696.0 | 56.0 | 2111.0 | 201.9 |
| metaviralSPAdes | – | – | – | – | – | – | – | – | – | – | – | – |
| PEHaplo | – | – | – | – | – | – | – | – | – | – | – | – |
| QuRe | 8145.92 | 19.898 | 693.0 | 31.0 | 4528.0 | 54.5 | 641.0 | 164.5 | 4528.0 | 54.5 | 641.0 | 164.5 |
| QVG | 123.54 | 1.489 | 151.0 | 3.0 | 498530.0 | 155156.0 | 171687.0 | 166176.7 | 363660.0 | 36817.0 | 171687.0 | 121220.0 |
| SPAdes | 27.73 | 0.327 | 429.0 | 905.0 | 248396.0 | 78.0 | 2171.0 | 274.5 | 248396.0 | 78.0 | 2171.0 | 274.5 |
| SSAKE | 22.655 | 0.102 | 100.0 | 165.0 | 37364.5 | 113.0 | 748.0 | 226.55 | 37364.5 | 113.0 | 748.0 | 226.55 |
| TRACESPipe | 170.025 | 9.768 | 369.5 | 5.0 | 342114.0 | 4937.0 | 171704.0 | 68422.8 | 115285.0 | 337.0 | 94675.0 | 23057.0 |
| TRACESPipeLite | 44.55 | 2.847 | 325.5 | 7.0 | 361404.0 | 3000.0 | 171694.0 | 51629.1 | 104462.0 | 242.0 | 82985.0 | 14923.15 |
| VirGenA | 17223.62 | 5.47 | 783.0 | 71.5 | 22776.5 | 136.0 | 1308.0 | 318.5 | 22776.5 | 136.0 | 1308.0 | 318.5 |
| ViSpA | 146.355 | 4.966 | 92.5 | 101.0 | 1820439.0 | 0.0 | 56546.0 | 18024.1 | 1818763.0 | 0.0 | 56485.0 | 18007.6 |
| V-pipe | 368.615 | 0.112 | 117.0 | 36.0 | 974599.0 | 1679.0 | 171685.0 | 27072.2 | 46033.0 | 0.0 | 40509.0 | 1278.7 |

Table S135: Results obtained for SRR23101235 using the benchmark proposed. The execution time was measured in seconds, the RAM usage was measured in GB and the CPU usage is presented as a percentage. The executions were, when possible, capped at 6 threads and 48 GB of RAM.

| Reconstruction tool | Execution time | RAM usage | CPU usage | Number of scaffolds | Reconstructed bases | Minimum scaffold length | Maximum scaffold length | Average scaffold length | Reconstructed bases (excluding N) | Minimum scaffold length (excluding N) | Maximum scaffold length (excluding N) | Average scaffold length (excluding N) |
|---|---|---|---|---|---|---|---|---|---|---|---|---|
| coronaSPAdes | 204.76 | 1.959 | 491.5 | 16794.0 | 3955512.0 | 74.0 | 1302.0 | 235.5 | 3955512.0 | 74.0 | 1302.0 | 235.5 |
| Haploflow | – | – | – | – | – | – | – | – | – | – | – | – |
| IRMA | 1802.825 | 0.389 | 690.5 | 28.0 | 131421.5 | 44.5 | 74364.5 | 4892.5 | 131420.0 | 44.5 | 74363.5 | 4892.45 |
| LAZYPIPE | 47.925 | 2.434 | 432.0 | 167.0 | 63442.0 | 272.0 | 1219.0 | 379.9 | 63442.0 | 272.0 | 1219.0 | 379.9 |
| metaSPAdes | 559.805 | 1.353 | 484.5 | 82410.0 | 8856513.0 | 56.0 | 799.0 | 107.5 | 8856513.0 | 56.0 | 799.0 | 107.5 |
| metaviralSPAdes | – | – | – | – | – | – | – | – | – | – | – | – |
| PEHaplo | – | – | – | – | – | – | – | – | – | – | – | – |
| QuRe | 49325.69 | 49.125 | 695.0 | 80.5 | 4810.5 | 13.0 | 177.0 | 62.95 | 4810.5 | 13.0 | 177.0 | 62.95 |
| QVG | 448.425 | 2.406 | 250.0 | 4.0 | 40887.0 | 5392.0 | 16569.0 | 10221.8 | 31538.0 | 114.0 | 16569.0 | 7884.5 |
| SPAdes | 675.875 | 1.845 | 496.0 | 6270.0 | 1572536.0 | 78.0 | 3386.0 | 250.8 | 1572436.0 | 78.0 | 3386.0 | 250.8 |
| SSAKE | 352.905 | 0.444 | 99.0 | 199.5 | 34077.0 | 104.5 | 314.5 | 170.8 | 34077.0 | 104.5 | 314.5 | 170.8 |
| TRACESPipe | 258.065 | 9.768 | 392.5 | 5.0 | 178452.0 | 4926.0 | 153083.0 | 35690.4 | 11897.0 | 377.0 | 7241.0 | 2379.4 |
| TRACESPipeLite | 68.295 | 2.85 | 251.5 | 6.0 | 194720.0 | 4926.0 | 153080.0 | 32453.3 | 16974.0 | 377.0 | 10064.0 | 2829.0 |
| VirGenA | – | – | – | – | – | – | – | – | – | – | – | – |
| ViSpA | 254.74 | 0.17 | 272.0 | 32.0 | 1459.0 | 0.0 | 1459.0 | 45.6 | 1459.0 | 0.0 | 1459.0 | 45.6 |
| V-pipe | 1805.84 | 0.11 | 109.5 | 32.0 | 953074.0 | 1684.0 | 177314.0 | 29783.6 | 3301.0 | 0.0 | 2343.0 | 103.2 |

Table S136: Results obtained for SRR23101259 using the benchmark proposed. The execution time was measured in seconds, the RAM usage was measured in GB and the CPU usage is presented as a percentage. The executions were, when possible, capped at 6 threads and 48 GB of RAM.

| Reconstruction tool | Execution time | RAM usage | CPU usage | Number of scaffolds | Reconstructed bases | Minimum scaffold length | Maximum scaffold length | Average scaffold length | Reconstructed bases (excluding N) | Minimum scaffold length (excluding N) | Maximum scaffold length (excluding N) | Average scaffold length (excluding N) |
|---|---|---|---|---|---|---|---|---|---|---|---|---|
| coronaSPAdes | 78.37 | 1.101 | 483.5 | 24083.0 | 6805865.0 | 74.0 | 1797.0 | 282.6 | 6805865.0 | 74.0 | 1797.0 | 282.6 |
| Haploflow | 93.365 | 3.31 | 99.5 | 5.0 | 2663.0 | 508.0 | 588.0 | 532.6 | 2663.0 | 508.0 | 588.0 | 532.6 |
| IRMA | 1074.31 | 0.292 | 681.0 | 28.0 | 131803.0 | 44.5 | 74366.0 | 4906.25 | 131802.0 | 44.5 | 74365.0 | 4906.2 |
| LAZYPIPE | 58.99 | 1.891 | 470.5 | 3451.0 | 1284821.5 | 272.0 | 1655.0 | 372.3 | 1284821.5 | 272.0 | 1655.0 | 372.3 |
| metaSPAdes | 189.725 | 1.16 | 469.0 | 34078.0 | 8684069.0 | 56.0 | 1706.0 | 254.8 | 8683669.0 | 56.0 | 1706.0 | 254.8 |
| metaviralSPAdes | – | – | – | – | – | – | – | – | – | – | – | – |
| PEHaplo | – | – | – | – | – | – | – | – | – | – | – | – |
| QuRe | 19822.305 | 38.659 | 694.5 | 40.0 | 2739.0 | 20.0 | 192.5 | 74.5 | 2739.0 | 20.0 | 192.5 | 74.5 |
| QVG | 326.36 | 1.527 | 308.0 | 3.0 | 42610.0 | 9472.0 | 16569.0 | 14203.3 | 32806.0 | 6765.0 | 16569.0 | 10935.3 |
| SPAdes | 260.475 | 2.138 | 445.5 | 22812.0 | 6582169.0 | 78.0 | 2126.0 | 288.5 | 6582169.0 | 78.0 | 2126.0 | 288.5 |
| SSAKE | 21.545 | 0.353 | 96.5 | 260.0 | 67590.5 | 122.0 | 547.5 | 260.0 | 67590.5 | 122.0 | 547.5 | 260.0 |
| TRACESPipe | 326.75 | 9.768 | 419.0 | 2.0 | 15060.0 | 5588.0 | 9472.0 | 7530.0 | 3369.0 | 1673.0 | 1696.0 | 1684.5 |
| TRACESPipeLite | 64.295 | 2.85 | 257.0 | 3.0 | 31413.0 | 5363.0 | 16578.0 | 10471.0 | 16343.0 | 200.0 | 14983.0 | 5447.65 |
| VirGenA | – | – | – | – | – | – | – | – | – | – | – | – |
| ViSpA | 226.925 | 2.518 | 244.0 | 30.0 | 126910.0 | 0.0 | 126033.0 | 4230.3 | 126895.0 | 0.0 | 126018.0 | 4229.8 |
| V-pipe | 1373.24 | 0.11 | 111.0 | 30.0 | 635696.0 | 1677.0 | 177314.0 | 21189.9 | 7308.0 | 0.0 | 7308.0 | 243.6 |

Table S137: Results obtained for SRR23101276 using the benchmark proposed. The execution time was measured in seconds, the RAM usage was measured in GB and the CPU usage is presented as a percentage. The executions were, when possible, capped at 6 threads and 48 GB of RAM.

| Reconstruction tool | Execution time | RAM usage | CPU usage | Number of scaffolds | Reconstructed bases | Minimum scaffold length | Maximum scaffold length | Average scaffold length | Reconstructed bases (excluding N) | Minimum scaffold length (excluding N) | Maximum scaffold length (excluding N) | Average scaffold length (excluding N) |
|---|---|---|---|---|---|---|---|---|---|---|---|---|
| coronaSPAdes | 74.27 | 0.92 | 469.0 | 4662.0 | 1049277.0 | 74.0 | 1849.0 | 225.1 | 1049277.0 | 74.0 | 1849.0 | 225.1 |
| Haploflow | – | – | – | – | – | – | – | – | – | – | – | – |
| IRMA | 1061.895 | 0.202 | 661.5 | 28.0 | 128546.0 | 44.5 | 74366.0 | 4780.85 | 128545.0 | 44.5 | 74365.0 | 4780.85 |
| LAZYPIPE | 35.055 | 1.948 | 436.0 | 109.0 | 45900.0 | 269.0 | 1407.0 | 421.1 | 45900.0 | 269.0 | 1407.0 | 421.1 |
| metaSPAdes | 269.395 | 0.818 | 472.5 | 20370.0 | 2548210.0 | 56.0 | 823.0 | 125.1 | 2548210.0 | 56.0 | 823.0 | 125.1 |
| metaviralSPAdes | 292.695 | 0.866 | 477.0 | 1.0 | 1435.0 | 1435.0 | 1435.0 | 1435.0 | 1435.0 | 1435.0 | 1435.0 | 1435.0 |
| PEHaplo | – | – | – | – | – | – | – | – | – | – | – | – |
| QuRe | 26099.33 | 29.646 | 705.5 | 114.5 | 10380.5 | 32.0 | 198.0 | 90.7 | 10380.5 | 32.0 | 198.0 | 90.7 |
| QVG | – | – | – | – | – | – | – | – | – | – | – | – |
| SPAdes | 365.18 | 1.259 | 478.5 | 2174.0 | 474636.0 | 78.0 | 1946.0 | 218.3 | 474636.0 | 78.0 | 1946.0 | 218.3 |
| SSAKE | 90.685 | 0.262 | 99.0 | 118.5 | 19769.0 | 108.0 | 266.0 | 166.8 | 19769.0 | 108.0 | 266.0 | 166.8 |
| TRACESPipe | 209.23 | 9.768 | 430.5 | 2.0 | 162552.0 | 9472.0 | 153080.0 | 81276.0 | 16854.0 | 2185.0 | 14669.0 | 8427.0 |
| TRACESPipeLite | 44.895 | 2.849 | 317.5 | 3.0 | 179122.0 | 9472.0 | 153081.0 | 59707.3 | 12420.0 | 1049.0 | 7985.0 | 4140.0 |
| VirGenA | – | – | – | – | – | – | – | – | – | – | – | – |
| ViSpA | 177.1 | 0.175 | 233.5 | 31.0 | 3901.0 | 0.0 | 3553.0 | 125.8 | 3898.0 | 0.0 | 3550.0 | 125.7 |
| V-pipe | 1036.955 | 0.109 | 111.0 | 31.0 | 941992.0 | 1683.0 | 171693.0 | 30386.8 | 3824.0 | 0.0 | 1990.0 | 123.4 |

Table S138: Results obtained for SRR23101228 using the benchmark proposed. The execution time was measured in seconds, the RAM usage was measured in GB and the CPU usage is presented as a percentage. The executions were, when possible, capped at 6 threads and 48 GB of RAM.

| Reconstruction tool | Execution time | RAM usage | CPU usage | Number of scaffolds | Reconstructed bases | Minimum scaffold length | Maximum scaffold length | Average scaffold length | Reconstructed bases (excluding N) | Minimum scaffold length (excluding N) | Maximum scaffold length (excluding N) | Average scaffold length (excluding N) |
|---|---|---|---|---|---|---|---|---|---|---|---|---|
| coronaSPAdes | 212.26 | 2.317 | 477.5 | 34330.0 | 9029188.0 | 74.0 | 16749.0 | 263.0 | 9029168.0 | 74.0 | 16749.0 | 263.0 |
| Haploflow | 195.975 | 7.266 | 99.0 | 5.0 | 5910.0 | 554.0 | 2669.0 | 1182.0 | 5910.0 | 554.0 | 2669.0 | 1182.0 |
| IRMA | 2532.53 | 0.526 | 704.0 | 34.0 | 157990.5 | 36.0 | 74366.0 | 4650.95 | 157989.5 | 36.0 | 74365.0 | 4650.9 |
| LAZYPIPE | 161.8 | 2.041 | 521.5 | 3700.0 | 1548550.0 | 200.0 | 16621.0 | 418.55 | 1548550.0 | 200.0 | 16621.0 | 418.55 |
| metaSPAdes | 705.85 | 2.226 | 485.0 | 67109.0 | 13606179.0 | 56.0 | 9896.0 | 202.7 | 13605629.0 | 56.0 | 9896.0 | 202.7 |
| metaviralSPAdes | 650.015 | 2.226 | 475.5 | 2.0 | 21945.0 | 5248.0 | 16697.0 | 10972.5 | 21945.0 | 5248.0 | 16697.0 | 10972.5 |
| PEHaplo | – | – | – | – | – | – | – | – | – | – | – | – |
| QuRe | 56292.89 | 41.919 | 686.5 | 71.5 | 15601.5 | 26.0 | 4964.0 | 218.25 | 15601.5 | 26.0 | 4964.0 | 218.25 |
| QVG | 542.285 | 1.63 | 340.5 | 11.0 | 244664.0 | 4628.0 | 162114.0 | 22242.2 | 230785.0 | 397.0 | 162114.0 | 20980.45 |
| SPAdes | 725.575 | 3.188 | 487.0 | 29991.0 | 8279425.0 | 78.0 | 16647.0 | 276.1 | 8279325.0 | 78.0 | 16647.0 | 276.1 |
| SSAKE | 50.395 | 0.591 | 98.0 | 184.0 | 50034.5 | 110.0 | 4693.5 | 271.15 | 50034.5 | 110.0 | 4693.5 | 271.15 |
| TRACESPipe | 825.77 | 9.768 | 398.5 | 6.0 | 349944.0 | 5142.0 | 162115.0 | 58324.0 | 36105.0 | 1038.0 | 11452.0 | 6017.5 |
| TRACESPipeLite | 109.39 | 2.85 | 202.5 | 6.0 | 204008.5 | 5121.0 | 162115.0 | 34001.45 | 35548.5 | 62.0 | 16565.0 | 5924.75 |
| VirGenA | – | – | – | – | – | – | – | – | – | – | – | – |
| ViSpA | 356.615 | 0.905 | 294.5 | 30.0 | 44751.0 | 0.0 | 17921.0 | 1491.7 | 44738.0 | 0.0 | 17920.0 | 1491.3 |
| V-pipe | 2351.975 | 0.124 | 109.5 | 30.0 | 709217.0 | 2538.0 | 235154.0 | 23640.6 | 26192.0 | 0.0 | 16158.0 | 873.1 |

Table S139: Results obtained for SRR12175231 using the benchmark proposed. The execution time was measured in seconds, the RAM usage was measured in GB and the CPU usage is presented as a percentage. The executions were, when possible, capped at 6 threads and 48 GB of RAM.

| Reconstruction tool | Execution time | RAM usage | CPU usage | Number of scaffolds | Reconstructed bases | Minimum scaffold length | Maximum scaffold length | Average scaffold length | Reconstructed bases (excluding N) | Minimum scaffold length (excluding N) | Maximum scaffold length (excluding N) | Average scaffold length (excluding N) |
|---|---|---|---|---|---|---|---|---|---|---|---|---|
| coronaSPAdes | 118.965 | 1.407 | 482.5 | 24345.0 | 6100374.0 | 76.0 | 3747.0 | 250.6 | 6100374.0 | 76.0 | 3747.0 | 250.6 |
| Haploflow | – | – | – | – | – | – | – | – | – | – | – | – |
| IRMA | 754.165 | 0.263 | 686.5 | 35.0 | 144368.0 | 36.0 | 74366.0 | 4124.8 | 144366.0 | 36.0 | 74365.0 | 4124.7 |
| LAZYPIPE | 46.445 | 2.021 | 459.0 | 556.0 | 219356.0 | 271.0 | 1579.0 | 394.5 | 219356.0 | 271.0 | 1579.0 | 394.5 |
| metaSPAdes | 489.01 | 1.208 | 437.5 | 42325.0 | 7083013.0 | 56.0 | 3724.0 | 167.3 | 7082803.0 | 56.0 | 3724.0 | 167.3 |
| metaviralSPAdes | – | – | – | – | – | – | – | – | – | – | – | – |
| PEHaplo | – | – | – | – | – | – | – | – | – | – | – | – |
| QuRe | 14239.35 | 27.03 | 699.0 | 29.0 | 1702.5 | 29.5 | 104.5 | 63.25 | 1702.5 | 29.5 | 104.5 | 63.25 |
| QVG | 253.02 | 1.685 | 263.0 | 2.0 | 18944.0 | 9472.0 | 9472.0 | 9472.0 | 9785.0 | 313.0 | 9472.0 | 4892.5 |
| SPAdes | 476.85 | 2.237 | 499.0 | 11734.0 | 3112349.0 | 78.0 | 3646.0 | 265.2 | 3112349.0 | 78.0 | 3646.0 | 265.2 |
| SSAKE | 185.6 | 0.376 | 99.0 | 113.5 | 22480.0 | 124.5 | 372.0 | 198.05 | 22480.0 | 124.5 | 372.0 | 198.05 |
| TRACESPipe | 235.165 | 9.768 | 446.5 | 1.0 | 9472.0 | 9472.0 | 9472.0 | 9472.0 | 3363.0 | 3363.0 | 3363.0 | 3363.0 |
| TRACESPipeLite | 45.58 | 2.85 | 315.5 | 2.0 | 26042.0 | 9473.0 | 16569.0 | 13021.0 | 5703.0 | 1686.0 | 4017.0 | 2851.5 |
| VirGenA | – | – | – | – | – | – | – | – | – | – | – | – |
| ViSpA | 171.445 | 0.176 | 246.5 | 26.0 | 584.0 | 0.0 | 379.0 | 22.5 | 584.0 | 0.0 | 379.0 | 22.5 |
| V-pipe | 1036.14 | 0.106 | 111.0 | 26.0 | 293390.0 | 1684.0 | 153080.0 | 11284.2 | 486.0 | 0.0 | 325.0 | 18.7 |

### 3 Software and Hardware recommendations

The benchmark was executed on a computer running Linux Ubuntu 22.04.4 LTS with an Intel® Core™ i7-6700 CPU @ 3.40GHz × 8 processor and with 64 GB of RAM and the computational resources were limited, when possible, to 48 GB of RAM and 6 CPU threads. For each program, the execution time was capped at six hours, either overall or per reference genome provided. If the computer running the benchmark has different specifications, it is recommended that the options --threads and --memory are set accordingly when executing "Reconstruction.sh".

### 4 Reproducibility

#### 4.1 Installation

##### 4.1.1 Downloading the benchmark

To download the benchmark files, Git must be installed. Git can be installed using:

```
sudo apt install git -y
```

To retrieve and give permissions to all the files necessary to run the benchmark, the following commands should be used:

```
git clone https://github.com/viromelab/HVRS.git
cd HVRS/src/
chmod +x *.sh
```

##### 4.1.2 Miniconda installation

To install and execute the benchmark, Miniconda [6] should be installed. If Miniconda is not installed, please execute the following command:

```
./Installation.sh --miniconda
```

Accept the terms and conditions and finish the installation of Miniconda. After the installation, the execution of the program pauses and the message "Please close and reopen this terminal" will appear. When that happens, press any key, close and reopen the terminal so the changes are applied.
If Miniconda was not initialized, please use the following command:

```
conda init
```

##### 4.1.3 Changing paths

The options "PYTHON2_PATH" and "CONDA_PREFIX" present in the file "Installation.sh" should be in accordance with the paths where they are installed on the computer. If they are not in accordance, they should be altered to ensure the proper execution of the benchmark.

After making the changes, if necessary, return to the path to which the benchmark was downloaded using the command:

```
cd HVRS/src/
```

##### 4.1.4 Reconstruction tools installation

To install all of the reconstruction tools used in the benchmark, as well as the tools used in the simulation, classification and evaluation processes, the following step will need to be performed:

```
./Installation.sh --all
```

There are options available to install only some of the programs included in the benchmark, however, it is recommended that all programs be installed. It is also recommended that after the installation process, the terminal is closed and opened again.

Along the installation process, the execution may pause and a prompt may appear on the screen. Please follow the instructions given by the prompt and press any key to resume the installation process.

##### 4.1.5 Installation options

> **-h, --help**
> Displays the help menu and exits the program.

> **--miniconda**
> Installs Miniconda.

> **--all**
> Installs all reconstruction programs, as well as the dataset-generating, classification and evaluation tools.

> **--coronaspades**
> Installs coronaSPAdes [7].

> **--haploflow**
> Installs Haploflow [8].

> **--irma**
> Installs IRMA [9].

> **--lazypipe**
> Installs LAZYPIPE (version 2) [10].

> **--metaspades**
> Installs metaSPAdes [11].

> **--metaviralspades**
> Installs metaviralSPAdes [12].

> **--pehaplo**
> Installs PEHaplo [13].

> **--qure**
> Installs QuRe [14].

> **--qvg**
> Installs QVG [15].

> **--spades**
> Installs SPAdes [16].

> **--ssake**
> Installs SSAKE [17].

> **--tracespipe**
> Installs TRACESPipe [18].

> **--tracespipelite**
> Installs TRACESPipeLite [18].

> **--virgena**
> Installs VirGenA [19].

> **--vispa**
> Installs ViSpA [20].

> **--vpipe**
> Installs V-pipe [21].

> **--tools**
> Installs other tools used in the benchmark, including the dataset-generating, classification and evaluation tools.

#### 4.2 Simulation of the datasets

To generate and retrieve the default datasets, the following command should be used:

```
./Simulation.sh
```

In this file, each of the datasets can be changed. If datasets are added or removed, those changes should be reflected in the files "Reconstruction.sh" and "Evaluation.sh", to ensure the results are correct.

#### 4.3 Verification

To verify whether or not the tools were correctly installed, the datasets were successfully generated and the reference viruses were extracted, the following command can be executed.

```
./Verification.sh --all
```

##### 4.3.1 Verification options

> **-h, --help**
> Displays the help menu and exits the program.

> **-t, --tools**
> Check if all of the reconstruction programs are installed.

> **-d, --datasets**
> Check if all of the datasets and reference genomes were successfully generated.

> **--all**
> Check if the tools were successfully installed and if the datasets and reference genomes were generated.

> **--coronaspades**
> Checks if coronaSPAdes [7] is installed.

> **--haploflow**
> Checks if Haploflow [8] is installed.

> **--irma**
> Checks if IRMA [9] is installed.

> **--lazypipe**
> Checks if LAZYPIPE (version 2) [10] is installed.

> **--metaspades**
> Checks if metaSPAdes [11] is installed.

> **--metaviralspades**
> Checks if metaviralSPAdes [12] is installed.

> **--pehaplo**
> Checks if PEHaplo [13] is installed.

> **--qure**
> Checks if QuRe [14] is installed.

> **--qvg**
> Checks if QVG [15] is installed.

> **--spades**
> Checks if SPAdes [16] is installed.

> **--ssake**
> Checks if SSAKE [17] is installed.

> **--tracespipe**
> Checks if TRACESPipe [18] is installed.

> **--tracespipelite**
> Checks if TRACESPipeLite [18] is installed.

> **--virgena**
> Checks if VirGenA [19] is installed.

> **--vispa**
> Checks if ViSpA [20]is installed.

> **--vpipe**
> Checks if V-pipe [21] is installed.

> **--other**
> Checks if the programs necessary to generate and classify the datasets and evaluate the results are installed.

#### 4.4 Reconstruction

To reconstruct the datasets generated in the simulation step, the following command should be used:

```
./Reconstruction.sh --all --threads THREADS --memory MEMORY --virgena-timeout VIRGENA_TIMEOUT
```

If you wish to modify the datasets and/or reference genomes considered by default, the file "Reconstruction.sh" should be modified accordingly. There are also options available if only some of the programs available on the benchmark should be executed.

The values of "THREADS" and "MEMORY" should be altered to suit the computer's specifications. Additionally, there is a timeout implemented for the execution of the tool VirGenA [19], where the processing time for each reference considered is capped to the value of "VIRGENA_TIMEOUT", which is set by default to 15 minutes. Depending on the computer running the benchmark and on the dataset considered, this value may have to be altered to ensure that the timeout does not occur too soon, preventing the tool from reconstructing the datasets. Additionally, there is a timeout for all tools, set by default at six (6) hours per dataset (for tools that do not require a reference to be given) and six (6) hours per reference (for tools that require a reference to be given). This parameter can be altered as well to fit specifications.

Note: The installation of the reconstruction programs is verified at the beginning of the execution process and tools that are considered to not be installed will be skipped.

If the reference genomes are unknown, the following command can be executed:

```
./Reconstruction.sh --all --threads THREADS --memory MEMORY --virgena-timeout VIRGENA_TIMEOUT --
    reads READS_NAME --top_falcon TOP_FALCON
```

The "READS_NAME" must be substituted by the name of the files to be reconstructed before the _1.fq and _2.fq. For example, "READS_NAME" should be substituted by "dataset" if the files containing the reads are named "dataset_1.fq" and "dataset_2.fq".

The "TOP_FALCON" parameter should be changed accordingly depending on the number of references that must be retrieved by FALCON-meta [5]. It should be noted that this number does not correspond to the actual number of references to be used but to the value of top of similarity considered by FALCON-meta which should be overestimated.

##### 4.4.1 Reconstruction options

- **-h, --help**
  Displays the help menu and exits the program.
- **--all**
  Reconstructs the genomes using all of the reconstruction programs available.
- **--coronaspades**
  Reconstructs the genomes using coronaSPAdes [7].
- **--haploflow**
  Reconstructs the genomes using Haploflow [8].
- **--irma**
  Reconstructs the genomes using IRMA [9].
- **--lazypipe**
  Reconstructs the genomes using LAZYPIPE (version 2) [10].
- **--metaspades**
  Reconstructs the genomes using metaSPAdes [11].
- **--metaviralspades**
  Reconstructs the genomes using metaviralSPAdes [12].
- **--pehaplo**
  Reconstructs the genomes using PEHaplo [13].
- **--qure**
  Reconstructs the genomes using QuRe [14].
- **--qvg**
  Reconstructs the genomes using QVG [15].
- **--spades**
  Reconstructs the genomes using SPAdes [16].

> **--ssake**
> Reconstructs the genomes using SSAKE [17].

> **--tracespipe**
> Reconstructs the genomes using TRACESPipe [18].

> **--tracespipelite**
> Reconstructs the genomes using TRACESPipeLite [18].

> **--virgena**
> Reconstructs the genomes using VirGenA [19].

> **--vispa**
> Reconstructs the genomes using ViSpA [20].

> **--vpipe**
> Reconstructs the genomes using V-pipe [21].

> **-t, --threads**
> Sets the maximum amount of threads used (when possible).

> **-m, --memory**
> Sets the maximum amount of RAM used (when possible).

> **--virgena-timeout**
> Sets the maximum time used by VirGenA [19] to reconstruct a genome for each reference.

> **--timeout**
> Sets the maximum time used by all reconstruction programs. If the reconstruction program requires a reference, the maximum time corresponds to the maximum time to reconstruct a genome for each reference. If this value is larger than the virgena-timeout, the virgena-timeout is overridden.

> **-r, --reads**
> Name of the files containing the FASTQ reads. The string must be the name before _1 and _2.fq. The references are retrieved using FALCON-meta [5].

> **-y, --yes**
> Assumes the answer to all prompts is yes. This may lead to loss of data, specifically regarding the reconstructed directory and the files containing viral references.

> **--top_falcon**
> Sets the top of similarity value for FALCON-meta [5].

##### 4.5 Evaluation

To evaluate the reconstruction made by each of the tools, the following command should be used.

```
./Evaluation.sh
```

The evaluation process is made using dnadiff from the MUMmer4 package [4], GeCo3 [22] and SeqKit [23]. The results of the evaluation process are available as 2 files: total_stats.tsv and total_stats.tex. Additionally, results regarding the execution of dnadiff are stored in 2 files: dnadiff_stats.tsv and dnadiff_stats.tsv. These 4 files contain information about the performance of each of the tools tested and are present in the directory "reconstructed". If the default datasets were modified, the "Evaluation.sh" file should be changed accordingly.

##### 4.6 Plots

To generate plots, the following command should be used:

```
./Plots.sh
```

All of the plots generated are available in the directory "Graphs". If tools or datasets were added to the benchmark, the files "Plots.sh" and "Plots2.sh" should be altered accordingly.

##### 4.7 Cleaning

To erase all of the files generated for the datasets, the following command should be used:

```
./Cleaning.sh
```

#### 4.8 Run all steps

To run all of the steps of the benchmark, after downloading the benchmark following the instructions described in Section 4.1.1 of the supplement, please change the variables "PYTHON2_PATH" and "CONDA_PREFIX" to the paths where Python 2 and Miniconda are installed, on the file "Installation.sh", if necessary. Additionally, the values of the options --threads and --memory should be changed to fit the specifications of the computer running the benchmark. Afterwards, the following command should be used:

```
./Run_benchmark.sh
```

[6] Anaconda, Inc. Miniconda, 2013.
